# Connectomic reconstruction predicts the functional organization of visual inputs to the navigation center of the *Drosophila* brain

**DOI:** 10.1101/2023.11.29.569241

**Authors:** Dustin Garner, Emil Kind, Aljoscha Nern, Lucy Houghton, Arthur Zhao, Gizem Sancer, Gerald M. Rubin, Mathias F. Wernet, Sung Soo Kim

**Affiliations:** Molecular, Cellular, and Developmental Biology, University of California Santa Barbara, Santa Barbara, CA, USA; Department of Biology, Freie Universität Berlin, Berlin, Germany; Janelia Research Campus, Howard Hughes Medical Institute, Ashburn, VA, USA

## Abstract

Many animals, including humans, navigate their surroundings by visual input, yet we understand little about how visual information is transformed and integrated by the navigation system. In *Drosophila melanogaster*, compass neurons in the donut-shaped ellipsoid body of the central complex generate a sense of direction by integrating visual input from ring neurons, a part of the anterior visual pathway (AVP). Here, we densely reconstruct all neurons in the AVP using FlyWire, an AI-assisted tool for analyzing electron-microscopy data. The AVP comprises four neuropils, sequentially linked by three major classes of neurons: MeTu neurons, which connect the medulla in the optic lobe to the small unit of anterior optic tubercle (AOTUsu) in the central brain; TuBu neurons, which connect the anterior optic tubercle to the bulb neuropil; and ring neurons, which connect the bulb to the ellipsoid body. Based on neuronal morphologies, connectivity between different neural classes, and the locations of synapses, we identified non-overlapping channels originating from four types of MeTu neurons, which we further divided into ten subtypes based on the presynaptic connections in medulla and postsynaptic connections in AOTUsu. To gain an objective measure of the natural variation within the pathway, we quantified the differences between anterior visual pathways from both hemispheres and between two electron-microscopy datasets. Furthermore, we infer potential visual features and the visual area from which any given ring neuron receives input by combining the connectivity of the entire AVP, the MeTu neurons’ dendritic fields, and presynaptic connectivity in the optic lobes. These results provide a strong foundation for understanding how distinct visual features are extracted and transformed across multiple processing stages to provide critical information for computing the fly’s sense of direction.

## Introduction

During navigation, vision provides fast and multimodal information that animals use to construct a robust internal representation of the surrounding world (e.g., landmarks, optic flow, intensity gradients, color, celestial bodies, skylight polarization, etc. [1–13]). However, not all these visual features contribute equally to forming this internal representation, and how important features are extracted and used by the navigation circuits in the brain remains incompletely understood. In rodents, for instance, the layout of the surrounding space has a crucial impact on recalling the location of food, but chromatic information is much less important [14, 15]. Although the retrosplenial cortex (RSC) appears to mediate the transformation of information from the visual cortex into an internal representation of the world [16–21], RSC neurons’ responses to visual stimuli have not been characterized in detail because of the sheer number of neurons involved and the complex, distributed nature of the network. Insects, on the other hand, have much smaller brains and rely on landmarks, celestial bodies, color, and skylight polarization to glean critical information for navigation [1, 2, 11, 22–24]. These features are integrated by compass neurons, which are analogous to ‘head direction’ cells in rodents [1, 25, 26]. How these features are extracted has not been studied.

To elucidate how the fly brain processes visual information during navigation, we leveraged recently available EM data for *Drosophila melanogaster* [27] to comprehensively investigate anatomical structure of the neural pathway that conveys and transforms visual information from the retina to the central complex [28], the brain area where compass neurons reside and which is conserved across arthropods [6, 28–31].

Here, we present a complete reconstruction of all neurons in the Anterior Visual Pathway (AVP) of the fly, encompassing three levels of processing from the medulla in the optic lobes through downstream targets that synapse on compass neurons in the central complex (Fig. 1a-b). We reconstructed neurons using a collaborative platform for analyzing a previously published EM dataset that contains the entire adult fly brain [27]. Based on their connectivity and innervation patterns, we categorized the neurons that connect the medulla to the anterior optic tubercule [28, 32–36] (Fig. 1c-g). We also compared the connectivity pattern of the entire AVP across both the left and right hemispheres as well as between different EM datasets [28, 37, 38] when possible (Fig. S1h-m). Finally, we traced the upstream inputs of the ring neurons, which provide visual information to the compass neurons [2, 39–43], all the way to the medulla. This enabled us to predict connectivity-based “receptive fields” of these neurons and provided insight into potential visual features or modalities detected by these neurons. Our analysis represents an important step toward building a mechanistic model of how visual information is processed and transformed across multiple stages to guide navigational decisions.

**Fig. 1.**
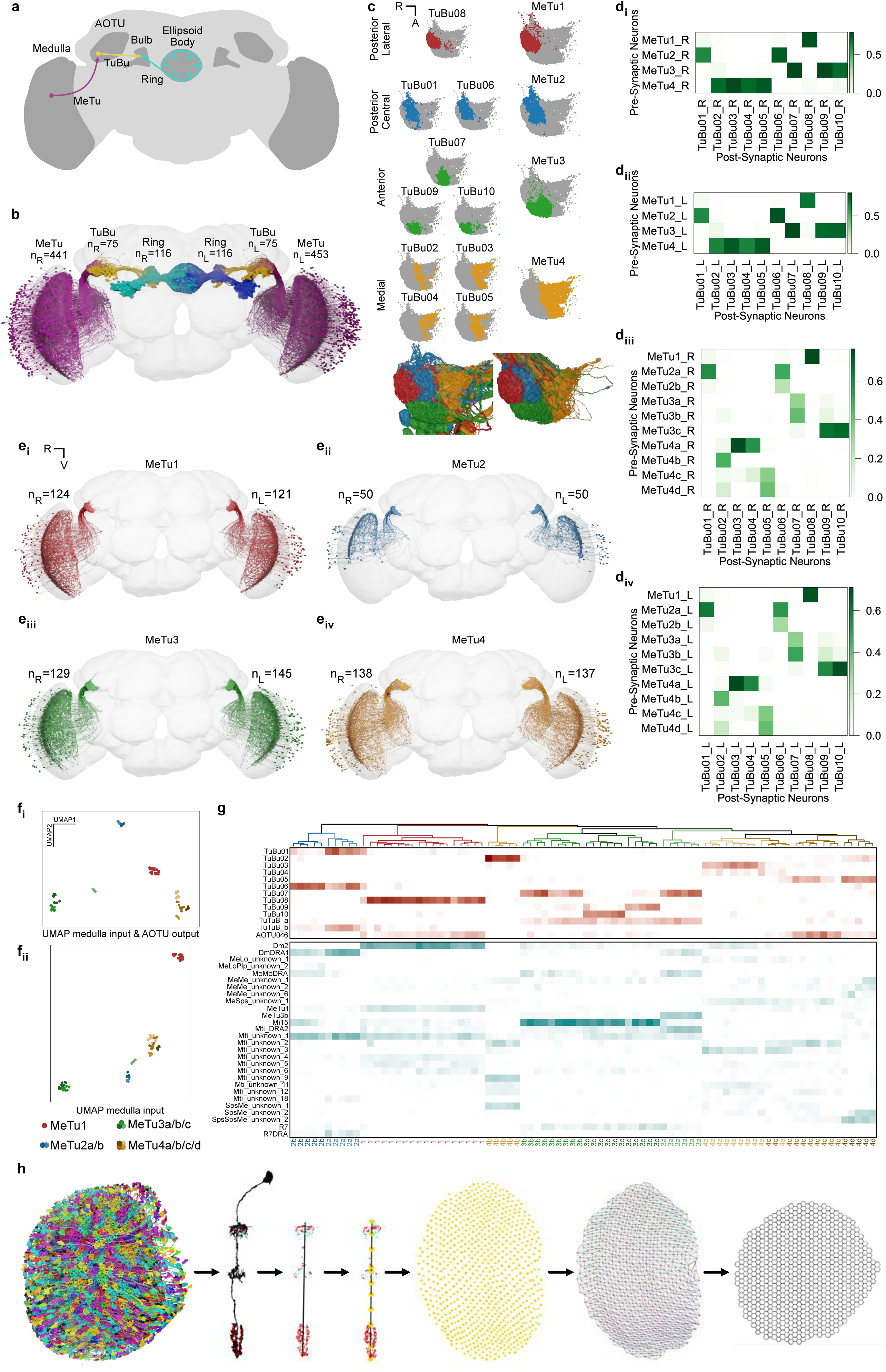
Identification and classification of MeTu neurons within the Anterior Visual Pathway. **a**, Diagram of the *Drosophila melanogaster* central brain, emphasizing the Anterior Visual Pathway (AVP). Important regions are darker gray, including medulla, AOTU, mulb, and EB (the former three have counterparts in both hemispheres). The three crucial neurons of the AVP are MeTu (from the Medulla to the AOTU_SU, purple), TuBu (from the AOTU_SU to the bulb, yellow), and Ring/ER (from the bulb to the EB, cyan). **b**, All MeTu (n=451 on the left, n=441 on the right), TuBu (n=75 on the left, n=75 on the right), and visual Ring (n=116 on the left, n=116 on the right) neurons. **c**, Top: synapse plots of TuBu (left) and MeTu (right) neurons in the posterior lateral (red), posterior central (blue), anterior (green), and medial (yellow) region of the AOTU_SU. Bottom: all TuBu (left) and MeTu (right) in the AOTU_SU_R. **d_i-ii_**, Synaptic weight matrix of MeTu type to TuBu type connectivity in the right (**d_i_**) and left (**d_ii_**) hemisphere. **d_iii-iv_**, Synaptic weight matrix of MeTu subtype to TuBu type connectivity in the right (**d_iii_**) and left (**d_iv_**) hemisphere. **e_i-iv_**, All neurons of types MeTu1 (**e_i_**), MeTu2 (**e_ii_**), MeTu3 (**e_iii_**), and MeTu4 (**e_iv_**). **f_i-ii_**, UMAPs of all MeTu neurons with identified upstream partners based on the synaptic weight of the top 5 medulla input neuron types and AOTU output neuron types (**f_i_**) or just synaptic weight of the top 5 medulla input neuron types (see Methods for details)(**f_ii_**). Groupings are generally consistent with MeTu1-4 groups in the main text with a notable exception that MeTu3a neurons are closer to MeTu2 neurons (because of the similar polarization input) than other MeTu3 neurons. **g**, Synaptic weight matrix of all MeTu neurons with identified upstream partners (columns) and their AOTU output partners (top rows in red) and top5 medulla input neuron types (bottom rows in dark cyan). Dendrogram branches and column labels are color-coded according to MeTu. **h**, Process of defining medulla columns and layers from all Mi1 neurons, a uni-columnar cell type, shown for the right optic lobe. From left to right: Render of all Mi1neurons of the right optic lobe, a single Mi1 neuron with pre- (red) and postsynaptic (cyan) sides, distal-proximal axis of a column is given by PC1 of a PCA on all synaptic sides of the corresponding Mi1 neuron, defining layer markers based on the upper and lower bound of the distal-proximal axis, m6 layer marker of all columns, manual assignment of neighboring columns along the horizontal (black), vertical (red), p (blue) and q (green) axis, and the resulting hex grid.

## Results

Information from the fly’s retina is processed in the medulla, the largest neuropil in the fly visual system, whose neuron types have been described extensively in numerous light [32, 33, 44–50] and electron microscopy (EM) studies [28, 51–53]. This processed information is conveyed to the central complex (where the sense of direction is computed) via a pathway called the “anterior visual pathway” (AVP), a conserved architecture across insects (Fig. 1a) [34, 54, 55].

Most neurons that leave the medulla in the AVP directly innervate a single anatomical structure called the “small unit” of the anterior optic tubercle (AOTUsu; Fig. 1a-b) [28, 32–36]. Since these neurons project from medulla to tubercle, they are named MeTu neurons [32]. Although prior anatomical studies of various MeTu neuron types have generally agreed at the macroscale (Fig. S1d) [34, 35, 56], the insufficient resolution of light microscopy resulted in considerable inconsistencies in grouping MeTu types and predicting their connectivity towards the central complex. Hence, we sought to provide a comprehensive view of this pathway in synapse-level detail.

The information conveyed by MeTu neurons is further processed by the neurons that connect the tubercle (AOTUsu) to the bulb (BU), called TuBu neurons [2, 34, 36] (Fig. 1a-d, S1e). There are 10 classes of TuBu neurons (Fig. 1c), each synapsing onto the dendrites of a distinct class of ‘ring’ (or ER) neurons (Fig. 1a-b, S1f), which are aptly named for their ring-like morphology (Fig. 1a) [28, 31, 41]. Ring neurons send their major processes to a donut-shaped structure called the ellipsoid body [28, 29], where they together form a complex recurrent neural network (Fig. 1b, S1g) [1, 57–60]. Finally, this visual information––along with other sensory modalities [61, 62]––is compiled by ‘compass’ neurons (also called EPG neurons [1, 28, 29]), the fly analog of the mammalian head direction cells [63].

Recently, the connectivity from TuBu to Ring to Compass neurons was studied at synaptic resolution [28], creating a dataset we refer to here as hemibrain data. However, it contains only one hemisphere and lacks upstream medulla neuropils and photoreceptor terminals. Consequently, it was impossible to confirm the results were general across hemispheres and animals, to provide information about MeTu neuron types, their inputs, and, as a consequence, to predict how visual information may be transformed on the way to the central complex. Therefore, we sought to reconstruct the entire AVP in both hemispheres (Fig. S1g-m), which will provide a solid foundation to reveal its general functions.

### Reconstructing the AVP

We densely proofread all neurons in AOTUsu using the FlyWire interface [64] (Fig. S1a-c) and the automatic synapse detection algorithm [65]. This allowed us to reconstruct the entire anterior visual pathway (AVP, Fig. 1b) from both hemispheres of the full adult female brain EM dataset (FAFB), including all MeTu neurons (453 on the left hemisphere, 441 on the right; Fig. 1a-b), TuBu neurons (75 on the left, 75 on the right), and ring neurons (116 on the left, 116 on the right). We further reconstructed all medulla-intrinsic Mi1 neurons in the medulla (782 on the left and 792 on the right; [44]) from both hemispheres (Fig. 1h) to map the exact locations of all reconstructed neurons relative to the retinotopic columns in the medulla.

To assess our proofreading quality [64], we selected 113 (of 441) MeTu neurons from the right hemisphere and performed multiple rounds of proofreading (Fig. S1b). We found that additional volume reconstructed in each round dramatically decreased after the first round (Fig. S1b_ii_). Furthermore, all MeTu neurons (894 neurons – both hemispheres and including neurons with a single round of proofreading) shared stereotypical arborization patterns in the medulla. Therefore, we were confident that our reconstruction quality of the 894 MeTu neurons was sufficiently accurate for categorization and morphological and connectivity analyses.

We focused our analyses on the detailed connectivity of MeTu neurons (Fig. 1-5, S1-S5) since the logic of their connections between optic lobes and the central brain was missing in previous studies [28]. We also included results from both hemispheres in most analyses, but, where indicated, some detailed analyses were restricted to the right hemisphere because the left hemisphere had an incomplete lamina and (minor) EM image alignment issues [27, 64, 66].

### Variance across hemispheres and brains

Our FAFB reconstruction of AVPs from both hemispheres [27, 66], together with hemibrain data [28], allowed us to make a first estimate of the variance within and across animals (Fig.S1h-m). Although the number of neurons of each type along the AVP was generally similar across hemispheres in the FAFB and hemibrain, they were more similar between hemispheres of the same brain than across brains (Fig. S1h-m) with notable exceptions for a few ring neuron types (Fig. S1h-m).

### MeTu connectivity defines four subregions in the AOTUsu and four major MeTu classes

The AOTUsu is mainly innervated by only four neuron types: the axons of MeTu neurons, the dendrites of downstream TuBu neurons (Fig. 1c), as well as synaptic terminals of AOTU046 and tubercle-to-tubercle (TuTu) neurons [28]. Drawing on the axonal arborization pattern of MeTu and the dendritic arborization pattern of TuBu, we divided the AOTUsu into four major subregions: posterior lateral (AOTUsu_PL), posterior central (AOTUsu_PC), anterior (AOTUsu_A), and medial (AOTUsu_M; Fig. 1c). These anatomical divisions led us to categorize them into four major classes (MeTu 1-4, Fig. 1c-e). Downstream TuBu neurons were categorized into 10 types, consistent with previous works (Fig. 1c; TuBu1–TuBu10; numbering follows the nomenclature of Hulse et al. [28]).

The *posterior lateral* AOTUsu (AOTUsu_PL) comprises the outermost lateral volume of the AOTUsu, facing the posterior side (Fig. 1c, Fig. S2e-f). All MeTu axons and TuBu dendrites found in this area arborize solely there and do not extend processes to other regions of the AOTUsu. We therefore designated all MeTu neurons arborizing in AOTUsu_PL as MeTu1 (Fig. 1c and e). All MeTu1 neurons form synapses with TuBu08 neurons (Fig. 1d, S1e). The dendrites of any given TuBu08 neuron partially overlap with those of neighboring TuBu08 neurons (Fig. S2e). As a population, the dendrites of TuBu08 neurons therefore roughly form a one-dimensional line along the dorsal-ventral axis, with a small positional variation along the medial-lateral axis (Fig. S2e). The AOTUsu_PL contains only one more cell type, being sparsely innervated by all four AOTU046 neurons (Fig. S2h), which may provide motor context from superior posterior slope (SPS) and also potentially mediate bilateral communication between hemispheres (Fig. S1e-f, S2g, k-l).

The *posterior central* AOTUsu (AOTUsu_PC) is located directly medial to AOTUsu_PL (Fig. 1c, Fig. S3e-f). We designated all MeTu neurons that innervate AOTUsu_PC as MeTu2. Their dendrites in medulla were limited to the dorsal rim area (DRA), where input from photoreceptors that are sensitive to skylight polarization is processed [10, 67]. MeTu2 neurons make synapses onto TuBu01 and TuBu06 (Fig. 1d, Fig. S1e). The dendrites of individual TuBu01 neurons do not overlap with neighboring neurons of the same type (Fig. S3e). As a population, TuBu01 and TuBu06 neurons form a one-dimensional line along the dorsal-ventral axis (Fig. S3e). AOTUsu_PC contains two more types of neurons TuTuB_a and TuTuB_b, both of which interconnect the two hemispheres (Fig. S2g, S3g).

The *anterior* AOTUsu (AOTUsu_A) is located in front of both AOTUsu_PL and AOTUsu_PC (Fig. 1c, Fig. S4f, h). We designated all MeTu neurons that innervate AOTUsu_A as MeTu3. MeTu3 neurons synapse onto TuBu07, TuBu09, and TuBu10 neurons. These TuBu neurons are located in the medial, central, and lateral portions of the AOTUsu_A, respectively (Fig. S4f, h). Of these three TuBu types, dendrites of the same type partially overlap each other but do not overlap with dendrites of other types (Fig. 1c, S4h). Additionally, they do not form clear one-dimensional lines along the dorsal-ventral axis as do the TuBu neurons in the AOTUsu_PL/PC. Finally, the entire anterior area is covered by dendritic and axonal processes of TuTuB_a (Fig. S3g), and innervated by axonal boutons of AOTU046 (Fig. S3h) exclusively in the TuBu09 location (Fig. S2g).

The *medial* AOTUsu (AOTUsu_M) is adjacent to both AOTUsu_PC and AOTUsu_A (Fig. 1c, Fig. S5e-f). We designated all MeTu neurons that innervate AOTUsu_M as MeTu4. Their axonal boutons tile the entire volume with varying densities and spans (Fig. S5f). MeTu4 neurons synapse onto TuBu02, TuBu03, TuBu04, and TuBu05 neurons. TuBu02 dendrites are wide along the anterior-posterior axis and have thin dendritic clumps (Fig. S5e). Dendrites of TuBu03 neurons form a rough one-dimensional line (Fig. S5e). TuBu04 dendrites are sparse but cover an exceptionally wide area, with some filling the entire volume. As a population, they are not linearly arranged (Fig. S5e). Dendrites of TuBu05 branch widely along the anterior-posterior axis but are narrower along the dorsal-ventral axis, forming a rough one-dimensional line (Fig. S5e). The AOTUsu_M has a thin two-layer structure: TuBu02 and TuBu05 both cluster along the border between the AOTUsu_M and the other AOTUsu areas, while TuBu03 and TuBu04 cluster more medially. Finally, only AOTU046 neurons, but not TuTuB_a/b neurons, innervate AOTUsu_M, thus potentially conveying motor information from SPS. (Fig. S2h).

### Synaptic connectivity defines ten MeTu subtypes

Four MeTu classes exhibit diverse morphologies and anatomical innervations, suggesting multiple parallel channels for visual features [33, 35, 41, 56], which we corroborated with genetic lines. To systematically categorize all possible subtypes of MeTu neurons, we focused our analysis on the five strongest synaptic partner types; this resulted in 28 types of upstream neurons, including a few MeTu types (Table 1). Applying a nonlinear dimensionality reduction analysis (UMAP) revealed four major patterns of presynaptic inputs mostly consistent with the four major MeTu classes defined in the previous section (Fig. 1f). We further performed categorization analyses (Fig. 1g) and found 10 subtypes.

**Table 1.**
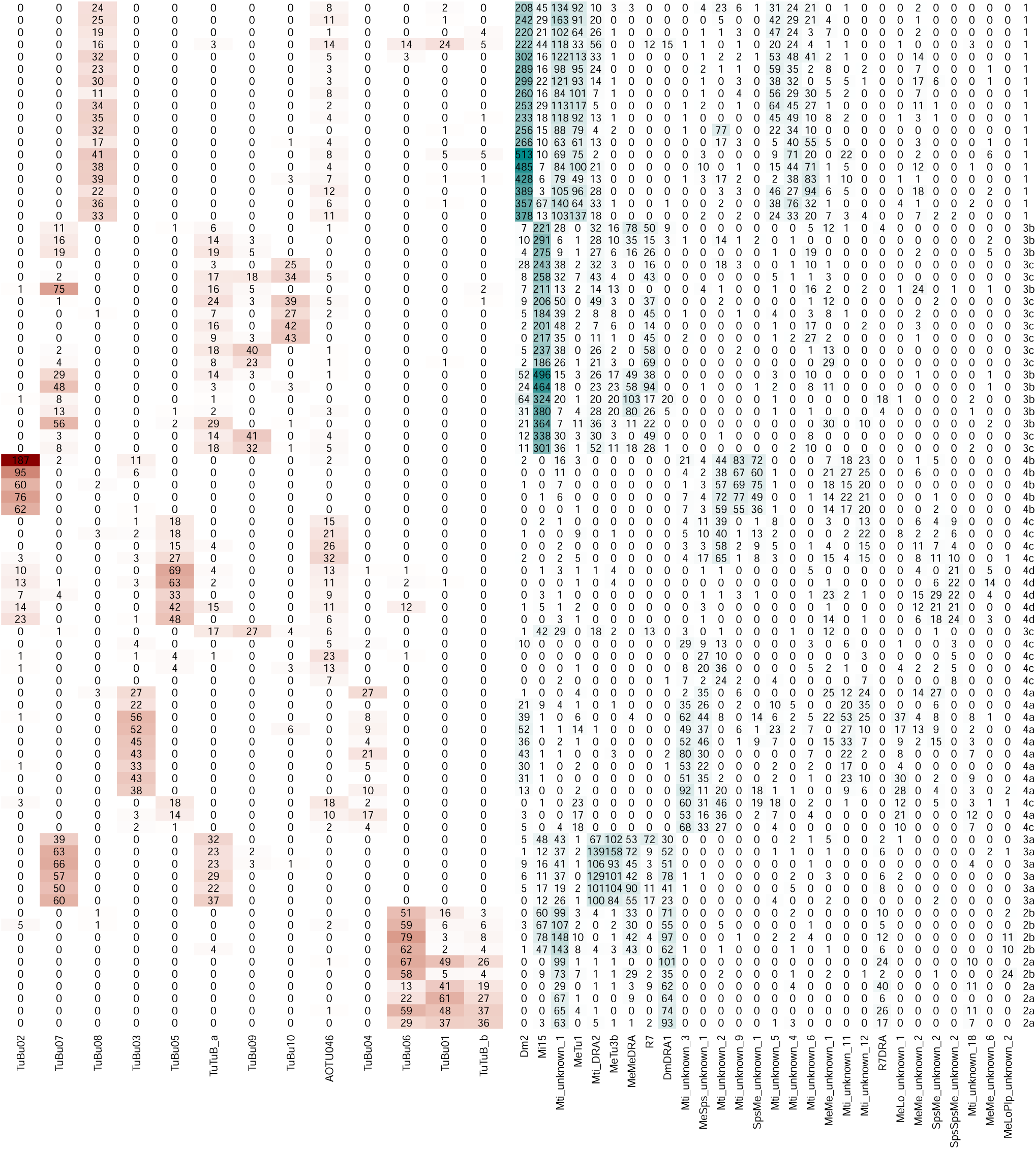
Connectivity of selected MeTu4 neurons. Each row represents a MeTu neuron. The type is shown on the right side of the table. Each column represents a neuron type making synapses with MeTu neurons. Numbers represent the total number of synapses. Red boxes represent connections from MeTu neurons and blue boxes represent connections to MeTu neurons.

#### MeTu1 cells form a homogeneous group

The compass neurons are strongly influenced by vertical stripes and their locations in azimuth [1, 59, 68], whose information is conveyed via ring neurons, likely ER4d [39–42]. This ring neuron type is the only partner downstream of TuBu08, which is, in turn, the only neuron type downstream of MeTu1 neurons (Fig. 2 and S2): Our analysis of the anatomy and connectivity of MeTu1 neurons helps explain the mechanisms that underlie ER4d neurons’ selectivity.

**Fig. 2.**
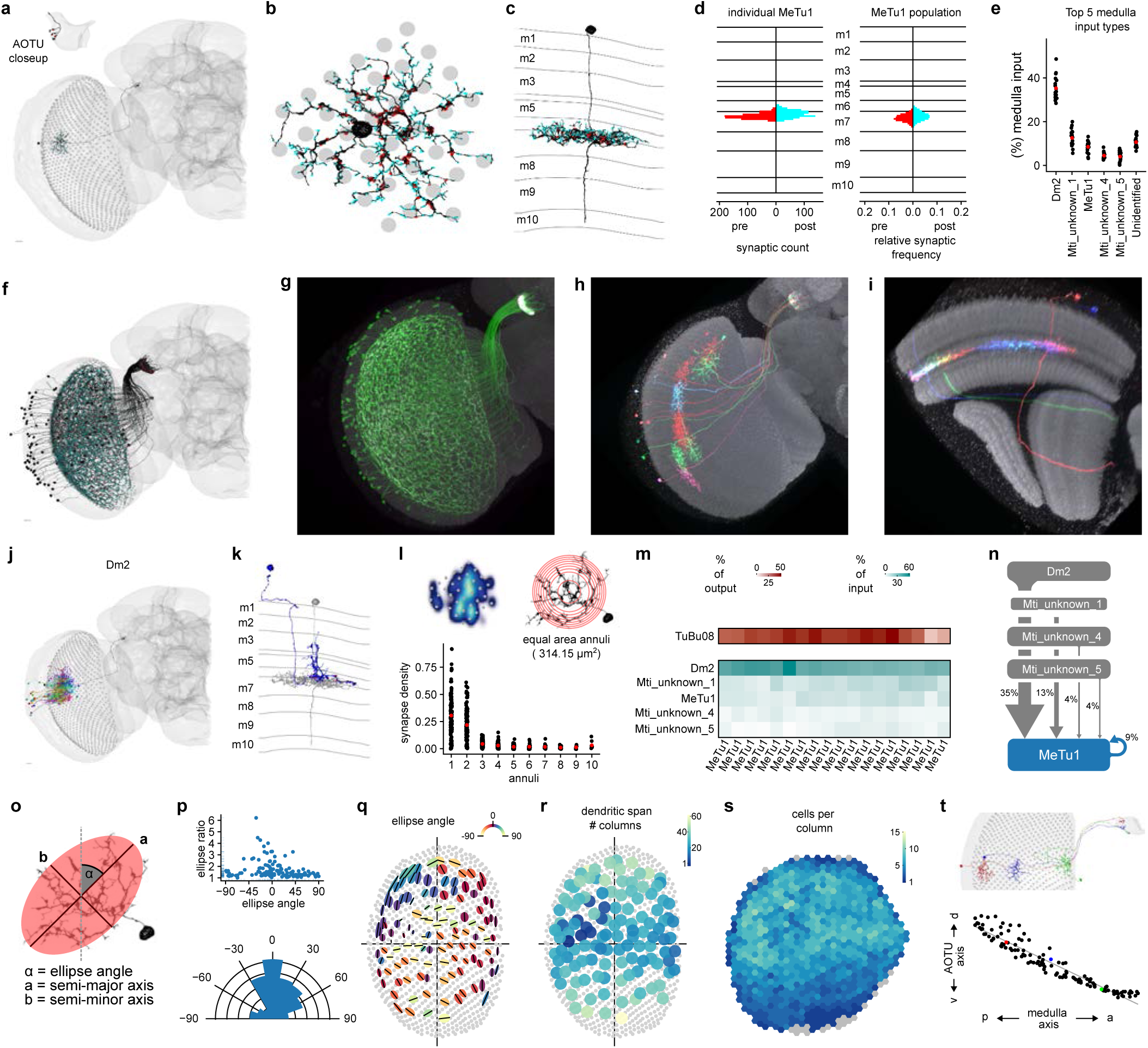
MeTu1 cells form a homogeneous group. **a**, Single MeTu1 neuron with presynapses in red and postsynapses in cyan, with a closeup of the AOTU portion in the top-left corner. Scale bar: 10 μm. **b**, Top view in the medulla of MeTu1 neuron in (**a**), with the medulla columns it spans as gray circles. **c**, Side view of same neuron as in (**a & b**), with the medulla layers labeled on the left. **d**, Left: Synapse distribution of presynapses (left) and postsynapses (right) of the MeTu1 neuron in (**a - c**) with reference to the medulla layers (as count / 100 nm). Right: Synapse distribution of all MeTu1 with reference to the medulla layers (relative frequency). **e**, Percentage of medulla input (%) of top 5 synaptic input types and unidentified types for all analyzed MeTu1neurons (see methods). Red point: population average. **f**, All MeTu1 neurons of the right optic lobe, with presynapses (red) and postsynapses (cyan). **g**, Confocal image of a MeTu1 specific split-gal4 driver (SS0038). **h-i**, MCFO images of MeTu1 neurons, from the anterior side (**h**) and the side of the medulla (**i**). **j**, MeTu1 shown in (**a**) in black, along with all upstream Dm2 partners in the medulla. **k**, MeTu1 shown in (**a**) in gray, along with a single Dm2 partner in blue, viewed from the side of the medulla with layers labeled on the left. Presynapses from the Dm2 are in red. **l**, Top-left: synapse density of a single MeTu1 neuron. Top-right: Illustration of equal area annuli with an area of 314.15µm^2^. Bottom: Synapse density for each annulus. Red point: population average. **m**, percentage of output (red) and input (cyan) of all MeTu1 neurons whose synaptic connectivity was analyzed to their top TuBu and upstream partners respectively. **n**, Diagram of the top 5 inputs of MeTu1 in the medulla, with line thickness denoting synaptic weight. **o**, Illustration of the elliptical measurement of a MeTu1 neuron’s dendritic span, with its ellipse angle (α), semi-major axis (a), and semi-minor axis (b) (see methods for details).. **p**, Top: Right optic lobe MeTu1 populations ellipse ratios (semi-major axis to semi-minor axis) as a function of the ellipse angles. Bottom: Right optic lobe MeTu1 populations relative frequency of ellipse angles. **q**, Ellipses of all MeTu1neurons of the right optic lobe with their semi-major axes as black lines and the color of the ellipse as a function of the ellipse angle. **r**, Number of columns spanned by each MeTu1 neuron of the right optic lobe. **s**, Number of MeTu1 cells each medulla column contains within its bounds. Gray columns have zero cells. **t**, Illustration of MeTu neuron topography. Top: Three MeTu1 neurons with the same dendritic position along the dorsal-ventral axis in the medulla, along with their resulting axon locations in the AOTU. Bottom: A graph of all MeTu1 neurons of the right optic lobe with their dorsal-ventral positions in the AOTU plotted against their posterior-anterior positions in the medulla. The three neurons from the top figure are highlighted in red, blue, and green.

MeTu1 neurons (N=121 left and N=124 right; Fig. 2, S2) form thick dendritic branches in the medulla layer 7 with small vertical protrusions extending to layer 6 (Fig. 2c-d). Dendrites span about 30-40 medulla columns (Fig. 2b, r), and each medulla column is innervated by multiple MeTu1 neurons (Fig. 2s). MeTu1 neurons receive the strongest input from Dm2 neurons covering the entire visual field (Fig. 2e, m-n; on average, 36 Dm2 neurons make 311 synapses per MeTu1 neuron), followed by Mti_unknown_1 (see Methods and ‘Codex Naming History.xlsx’ for the reasoning of nomenclature and other names of Mti neurons), MeTu1 (Fig. S2a), Mti_unknown_4, and Mti_unknown_5 (Fig. 2e, m-n). The density of synapses drops at 20 to 30 µm away from the medulla centroid of a MeTu1 neuron (Fig. 2l lower panel). The recurrent connection between MeTu1 neurons in medulla is unique among all MeTu types, but the functional implication is yet to be explored. The orientation of MeTu1 dendritic span, when fitted with a two-dimensional Gaussian function, tends to be vertical. We observe that near the anterior and posterior edges, MeTu neurons’ dendritic spans narrow (Fig. 2p-q), but whether this change has functional implications remains to be explored.

MeTu1 neurons project axons to the AOTUsu_PL, where they synapse with TuBu08 neurons (Fig. 1d, m, 2m, S2c), among other connections (Fig. S1e, S2g). The connection from MeTu1 neurons to TuBu08 neurons is retinotopic; the more anteriorly or posteriorly MeTu1 dendrites are located in the medulla, the more ventrally or dorsally they project in the AOTUsu_PL, respectively (Fig. 2t). In other words, each TuBu08 neuron receives input from a group of MeTu1 neurons at a particular azimuth, regardless of their elevation in medulla. Such one-dimensional mapping serves as a potential anatomical basis for TuBu08 neurons to be selective to vertical bars or the azimuthal location of visual stimuli but not the elevation (Video 1), an anatomical structure similar to the classic Hubel & Wiesel model of how simple cells in the mammalian primary visual cortex receive input from the lateral geniculate nucleus in the thalamus [69].

#### The two MeTu2 subtypes both process polarized skylight

Many insects navigate relying on skylight polarization [5, 9, 12, 70, 71]. In *Drosophila*, ER4m and ER5 neurons are the prominent ring neurons that process skylight polarization [2, 28]. MeTu2 neurons (Fig. 3, S3) are notable as the only upstream inputs of these ring neurons (Fig. S1g; via TuBu01 for ER4m; via TuBu06 for ER5) [28]. They are clustered in the dorsal half of the medulla with dendrites mainly tiling the dorsal rim area (DRA) (Fig. 3a, r), where neurons process skylight polarization [2, 13, 48, 49].

**Fig. 3.**
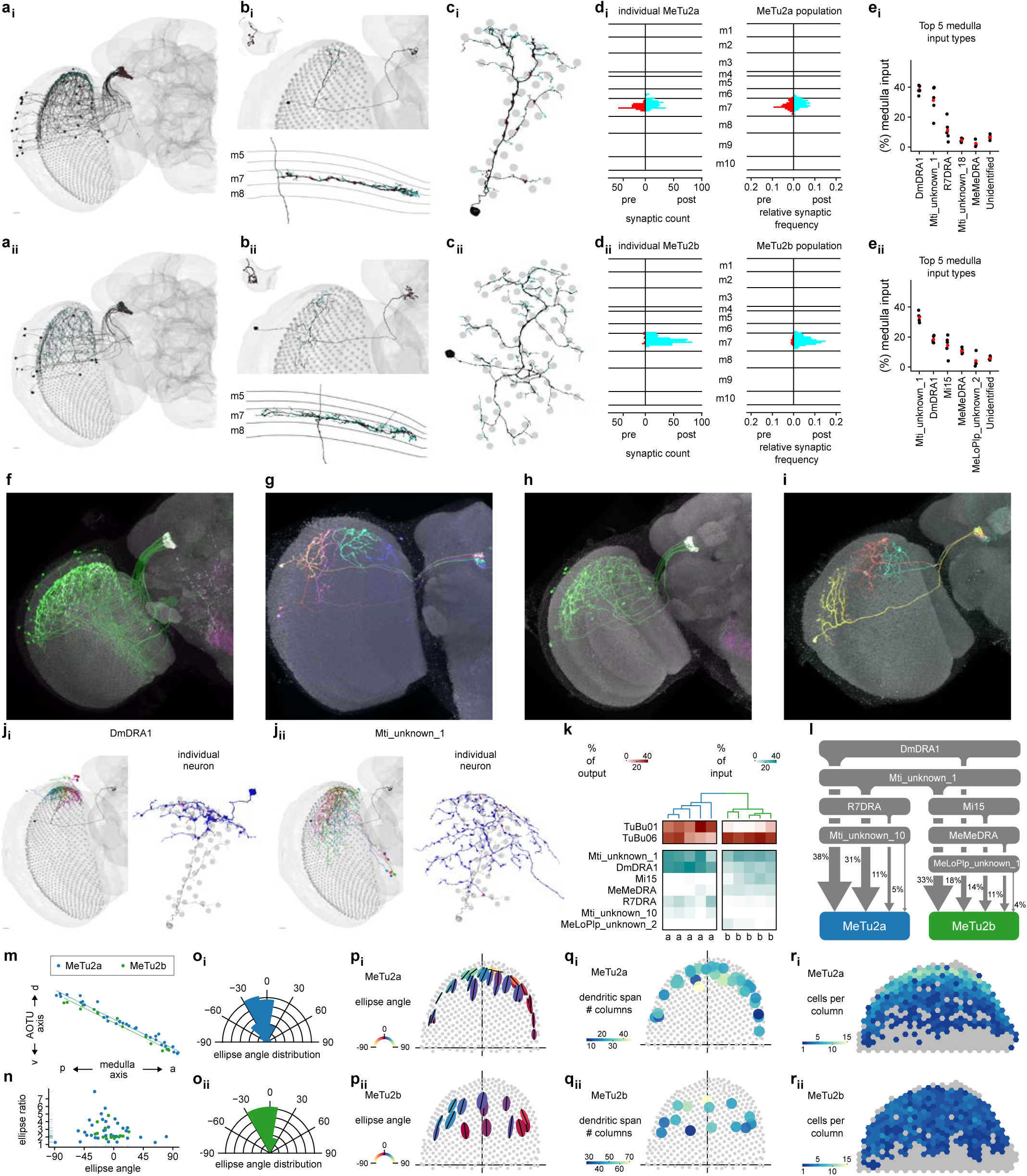
The two MeTu2 subtypes both process polarized skylight. **a**, AllMeTu2a (**a_i_**) and MeTu2b (**a_ii_**) neurons of the right optic lobe. Presynapses are red and postsynapses are cyan. **b**, Top: Single MeTu2a (**b_i_**) or MeTu2b (**b_ii_**) neuron, with a closeup of the AOTU portion in the top-left corner. Bottom: Side view of the same neuron, with the medulla layers labeled on the left. **c**, Top view of the same neurons as in (**b**), with the medulla columns it spans as gray circles. **d**, Left: Synapse distribution of presynapses (red) and postsynapses (cyan) of the neurons in (**b**) with reference to the medulla layers. Right: Synapse distribution of all MeTu2a/b respectively with reference to the medulla layers (relative frequency). **e**, Percentage of medulla input (%) of top 5 synaptic input types and unidentified types for all analyzed MeTu2a (**e_i_**) or MeTu2b (**e_ii_**) neurons. **f**, Confocal image of a MeTu2a specific split-gal4 driver (SS00336). **g**, MCFO image of MeTu2a neurons (SS00336).. **h**, Confocal image of a MeTu2a specific split-gal4 driver (SS03744). **i**, MCFO image of MeTu2b neurons (SS03744). **j**, MeTu2 main presynaptic partners. **j_i_**,Left: MeTu2a from (**b_i_**) in black, with all presynaptic DmDRA1 partners. Right: Same MeTu2a in gray, along with a single DmDRA1 partner in blue. Presynapses from the DmDRA1 are red. **j_ii_**, Left: MeTu2a from (**b_i_**) in black, with all presynaptic Mti_unknown_1 partners. Right: Same MeTu2a shown in gray, along with a single Mti_unknown_1 partner in blue.. Presynapses from the Mti_unknown_1 are red. **k**, Connectivity dendrogram of all analyzed MeTu2a/b neurons (labeled on the bottom). Percentage of output (red) and input (cyan) to their top TuBu and upstream partners respectively. **l**, Diagram of the top inputs of MeTu2a (blue) and MeTu2b (green) in the medulla, with line thickness denoting synaptic weight. **m**, Dorsal-ventral positions in the AOTU of all MeTu2a (blue) and MeTu2b (green) neurons in the right hemisphere as a function of their posterior-anterior positions in the medulla. **n**, Ellipse ratios (semi-major axis to semi-minor axis) of allMeTu2a (blue) and MeTu2b (green) of the right hemisphere as a function of the ellipse angles. **o**, Relative frequency of ellipse angles for all MeTu2a (**o_i_**) and MeTu2b (**o_ii_**) neurons in the right hemisphere. **p**, Ellipses of all MeTu2a (**p_i_**) and MeTu2b (**p_ii_**) neurons of the right optic lobe with their semi-major axes as black lines and the color of the ellipse as a function of the ellipse angle. **q**, Number of columns spanned by each MeTu2a (**q _i_**) and MeTu2b (**q_ii_**) neuron of the right optic lobe. **r**, Number of MeTu2a (**r_i_**) and MeTu2b (**r_ii_**) neurons each medulla column of the right optic lobe contains within its bounds. Gray columns have zero cells.

Our clustering analysis identified two MeTu2 subtypes (Fig. 3k) with distinct ramifications in the medulla and strikingly different connectivity patterns in the AOTUsu_PC (S3c), which we named MeTu2a (N=33 left and N=36 right; Fig. 3l) and MeTu2b (N=17 left and N=14 right; Fig. S3c-d). Both subtypes exhibit generally vertical arborizations and similar dendritic spans (Fig. 3n-p), but the interconnectivity between MeTu2b in medulla is much stronger than that within MeTu2a (Fig. S3a-b). Furthermore, MeTu2b neurons appear to receive more input from MeTu2a than they provide input to MeTu2a. Finally, although both MeTu2 subtypes mainly stratify within medulla layer 7 (Fig. 3b, d), MeTu2a neurons are postsynaptic to local interneurons Dm-DRA1 and Mti_unknown_1 that are potentially sensitive to light polarization [32], whereas MeTu2b inputs additionally include Mi15, whose function remains unknown, and a stronger connection from interhemispheric MeMe-DRA (Fig. 3e, k-l) [32]. Hence, MeTu2b may integrate additional inputs from the contralateral hemisphere, enabling the processing of a more global skylight polarization pattern.

MeTu2 innervation of the AOTUsu respects the same topographic rules as described for MeTu1 (Fig. 3m; MeTu2 neurons with dendrites in the anterior medulla project axons ventrally; those with posteriorly localized dendrites project dorsally in the AOTUsu_PC). Also, as in the medulla, MeTu2b are more strongly interconnected in AOTUsu than are MeTu2a neurons (Fig. S3b). Furthermore, while MeTu2a and MeTu2b both synapse onto TuBu01 and TuBu06, MeTu2a are more strongly connected to TuBu01 than MeTu2b, which are strongly presynaptic to TuBu06 but only weakly to TuBu01 (Fig. 3k, S3c). Finally, MeTu2a and MeTu2b are differently connected to the bilateral neurons TuTuB_a and TuTuB_b (Fig. S1e, S2g, S3g; [28, 72]). Overall, these connectivity differences in the AOTUsu_PC, combined with their distinct anatomical features in the medulla, indicate that MeTu2a and MeTu2b likely convey distinct features of skylight polarization to downstream circuits, including compass neurons, with MeTu2b processing potentially more complex and global polarization pattern.

#### Three subtypes of MeTu3 are functionally segregated

The *Drosophila* compass neurons can use the two-dimensional organization of the surrounding world to compute the head direction [68], but the source of this information was unclear. Here, we provide evidence that MeTu3 (Fig. 4 and S4) and its downstream neurons process, in addition to skylight polarization (via ER3w_ab), the two-dimensional organization of the scene (via ER2_ad/b/c, which was proposed to be functionally similar to ER4d).

**Fig. 4.**
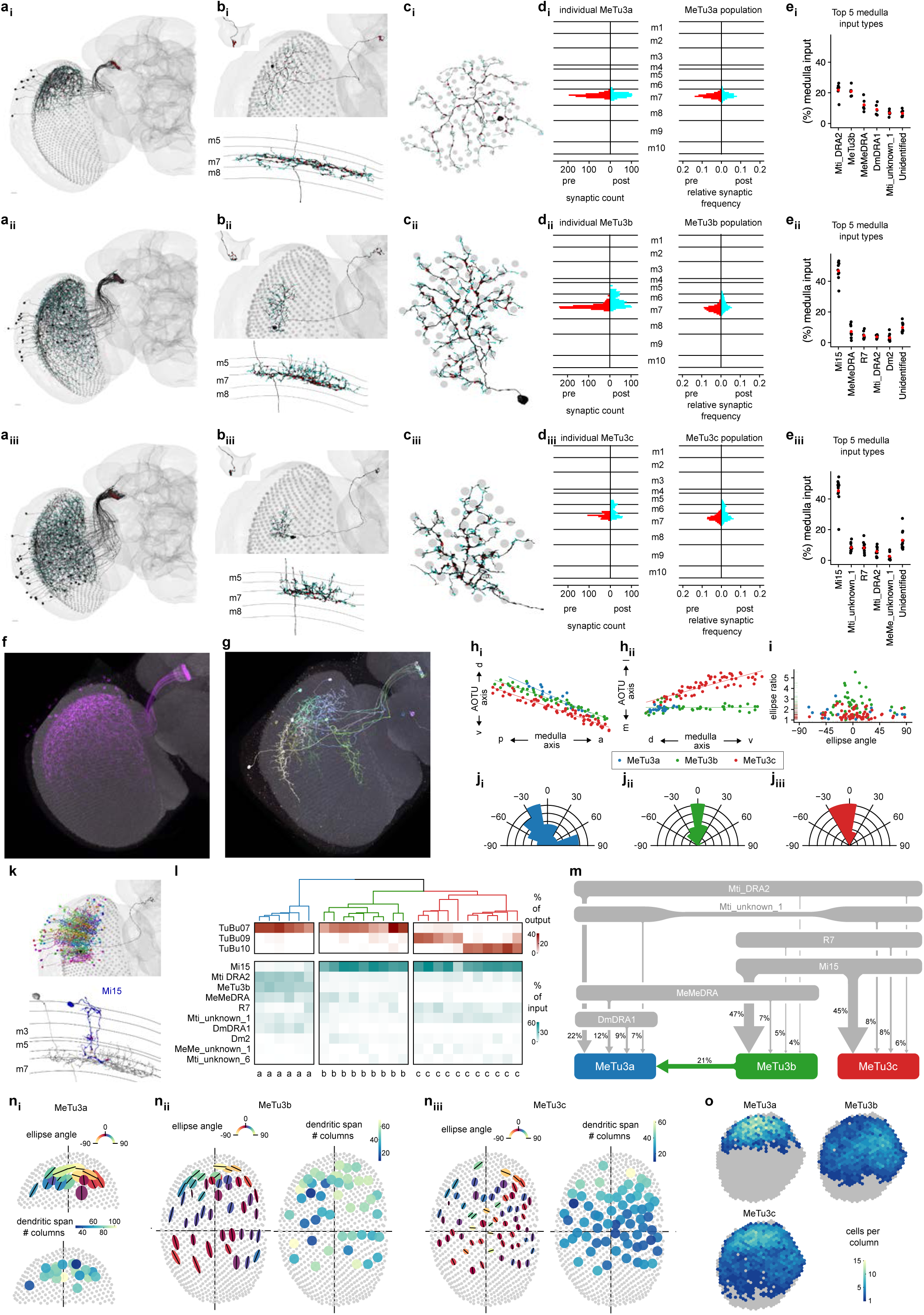
Three subtypes of MeTu3 are functionally segregated. **a_i-iii_**, Entire population of MeTu3a/b/c neurons respectively of the right hemisphere. Presynapses are red and postsynapses are cyan. **b_i-iii_**, Top: Single MeTu3a/b/c neuron respectively, with a closeup of the AOTU portion in the top-left corner. Bottom: Side view of the same, with the medulla layers labeled on the left. **c_i-iii_**, Top view of the same neurons in (**b_i-iii_**), with the medulla columns it spans as gray circles. **d_i-iii_**, Synapse distribution of presynapses (red) and postsynapses (cyan) of the neurons in (**b_i-iii_**) with reference to the medulla layers. Right: Synapse distribution of all MeTu3a/b/c respectively with reference to the medulla layers (relative frequency). **e_i-iii_**, Percentage of medulla input (%) of top 5 synaptic input types and unidentified types for all analyzed MeTu3a/b/c neurons respectively. **f**, Confocal image of a MeTu3b specific split-gal4 driver (SS00988). **g**, MCFO image of MeTu3b neurons (SS00988). **h**, Dorsal-ventral positions in the AOTU of all MeTu3a (blue), MeTu3b (green), and MeTu3c (red) neurons in the right hemisphere as a function of their posterior-anterior positions in the medulla. **i**, Ellipse ratios (semi-major axis to semi-minor axis) of all MeTu3 neurons of the right hemisphere as a function of the ellipse angles. **j_i-iii_**, Relative frequency of ellipse angles for all MeTu3a/b/c neurons in the right hemisphere respectively. **k,** Top: MeTu3b from (**b_ii_**) in black, along with all presynaptic Mi15 partners. Bottom: Side view of same MeTu3b in gray, with a single Mi15 partner in blue. Mi15 to MeTu3b synapses in red. **l**, Connectivity dendrogram of all analyzed MeTu3a/b/c neurons (labeled on the bottom). Percentage of output (red) and input (cyan) to their top TuBu and upstream partners respectively. **m**, Diagram of the top inputs of MeTu3a (blue), MeTu3b (green), and MeTu3c (red) in the medulla, with line thickness denoting synaptic weight. **n,** Morphometric analysis of MeTu3a (**n_i_**), MeTu3b (**n_ii_**), and MeTu3c (**n_iii_**) neurons. First plot: Fitted ellipses with semi-major axes as black lines and the color of the ellipse as a function of the ellipse angle. Second plot: Number of columns spanned by each MeTu3 neuron of the right optic lobe. **o**, Number MeTu3a (top-left), MeTu3b (top-right), and MeTu3c (bottom-left) neurons each medulla column in the right optic lobe contains within its bounds. Gray columns have zero neurons.

Our connectivity analysis identified three distinct MeTu3 subtypes (MeTu3a/b/c; Fig. 4l-m) with regionalized clusters of dendrites (Fig. 4a), each of which is vertically elongated (Fig. 4n). MeTu3a (N=20 left and N=19 right, Fig. 4a_i_-e_i_) has dendrites that cluster in the dorsal third of the medulla similar to MeTu2 (suggesting processing of skylight polarization), are confined to layer 7 (Fig. 4b_i_, d_i_), and specifically lack vertical protrusions across medulla layers (Fig. 4b_i_, d_i_). Their dendrites manifest more radial branches than those of MeTu2a/b, which are confined in a more elliptical area and elongated along the ventral-dorsal direction (Fig. 3p, 4n_i_). MeTu3b cells (N=53 and N=46 right) have dendrites clustered most densely in the dorsal half of the medulla but also extend to the ventral two-thirds, with pronounced vertical protrusions that cover layers 5, 6, and 7 (Fig. 4b_ii_, d_ii_). They receive direct inhibitory input from UV-sensitive R7 photoreceptors and indirect input from blue/green-sensitive R8 [32] via Mi15, suggesting they process chromatic information (Fig. 4e_i__-ii_, l-m, S4g). MeTu3c cells (N=72 left and N=64 right) have dendrites lower than those of MeTu3b, covering the equator and some of the ventral part of medulla (Fig. 4b_iii_, e_iii_). Dendritic processes innervate the same layers 5, 6, and 7 as MeTu3b, and receive the same inhibitory input from R7 and indirect input from R8 via Mi15 (Fig. 4e_iii_), suggesting similar potential chromatic coding as MeTu3b.

MeTu3 innervation of the AOTUsu_A respects the same topographic rules as described for MeTu1 and MeTu2 (anterior-posterior axis in medulla to ventral-dorsal axis in AOTUsu_A; Fig. 4h). Axons of MeTu3a/b/c are not well segregated in the AOTUsu_A, despite the downstream TuBu neurons (TuBu07, TuBu09, and TuBu10; Fig. S4c) having well-segregated dendrites (Fig. 1c, Fig. S4h) [28]. Consequently, some MeTu3 neurons are specifically connected to only one of the three TuBu types––TuBu07––while others are connected to both TuBu07 and TuBu09, or to both TuBu09 and TuBu10 (Fig. 4l, Fig. S4c). All three MeTu3 subtypes are strongly and reciprocally connected to TuTuB_a neurons (Fig. S1e, S2g). TuTuB_a neurons receive most of their input from MeTu3 neurons. Likewise, MeTu3 neurons receive most of their input from TuTuB_a. Since the neurotransmitter is predicted likely to be inhibitory [65], MeTu3 neurons may exhibit strong bilateral inhibitory interactions across the entire visual field.

MeTu3a and MeTu3b neurons are mainly connected to TuBu7, upstream of ER3w_ab. This convergence of MeTu3a and MeTu3b suggests that ER2w_ab may encode a combination of skylight polarization and chromatic information of the sky. On the other hand, MeTu3c neurons are mostly presynaptic to both TuBu09 and TuBu10 (Fig. 4l, S4c). Intriguingly, TuBu09 neurons receive input from MeTu3c neurons with dendrites located more dorsally in the medulla, whereas TuBu10 neurons receive input from MeTu3c neurons with dendrites located more ventrally in the medulla (Fig. 4h_ii_, red dots). Thus, the neurons downstream of MeTu3c can encode the elevation of visual stimuli. This allows encoding the elevation of the sun or a 2-D organization of visual objects in a surrounding scene, a unique capability among all MeTu neurons.

#### Four MeTu4 subtypes convey widefield visual inputs

Compass neurons receive diverse input from ring neurons, some of which exhibit responses to the contralateral visual field and self-generated motion signals [40]. Considering that these ring neurons, whose dendrites are in the inferior BU, are downstream of MeTu4 neurons (Fig. 5 and S5) that originate in the ipsilateral optic lobe, their response pattern was mysterious. Our analyses show that MeTu4 neurons receive inputs from distinct parts of the visual world (dorsal, frontal, ventral), with virtually no input from a columnar medulla cell type (with the only weakly detectable columnar input from Dm2; Fig. 5e, k-l), but mostly from Mti neurons that arborize within many medulla columns, called and from others that convey potential motor information from SPS to medulla (Fig. 4e). These unique properties of MeTu4 may explain the mysterious properties of ring neurons in the inferior BU.

**Fig. 5.**
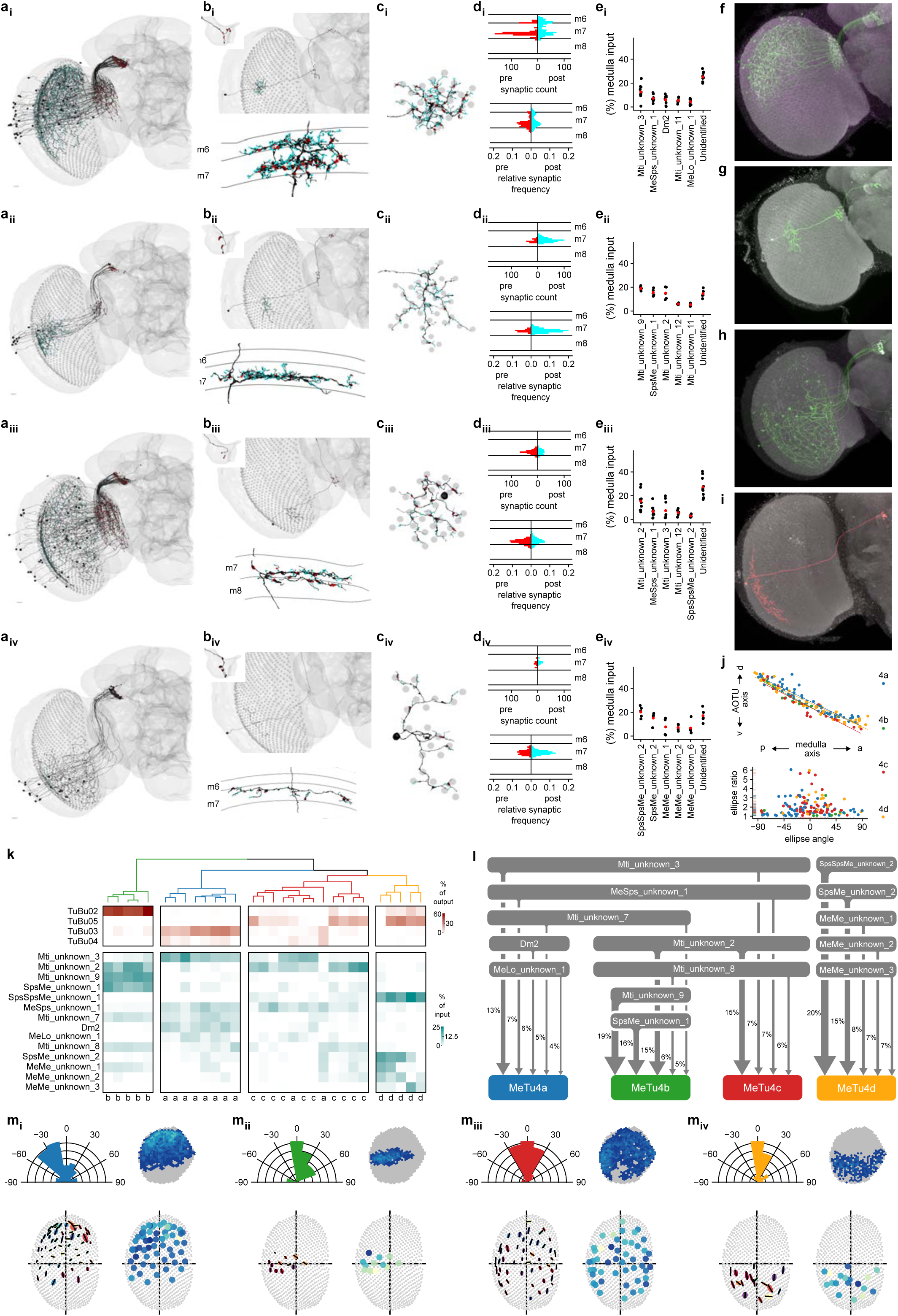
Four MeTu4 subtypes convey widefield visual inputs. **a_i-iv_**, Entire population of MeTu4a/b/c/d neurons respectively of the right hemisphere. Presynapses are red and postsynapses are cyan. **b_i-iv_**, Top: Single MeTu4a/b/c/d neuron respectively, with a closeup of the AOTU portion in the top-left corner. Bottom: Side view of the same neuron, with the medulla layers labeled on the left. **c_i-iv_**, Side view of the same neurons in (**b_i-iv_**, with the medulla columns it spans as gray circles. **d_i-iv_**, Top: Synapse distribution of presynapses (red) and postsynapses (cyan) of the neurons in (**b_i-iv_**) with reference to the medulla columns. Bottom: Synapse distribution of all MeTu4a/b/c/d respectively with reference to the medulla layers (relative frequency). **e_i-iv_**, Percentage of medulla input (%) of top 5 synaptic input types and unidentified types for all analyzed MeTu4a/b/c/d neurons. **f**, Confocal image of a MeTu4a specific split-gal4 driver (SS03719). **g**, MCFO image of MeTu4a neuron (SS03719). **h**, Confocal image of a MeTu4d specific split-gal4 driver (SS23880). **i**, MCFO image of MeTu4d neuron (SS23880).. **j**, Top: dorsal-ventral positions in the AOTU ofall MeTu4a (blue), MeTu4b (green), MeTu4c (red), and MeTu4d (yellow) neurons in the right hemisphere as a function of their posterior-anterior positions in the medulla. Bottom: Ellipse ratios (semi-major axis to semi-minor axis) of all MeTu4 neurons of the right hemisphere as a function of their ellipse angles. **k**, Connectivity dendrogram of all analyzed MeTu4a/b/c/d neurons (labeled on the bottom). Percentage of output (red) and input (cyan) to their top TuBu and upstream partners respectively. **l**, Diagram of the top inputs of MeTu4a (blue), MeTu4b (green), MeTu4c (red), and MeTu4d (yellow) in the medulla, with line thickness denoting synaptic weight. **m,** Morphometric analysis of MeTu4a (**m_i_**), MeTu4b (**m_ii_**), MeTu4c (**m_iii_**), and MeTu4d (**m_iv_**) neurons. Top-left: Relative frequency of ellipse angles. Top-right: Number of respective MeTu4a/b/c/d neurons each medulla column in the right optic lobe contains within its bounds. Gray columns have zero cells. Bottom-left: Fitted ellipses with their semi-major axes as black lines and the color of the ellipse as a function of the ellipse angle. Bottom-right: Number of columns spanned by each MeTu4 neuron in the right hemisphere.

Based on the connectivity in medulla and AOTUsu_M, we categorized MeTu4 into four subgroups, named MeTu4a/b/c/d (Fig. 5a). The dendrites of MeTu4a cells (N=69 left and N=60 right) cluster densely in the dorsal half of the medulla but also extend ventrally (Fig. 5a_i_, m_i_) with unique arborization in two medulla layers (M6 and M7; Fig. 5b, d). Despite their dorsal location, they form no synaptic connections with polarized light-sensitive photoreceptors or DRA neurons. MeTu4b neurons (N=8 left and N=12 right) are remarkable for their unique dendritic arrangement: they span a rather small area in the equator, mostly in the posterior-medial part of the medulla that represents the frontal central visual field (because of the cross-over connections from lamina to medulla along the anterior-posterior axis; Fig. 5a_ii_, m_ii_). This unique anatomical feature suggests that MeTu4b neurons are specialized for detecting a currently unknown visual feature that is meaningful only in front of the fly. MeTu4c neurons (N=41 left and N=48 right) span the entire dorsal half of the Medulla (Fig. 5a_iii_, m_iii_), whereas MeTu4d neurons (N=19 left and N=18 right) cluster exclusively in the ventral half of the medulla (Fig. 5a_iv_, m_iv_), ideally positioned to detect features in the ventral visual field. Both MeTu4c and MeTu4d receive nearly identical input from a wide variety of interneurons, including those conveying information from other brain areas, e.g., the SPS (Fig. 5e_iii-iv_, k-l).

Like all other MeTu types, axonal projections of all MeTu4 neurons maintain anterior-posterior retinotopy in the AOTUsu_M along the ventral-dorsal axis (Fig. 5j), in contrast to a previous report [35]. MeTu4a/b/c also have presynaptic connections in lobula (Fig. 5a_i__-iii_, S5h), but these connections do not contribute to the AVP and were thus excluded from further analyses. In the AOTUsu_M, all MeTu4a neurons are presynaptic to TuBu03; some are also presynaptic to TuBu04. MeTu4b neurons are presynaptic to TuBu02 neurons. Both MeTu4c and MeTu4d subtypes are primarily presynaptic to TuBu05 (Fig. S5k), but MeTu4d additionally makes presynaptic connections with TuBu02. MeTu4b and MeTu4c receive the main interhemispheric connections within AOTUsu_M (Fig. S1e, S2g-h). MeTu4b receives strong input from AOTU046 but does not provide reciprocal input into AOTU046. In contrast, MeTu4c is strongly and reciprocally connected to AOTU046. Finally, MeTu4d receives no input from AOTU046, and provides only weak input to AOTU046. Additionally, MeTu4d receives weak input from TuTuB_a.

### Linking visual features to information channels along the AVP

A common pattern across all AVP channels is the convergence of MeTu neurons onto a significantly smaller number of TuBu neurons (Fig. S1l, S2c, S3c, S4c, S5c, S6b). In this transformation, each TuBu neuron integrates information from a large area of the visual field, which is a prerequisite for extracting simple spatial features. TuBu neurons also receive strong input about contralateral visual field from TuTu neurons (Fig. S2g-i, S2g-i). The next step in processing––from TuBu to ring neurons––exhibits a re-expansion in the number of neuronal types (from 10 TuBu types to potentially 18 ring neuron types in hemibrain, or 14 ring neuron types in FAFB). The ratio of connections from TuBu to ring neurons (Fig.S1m, S2d, S3d, S4d, S5d) varies between 0.33 and, for some neurons, 4. Thus, the transformation from TuBu to ring may extract several more visual features, which might be more complex than simply summing features that TuBu neurons encode.

With the full synaptic connectome of the AVP from the optic lobe to the central complex, we inferred the visual features that are potentially encoded by any given ring neuron class. This, in turn, represents information compass neurons integrate to develop the fly’s sense of direction.

#### Reconstructing putative receptive fields for all ring neuron classes

To link visual features to channels along the AVP, we reconstructed the putative receptive fields for all ring neuron classes and categorized their visual inputs based on the visual information they process.

We traced the synaptic connections of individual ring neurons of each type backwards along the pathway to the level of MeTu neuron dendrites in the medulla (Fig. 6, Video 1-3). To quantify the putative visual area to which each neuron likely responds, we defined every medulla column based on Mi1 neurons (Fig. 1h) and transformed this map into putative eye maps of *Drosophila melanogaster* [73, 74]. Then, for each ring neuron, we back-traced the upstream connections in two ways: One followed TuBu to MeTu connections (putative excitatory direct pathway; Fig. 6a, S6c) and the other followed TuBu to TuTu to ipsi- and contralateral MeTu connections (putative inhibitory indirect pathway; Fig. 6b, S3g-i). We used the dendritic arborization in the medulla for each pathway to estimate the area of excitation (direct pathway) or inhibition (indirect pathway). We combined them into a single putative receptive field (Fig. 6c), which we further analyzed to obtain the outline of the excitatory field. We then combined outlines of the same type of ring neurons to illustrate the visual area that the population of ring neurons covers (Fig. 6e). Finally, we determined the degree of overlap of the excitatory field (Fig. 6e). Figure 6e is a compilation of all ring neuron types in the AVP.

**Fig. 6.**
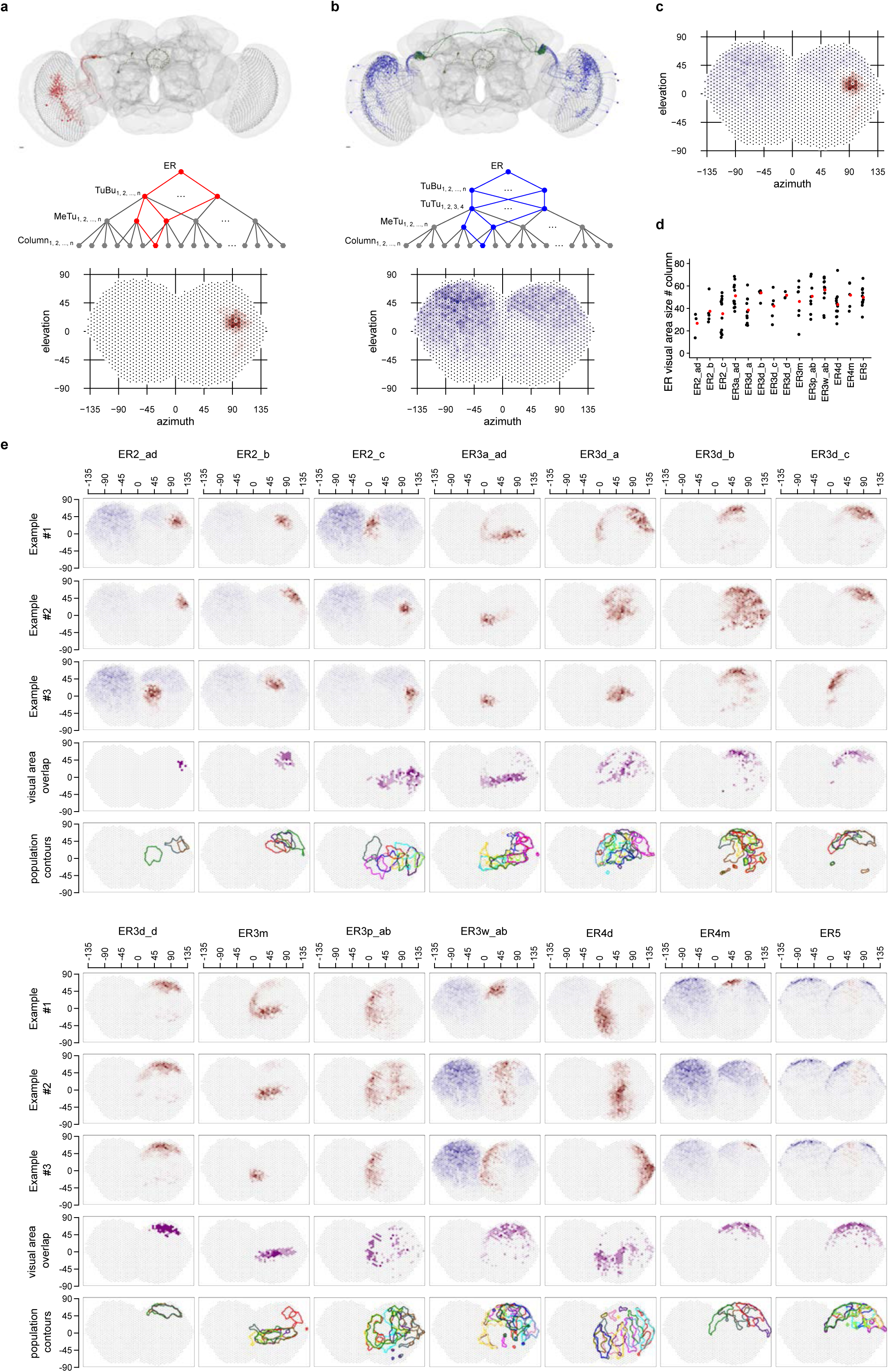
Putative visual areas of ER neurons in the right hemisphere. **a**, Putative excitatory direct pathway. Top: all connected TuBu and upstream MeTu neurons for on given exemplary ER neuron. Middle: Connectivity graph illustration of the excitatory direct pathway. In red are all branches connecting to one given column. Bottom: Resulting putative visual area. **b**, Putative inhibitory indirect pathway. Top: all connected TuBu, upstream TuTu and upstream MeTu neurons for on given exemplary ER neuron. Middle: Connectivity graph illustration of the inhibitory indirect pathway. In blue are all branches connecting to one given column. Bottom: Resulting putative visual area. We did not analyze the AOTU046 pathway because its neurotransmitter was not conclusive [65]. **b**, Summed visual areas from a and b. **d**, Visual area size as number of covered columns for all visual ER neurons of the right hemisphere. Red point: population average. **e**, For all visual ER types (columns) of the right hemisphere we show three exemplary visual areas of individual neurons (first three rows), visual area overlap as number of ER neurons present in a given “visual column” (fourth row) and contour outline of the visual area of all neurons of a given type in the right hemisphere (fifth row).

Back-tracing the synaptic pathway starting from ring neuron ER4d revealed that its upstream MeTu1 neurons are aligned vertically in the medulla (Video 1). They cover about 40° azimuth and the entire vertical span (Fig.6e). This vertical arrangement of MeTu1 neurons was consistent across ER4d neurons and covered the entire visual field as a population, like an array of vertical bars (Fig.6e, Fig 7). It suggests that ER4d neurons are selective to vertically elongated visual stimuli or to the location of visual stimuli along the horizontal plane, regardless of the elevation. Such a pathway would be best suited for detecting visual landmarks suitable as a reference for setting a heading.

**Fig. 7.**
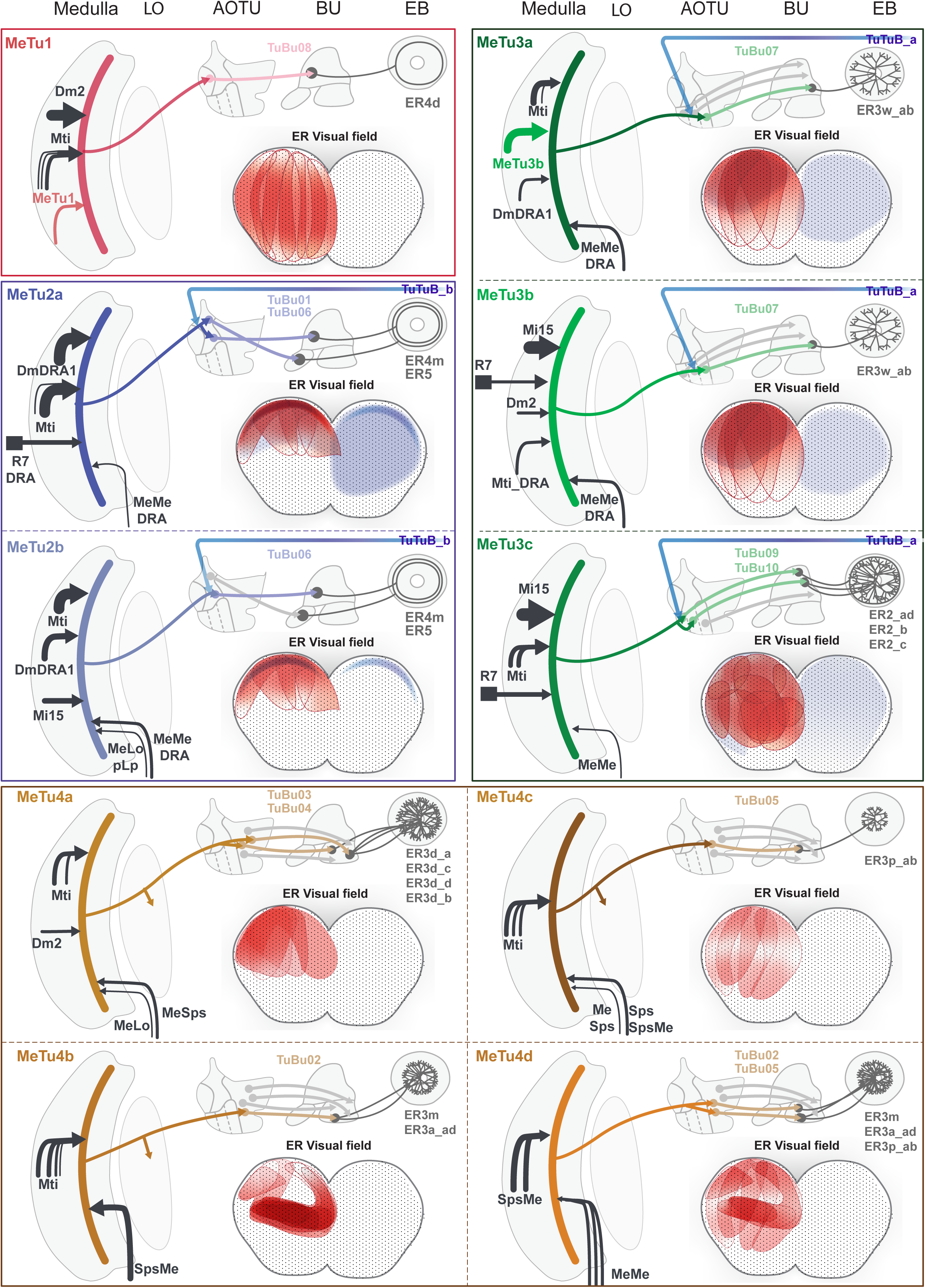
Overviews of Parallel Anterior Visual Pathways. Each panel shows an generalized neural pathway and receptive field of a MeTu subtype. From the left to top right side, there is a diagram of the pathway from the medulla to the EB. Medulla inputs to MeTu are shown on the left of the medulla if they come from the retina or medulla, and on the right if they come from the central brain. Photoreceptor inputs in the retina are shown as squares. The AVP from the MeTu to the TuBu and Ring neurons are shown, as well as if the MeTu has outputs in the lobula or synapses with TuTuB neurons in the AOTU. Receptive fields of the relevant Ring neurons are on the bottom right. Red indicates excitatory input from MeTu neurons, whereas blue indicates putative inhibitory input from TuTuB neurons. Only inhibitory input from TuTuB is shown; AOTU046 is not included because the excitatory/inhibitory nature of the neuron is unknown [65]. The interaction between AVP channels appears to be minimal. In other words, direct interaction between the four major MeTu types is negligible downstream of the medulla.

In contrast, back-tracing starting from single ER2_ad and ER2_b ring neurons revealed that they receive information from MeTu3c neurons with dendrites located in the dorsal medulla (Fig. 6e, Video 2), whereas individual ER2_c neurons receive inputs exclusively from MeTu3c neurons with dendrites in the central medulla (Video 3). Similar to the channel converging into ER4d neurons, the MeTu3c populations upstream of ER2_ad, ER2_b, and ER2_c neurons tile the entire Mi1 map, amounting to approximately 300-330° of visual space [75]. In other words, the ER2 population tiles the visual field two-dimensionally, providing more organizational details of the scene than ER4d neurons (Fig. 6e).

#### Inputs to MeTu provide further information about the processed cues

To distinguish between the potential visual features processed by ER4d and ER2 neurons, we used the previously reconstructed photoreceptor inputs upstream of a small number of MeTu1 and MeTu3c neurons [32]. As described before, the main upstream input of MeTu1, the distal medulla cell Dm2, receives preferentially inputs from UV-sensitive pale photoreceptors (81%) [32], suggesting that ER4d neurons process UV stimuli. On the other hand, the main upstream input of MeTu3c, the medulla intrinsic cell type Mi15, receives input from green-sensitive yellow R8 photoreceptors (67%) [32], suggesting that ER2 neurons process longer wavelength stimuli.

Unlike the ER4d and ER2 populations, ER4m and ER5 neurons receive strong input from polarization-sensitive channels involving the DRA-specific MeTu2a and MeTu2b neurons, respectively. As described before, MeTu2b exhibits more complex connectivity than MeTu2a and therefore may encode complex features of polarized light, either by combining it with other features or/and by integrating across both hemispheres. Hence, ER5 may process more complex features of polarized skylight, whereas ER4m appears to process skylight polarization alone as a navigational cue, consistent with a previous report [2]. This difference is intriguing because ER5 is involved in circadian rhythms [76, 77] (but see [78]).

ER3w receives input from MeTu3a and MeTu3b neurons, potentially combining skylight polarization information (both from MeTu3a and MeTu3b, Fig.4A and B) and localized visual feature information from some of the dorsal visual field (Fig. 6e). On the other hand, most other ER3 subtypes with dendrites in inferior BU are downstream of MeTu4 subtypes that do not receive columnar input in the medulla. Therefore, the features that putative receptive fields encode are unclear.

Overall, we predict that ring neurons downstream of MeTu1-3 neurons encode diverse information including polarized light (ER4m, ER5, and ER3w), vertical stimuli or azimuthal location of visual features (ER4d), and the 2-D organization of visual scenes, including azimuth and elevation (ER2), a system suitable for processing both the elevation of celestial body (e.g., the Sun) [79] and surrounding 2D environment [68].

## Discussion

Here we report, for the first time, the synapse-level reconstruction of the entire anterior visual pathway from peripheral visual system (medulla) to central complex neurons that compute an abstract internal representation of the world (compass neurons). Across three levels of processing, we observed convergent and divergent connections, and, by back-tracing from ring neurons to MeTu neurons, we inferred the diverse visual features the compass neurons may use to compute the head direction (Fig. 7).

Our analysis of the reconstructed synaptic pathways reveals a subdivision of the fly’s visual field roughly into three regions: A narrow band in the dorsal-most visual field (DRA) [2, 13, 48, 49], the remaining upper visual field (both of which are facing the sky), and the rest of the visual field, which includes the equator and below the equator. Interestingly, the DRA and upper visual fields are occupied by the large majority of MeTu neuron types with overlapping receptive fields (MeTu1, MeTu2a/b, MeTu3a/b, MeTu4a/b); in contrast, the lower and frontal visual fields are subserved by lower numbers of cells and subtypes (innervated by MeTu1, MeTu3c and MeTu4c/d). This distribution suggests that, during navigation, even a behavioral generalist like *Drosophila melanogaster* prioritizes celestial cues, including the skylight polarization pattern, over cues in the lower visual field.

EM connectomic information can be used to predict neural functions [3, 28, 58, 80–85]. Our predictions of ring neuron receptive fields suggest the visual features encoded by ring neurons are coarse in spatial resolution, which may indicate that detailed features or minute variance in the scene are not important to compute the head direction. These results are consistent with previous reports of the several classes of ring neurons that have been physiologically characterized [39–42] and enlarge our knowledge about the visual responses of various ring neuron classes. For example, ER4d and ER2ad/b/c neurons were thought to have similar response patterns to visual input, but our results strongly suggest ER4d neurons are tuned to vertically elongated areas, whereas ER2ad/b/c are sensitive to smaller circular areas two-dimensionally tiling the visual field, while synaptic inputs suggest chromatic sensitivity to differfrom ER4d.

Our comprehensive AVP connectome lays the groundwork for understanding integration across sensory modalities that can now be tested through standard behavioral or physiological assays. For example, although we focused our receptive field prediction on visual cues, receptive fields may also be influenced by other sensory or motor modalities. ER3 neurons in the inferior bulb, for example, were previously reported to be sensitive to self-generated motor signals [40]. Consulting the connectome (Fig. 5 and 7), we find that only MeTu4 neurons upstream of ER3 neurons receive inputs from projection neurons in the superior posterior slope (SPS), which is densely innervated by descending neurons that may send motor commands to the ventral nerve cord. Therefore, it is plausible that the connections from SPS provide an efference copy of motor commands to MeTu4 in medulla, which is conveyed to ER3 ring neurons.

Our analyses also revealed general organizational principles of how visual features are processed. Nine out of the 10 parallel information channels formed by MeTu neurons appear to maintain only azimuth information while discarding information about elevation, a strategy that seems particularly efficient for choosing a heading. It is further corroborated by the fact that at least three of these channels are polarization sensitive, which would provide robust directional information and is an evolutionarily conserved strategy across many insect species [71]. Only the MeTu3c channel encodes both azimuth and elevation – a property that seems ideal for perceiving the 2D organization of the surrounding environment or tracking the celestial body’s position across the day.

Intriguingly, two channels exhibited completely unexpected specializations. First, MeTu4b sample exclusively the frontal part of the visual field, including the zone of binocular overlap; this would be ideal for fixation behavior during navigation. Second, MeTu4d cover only the ventral half of the visual field. We speculate that this might serve to processes optic flow [86] or orienting towards shiny surfaces (e.g., water) that produce horizontally polarized reflections [71, 87], which flies detect and use to adjust their body orientation [88, 89].

Considering the fundamental importance of navigation, it is not surprising that the anatomical structure of the AVP is largely conserved across such diverse insect species as flies, bees, wasps, and beetles [90–92]. It is likely that deep similarities exist in the basic logic of visual feature extraction [6, 22, 25, 54, 55, 72, 93, 94], but despite many studies of the AVP across species, our knowledge about the AVP neurons has been fragmented by the lack of a complete circuit diagram to frame systematic investigations. Our AVP connectome now provides this framework. Thus, we anticipate our results will be invaluable when designing and prioritizing physiological experiments to interrogate the AVP, not just for flies but also for other insects.

Finally, we have known for some time that visual features are processed hierarchically [28]; from dung beetles [4] to mammals [15], animals exhibit specific cue preferences during navigation. Thus, our work of the full AVP reconstruction is essential to mechanistically understand the circuit implementation, as well as the shared functional principles, that underlie the integration and transformation of this information into a heading signal. Moreover, with the ability to dissect detailed circuit dynamics of neural populations using rich genetic tools in flies, this connectome paves the way to probe and understand the computational mechanisms of visually guided navigation.

## Methods

### Flywire

#### Overview

Our study analyzed the Full Adult Fly Brain (FAFB), an adult female *Drosophila melanogaster* brain imaged at the synaptic level with serial section transmission electron microscopy [27]. We used the Flywire interface, which auto-segmented FAFB EM data to construct 3-dimensional segmentations of individual neurons [64]. To reconstruct desired neurons, we first identified relevant axons, dendrites, and branches. Possible errors by the auto-segmentation were mainly unfinished branches caused by missing EM slices or incorrect connections caused by shifted EM slices. Additionally, some neurons had darker cytosols in the EM data, possibly due to neuronal damage during the dissection process [95], and were therefore not as well-constructed by the auto-segmentation. We manually corrected each of these errors.

#### Dense EM Reconstruction

To find all MeTu, TuBu, TuTu, and AOTU046 neurons in the anterior visual pathway (AVP), we densely reconstructed the anterior optic tubercle (AOTU) small unit by scanning through every layer of EM in the neuropil volumes and proofreading all neurons composing them (disregarding twigs). Ring neurons were identified be means of TuBu downstream connectivity [65]. After each of these neurons was proofread, we classified them and compiled lists of their coordinates for further analysis.

#### Computational Tools

##### Region Boundaries

Regions were distinguished in our study in order to limit synapsesto specified neuropils. These regions included ME_L, ME_R, LO_L, LO_R, AOTU_L, AOTU_R, BU_L, BU_R, and EB. Using *SciPy*’s spatial module, we created Delaunay tessellations using a set of Flywire coordinates in order to determine whether synapses were contained within the given regions. The sets of points were not a comprehensive boundary box of individual neuropils, but rather formed polyhedra that contained the regions of interest of the relevant neurons. The coordinates were selected with the help of Flywire’s annotation lines to ensure that all neurons’ synapses were incorporated. In the case of the medullas, in which the Delaunay tessellation incorporated some lobula synapses as well, the ipsilateral lobula region was subtracted.

#### AOTUsu subdivision in comparison with previous studies

The connectome of the AVP, which revealed four major MeTu types, clarifies discrepancies in previous literature. MeTu_im in Omoto & Keles et al. [34] appears to be MeTu4 because DALcl2d TuBu neurons project from AOTU_im to BUi. In addition, based on spatial organization, MeTu_lc, lp, and la/il in Omoto & Keles et al. may correspond to MeTu1, 2, and 3, respectively, though the AOTUsu map is slightly different from our study. A critical discrepancy we could not resolve was TuBu_a in Omoto & Keles et al. They described that TuBu_a projects from AOTU_il to the Bua which we did not observe in the FAFB dataset. In Hulse, Haberkern, Franconville and Turner-Evans et al. [28], the TuBu neuron type innervating BUa is TuBu01, which are located in AOTUsu_PC, downstream of MeTu2. However, Omoto & Keles et al. claims these TuBu neurons project from the AOTUsu_A to the BUa. We believe this discrepancy is due to the lower resolution of light microscopy, and TuBu_a should be re-classified.

Timaeus et al. [35] divided the AOTUsu into five subdomains, separating AOTUsu_a into lateral and anterior central parts. They state that R7 may be upstream of MeTu_la, MeTu_ca, and MeTu_cp (MeTu3 and MeTu2, respectively), which agrees with what we found. However, they only found TuBu neurons projecting from the AOTUsu_la/lp/ca/cp to the BU_s and the AOTUsu_m to the BUi, meaning they did not discover TuBu01. Finally, they claimed that MeTu_m (MeTu4) dendrites also project to medulla layers 2 and 8, which was inconsistent with what we found in the FAFB dataset.

Tai et al. [56], unlike the other two papers, found four subdomains of the AOTUsu (L-AOTU_1-4_), which are connected linearly from the edge of the AOTUlu to the lateral-most edge of the AOTUsu. The respective MeTu neurons in these regions were called MT_1-4_ (not corresponding to our study’s MeTu1-4). This study only showed an anterior view of the AOTU, and as such, it is possible that they did not find the AOTUsu_PC, which is obscured by the AOTUsu_A from the anterior side. In this case, the corresponding regions are AOTUsu_M (L-AOTU_1_), AOTUsu_A (L-AOTU_2-3_), and AOTUsc_PL (L-AOTU_4_). The corresponding MeTu neurons are thus MeTu4a/b/c/d (MT_1_), MeTu3c (MT_2_), MeTu3a/b (MT_3_), and MeTu1 (MT_4_).

#### Synaptic Connectivity Matrices

Synaptic connectivity between neurons was found using automatic synapse detection [65]. For all our connectivity analysis we used a cleft score of >= 50 and excluded autapses and synapses to the background segmentation. Two types of connectivity matrices were generated throughout the study: individual neuron weight matrices (purple) and neural type weight matrices (green). For the individual neuron weight matrices, the number of synapses between each neuron was first calculated. To determine the relative weight within the given region, this quantity was divided by the postsynaptic neuron’s total number of synapses in the region. Neurons with fewer than 3 total regional connections were removed. Additionally, a few outliers found to connect to specific neurons than others were removed: Among MeTu interconnectivity, there were 1 MeTu1_R, 1 MeTu2a_R, 2 MeTu4a_L, and 5 MeTu4a_R outliers, and among Ring interconnectivity, there was 1 ER3d_b_L outlier.

The ordering of the neurons within the connectivity matrices was based on the location of TuBu neurons within the AOTU: TuBu neurons with dorsal dendrites were first (which corresponds to the front of the fly’s visual field), and TuBu neurons with ventral dendrites were last. Most TuBu types form a dorsal-ventral line within the AOTU, which made this ordering possible. Both MeTu and ring neurons were ordered in groups based on which of these TuBu neurons they were most connected to (MeTu neurons presynaptically in the AOTU and ring neurons postsynaptically in the bulb). Within the groups they were ordered by how many synapses they shared to that TuBu neurons.

Neural type weight matrices show the connections of whole classes of neurons. First, the total number of synapses between all presynaptic and postsynaptic neurons of the respective given types was calculated. Then, these quantities were divided by the total number of synapses of all postsynaptic neurons of the given type within the region. This gave a measure of the total synaptic weight between the two types.

#### 3-Dimensional Rendering

3D renderings were either generated in Blender with neuron meshes retrieved using the Python CloudVolume package or in R with the rgl and fafbseg package.

##### Medulla columns and layers

We identified all Mi1 neurons, a unicolumnar celltype, in both hemispheres as a proxy for individual medulla columns as Mi1 neurons are present in each medulla column and span the entire distal-proximal axis of the medulla from M1 to M10. For each Mi1 neuron we performed a PCA on all pre- and postsynaptic sides of the neuron (Fig. 1h). PC1 corresponds to the distal-proximal axis of the column. The upper and lower boundary of each column is defined as the 0.03 and 0.97 percentile of synapses on the distal-proximal axis.

Medulla layers are based on the average synapse distribution of Mi1, Mi4, L1, L2, L3, L5, Dm8 and T4 neurons along the distal-proximal axis in three exemplar columns. Layer M1: [-3.9% — 5.5%]; layer M2: [5.5% — 17.1%]; layer M3: [17.1% — 30.8%]; layer M4: [30.8% — 34.0%]; layer M5: [34.0% — 43.2%]; layer M6: [43.2% — 50.1%]; layer M7: [50.1% — 63.1%]; layer M8: [63.1% — 75.4%]; layer M9: [75.4% —92.4%]; layer M10: [92.4% — 102.2%].

#### MeTu Analysis

##### MeTu Classification

We describe MeTu types (labeled with numbers: MeTu1 - MeTu4) and MeTu subtypes (labeled with lowercase letters: exp. MeTu2a). MeTu1/2/3 were previously called MC61 [96] and MeTu4 was called MC64 [28]. MeTu2 was also called MeTu-DRA [32]. The location of axons and dendrites of MeTu (Fig. 1c), TuBu (Fig. 1c), TuTu (Fig. S3g), and AOTU046 (Fig. S2h) neurons maintain specific patterns of localizations within the AOTUsu [28], through which we determined four distinct regions (Posterior Lateral, Posterior Central, Anterior, and Medial). The axonal boutons of each MeTu neuron terminates within one of these four areas, so we classified MeTu1, MeTu2, MeTu3, and MeTu4 as types. Between the left and right hemispheres respectively, there are 121 and 124 MeTu1, 50 and 50 MeTu2, 145 and 129 MeTu3, and 137 and 138 MeTu4. There is one neuron on the right side whose axonal tract terminated before projecting to the medulla. It was labeled MeTu_incomplete_R and was excluded from further analysis.

Analysis of morphology, up- and downstream connectivity as well as spatial distribution in the medulla revealed distinct MeTu subpopulations within the MeTu2, MeTu3 and MeTu4 types which led us to define MeTu subtypes.

MeTu1 forms a homogenous neuron population in terms of morphology, and up- and downstream connectivity without any distinctive features which would allow any further subtyping(Fig. 2s). MeTu2 is upstream of two TuBu types, TuBu01 and TuBu06. MeTu2a is connected to both, TuBu01 and TuBu06, with a preference for TuBu01 whether MeTu2b is connected to TuBu06 with very few synapses onto TuBu01 (Fig. 3w).

We found three MeTu3 subtypes: MeTu3a/b/c. MeTu3a has flat dendrites and lacks presynaptic connections to Mi15, while MeTu2b/c has vertical dendritic protrusions and connects to Mi15. (MeTu3a was specifically classified as MeTu3 that has ≤13 synapses with Mi15 neurons.) Note that MeTu2a/b cell bodies are located closer to the medulla equator, whereas MeTu3a cell bodies are found above the center of the branching (data not shown). Within the AOTUsu, all MeTu3a project to TuBu07. MeTu3b is strongly connected to TuBu07, and MeTu3c is most strongly connected to TuBu09 and TuBu10. To further analyze this distinction, we compared their postsynaptic weights with Mi15, Mti_unknown_1, Mti_DRA2, and MeMeDRA. MeTu cells sensitive to skylight polarization have so far been physiologically characterized in Drosophila [2], and a careful comparison between their light microscopic data and our connectomic reconstruction identifies these cells as MeTu 2b and MeTu3a.

MeTu4 is generally morphologically distinct from other MeTu types because neurons contain boutons within the lobula. However, light microscopy suggested there is a subtype that does not have these boutons (Fig. 4k+5). We as well found a MeTu4 population without lobula boutons and few loblua synapses (15 pre- and postsynapse), which we named MeTu4d. MeTu4d additionally only arborizes within the ventral half of the medulla.

We further grouped MeTu4 neurons with lobula boutons in distinctive subtypes based on morphology, spatial distribution in the medulla, and downstream TuBu connectivity. MeTu4a has two layers of dendrites in the medulla, more densely arborizes in the dorsal half of the medulla, and are presynaptic to TuBu03 and TuBu04. MeTu4b has single-layered dendrites, only occupies the posterior-center of the medulla in both hemispheres, and are presynaptic to TuBu02. MeTu4c also has single-layered dendrites, arborizes across the entire medulla, and are presynaptic to TuBu05.

We sought to provide light microscopic evidence in the form of cell type-specific driver lines, corroborating the existence of genetically defined subclasses of visual projection neurons described in this study [46, 47, 97–99].

UMAP in Fig. 1f_i_ is based on connectivity to up- and downstream partners as features. We selected a total of 84 neurons and identified all their presynaptic partners. We selected these neurons to ensure that at least five neurons upstream of each TuBu type were included and that the selected specimens covered the medulla space evenly. We identified all upstream partners with 4 or more synapses. Downstream neurons include 13 types (all TuBu types, TuTuA, TuTuB and AOTU046), and Upstream neurons include 28 types (all top five connected neuron types of all MeTu subtypes; Table 1). UMAP in Fig. 1f_ii_ is only based on the 28 upstream types. All connectivity types are also shown in Fig. 1g.

##### Proofreading Rounds

For a subset of MeTu neurons we increased proofreading quality by increasing the rounds of detailed proofreading [64]. We used the right optic lobe because the left optic lobe has a partially detached lamina and parts of the posterior side of the medulla are distorted. [27]. We chose 113 of the 441 right MeTu neurons to undergo multiple rounds of proofreading. Originally, 101 neurons were chosen randomly with the same relative ratios of MeTu1-4s as in the population: 28 MeTu1, 12 MeTu2, 30 MeTu3, and 31 MeTu4. When we later discovered subcategories of the neurons, we wanted at least 5 of each subtype. In the end, we proofread the following 113 neurons: 29 MeTu1, 7 MeTu2a, 5 MeTu2b, 6 MeTu3a, 13 MeTu3b, 16 MeTu3c, 13 MeTu4a, 5 MeTu4b, 14 MeTu4c, and 5 MeTu4d.

Each of these neurons underwent three rounds of proofreading, and volumetric comparisons were performed to determine the differences in accuracy between the three rounds. The first round was the cursory proofreading that was done to all 441 MeTu neurons. The next two rounds were split between the two proofreaders (DG and EK). Each proofreader densely proofread half of the 113 for the second round, and then switched and worked on the other half for the third round.

##### Upstream Connections

We used automatic synapse detection to find presynaptic partners of the proofread MeTu neurons. As stated in the proofreading section, we picked them based on the ratio of the entire population, with a minimum of 5 neurons per type. Additionally, as with the proofreading rounds, we only looked at neurons on the right side. Because several neurons contain a darker cytosol and are not segmented well in Flywire, we left out any of those neurons in favor of normal neurons. Thus, we analyzed the following 84 neurons: 18 MeTu1, 5 MeTu2a, 5 MeTu2b, 6 MeTu3a, 9 MeTu3b, 11 MeTu3c, 10 MeTu4a, 5 MeTu4b, 10 MeTu4c, and 5 MeTu4d.

For each neuron, we identified each presynaptic partner with ≥4 synaptic connections. Many partners had been classified in previous studies, but we found new medulla tangential intrinsic (Mti) neurons. We also found additional neurons that linked multiple locations. We labeled them based on the regions they arborized in, beginning with their dendrites (MeSps, MeLo, SpsSpsMe). If the cells had not been previously characterized, we labeled them “_unknown_#” (Mti_unknown_1, Mti_unknown_2, etc.) based on morphological groups. The lower numbers had more connections to the MeTu population. We provide a spreadsheet of these neurons with corresponding connectomic names recently proposed by the FlyWire team and available in Codex [53].

##### Synapse Density

MeTu and TuBu synapse density maps in the AOTU were created from three angles: from the dorsal side looking towards the ventral side, from the anterior side looking towards the posterior side, and from the lateral side looking towards the medial side (Fig. S2f, S3f, S4h, S5f). Each of these views were rotated 30° along the anterior-posterior axis. Each map was created by finding all of the connections within small volumes, each 40nm by 40nm by the length of the AOTU along the viewpoint axis. When the number of connections were computed, they were subjected to a Gaussian blur with a sigma value of 10. Color maps were then created based on the relative values, with higher values having higher opacity. Demonstrative synapse maps were created as well (Fig. 1c). These were subjected to a Gaussian blur with a sigma value of 4, and did not vary in opacity based on synapse density.

##### Neurotransmitter Predictions

We use the neurotransmitter prediction described in a recent study [100]. We calculate the average neurotransmitter probability across all presynapses of an individual neuron (Fig. S2i, S3h_i-ii_, S6c_i-x_).

#### AVP Analysis

##### AVP Classification

The recent connectomic analysis [28] of the hemibrain [37, 38] provided full classifications of TuTu, TuBu, and ring neurons which we adopted in this study. This study gave detailed classifications to 17 bulbar Ring neurons and 5 LAL ring neurons. Of the 17 bulbar neurons, there are 11 distinct morphologies, and we classified the FAFB neurons as follows: ER2_abd, ER2_c, ER3a_ad, ER3d_acd, ER3d_b, ER3m, ER3p_ab, ER3w. ER4d, ER4m, ER5. The study also described patterns of interconnectivity between Ring neurons, and using synaptic analysis we distinguished ER2ad and ER2b, and ER3d_a, ER3d_c, and ER3d_d. There are multiple morphologies of ER2c neurons (which is consistent with hemibrain), but we did not further subcategorize these neurons. However, some connectivity patterns are not consistent between the hemibrain and FAFB, so we did not subclassify all neurons to the same level of detail. In the instances of ER2_a and ER2_d; ER3a_a and ER3a_d; ER3p_a and ER3p_b; and ER3w_a and ER3w_b, we maintained their names as ER2_ad, ER3a_ad, ER3d_acd, ER3p_ab, and ER3w_ab. In the hemibrain, TuBu neurons were classified based on their downstream ring neuron partners. After classifying all the corresponding ring neurons, we similarly grouped the TuBu neurons as TuBu01-10. There are three ambiguous TuBu neurons. One TuBu in the right hemisphere is upstream of an ER2c neuron but is located in line with other TuBu09 as opposed to TuBu10, which are generally upstream of ER2c. We labelled this neuron as TuBu09 because of its AOTU location. Another TuBu nbeuron in the right hemisphere has the dendritic morphology of a TuBu04 and is downstream of MeTu4a, but is upstream of ER3p_ab. We classified it as TuBu04 as opposed to TuBu05. One neuron in the left hemisphere has a normal microglomerulus partnered with an ER3a_ad and two ER3m neurons like TuBu02 neurons. However, this neuron projects to the SPS, as opposed to the AOTU. Because there is no other neuron in this dataset or hemibrain with this projection pattern, we determined that it may have been a developmental error and labelled it TuBu_misc_L, only including it in connectivity tables between TuBu and Ring neurons.

We identified TuTub_a and TuTub_b based on morphology. There is one of each type per hemisphere. There are four AOTU046 neurons, with dendrites in one SPS and axons in both AOTU and both bulbs. We used these eight neurons for bihemispheric type connections. The quantity of each of these neuron types is consistent with hemibrain [28].

##### Bihemispheric Connections

Connectivity diagrams of bihemispheric neurons are based on type connectivity matrices from the right hemisphere (Fig. S2g). Each arrow represents the weight of the postsynaptic neuron type’s connection to the presynaptic neuron type. Only weights ≥0.05 were represented as arrows. Arrow thickness was determined linearly based on the weight.

Bihemispheric neuron diagrams in Fig. S2h, S3gi-ii are made using neurons on from the right hemisphere. Pie charts within the figures show the relative amount of presynaptic (red) and postsynaptic connections (cyan) of the neuron. Within the AOTU, these only include connections between the bihemispheric neurons and MeTu and TuBu neurons, as those are the only neurons in the AOTU in which the subregion is known. Within the bulb, the connections shown are between AOTU046 and TuBu and ring neurons. Within the SPS, the connections are between AOTU046 and all SPS neurons. The relative size of the pie charts refers to the quantities of bihemispheric synapses in each subregion. In the case of AOTU046, these were calculated by averaging the two neurons on the right side. Lines are drawn to subregions that have ≥100 synapses.

#### Mapping medulla columns to ommatidia coordinates

For each medulla column we manual assigned the neighboring columns along the vertical, horizontal, p and q axis (Fig. 1h sixth panel). The column maps were mirrored along the dorsal ventral axis to reflect the inversion from retina to medulla along antero-posterior (A-P) retinotopic axis through the first optic chiasm (OCH1). We assume an interommatidial angle of 5.5°. The resulting retinal maps are centered on 0° elevation. Negative azimuth values correspond to the left eye and positive values to right eye. We assume a zone of binocular overlap of eight columns total (four columns of the left eye extend into the right hemisphere on the anterior side and four columns of the right eye extend into the left hemisphere on the anterior side).

##### Ring neuron visual area

For each “visual column” of a given ring neuron we calculate the putative excitatory value as the sum of all weighted branches connecting the ring neuron via TuBu and MeTu neurons to the “visual column” (Fig. 6a middle panel). Each branch is the product of synaptic weights of TuBu neuron to ring neuron connection, MeTu neuron to TuBu neuron connection and MeTu neuron medulla column occupancy. MeTu neuron medulla column occupancy is calculated as the fraction of presynaptic sides closed to the column. Putative inhibitory values are the sum all weighted branches connecting the ring neuron via TuBu, TuTuB and MeTu neurons to the “visual column” multiple by −20 (Fig. 6b middle panel). “Visual column” values for figure panels showing the combined putative excitatory and inhibitory visual areas were calculated as the sum of the inhibitory value and excitatory value (Fig. 6c and e).

#### Hemibrain Comparison

The hemibrain dataset contains the entire central complex of the *D. melanogaster* brain, but only extends to include the AOTU pf the left hemisphere. Therefore, it contains two sets of ring neurons, one set of TuBu neurons, and only the boutons of one set of MeTu neurons. The MeTu neurons were named MC64 or MC61.

We used the Python module *neuprint* to look at the MC61 and MC64 that are presynaptic to the previously defined TuBu neurons. We first distinguished MeTu1-4 based on their respective TuBu types. All MC64 are MeTu4, but a small population of MeTu4 ais MC61 instead. We plotted the number of synapses within the lobula among MC61 and MC64 to determine that this distinction was due to differences in the number of lobula connections (data not shown). We distinguished MeTu4 the same way as FAFB, where fewer than 15 synapses in the lobula signified MeTu4d. We classified all other MeTu subtypes using their downstream TuBu partners. The only classification we were unable to make was that of MeTu3a and MeTu3b, as they were separated using upstream connections in the medulla, which the dataset did not include. We labeled these neurons as MeTu3ab, and adjust the FAFB one in comparison plots. After classification, there are 127 MeTu1, 39 MeTu2a, 14 MeTu2b, 64 MeTu3ab, 86 MeTu3c, 68 MeTu4a, 13 MeTu4b, 41 MeTu4c, and 17 MeTu4d.

After obtaining all AVP neurons in Flywire and neuprint, we compared the relative numbers of neurons among four hemispheres with Ring and bihemispheric neurons, and three hemispheres with TuBu and MeTu neurons. We noticed discrepancies among TuBu and ring neuron counts, so we compared the ratios of ring to TuBu counts in the three hemispheres (Fig. S1n-s).

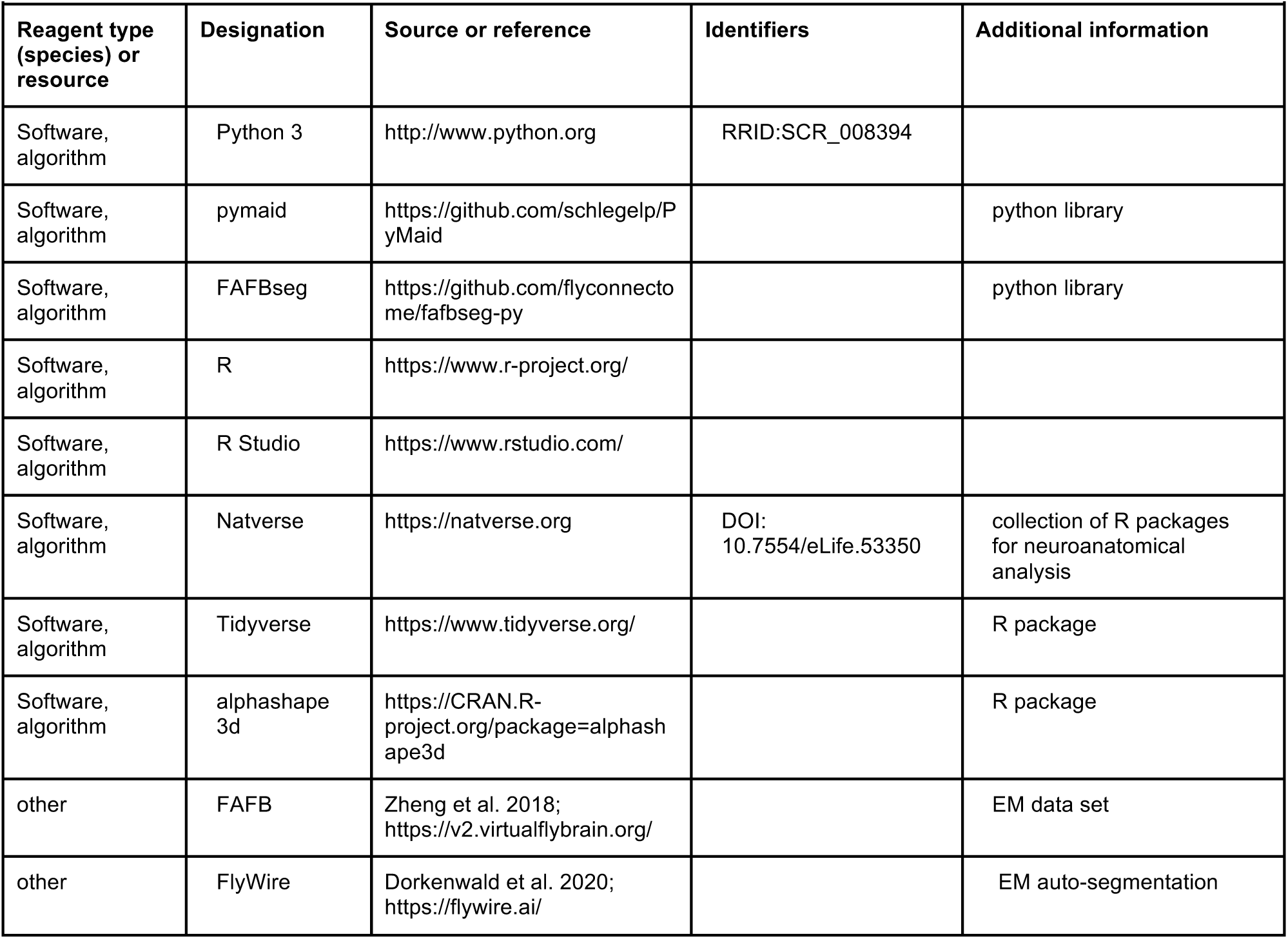
Key Resources Table.

## Supporting information

Data set 1

Data set 2

Neuron names

FlyWire consortium editing record

## Acknowledgments

DG, LH, and SK were supported by the National Eye Institute of the National Institutes of Health (DP2EY032737). The content is solely the responsibility of the authors and does not necessarily represent the official views of the National Institutes of Health. DG, LH, and SK were also supported by Searle Scholars Program, Sloan Research Fellowship, and Klingenstein-Simons Fellowship in Neuroscience. EK, GS, and MW were supported by Deutsche Forschungsgemeinschaft (DFG) grant WE 5761/4-1, SPP 2205, FOR 5289, and AFOSR grant FA9550-19-1-7005. AN and GR were supported by Howard Hughes Medical Institute.

We thank M. Reiser for helping out in the early stages of this project, J.-M. Knapp for valuable discussions and comments on the manuscript, and B. Gorko for rendering videos. We thank the Princeton FlyWire team and members of the Murthy and Seung labs, as well as members of the Allen Institute for Brain Science, for development and maintenance of FlyWire (supported by BRAIN Initiative grants MH117815 and NS126935 to Murthy and Seung). We also acknowledge members of the FlyWire consortium for their contribution to the reconstruction of neurons we used in this work. Specifically, F. Collman in the Collman lab; Nseraf, AzureJay, TR77, Krzysztof Kruk, st0ck53y, annkri, Kfay, bl4ckscor3, JousterL, Mavil, I. Georgiev, Andrearwen, a5hm0r in the Eyewire team; G. Linneweber in the Linneweber lab; V. Sane, A. Yadav, R. Rana, A. Pandey, I. Tamimi, G. Badalemente, L. Serratosa, Y. Yin, M. Santos, P. Schlegel, D. Kakadiya, Z. Vohra, S. Sisodiya, C. Nair, I. Salgarella, D. Sapkal, A. Javier, D. Patel, G. Jefferis, S. Fang, C. Dunne, Y. Patel, N. Patel, E. Munnelly in the Jefferis lab; L.S. Capdevila in the Jefferis and Wilson labs; J. Hsu in the Jefferis and Waddell labs; T. Yang, M. Flynn, A. T, S. Koskela in Jaenlia and the Reiser lab; hanetwo in the J. Kim lab; L. Guo in the Simpson lab; M. Bui, S. Cho in the Colodner lab; J. Eckhardt in the Murthy lab; Z. Zheng in the Seung lab; R.A. Candilada, N. Hadjerol, R. Tancontian, Z. Lenizo, J. Bañez, A. Dagohoy, S. Serona, S.M. Monungolh, R. Salem, A.T. Burke, D. Bland, K.P. Willie, A.J. Mandahay, J.A. Ocho, D.J. Akiatan, K.J. Vinson, N. Panes, J. Laude, J. Dolorosa, Philip, M. Lopez, Clyde, J. Salocot, M.L. Pielago, C. Martinez, B. Silverman, R. Willie, J. Saguimpa, A.M. Gogo, M. Manaytay, M. Albero, D. Bautista, J.D. Asis, C. Pilapil, J. Seguido, S. Yu, M. Pantujan, J. Hebditch, E. Tamboboy, J. Gager, Celia D, M. Sorek, M. Moore, C. McKellar in the Murthy and Seung labs; M. Selcho in the Selcho lab; J. Ch, M. Ioannidou, A. Oswald, L. Lörsch, A. Bast, S.M.M. Obando in the Silies lab; L. Walter, X. Zhong, P.G.A. de Antón, Emre D., Solenne P. in the Wernet lab; Q. Vanderbeck, T. Okubo in the Wilson lab; A.S. Diez in the Behnia lab; B. Huang, T. Crahan in the Kim lab.

## Author Contributions

SK and MW conceived the study. DG, LH, and EK collected data. DG and EK analyzed the data. GS drew AVP schematics. AZ helped spatially mapping medulla columns. AN and GR generated fly lines and light microscopy images. SK, MW, DG, and EK wrote the manuscript with input from everyone.

## Data availability

All raw data is available at flywire.ai. Our Data Sets 1 and 2 provide neuron IDs.

## Code availability

The code for analyzing the data will be provided as a github repository at the time of publication.

---

**Fig. S1.**
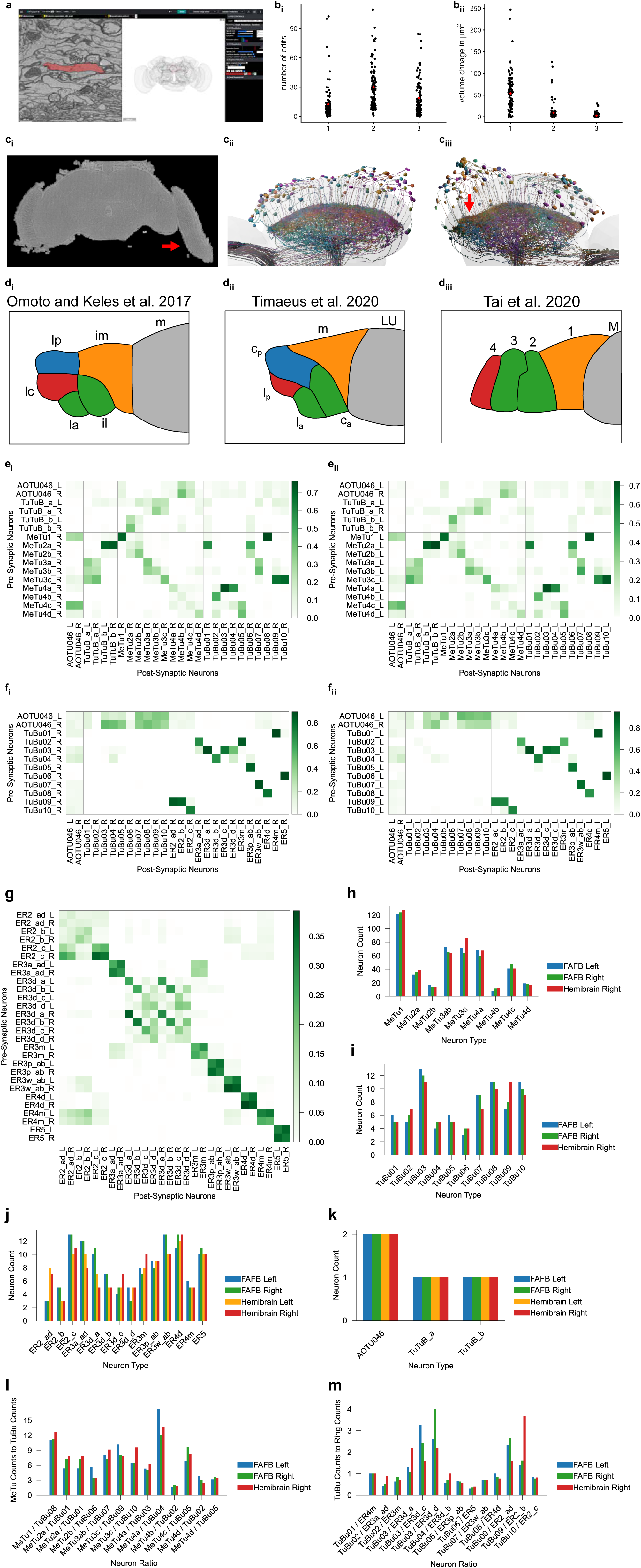
Methods of analysis of the AVP and AOTU_SU. **a**, Diagram of Flywire. Left: multiple stacked EM layers. Right: Image of the Flywire interface. **b**, Quality control through three rounds of proofreading among 113 MeTu neurons of the right hemisphere. The total number of edits per neuron per round (**b_i_**) and the change in volume from before to after each proofreading round (**b_ii_**). **C_i-iii_**, EM data quality of the left and right optic lobe . (**c_i_**) EM slice of the fly brain, note the partially detached lamina (arrow) on the left optic lobe. (**c_ii_**) and (**c_iii_**) MeTu1 neurons of the right and left optic lobe respectively viewed from the dorsal side, note the uneven image alignment on the posterior side (arrow) of the left optic lobe (**c_iii_**). **d_i-iii_**, Comparison of different illustrationsof the AOTUsu subregion [35, 41, 56]. **e_i-ii_**, Synaptic weight matrix of bihemispheric neuron types and MeTu subtypes to themselves and TuBu types on the right (**e_i_**) and left (**e_ii_**) hemisphere. **f_i-ii_**, Synaptic weight matrix of bihemispheric neuron types and TuBu types to themselves and Ring neurons, on the right (**f_i_**) and left (**f_ii_**) hemisphere. **g**, Synaptic weight matrix of all visual Ring neuron subtypes. **h-k**, Comparing neuron counts between both hemispheres in the FAFB and applicable hemispheres in the hemibrain datasets, of MeTu (**h**), TuBu (**i**), Ring (**j**), and bihemispheric (**k**) neurons. Note the difference between FAFB and hemibrain data. For example, in FAFB, there are only six ER2_a/d neurons (3/side) while the hemibrain has 15 of these neurons (eight on the left, seven on the right) (**j**). Because there were no verifiable differences among these neurons, we categorized them as a single group. Similarly, although the total number of ER3a_ad neurons was similar between FAFB and hemibrain data, we were unable to identify distinct features to differentiate ER3a_a and ER3a_d in FAFB (**j**). Thus, we combined them into a single group as well. We did the same with ER3p_ab and ER3w_ab, as potential subtypes were similarly indistinguishable (**j**). **l-m**, Comparing the ratios of MeTu to TuBu (**l**) and TuBu to Ring (**m**) between both datasets in applicable hemispheres.

**Fig. S2.**
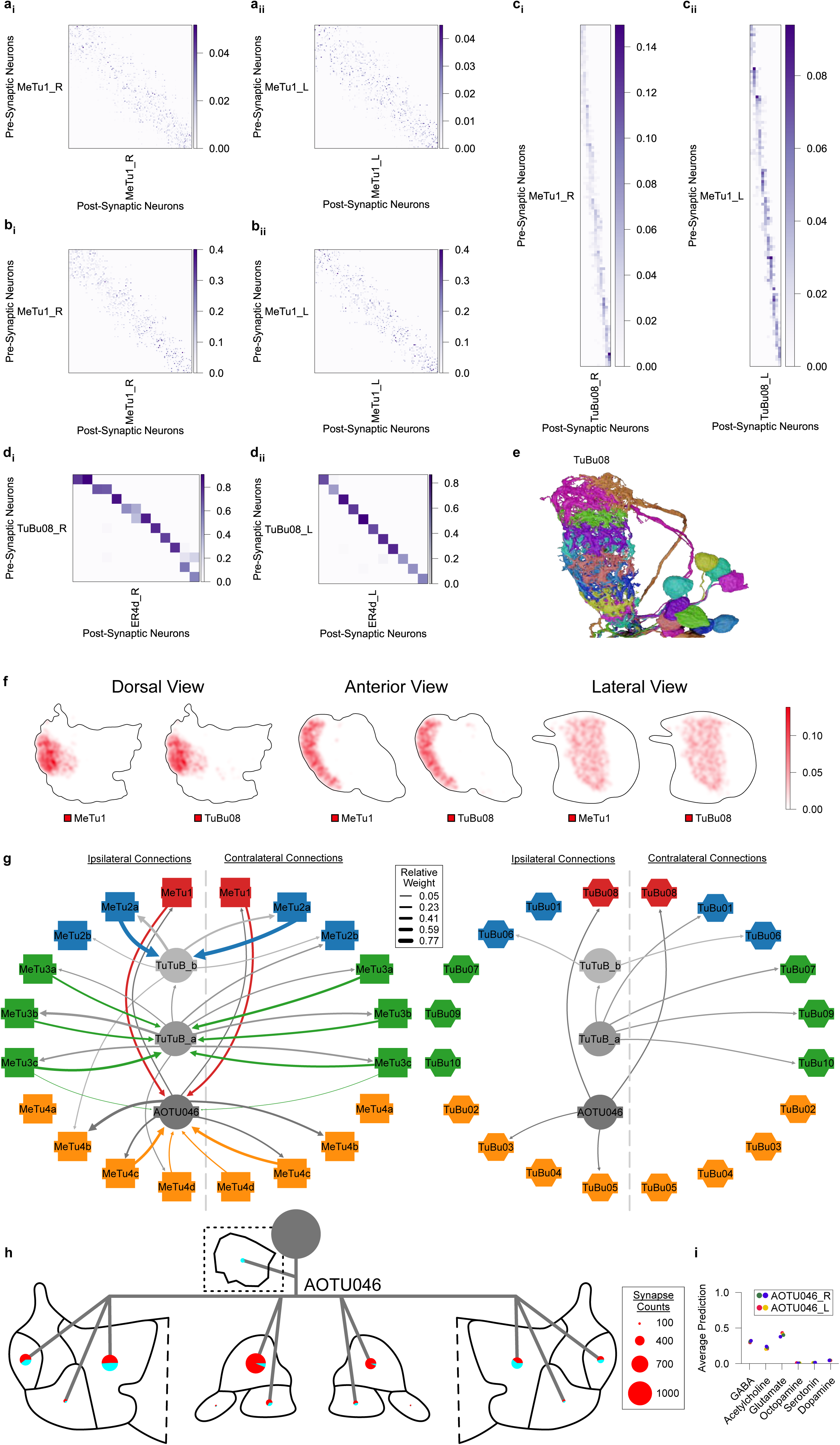
MeTu1 Pathway Connectivity. **a_i-ii_**, Synaptic weight matrices of MeTu1 interconnectivity within the medulla in the right (**a_i_**) and left (**a_ii_**) hemisphere. **b_i-ii_**, Synaptic weight matrices of MeTu1 interconnectivity within the AOTU in the right (**b_i_**) and left (**b_ii_**) hemisphere. **c_i-ii_**, Synaptic weight matrices of MeTu1 to TuBu08 in the AOTU in the right (**c_i_**) and left (**c_ii_**) hemisphere. **d_i-ii_**, Synaptic weight matrices of TuBu08 to ER4d in the Bulb in the right (**d_i_**) and left (**d_ii_**) hemisphere. **e**, All TuBu08 neurons of AOTUsu in the right hemisphere from the lateral perspective. Viewed from the anterior side, AOTUsu_PL is curved like a bow. Perpendicular cross sections through this domain appear elliptical, tapering along the ventral direction. **f**, Synapse density maps of connections between MeTu1 and TuBu08 neurons in the AOTU_SU from the dorsal, anterior, and lateral perspectives. All perspectives have been rotated 30° with respect to the anterior-posterior axis, and synapse densities were blurred with a Gaussian filter with a sigma value of 10. **g**, Left: Synaptic weight between bihemispheric neuron types and MeTu subtypes on the ipsilateral (left) and contralateral (right) sides. Right: Similar to the left, Synaptic weight between bihemispheric neuron types and TuBu types. MeTu2a receives strong synaptic inputs from TuTuB_b on both sides, but none from TuTuB_a. It also reciprocally provides strong inputs to TuTuB_b on both sides, but only very weak input to TuTuB_a. **h**, Diagram of an AOTU046 neuron. AOTU046 neurons innervates AOTUsu_M, where they send sparse axons along the anterior/posterior-lateral face and extend boutons toward the posterior-medial triangle vertex at only a single latitude halfway down the dorsal-ventral axis. Pie charts are the ratio of presynaptic (red) to postsynaptic (cyan) connections to AVP neurons in the AOTU and Bulb regions, and connections to all neurons in the SPS. The SPS depicted in a cutout. Pie chart sizes are based on the relative number of connections (legend on the right). **i**, Average neurotransmitter prediction score over all synapses in each AOTU046 neuron for each type of neurotransmitter.

**Fig. S3.**
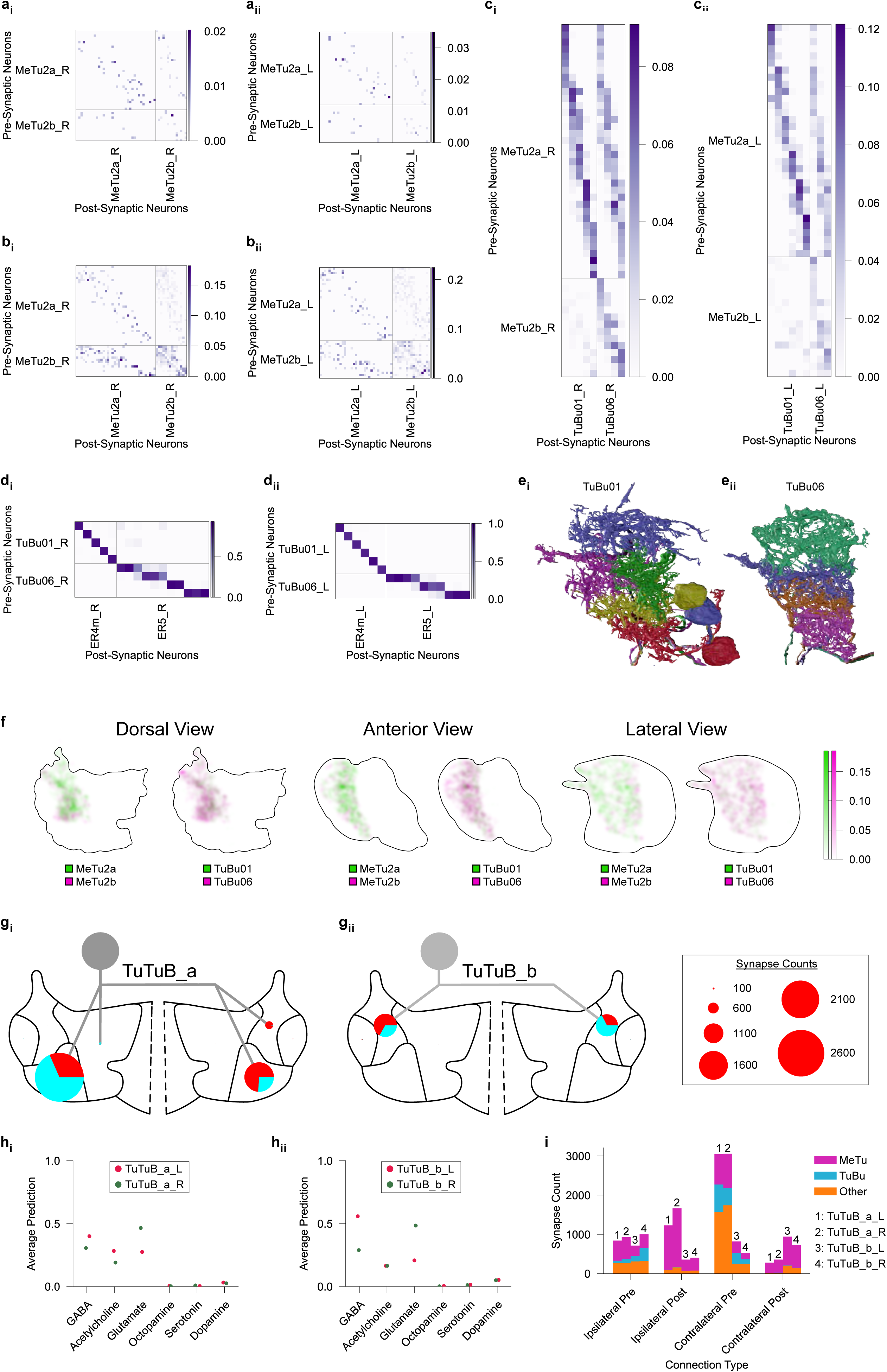
MeTu2 Pathway Connectivity. **a_i-ii_**, Synaptic weight matrices of MeTu2 interconnectivity within the medulla in the right (**a_i_**) and left (**a_ii_**) hemisphere. **b_i-ii_**, Synaptic weight matrices of MeTu2 interconnectivity within the AOTU of the right (**b_i_**) and left (**b_ii_**) hemisphere. **c_i-ii_**, Synaptic weight matrices of MeTu2 to TuBu1 and TuBu06 in the AOTU in the right (**c_i_**) and left (**c_ii_**) hemisphere. **d_i-ii_**, Synaptic weight matrices of TuBu01 and TuBu06 to relevant Ring neurons in the bulb in the right (**d_i_**) and left (**d_ii_**) hemisphere. **e_i-ii_**, All TuBu01 (**e_i_**) and TuBu06 (**e_i-ii_**) neurons from the lateral perspective in the AOTUsu. The cross-section of AOTUsu_PC is curved and slightly wraps around the AOTUsu_PL. Like the AOTUsu_PL, the area becomes thinner towards the ventral side. **f**, Synapse density maps of connections between MeTu2 and TuBu1 and TuBu06 neurons in the AOTU_SU from the dorsal, anterior, and lateral perspectives. All perspectives have been rotated 30° with respect to the anterior-posterior axis, and synapse densities were blurred with a Gaussian filter with a sigma value of 10. The distribution of presynaptic boutons of each MeTu2 neuron extends across the entire anterior-posterior axis of AOTUsu_PC, but the distribution is restricted along the dorsal-ventral axis. **g_i-ii_**, Diagrams of TuBuB_a (**g_i_**) and TuBuB_b (**g_ii_**) neurons. Boutons of TuTuB_a [28] sparsely protrude into the posterior central area only on the opposite side of the soma, while both axons and dendrites of TuTuB_b neurons innervate the entire AOTUsu_PC. Pie charts are the ratio of presynaptic (red) to postsynaptic (cyan) connections to AVP neurons in the AOTU and Bulb. Pie chart sizes are based on the relative amount of connections (legend on the right). **h_i-ii_**, The average neurotransmitter prediction score over all synapses in each TuTuB_a (**h_i_**) or TuTuB_b (**h_ii_**) neuron for each type of neurotransmitter. **i**, Number of synapses of each TuTuB neuron between the ipsilateral and contralateral hemisphere, based on the type of neuron it is connected to.

**Fig. S4.**
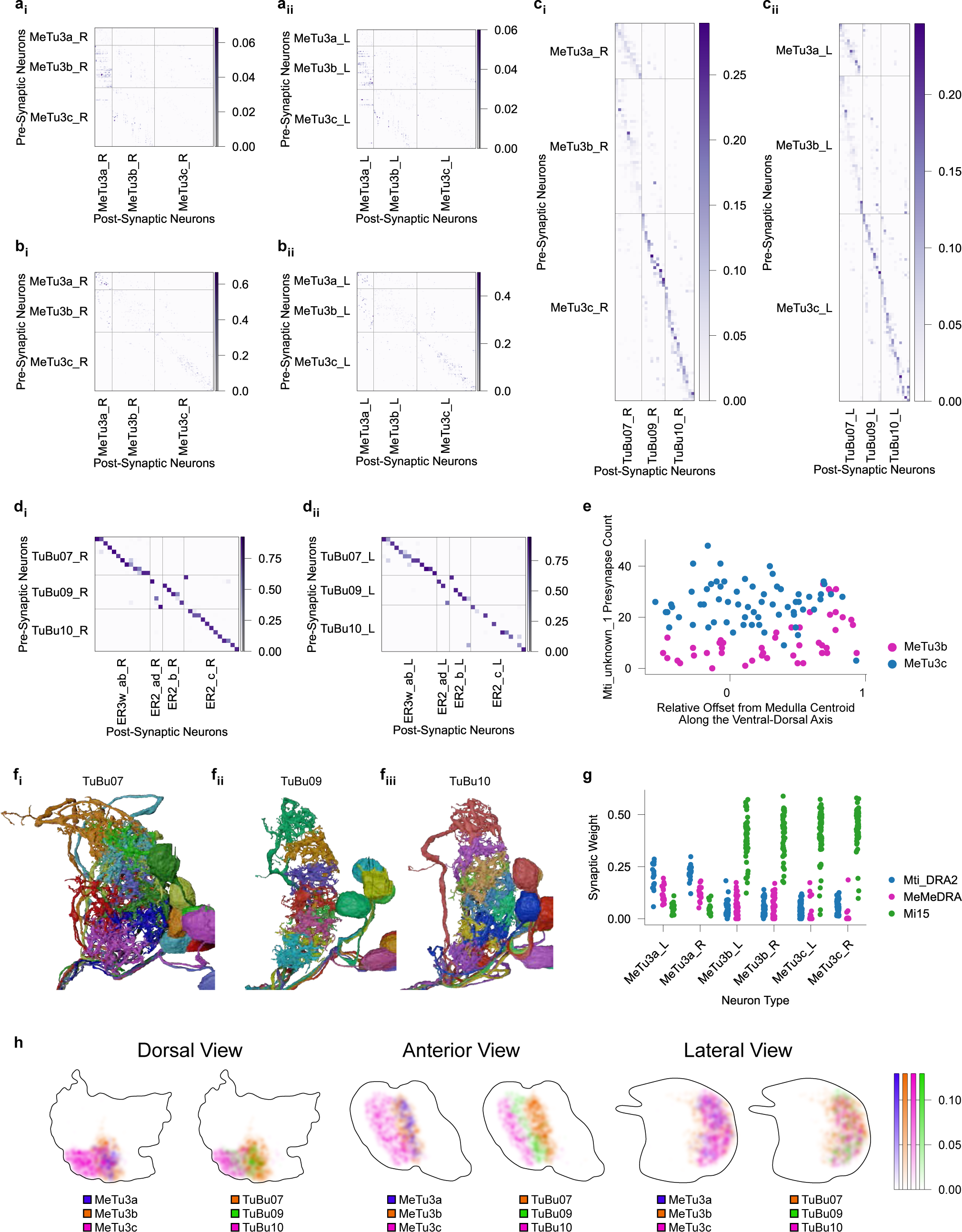
MeTu3 Pathway Connectivity. **a_i-ii_**, Synaptic weight matrices of MeTu3 interconnectivity within the medulla in the right (**a_i_**) and left (**a_ii_**) hemisphere. **b_i-ii_**, Synaptic weight matrices of MeTu3 interconnectivity within the AOTU in the right (**b_i_**) and left (**b_ii_**) hemisphere. **c_i-ii_**, Synaptic weight matrices of MeTu3 to TuBu_A in the AOTU in the right (**c_i_**) and left (**c_ii_**) hemisphere. **d_i-ii_**, Synaptic weight matrices of TuBu_A to relevant Ring neurons in the bulb in the right (**d_i_**) and left (**d_ii_**) hemisphere. **e**, Number of synapses from Mti_unknown_1 onto MeTu3b (pink) and MeTu3c (blue) neurons in the right hemisphere as a function of their relative position along the dorsal-ventral axis. 0 on the x-axis refers to the center of the Medulla, while positive values are more dorsal and negative values are more ventral. **f_i-iii_**, All TuBu07 (**f_i_**), TuBu09 (**f_ii_**), and TuBu10 (**f_iii_**) neurons of AOTUsu in the right hemisphere from the lateral perspective. The volume of AOTUsu_A is flat on the posterior and the medial sides and curves in a quarter-ellipse from the lateral edge to the anterior edge. **g**, Synaptic weight of upstream neurons to all neurons of the different subtypes of MeTu3. **h**, Synapse density maps of connections between MeTu3 and TuBu_A neurons in the AOTU_SU from the dorsal, anterior, and lateral perspectives. All perspectives have been rotated 30° with respect to the anterior-posterior axis, and synapse densities were blurred with a Gaussian filter with a sigma value of 10.

**Fig. S5.**
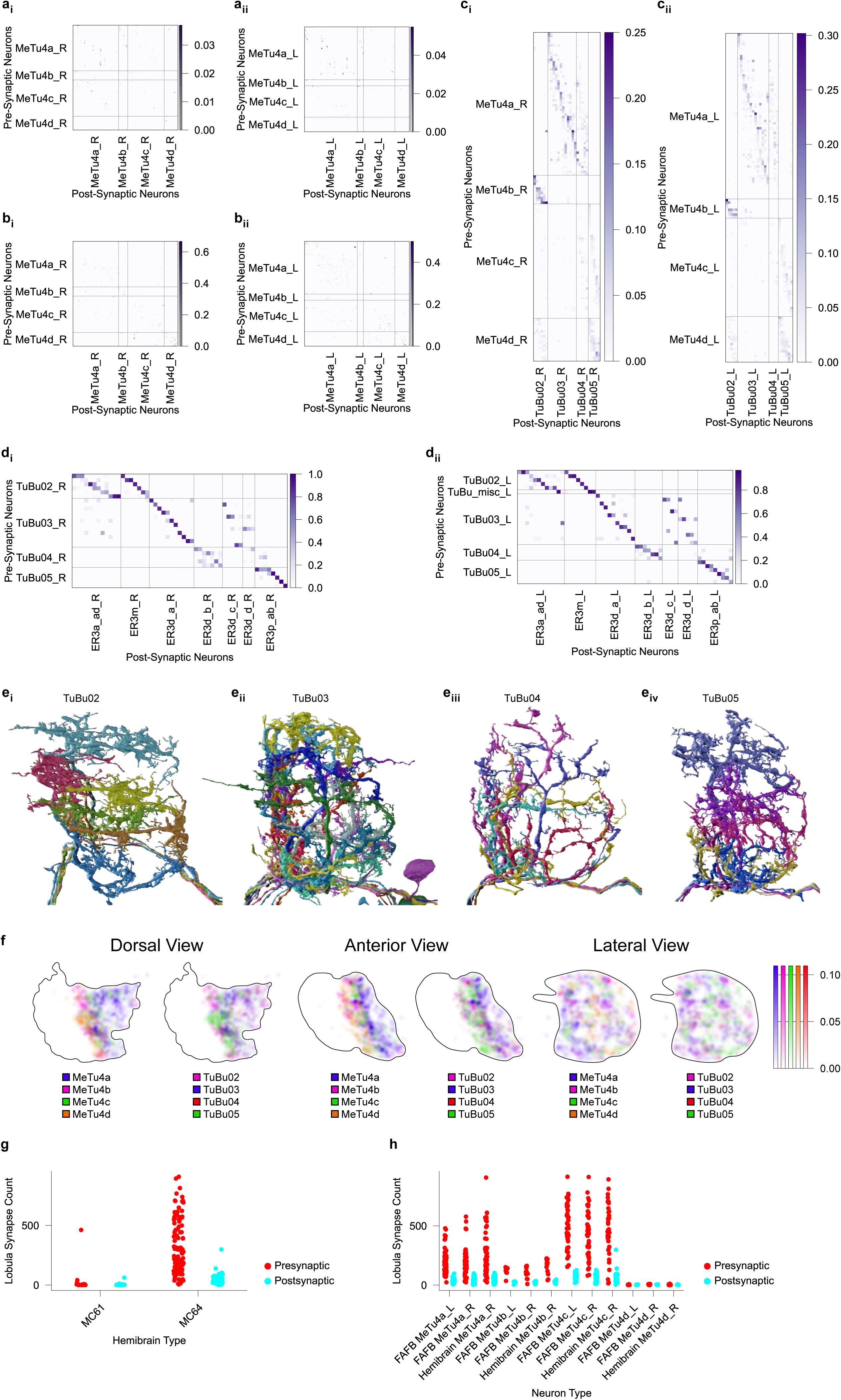
MeTu4 Pathway Connectivity. **a_i-ii_**, Synaptic weight matrices of MeTu4 interconnectivity within the medulla in the right (**a_i_**) and left (**a_ii_**) hemisphere. **b_i-ii_**, Synaptic weight matrices of MeTu4 interconnectivity within the AOTU in the right (**b_i_**) and left (**b_ii_**) hemisphere. **c_i-ii_**, Synaptic weight matrices of MeTu4 to TuBu02/03/04/05in the AOTU in the right (**c_i_**) and left (**c_ii_**) hemisphere. **d_i-ii_**, Synaptic weight matrices of TuBu TuBu02/03/04/05 to relevant Ring neurons in the bulb in the right (**d_i_**) and left (**d_ii_**) hemisphere. **e_i-iv_**, All TuBu02 (**e_i_**), TuBu03 (**e_ii_**), TuBu04 (**e_iii_**), and TuBu05 (**e_iv_**) neurons of AOTUsu in the right hemisphere from the lateral perspective. The cross-section of AOTUsu_M appears right-triangle-shaped. One edge of the triangle is the boundary between AOTUsu_M and both AOTUsu_PC and AOTUsu_A, while the opposite vertex extends medially outward on the posterior side. Some MeTu and TuBu types cluster only along the region border, while others fill the entire triangle. **f**, Synapse density maps of connections between MeTu4 and TuBu02/03/04/05neurons in the AOTU_SU from the dorsal, anterior, and lateral perspectives. All perspectives have been rotated 30° with respect to the anterior-posterior axis, and synapse densities were blurred with a Gaussian filter with a sigma value of 10. **g**, Number of lobula presynapses (red) and postsynapses (cyan) of all MeTu4 neurons in the hemibrain dataset, sorted by whether they were previously classified as MC61 or MC64. MeTu4d neurons were generally classified as MC61, due to not having synapses in the lobula. **h**, Number of lobula presynapses (red) and postsynapses (cyan) of all MeTu4 neurons in both datasets after subclassification.

**Fig. S6.**
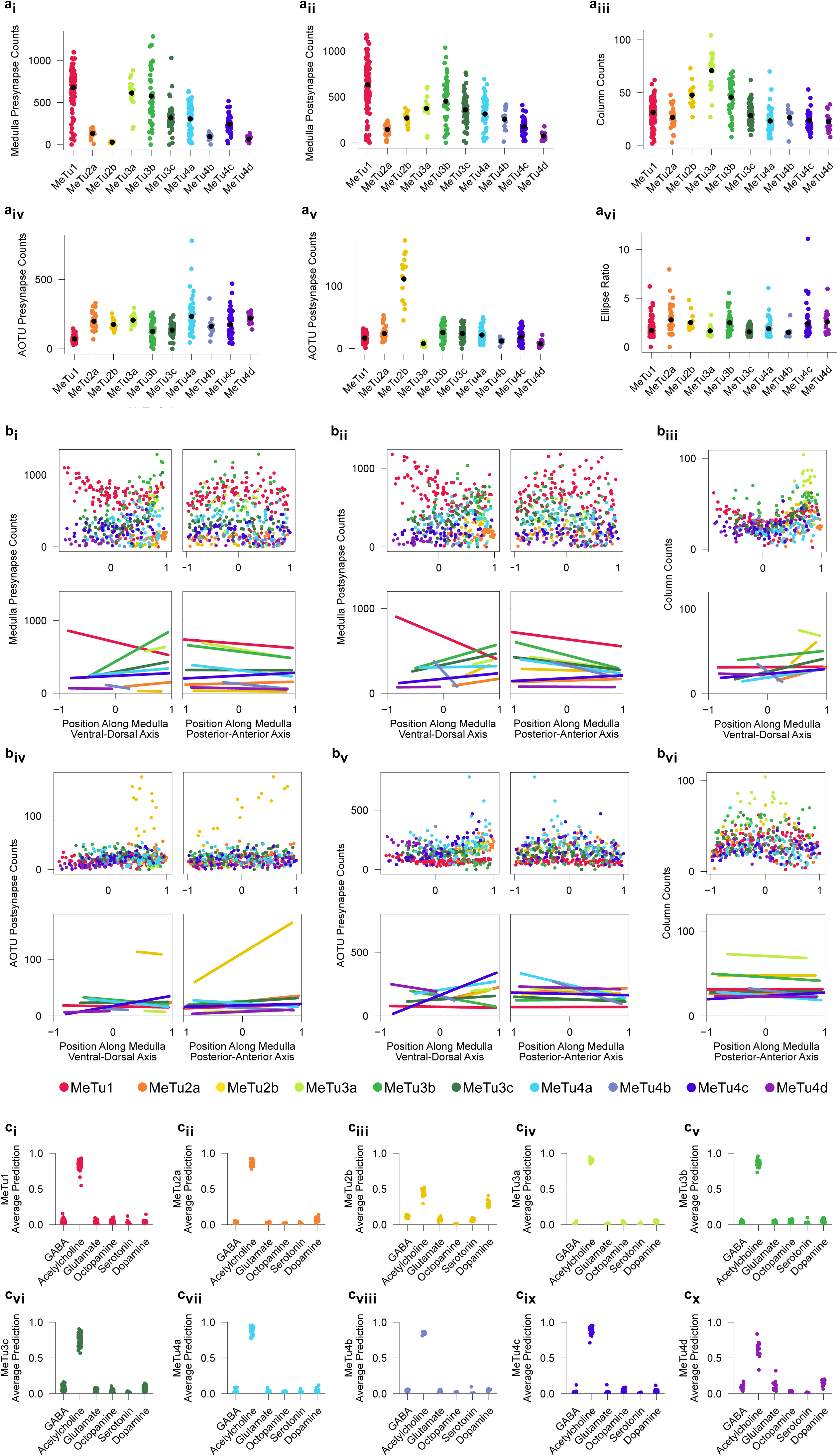
Comparisons Between MeTu Subtypes. **a**, Comparisons between all MeTu subtypes in the right hemisphere. Number of presynapses in the medulla (**a_i_**), number of postsynapses in the medulla (**a_ii_**), number of medulla columns that each neuron’s dendritic span occupies (**a_iii_**), number of presynapses in the AOTU (**a_iv_**), number of postsynapses in the AOTU (**a_v_**), and the ellipse ratio of the dendrites in the medulla (**a_vi_**). **b**, Comparisons among all MeTu subtypes in the right hemisphere with respect to the ventral-dorsal axis (negative values are ventral, positive values are dorsal) and the posterior-anterior axis (negative values are posterior, positive values are anterior) of the medulla. Scatter plot of all MeTu neurons (top) and the line of best fit (bottom). The color legend is at the bottom of (**b**). **b_i_**, Number of presynapses in the medulla as a function of the position along the ventral-dorsal axis (left) and posterior-anterior axis (right) of the medulla. **b_ii_**, Number of postsynapses in the medulla as a function of the position along the ventral-dorsal axis (left) and posterior-anterior axis (right) of the medulla. **b_iii_**, Number of medulla columns that each neuron’s dendritic span occupies as a function of the position along the ventral-dorsal axis of the medulla. **b_iv_**, Number of presynapses in the AOTU against the ventral-dorsal axis (left) and posterior-anterior axis (right) of the AOTU. **b_v_**, Number of postsynapses in the AOTU against the ventral-dorsal axis (left) and posterior-anterior axis (right) of the AOTU. **b_vi_**, Number of medulla columns that each neuron’s dendritic span occupies as a function of the position along the posterior-anterior axis of the medulla. **c_i-x_**, Average neurotransmitter prediction score over all synapses in each MeTu1/2a-b/3a-c/4a-d neuron respectively for each type of neurotransmitter. Note that the neurotransmitter of MeTu2b is not clearly predicted as cholinergic.

**Fig. S7.**
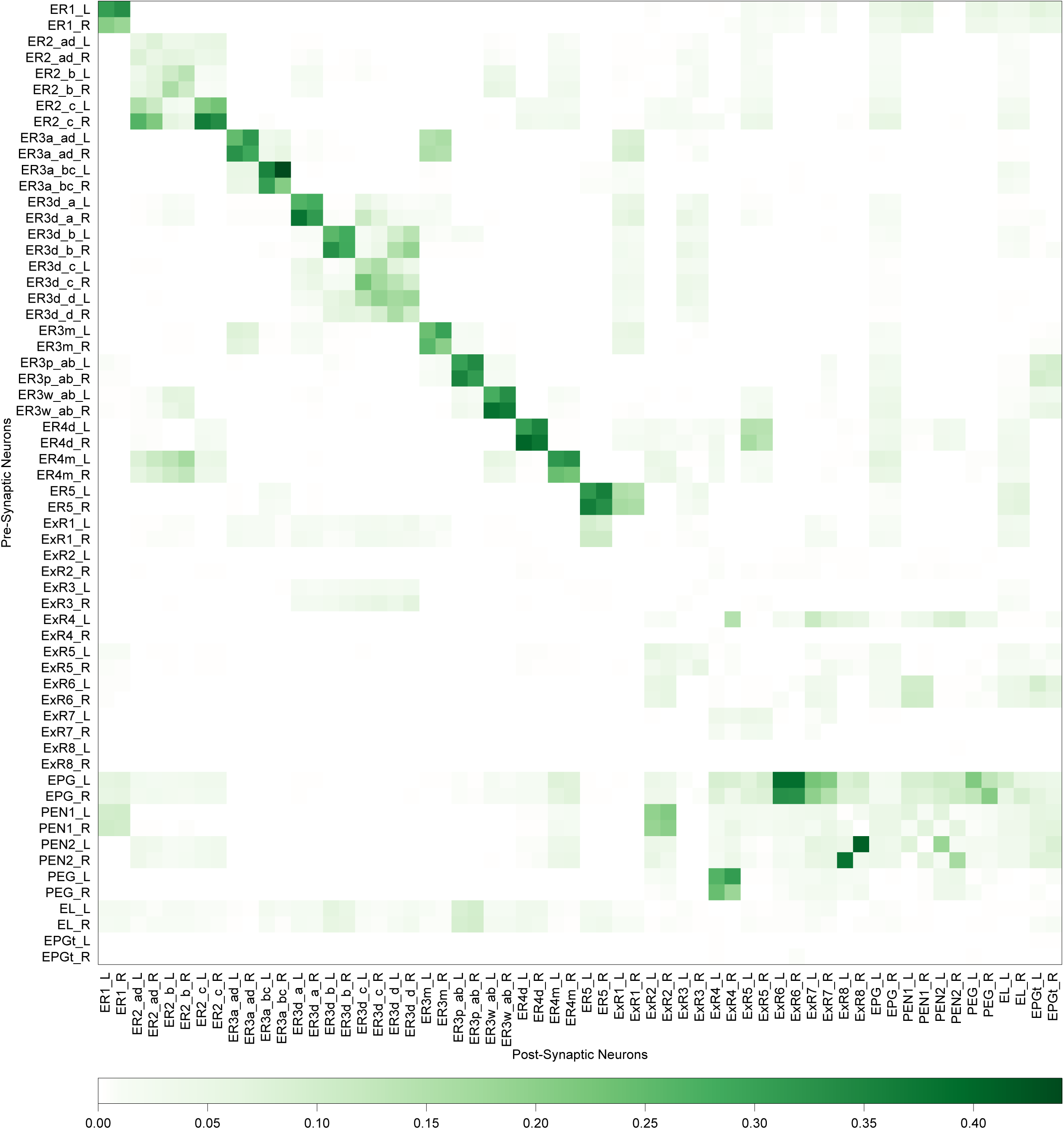
Synaptic Connections Between Ring and Columnar Neurons Within the Ellipsoid Body. Connection Matrix of various neuron types within the EB, including all visual ring neurons and lateral accessory lobe ring neurons [31, 61], extrinsic ring neurons [34], EPG neurons [1], PEN1 and PEN2 neurons [57, 101], and PEG, EL, and EPGt neurons [28, 29, 58, 102].

