## Supplementary material for "Connectomic reconstruction predicts the functional organization of visual inputs to the navigation center of the *Drosophila* brain": Data set 1

### MeTu1_720575940604885792

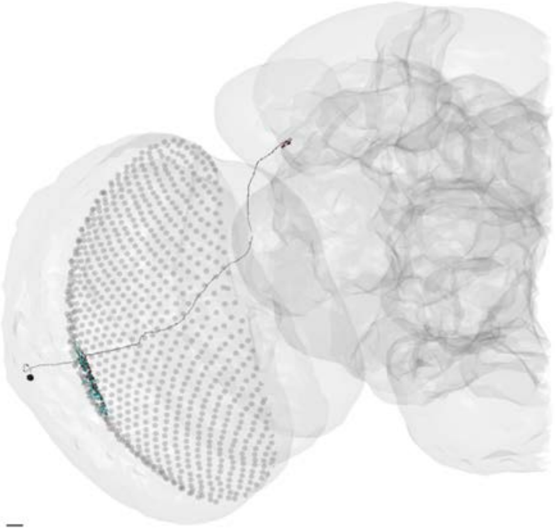

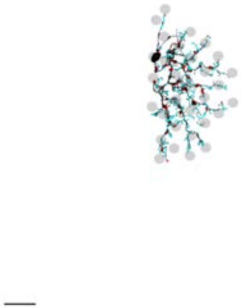

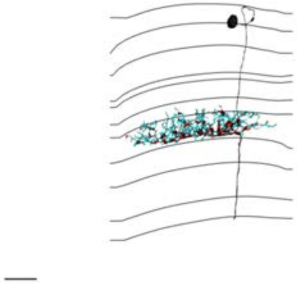

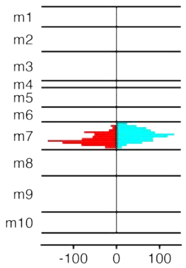


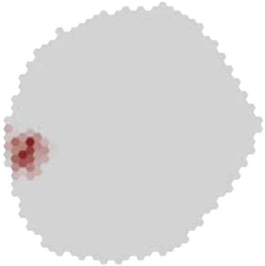

MeTu1_720575940607245186

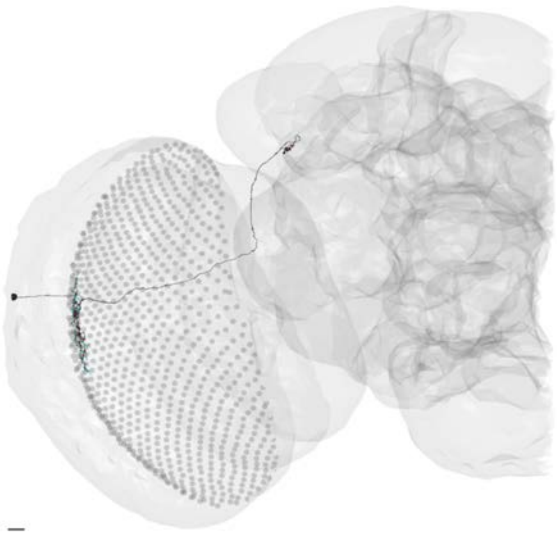

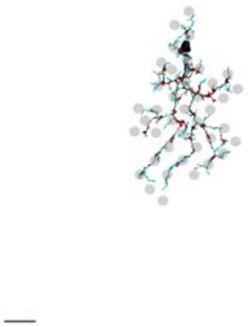

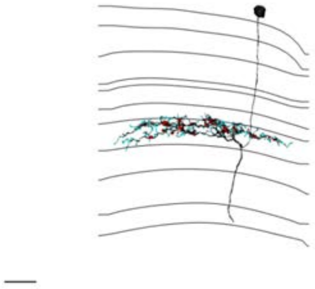

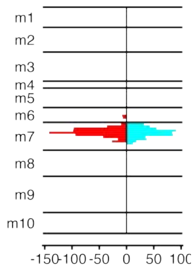


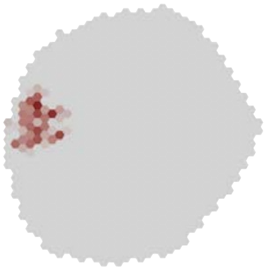

### MeTu1_720575940607509449

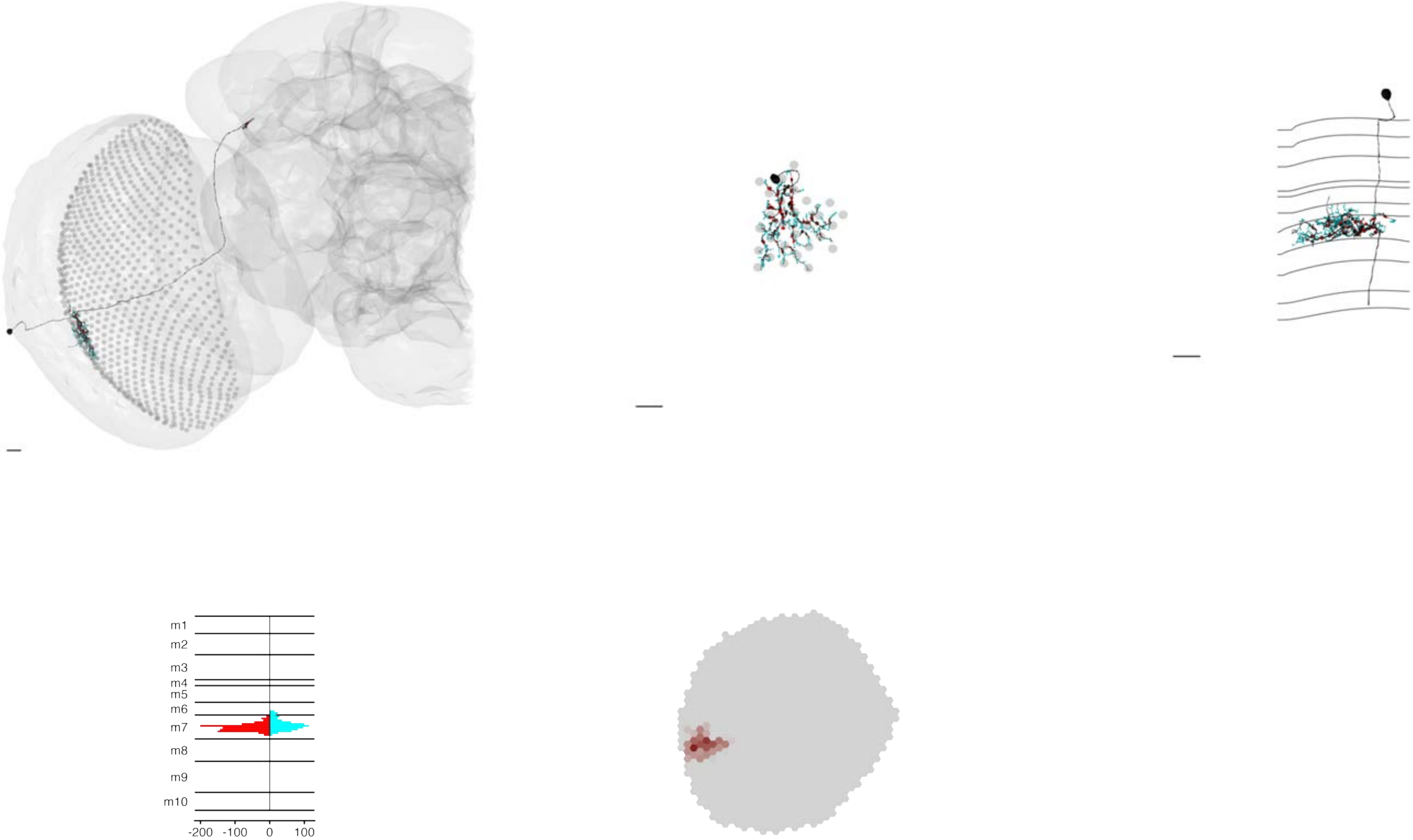

### MeTu1_720575940607582729

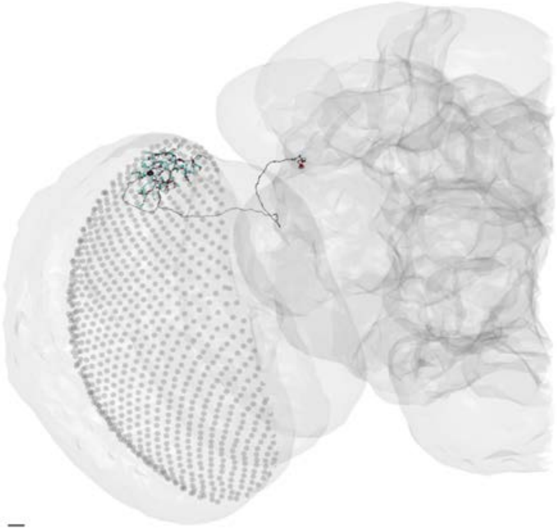

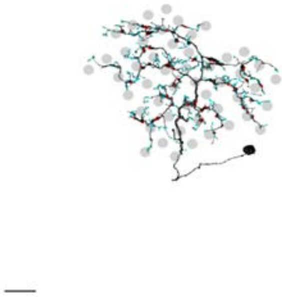

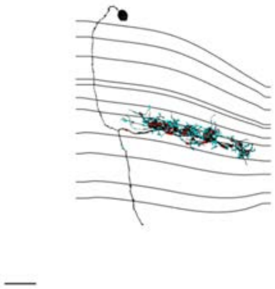

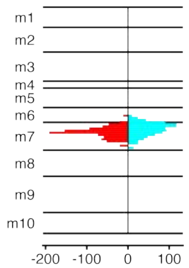


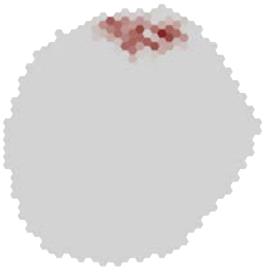

### MeTu1_720575940607798402

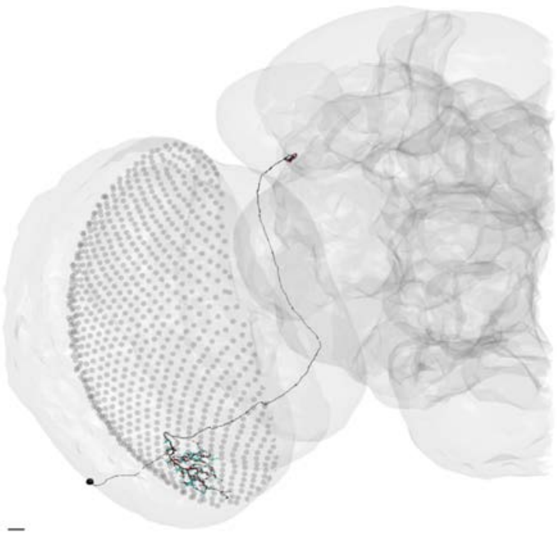

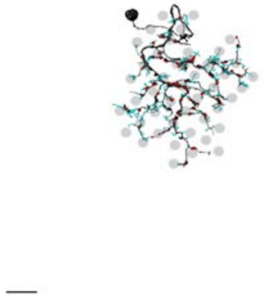

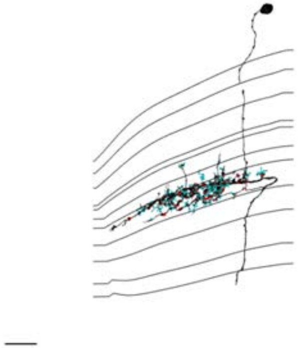

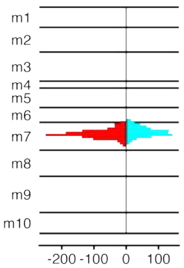


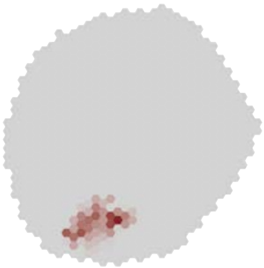

### MeTu1_720575940608907100

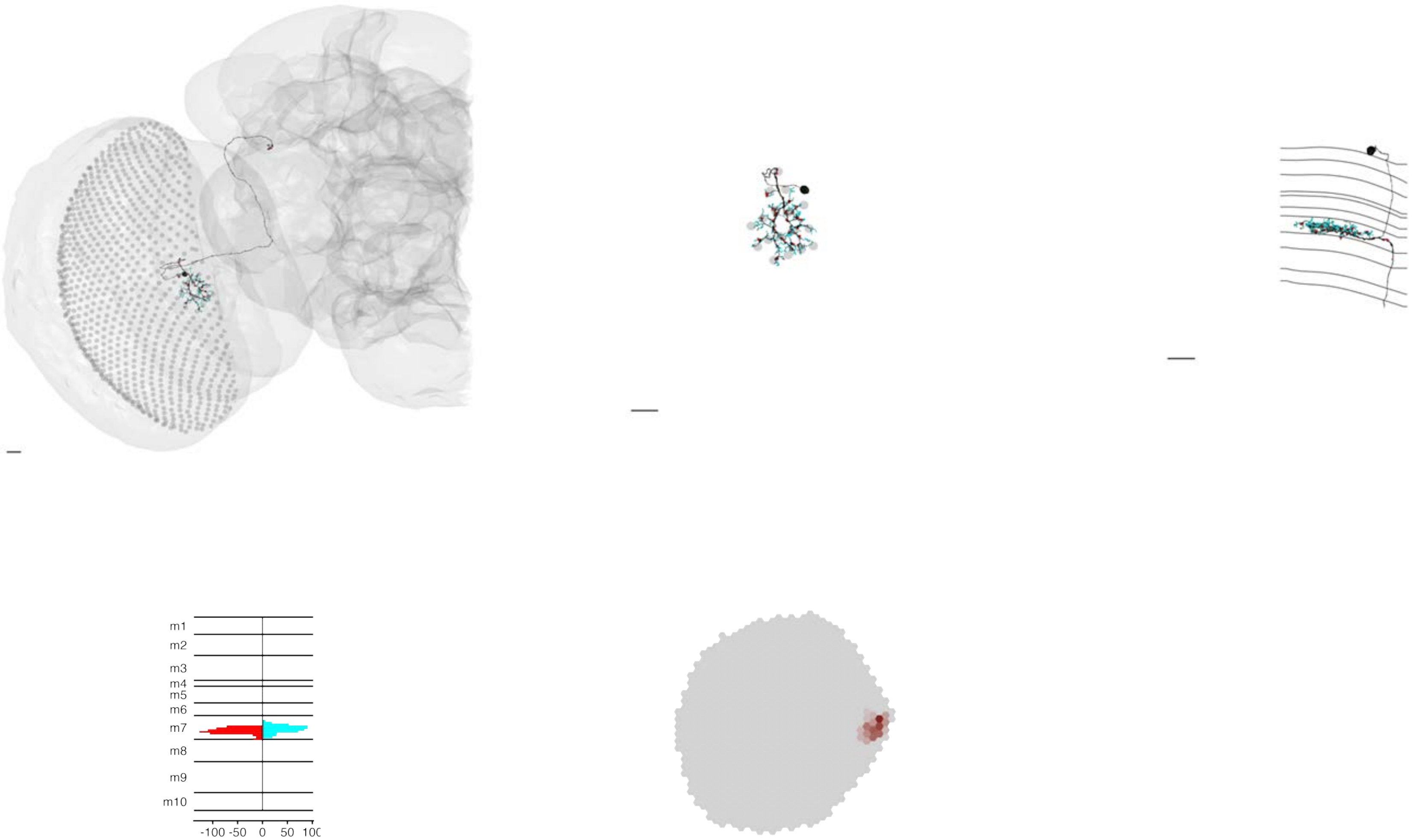

MeTu1_720575940609664963

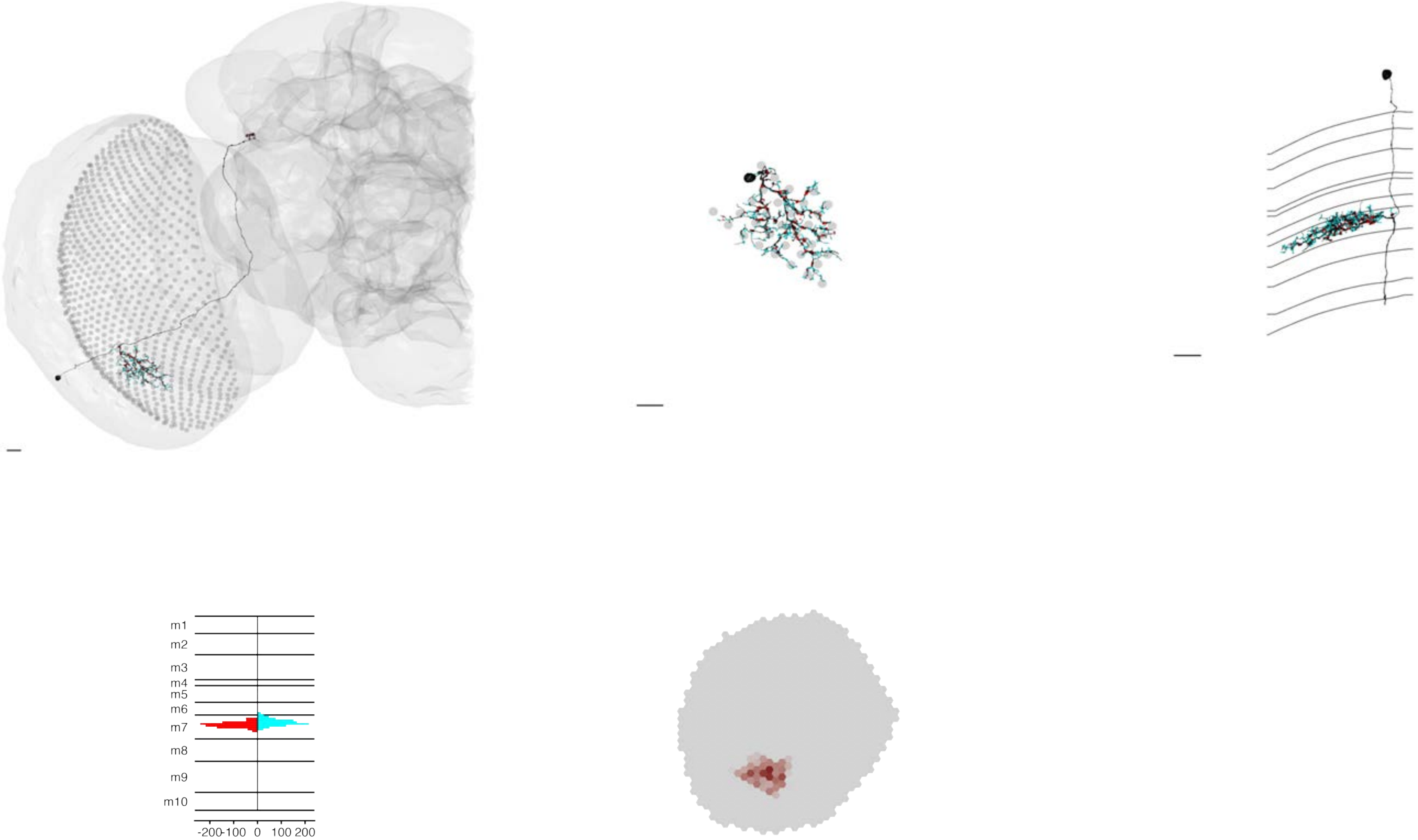

MeTu1_720575940609976533

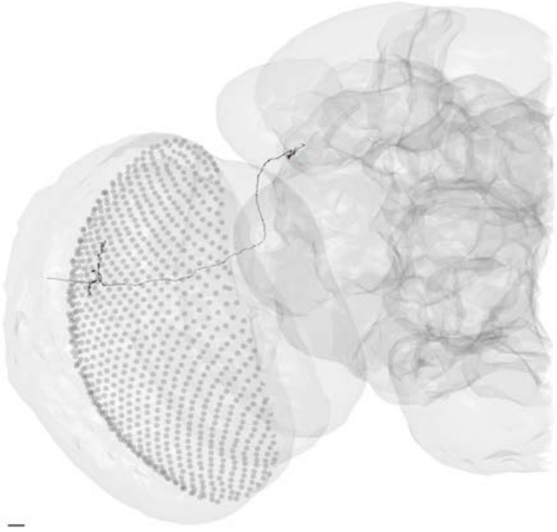

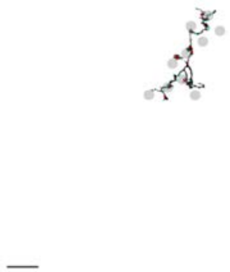

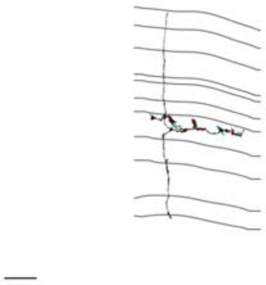

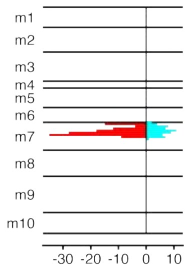


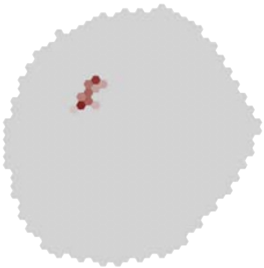

### MeTu1_720575940610257476

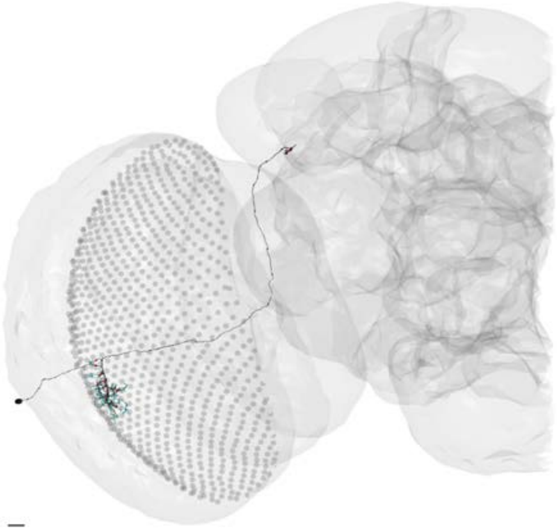

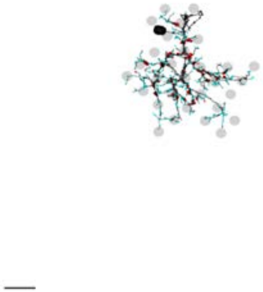

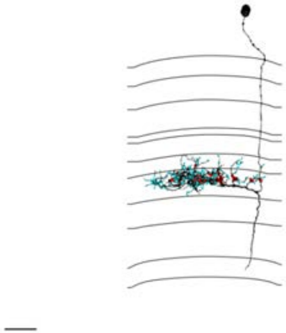

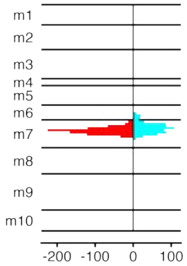


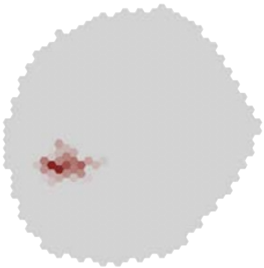

### MeTu1_720575940610266517

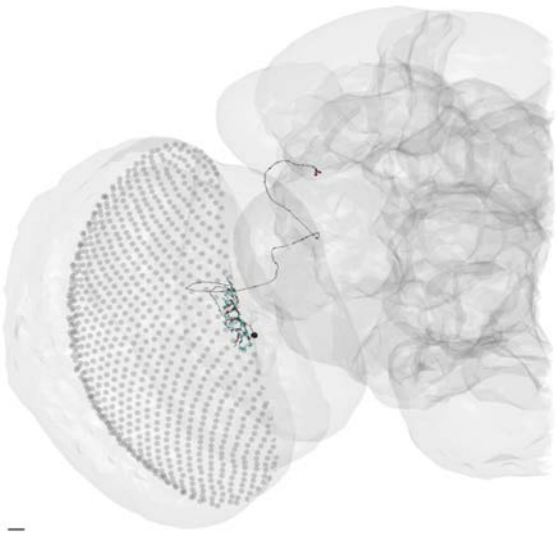

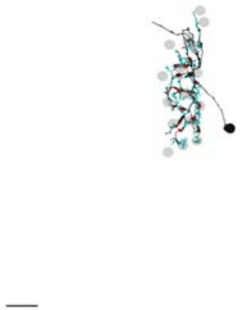

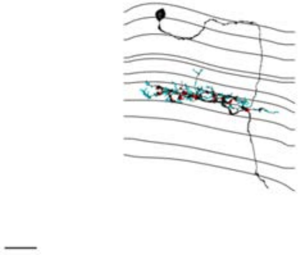

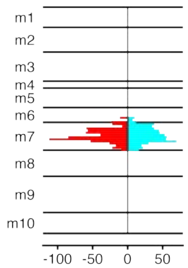


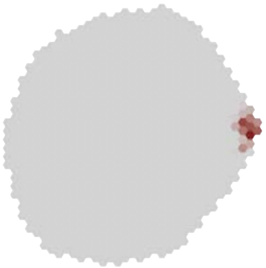

MeTu1_720575940611063921

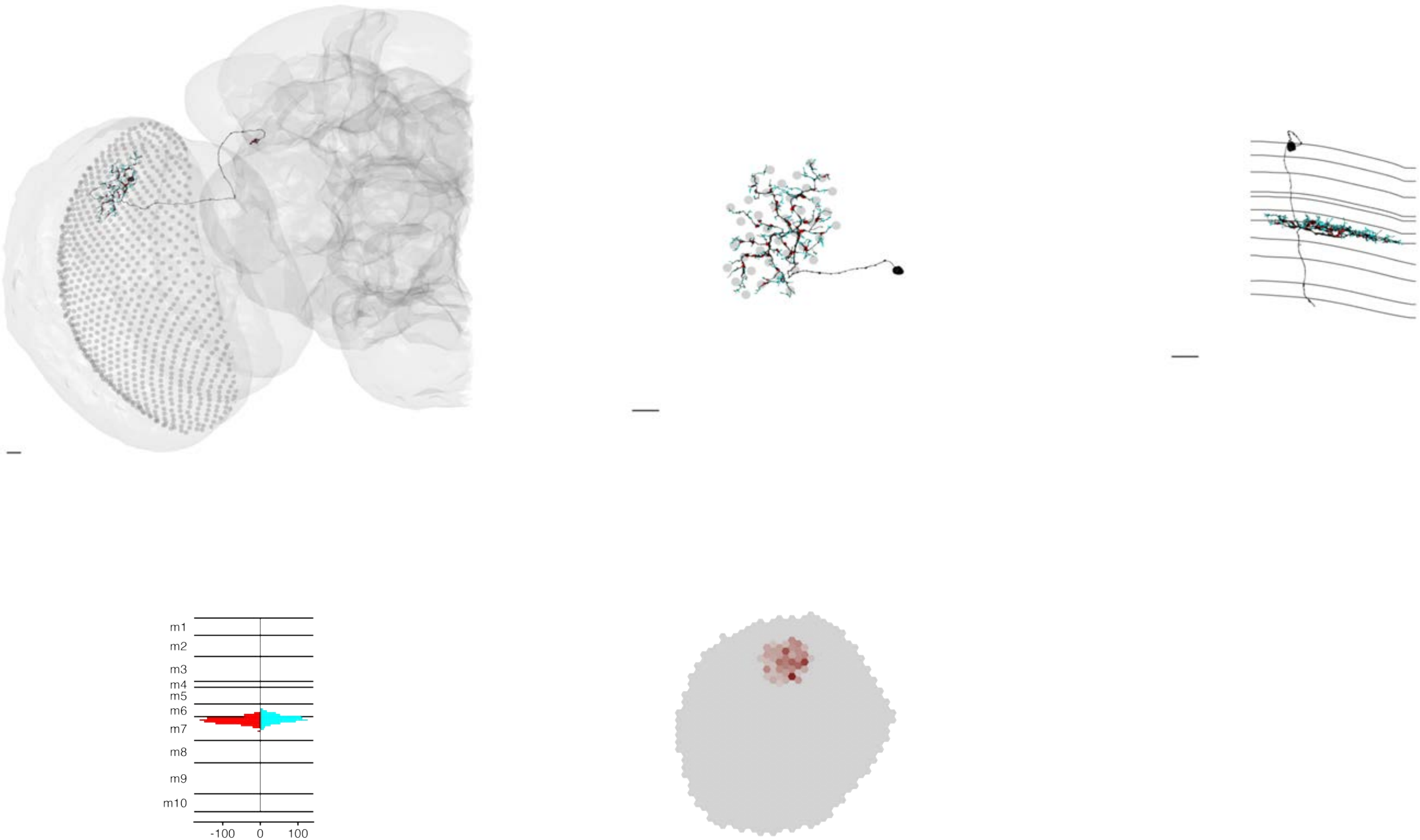

### MeTu1_720575940611335666

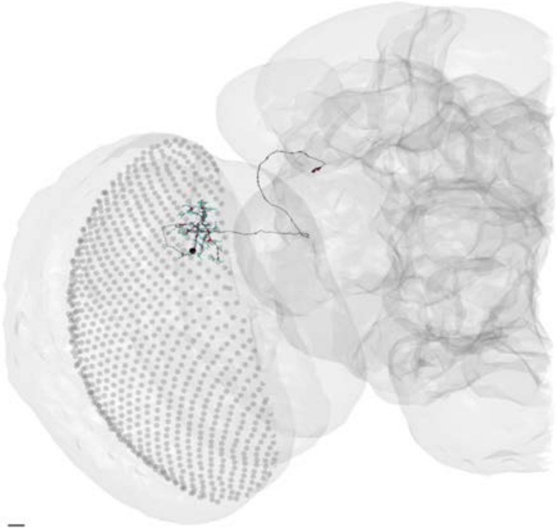

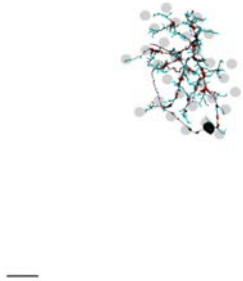

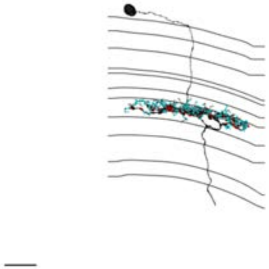

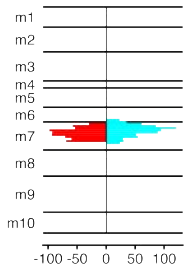


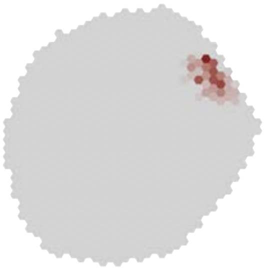

### MeTu1_720575940611700430

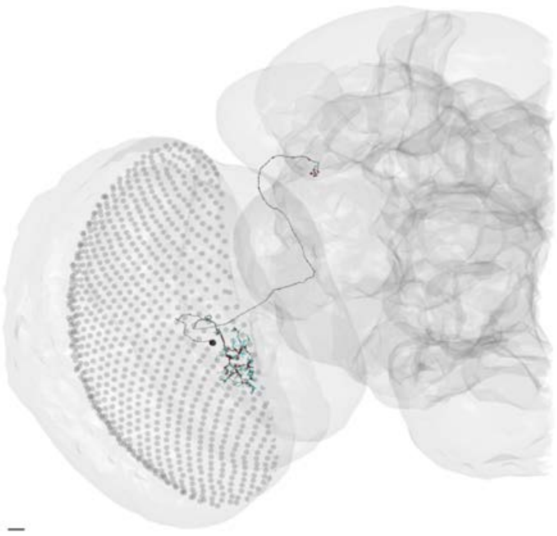

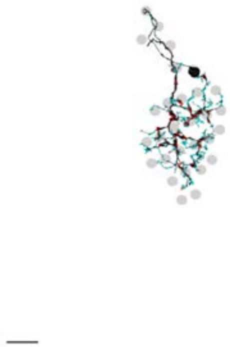

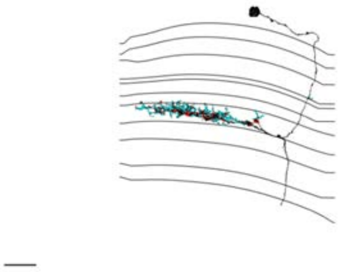

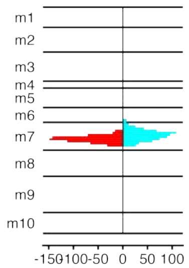


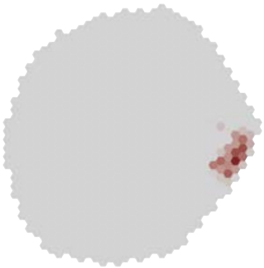

### MeTu1_720575940611780962

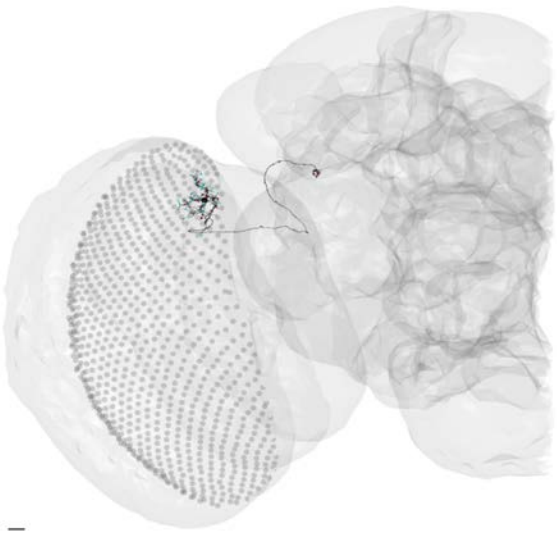

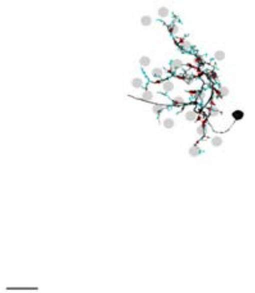

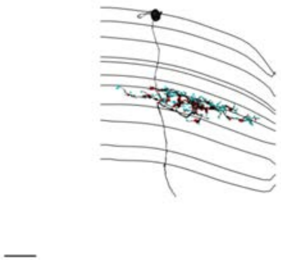

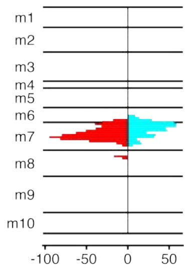


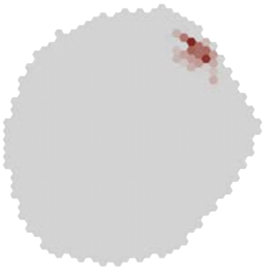

### MeTu1_720575940612340630

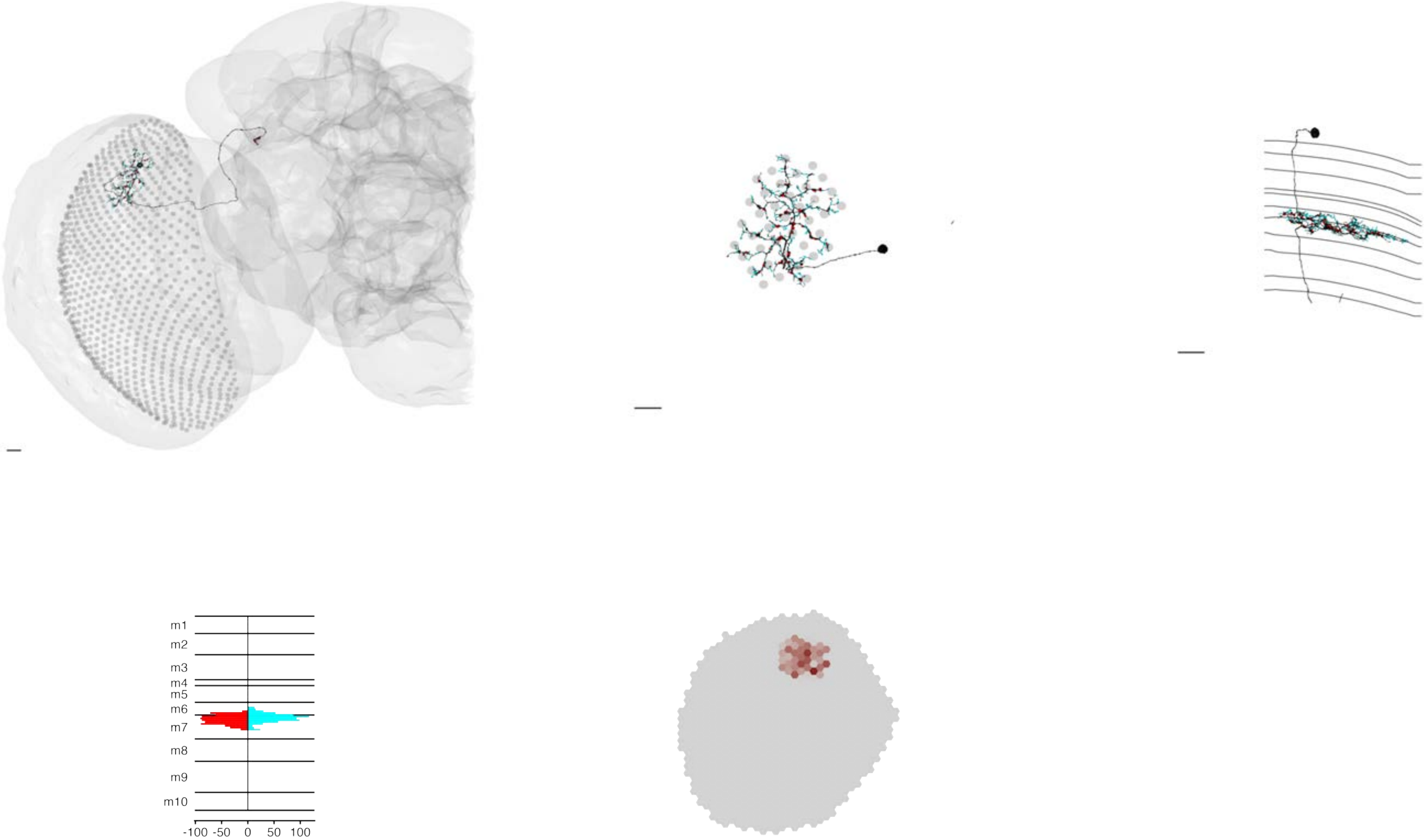

### MeTu1_720575940612572822

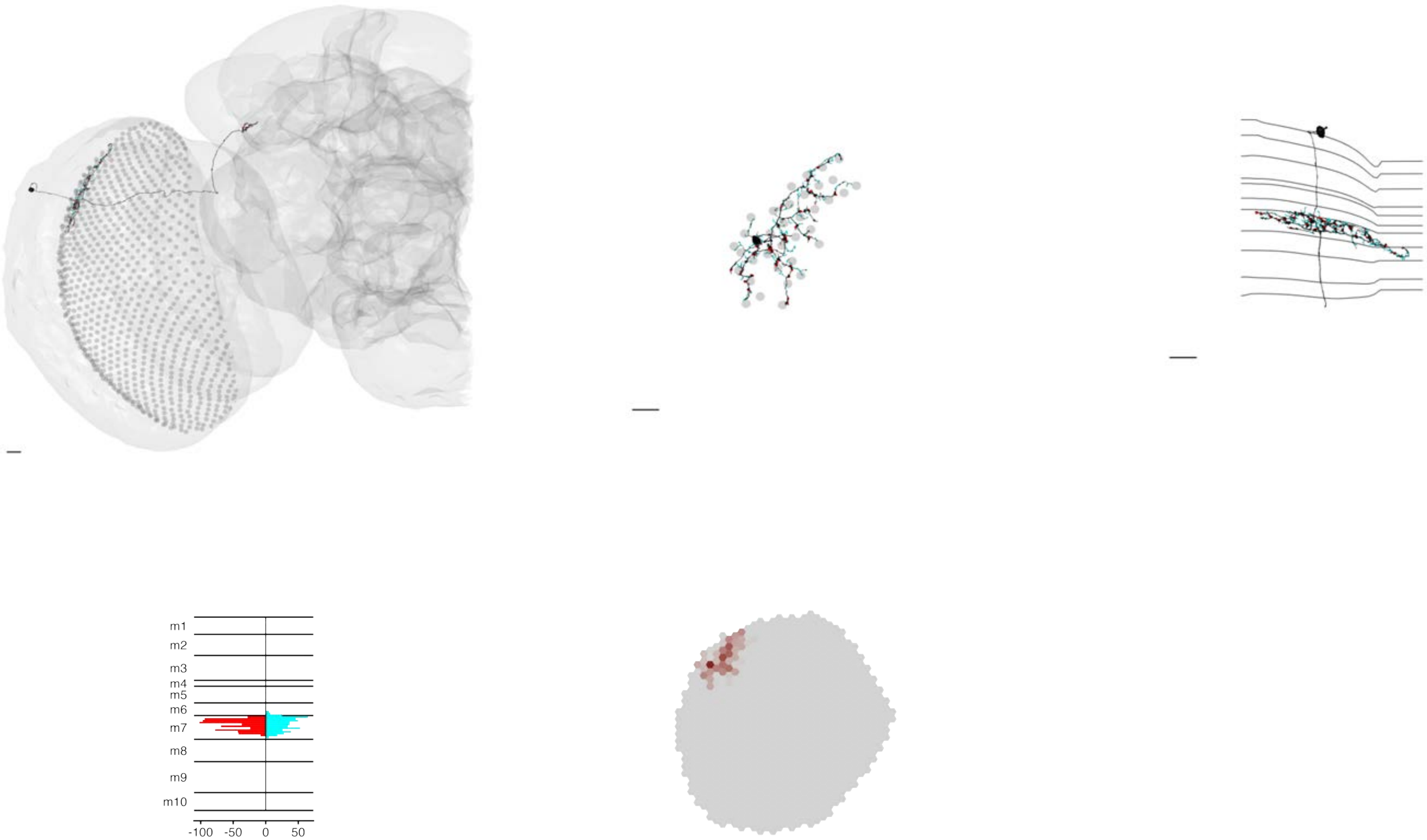

### MeTu1_720575940612637810

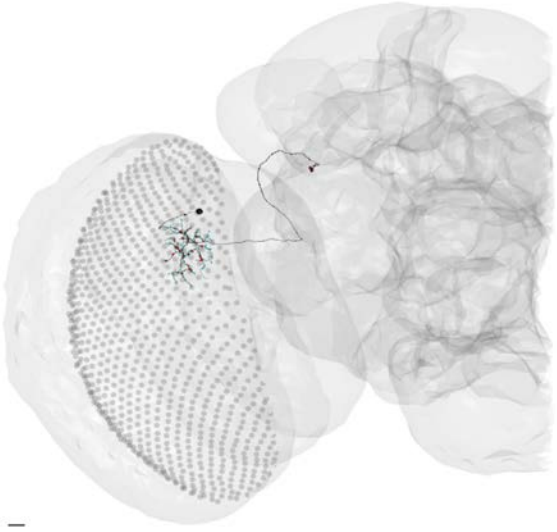

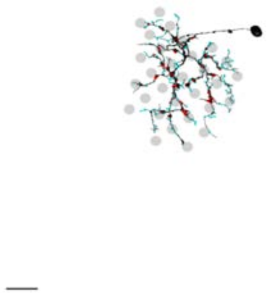

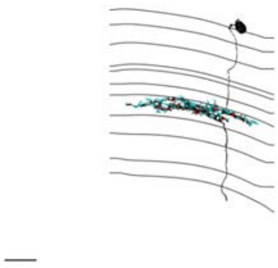

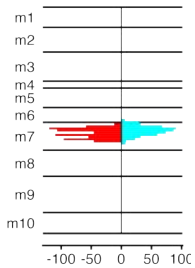


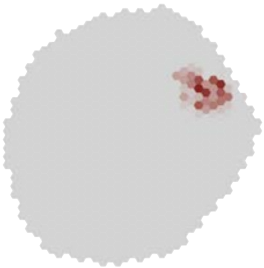

### MeTu1_720575940613739586

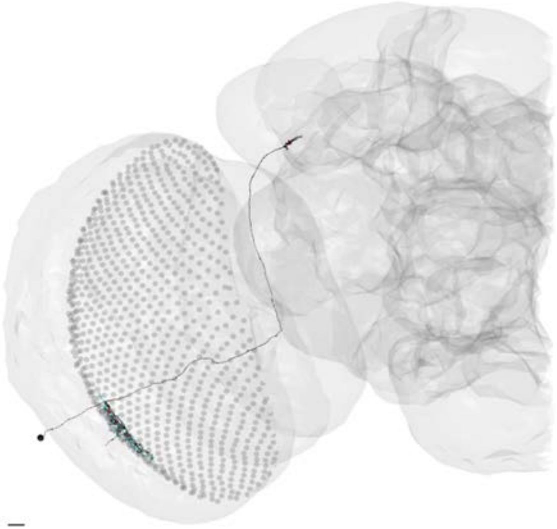

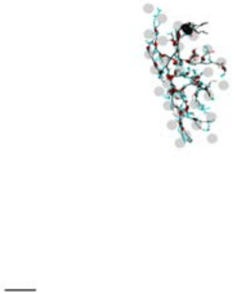

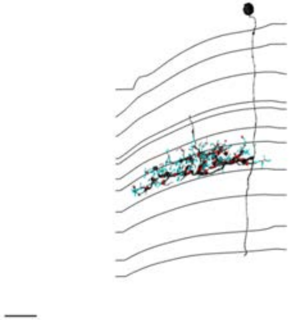

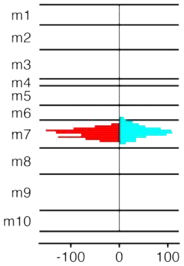


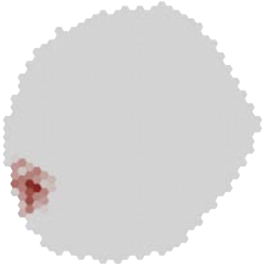

### MeTu1_720575940613834207

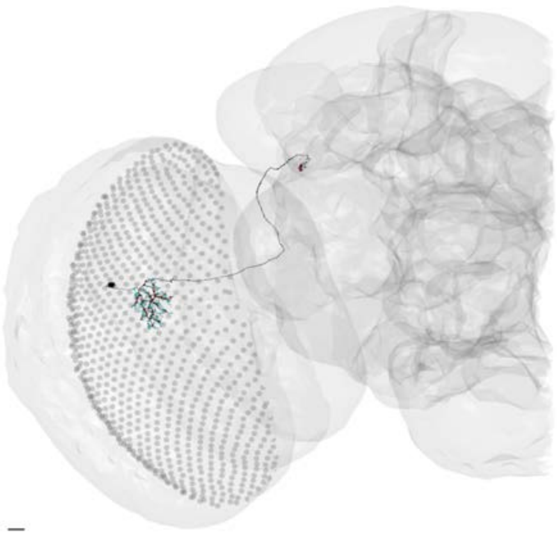

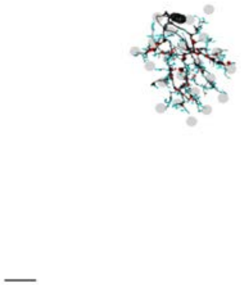

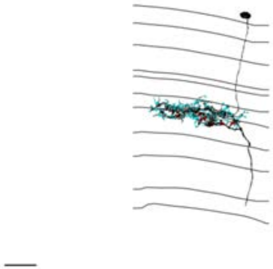

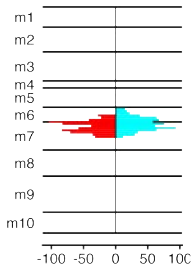


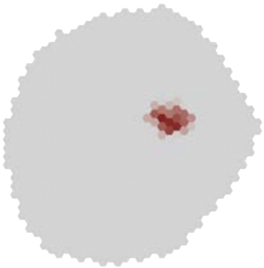

MeTu1_720575940614723189

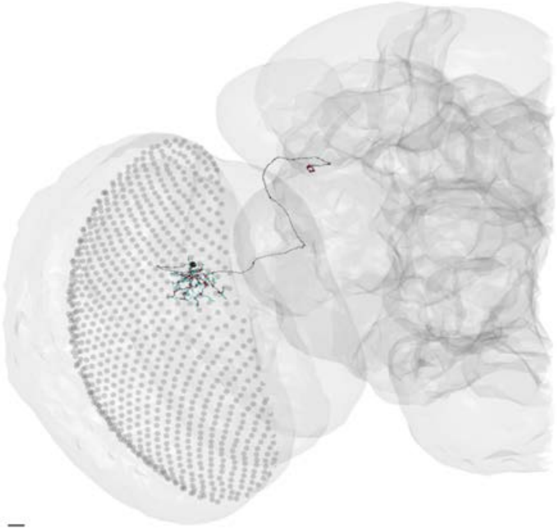

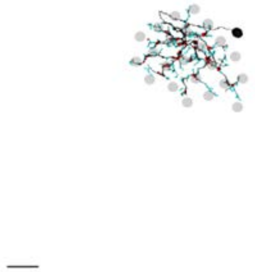

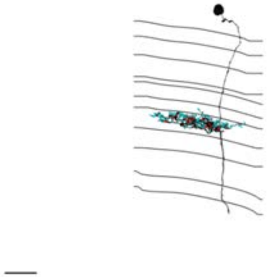

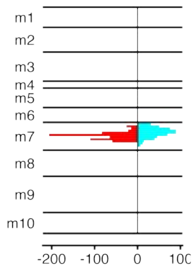


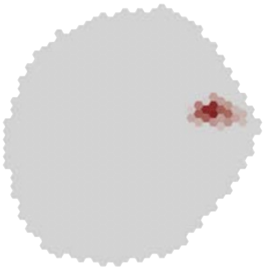

### MeTu1_720575940615291538

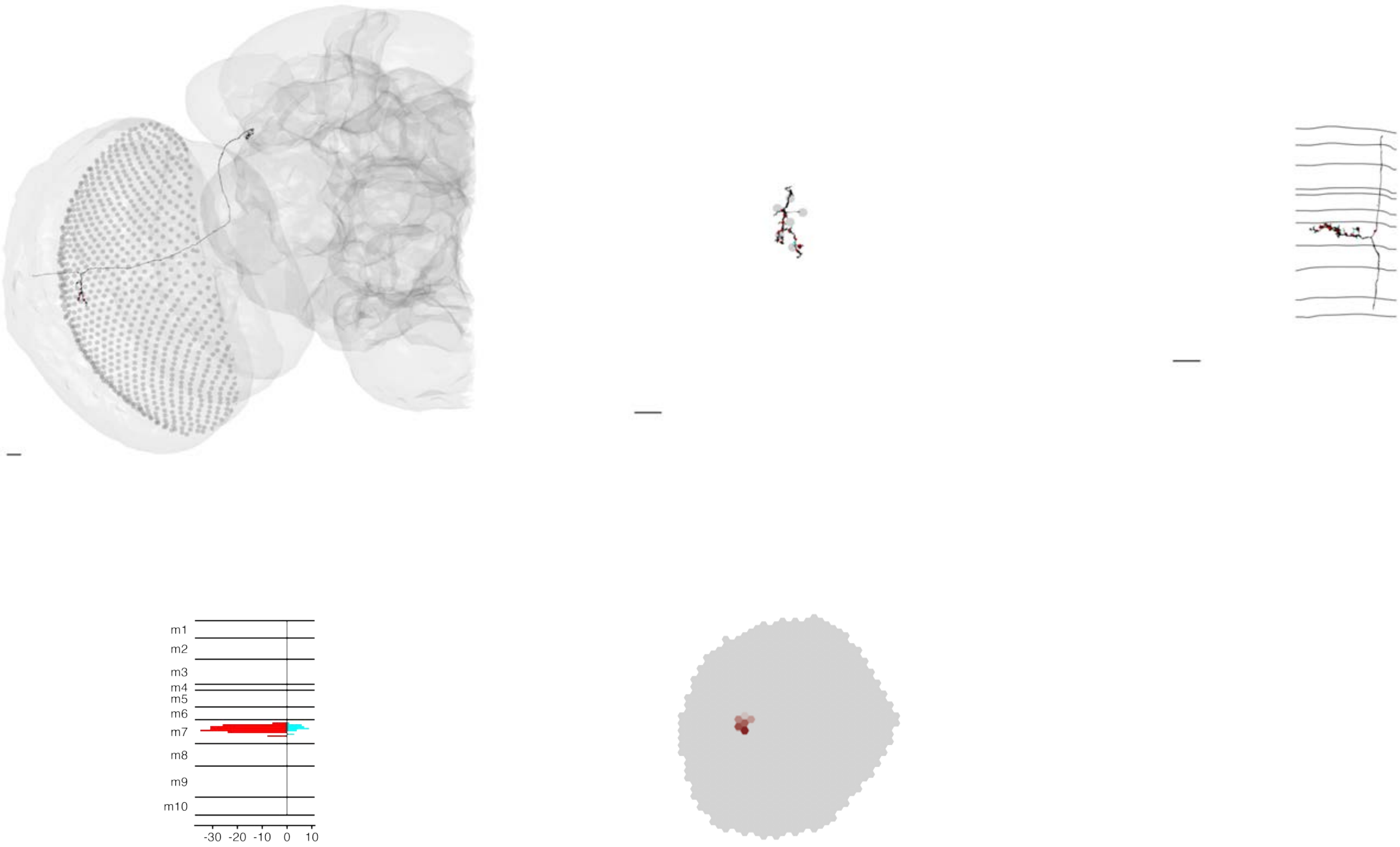

### MeTu1_720575940616017382

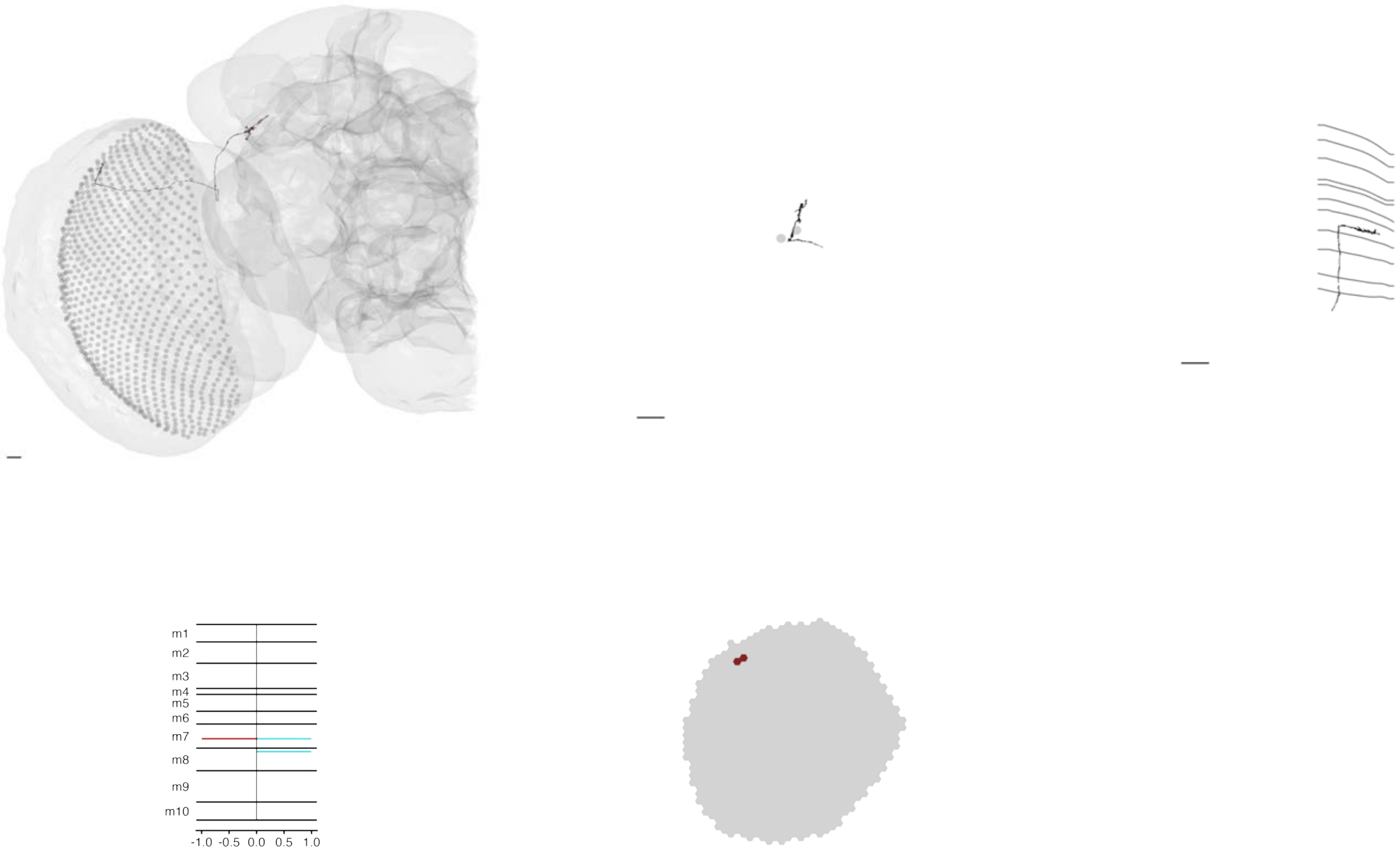

### MeTu1_720575940616078751

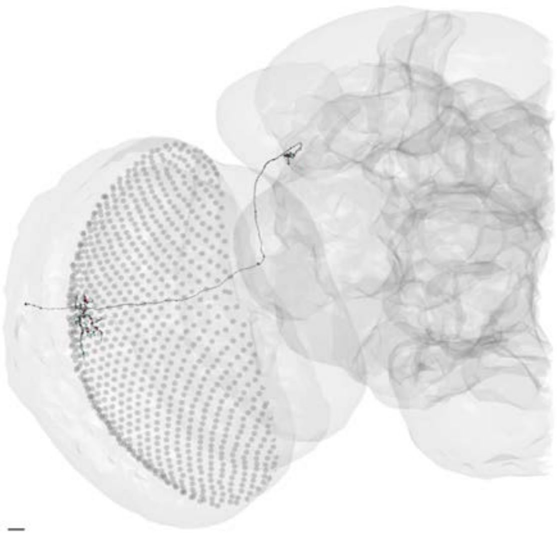

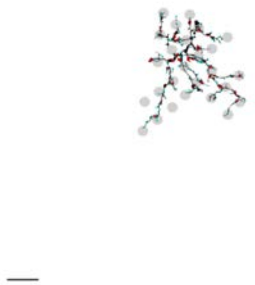

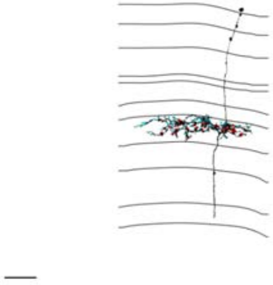

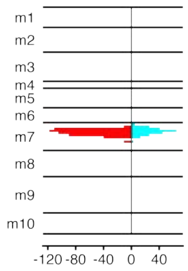


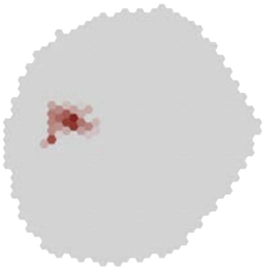

### MeTu1_720575940616190164

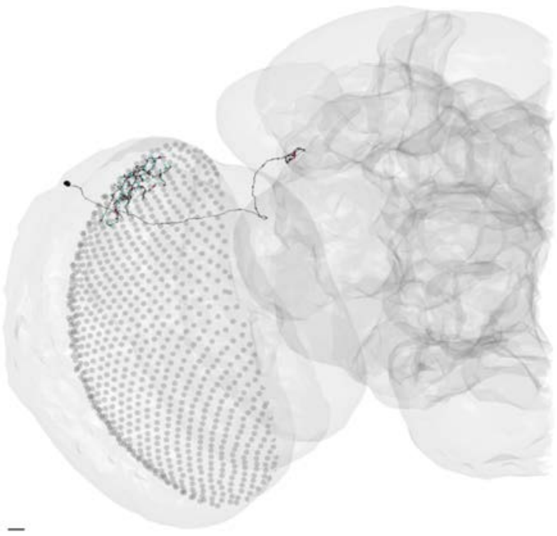

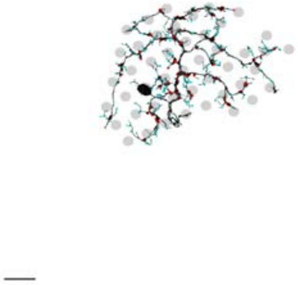

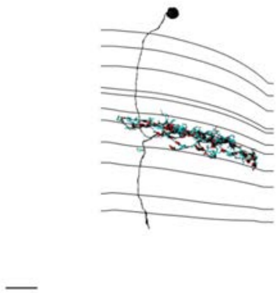

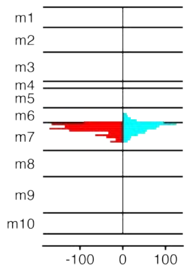


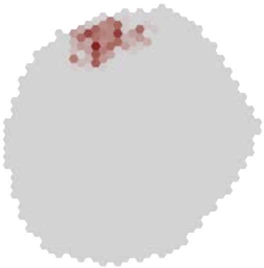

MeTu1_720575940616343380

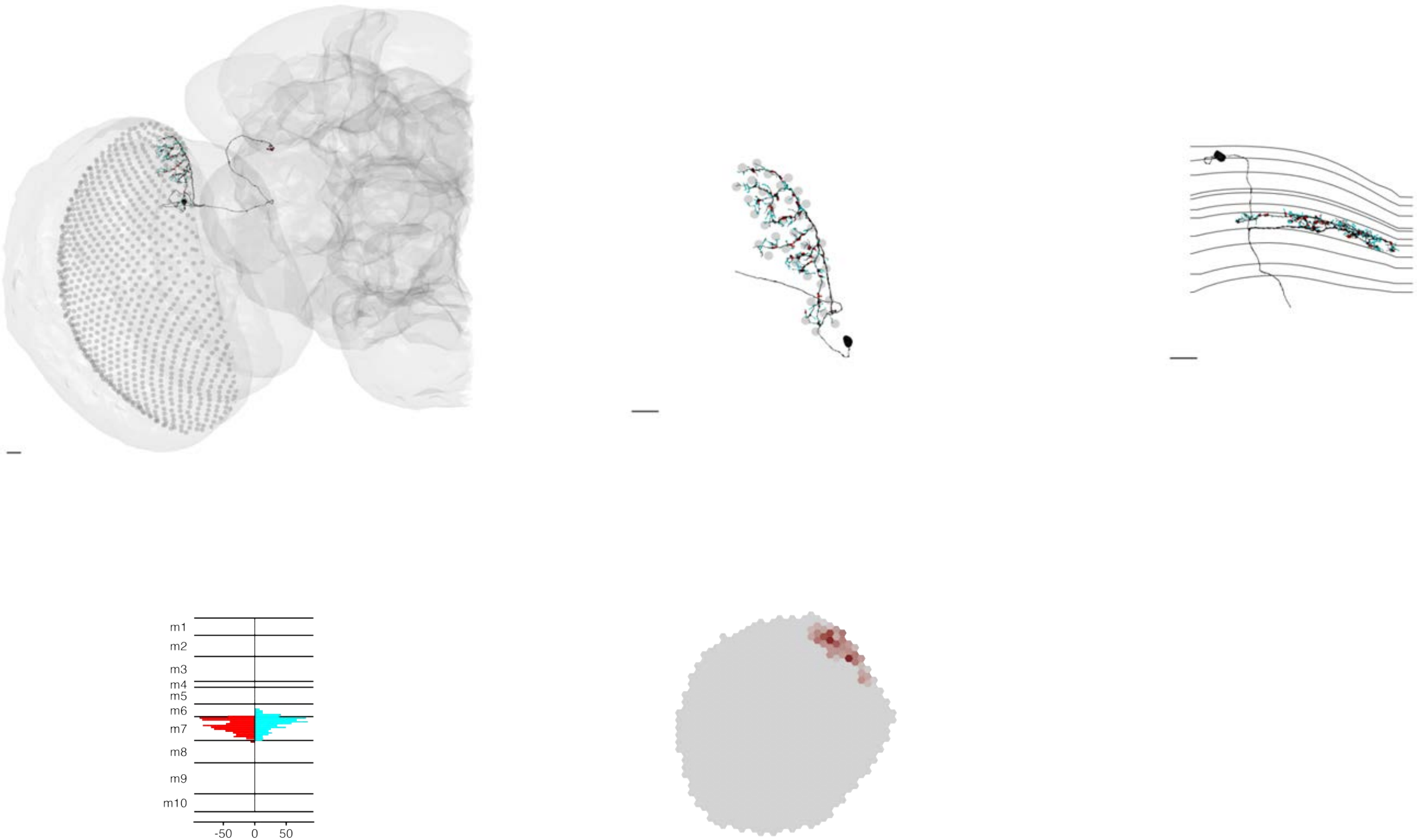

### MeTu1_720575940616672349

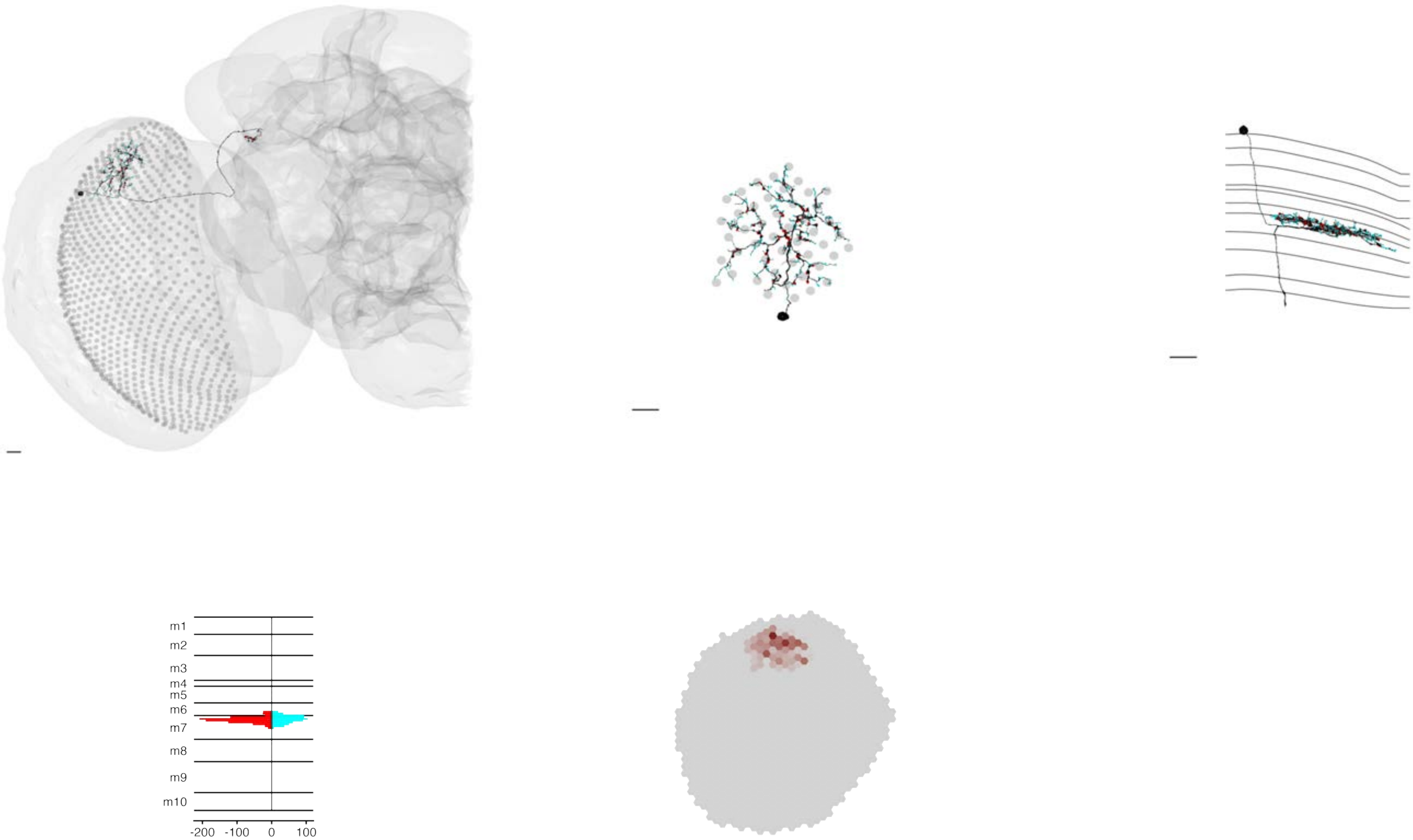

### MeTu1_720575940617183134

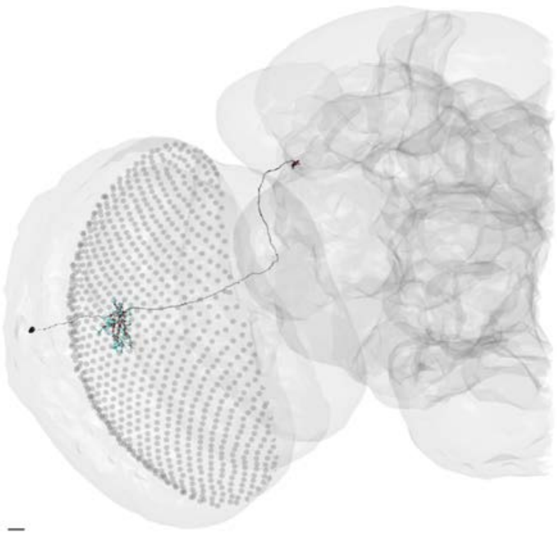

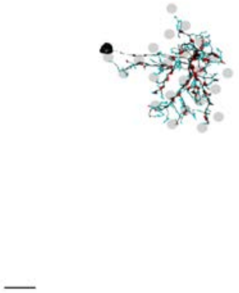

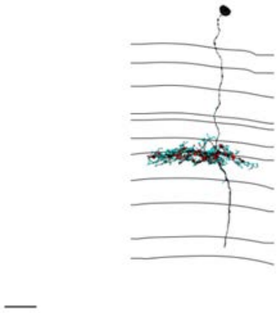

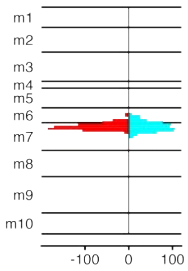


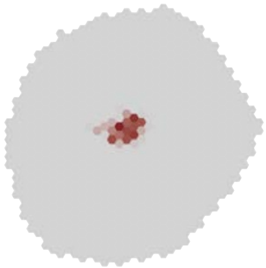

### MeTu1_720575940617282117

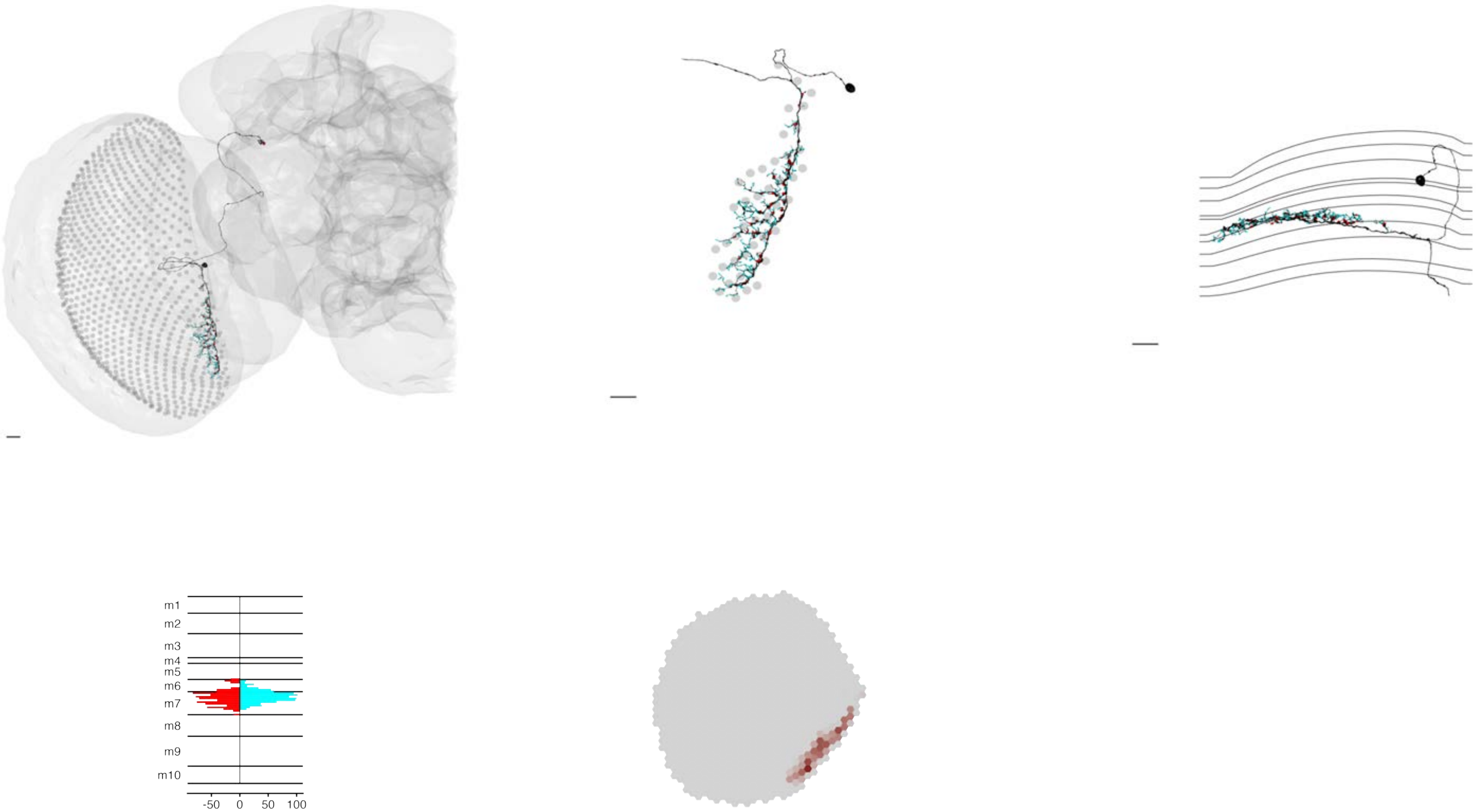

### MeTu1_720575940617283101

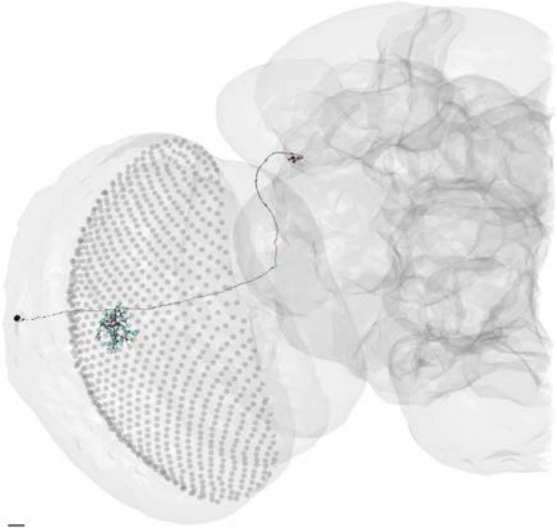

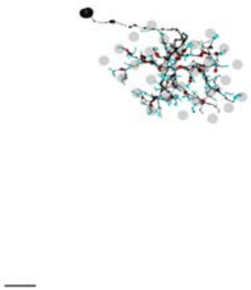

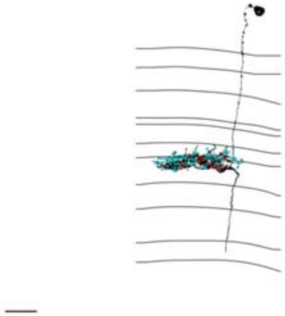

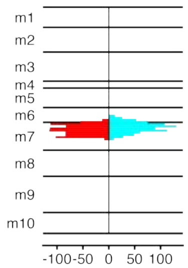


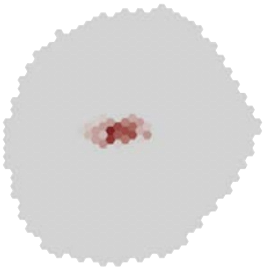

### MeTu1_720575940617366745

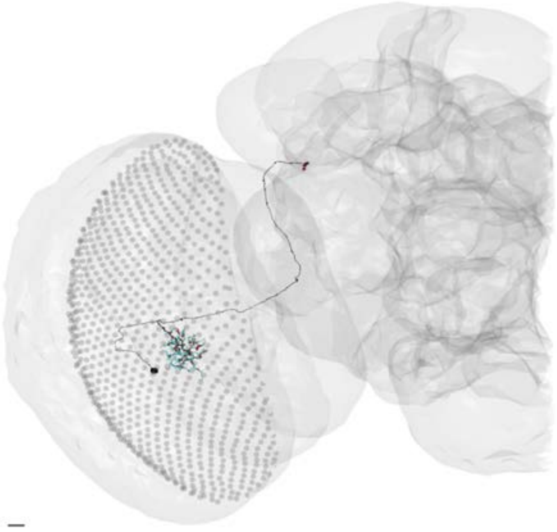

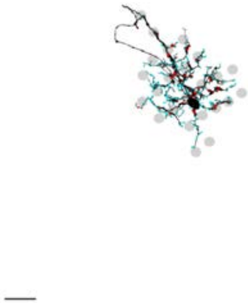

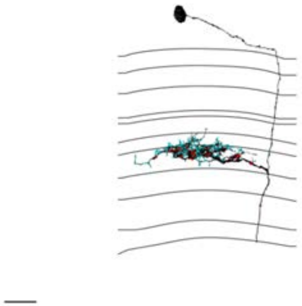

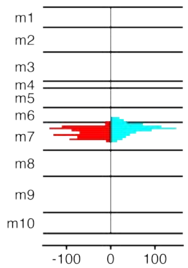


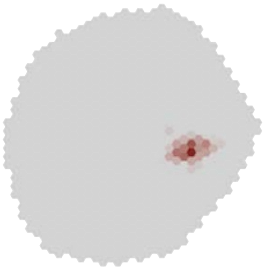

### MeTu1_720575940618408943

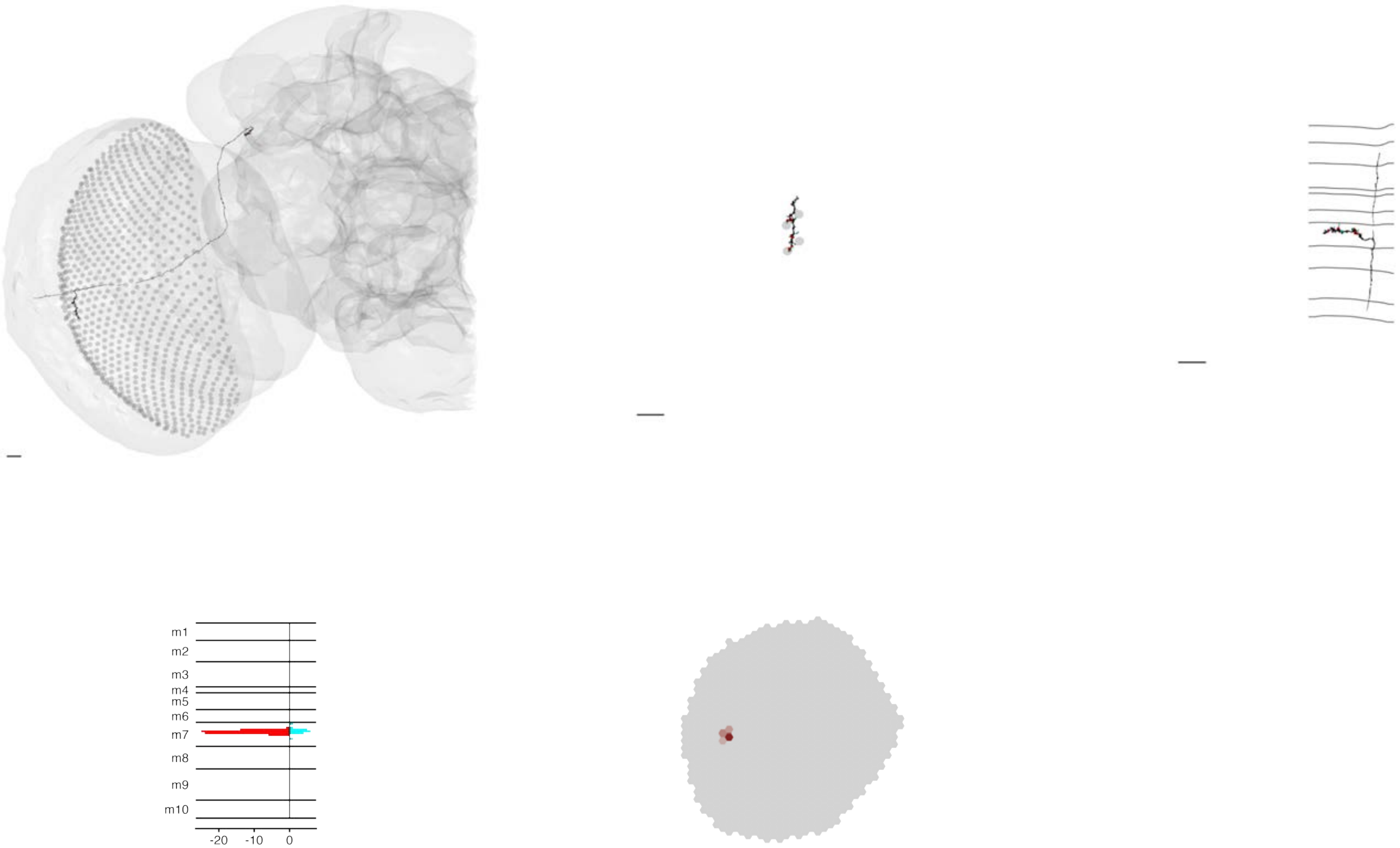

### MeTu1_720575940618971227

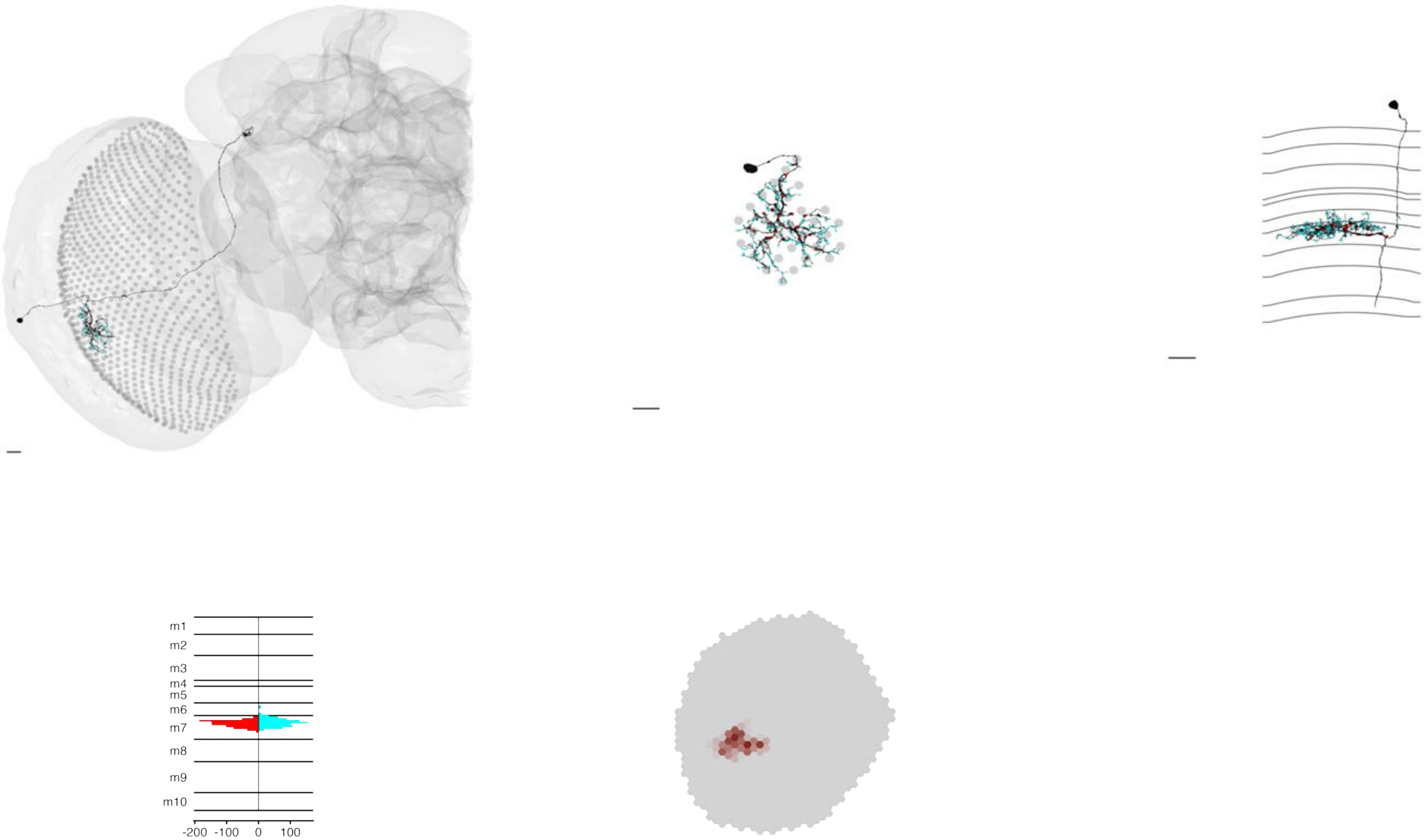

### MeTu1_720575940619080497

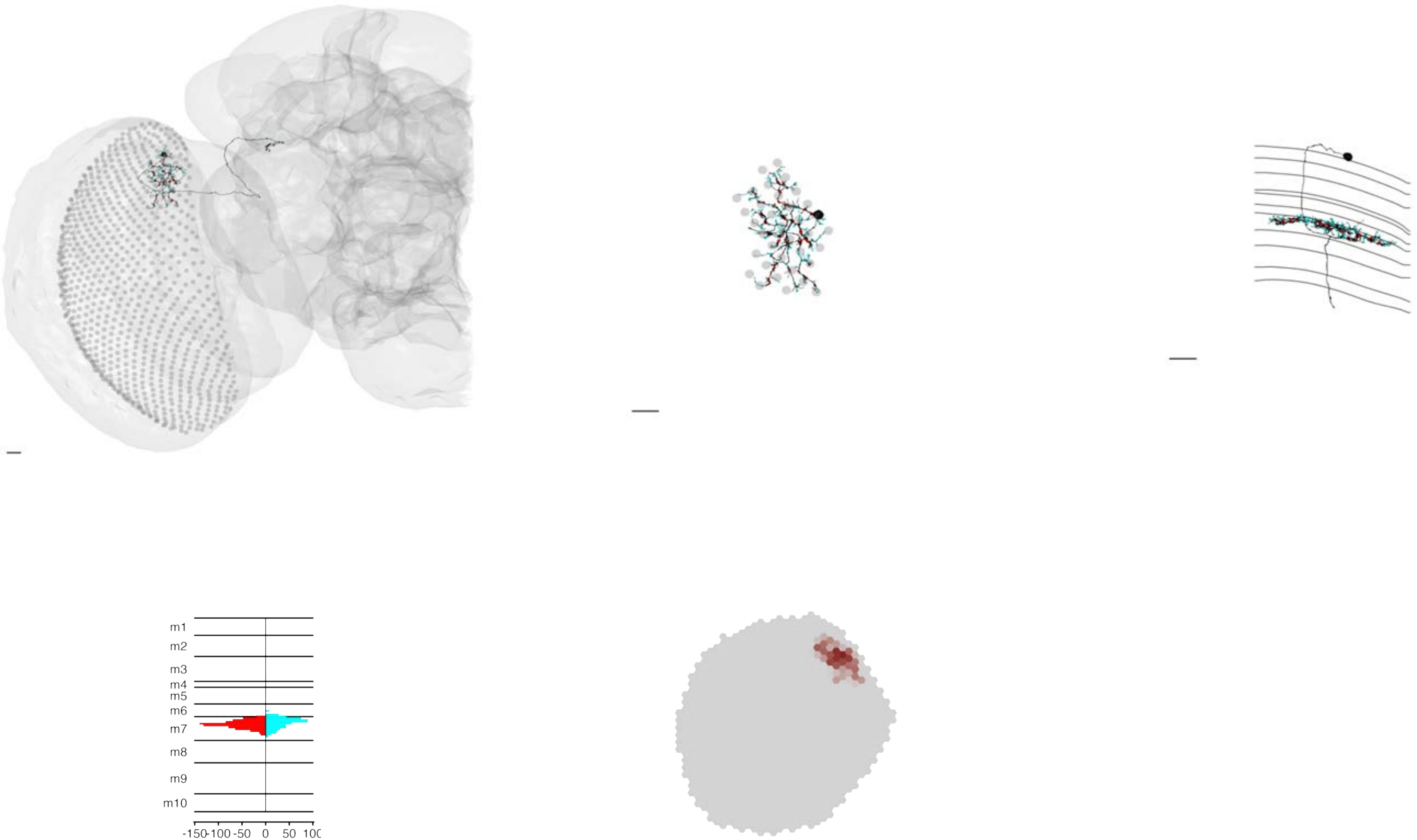

### MeTu1_720575940619191022

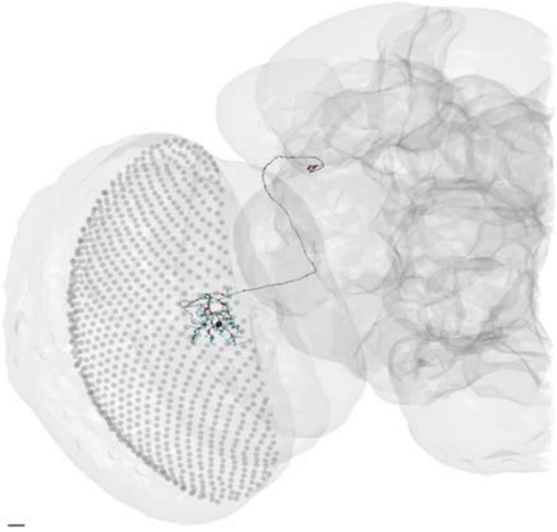

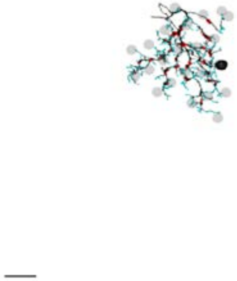

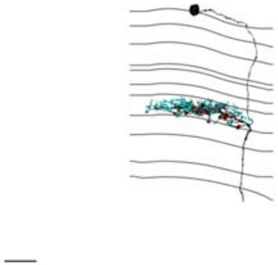

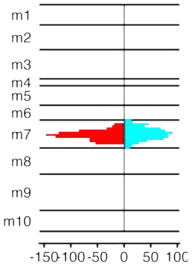


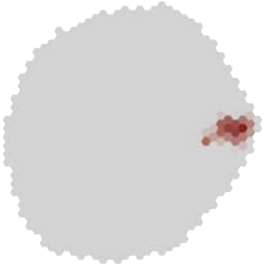

### MeTu1_720575940619661150

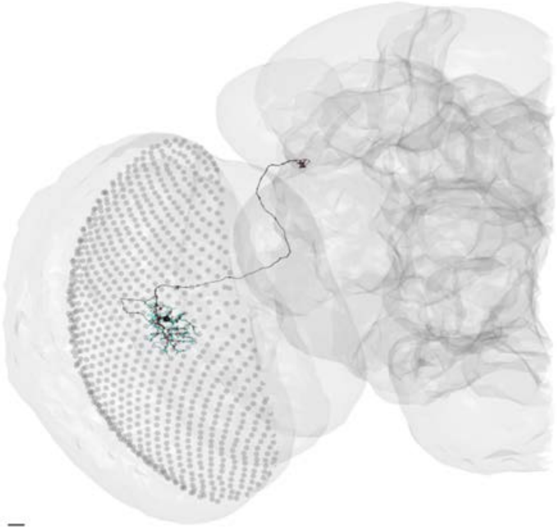

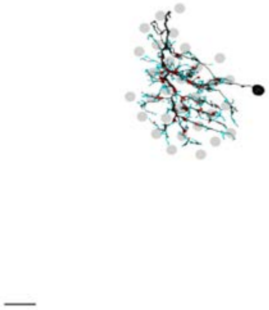

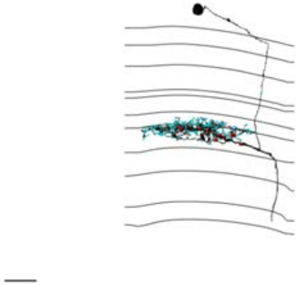

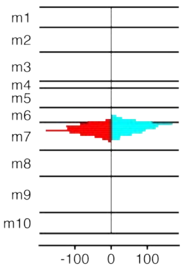


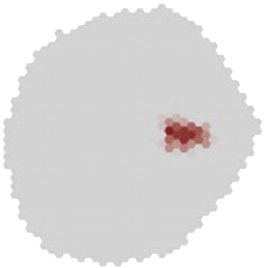

### MeTu1_720575940619687222

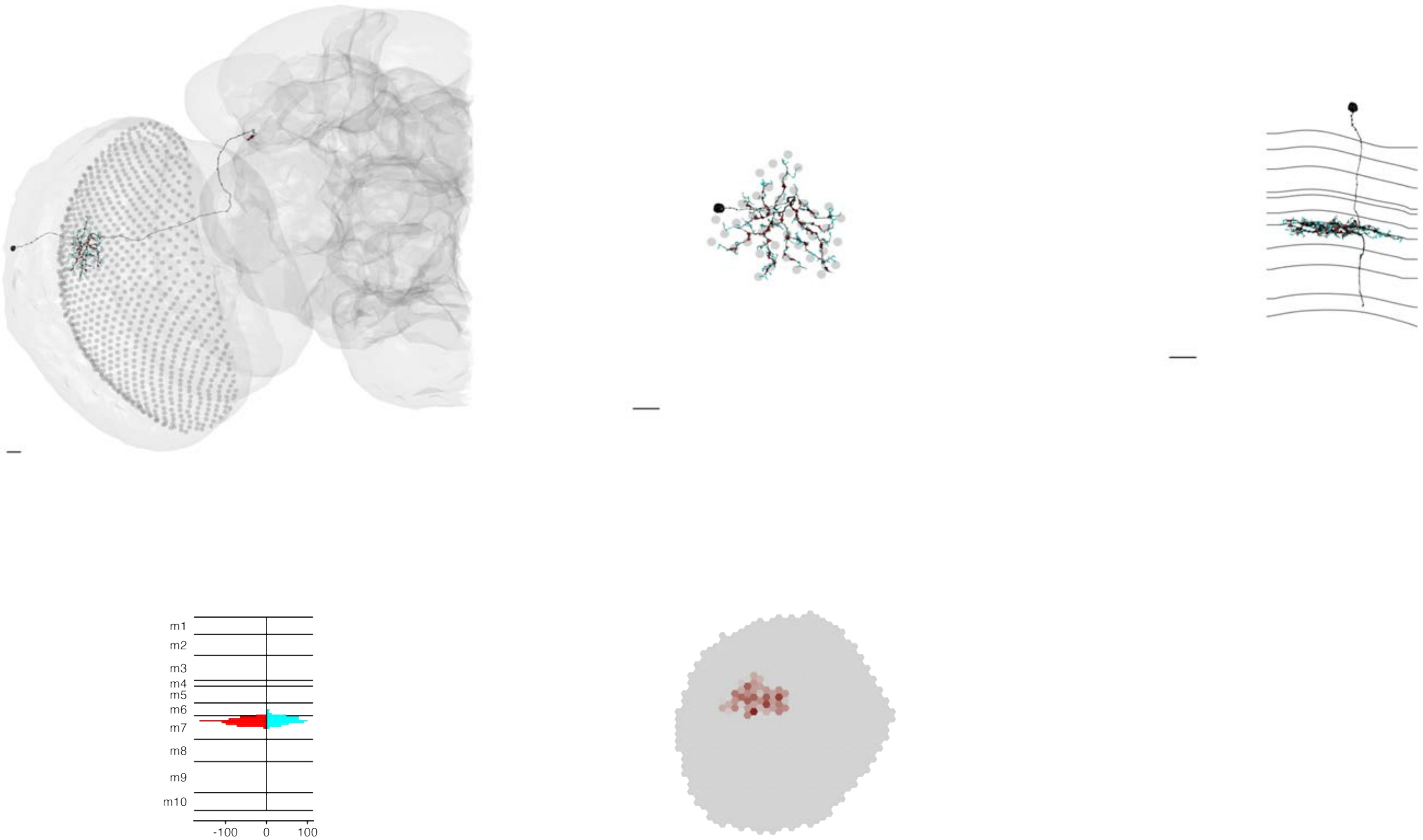

MeTu1_720575940619883758

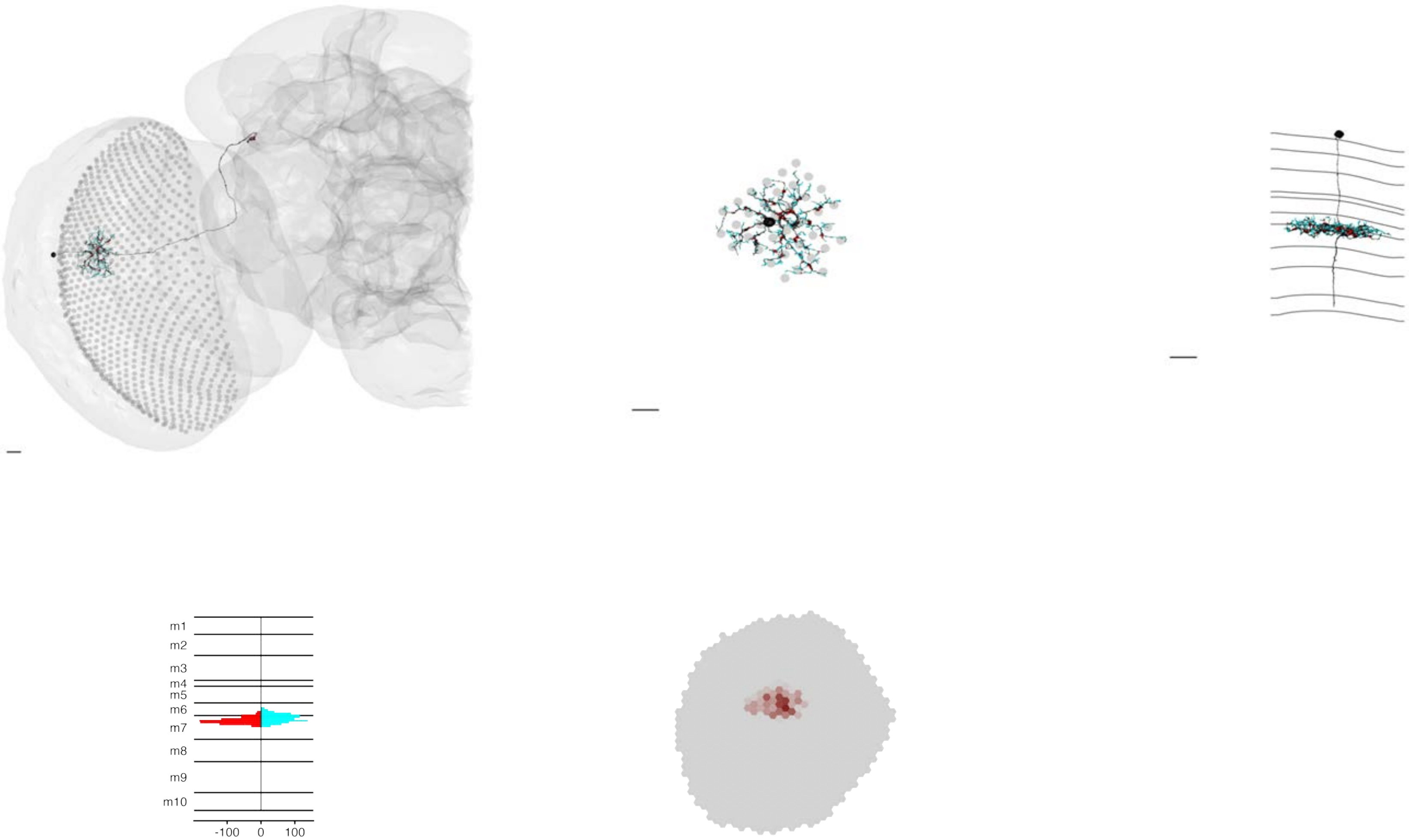

MeTu1_720575940620019265

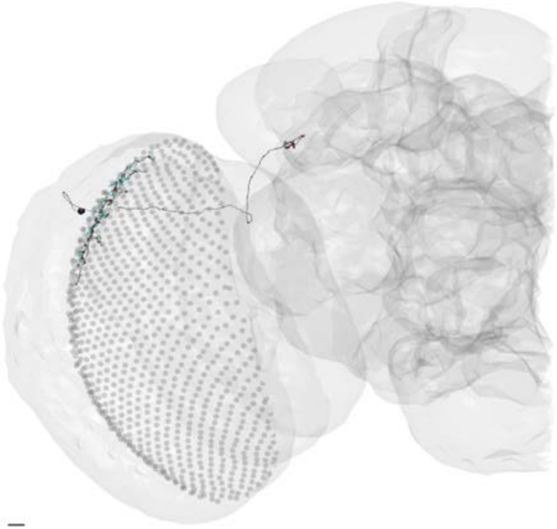

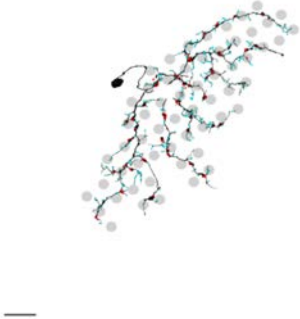

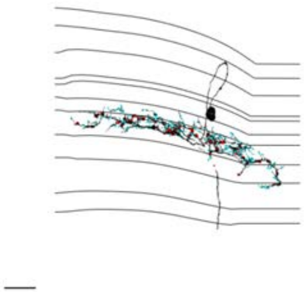

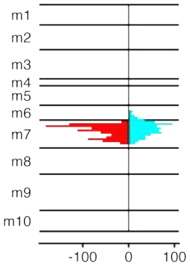


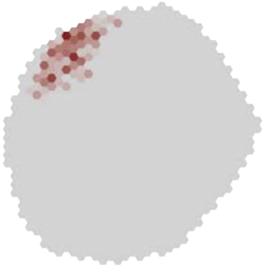

### MeTu1_720575940620183316

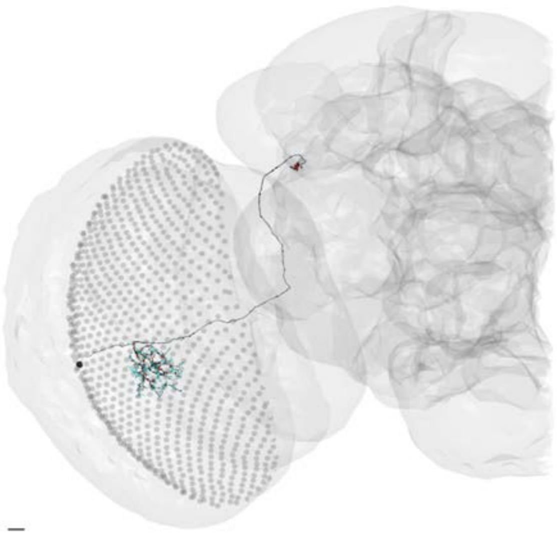

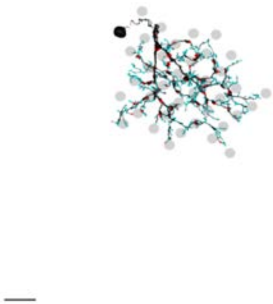

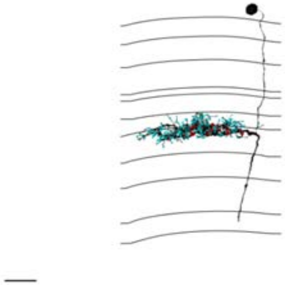

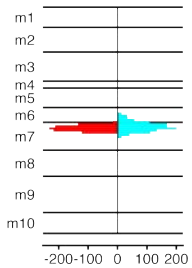


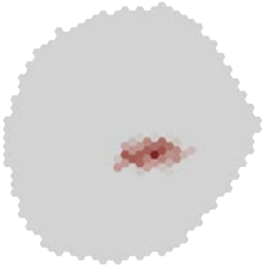

### MeTu1_720575940620336929

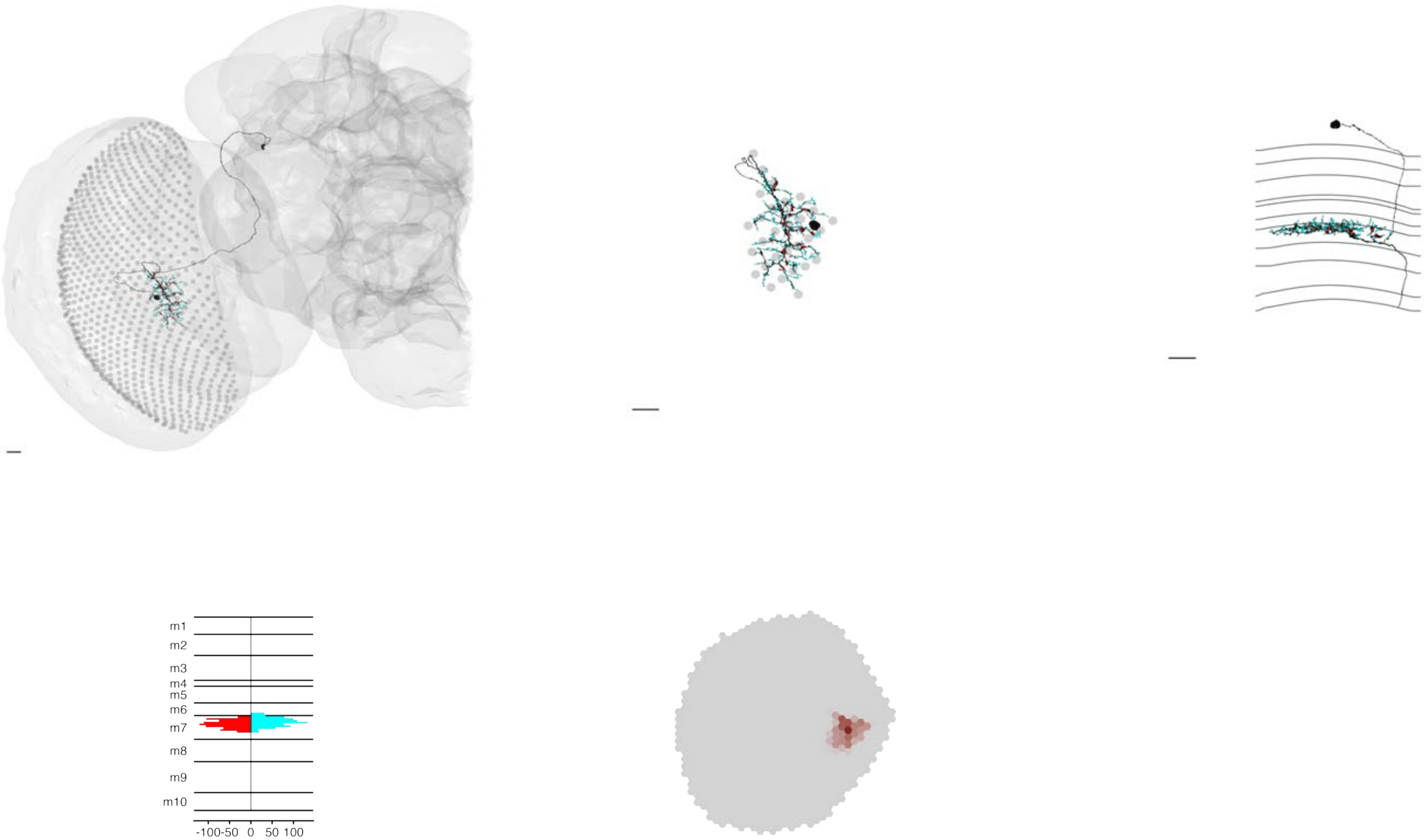

### MeTu1_720575940620553229

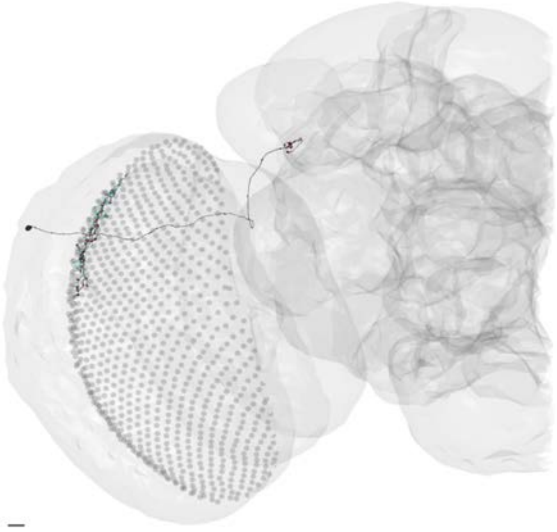

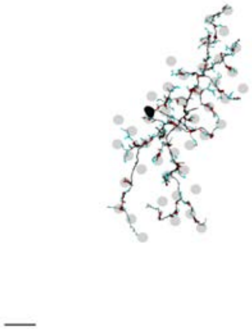

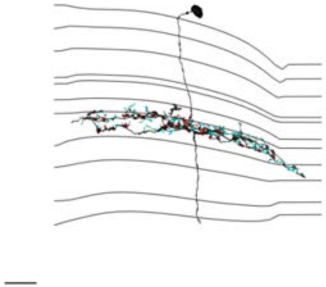

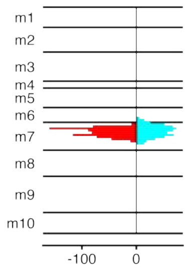


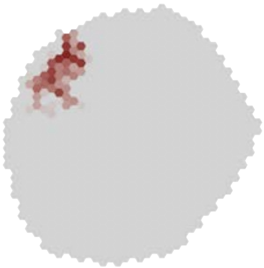

### MeTu1_720575940620650661

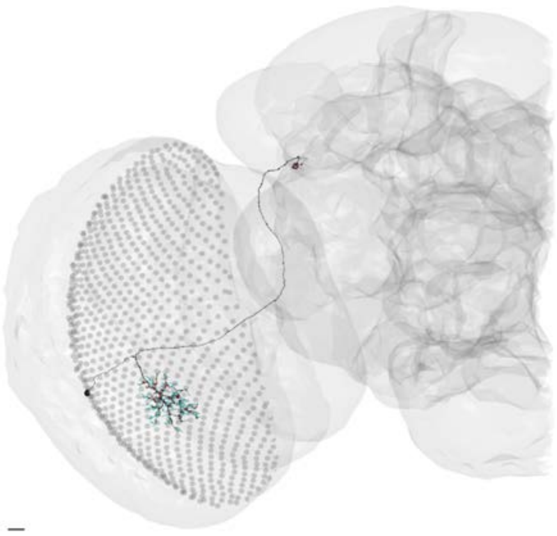

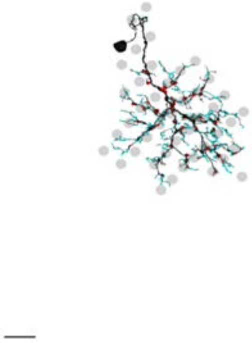

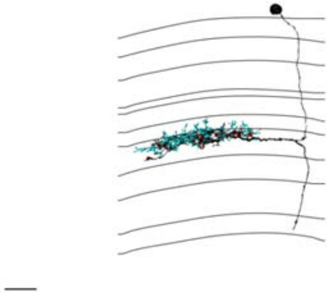

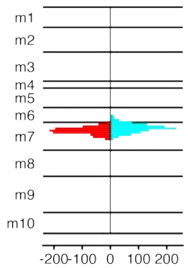


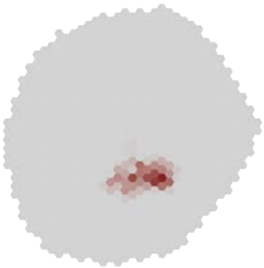

MeTu1_720575940620785030

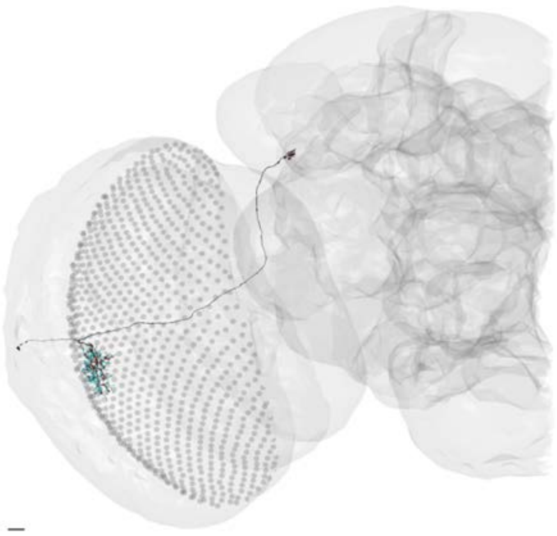

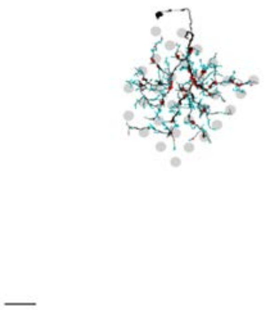

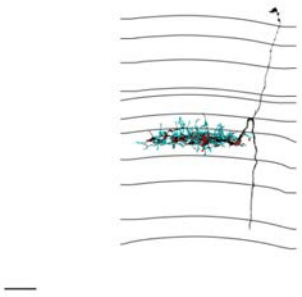

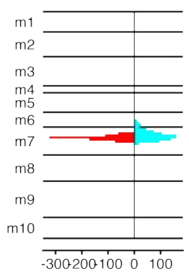


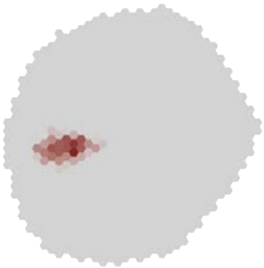

### MeTu1_720575940620986495

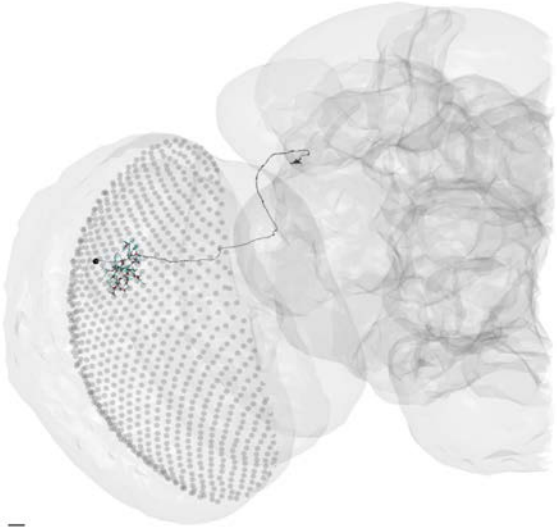

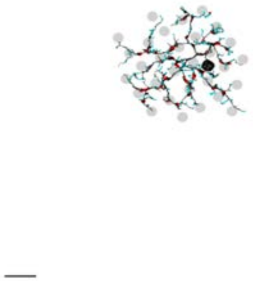

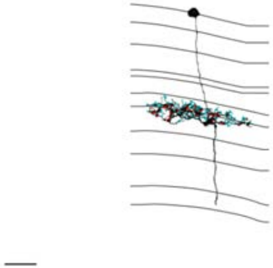

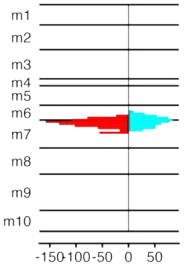


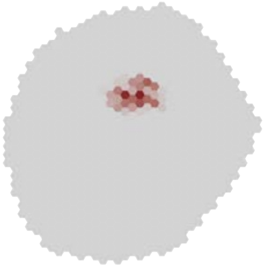

### MeTu1_720575940621116353

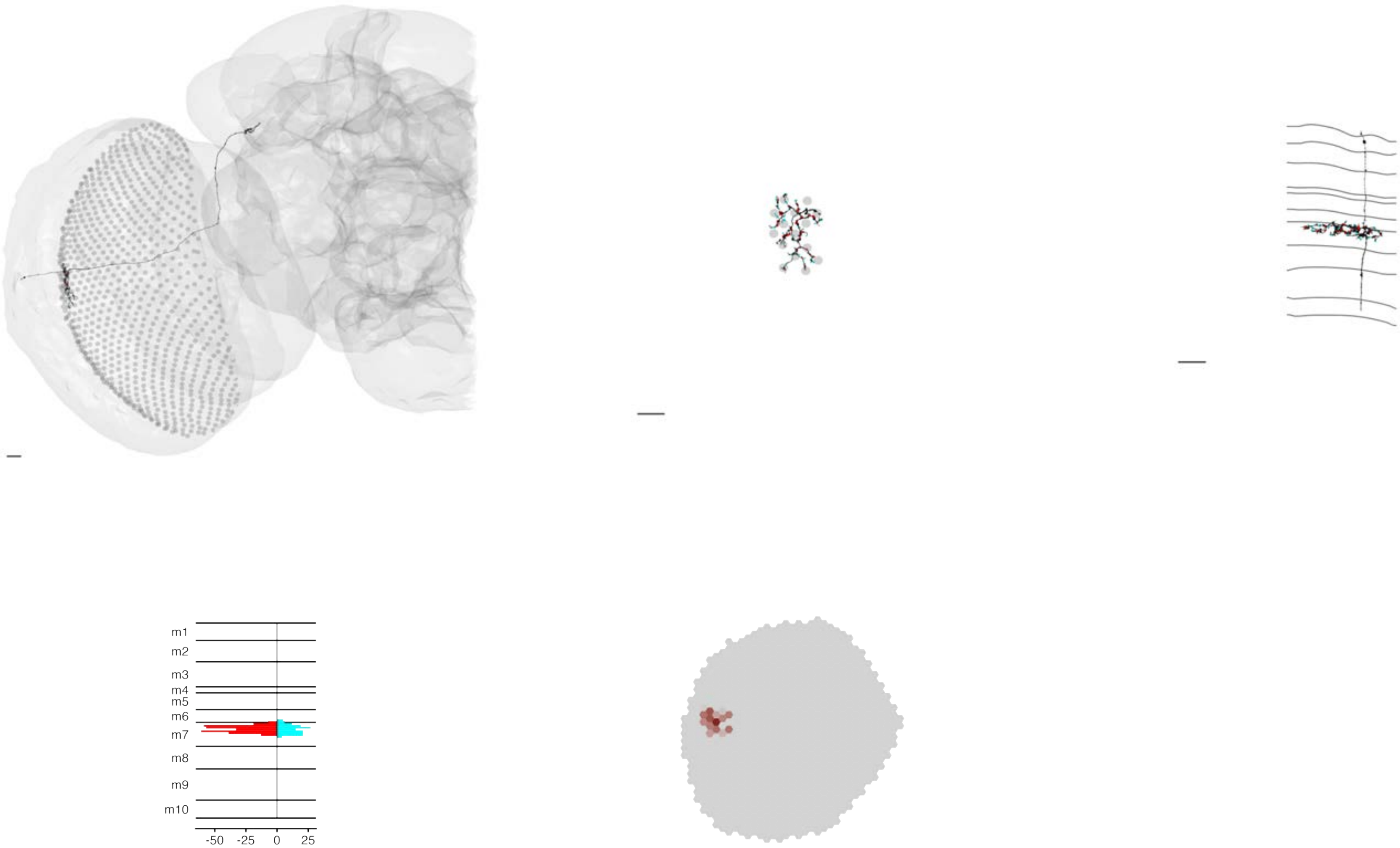

### MeTu1_720575940621299369

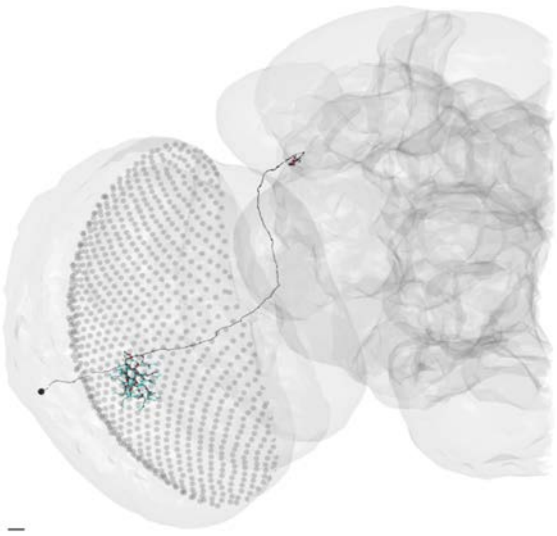

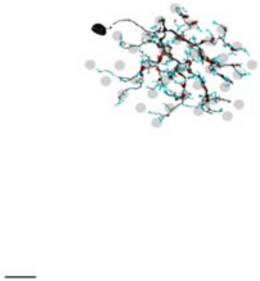

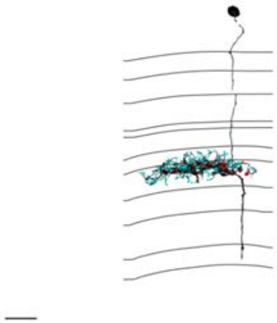

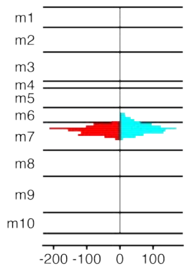


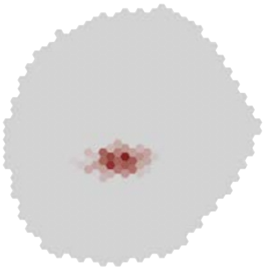

### MeTu1_720575940621434807

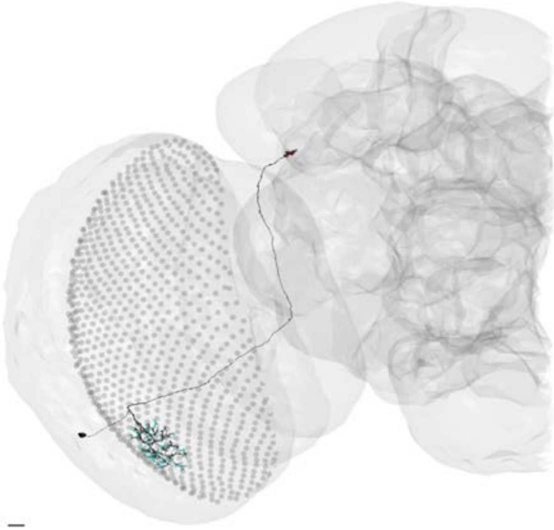

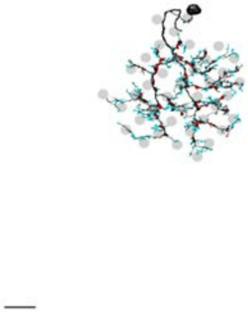

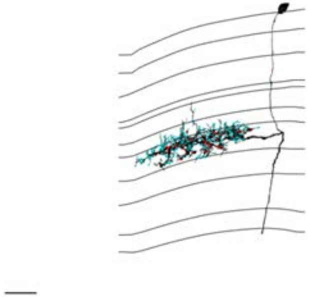

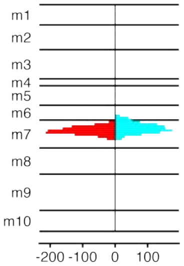


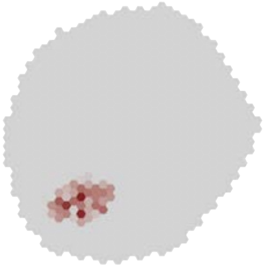

### MeTu1_720575940621485677

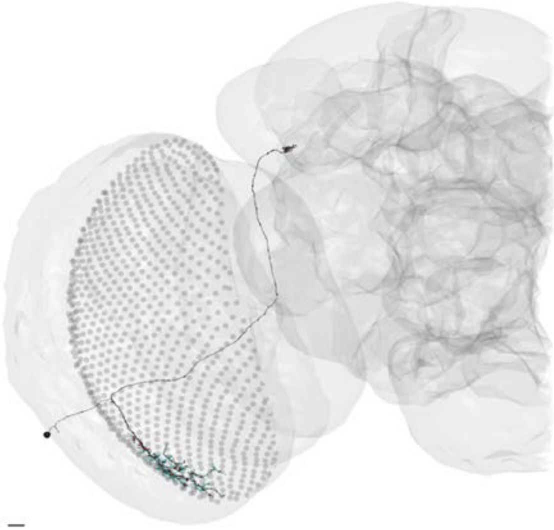

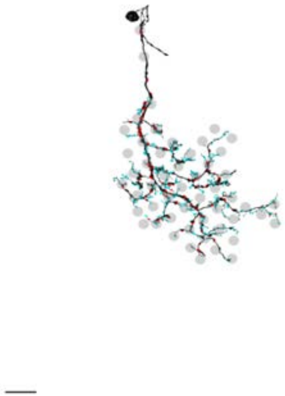

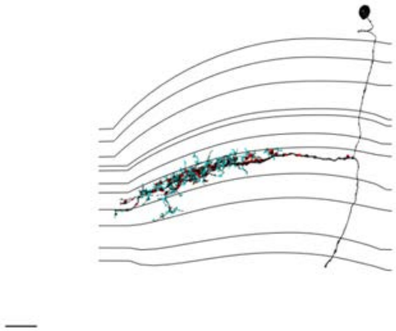

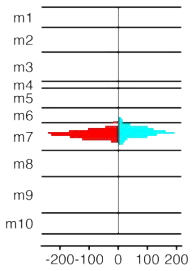


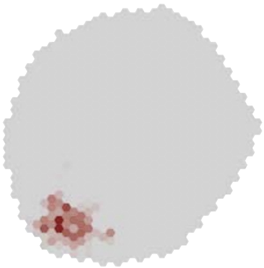

MeTu1_720575940621610572

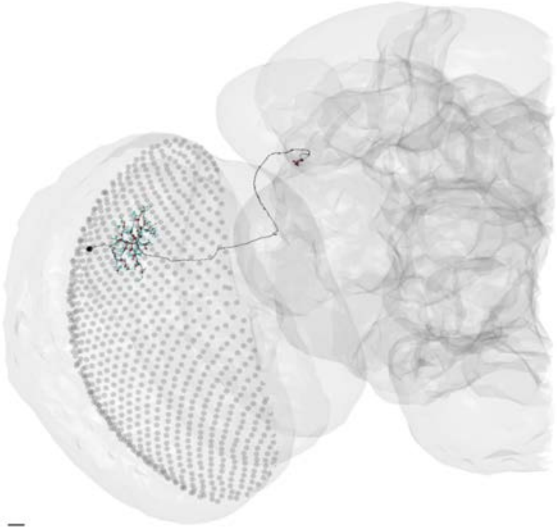

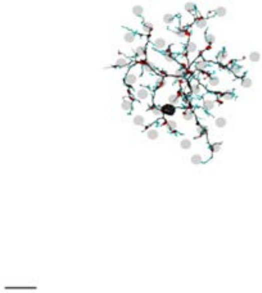

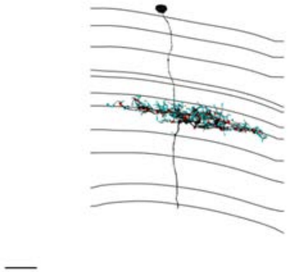

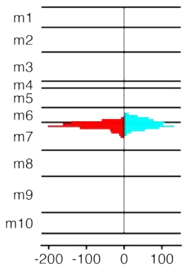


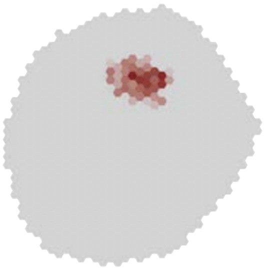

### MeTu1_720575940621804966

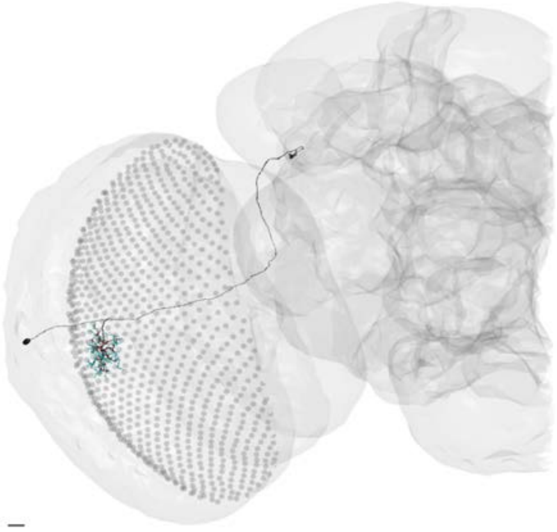

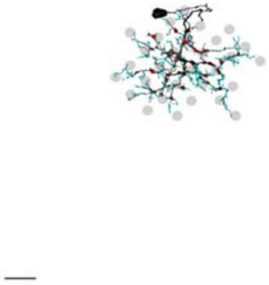

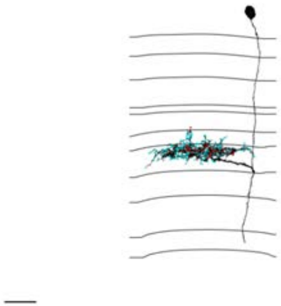

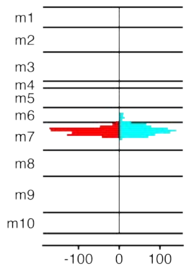


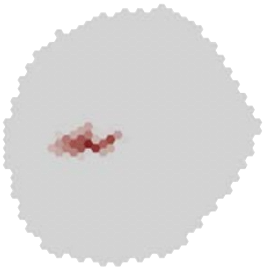

MeTu1_720575940621988829

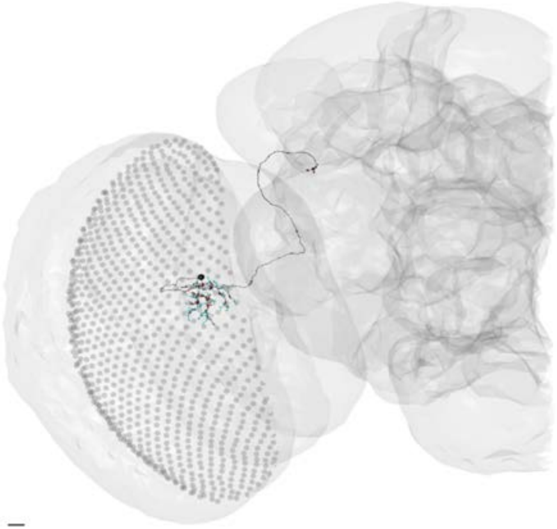

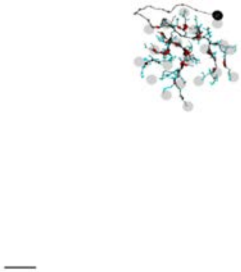

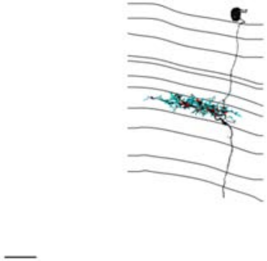

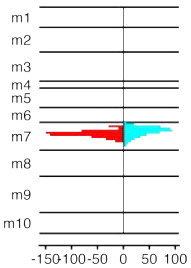


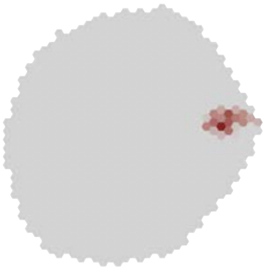

### MeTu1_720575940622147597

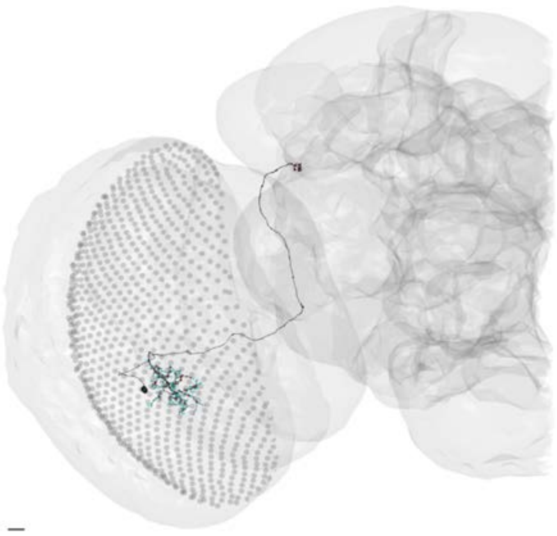

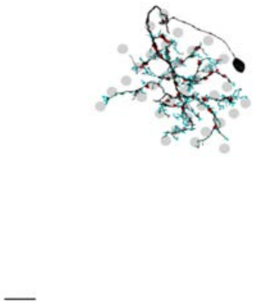

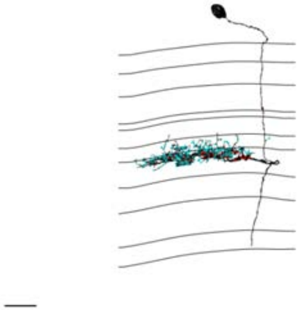

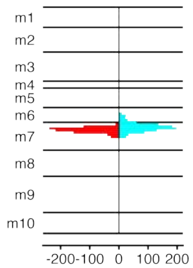


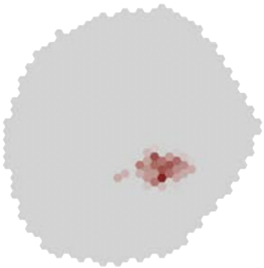

### MeTu1_720575940622287228

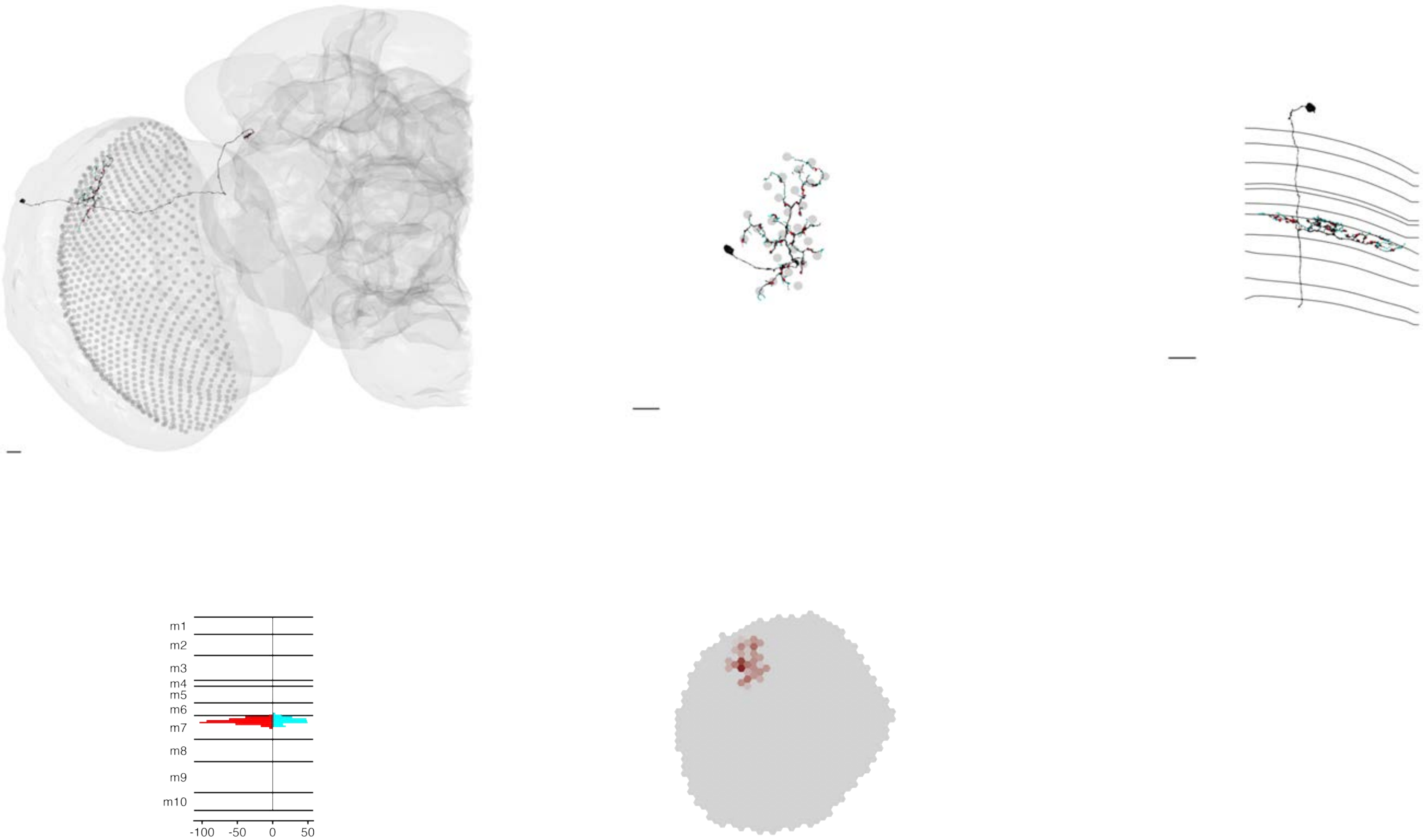

### MeTu1_720575940622449407

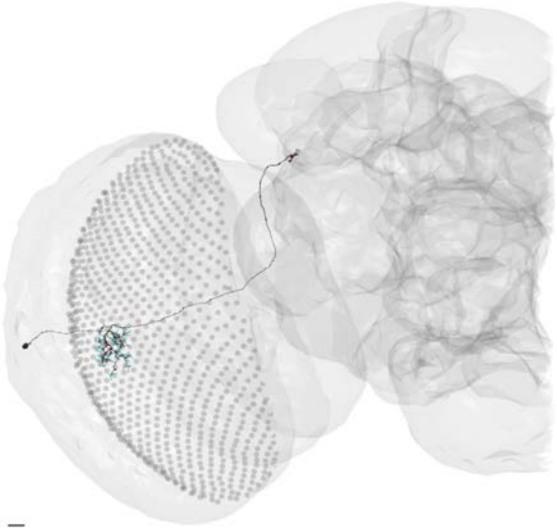

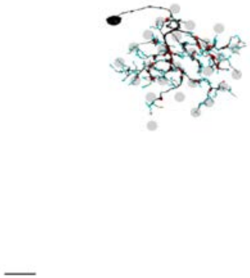

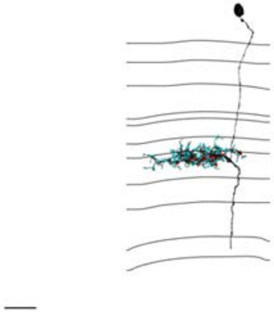

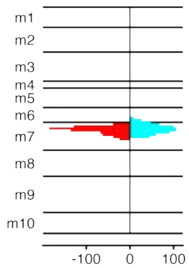


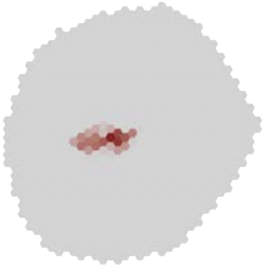

### MeTu1_720575940622647989

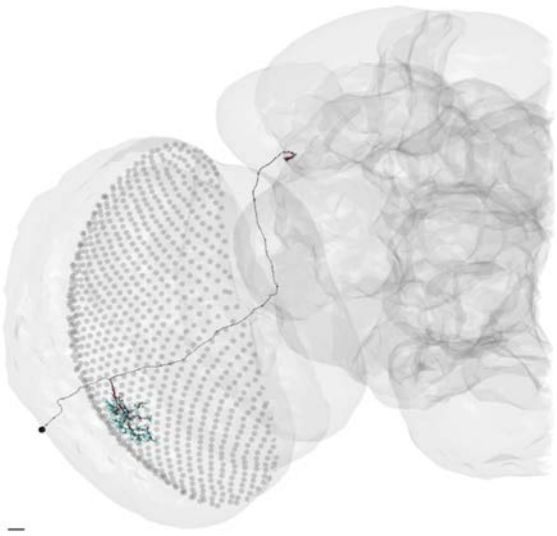

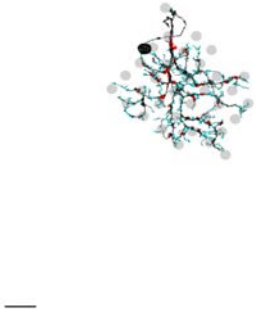

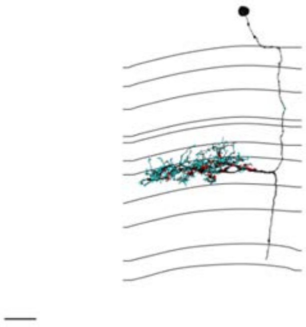

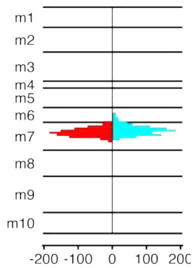


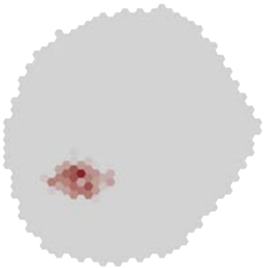

MeTu1_720575940622676488

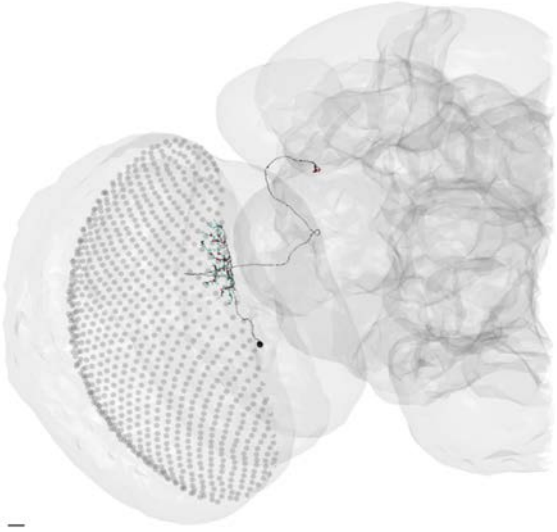

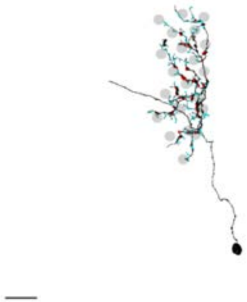

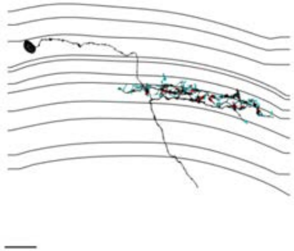

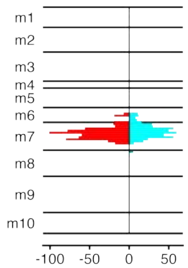


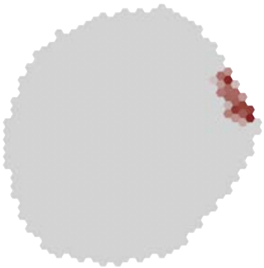

### MeTu1_720575940622739388

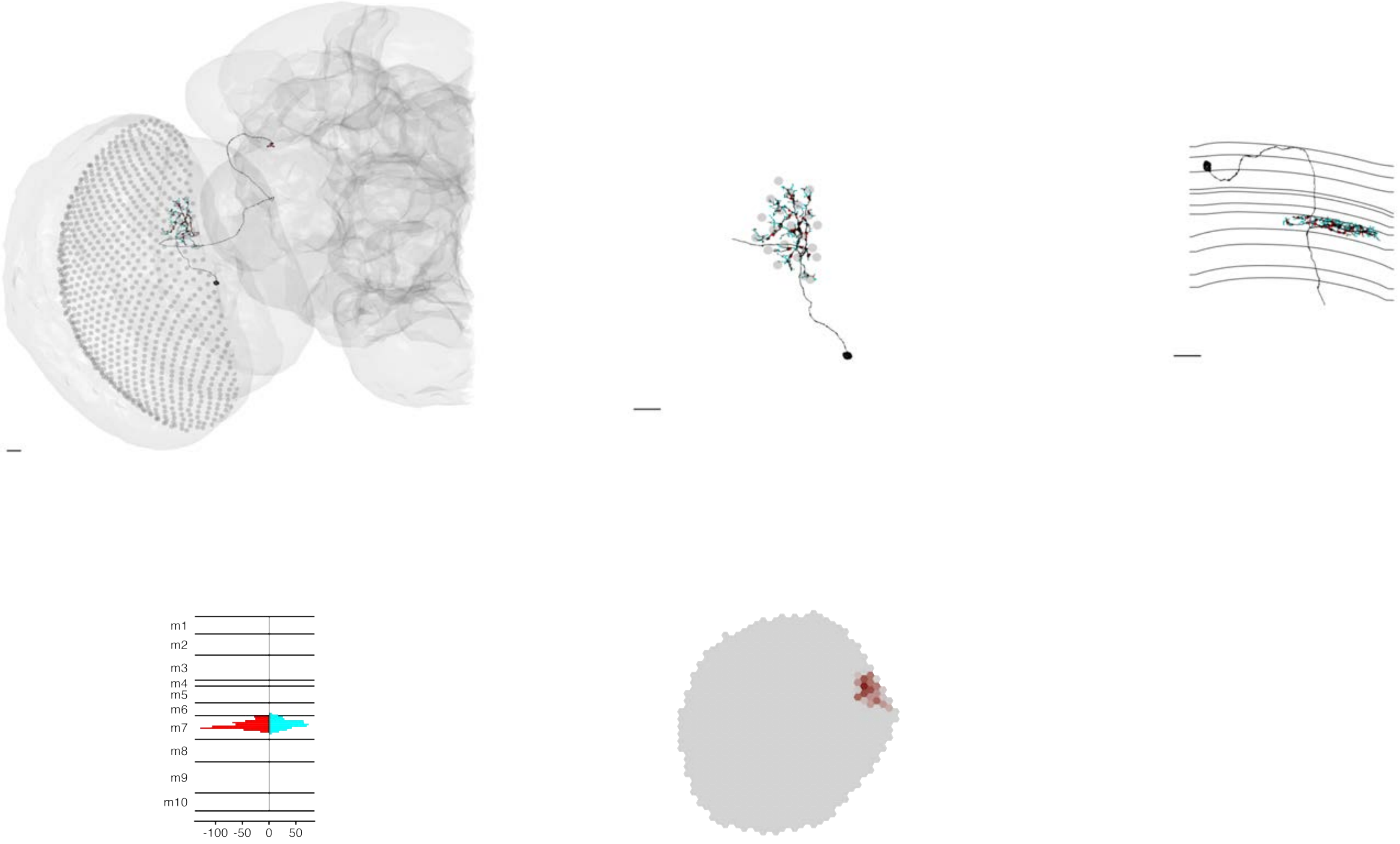

### MeTu1_720575940622745066

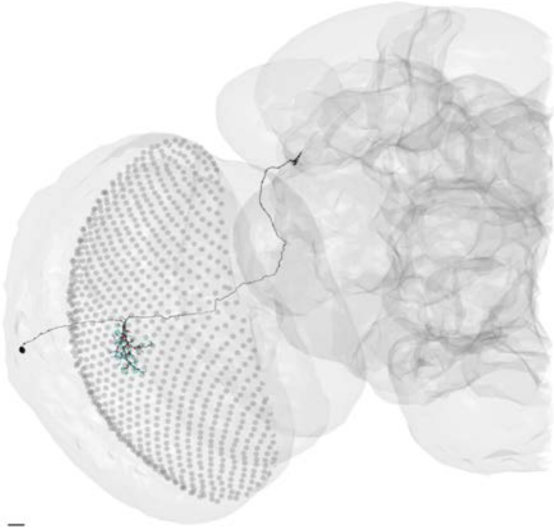

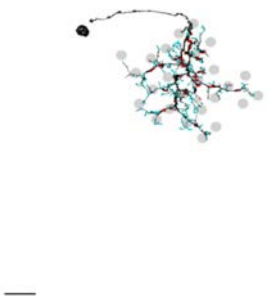

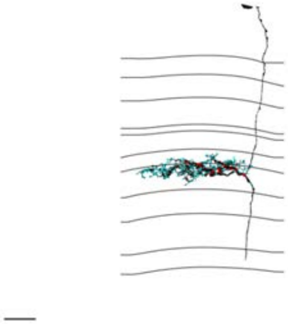

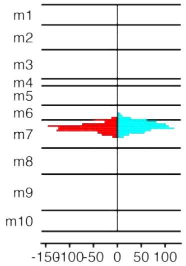


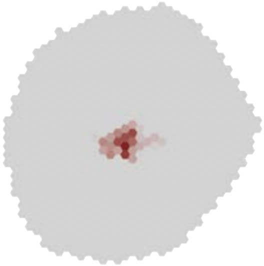

### MeTu1_720575940622759059

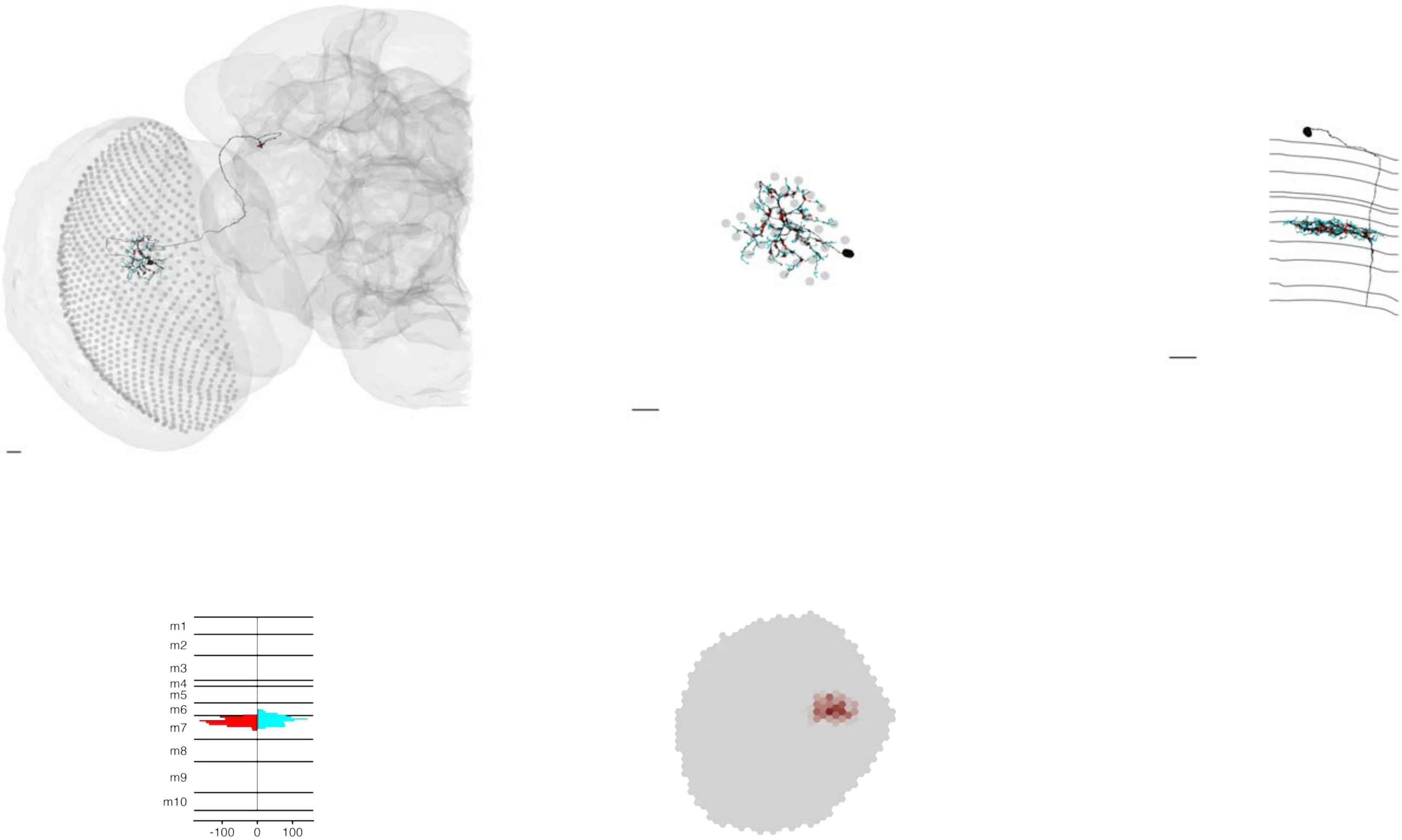

### MeTu1_720575940623246862

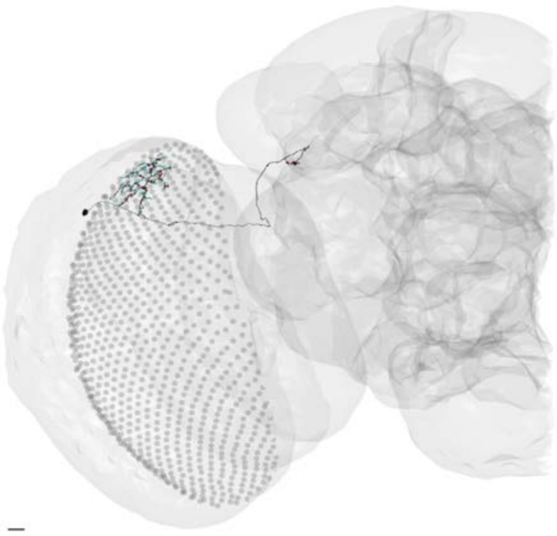

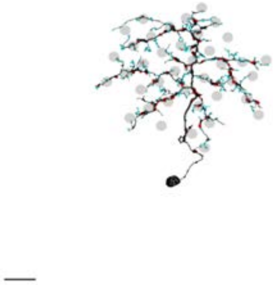

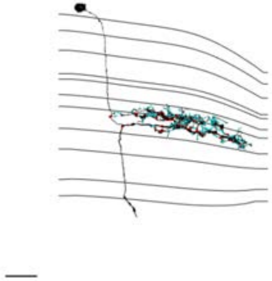

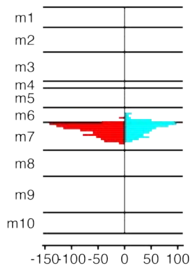


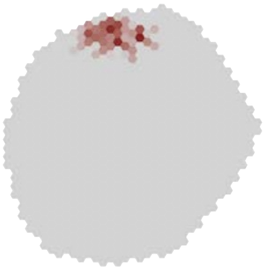

### MeTu1_720575940623262025

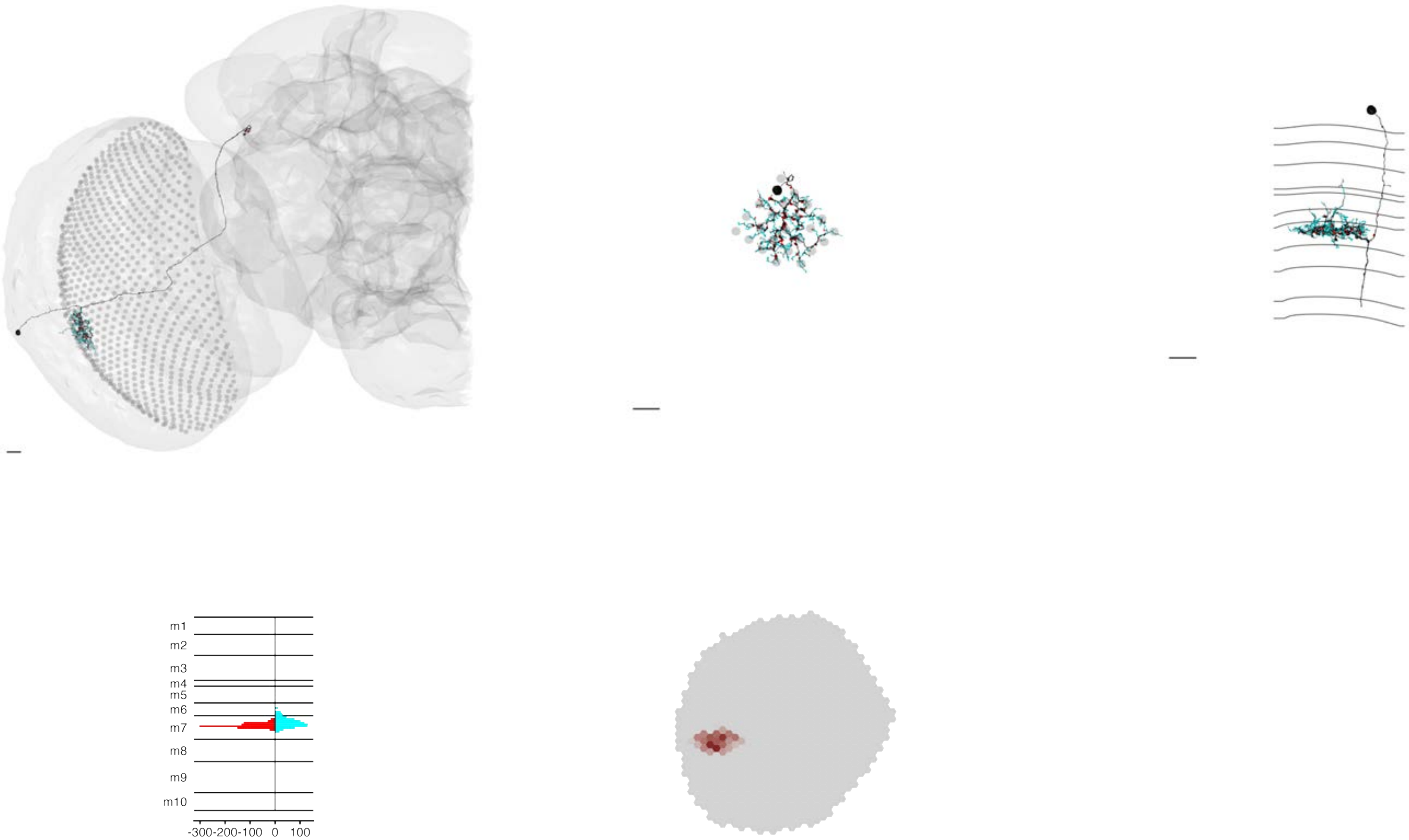

### MeTu1_720575940623295271

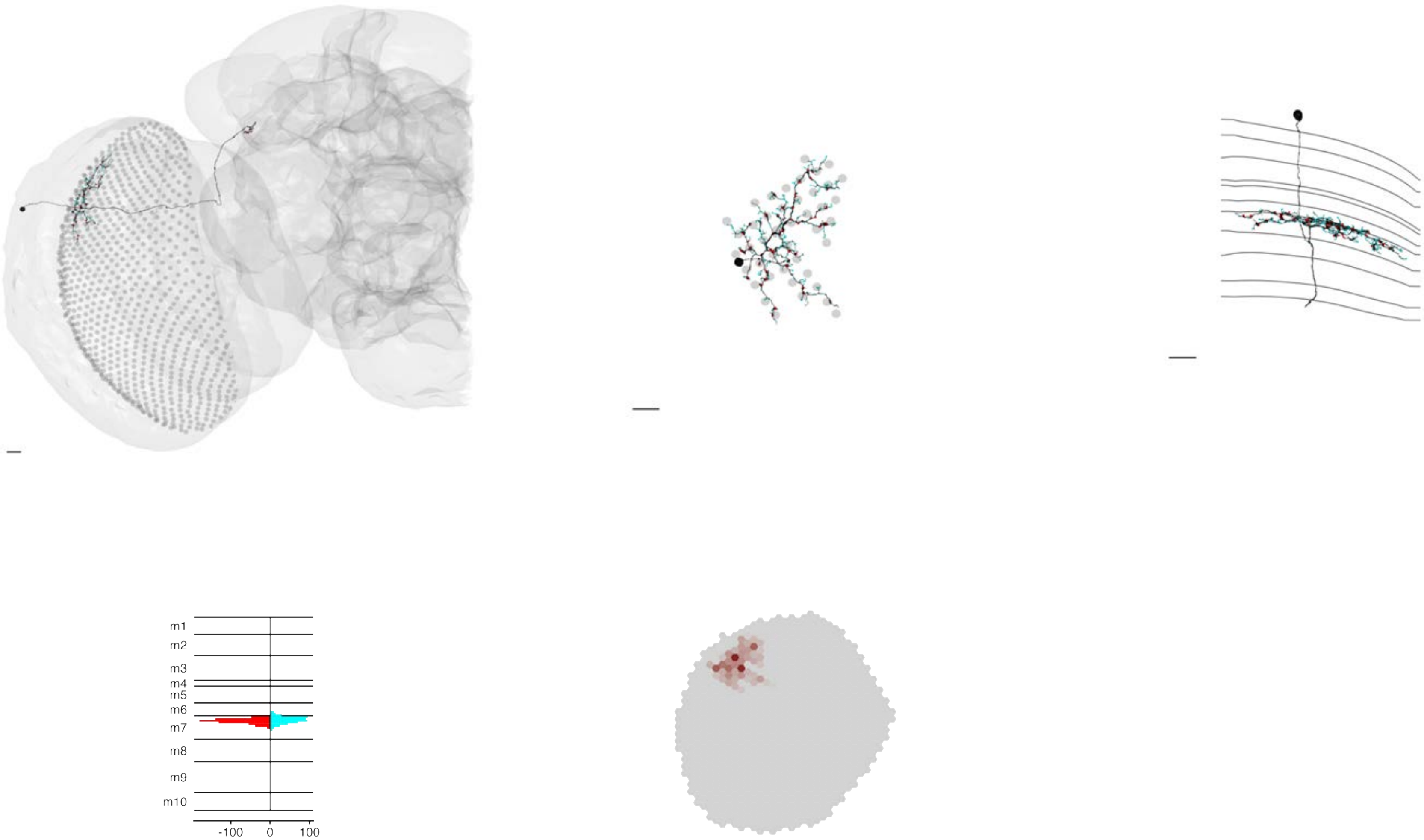

MeTu1_720575940623958271

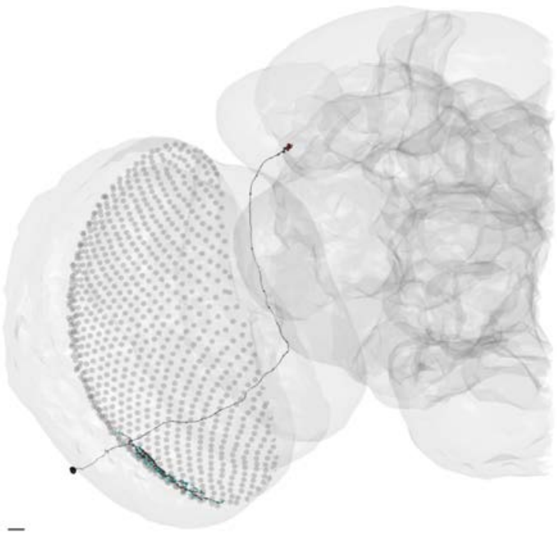

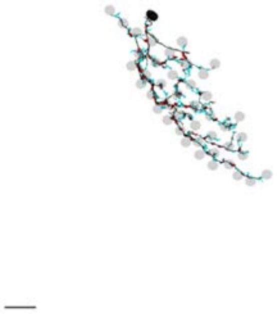

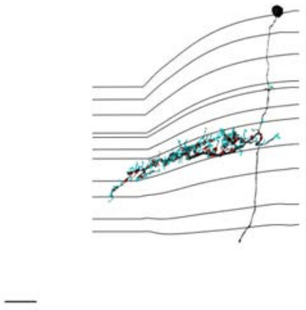

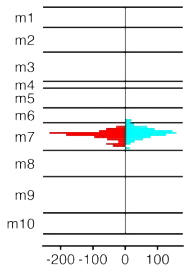


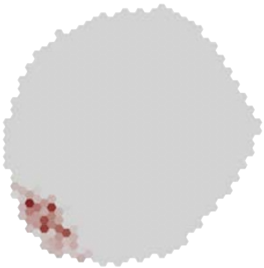

### MeTu1_720575940624007056

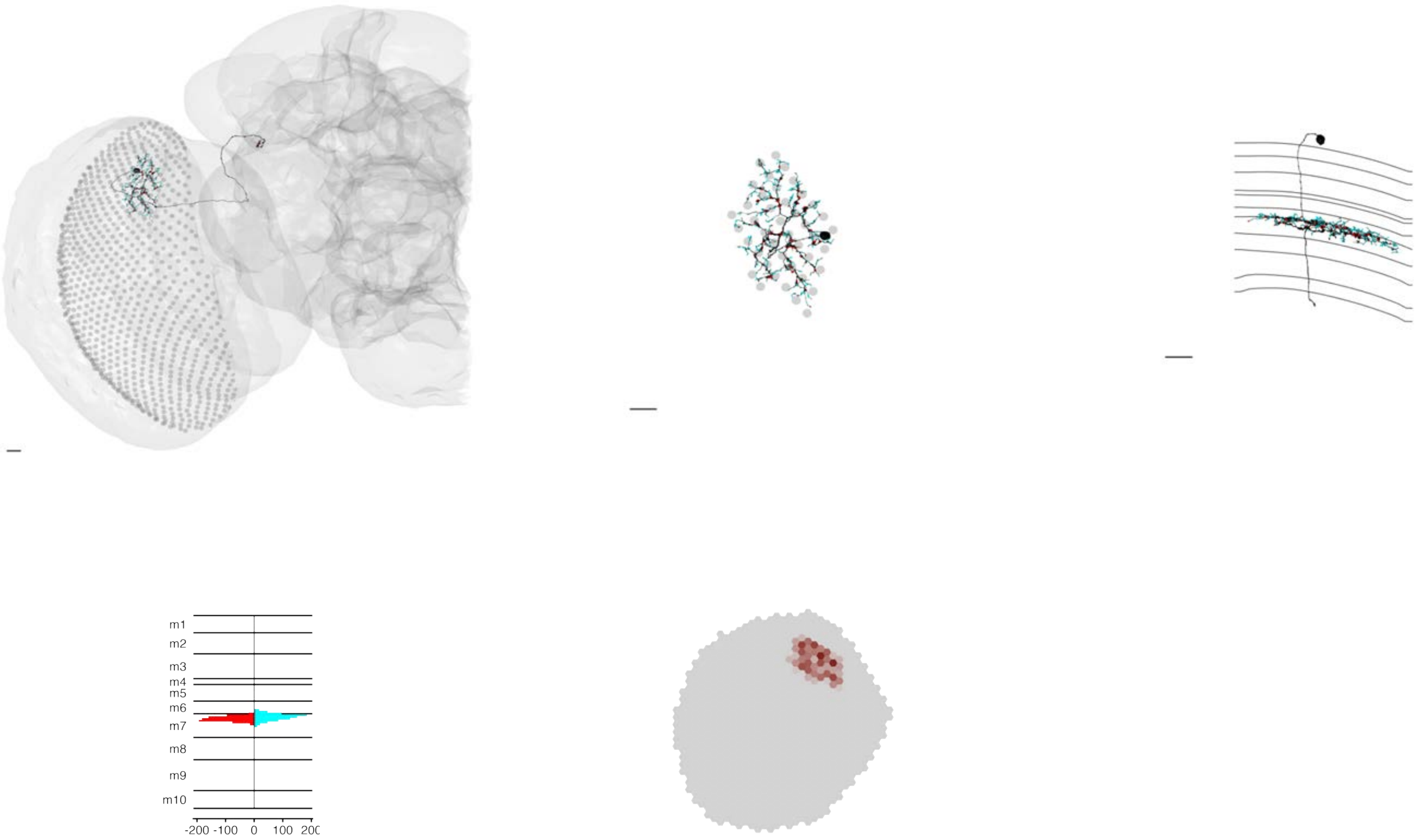

### MeTu1_720575940624063420

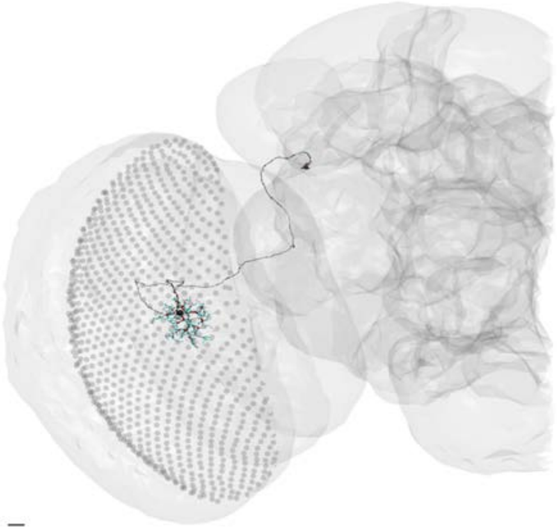

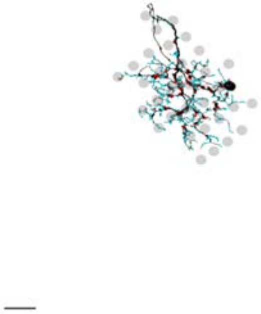

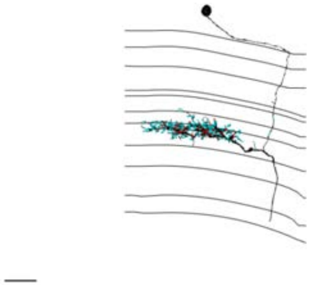

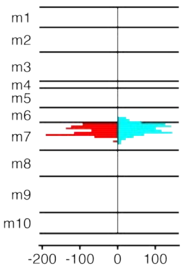


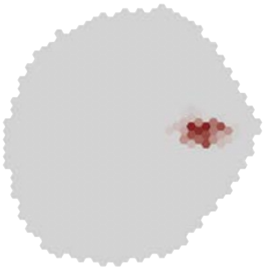

### MeTu1_720575940624176586

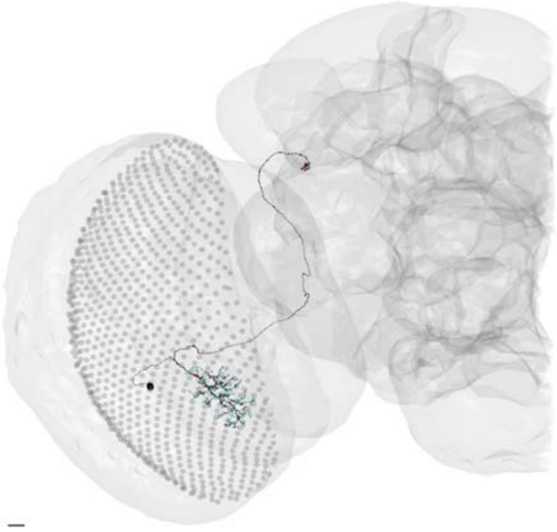

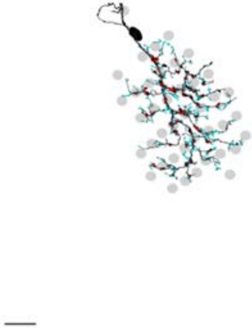

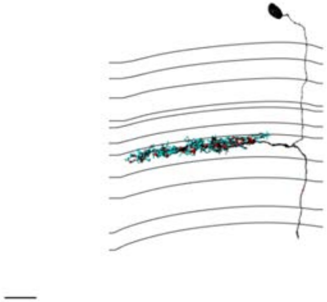

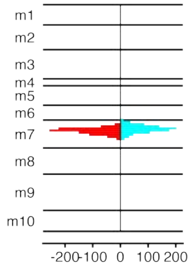


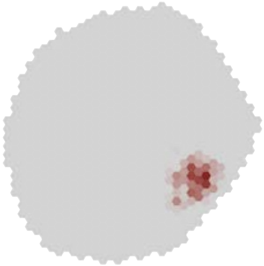

### MeTu1_720575940624187068

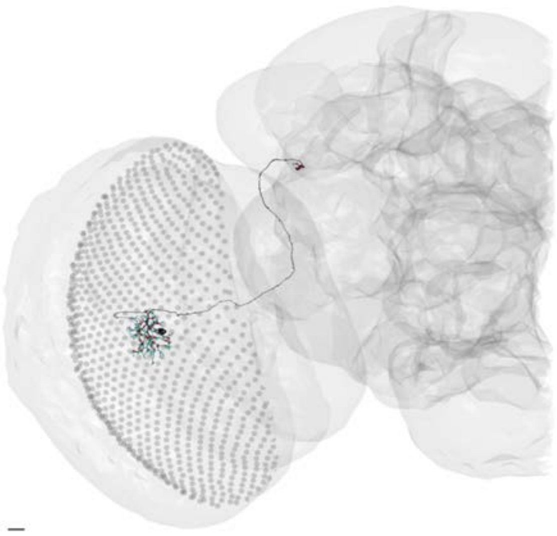

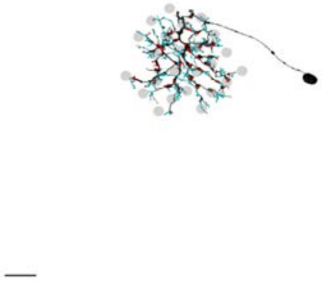

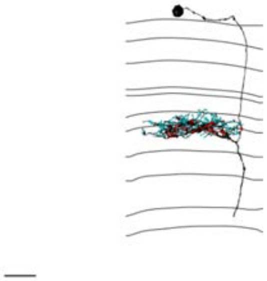

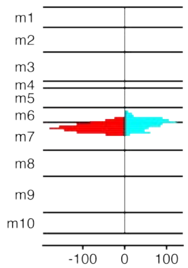


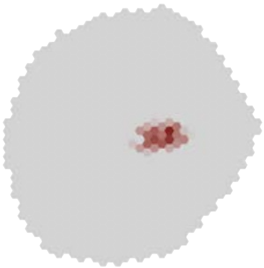

### MeTu1_720575940624225598

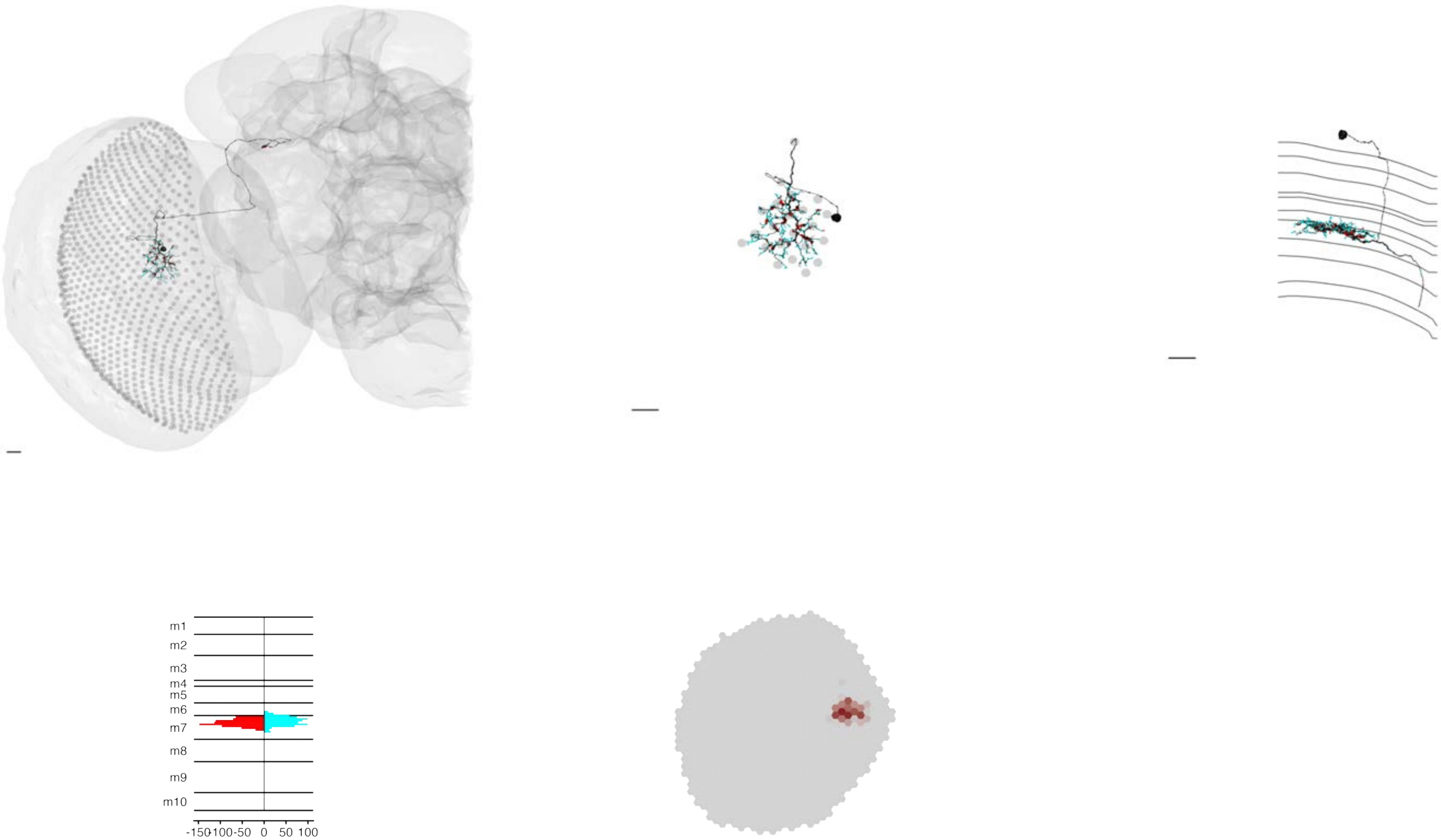

### MeTu1_720575940624322905

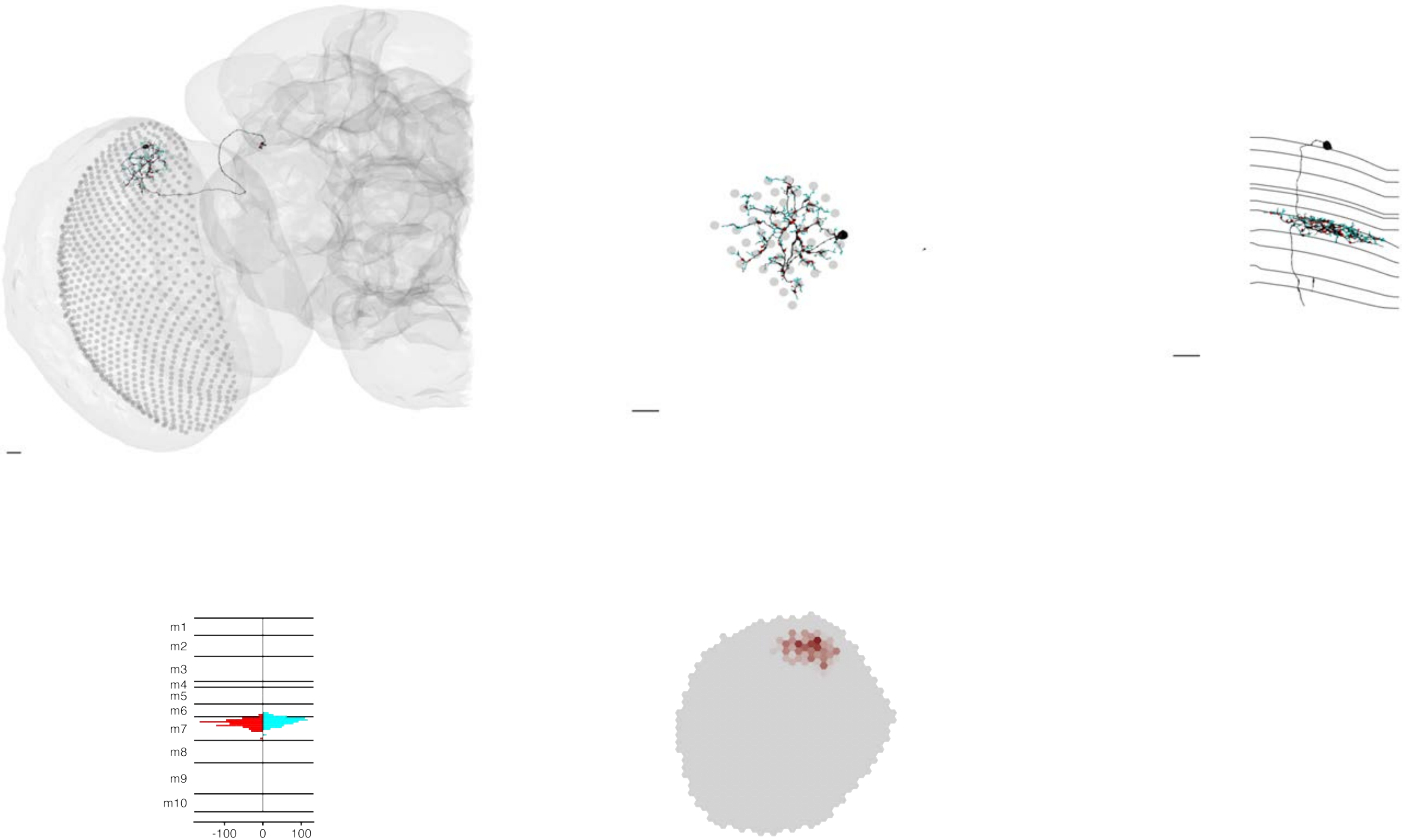

### MeTu1_720575940624739472

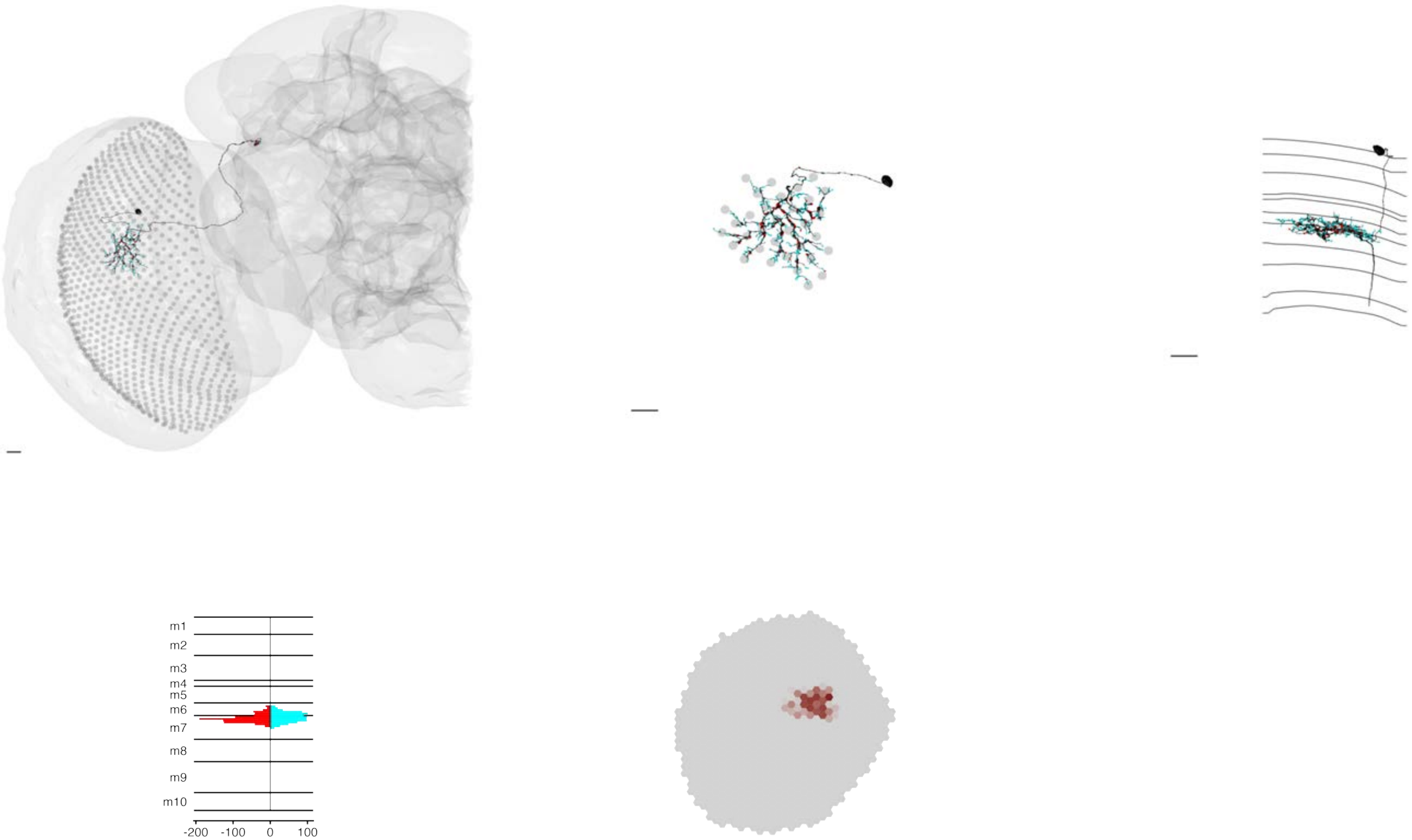

### MeTu1_720575940624742032

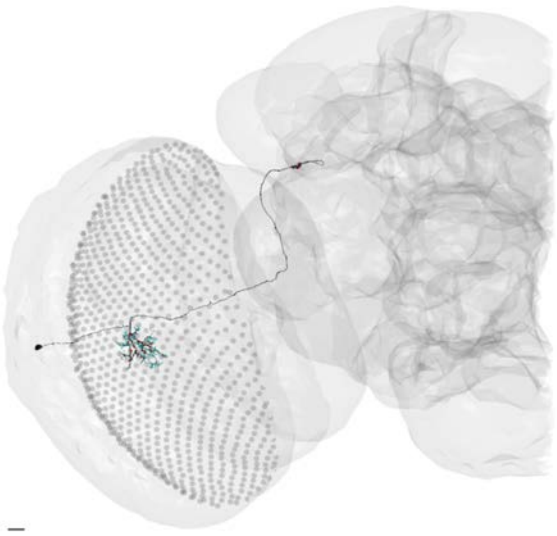

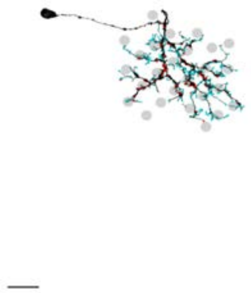

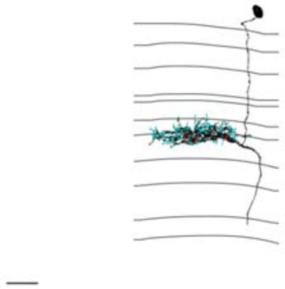

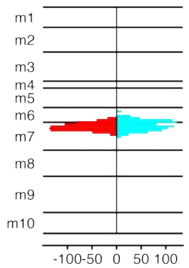


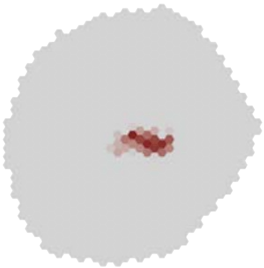

### MeTu1_720575940625228862

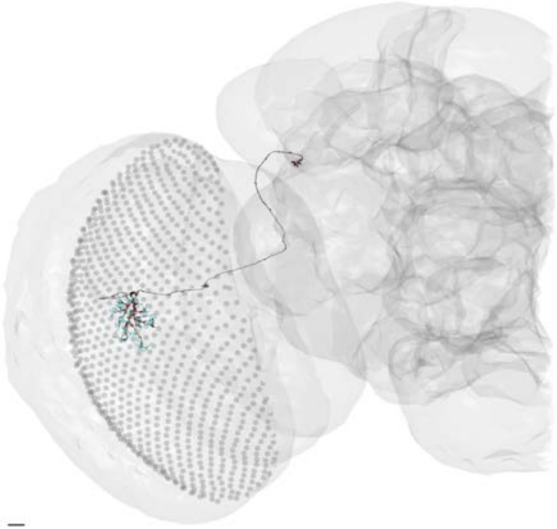

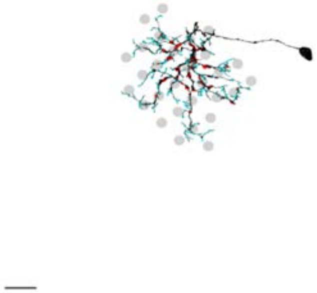

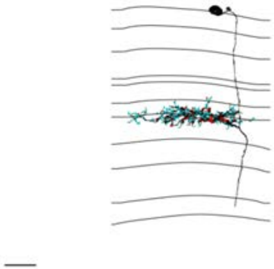

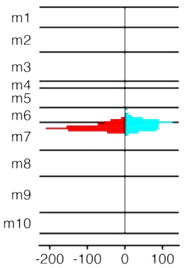


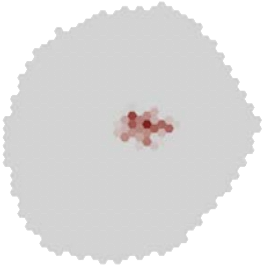

### MeTu1_720575940625597061

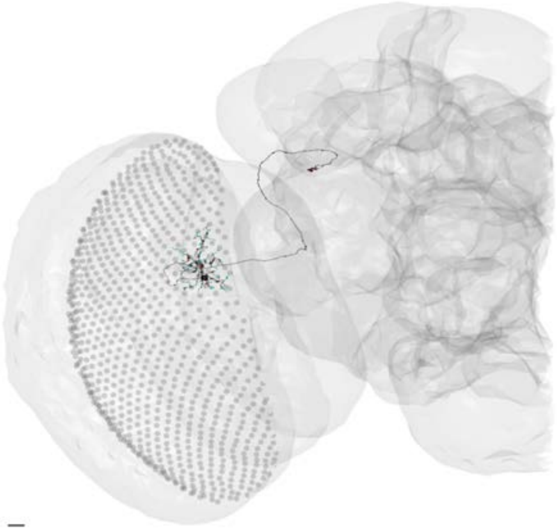

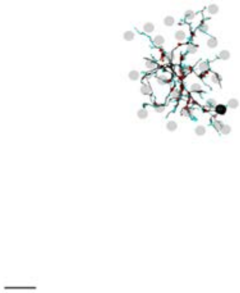

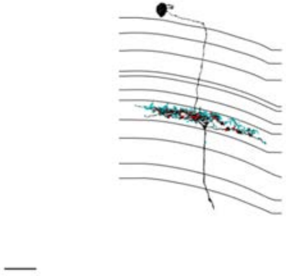

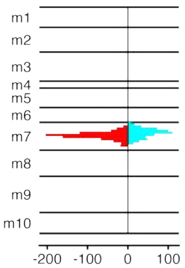


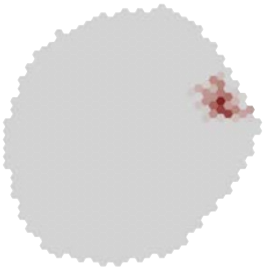

### MeTu1_720575940625656552

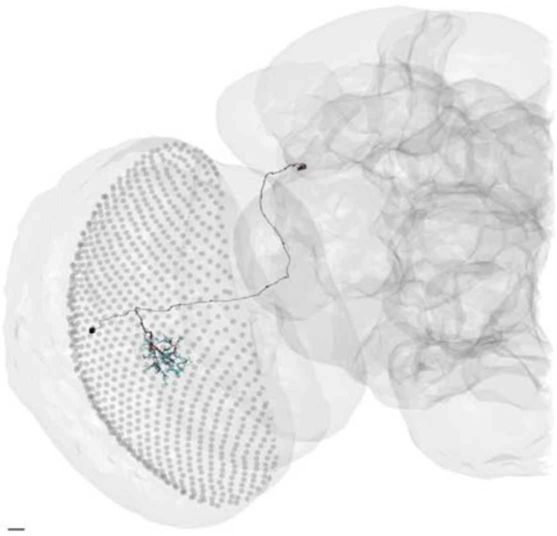

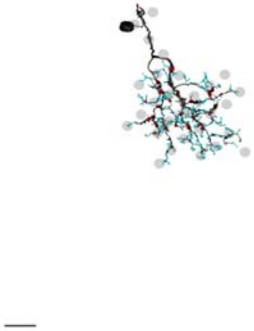

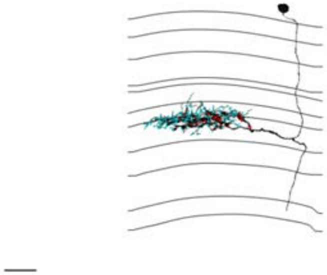

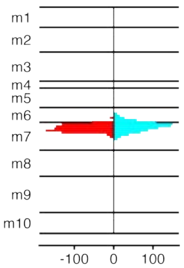


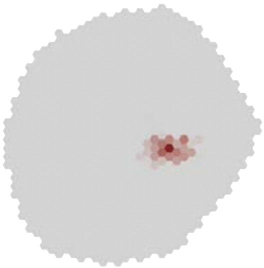

### MeTu1_720575940625704231

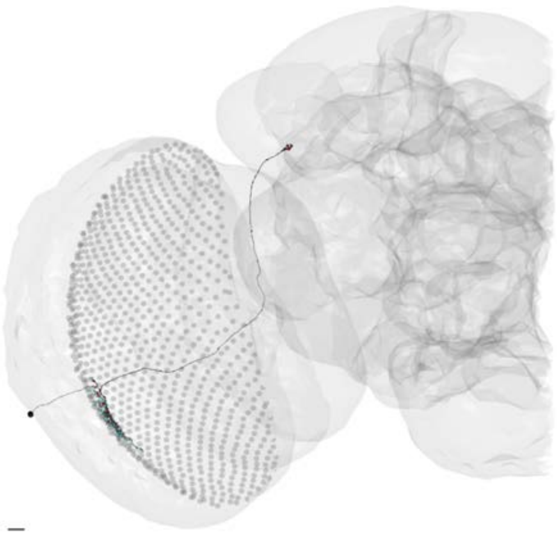

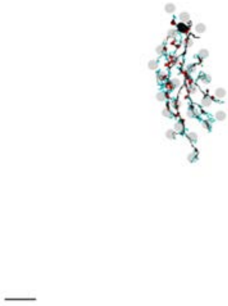

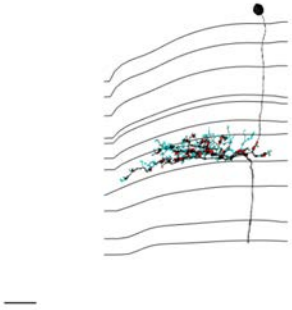

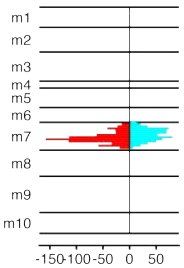


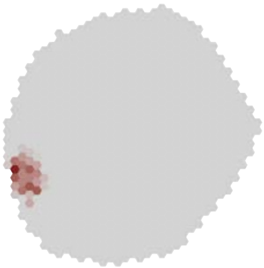

### MeTu1_720575940626285872

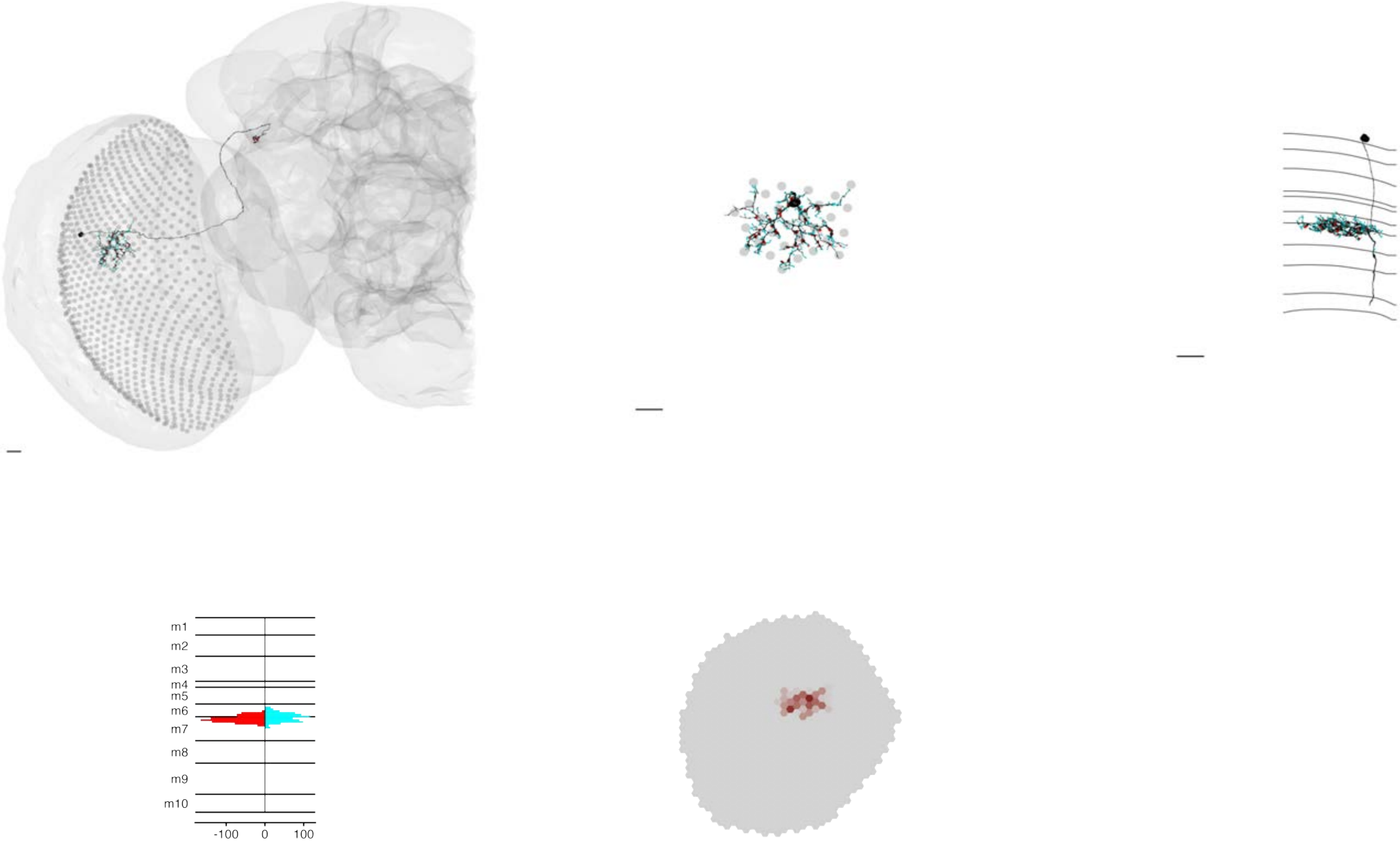

### MeTu1_720575940626330353

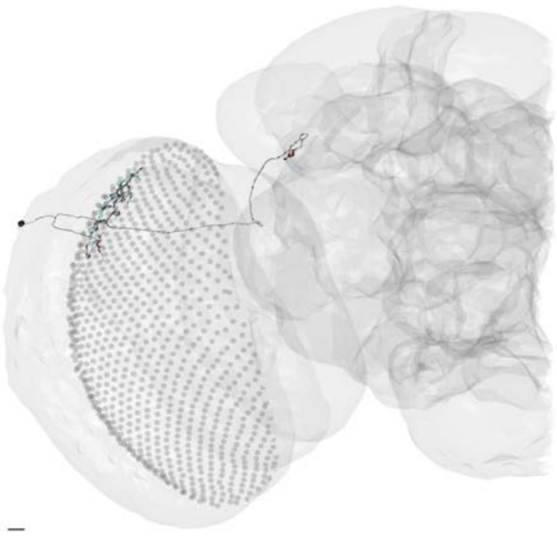

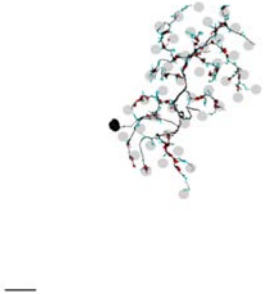

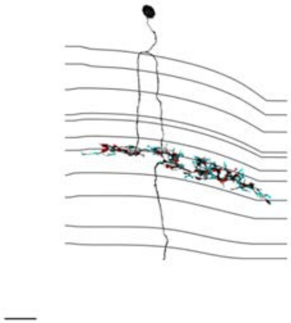

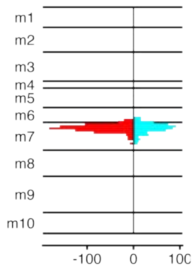


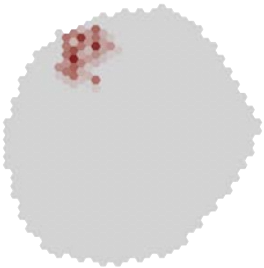

MeTu1_720575940626354057

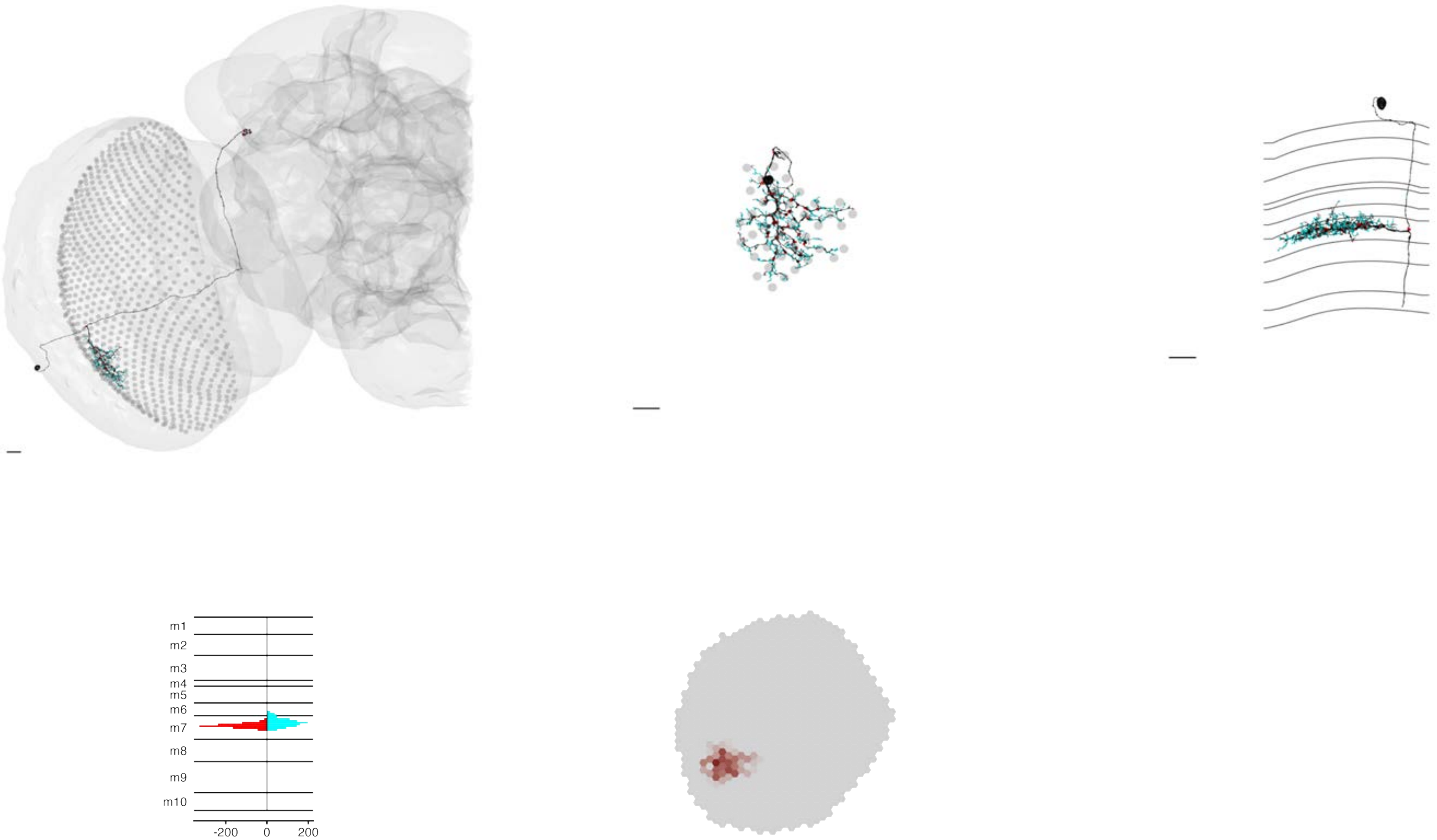

### MeTu1_720575940626509673

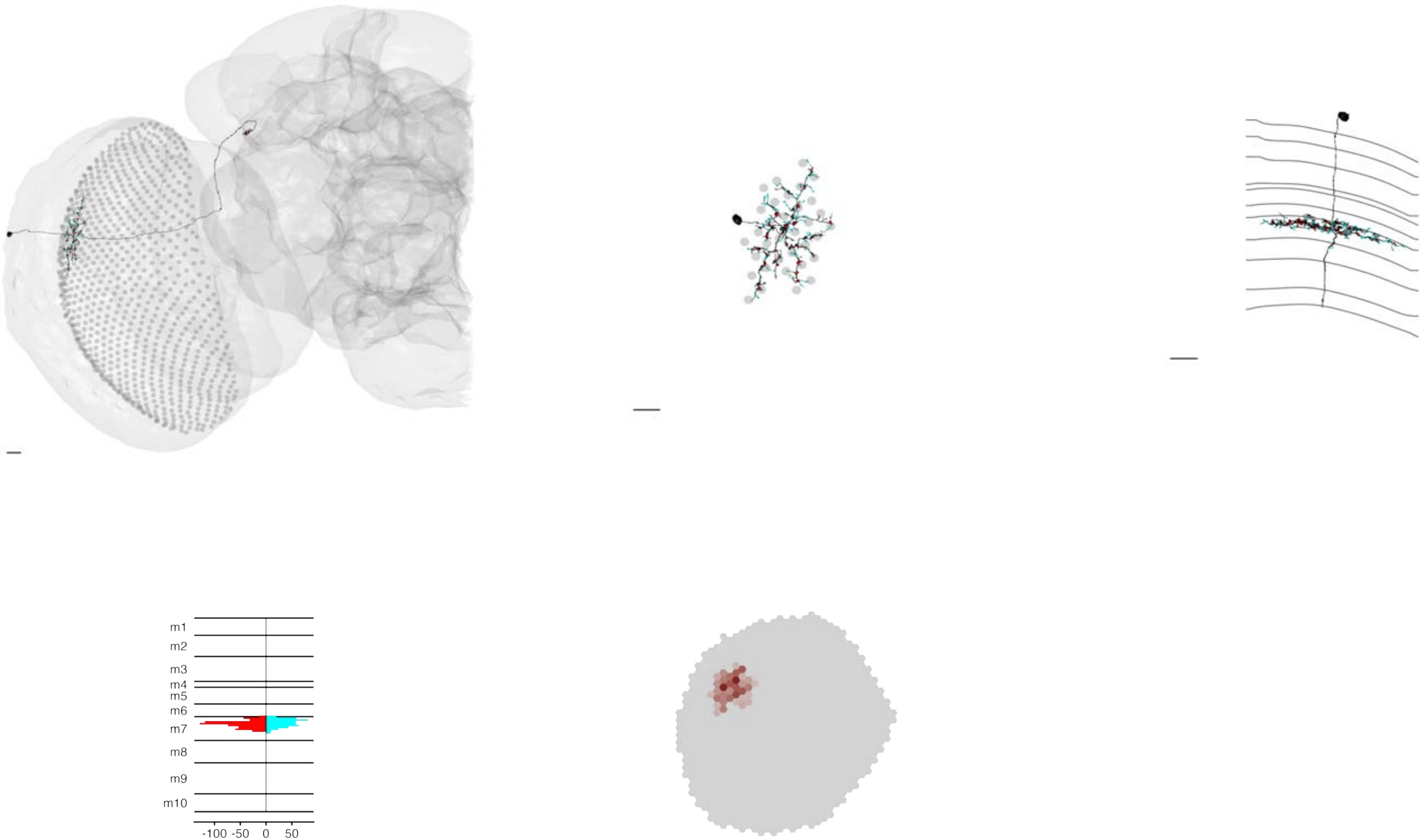

### MeTu1_720575940626557065

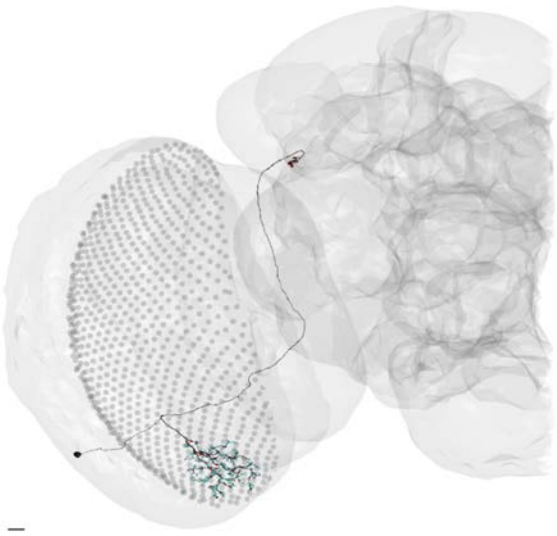

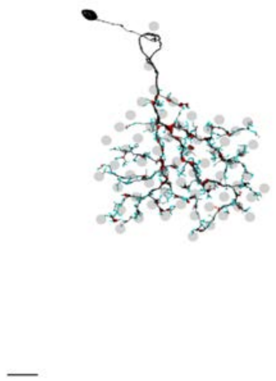

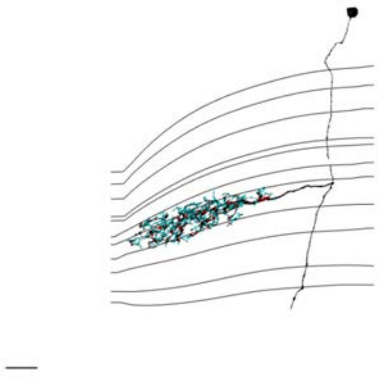

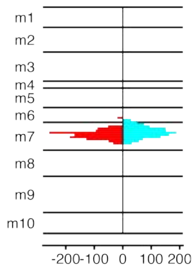


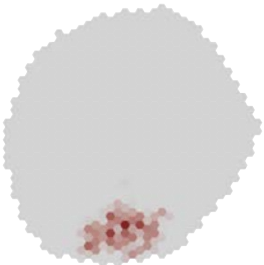

### MeTu1_720575940626596042

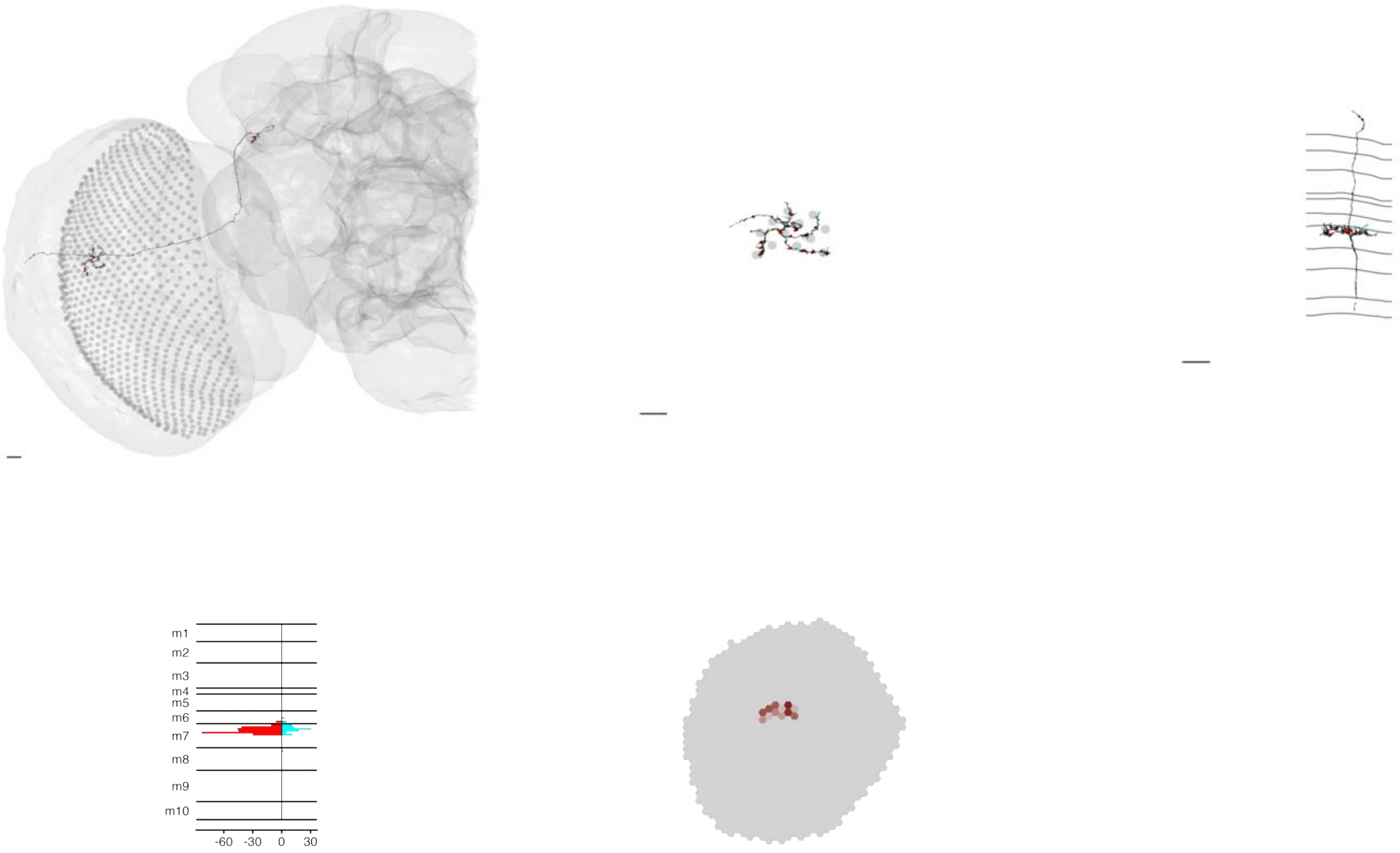

MeTu1_720575940626722448

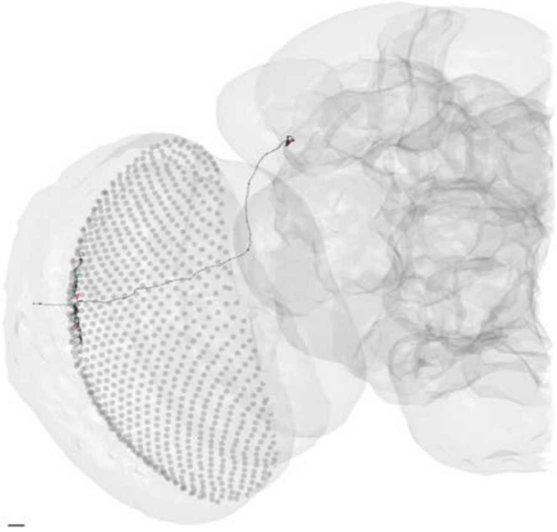

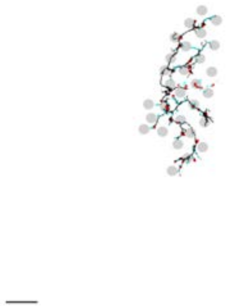

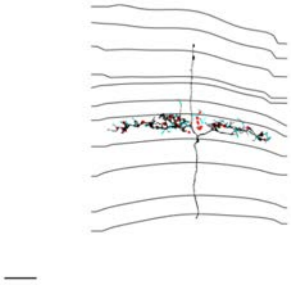

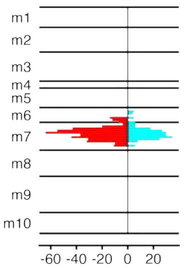


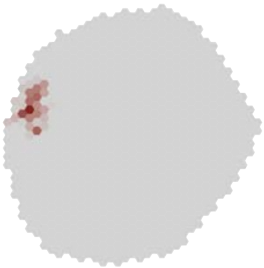

### MeTu1_720575940626784574

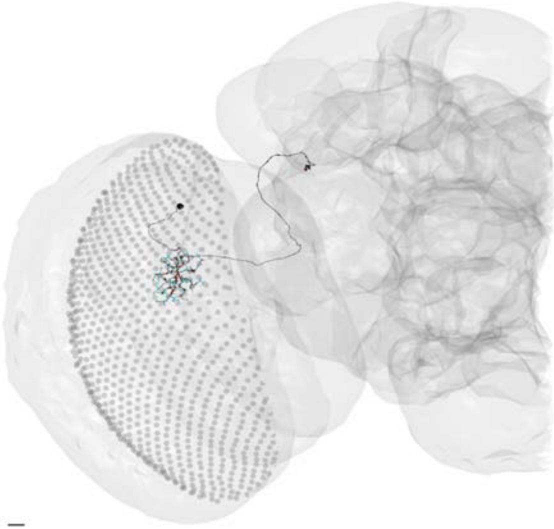

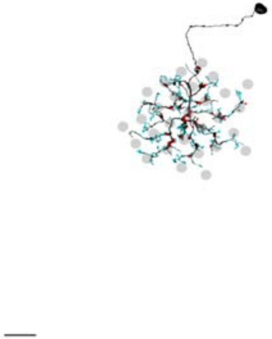

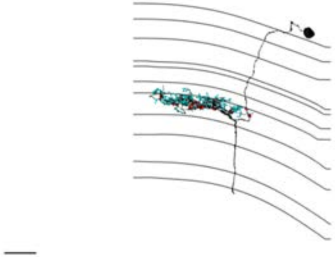

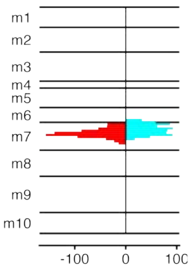


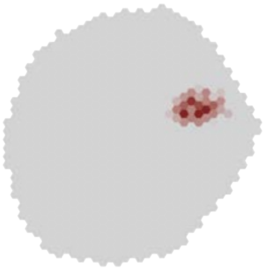

### MeTu1_720575940626878052

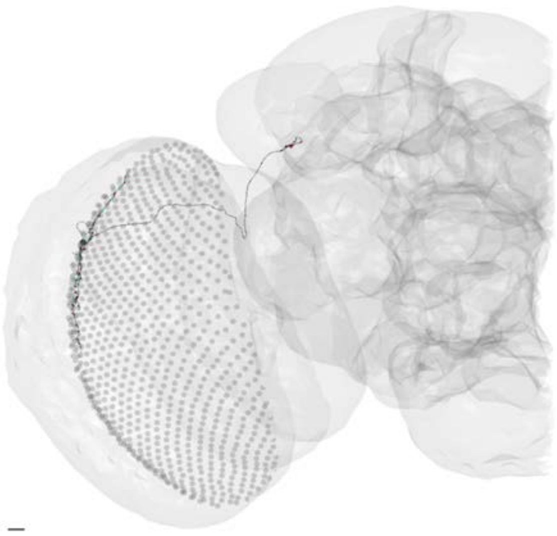

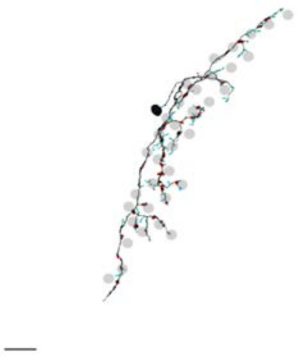

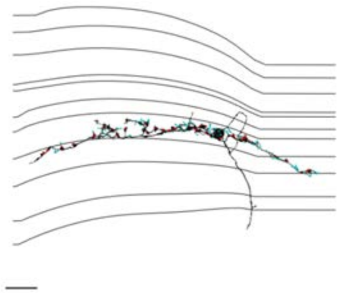

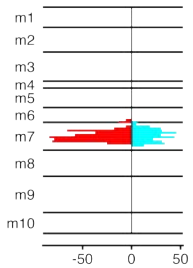


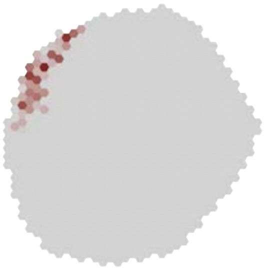

MeTu1_720575940627843594

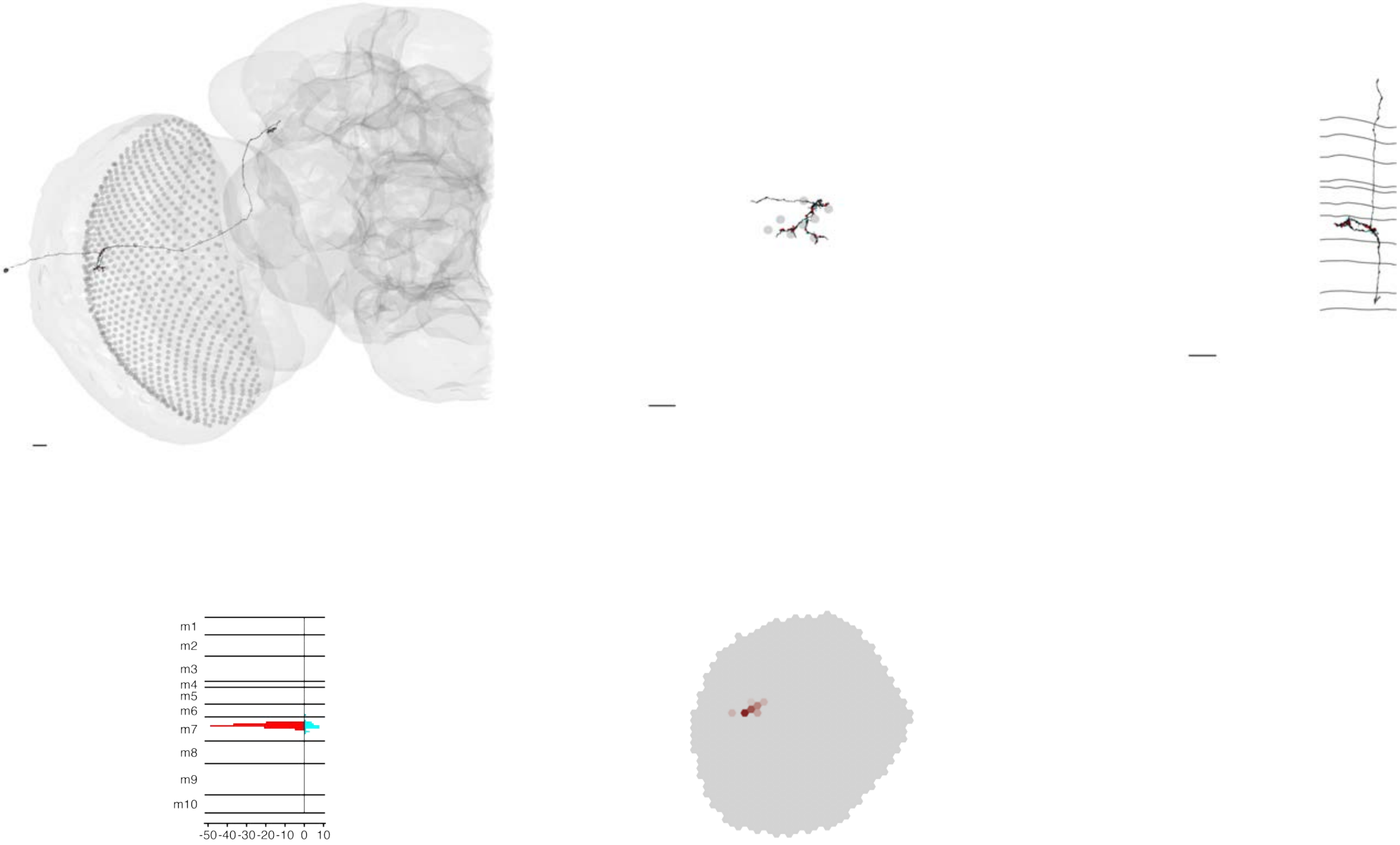

### MeTu1_720575940627845904

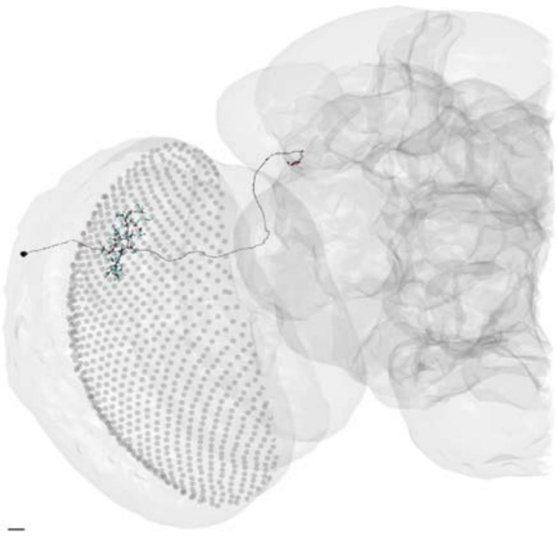

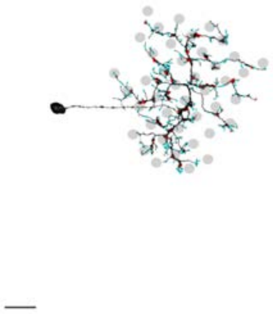

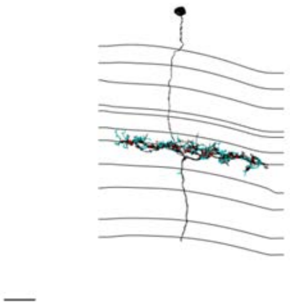

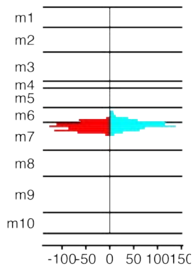


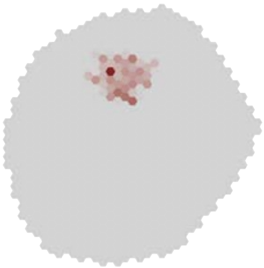

### MeTu1_720575940628221227

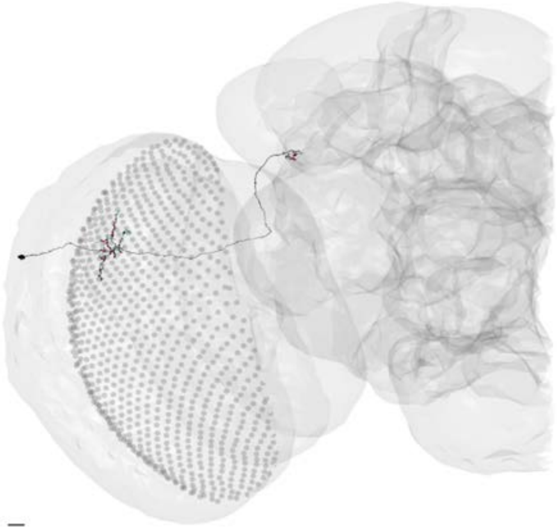

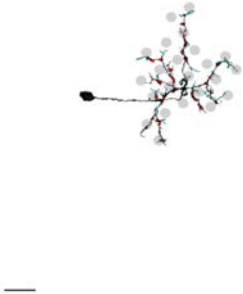

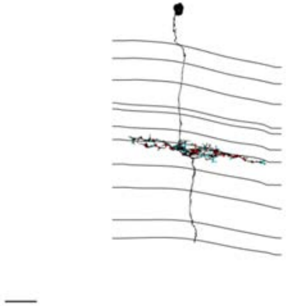

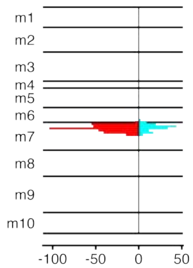


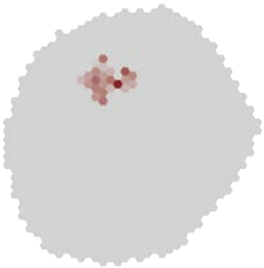

### MeTu1_720575940628319656

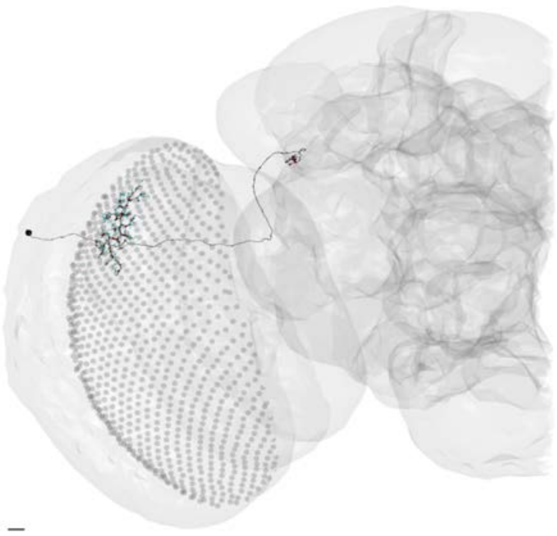

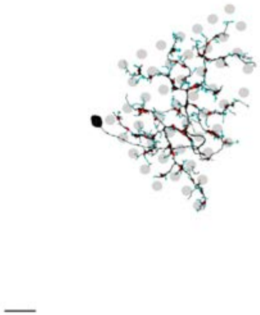

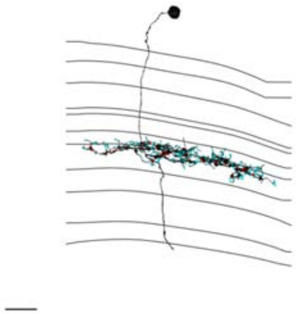

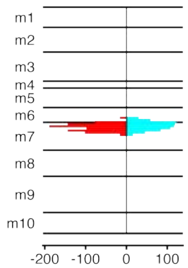


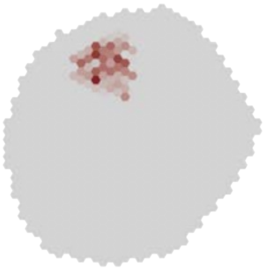

### MeTu1_720575940628403272

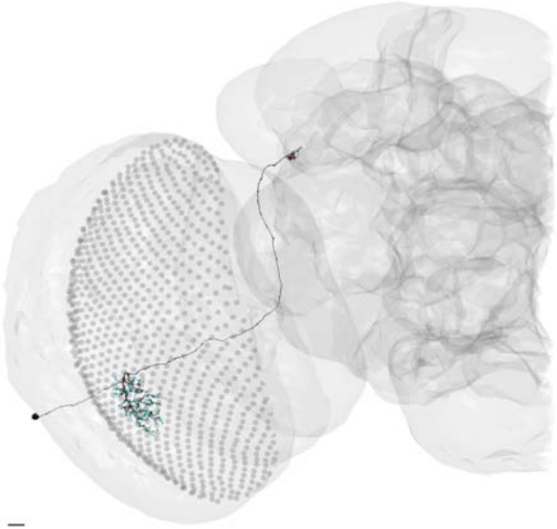

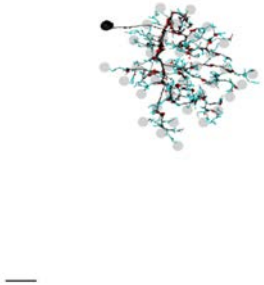

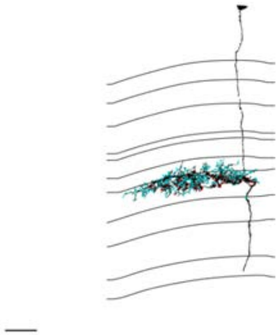

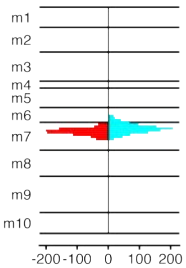


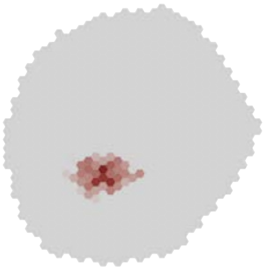

### MeTu1_720575940628594135

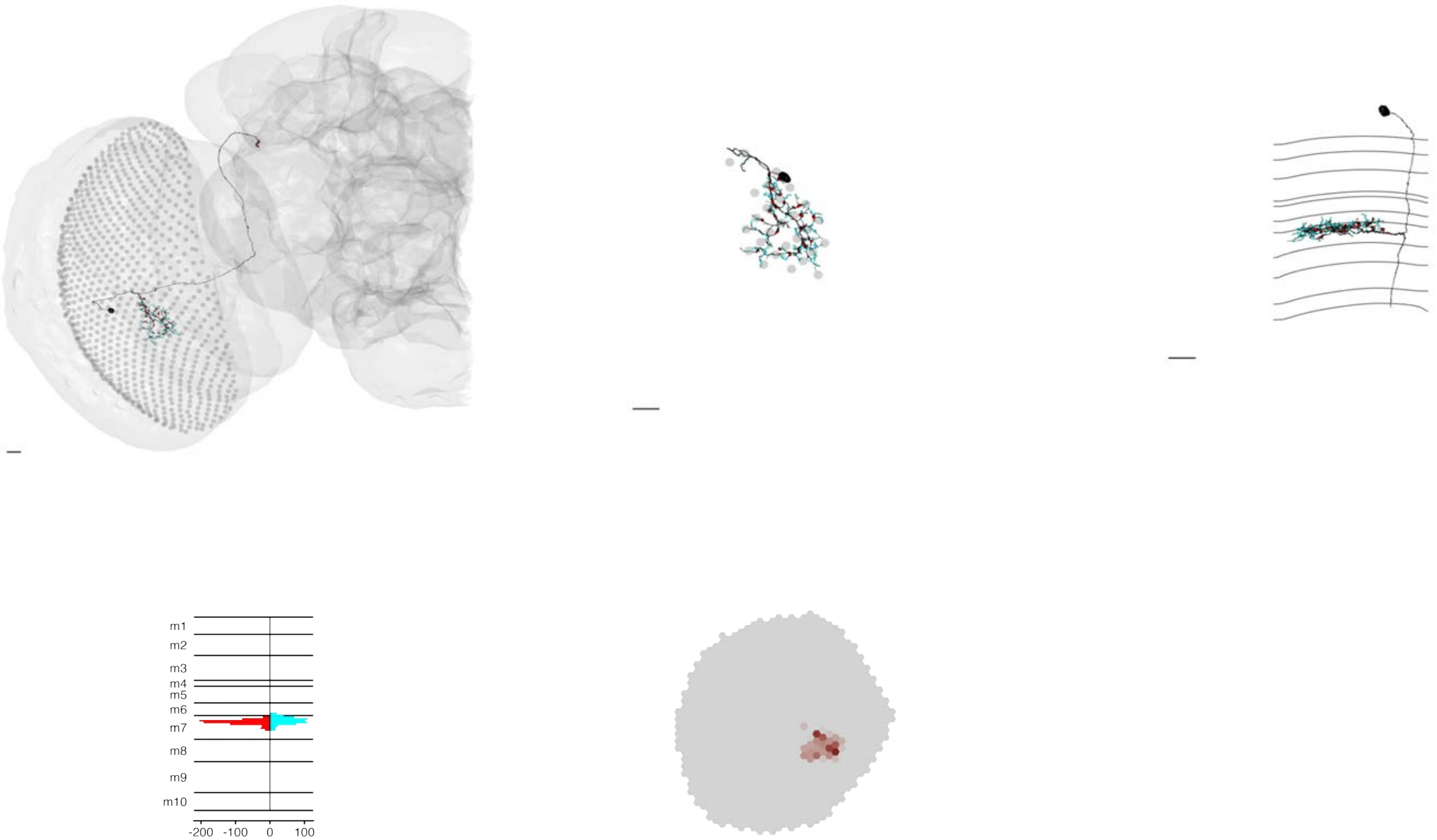

### MeTu1_720575940628623432

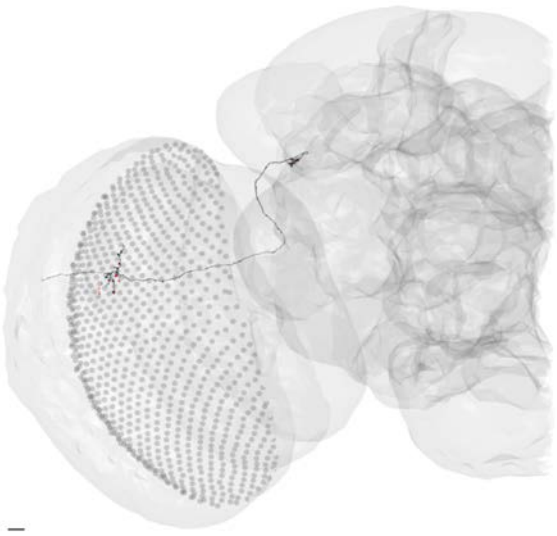

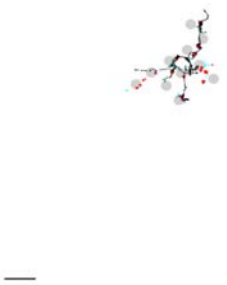

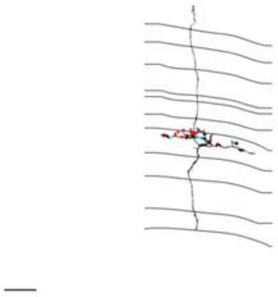

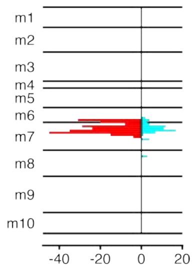


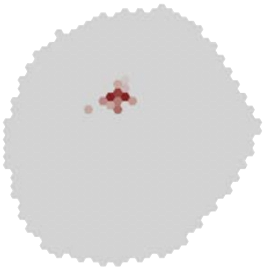

MeTu1_720575940629260855

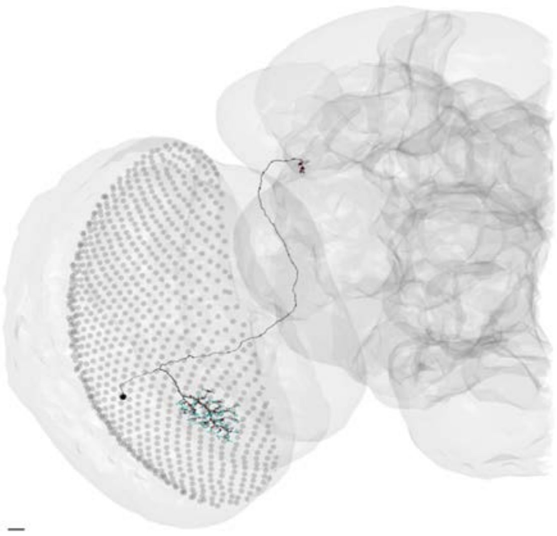

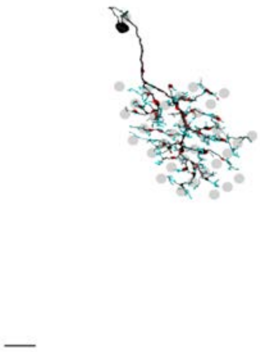

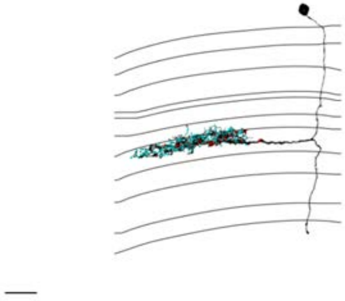

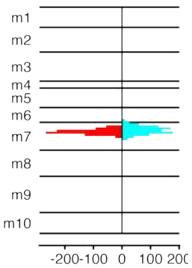


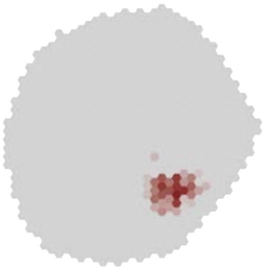

### MeTu1_720575940629421879

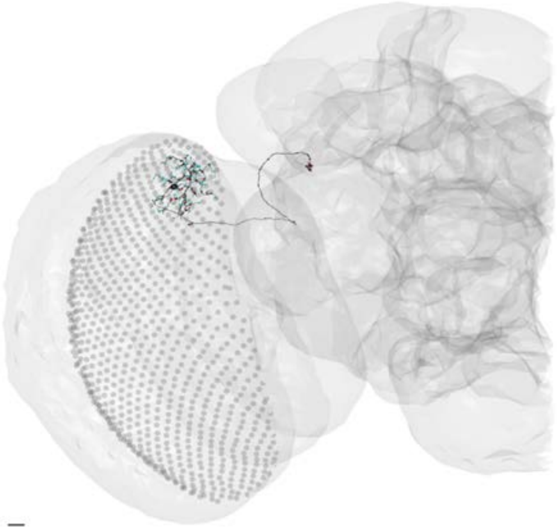

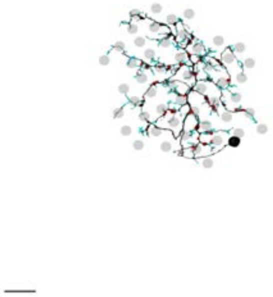

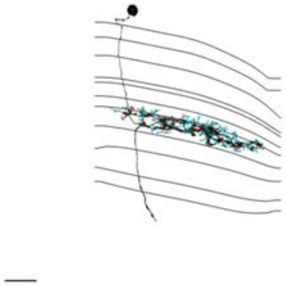

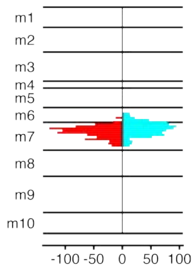


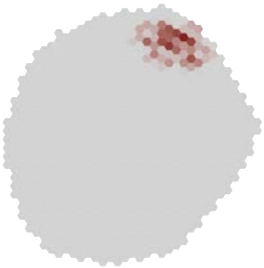

### MeTu1_720575940629443543

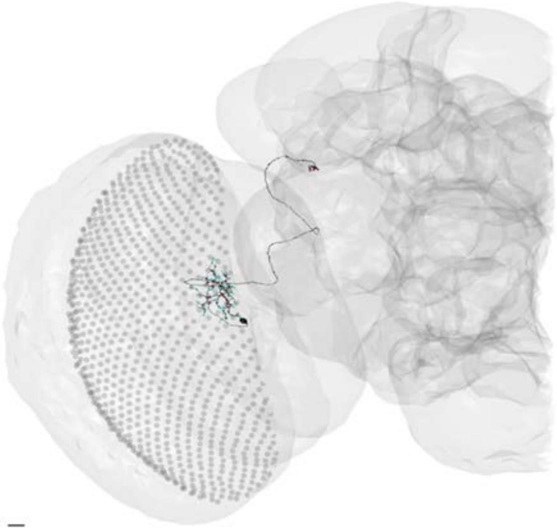

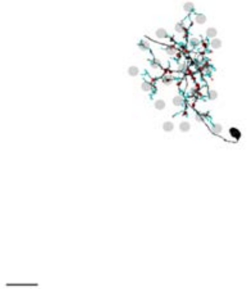

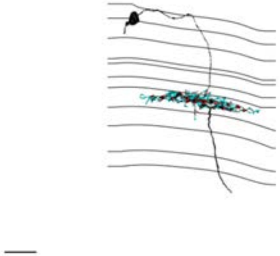

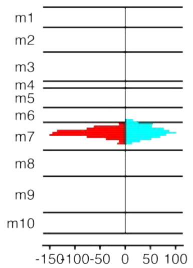


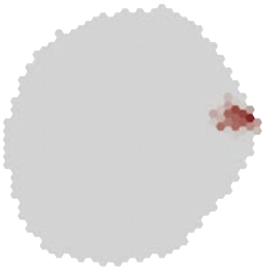

### MeTu1_720575940629765418

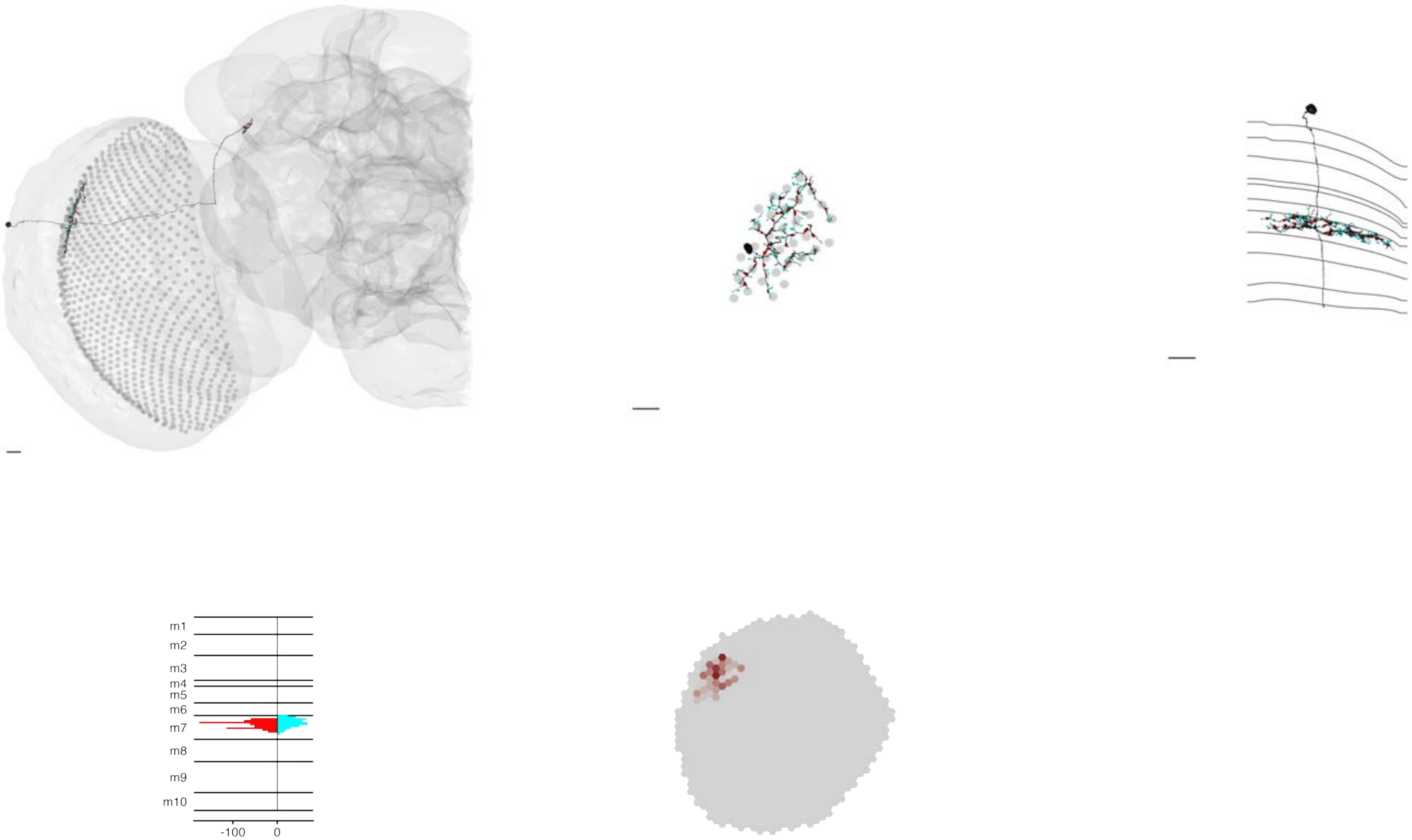

### MeTu1_720575940630006528

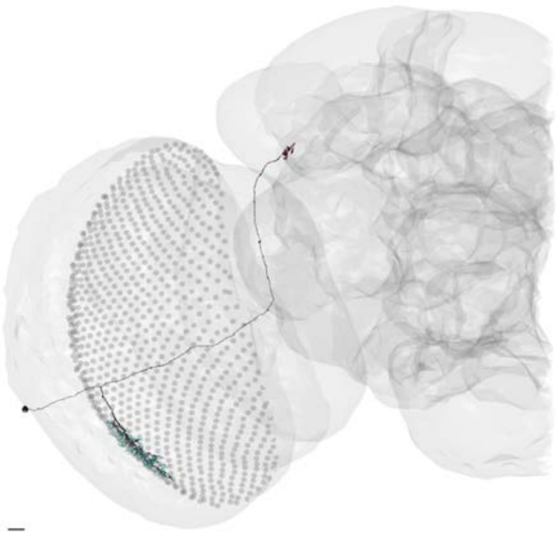

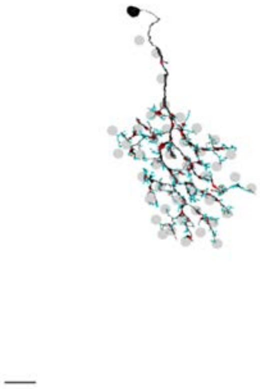

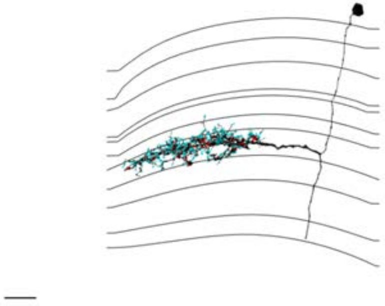

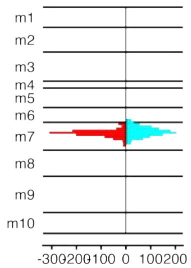


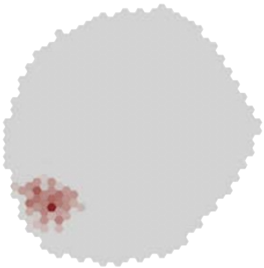

### MeTu1_720575940630562754

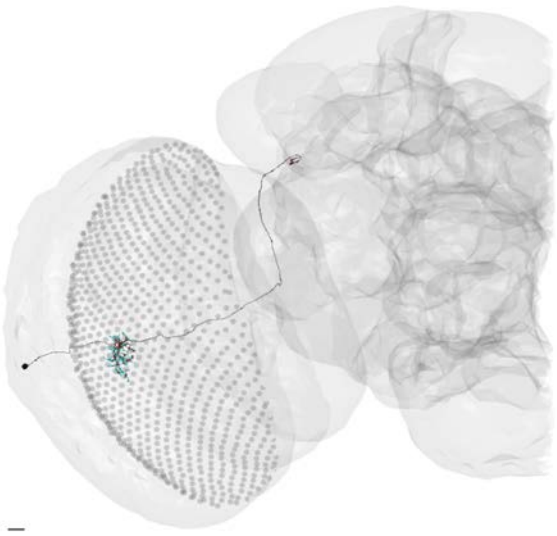

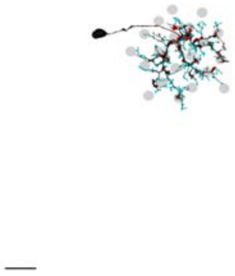

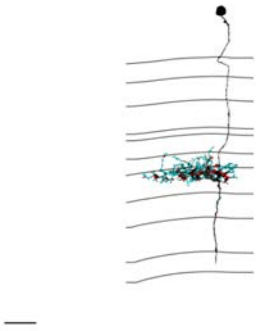

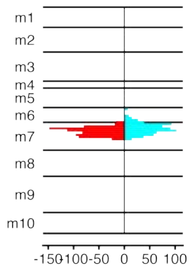


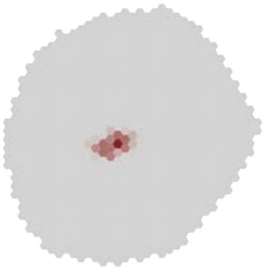

### MeTu1_720575940631066875

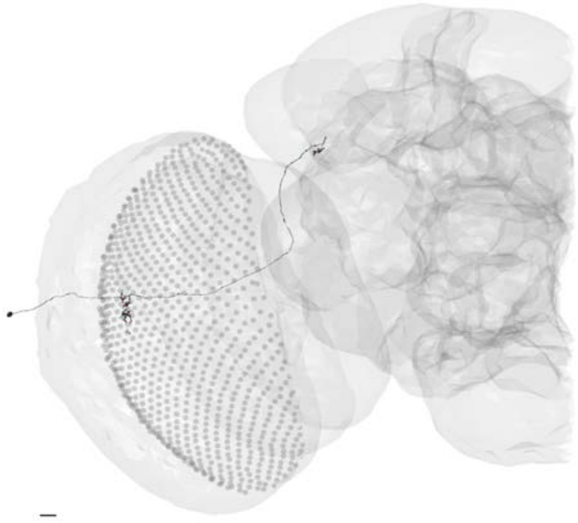

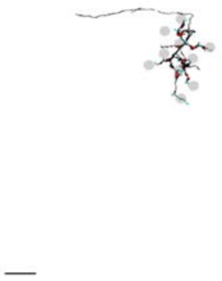

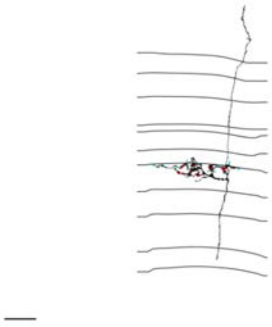

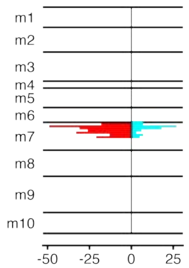


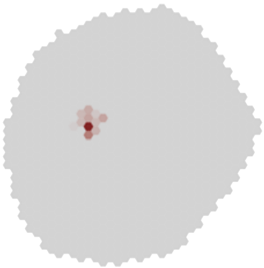

### MeTu1_720575940631102428

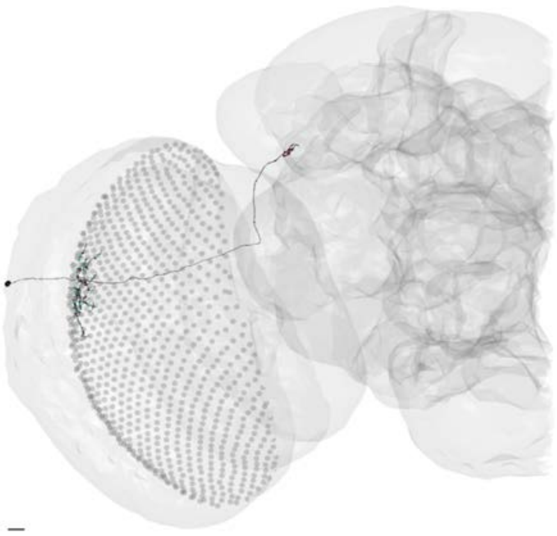

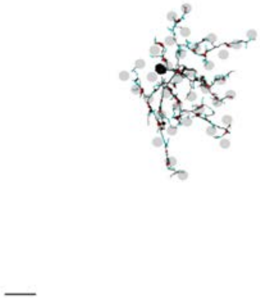

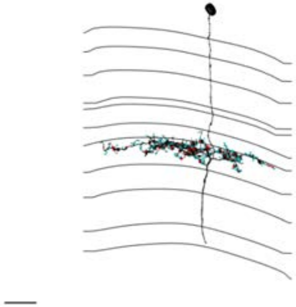

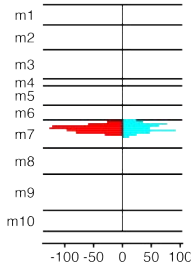


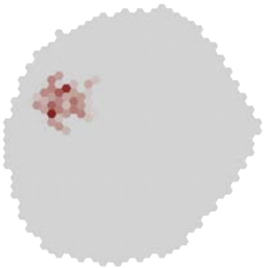

MeTu1_720575940631324077

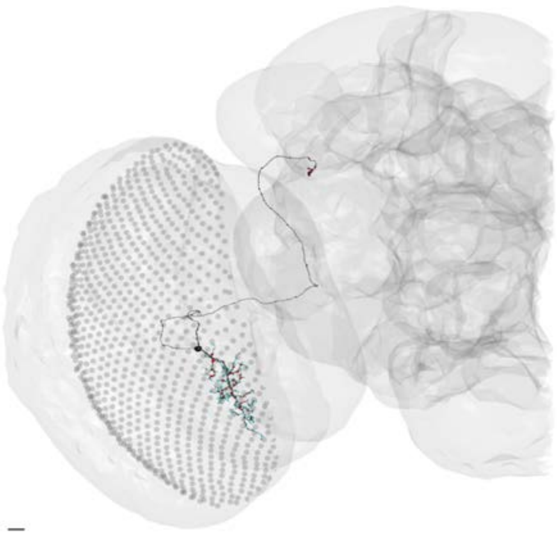

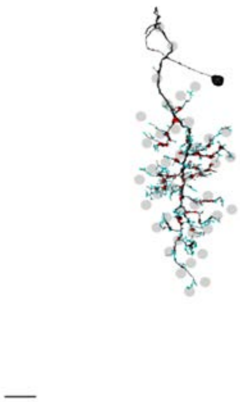

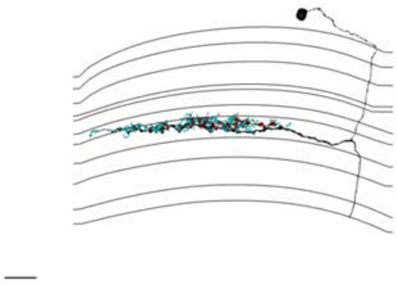

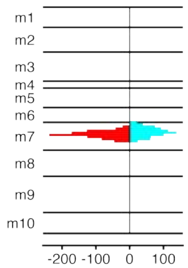


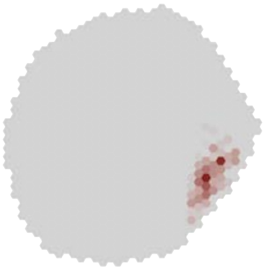

### MeTu1_720575940631694050

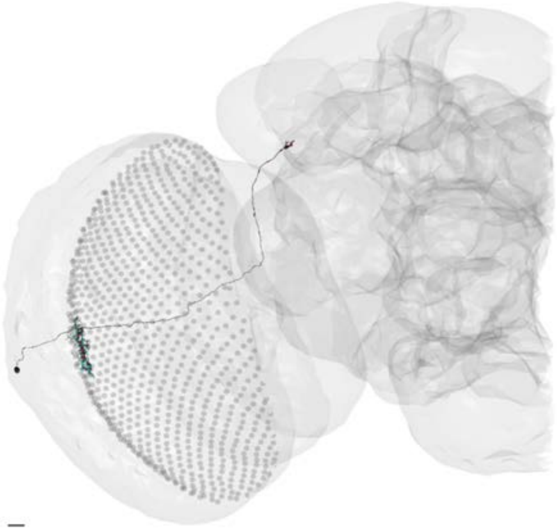

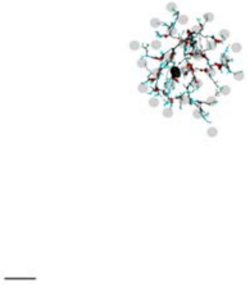

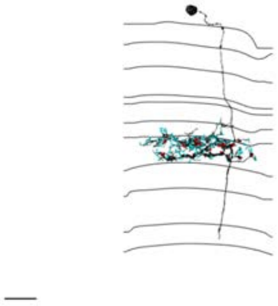

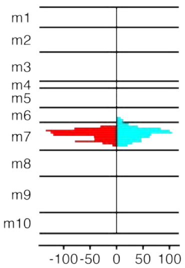


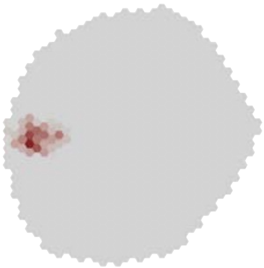

### MeTu1_720575940632772704

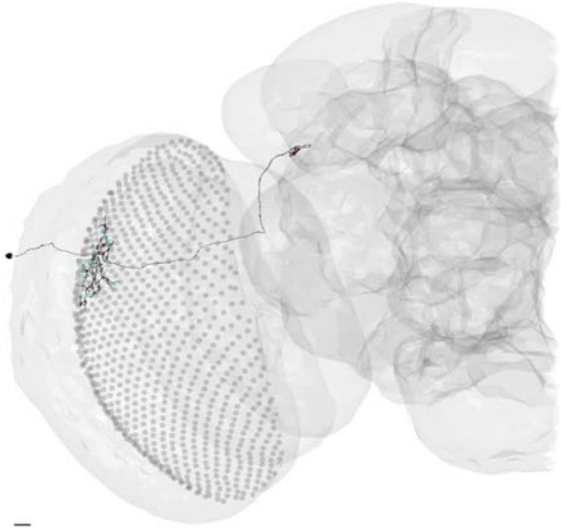

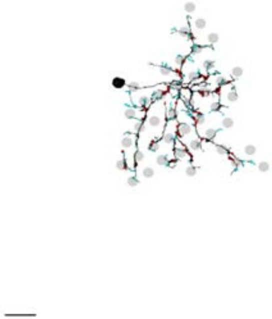

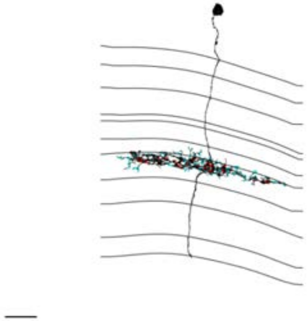

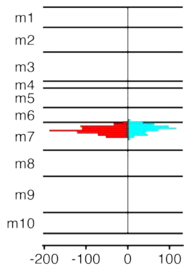


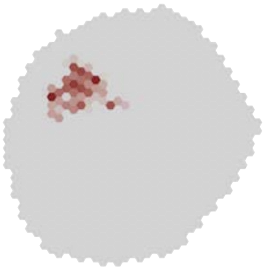

### MeTu1_720575940633046719

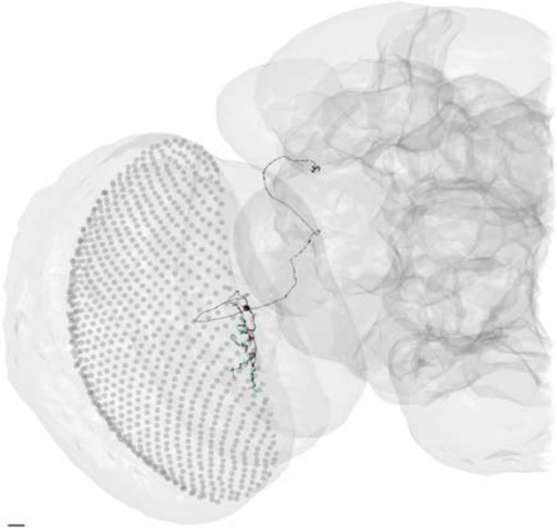

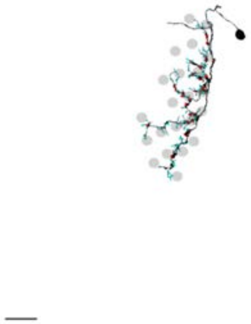

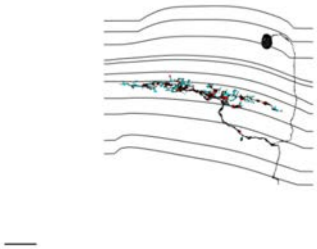

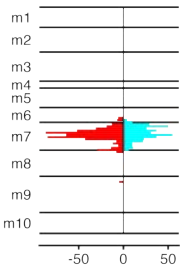


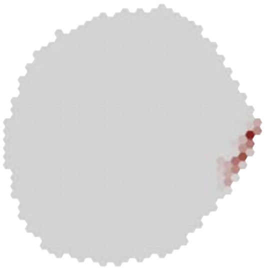

### MeTu1_720575940633185972

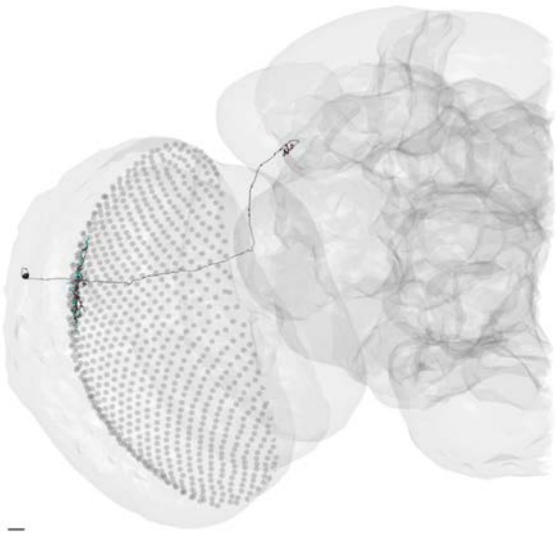

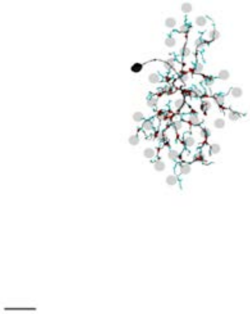

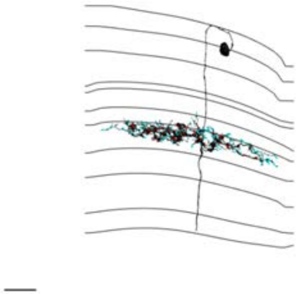

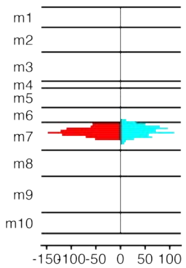


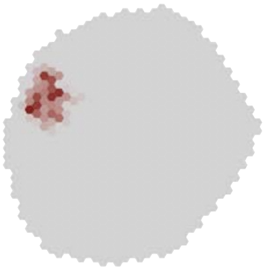

### MeTu1_720575940633207777

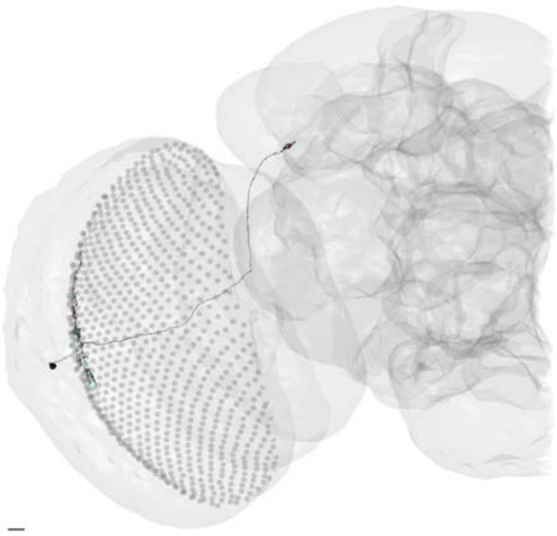

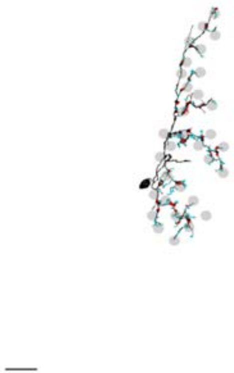

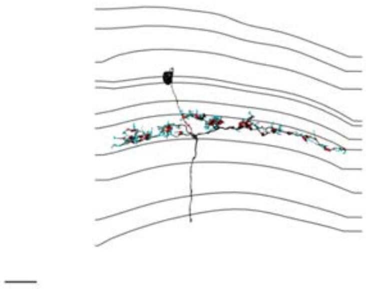

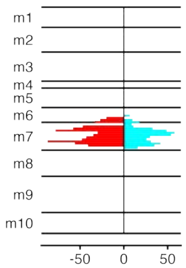


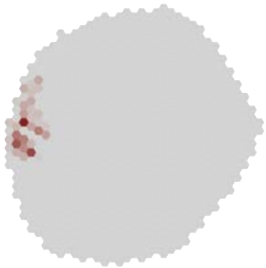

### MeTu1_720575940633453675

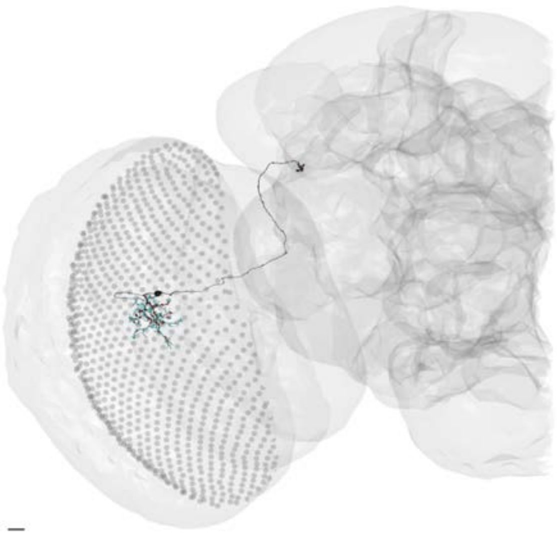

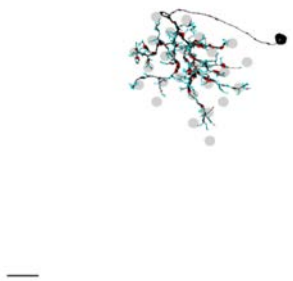

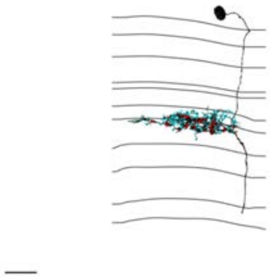

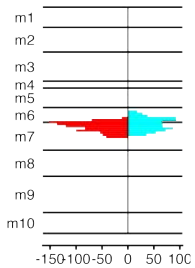


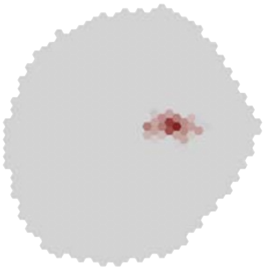

### MeTu1_720575940634124430

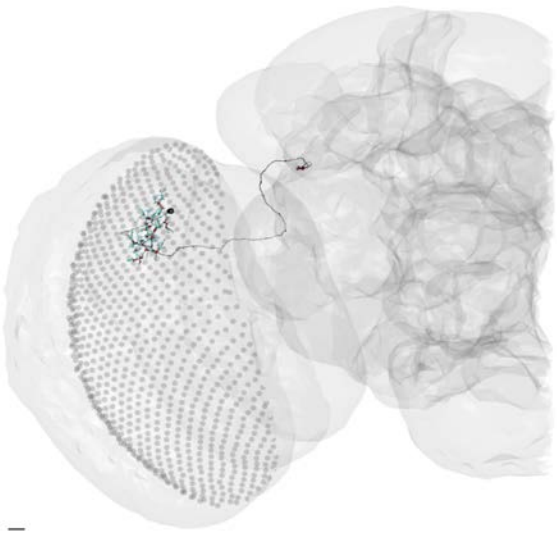

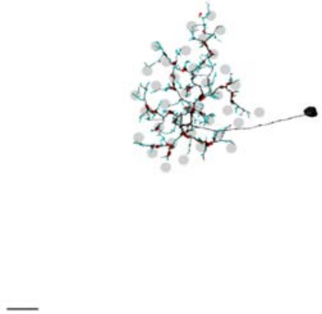

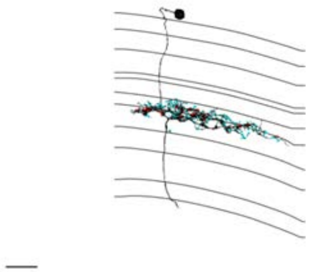

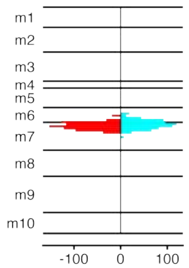


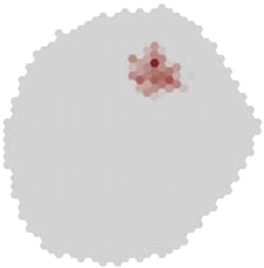

### MeTu1_720575940634745780

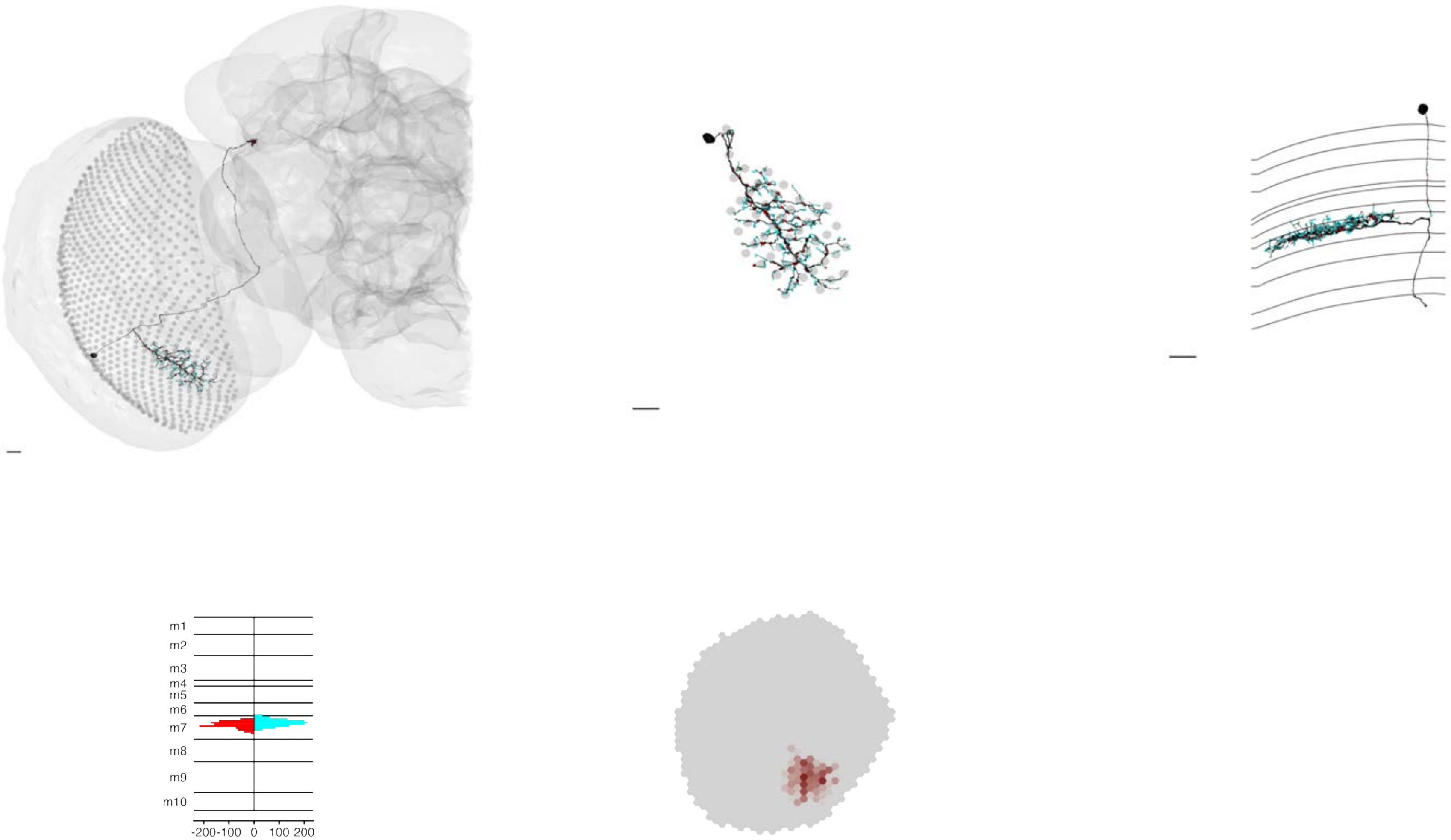

### MeTu1_720575940637215088

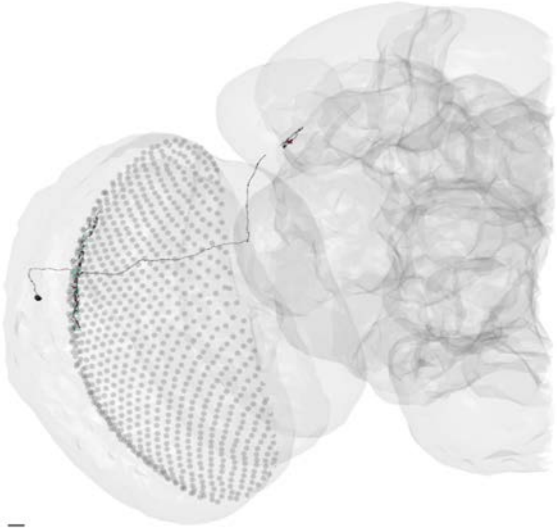

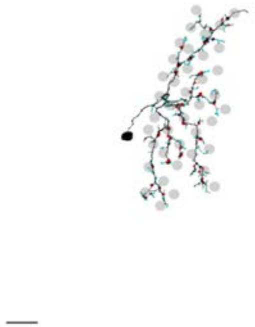

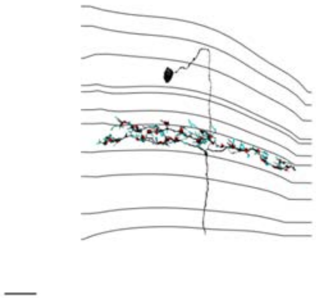

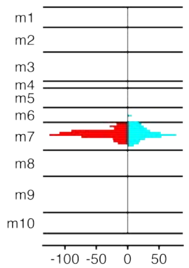


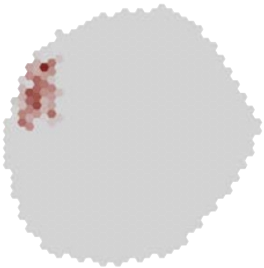

### MeTu1_720575940637535677

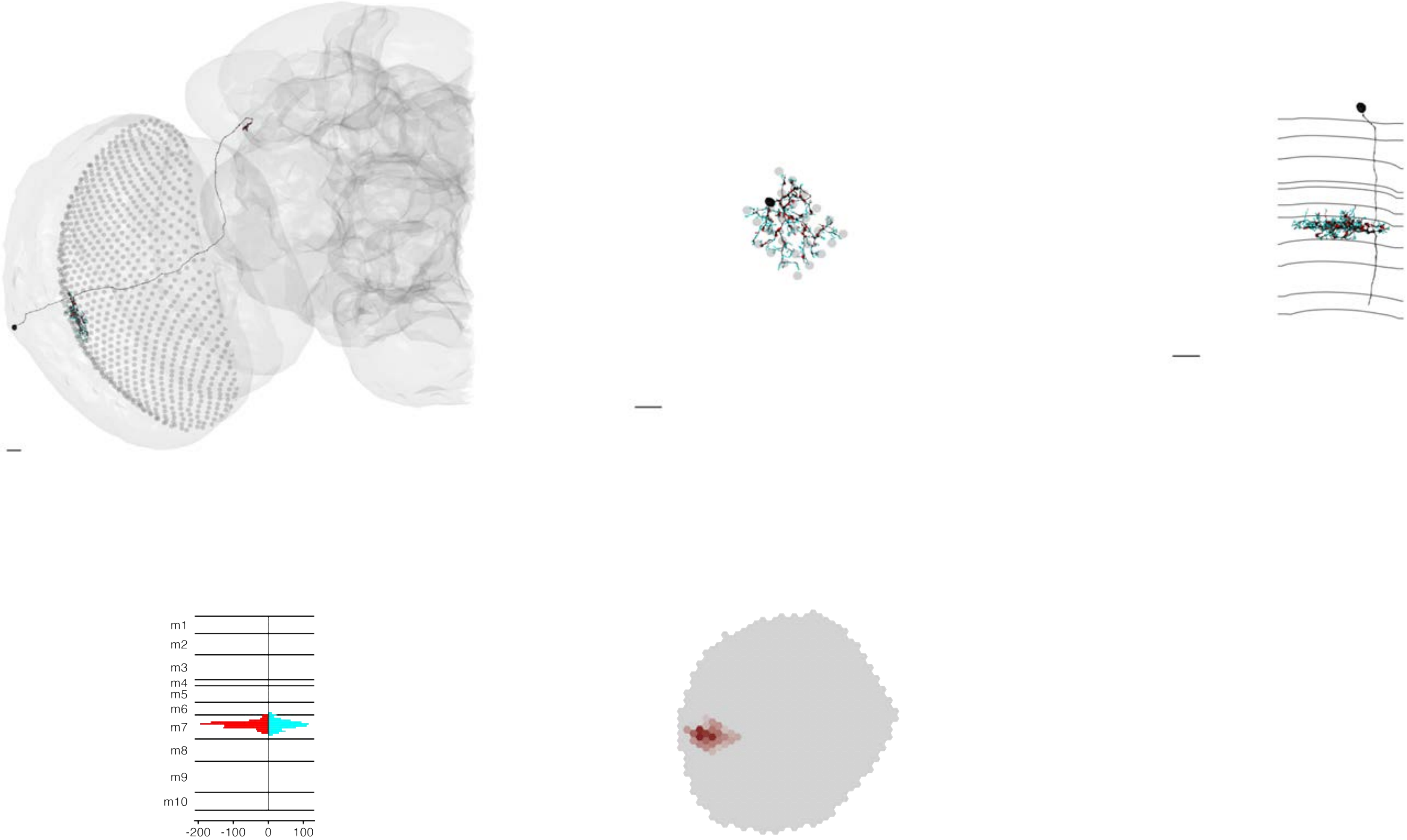

### MeTu1_720575940638033752

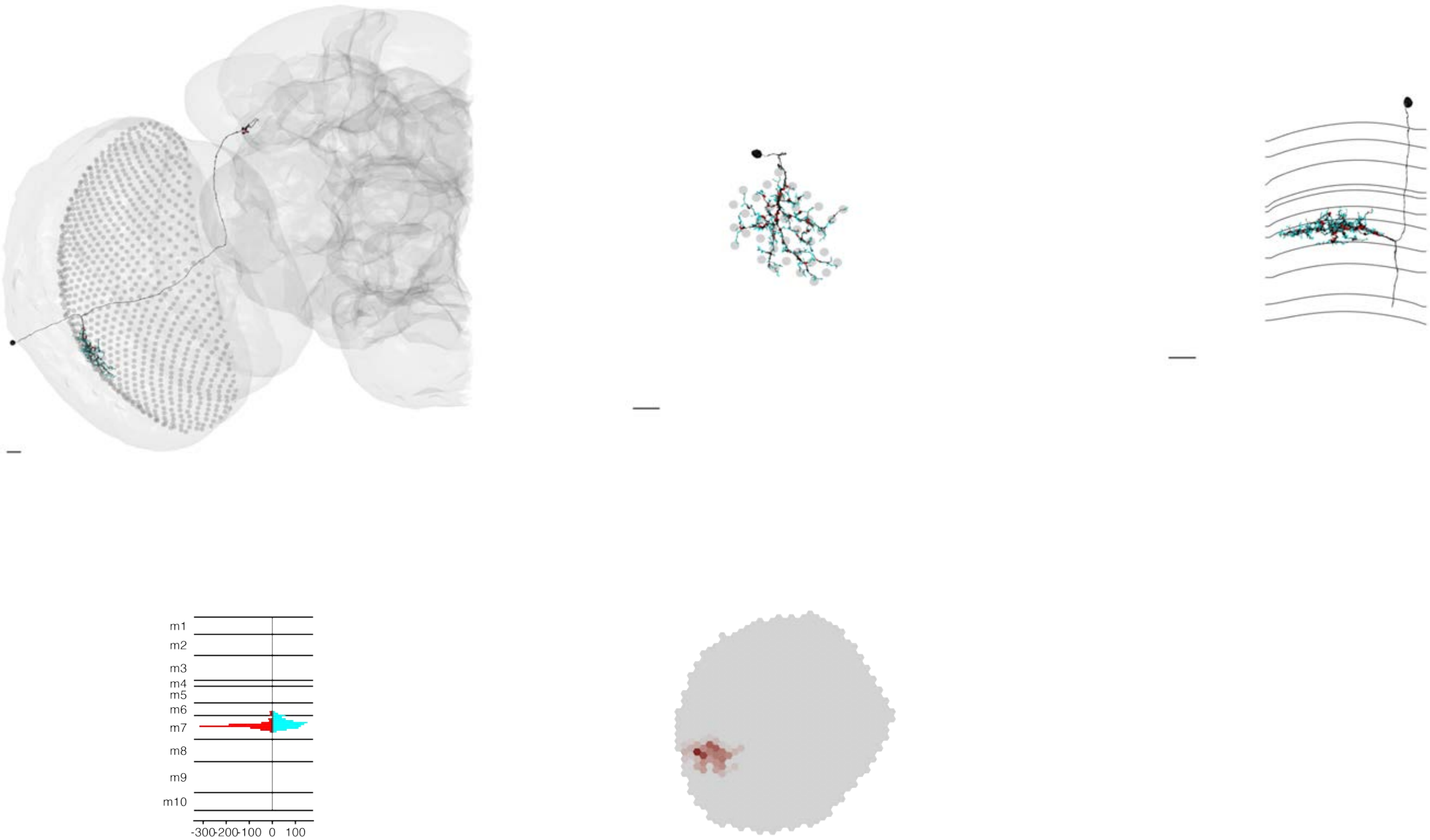

### MeTu1_720575940638131544

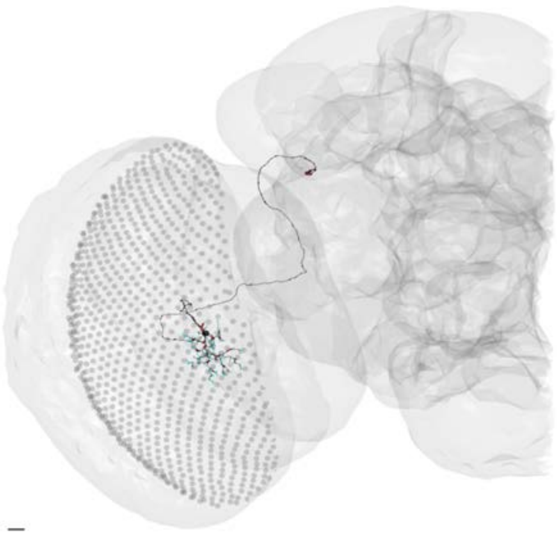

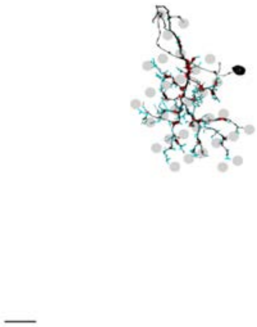

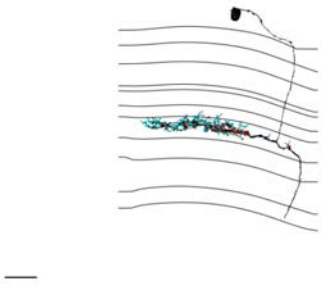

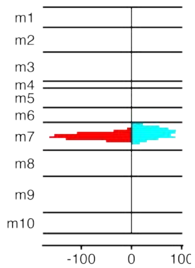


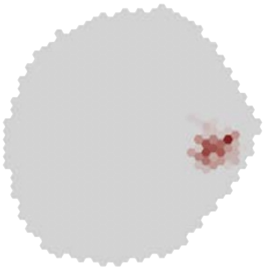

### MeTu1_720575940639590901

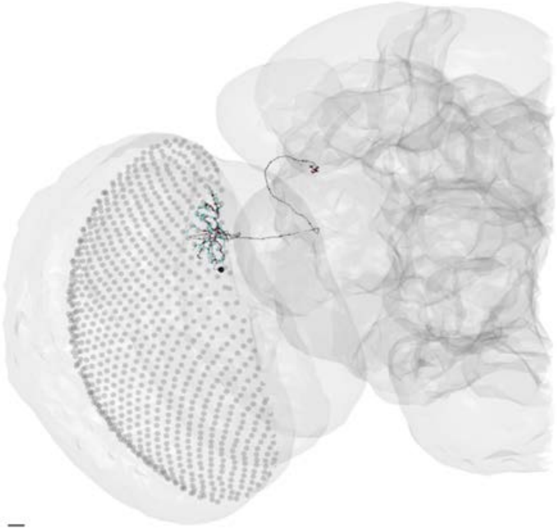

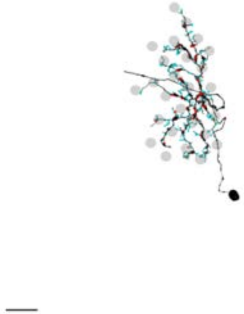

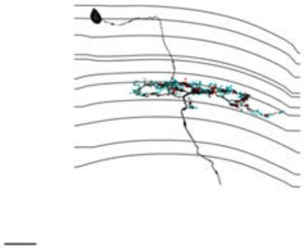

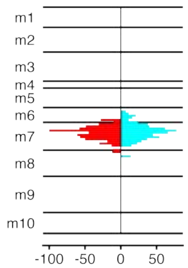


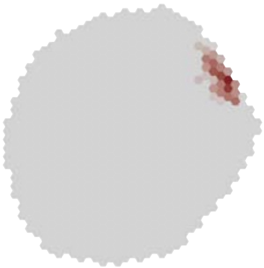

### MeTu1_720575940639611893

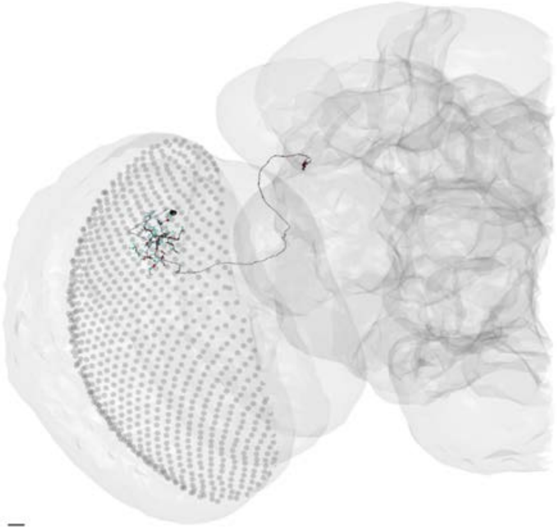

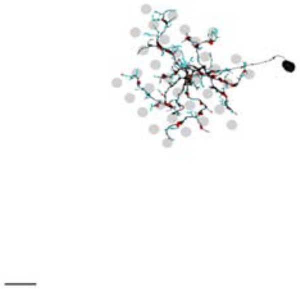

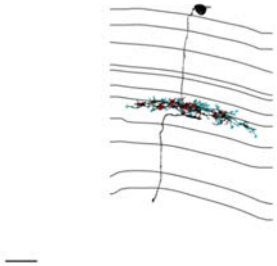

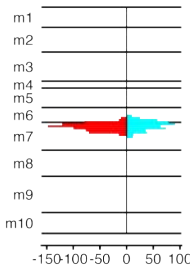


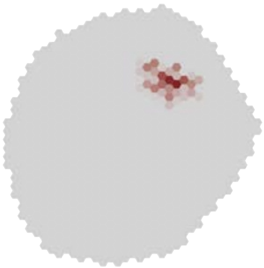

### MeTu1_720575940639670717

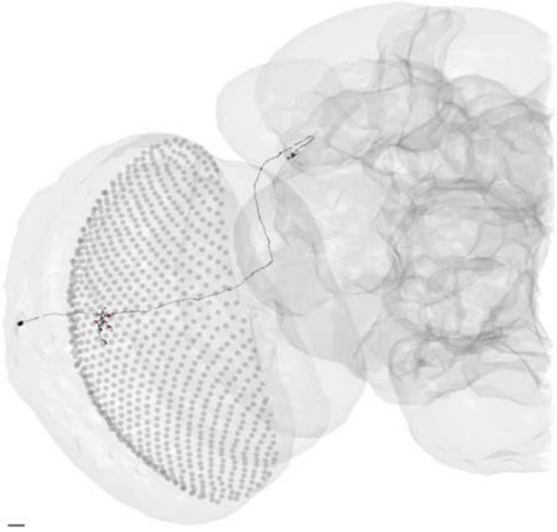

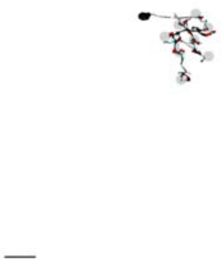

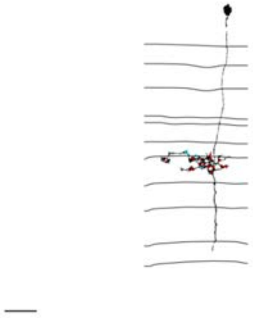

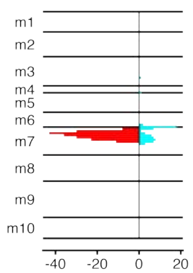


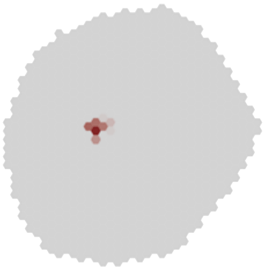

### MeTu1_720575940639708368

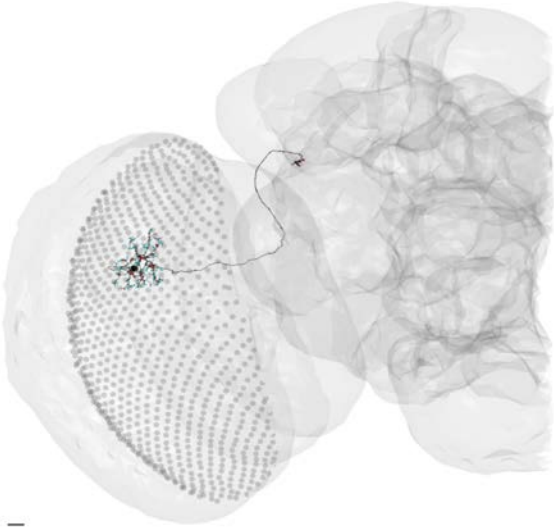

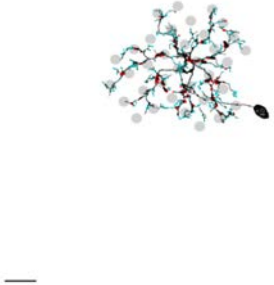

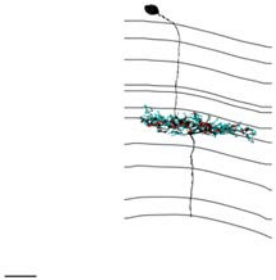

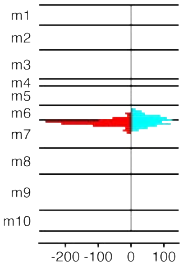


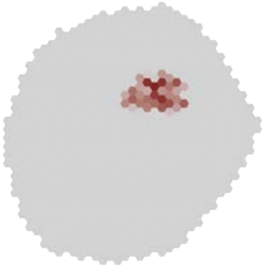

### MeTu1_720575940639872821

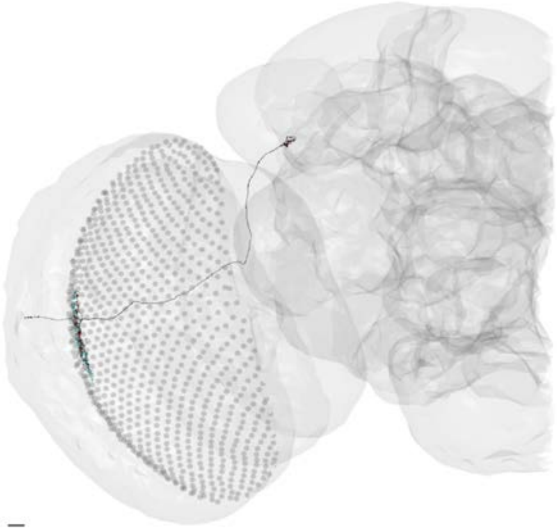

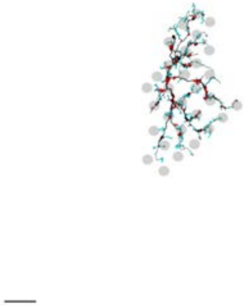

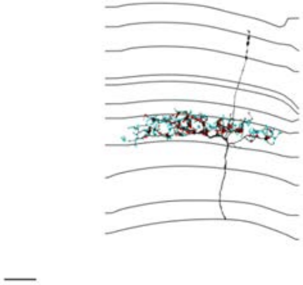

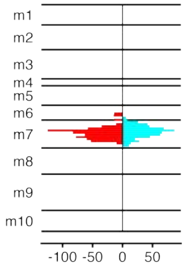


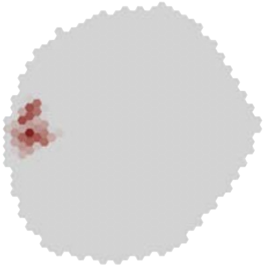

MeTu1_720575940641099149

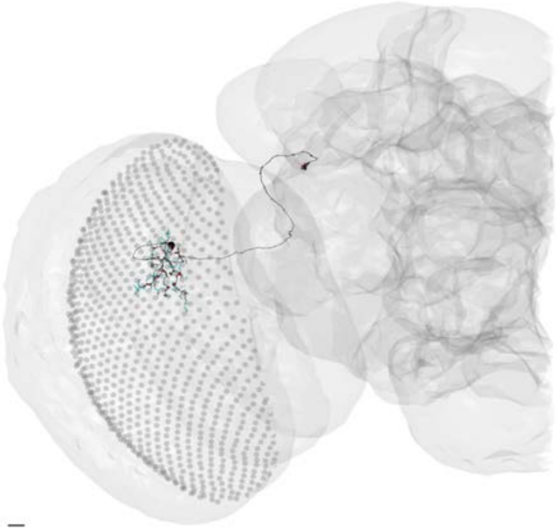

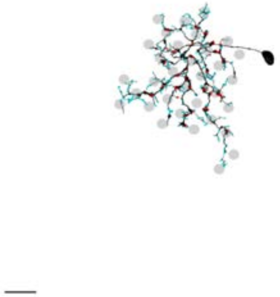

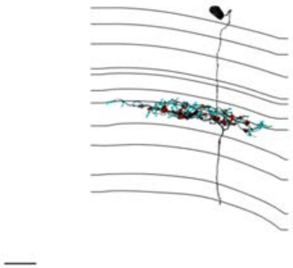

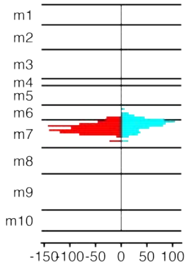


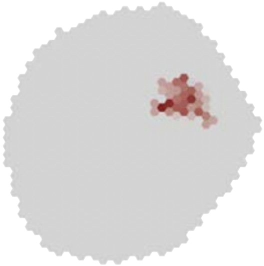

### MeTu1_720575940641954549

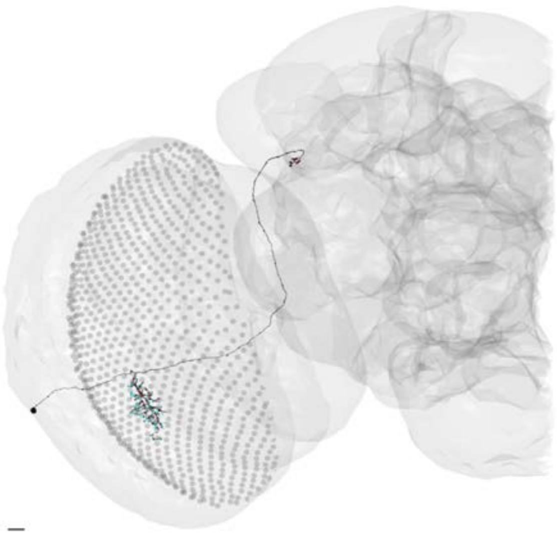

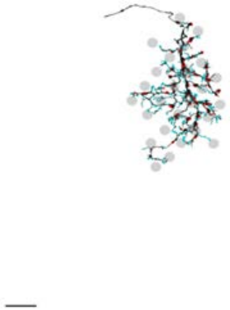

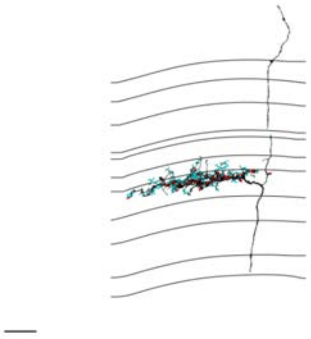

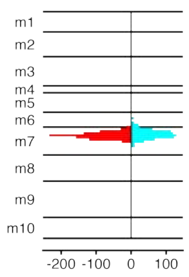


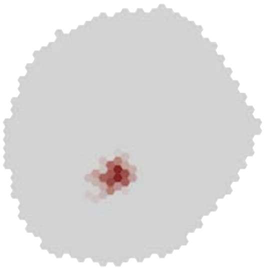

### MeTu1_720575940642371035

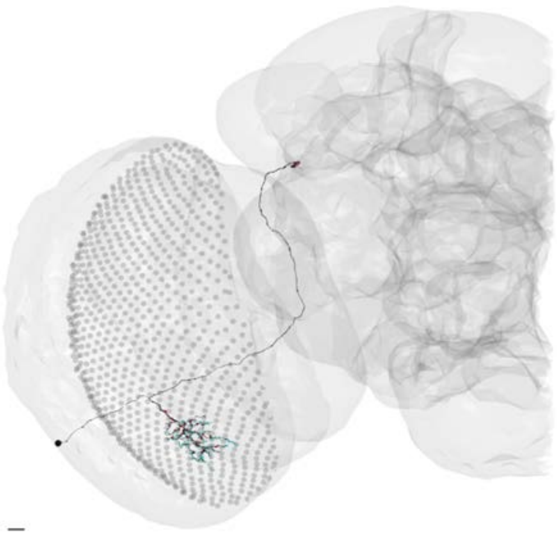

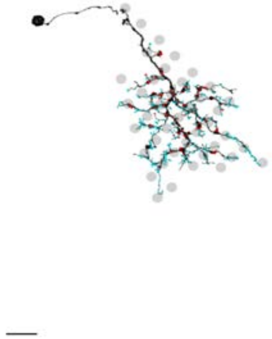

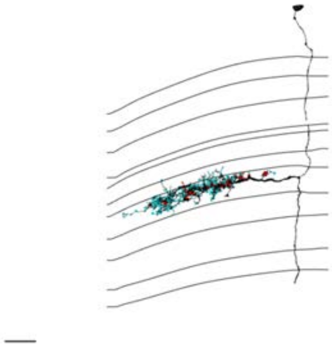

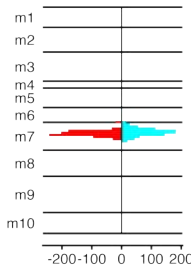


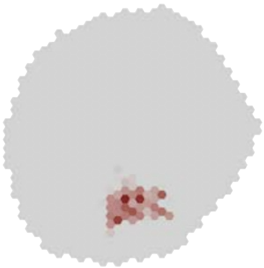

### MeTu1_720575940643613856

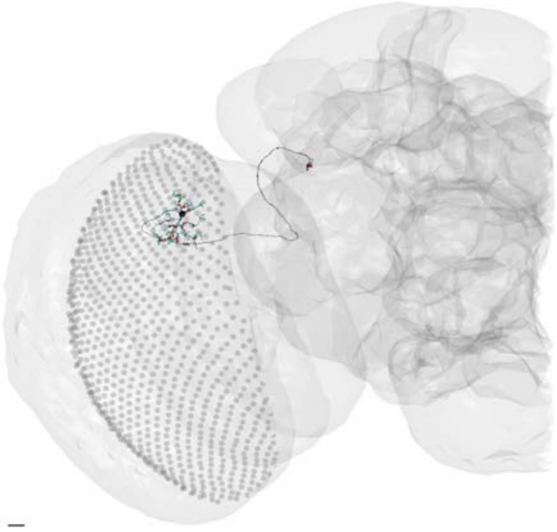

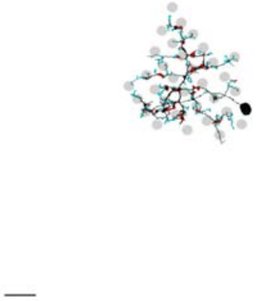

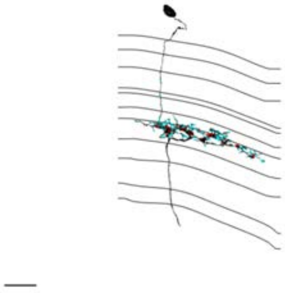

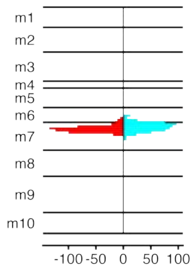


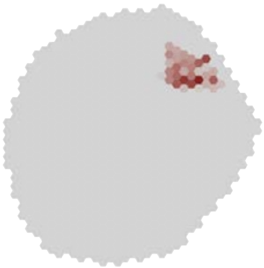

### MeTu1_720575940644670499

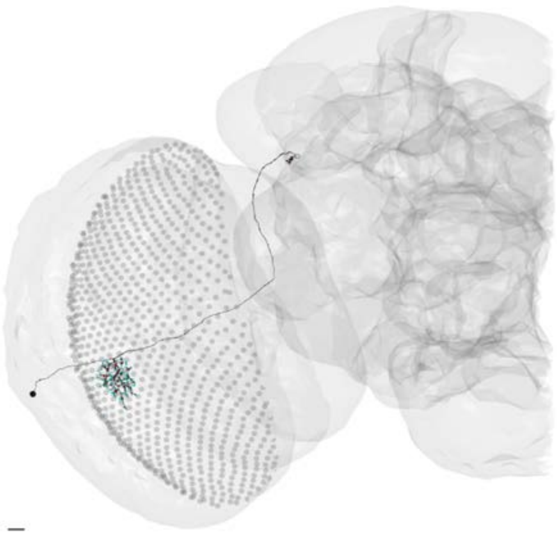

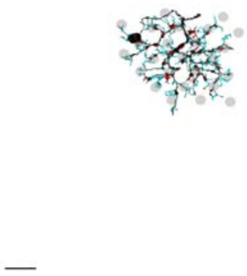

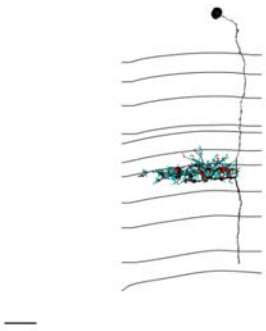

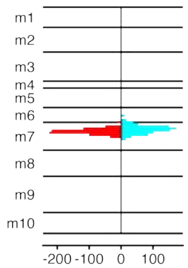


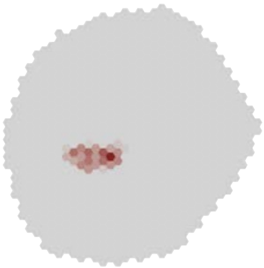

MeTu1_720575940645570327

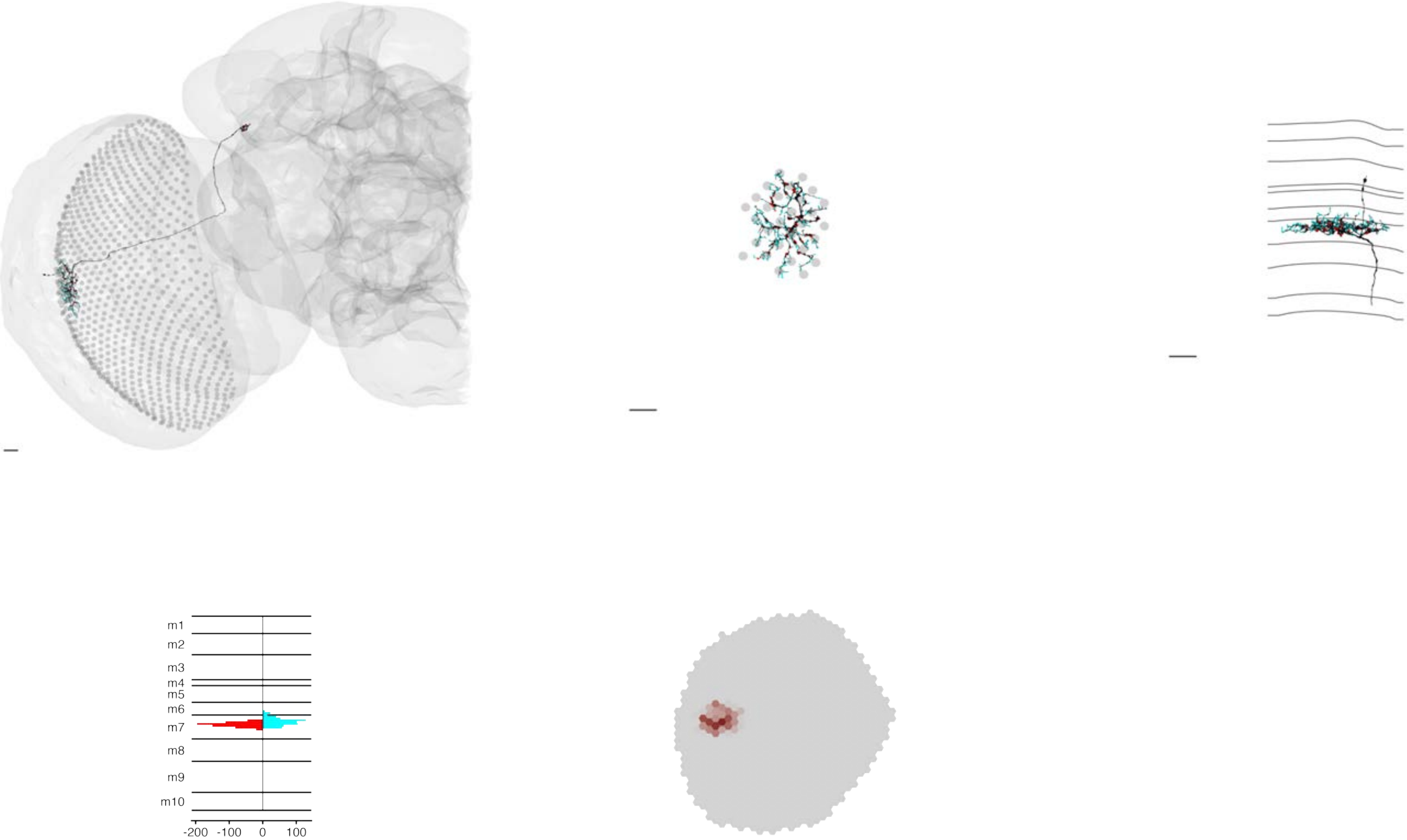

### MeTu1_720575940650407801

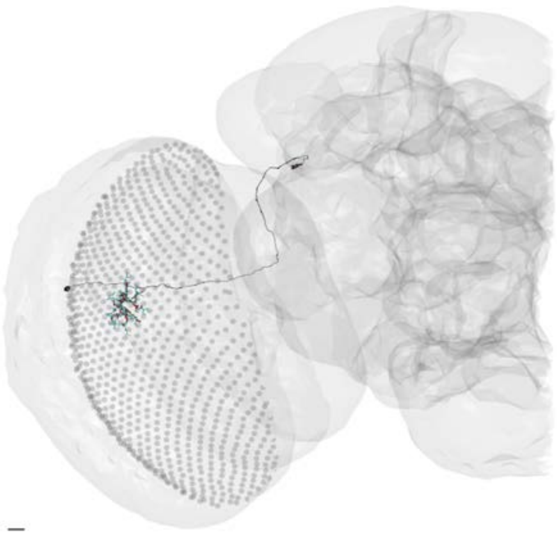

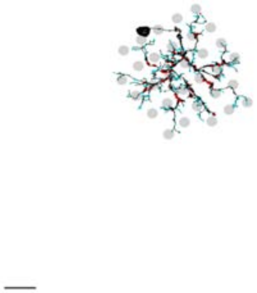

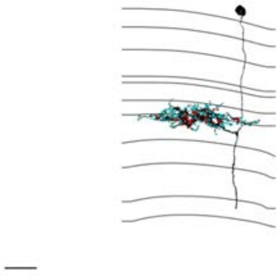

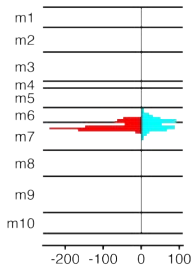


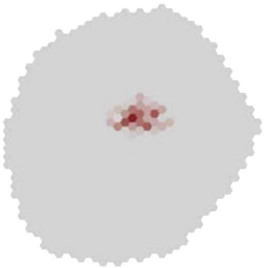

### MeTu2a_720575940604345632

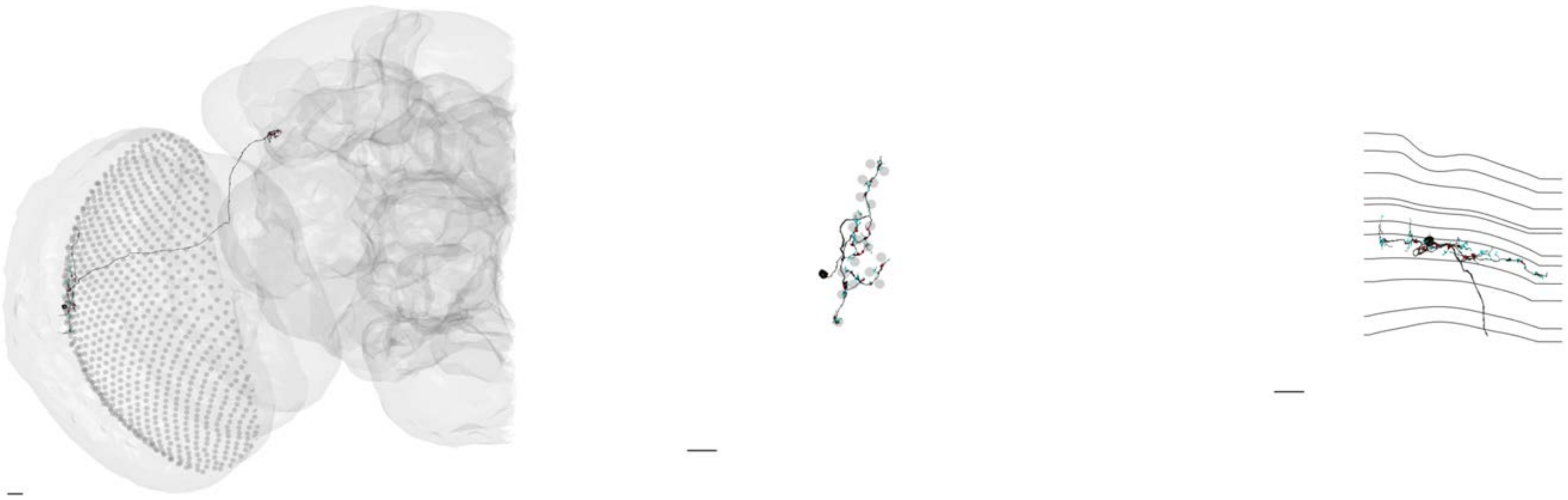

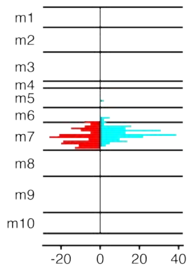


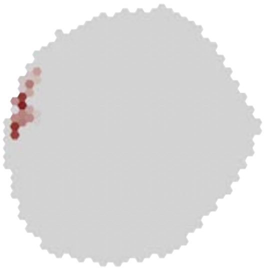

### MeTu2a_720575940605278386

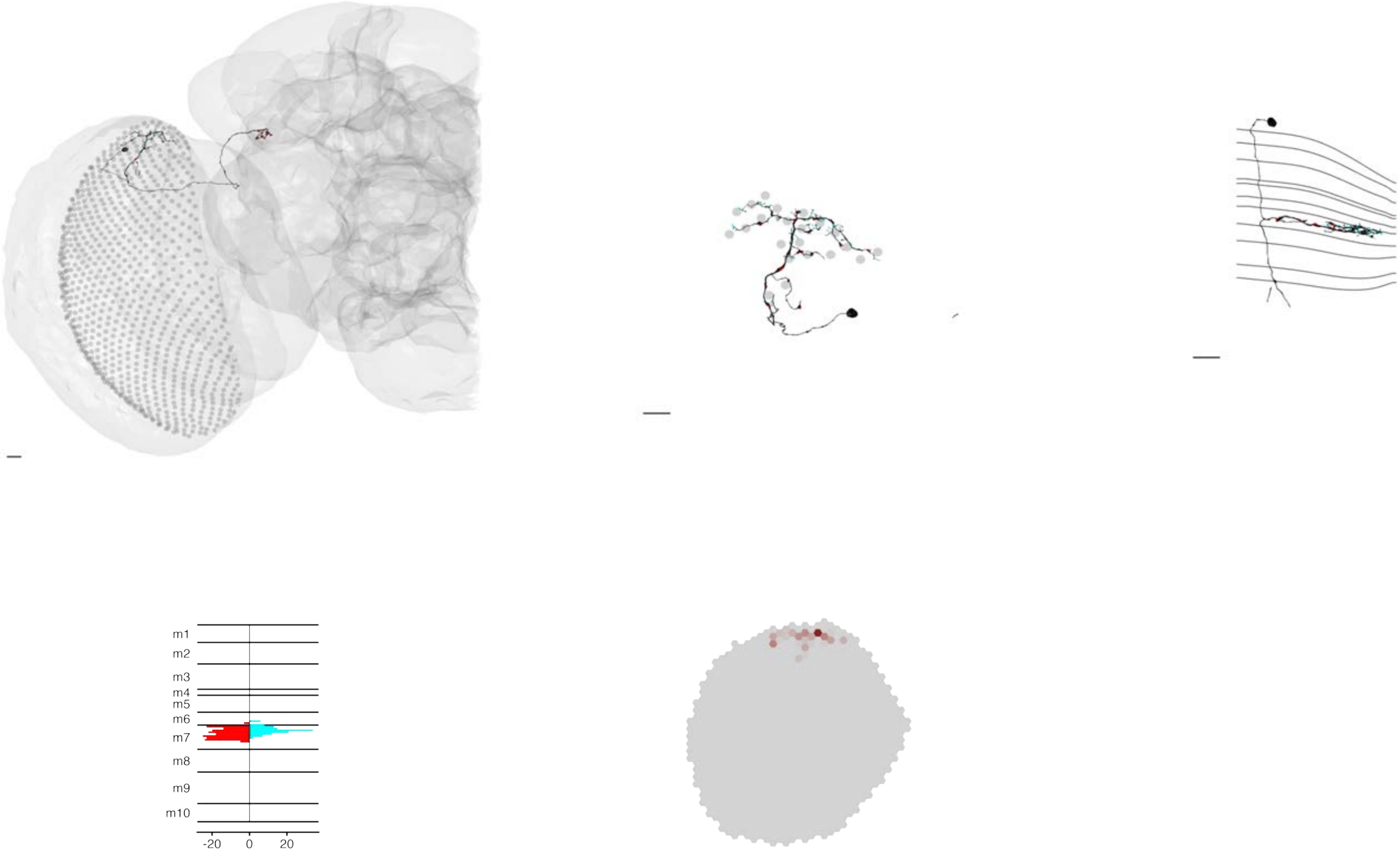

### MeTu2a_720575940606039072

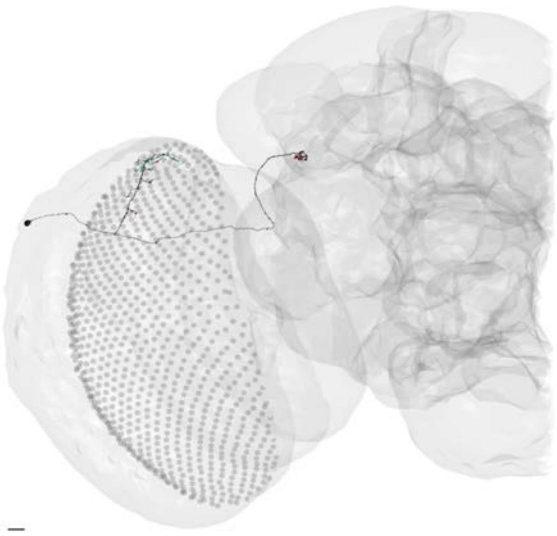

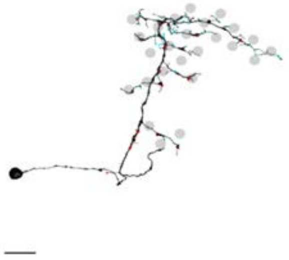

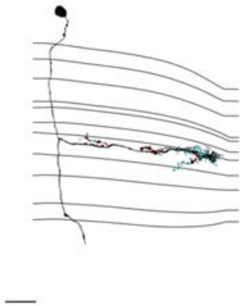

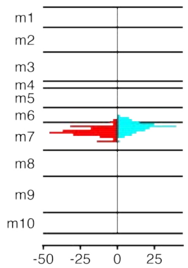


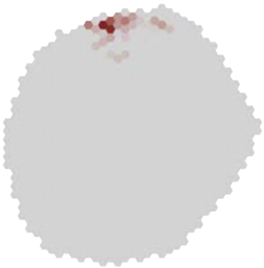

### MeTu2a_720575940606040352

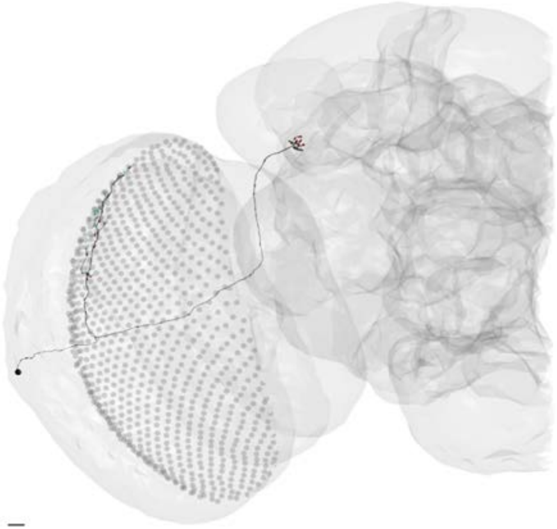

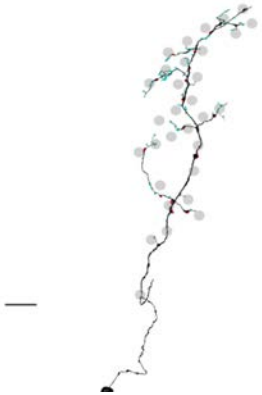

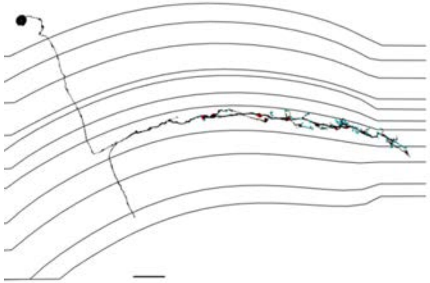

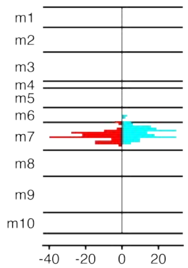


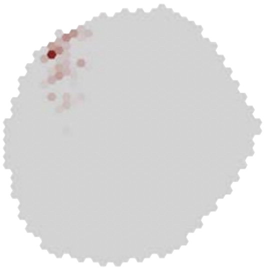

### MeTu2a_720575940606615474

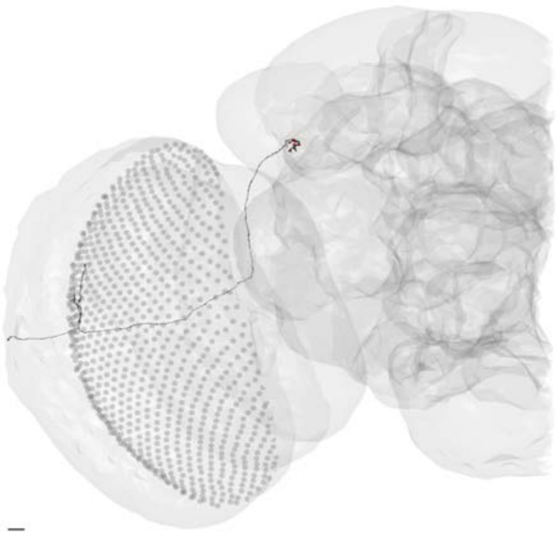

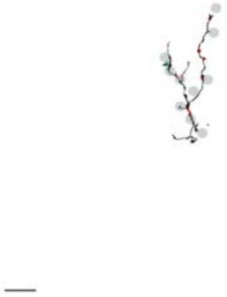

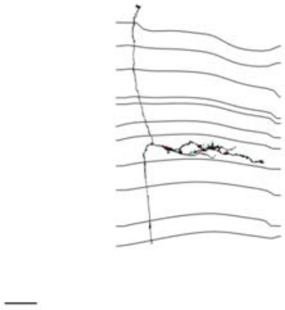

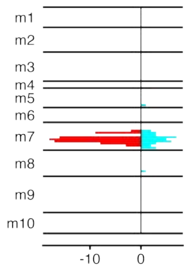


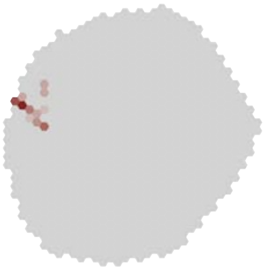

### MeTu2a_720575940610585366

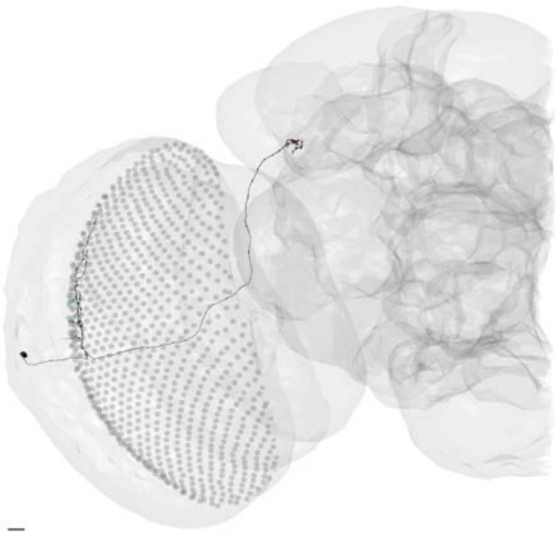

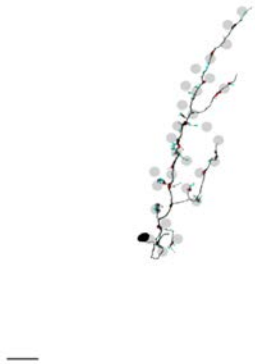

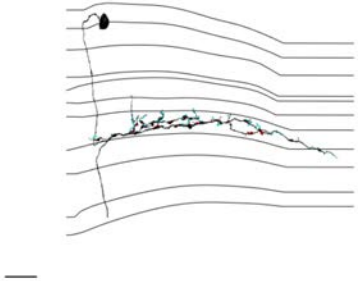

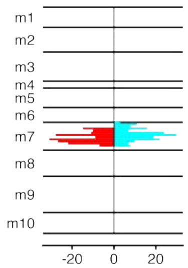


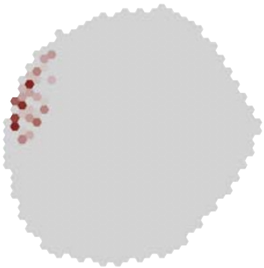

MeTu2a_720575940611532458

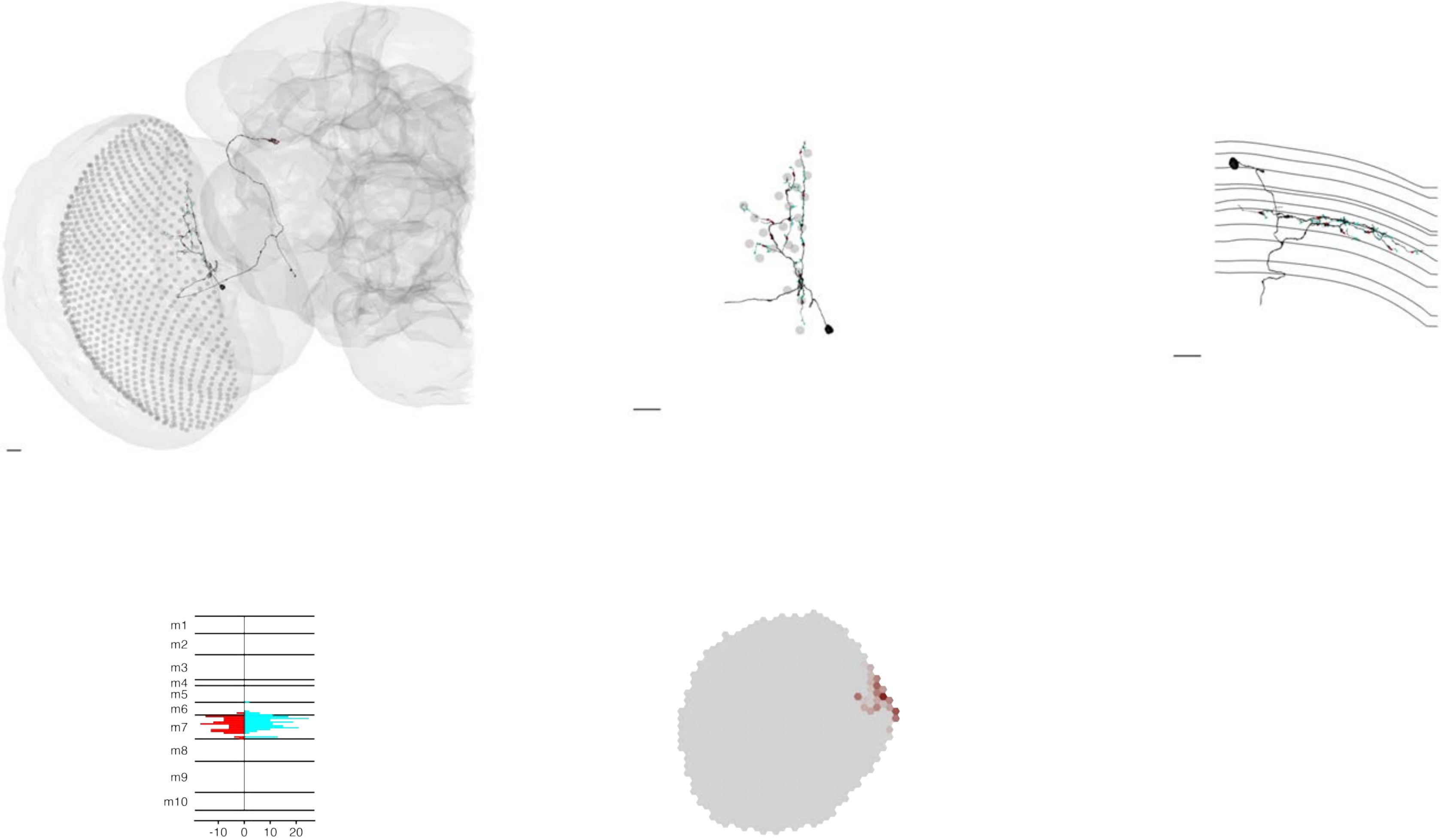

### MeTu2a_720575940612162814

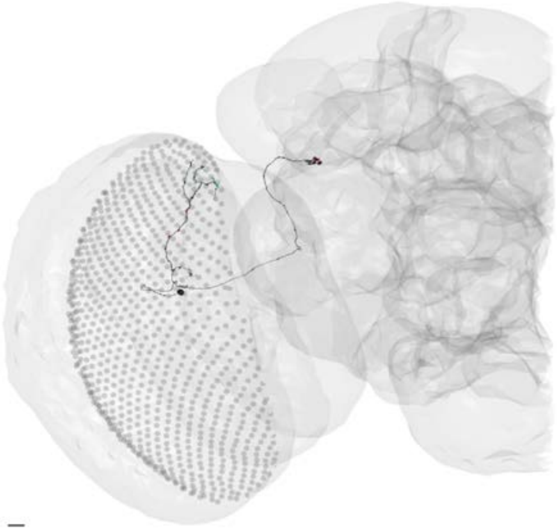

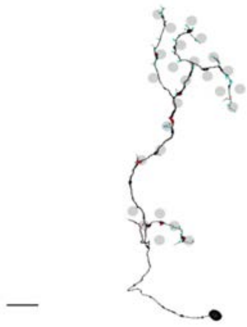

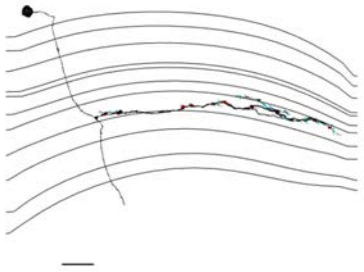

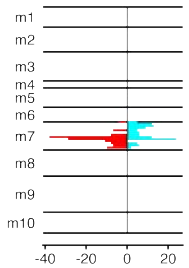


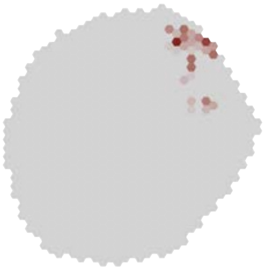

MeTu2a_720575940613145366

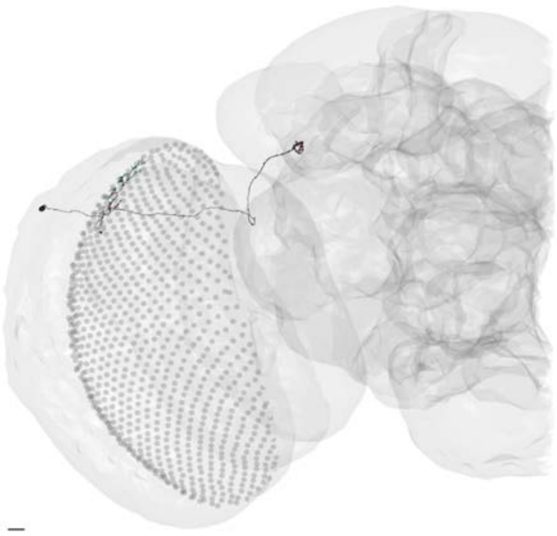

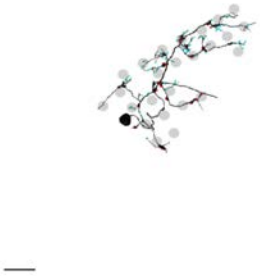

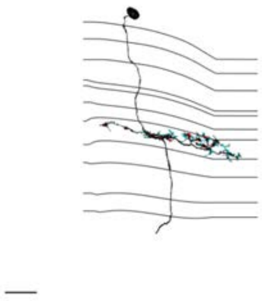

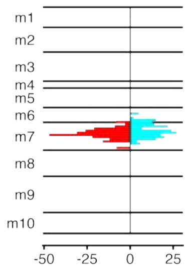


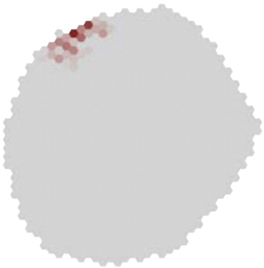

### MeTu2a_720575940614675921

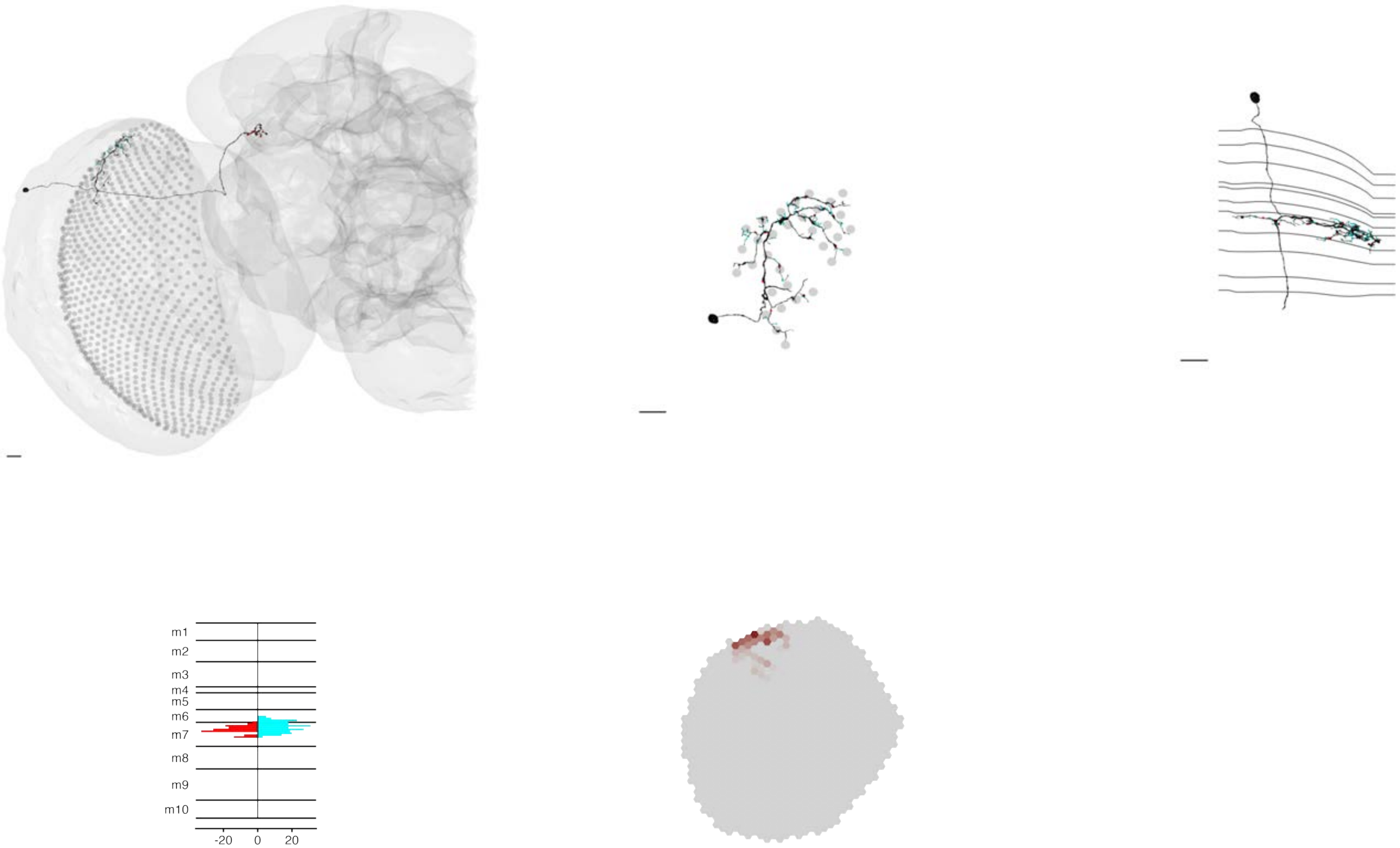

### MeTu2a_720575940615406954

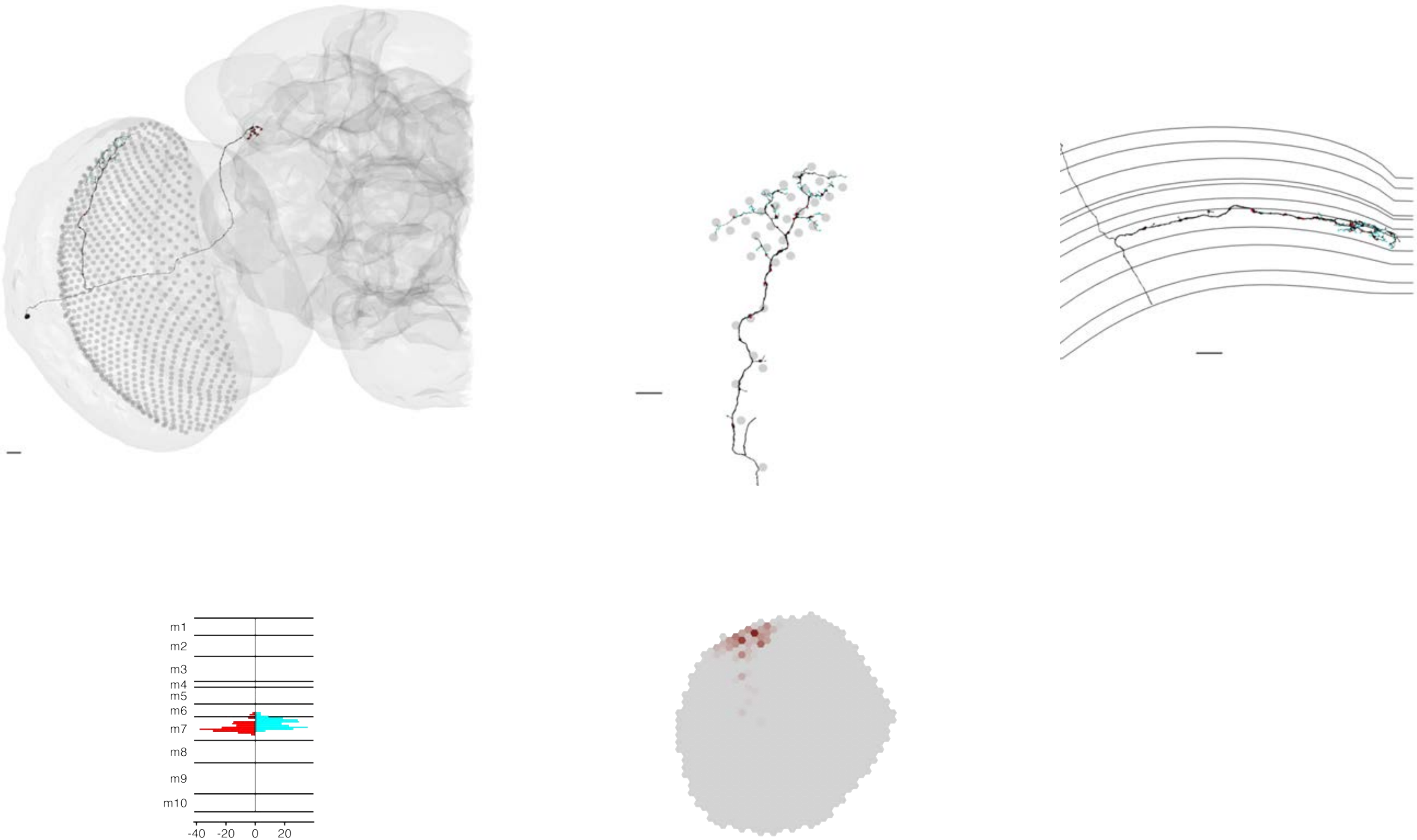

### MeTu2a_720575940615643754

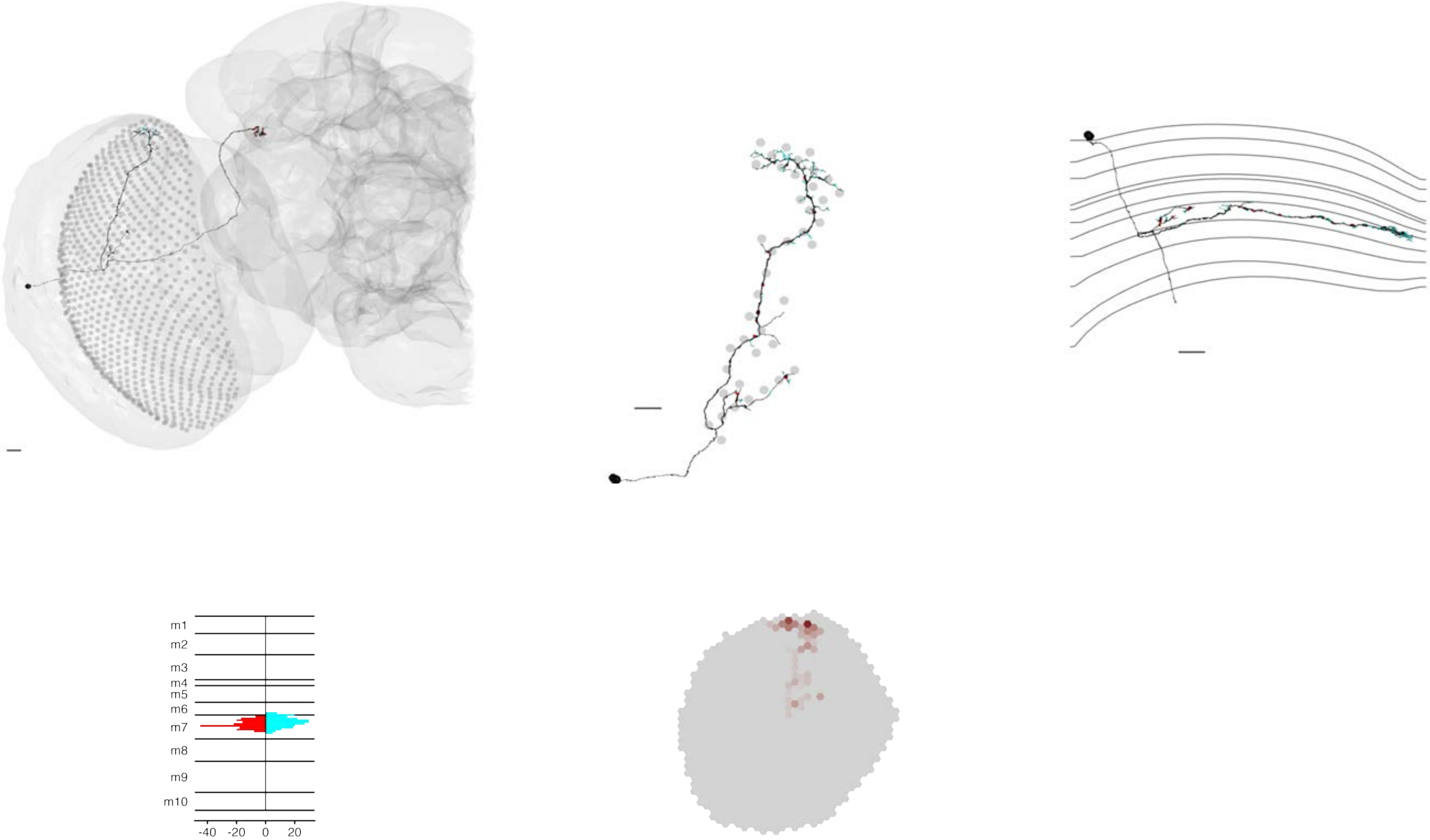

### MeTu2a_720575940616968059

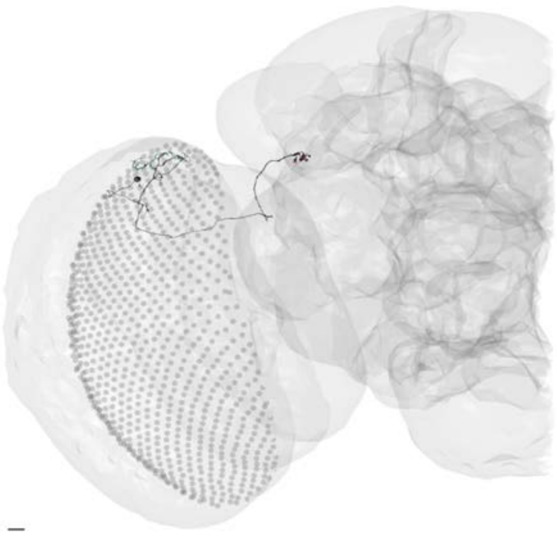

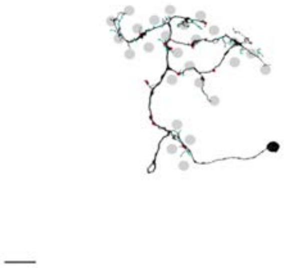

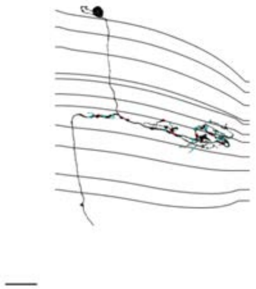

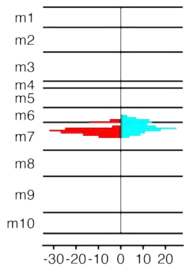


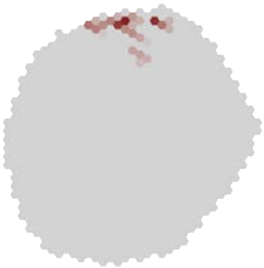

### MeTu2a_720575940617779781

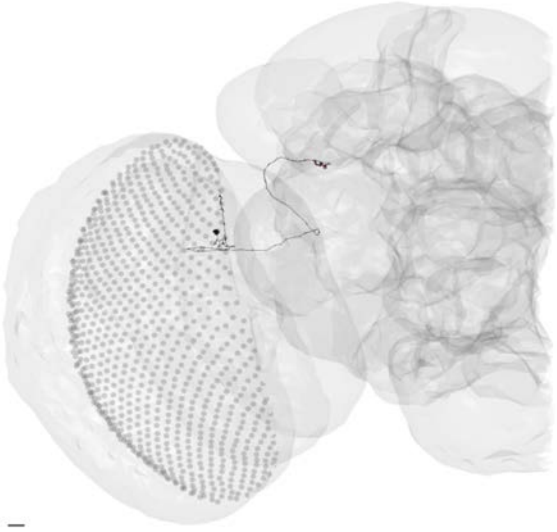

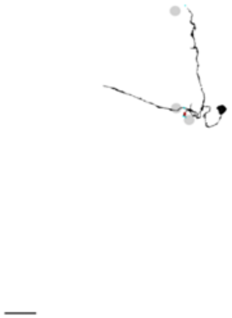

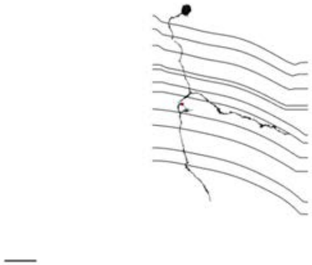

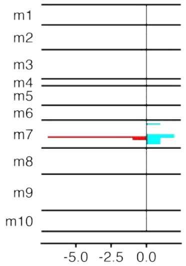


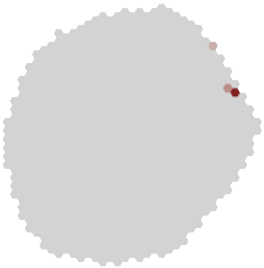

### MeTu2a_720575940618086841

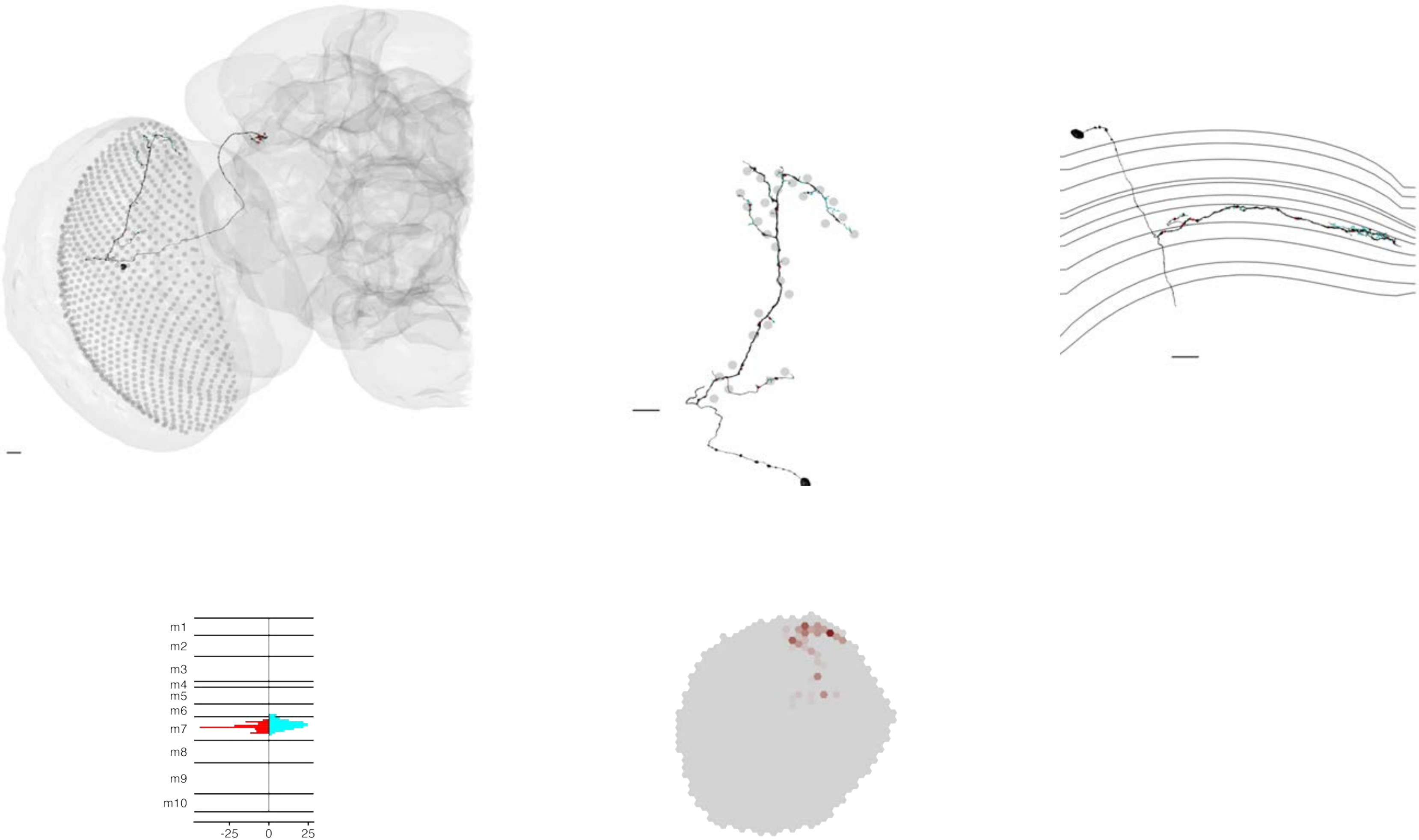

MeTu2a_720575940618961773

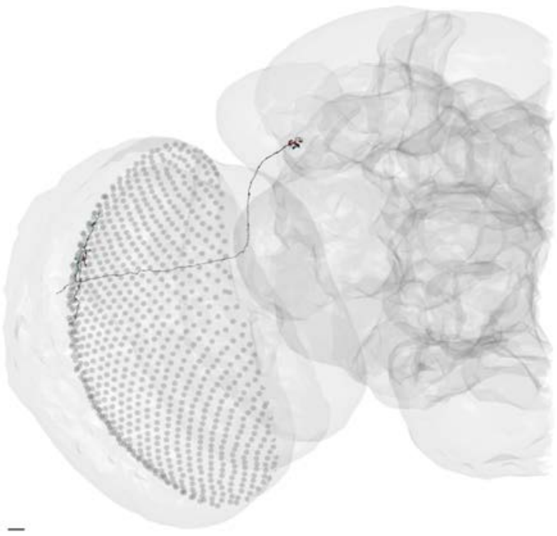

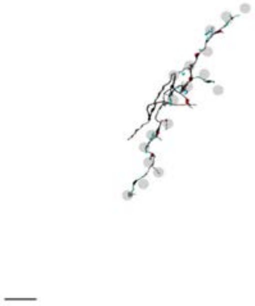

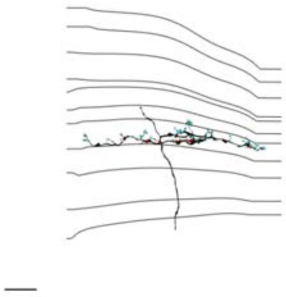

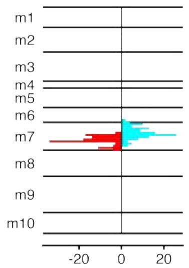


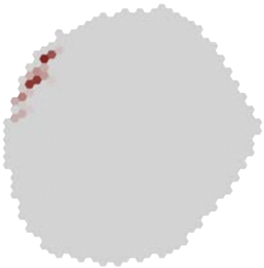

### MeTu2a_720575940619364724

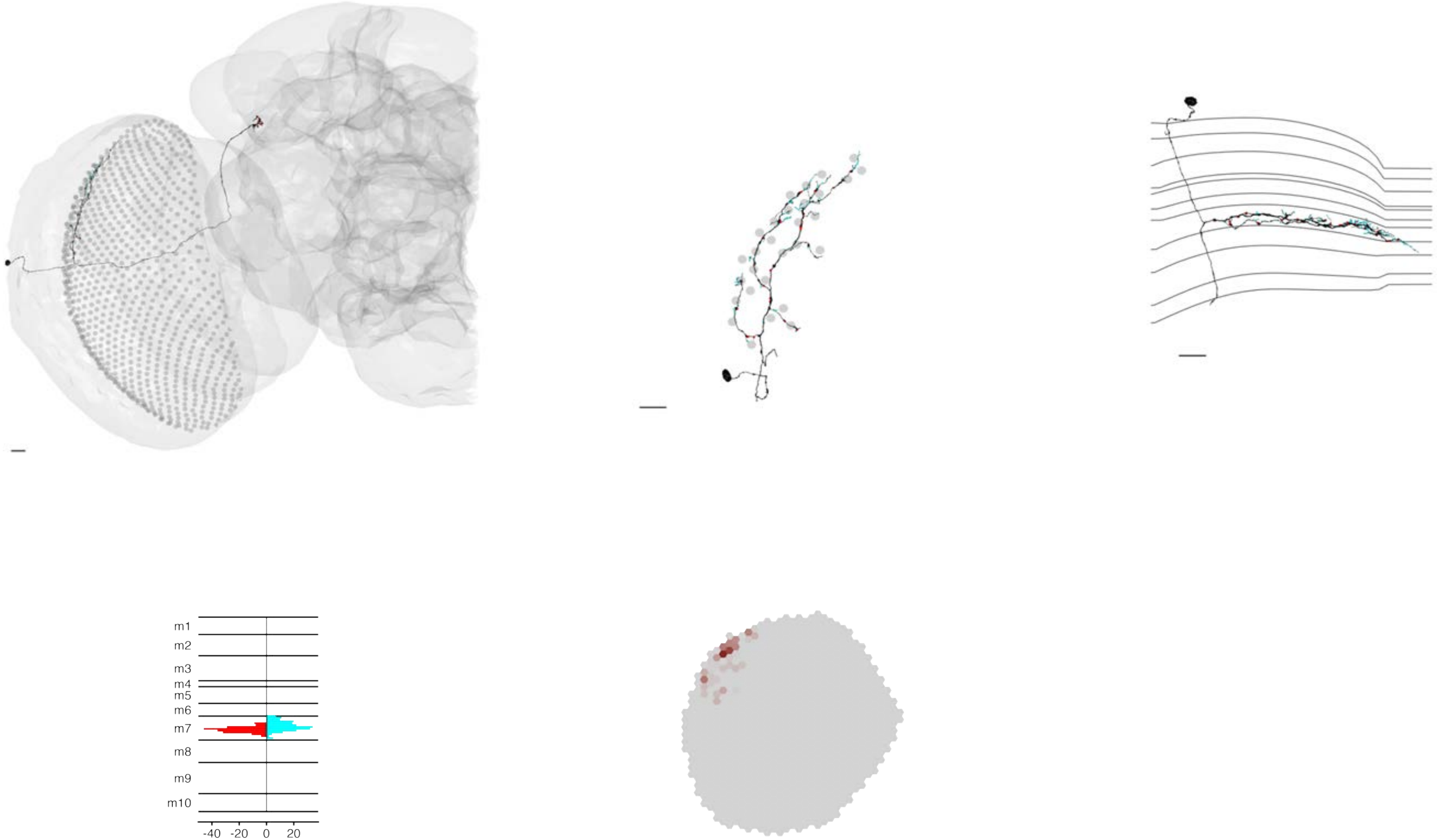

### MeTu2a_720575940620345429

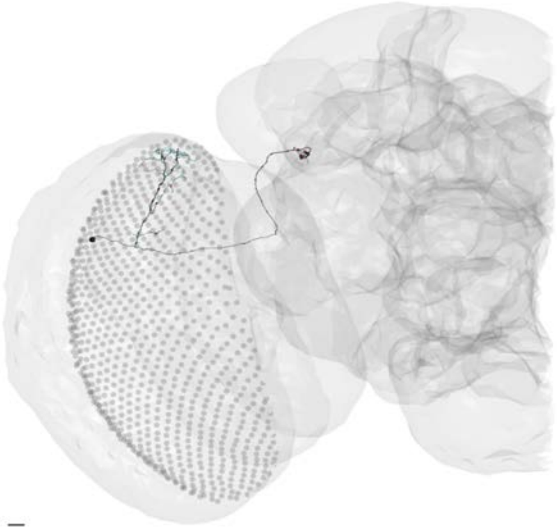

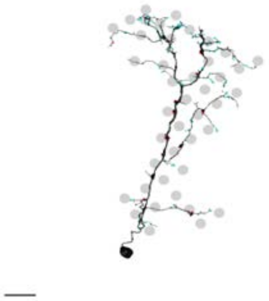

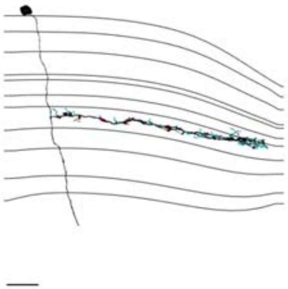

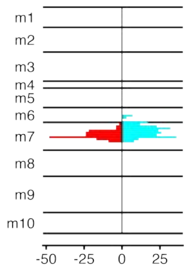


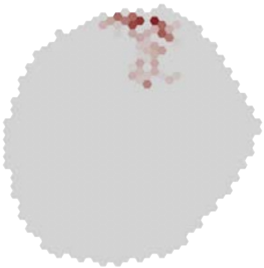

### MeTu2a_720575940621063406

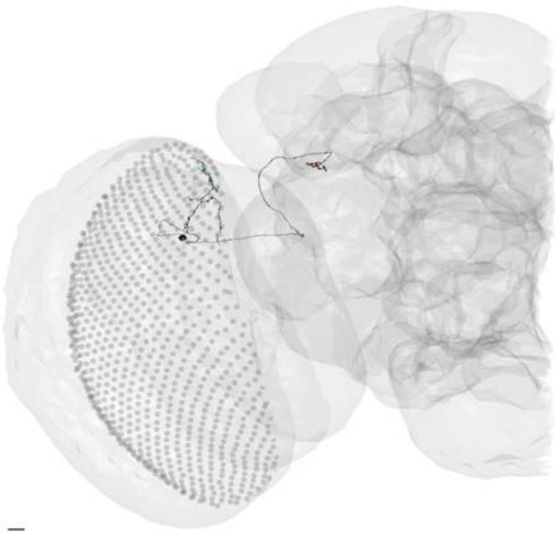

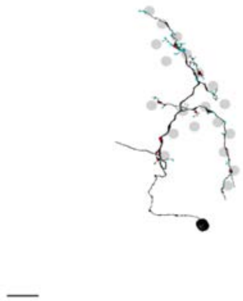

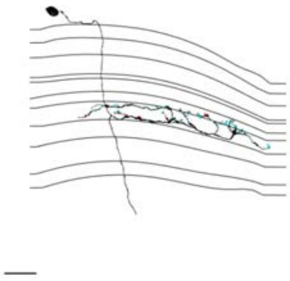

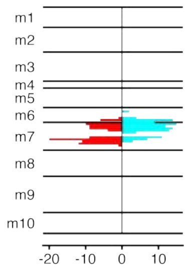


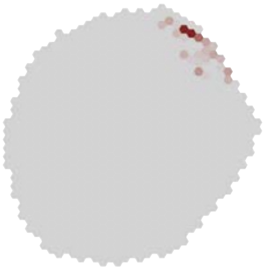

MeTu2a_720575940621545194

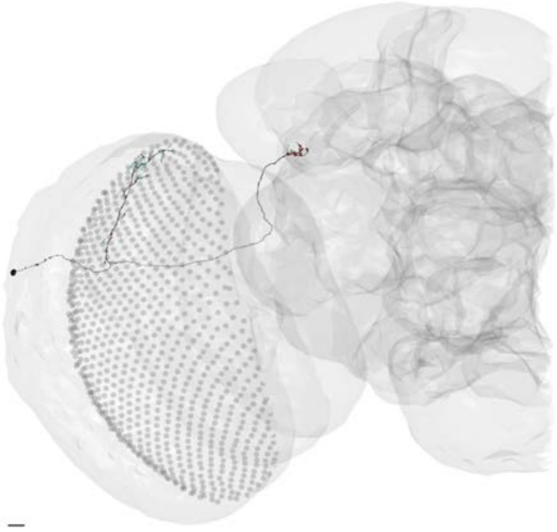

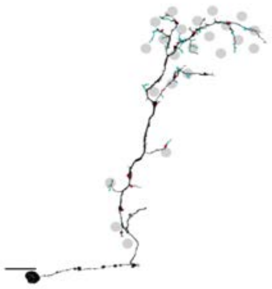

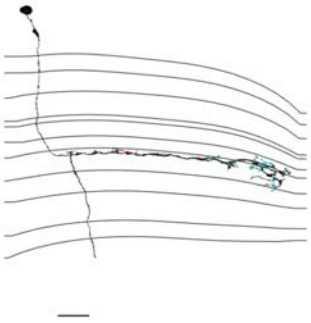

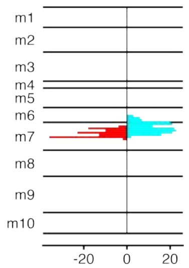


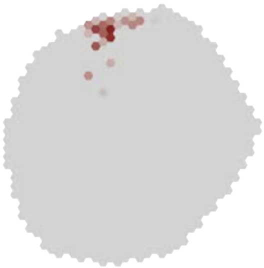

MeTu2a_720575940623830419

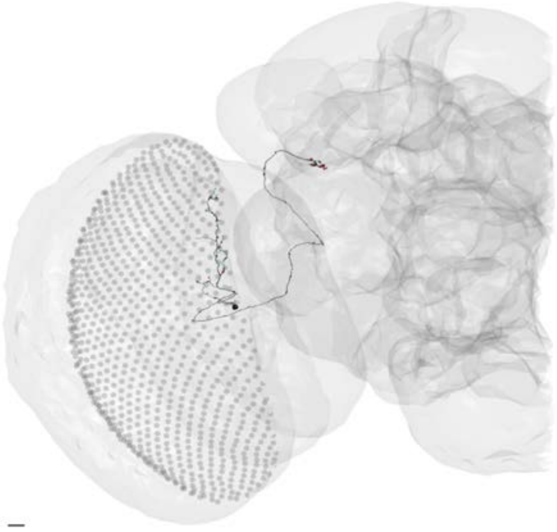

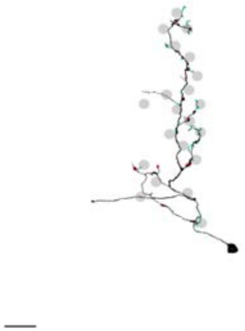

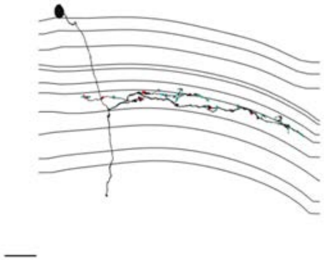

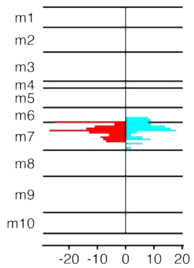


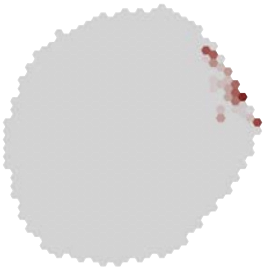

### MeTu2a_720575940625065195

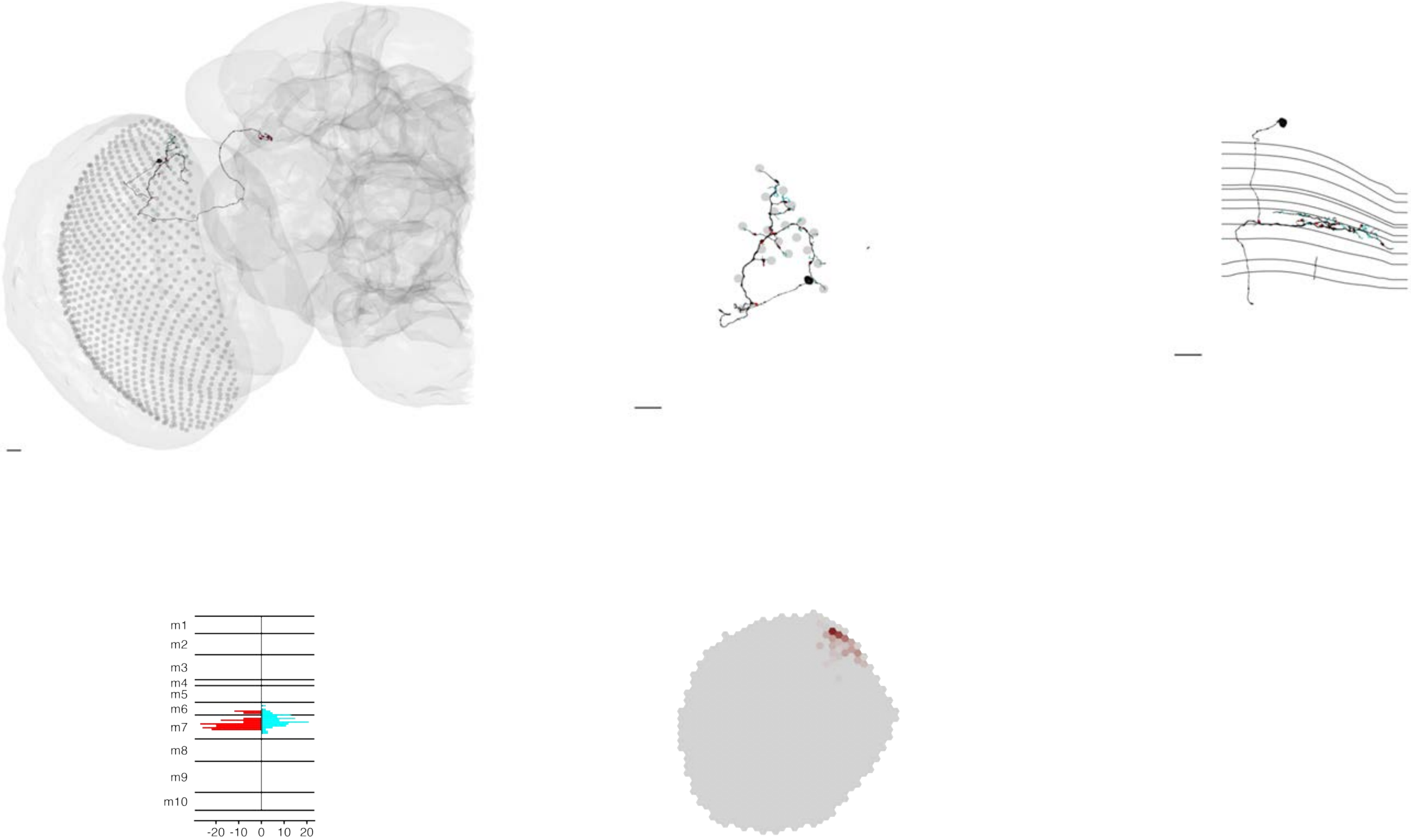

### MeTu2a_720575940627631515

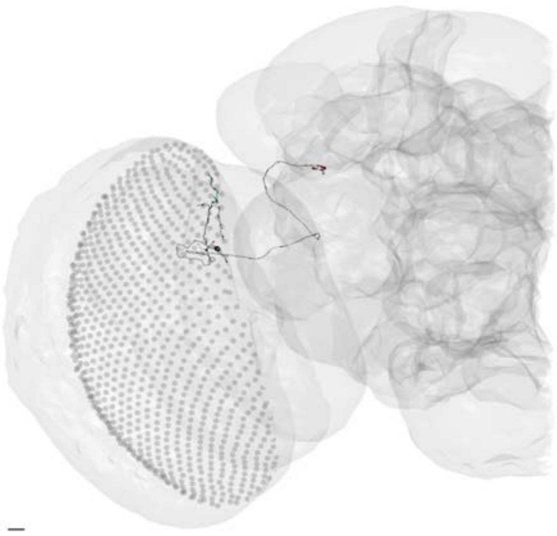

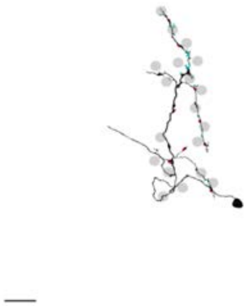

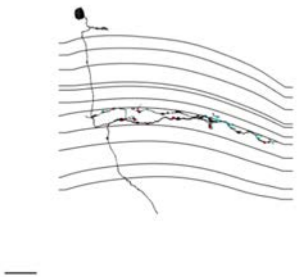

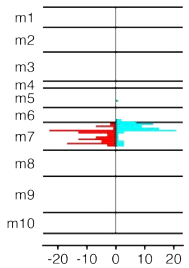


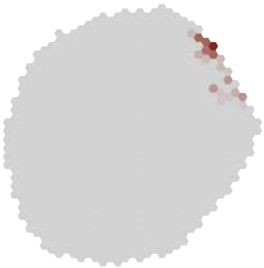

### MeTu2a_720575940627715706

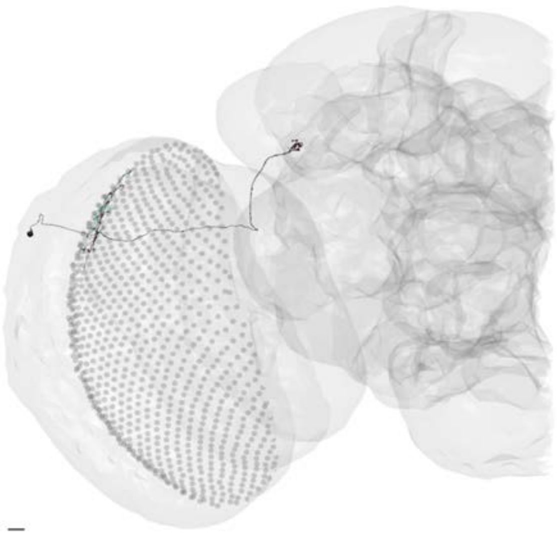

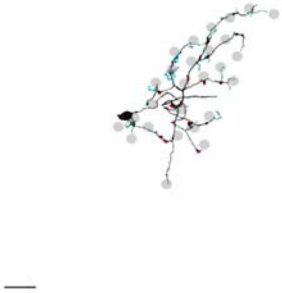

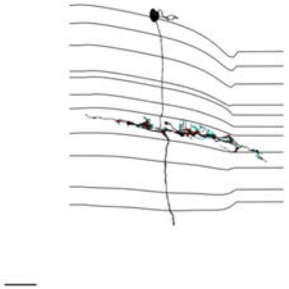

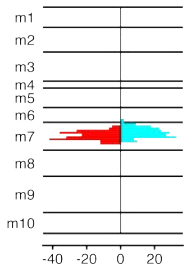


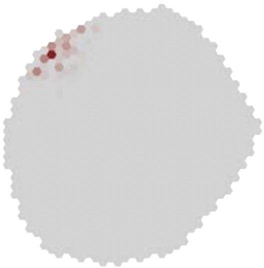

### MeTu2a_720575940628706280

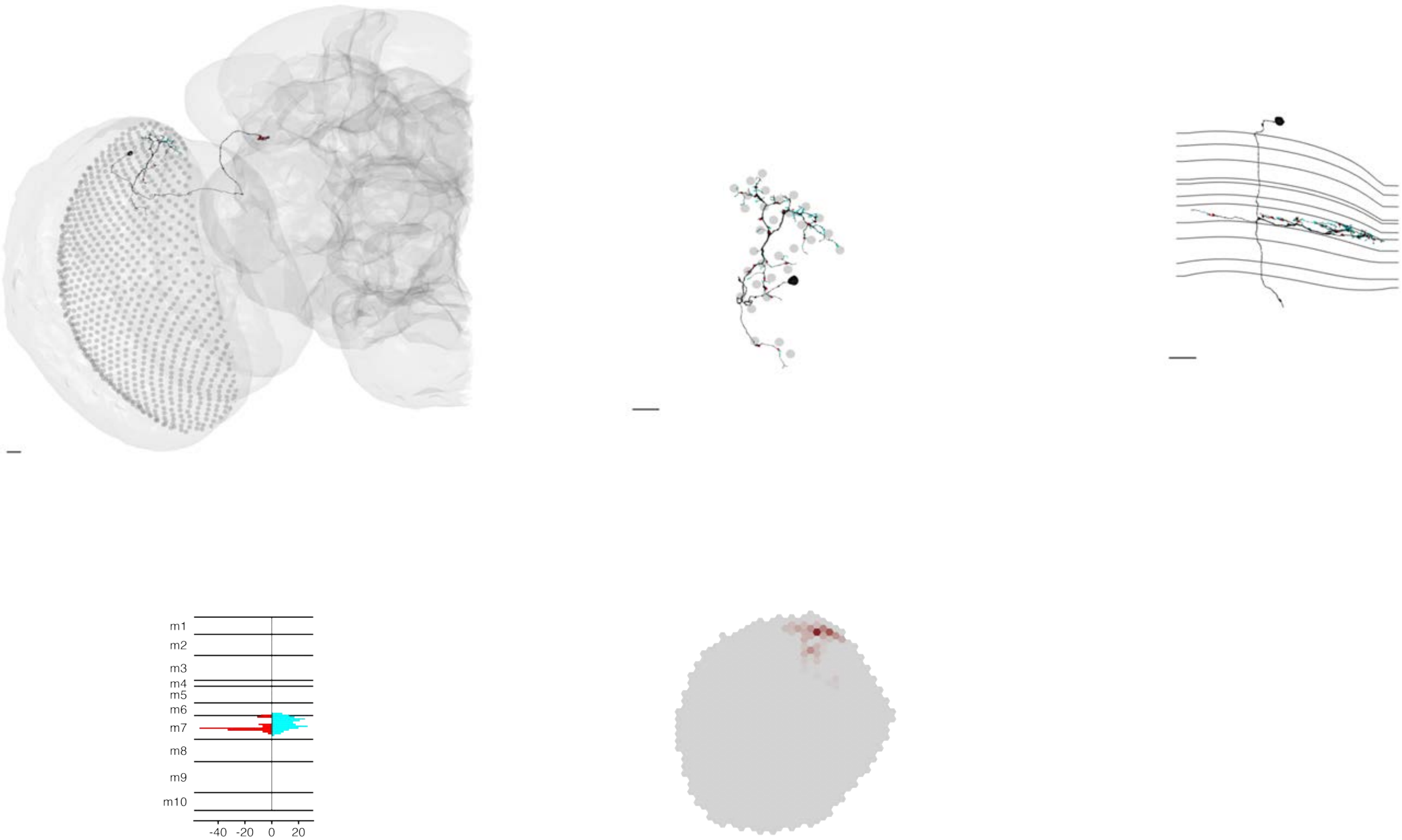

### MeTu2a_720575940629663800

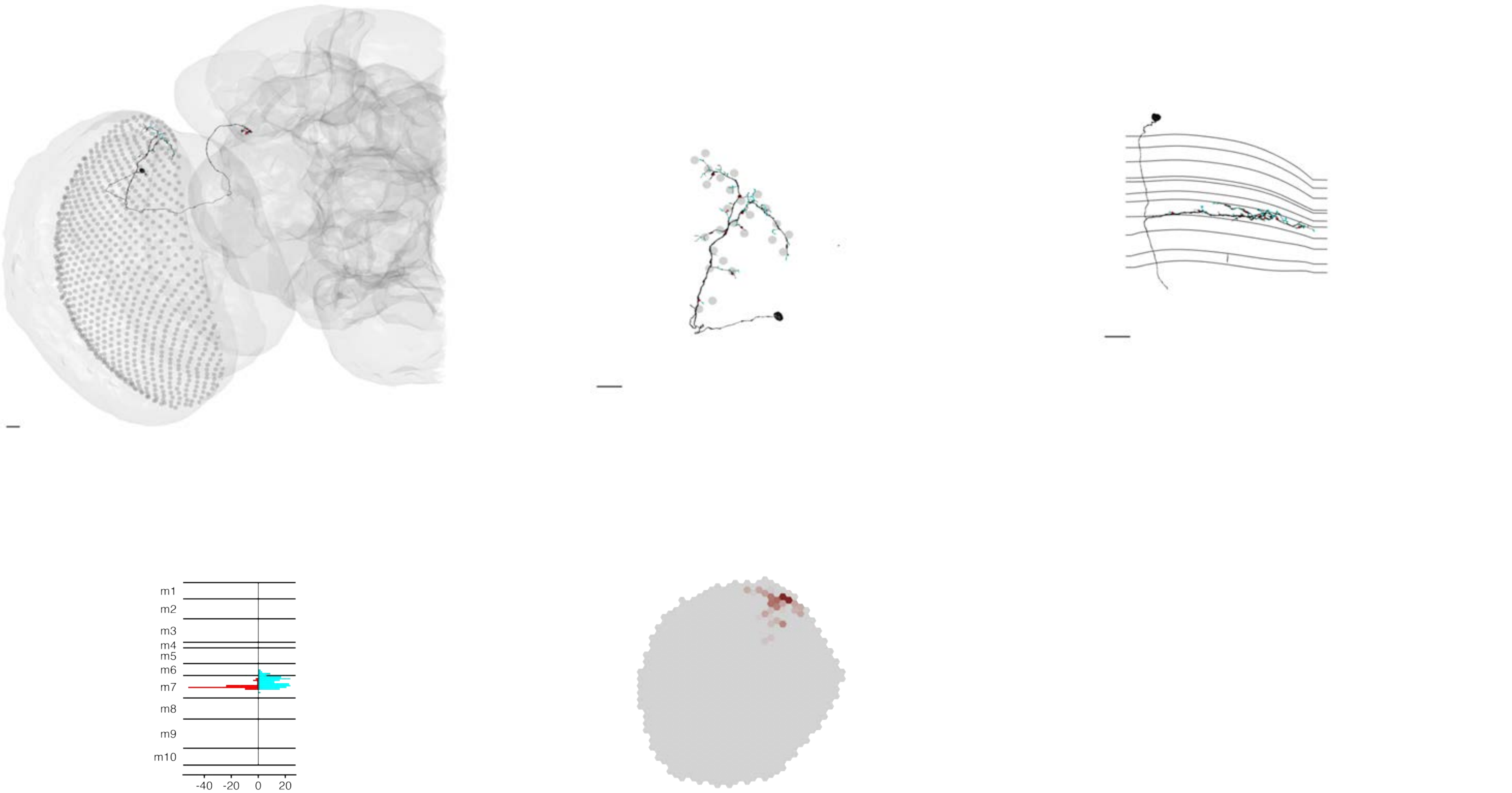

### MeTu2a_720575940630945644

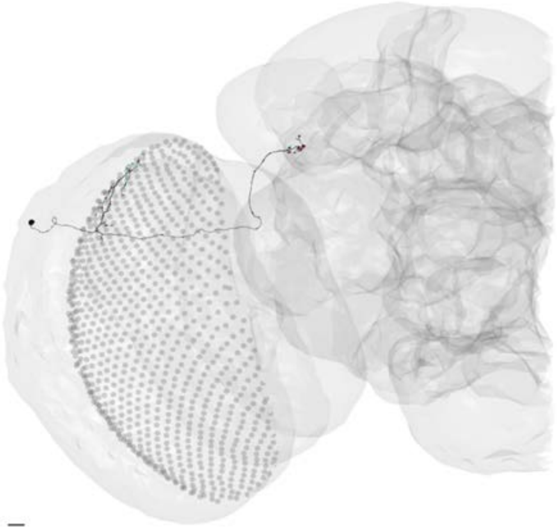

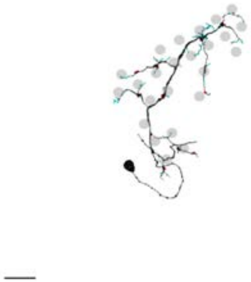

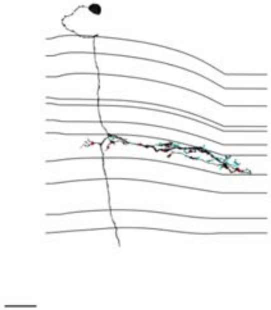

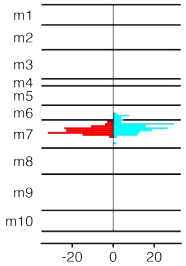


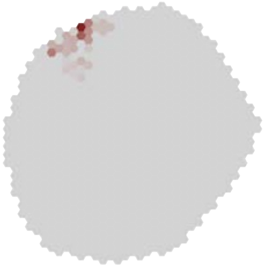

### MeTu2a_720575940635404919

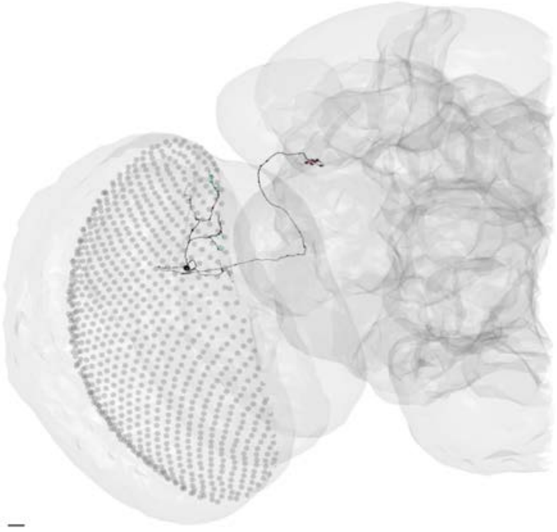

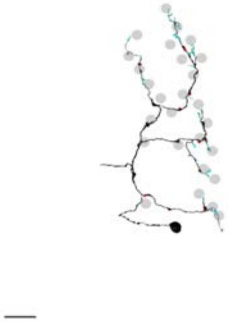

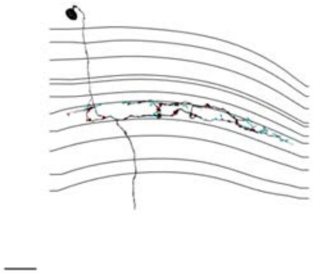

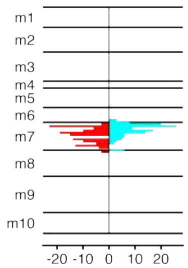


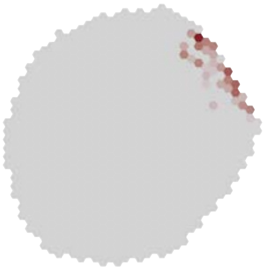

### MeTu2a_720575940635917236

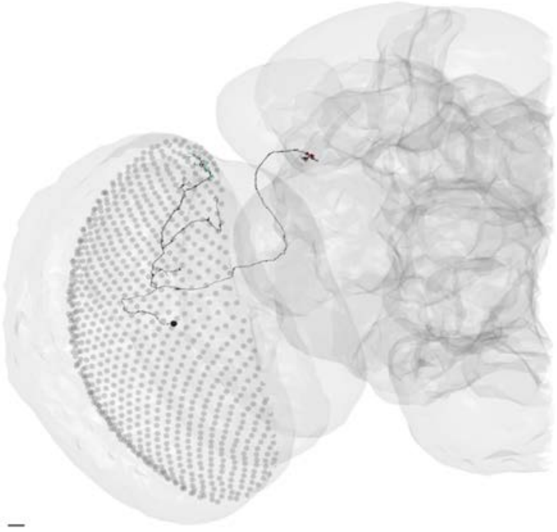

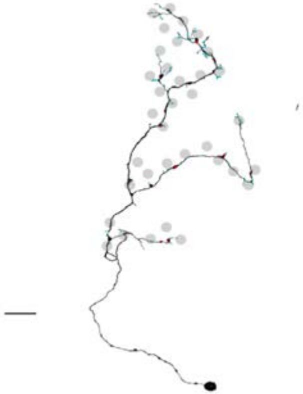

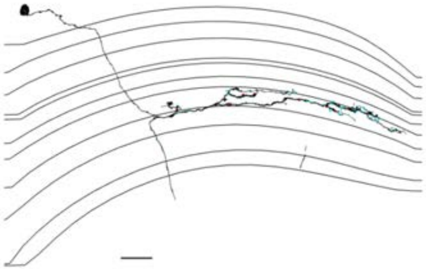

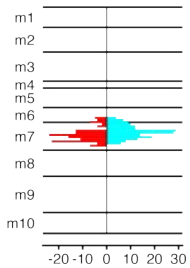


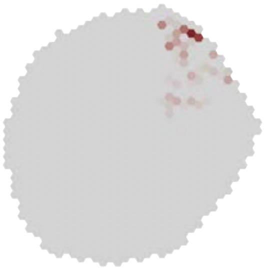

### MeTu2a_720575940636070836

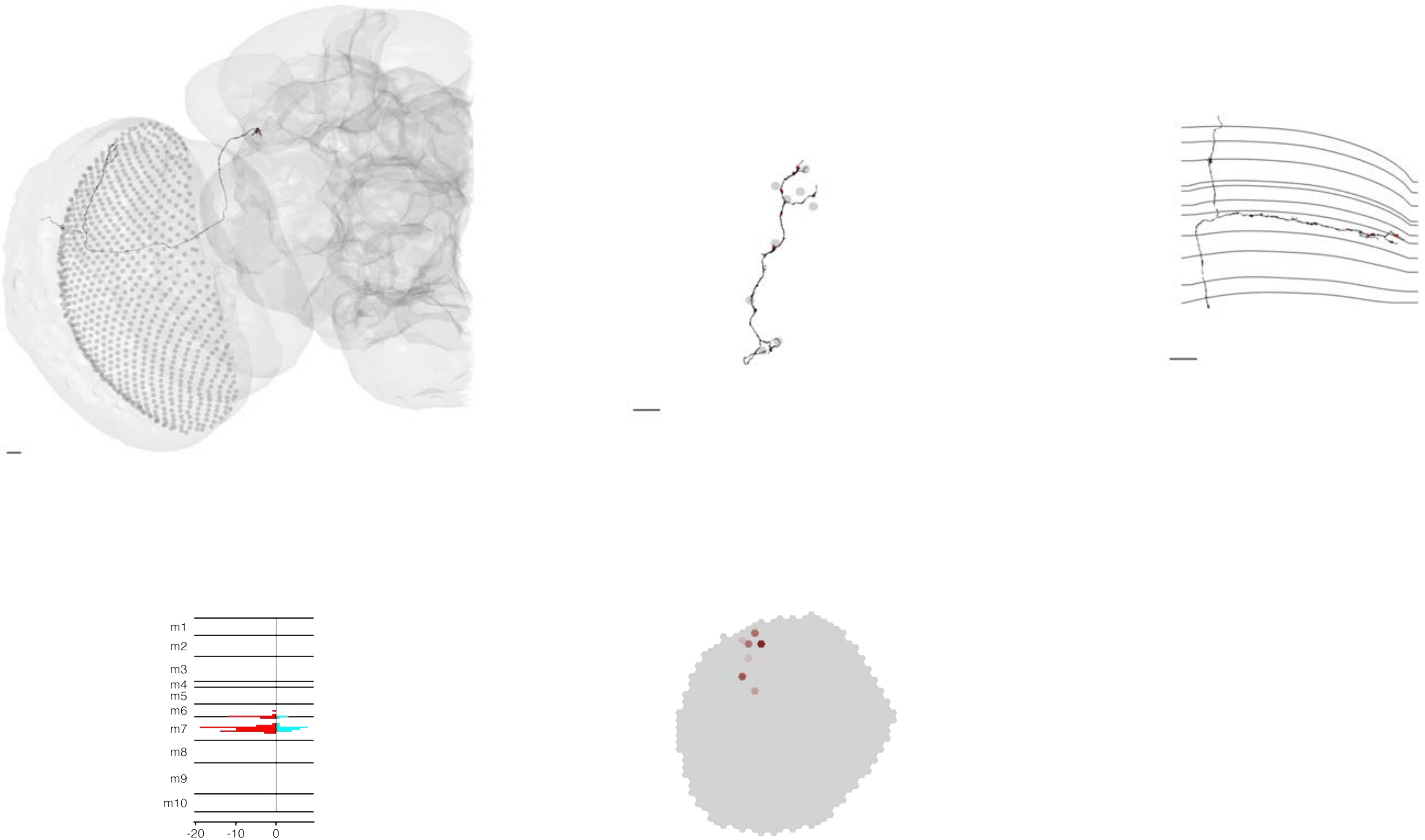

### MeTu2a_720575940636361839

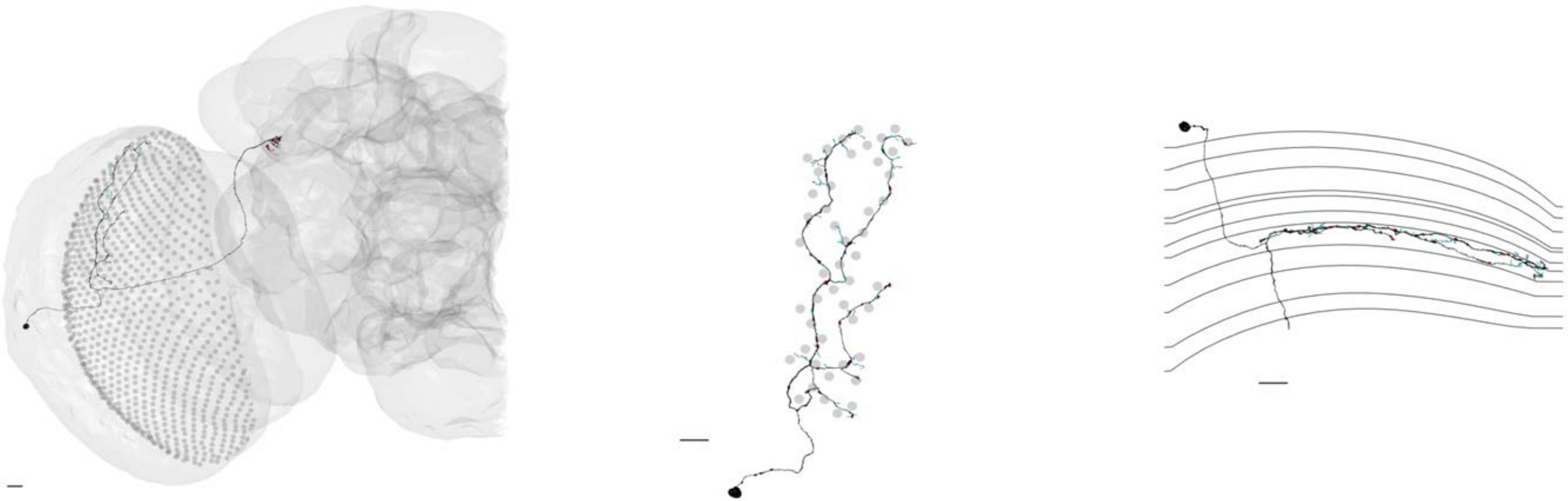

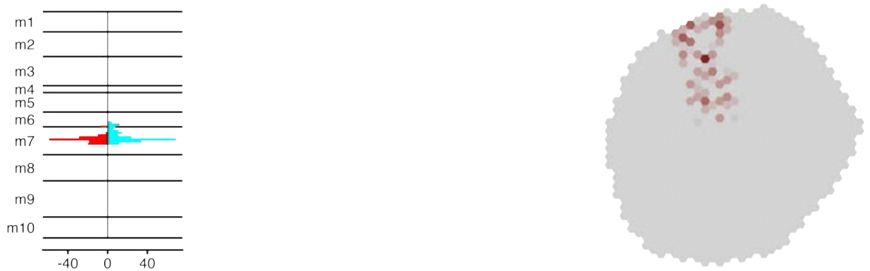

### MeTu2a_720575940637227438

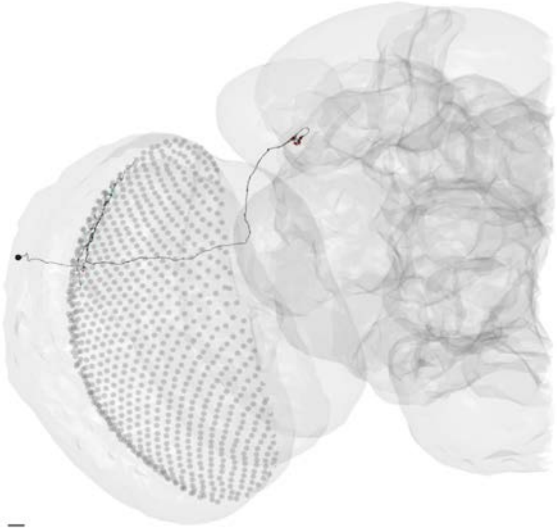

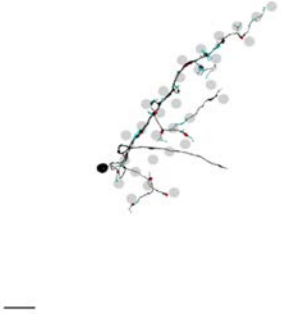

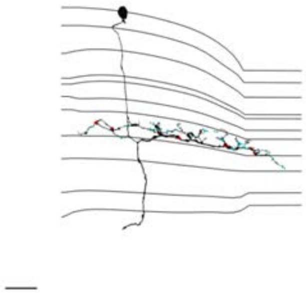

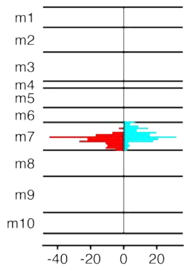


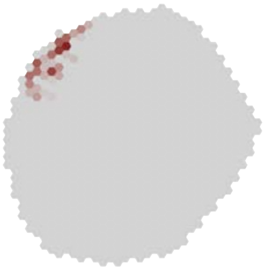

### MeTu2a_720575940638825193

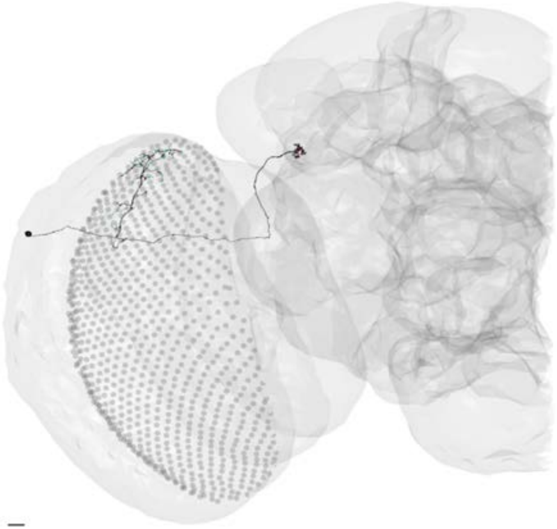

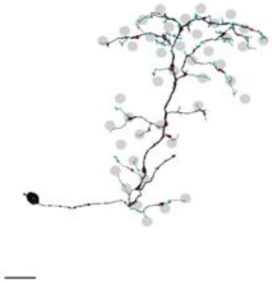

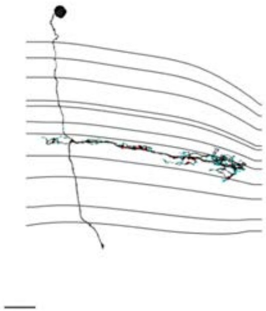

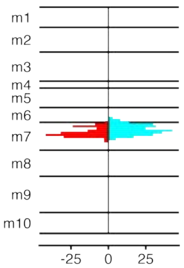


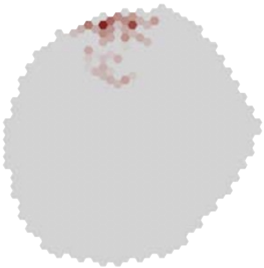

### MeTu2a_720575940641942472

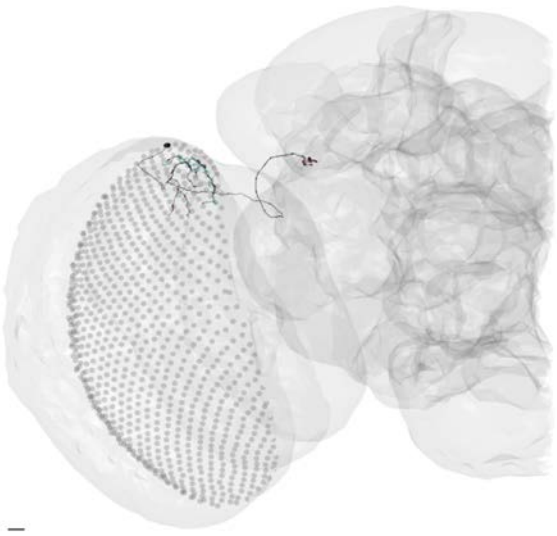

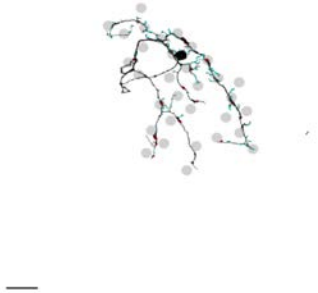

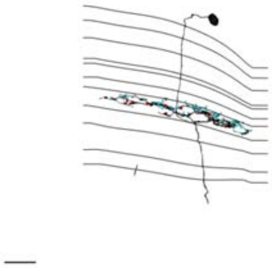

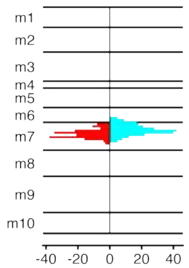


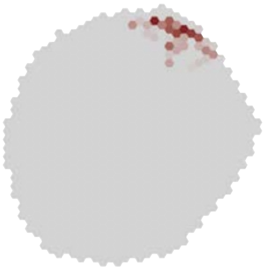

### MeTu2a_720575940646439044

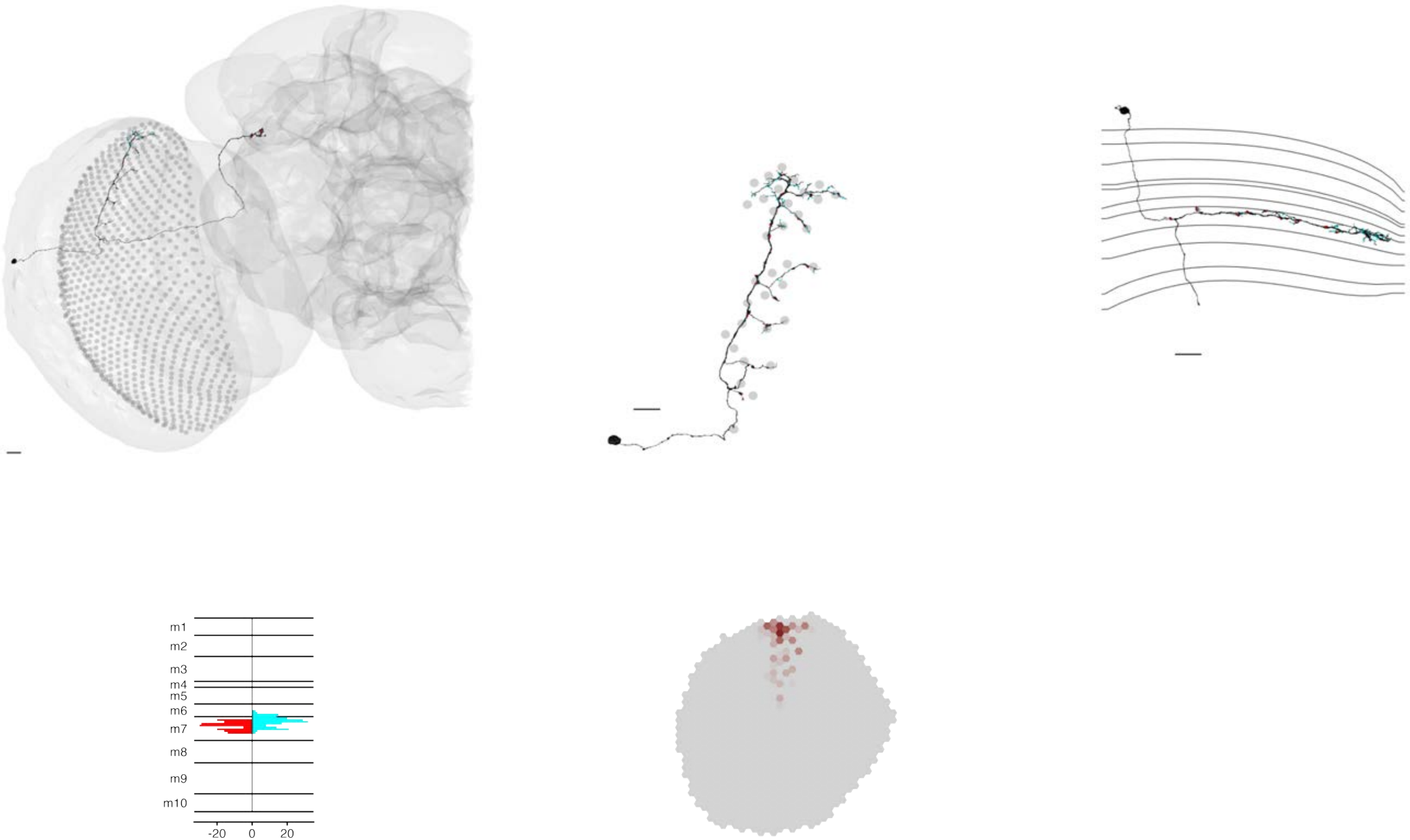

### MeTu2a_720575940658755201

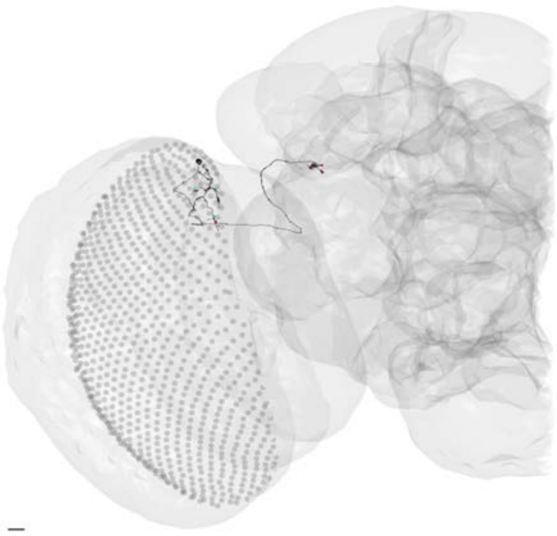

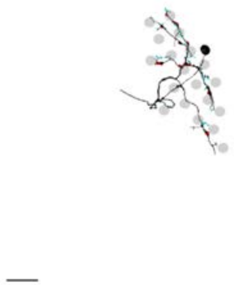

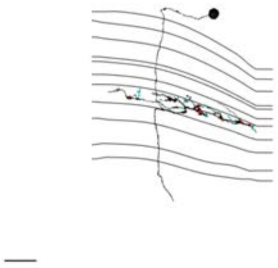

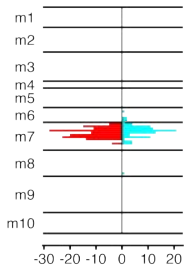


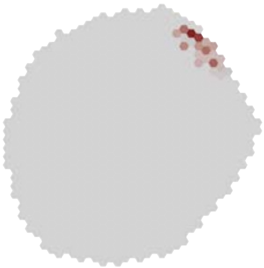

### MeTu2b_720575940605496777

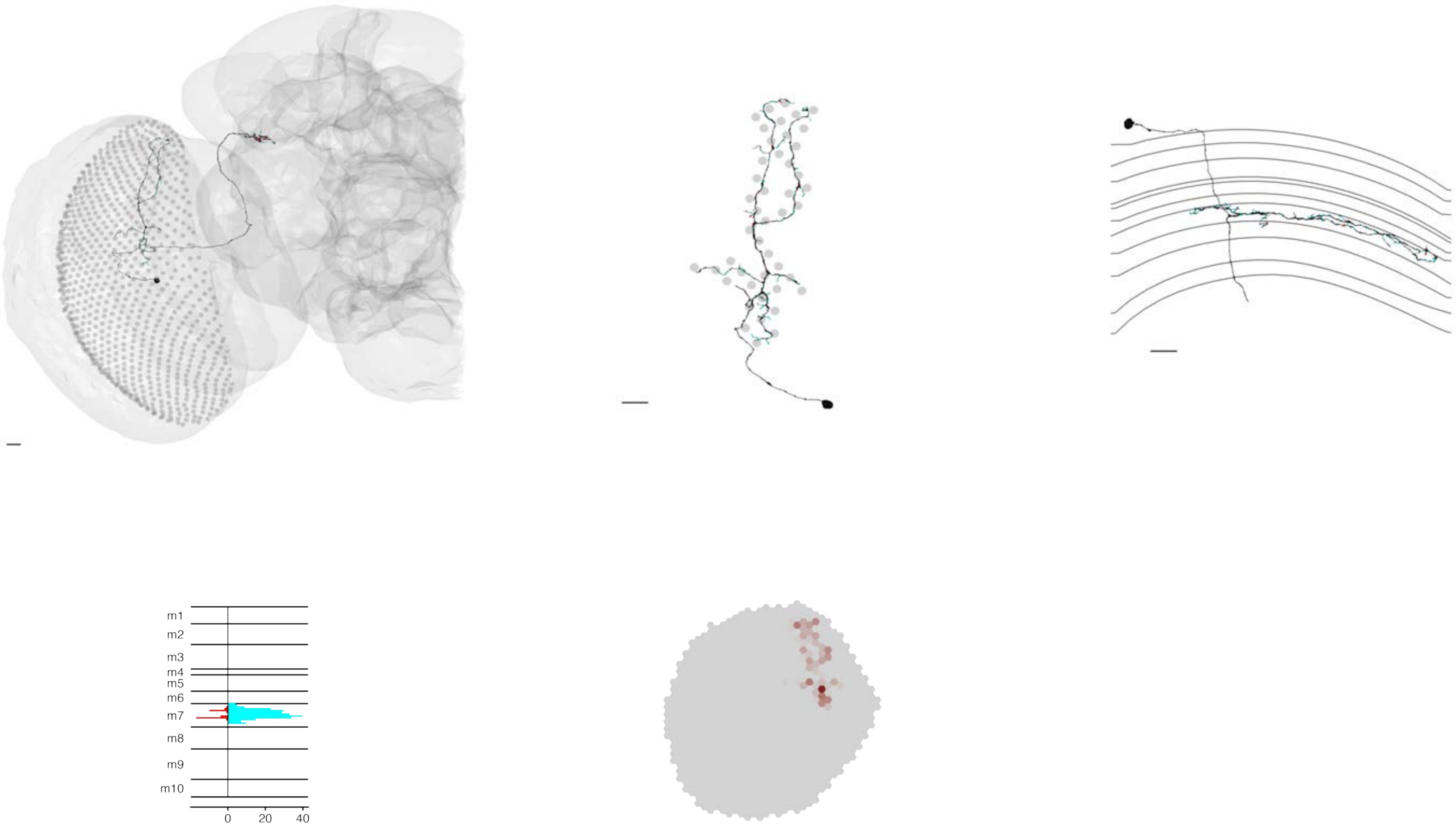

### MeTu2b_720575940609246276

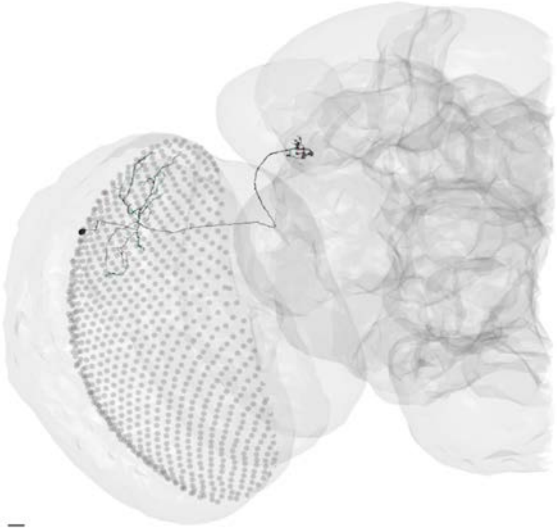

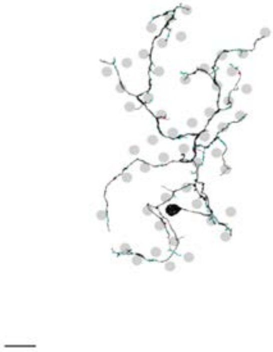

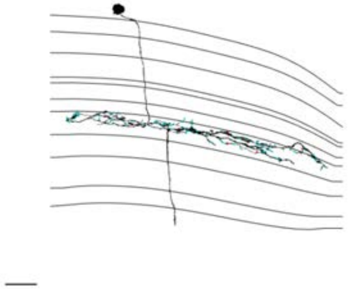

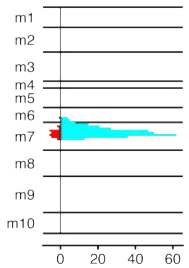


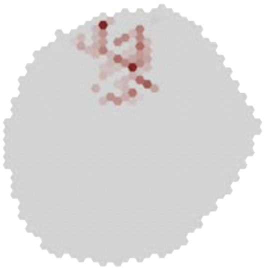

### MeTu2b_720575940614559554

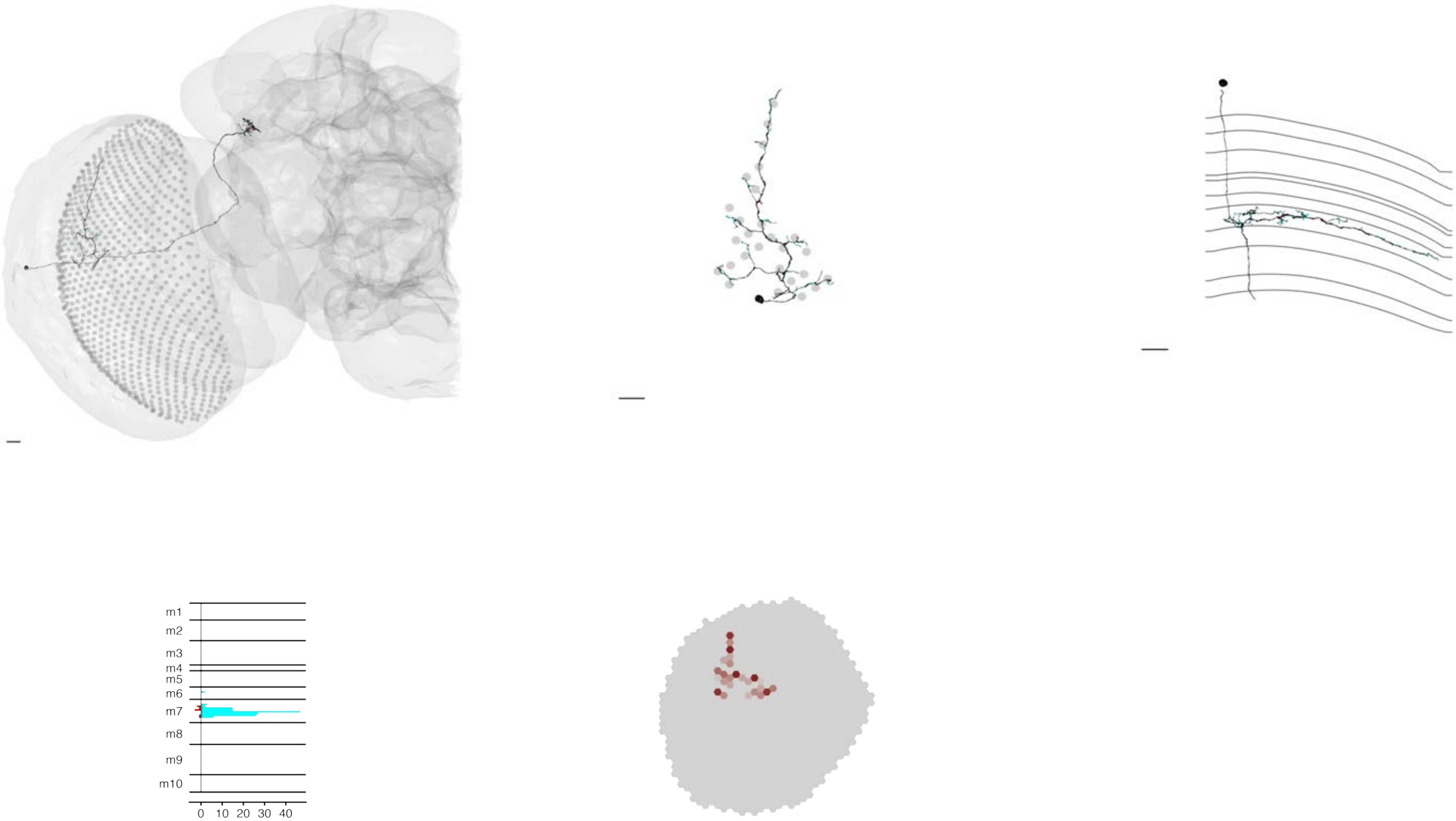

### MeTu2b_720575940618639105

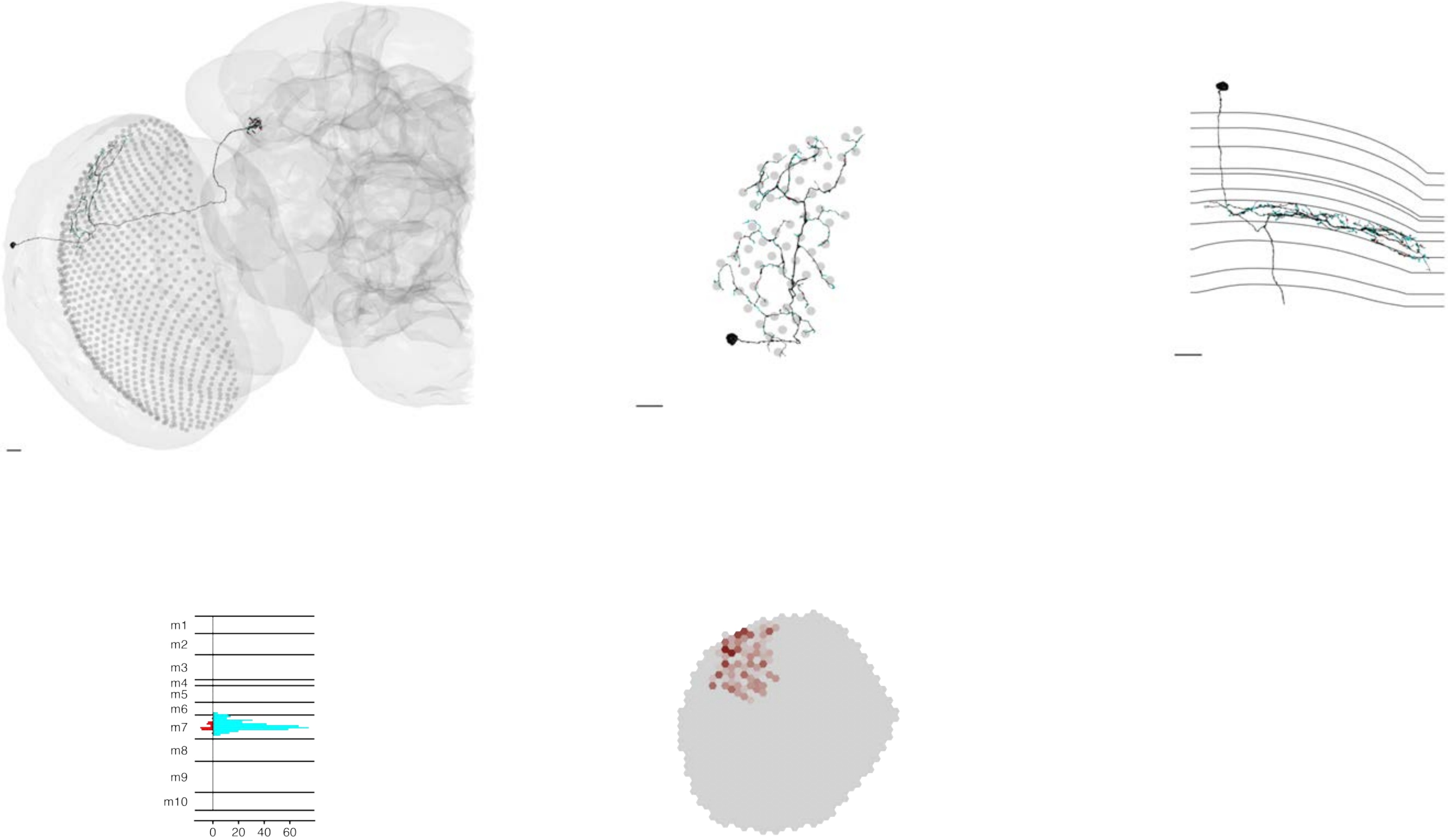

### MeTu2b_720575940619137600

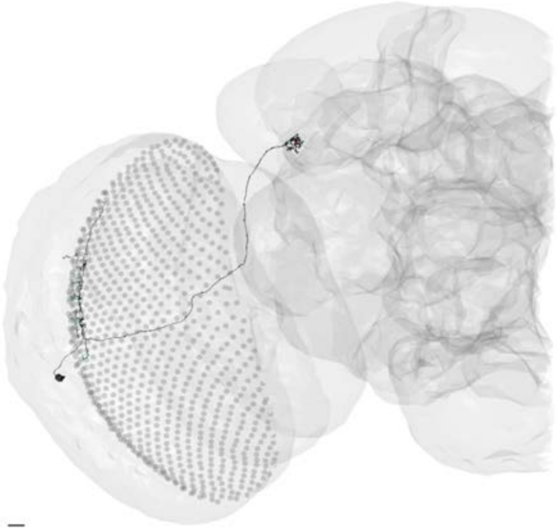

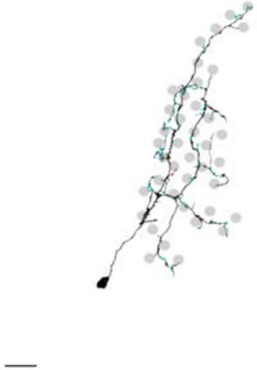

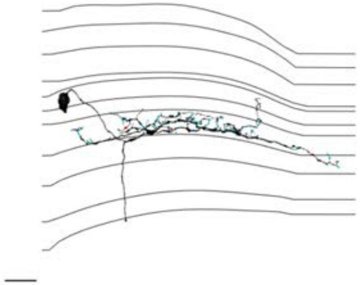

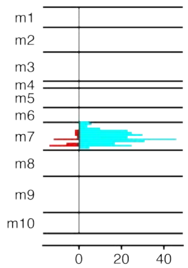


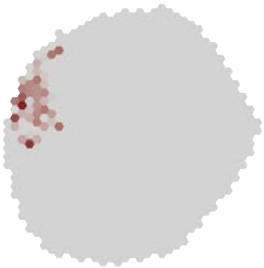

### MeTu2b_720575940619754166

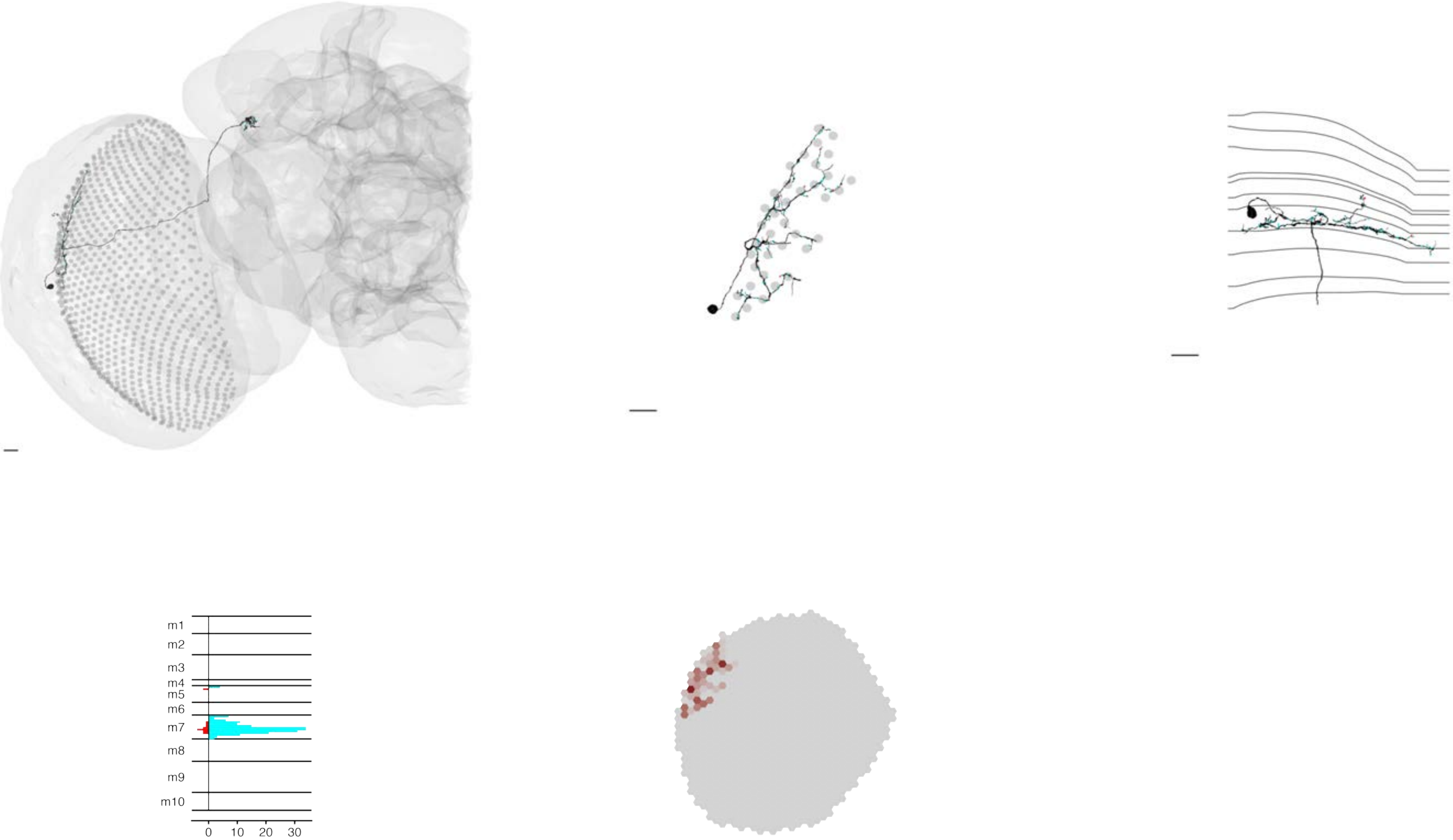

MeTu2b_720575940621062126

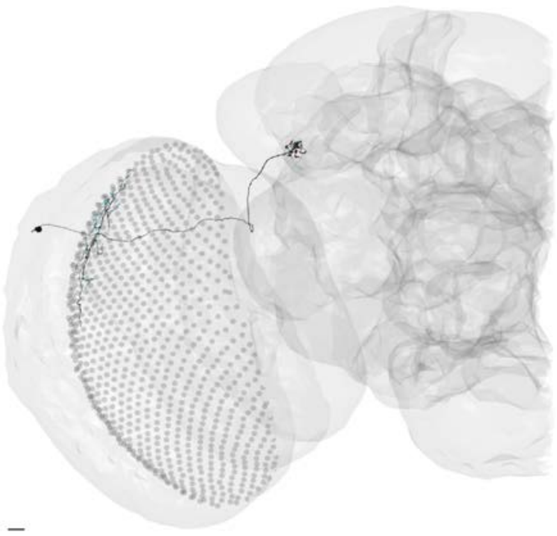

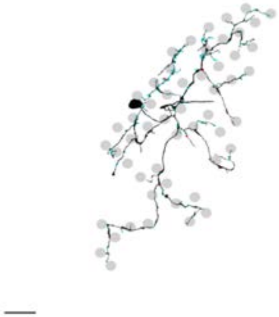

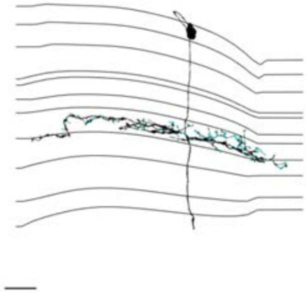

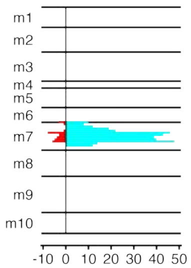


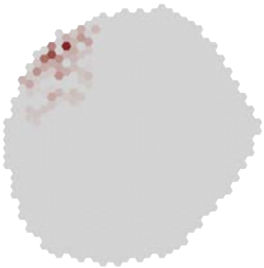

### MeTu2b_720575940625796622

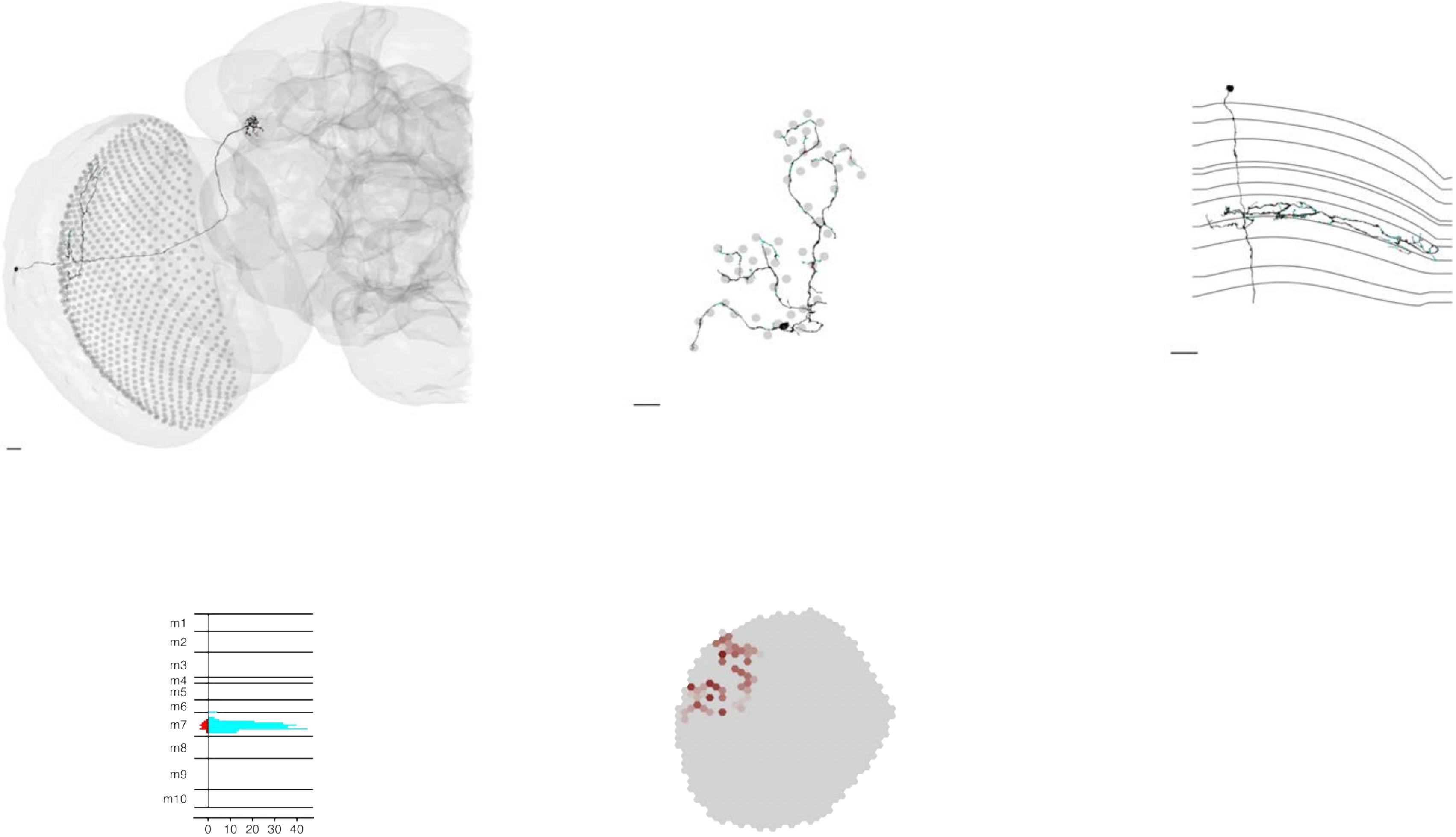

### MeTu2b_720575940627643633

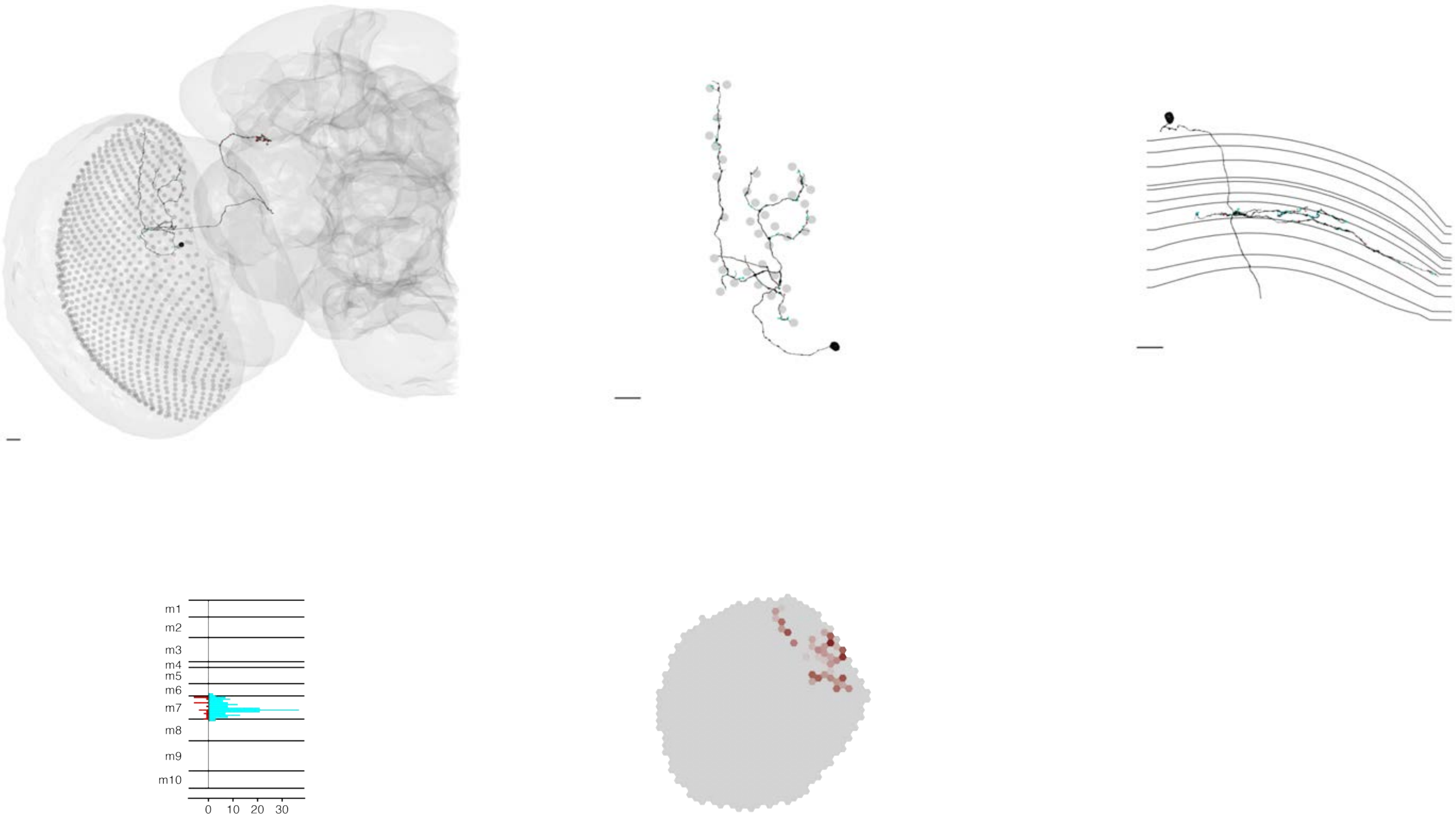

### MeTu2b_720575940628784762

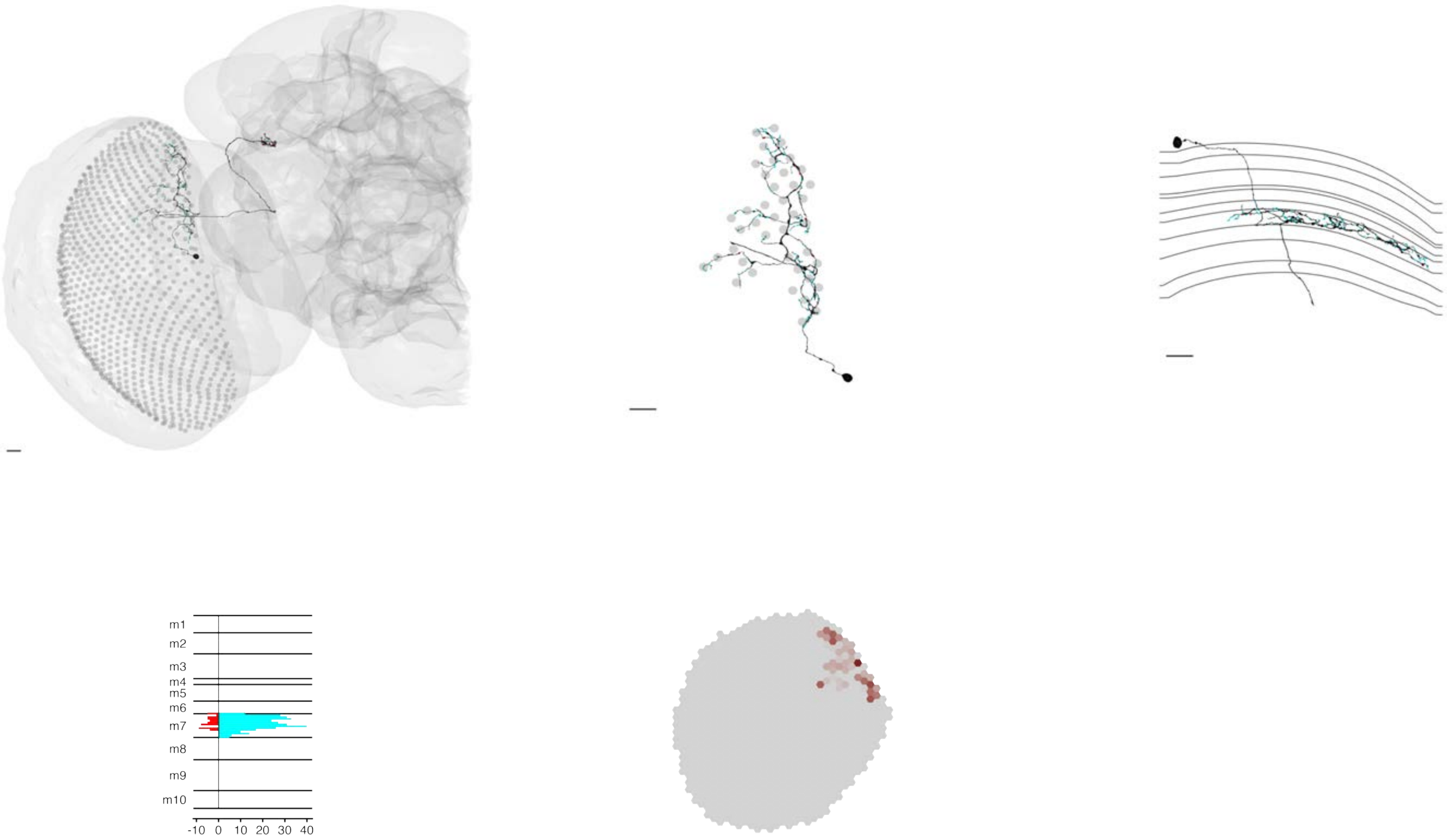

### MeTu2b_720575940632461196

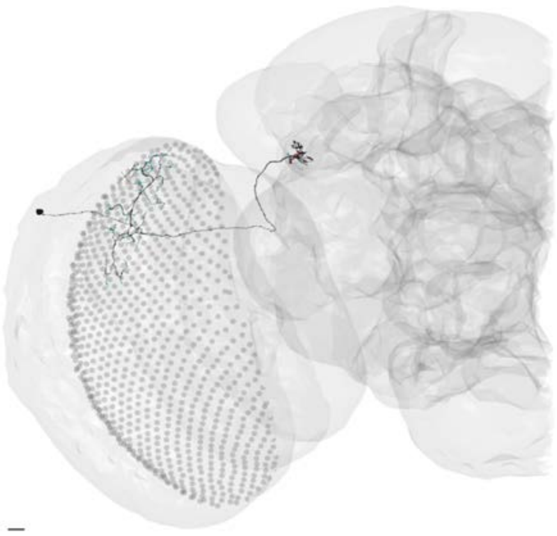

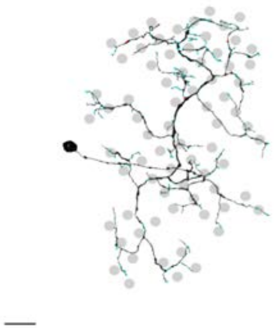

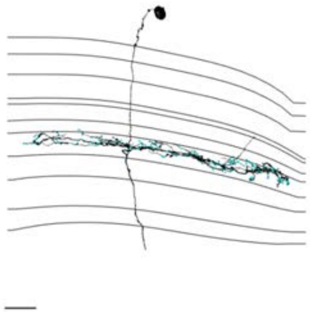

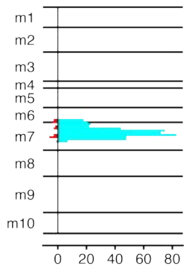


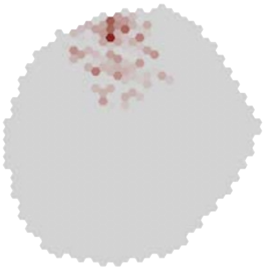

### MeTu2b_720575940637676376

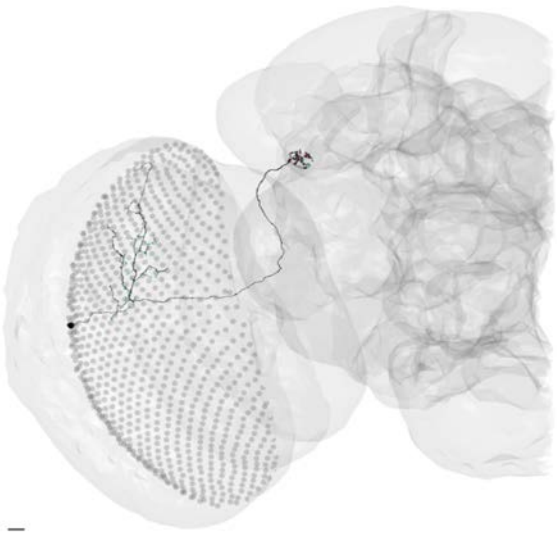

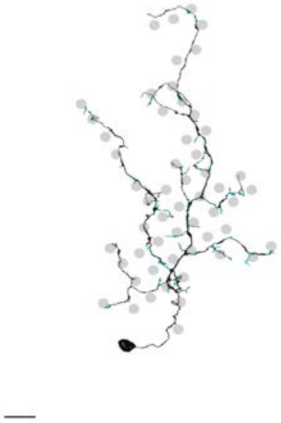

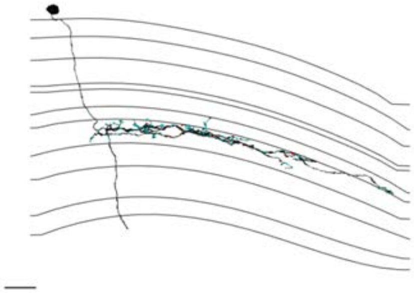

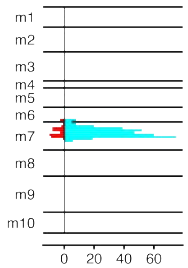


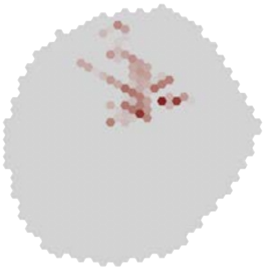

### MeTu2b_720575940643799332

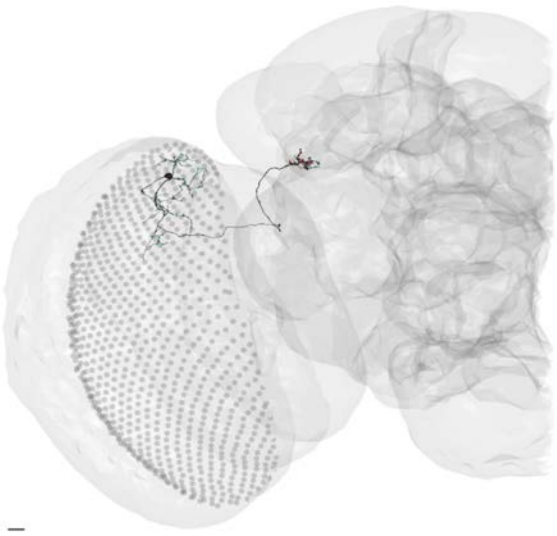

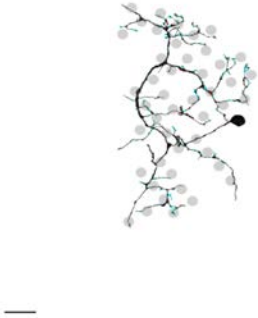

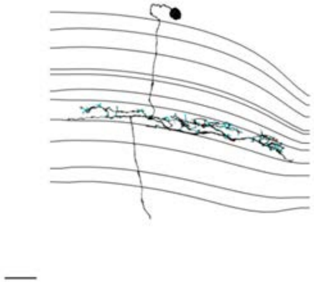

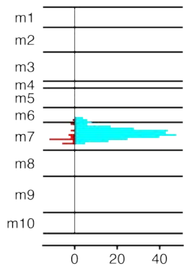


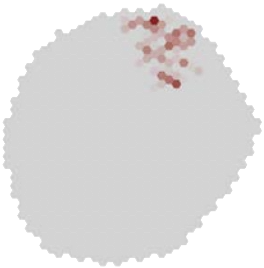

### MeTu2b_720575940660579201

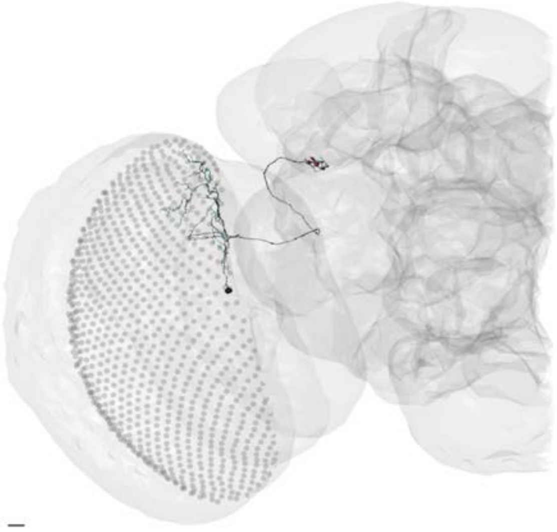

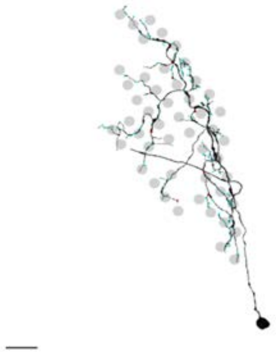

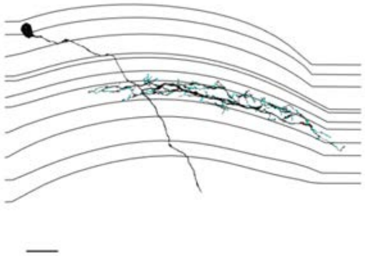

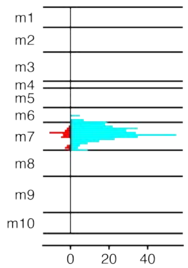


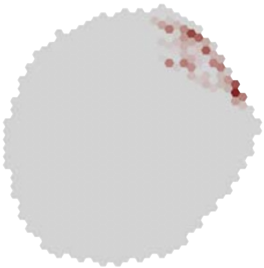

### MeTu3a_720575940612646883

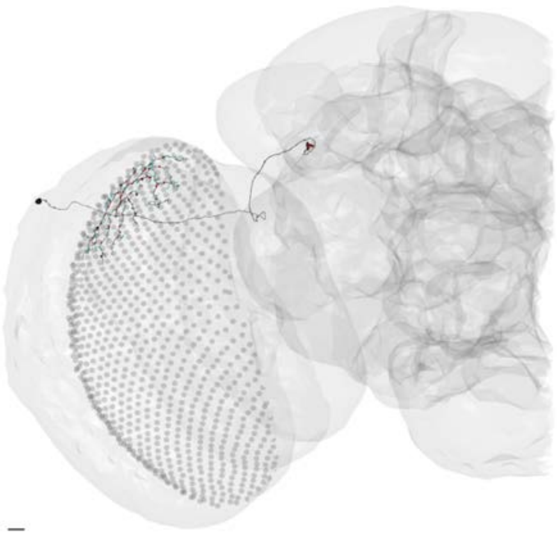

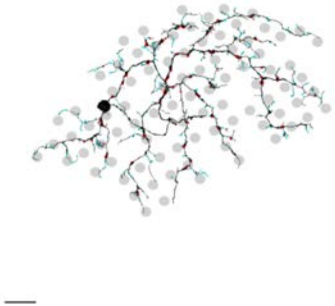

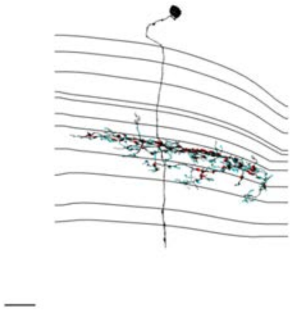

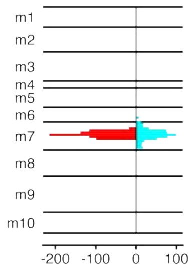


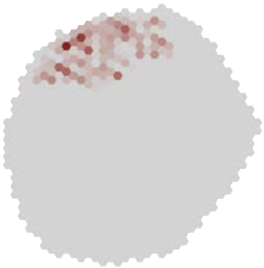

### MeTu3a_720575940613144086

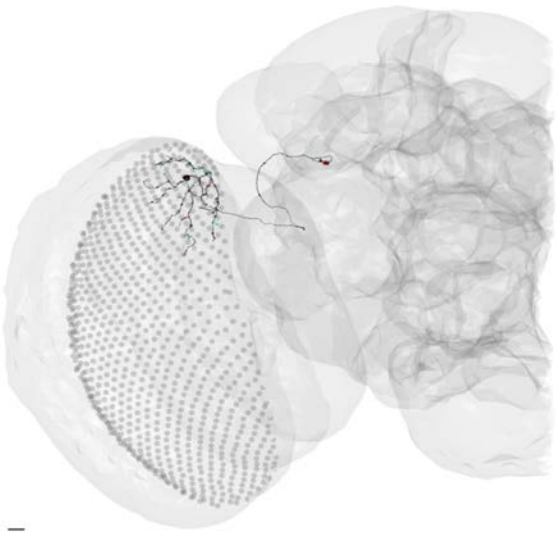

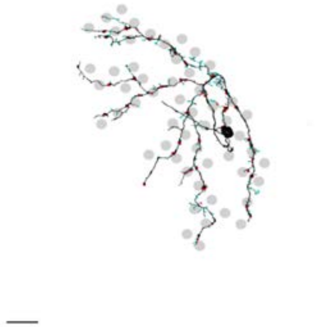

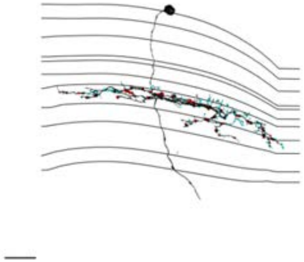

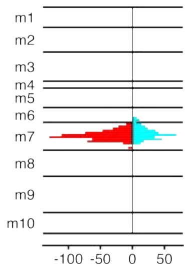


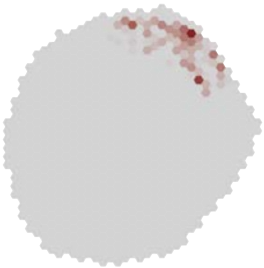

### MeTu3a_720575940613218418

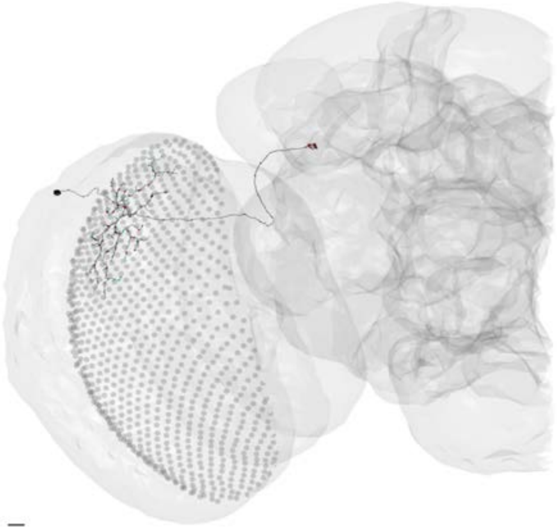

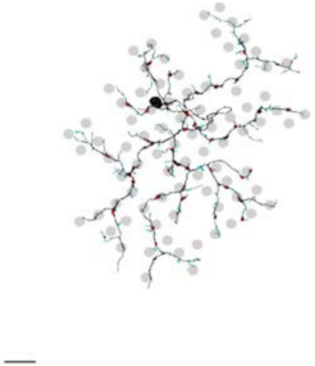

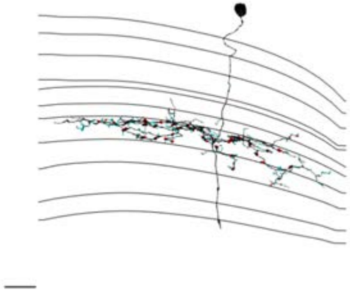

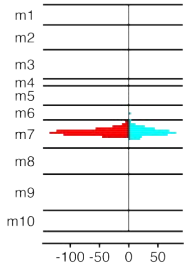


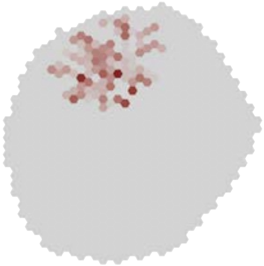

### MeTu3a_720575940615406698

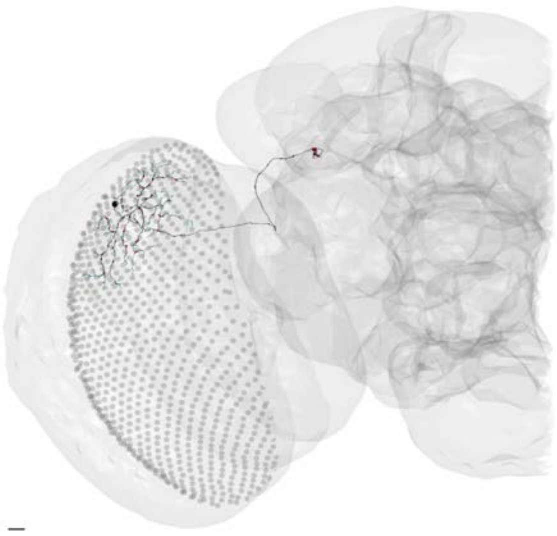

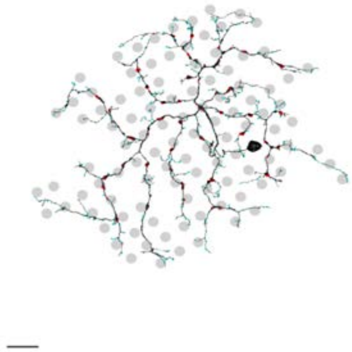

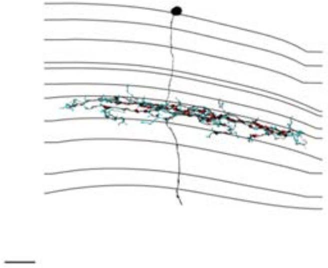

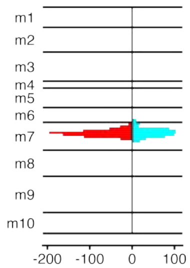


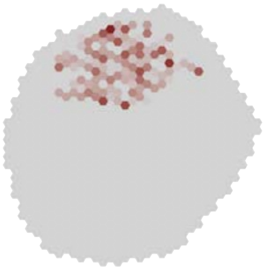

### MeTu3a_720575940615467170

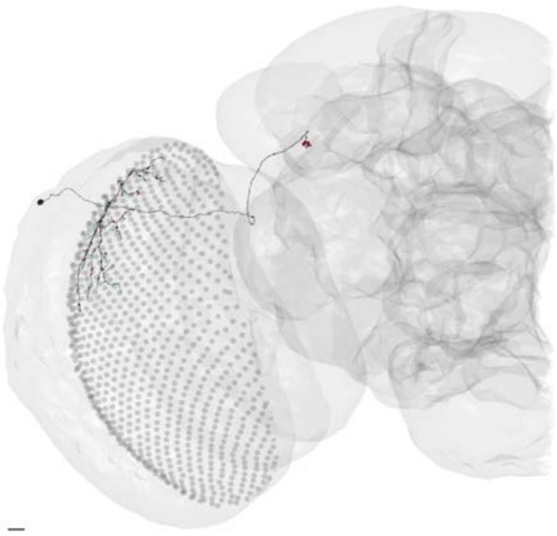

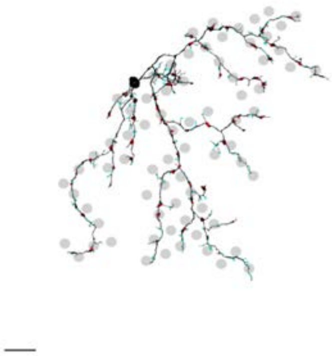

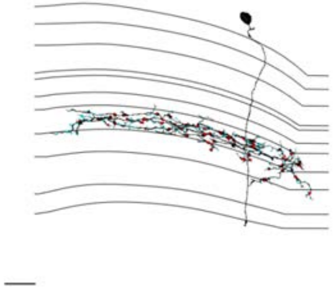

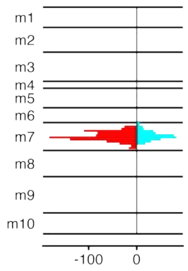


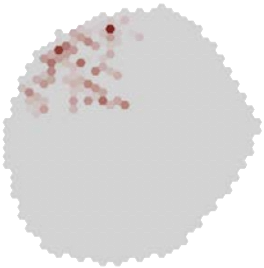

MeTu3a_720575940616499089

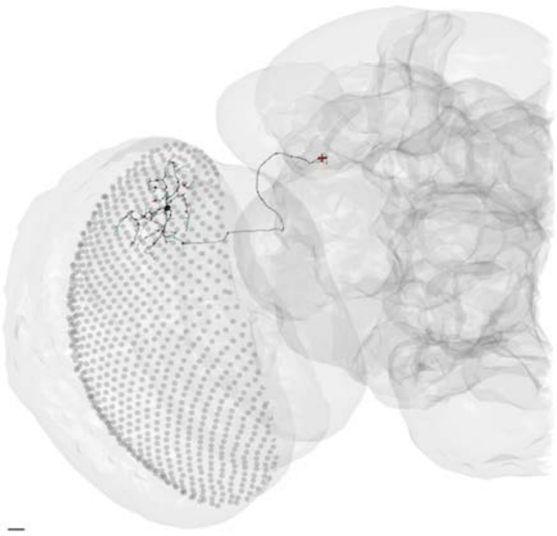

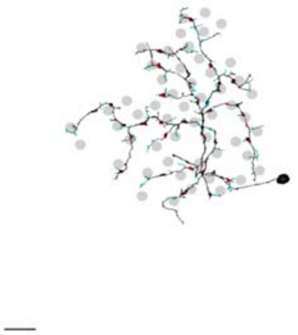

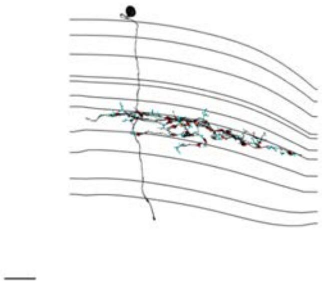

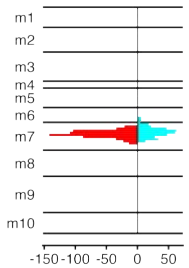


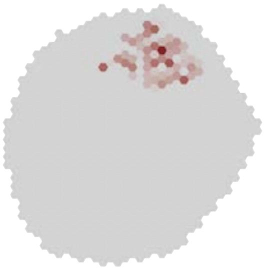

### MeTu3a_720575940619861918

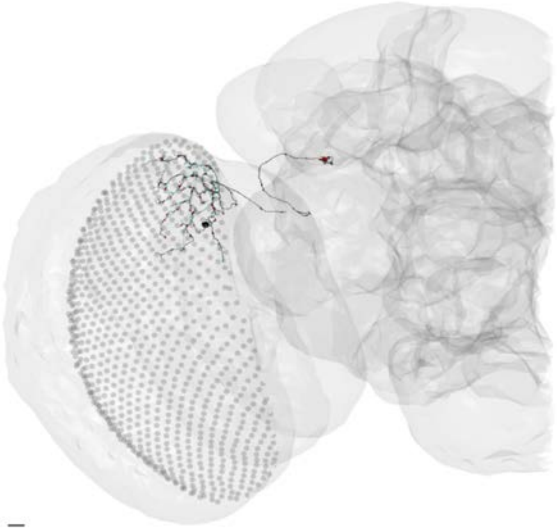

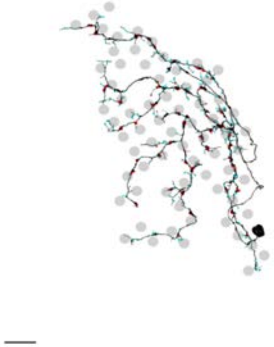

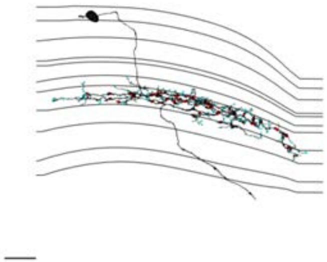

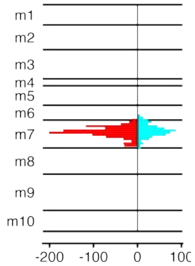


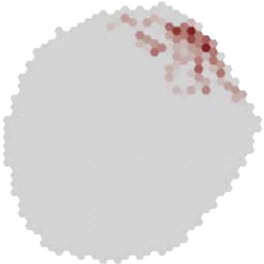

### MeTu3a_720575940620275565

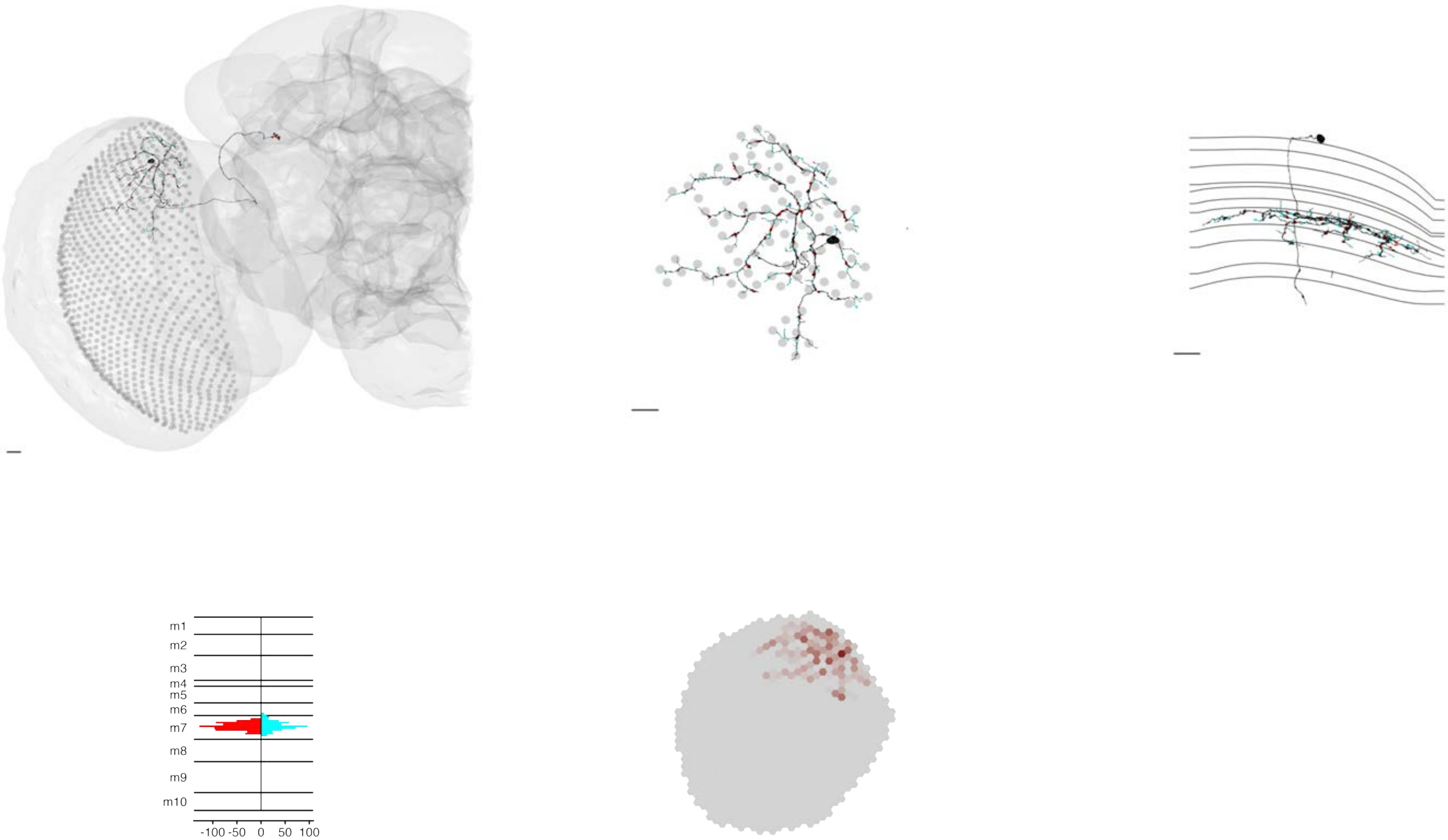

MeTu3a_720575940621036123

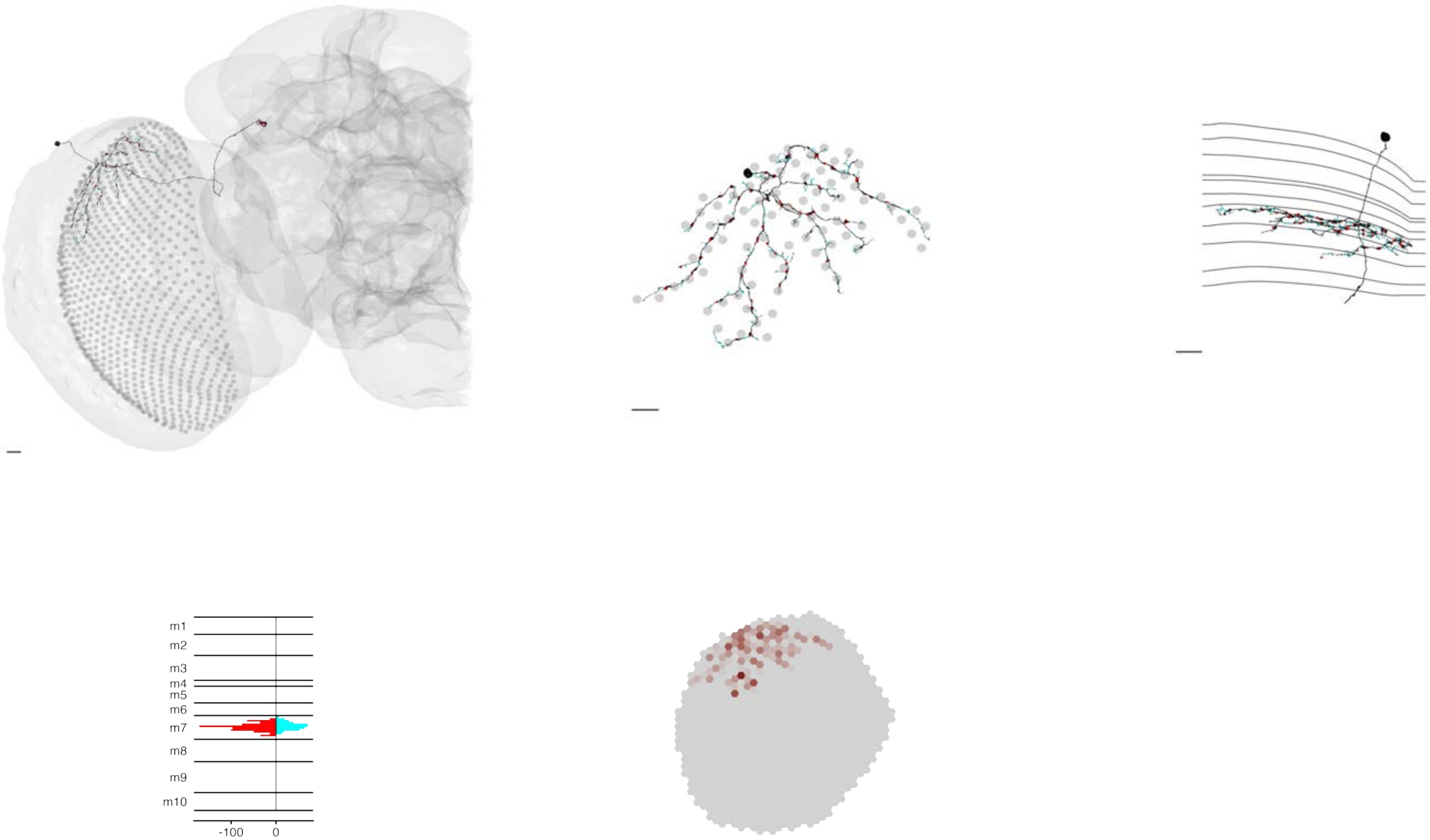

### MeTu3a_720575940623440363

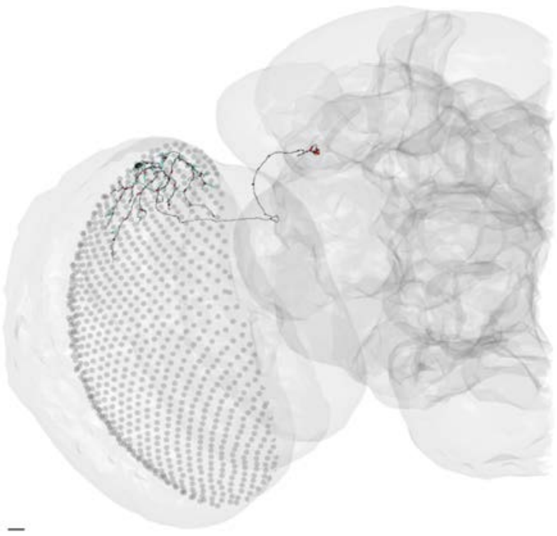

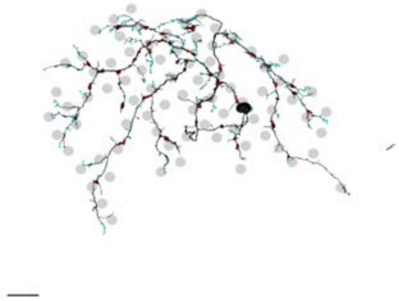

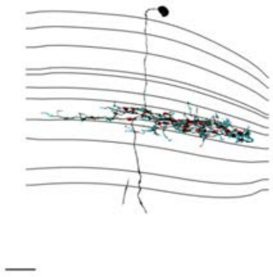

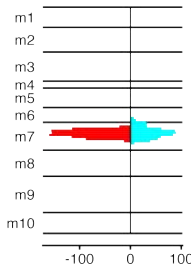


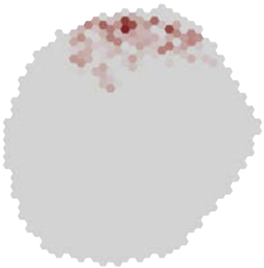

### MeTu3a_720575940624799467

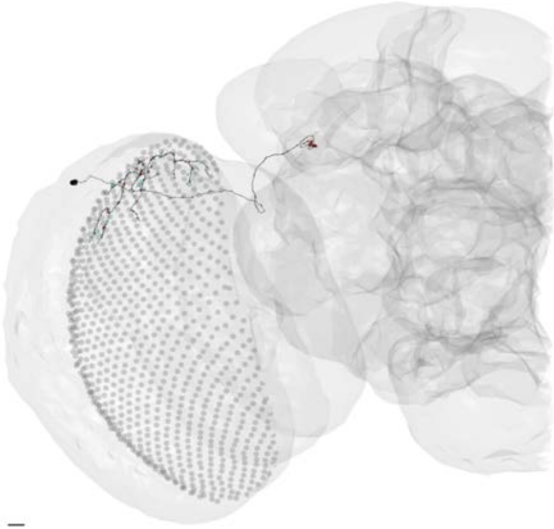

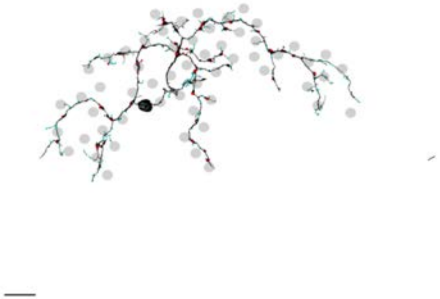

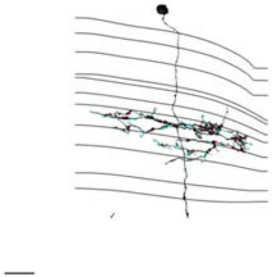

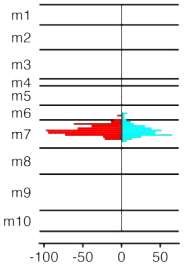


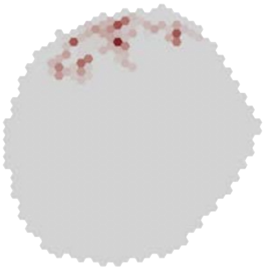

### MeTu3a_720575940626707518

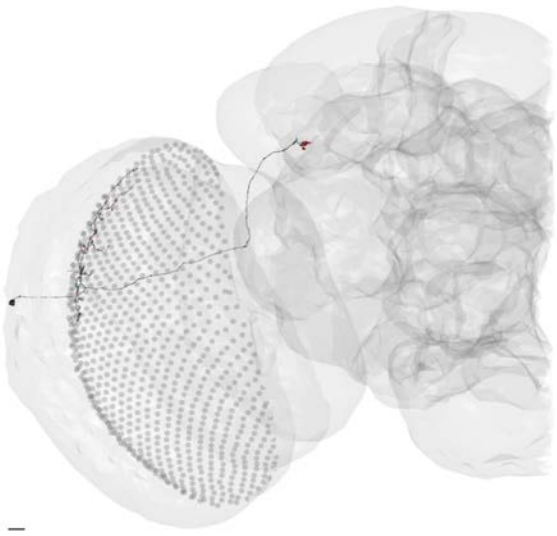

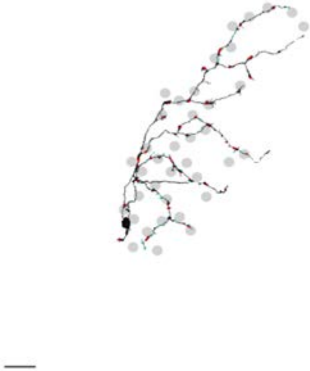

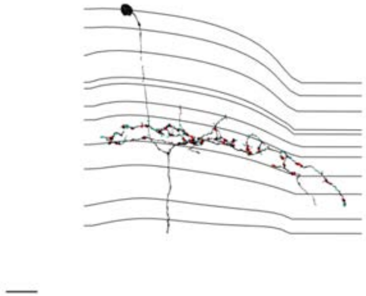

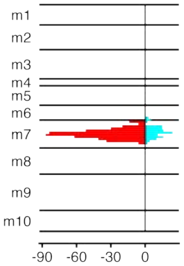


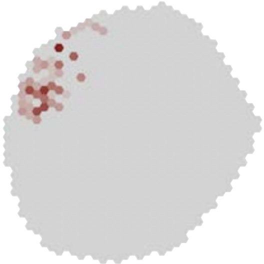

### MeTu3a_720575940628649290

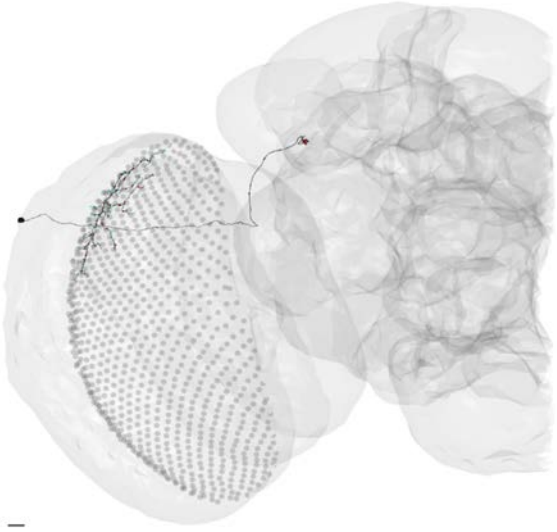

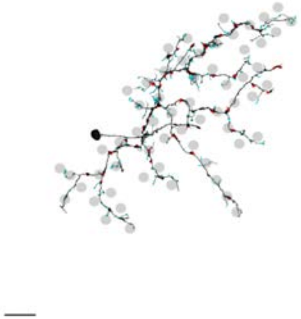

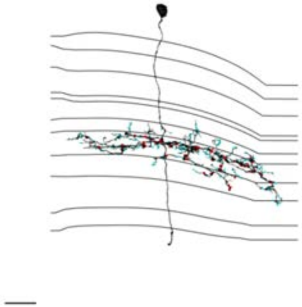

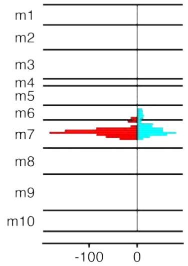


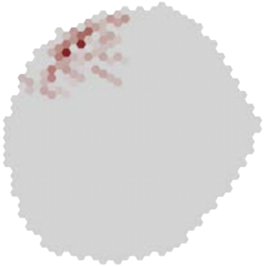

### MeTu3a_720575940629357135

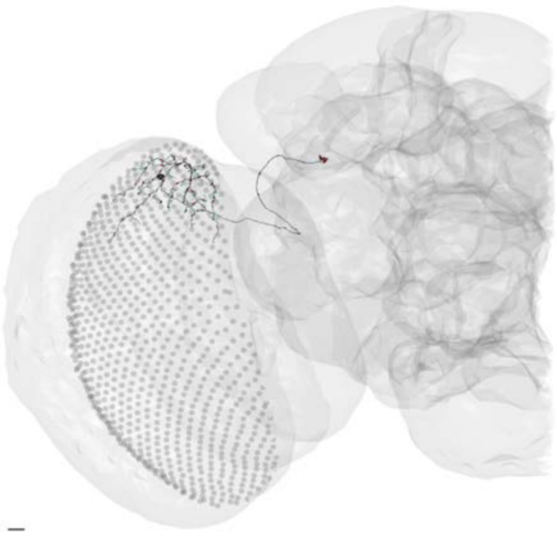

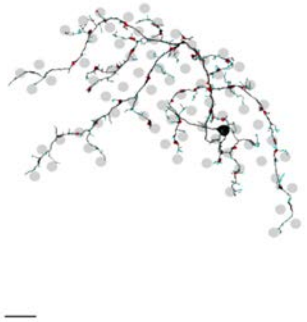

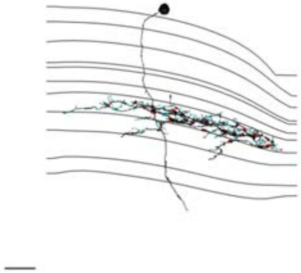

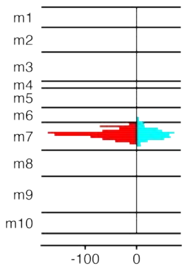


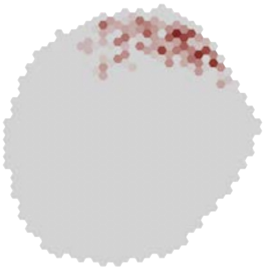

### MeTu3a_720575940629386496

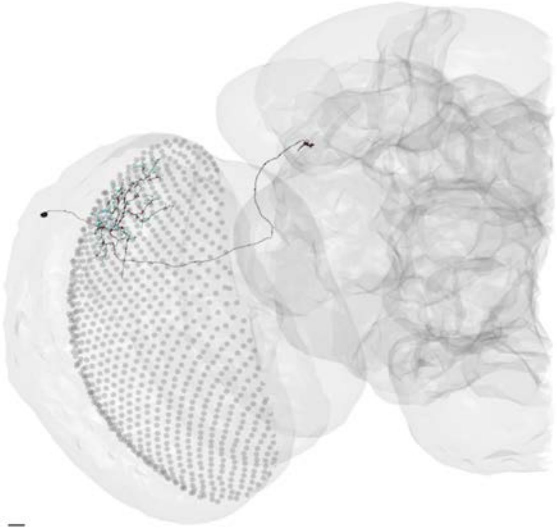

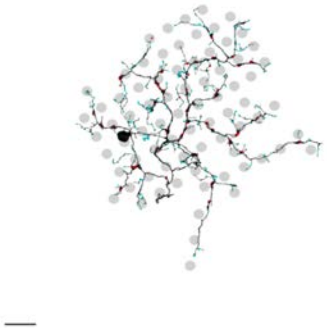

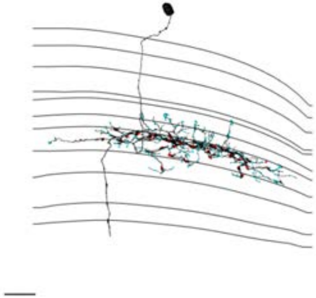

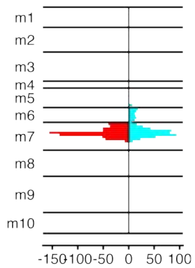


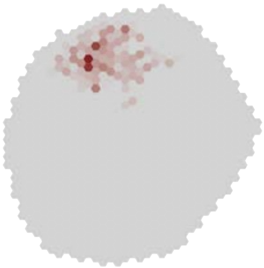

### MeTu3a_720575940631504591

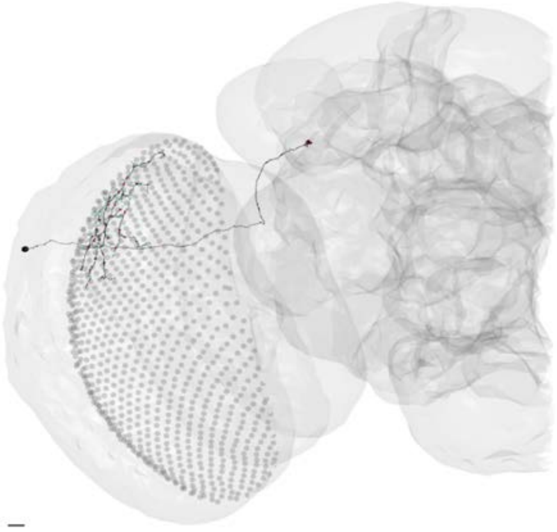

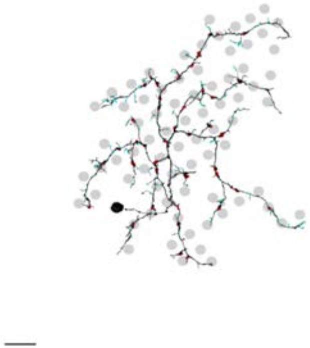

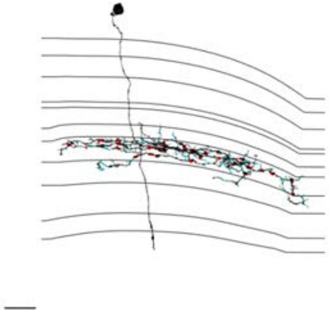

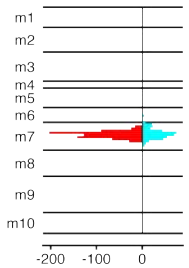


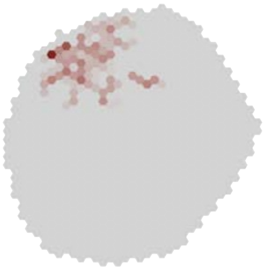

### MeTu3a_720575940631846698

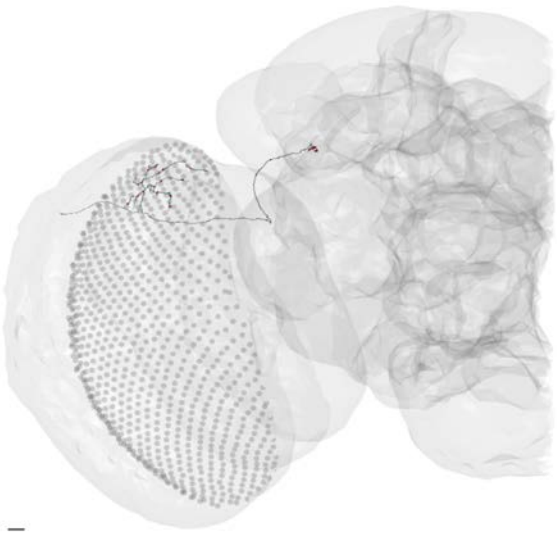

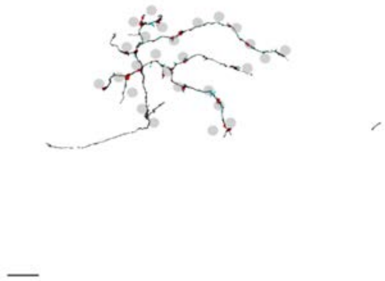

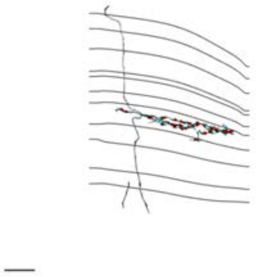

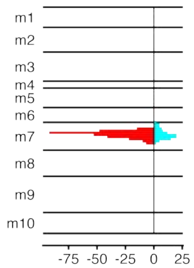


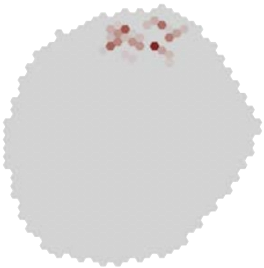

### MeTu3a_720575940633829783

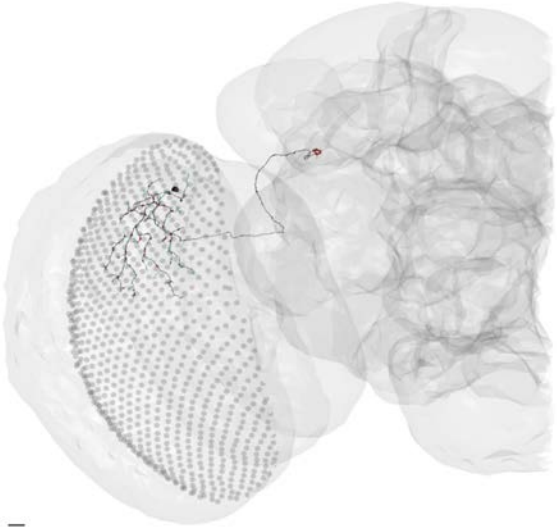

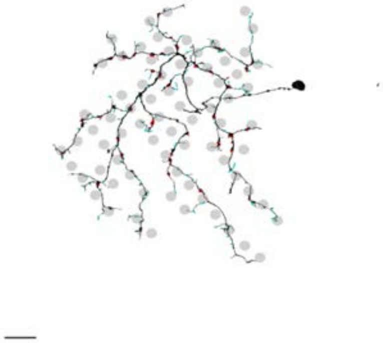

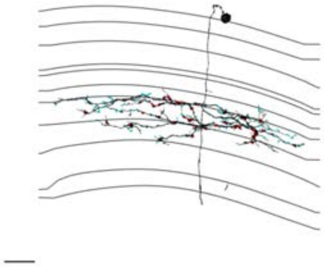

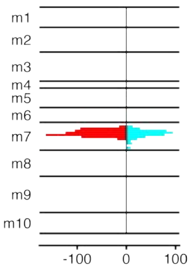


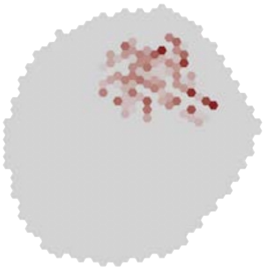

### MeTu3a_720575940655346337

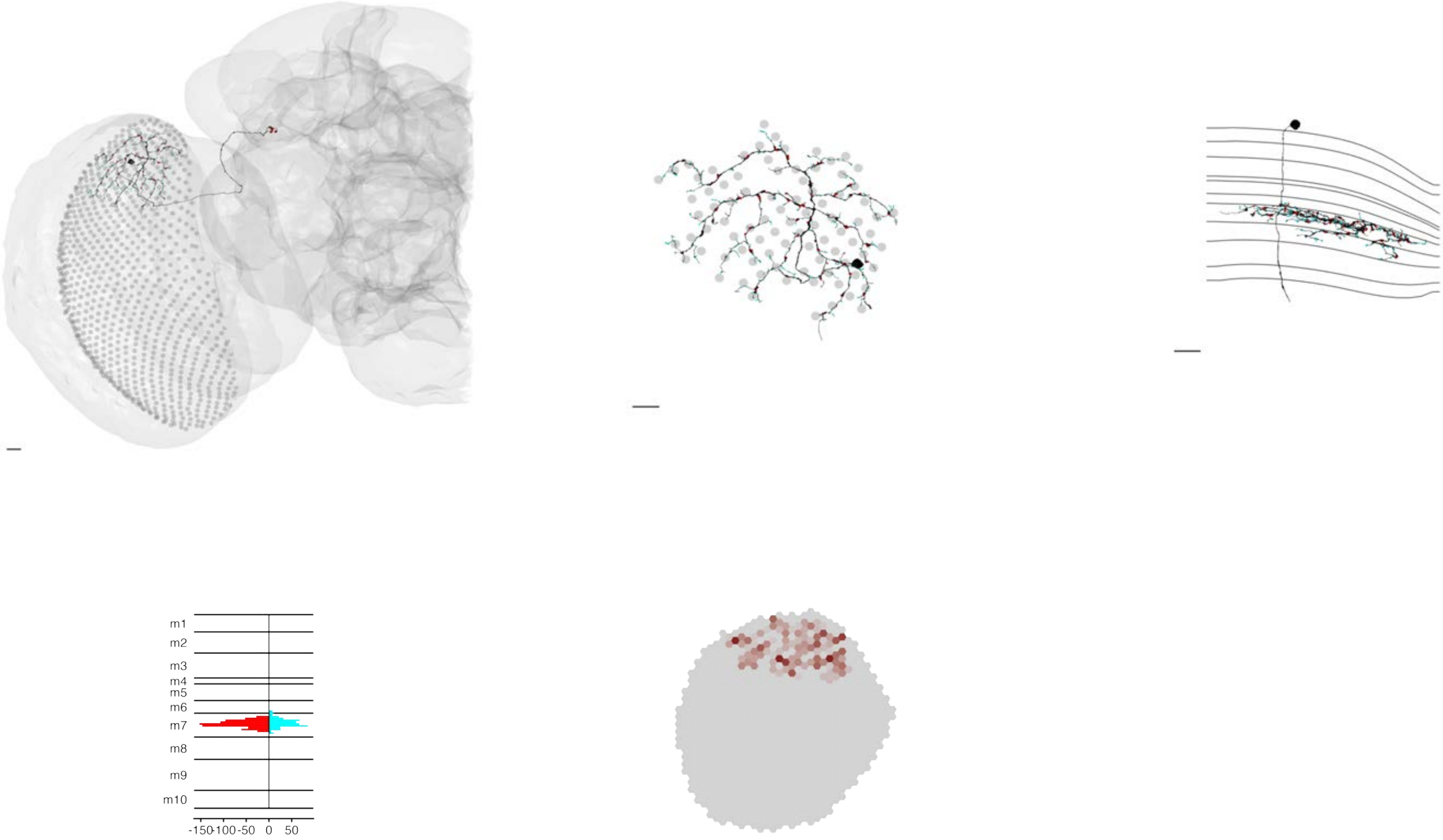

### MeTu3b_720575940604304742

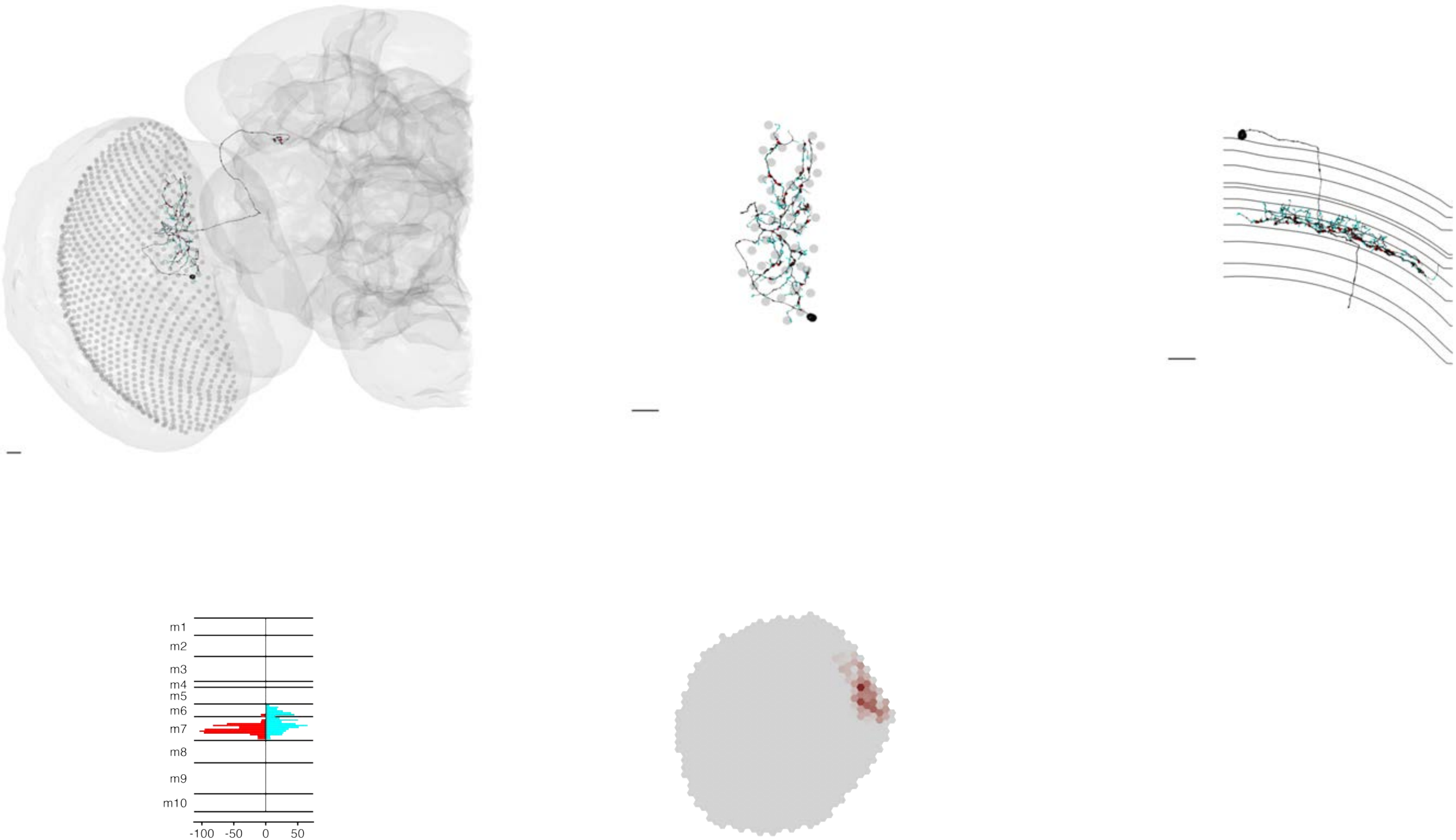

### MeTu3b_720575940606722140

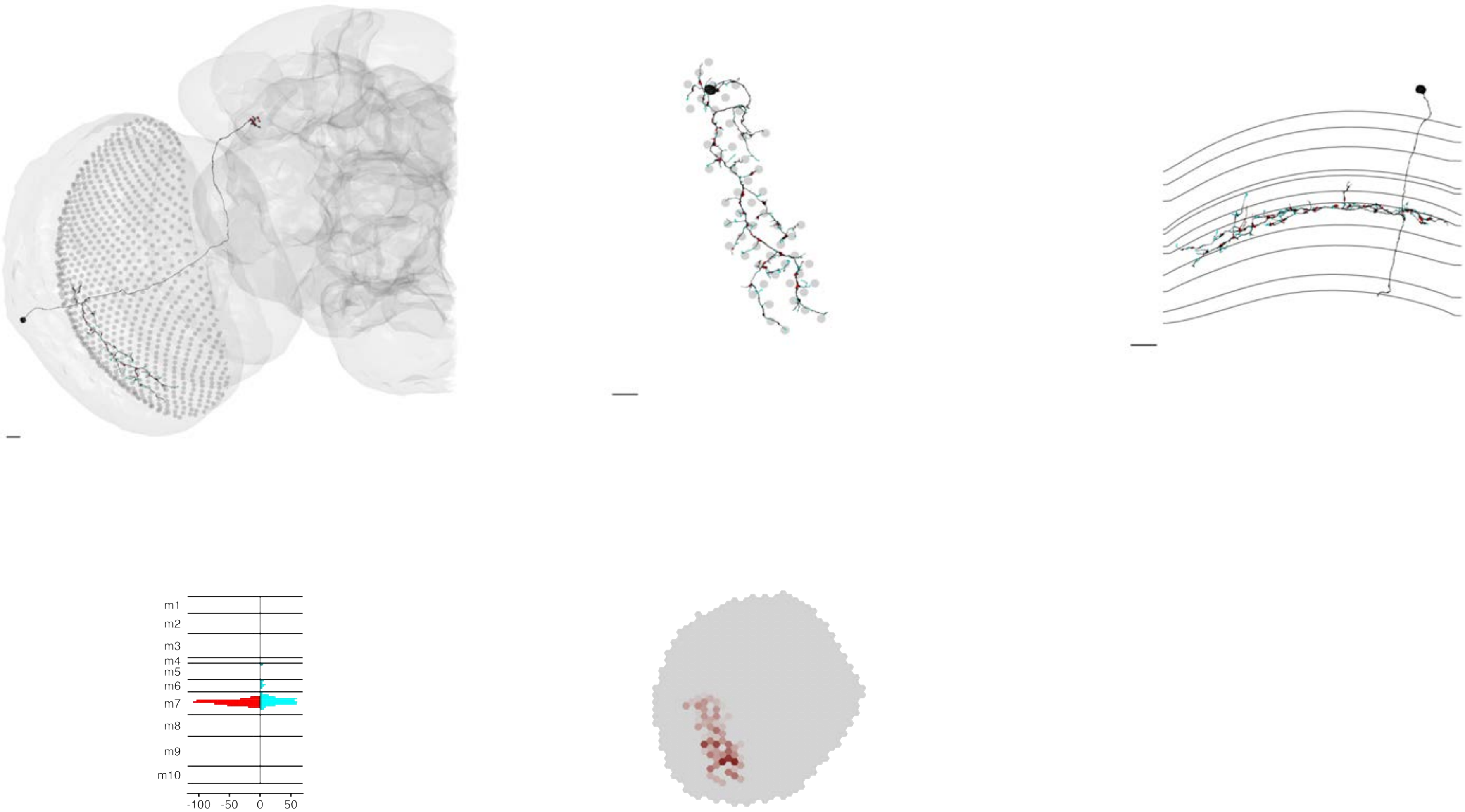

MeTu3b_720575940609304184

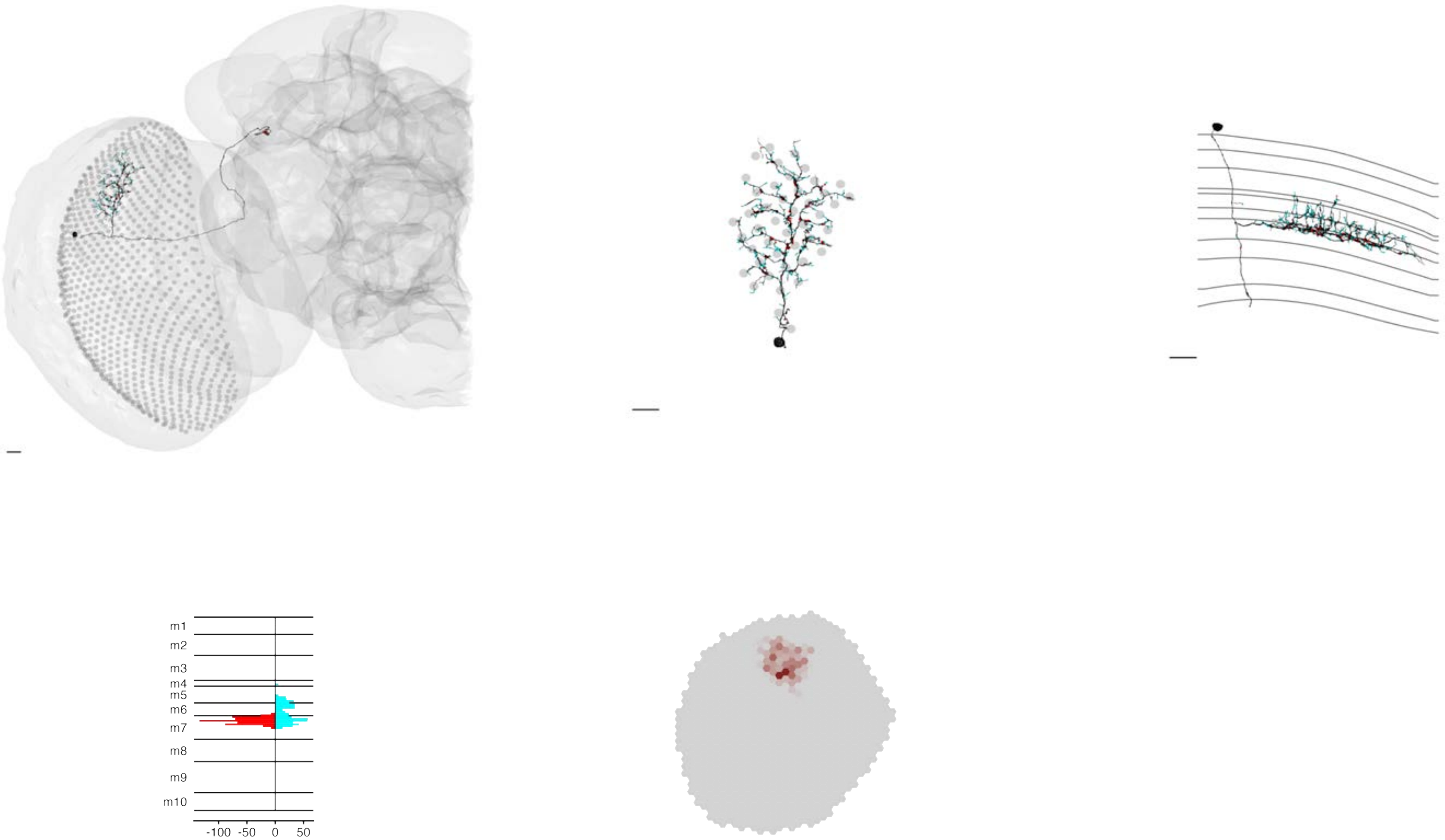

### MeTu3b_720575940611991062

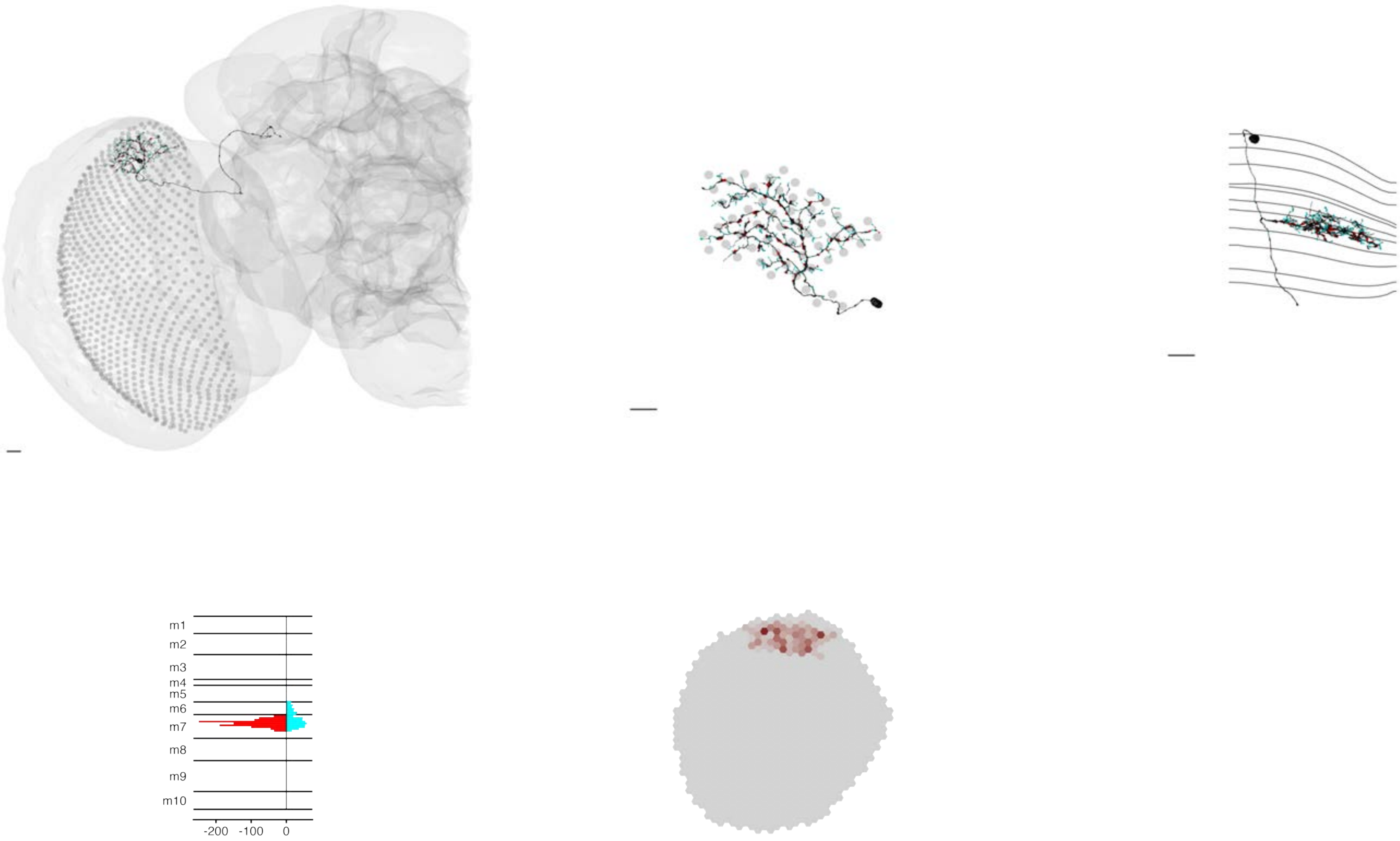

### MeTu3b_720575940612455599

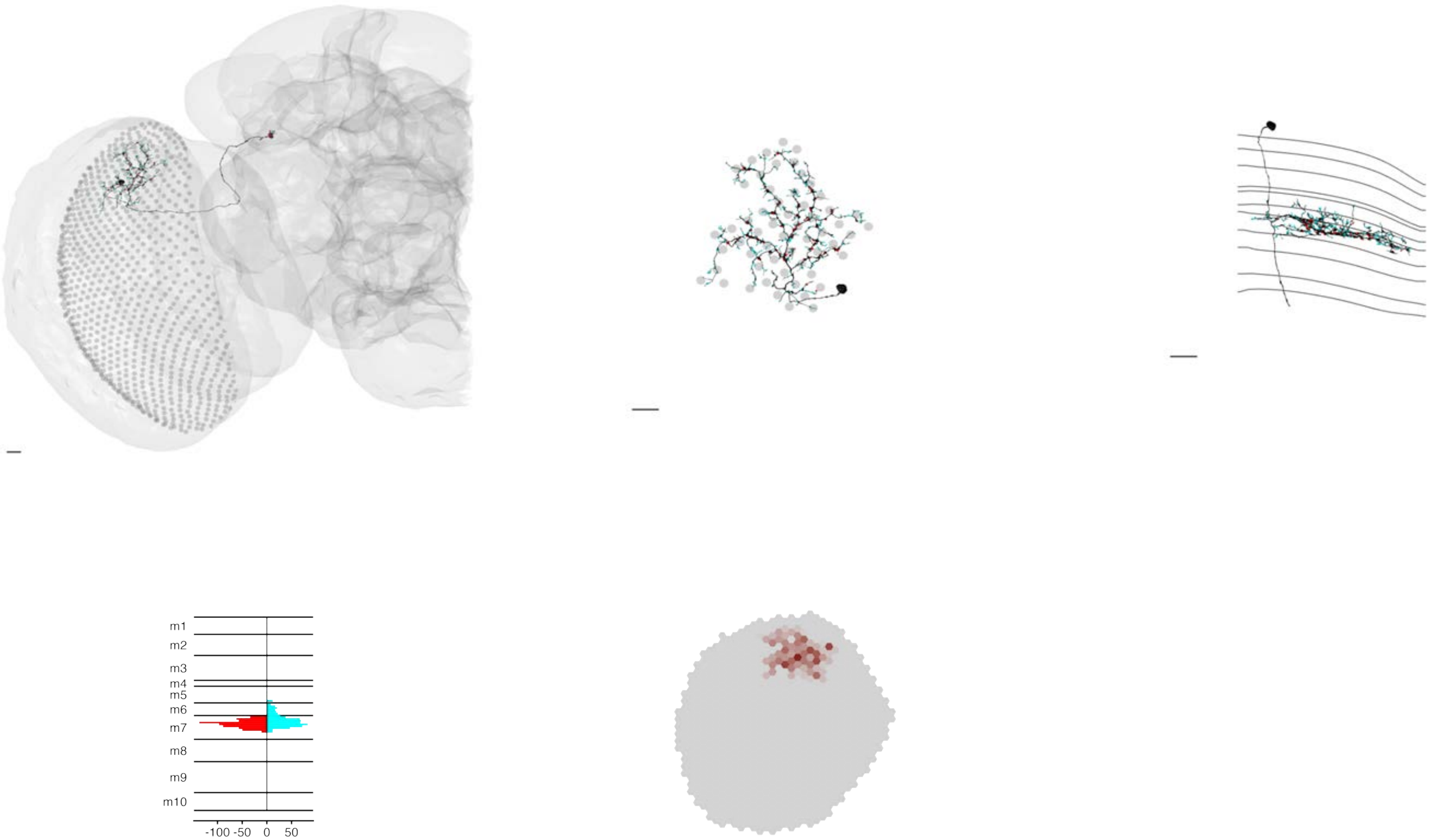

### MeTu3b_720575940614499634

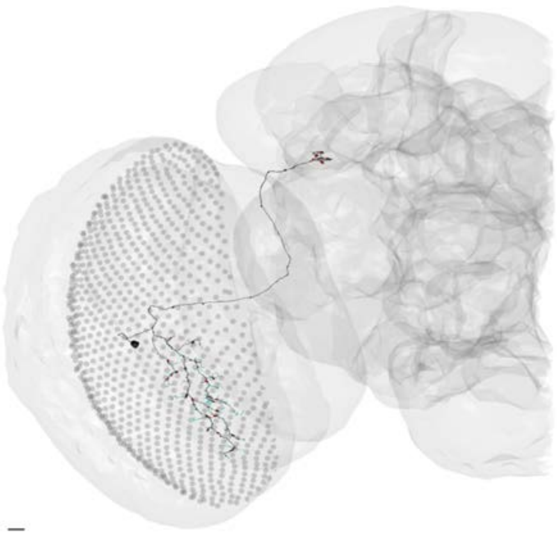

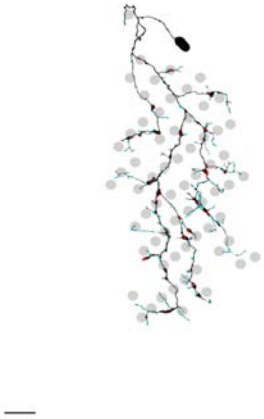

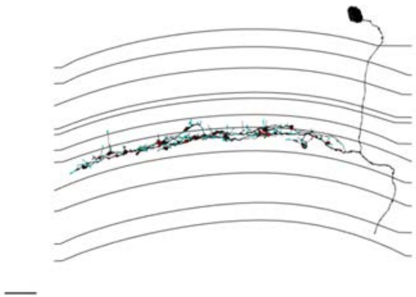

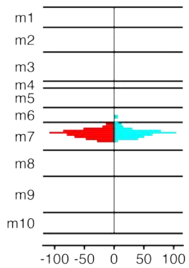


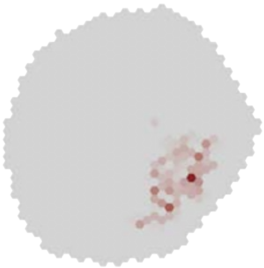

### MeTu3b_720575940614552170

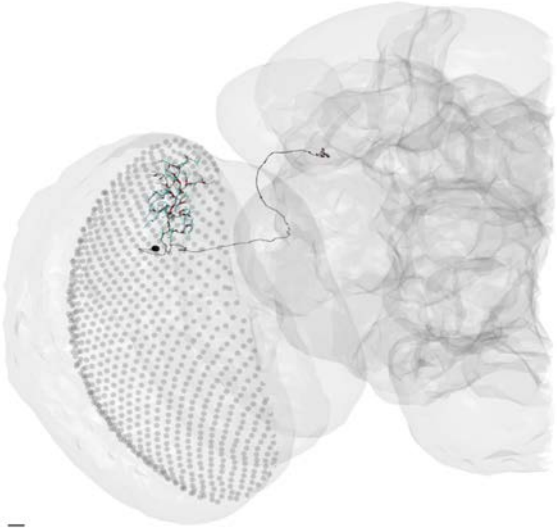

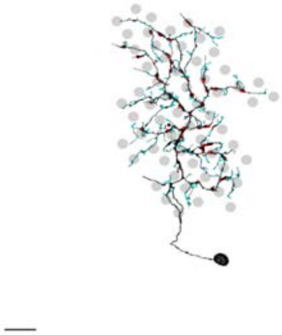

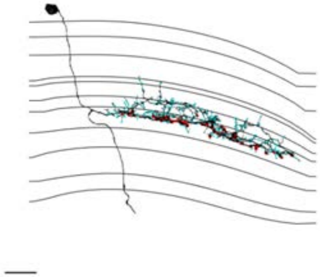

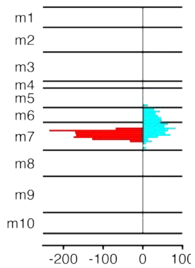


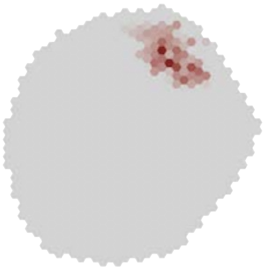

### MeTu3b_720575940615507506

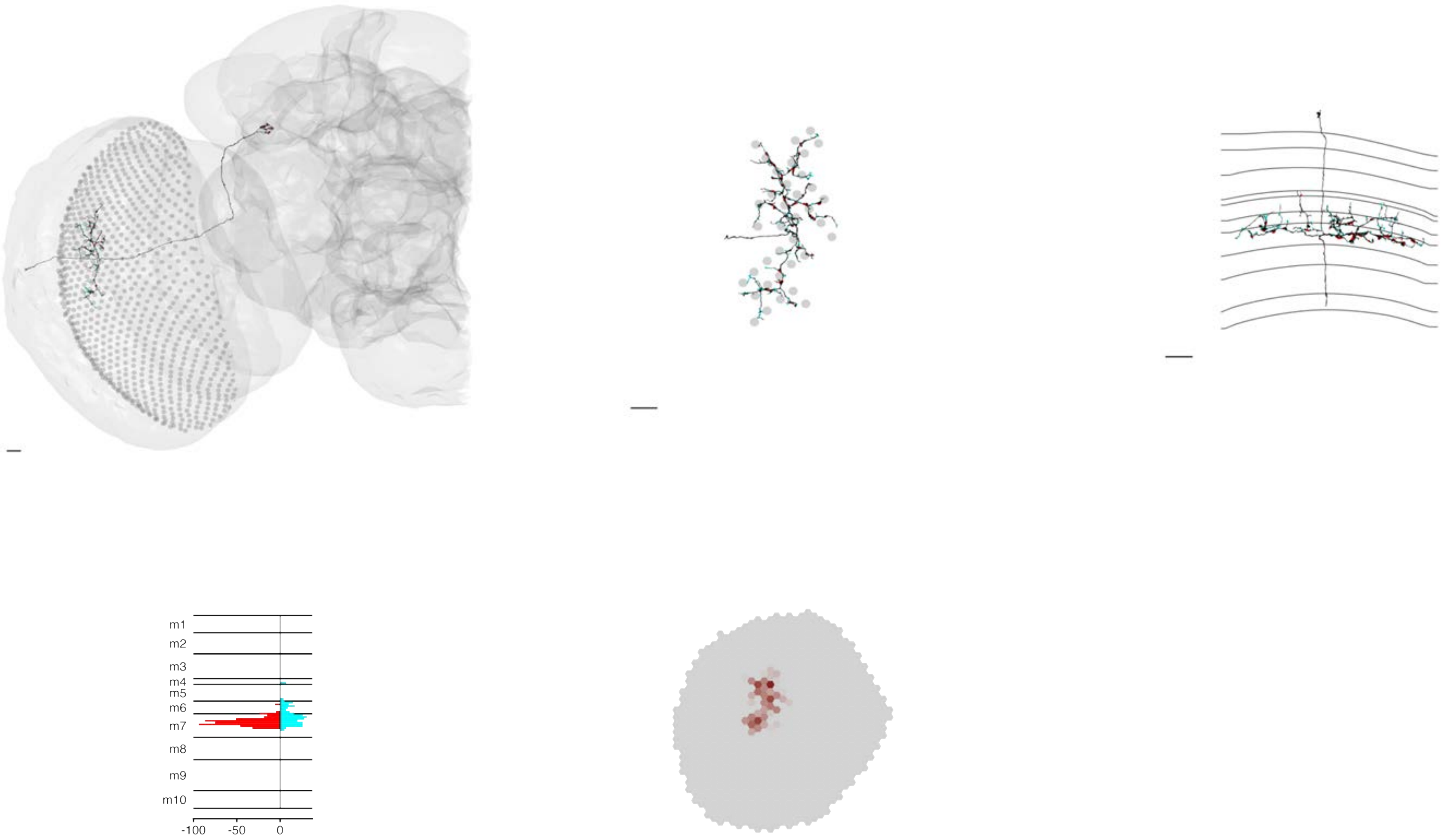

MeTu3b_720575940617793344

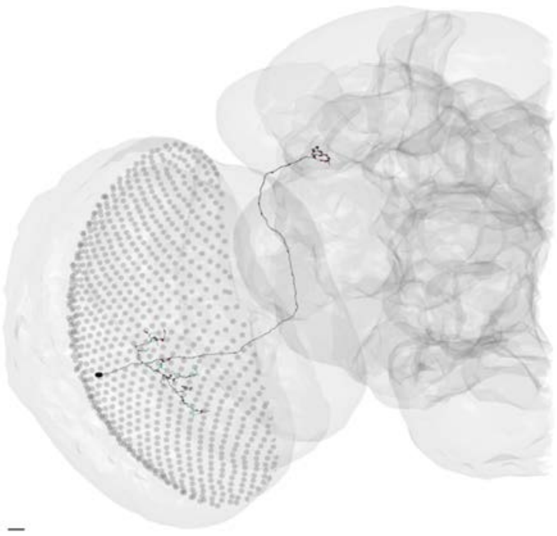

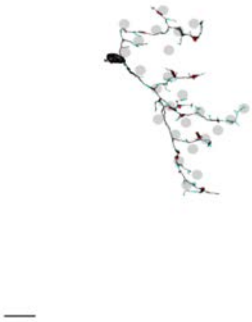

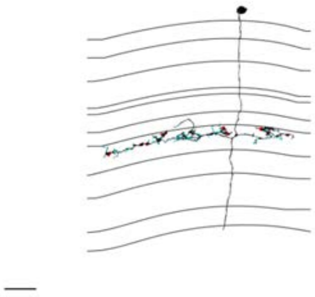

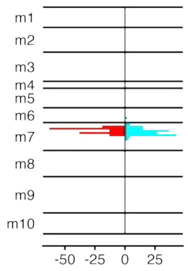


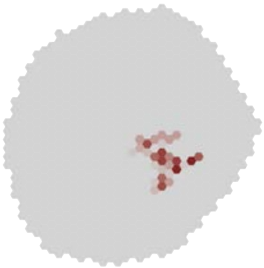

### MeTu3b_720575940618558292

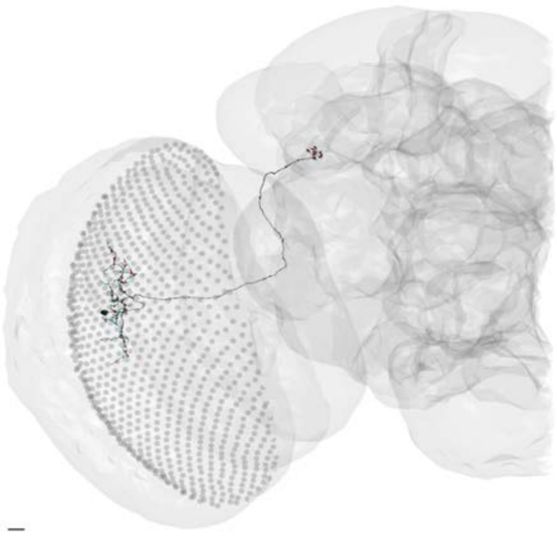

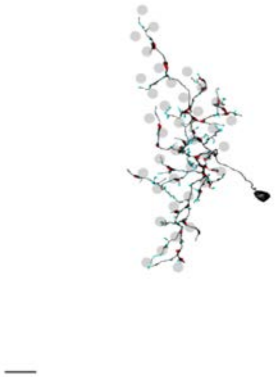

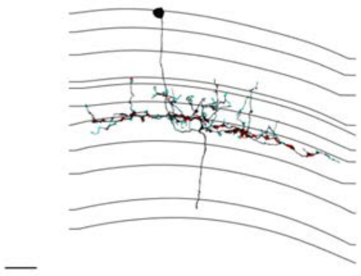

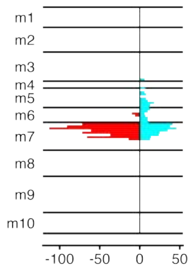


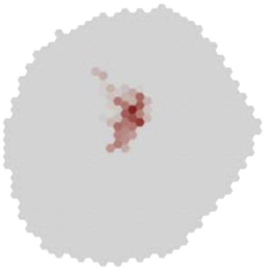

### MeTu3b_720575940620510657

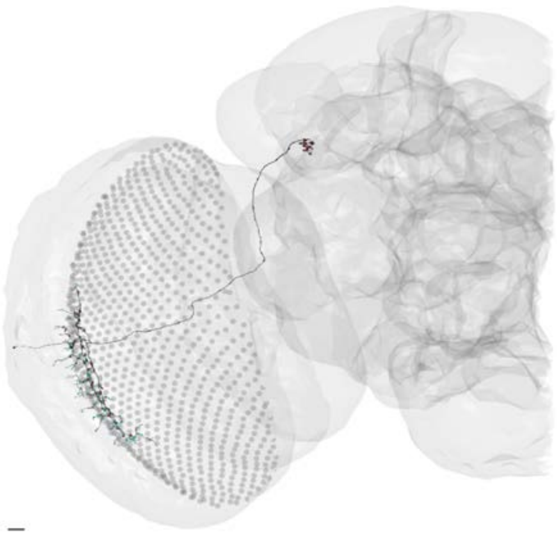

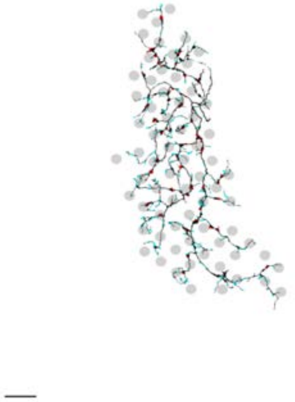

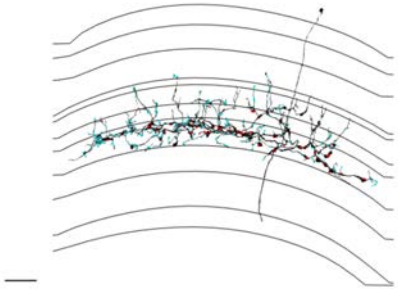

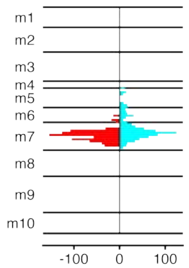


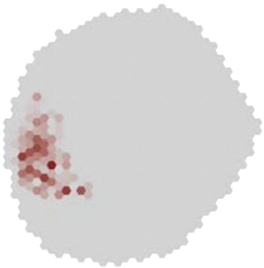

### MeTu3b_720575940620605099

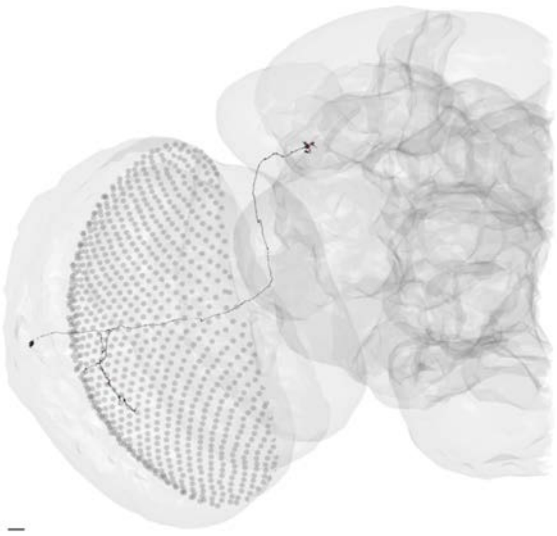

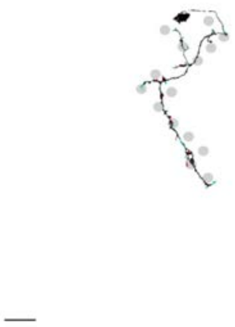

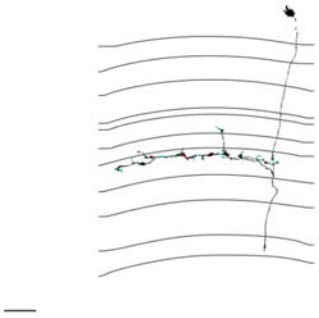

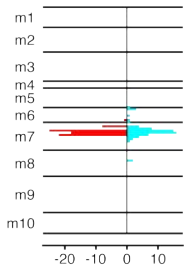


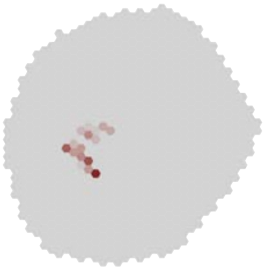

### MeTu3b_720575940620883956

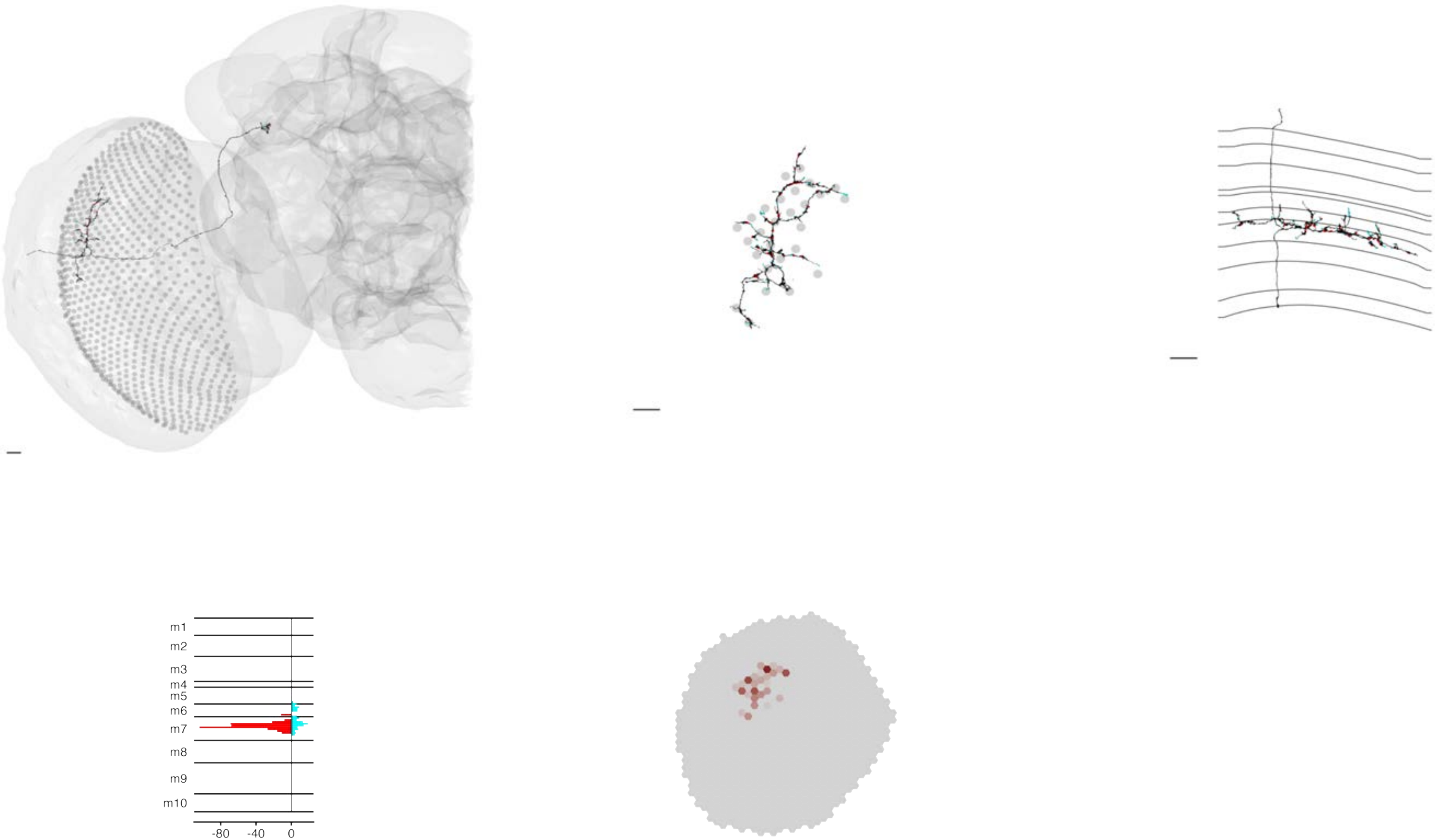

### MeTu3b_720575940621116705

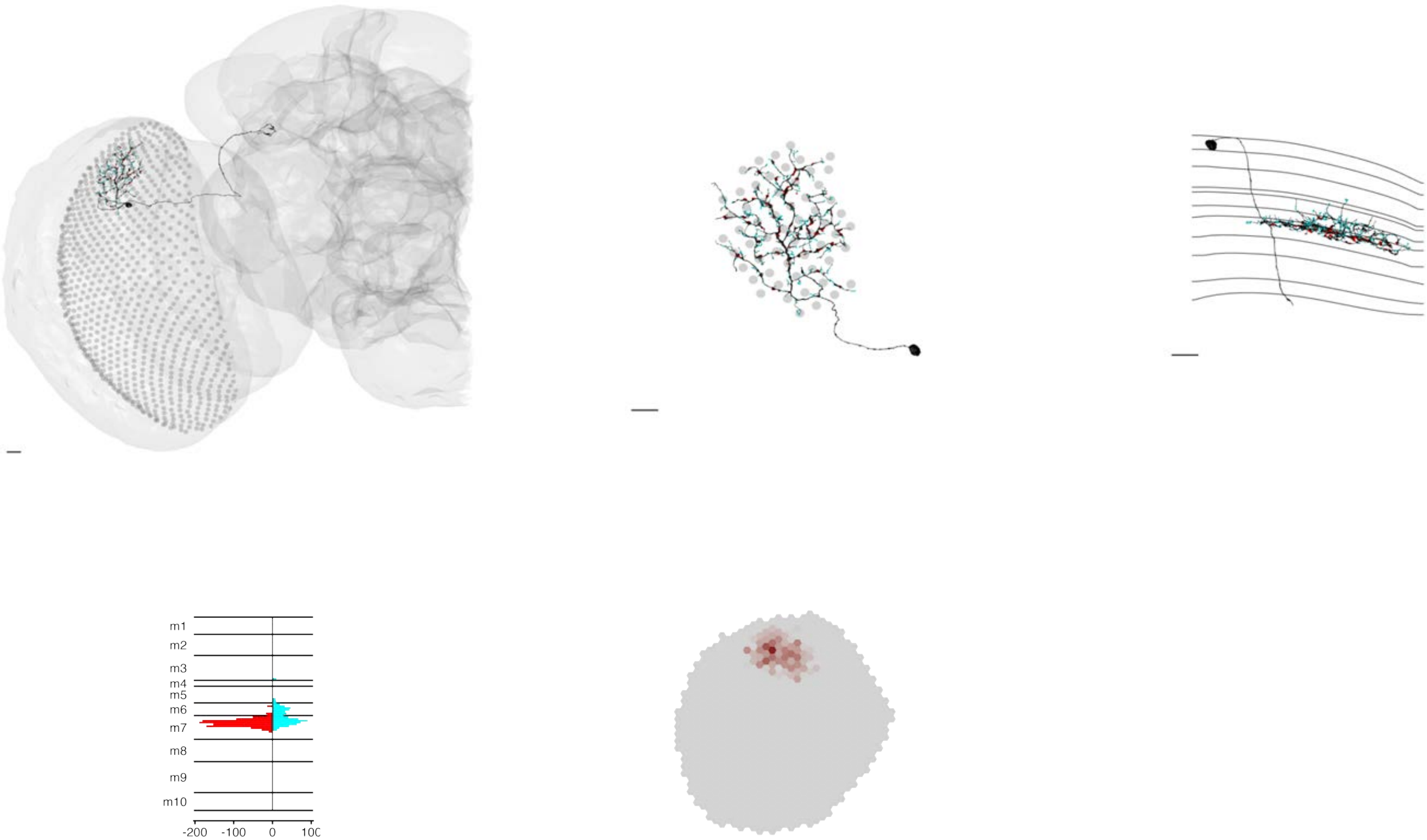

### MeTu3b_720575940621166319

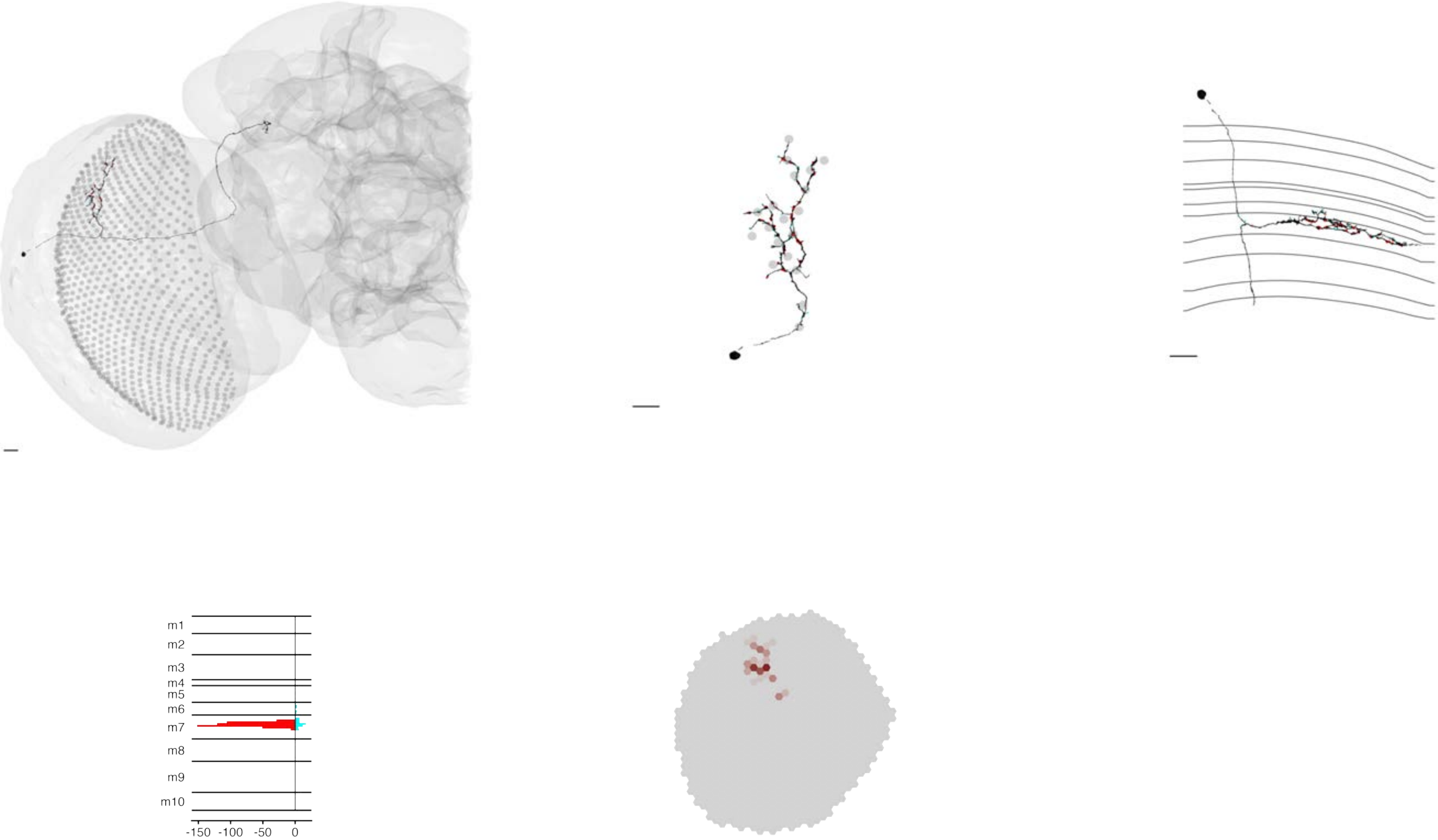

### MeTu3b_720575940621768769

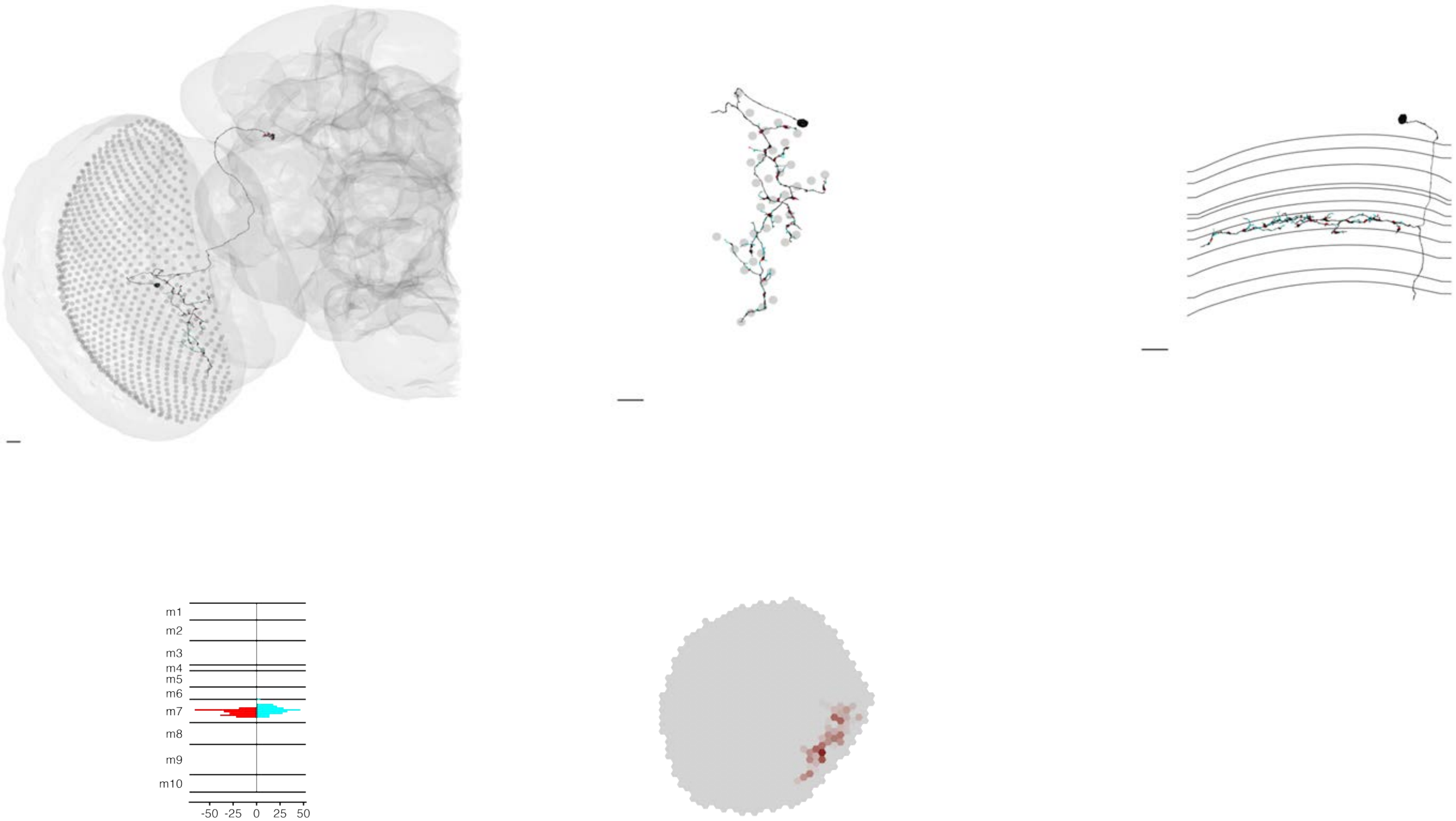

MeTu3b_720575940622172276

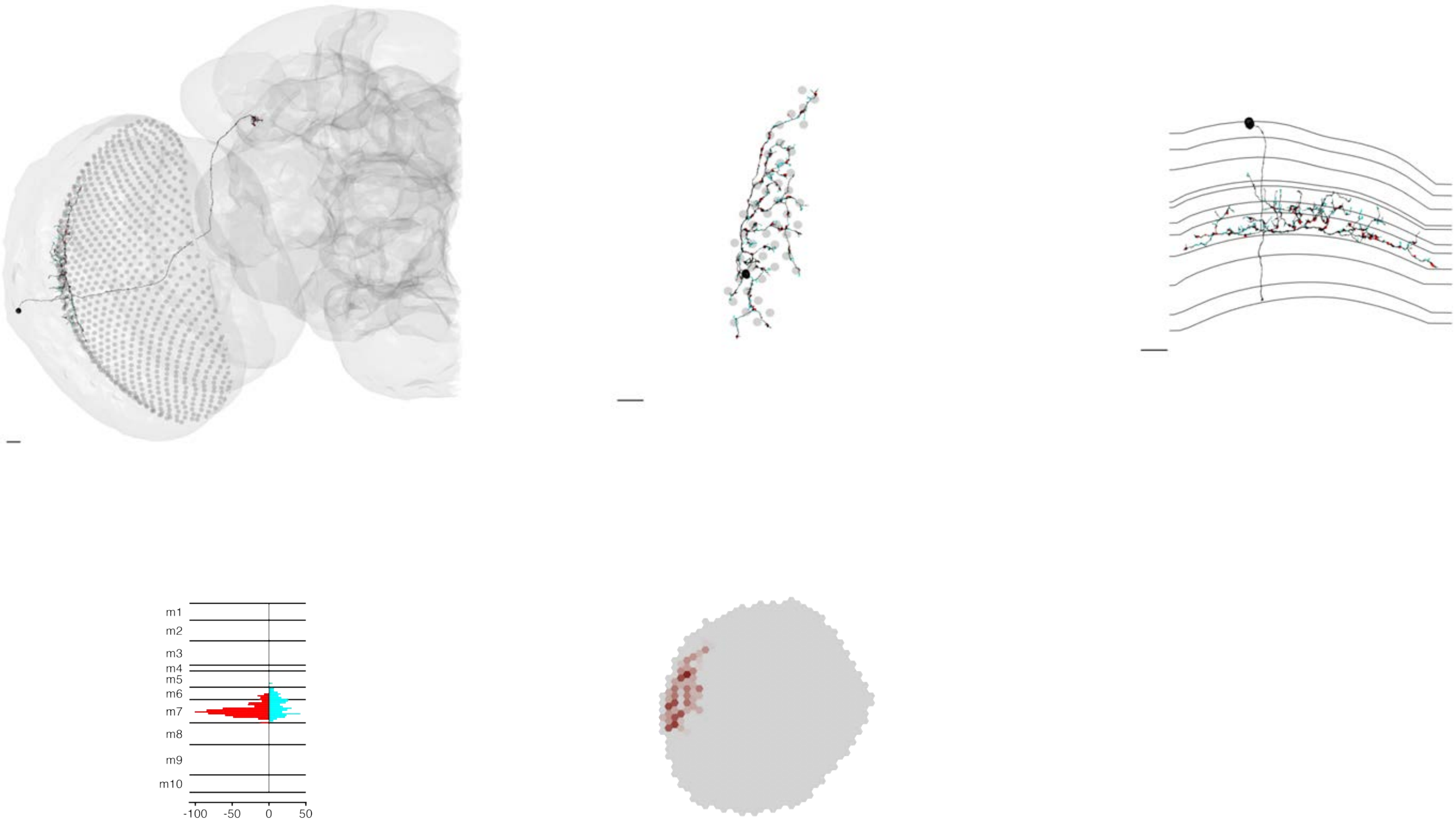

### MeTu3b_720575940622498756

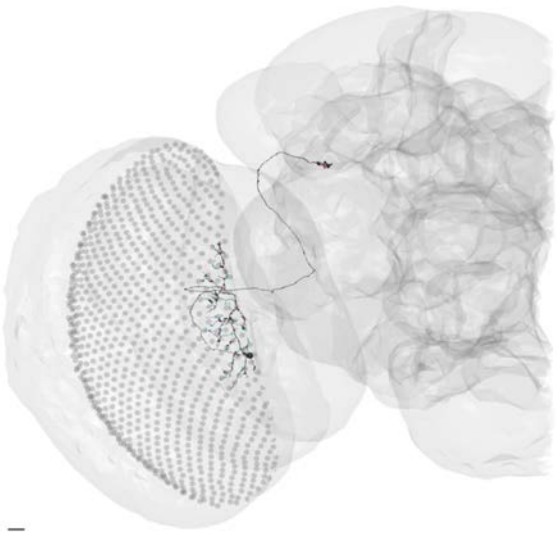

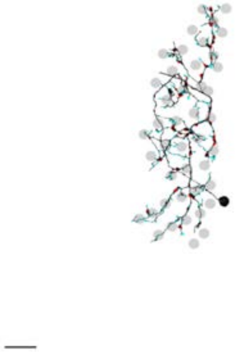

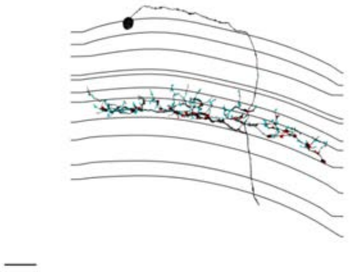

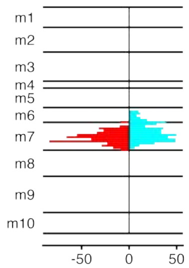


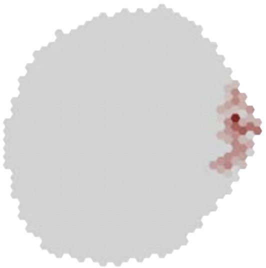

### MeTu3b_720575940622985385

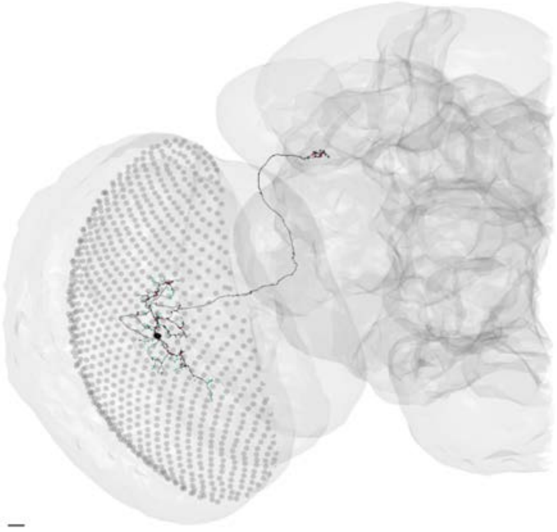

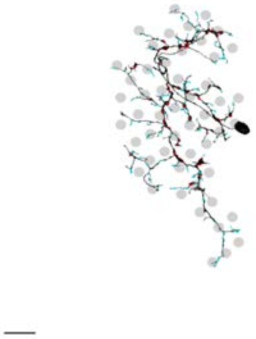

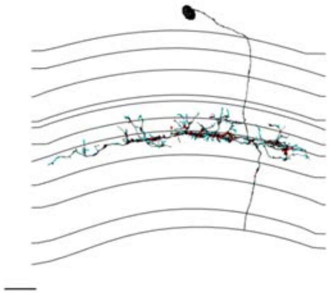

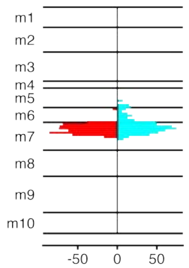


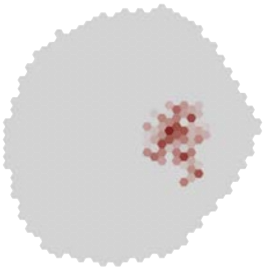

### MeTu3b_720575940623634570

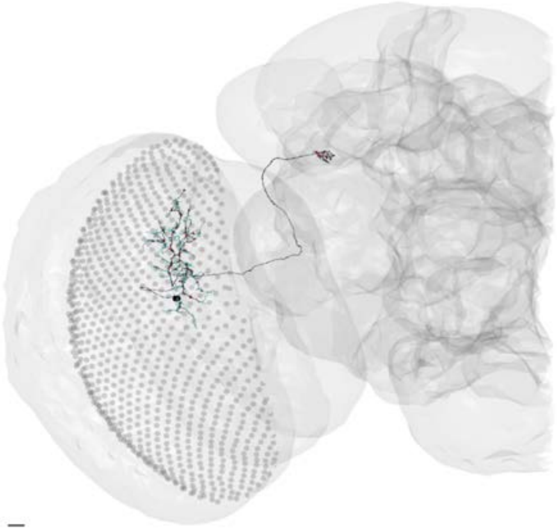

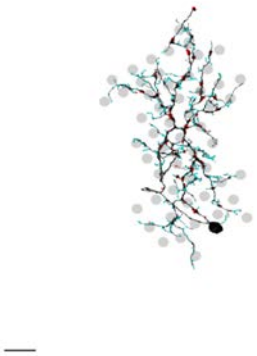

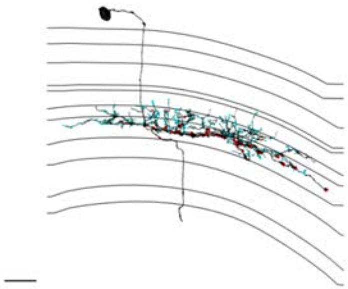

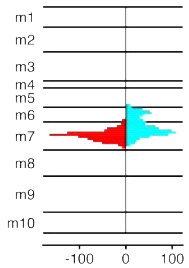


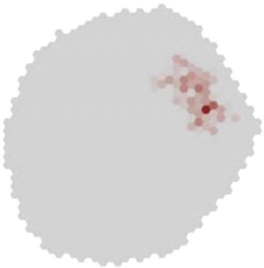

### MeTu3b_720575940623876071

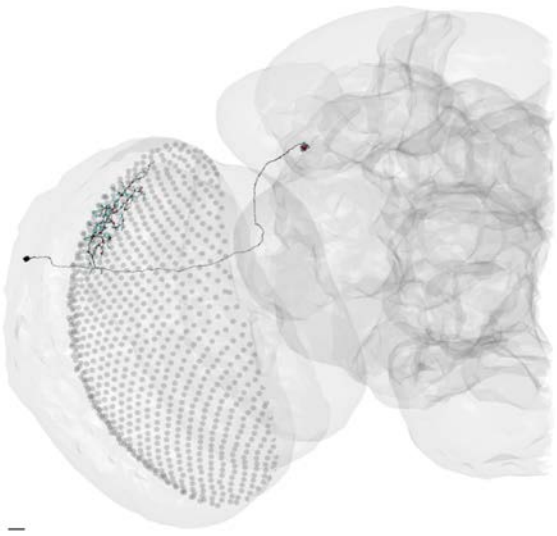

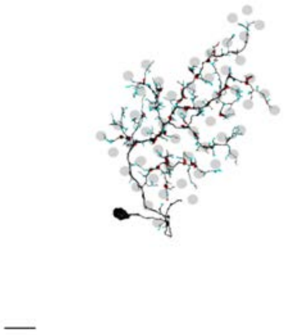

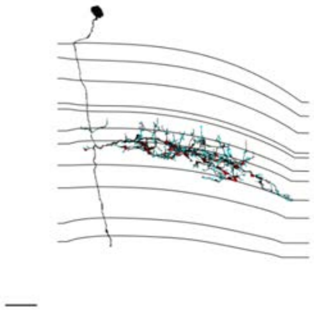

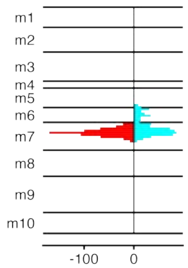


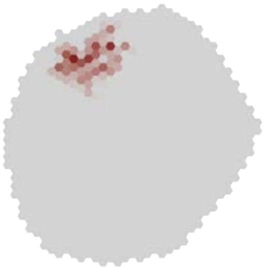

### MeTu3b_720575940624004843

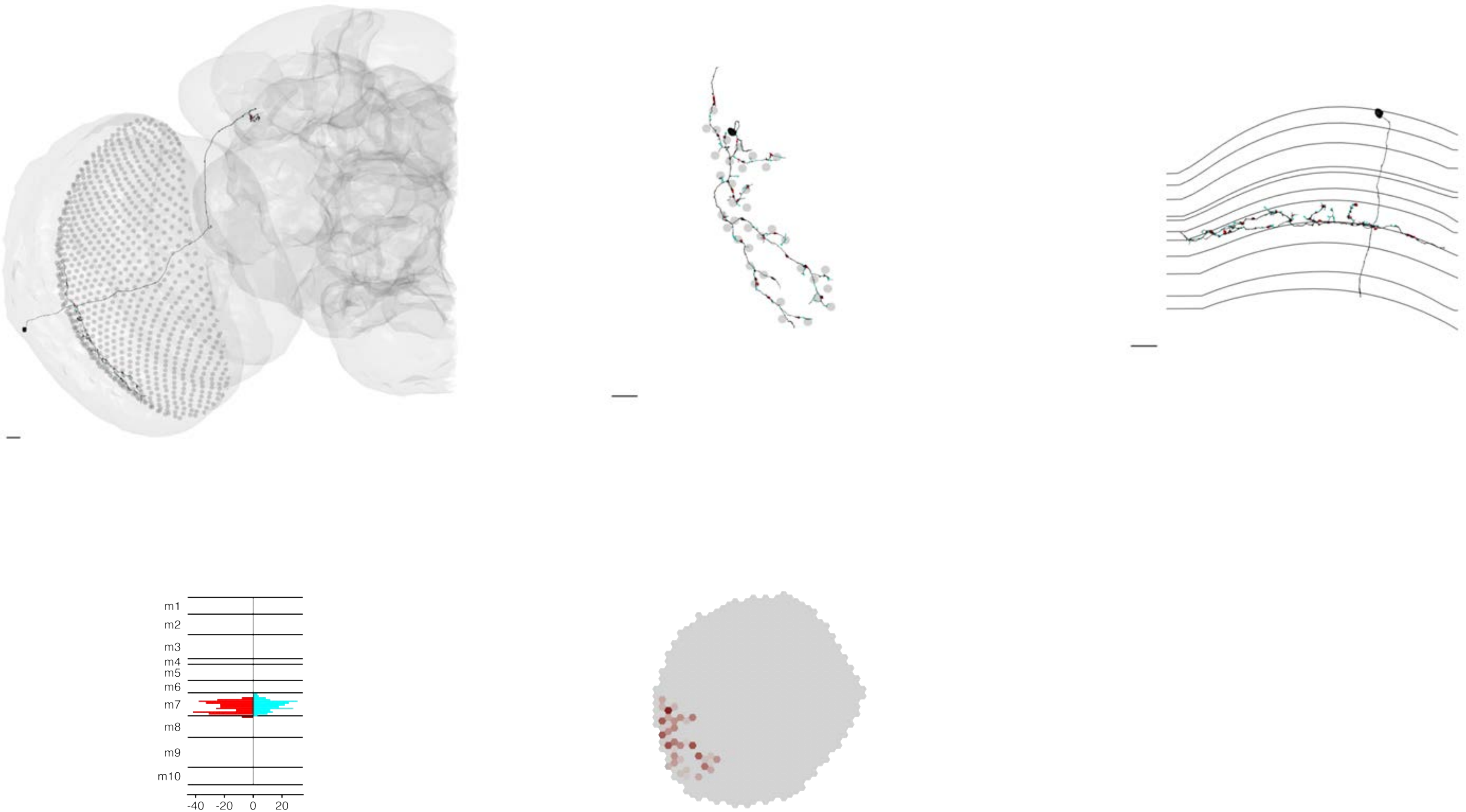

### MeTu3b_720575940624085944

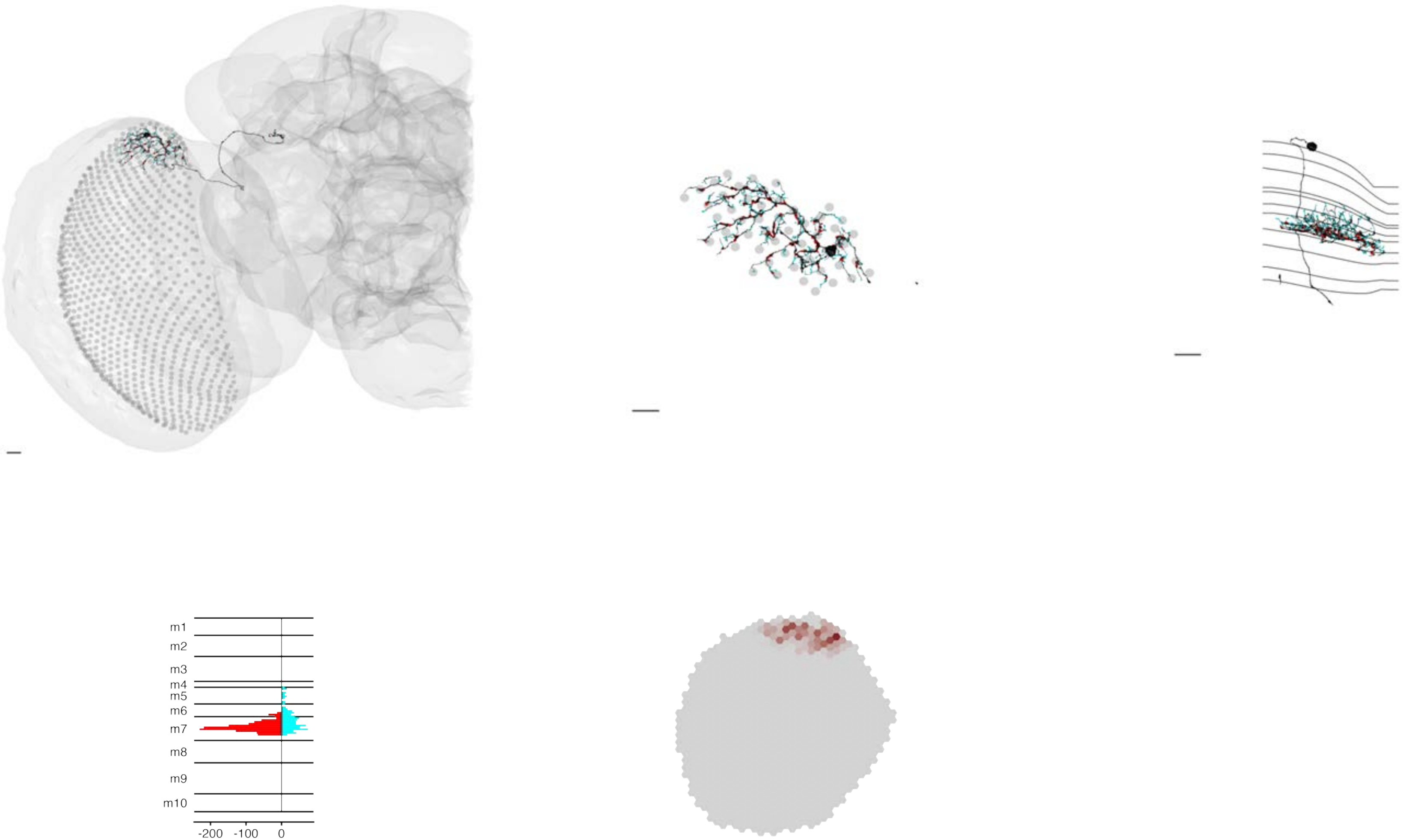

### MeTu3b_720575940625622274

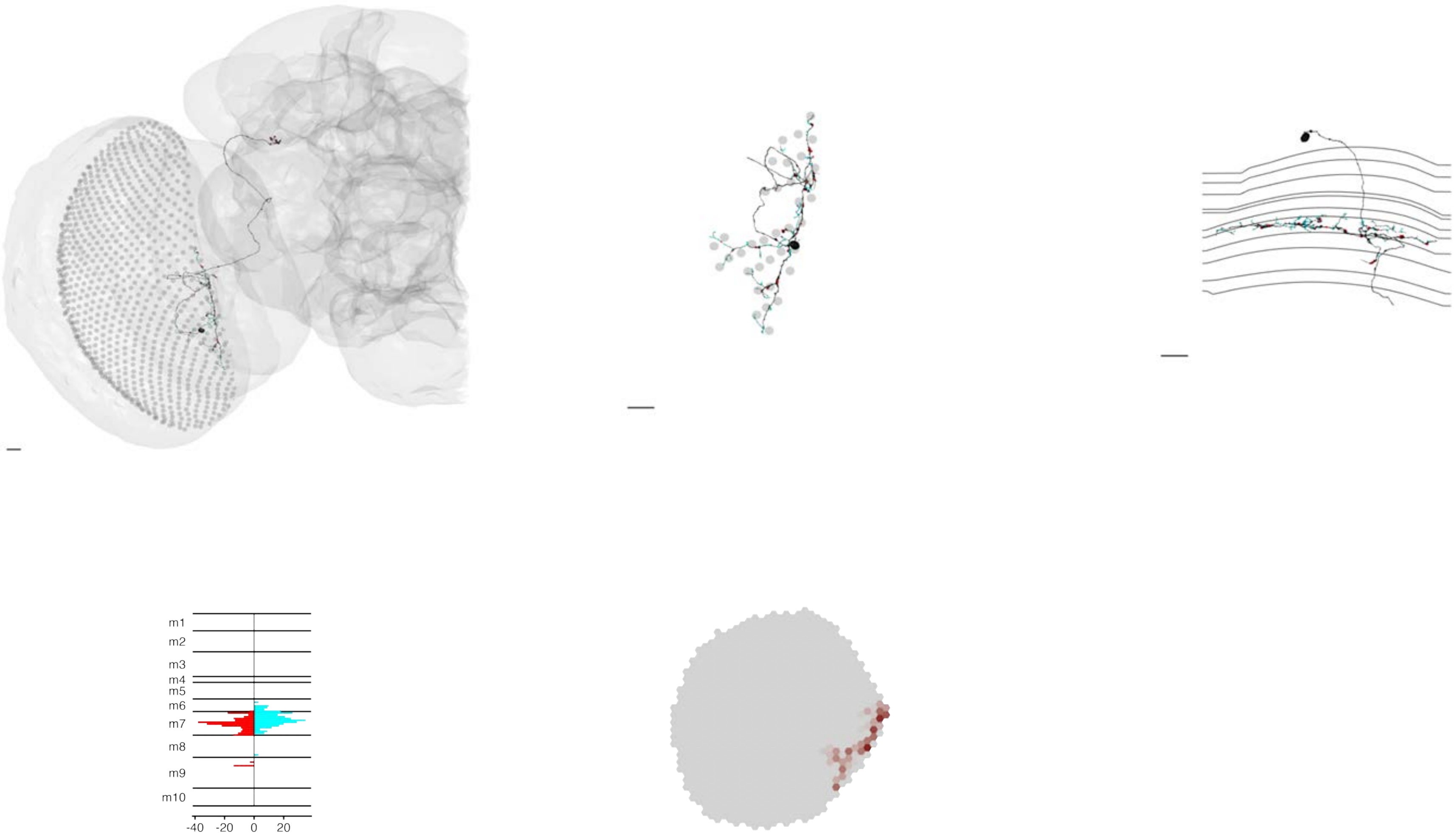

### MeTu3b_720575940626711358

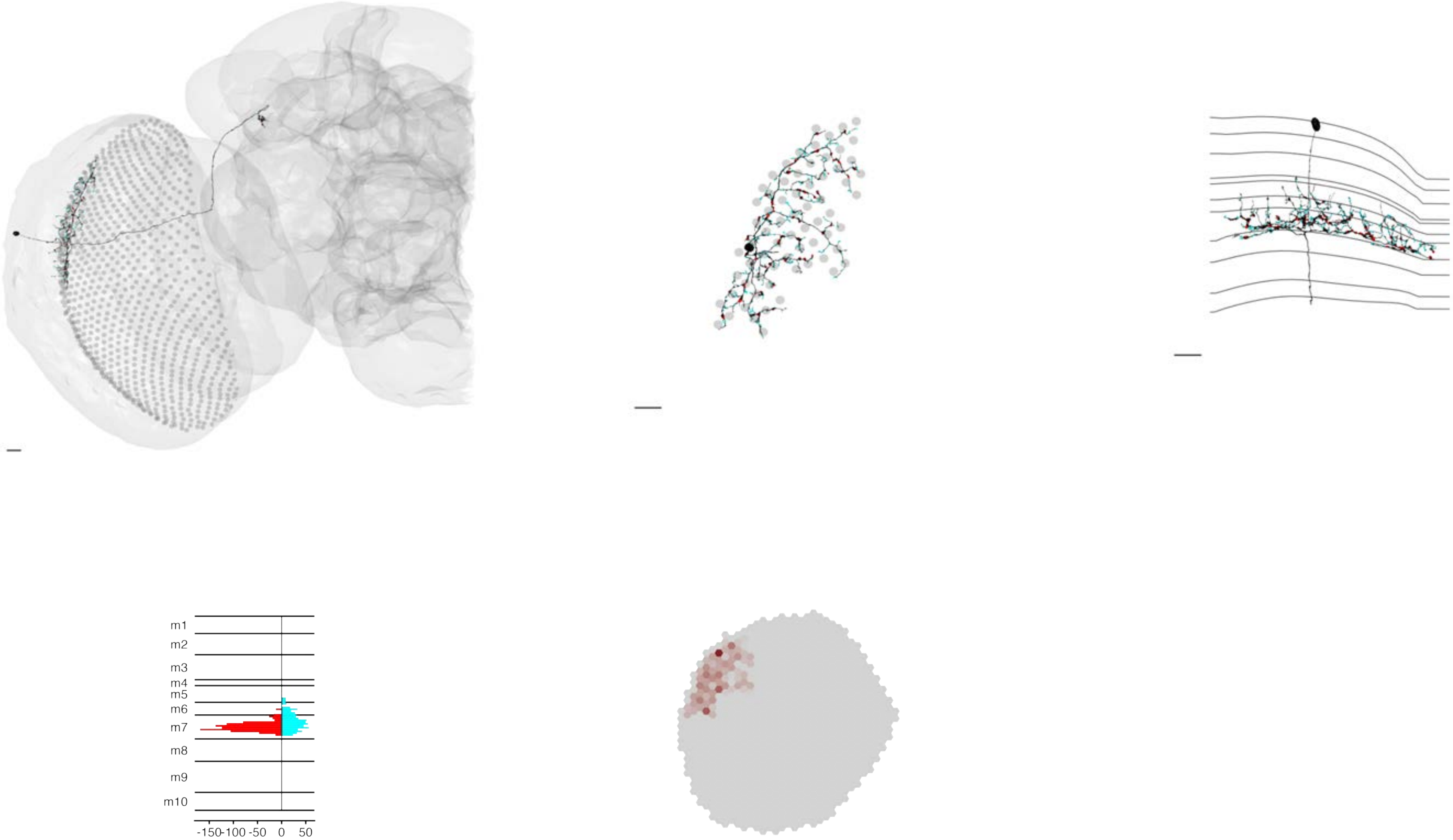

MeTu3b_720575940628221456

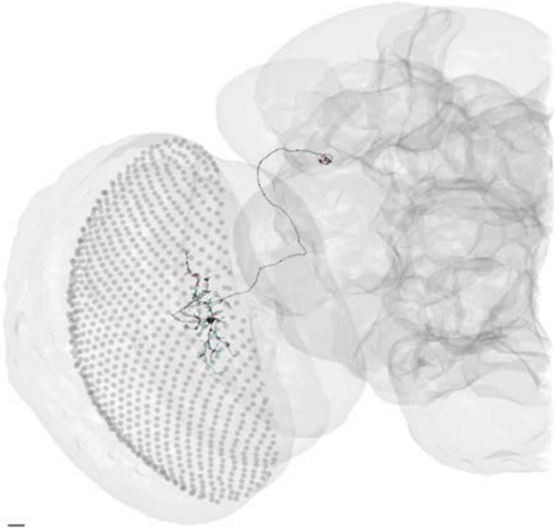

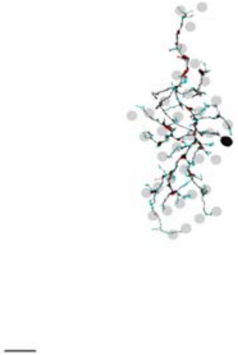

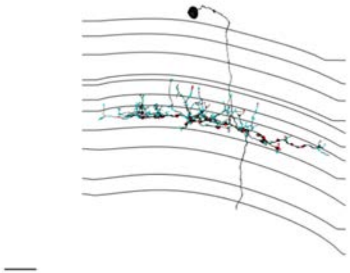

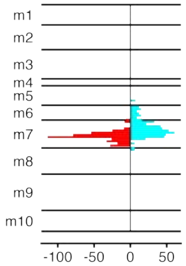


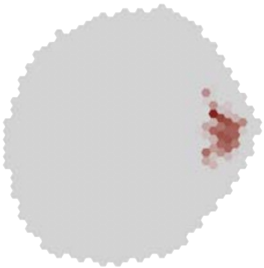

MeTu3b_720575940628531194

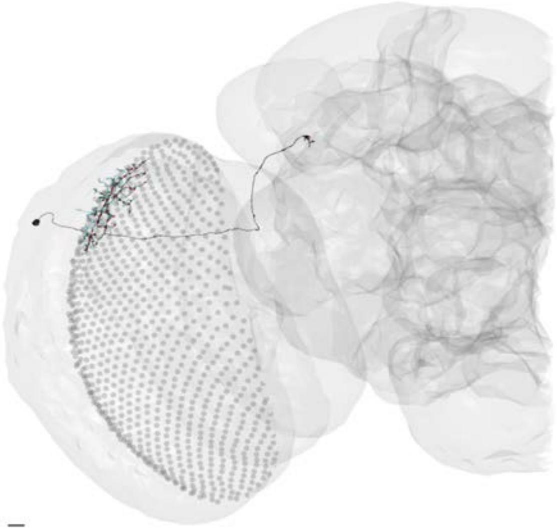

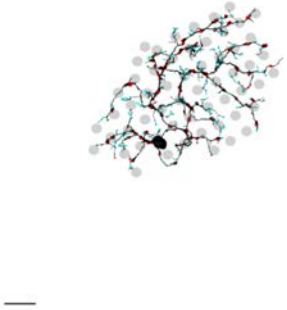

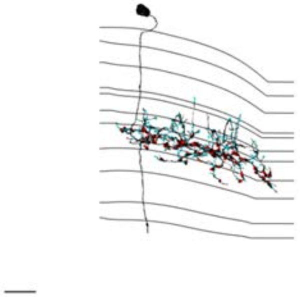

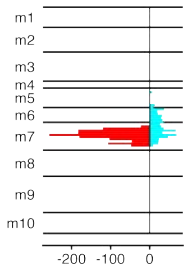


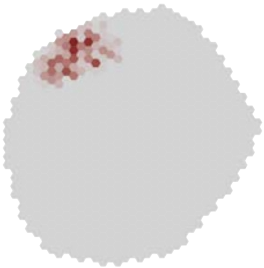

MeTu3b_720575940628752502

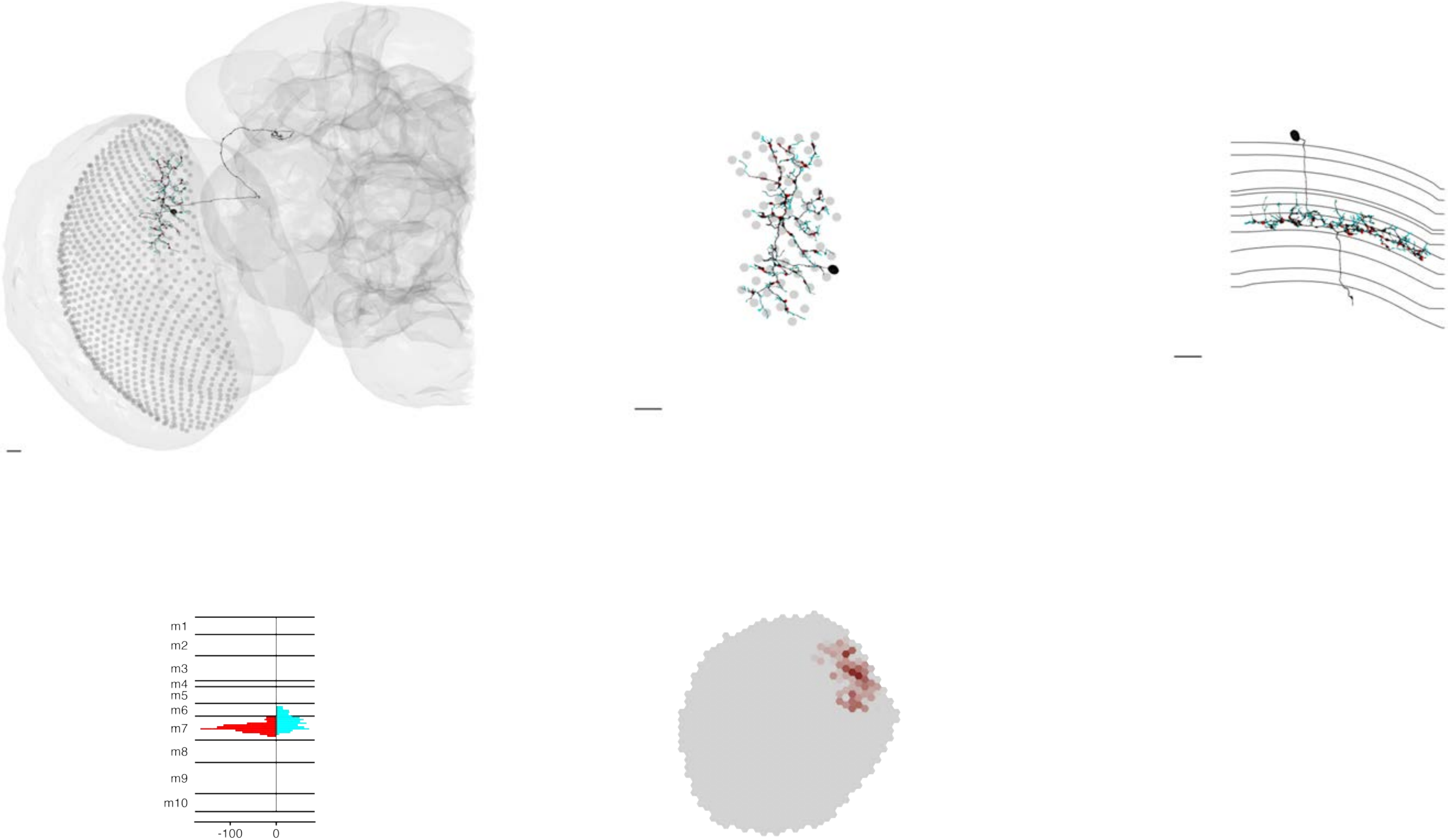

### MeTu3b_720575940629357903

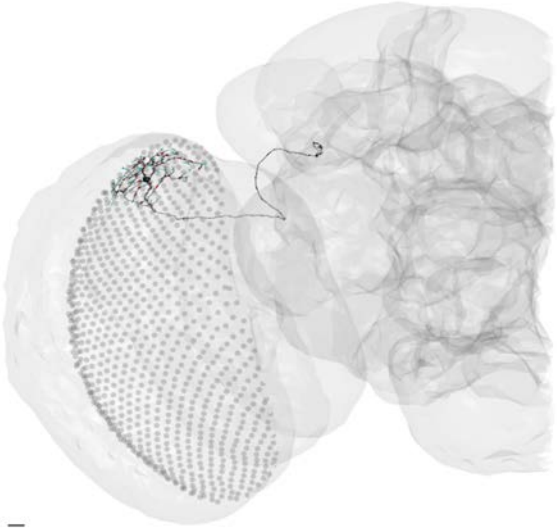

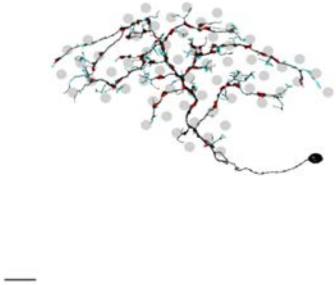

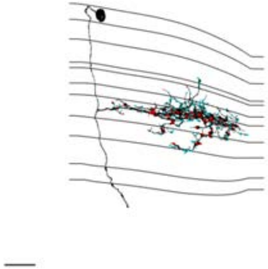

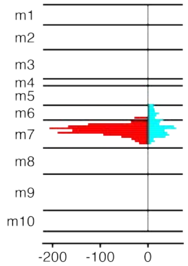


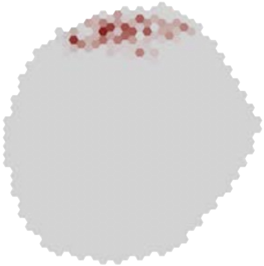

### MeTu3b_720575940631151478

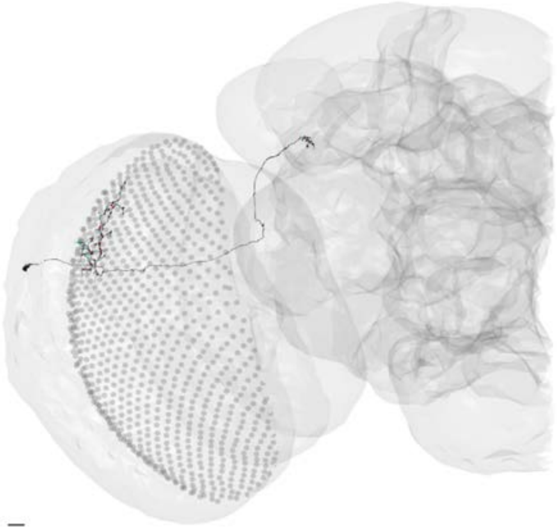

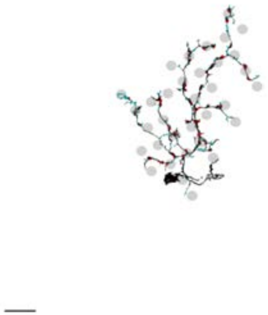

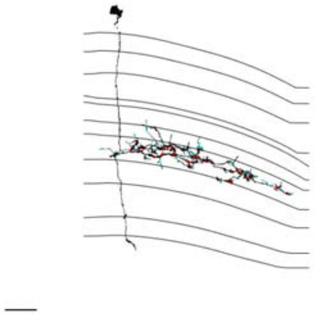

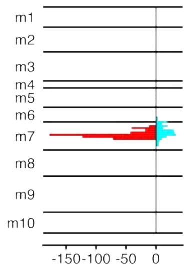


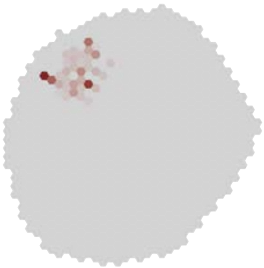

### MeTu3b_720575940631322193

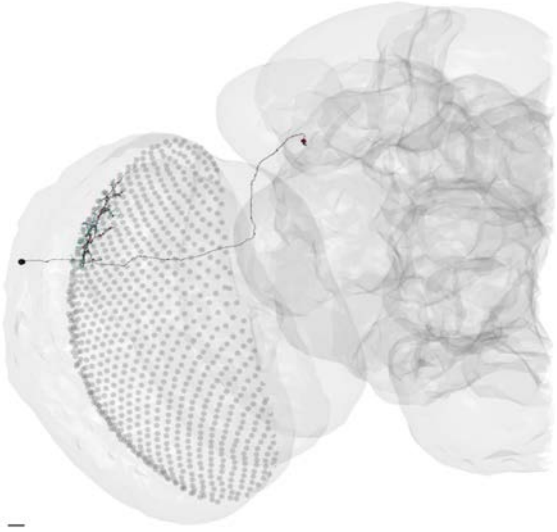

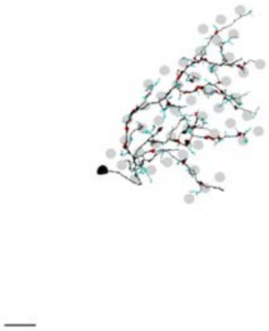

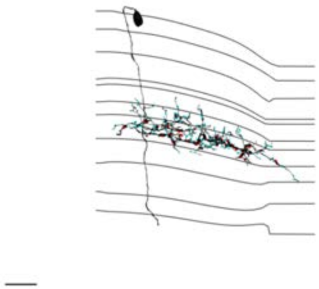

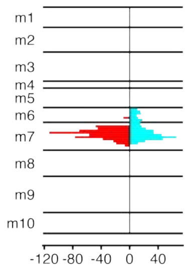


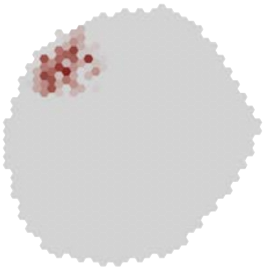

### MeTu3b_720575940631504847

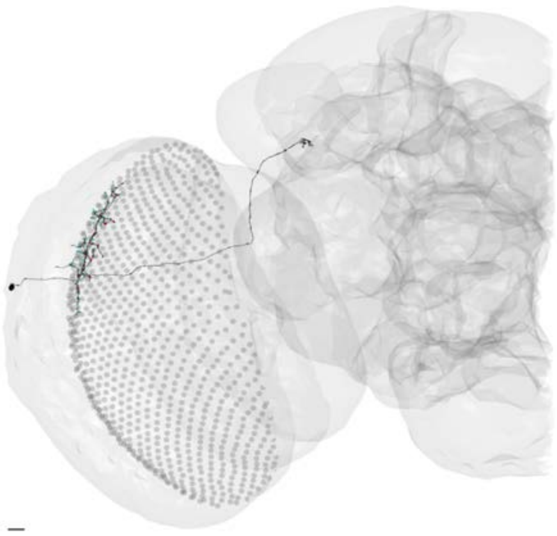

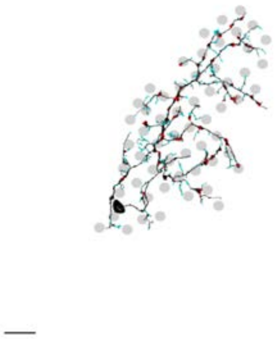

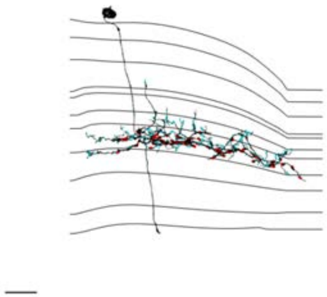

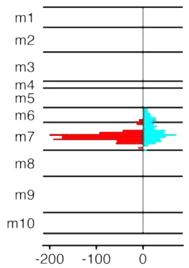


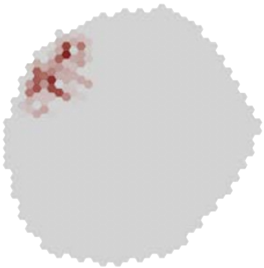

### MeTu3b_720575940632013357

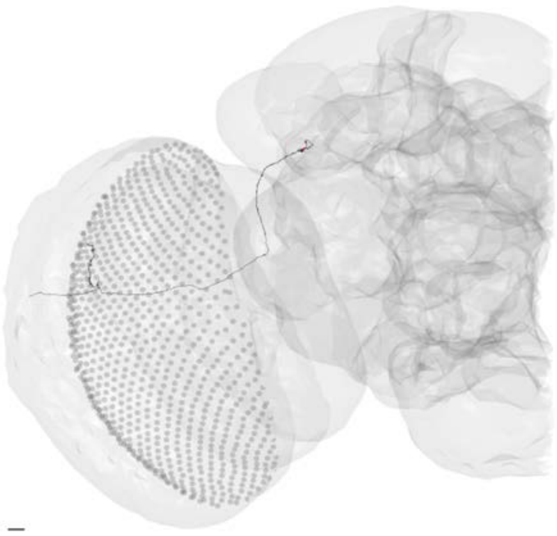

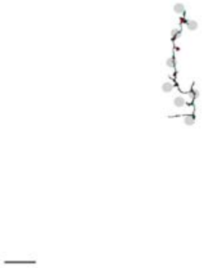

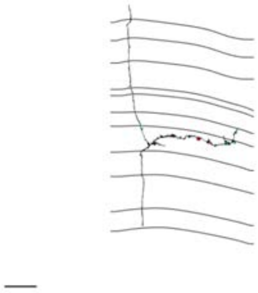

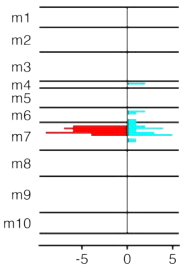


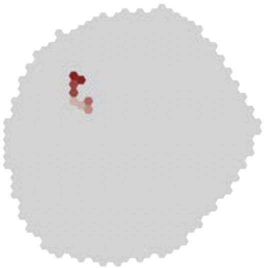

### MeTu3b_720575940632078561

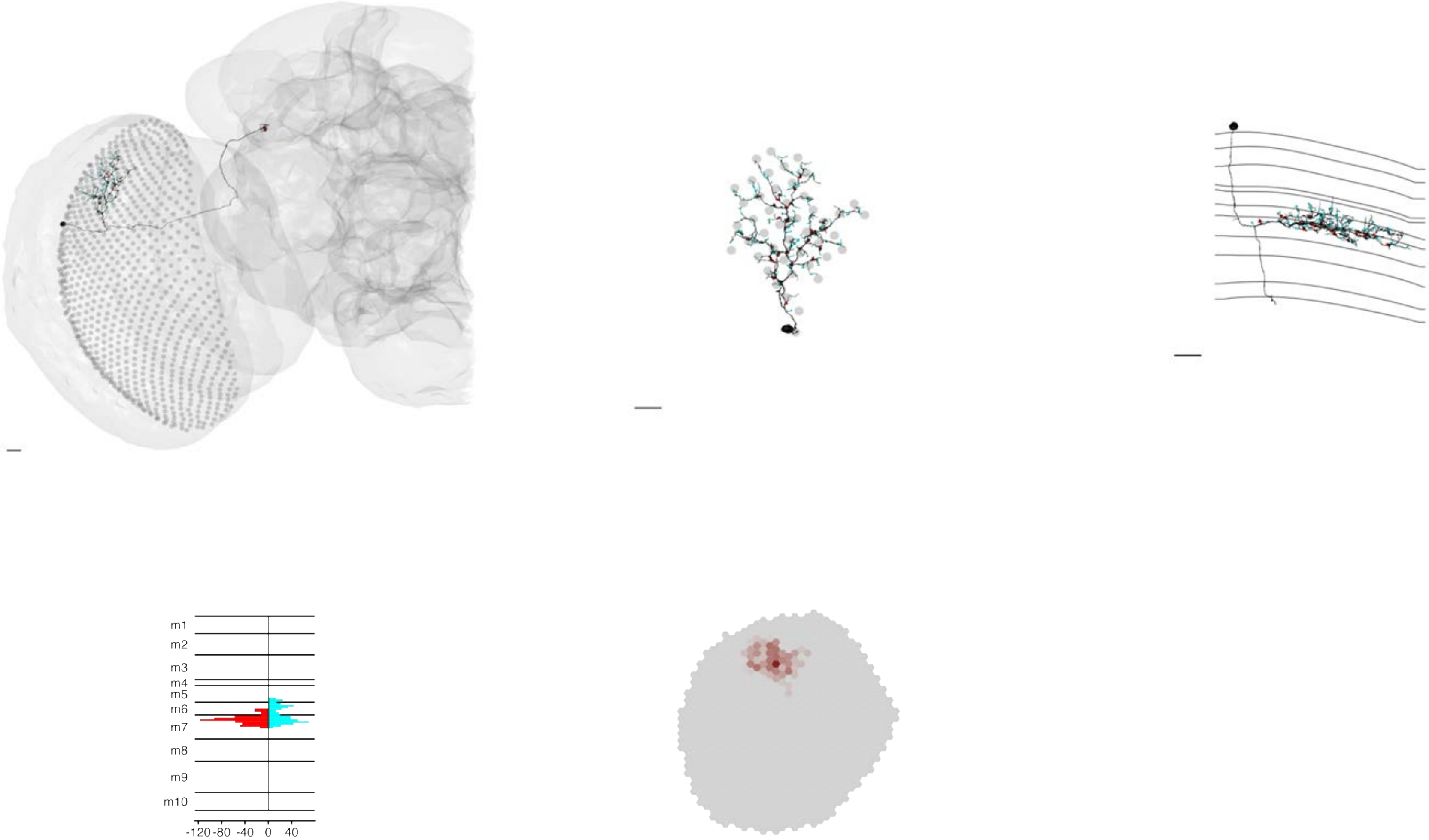

### MeTu3b_720575940632451245

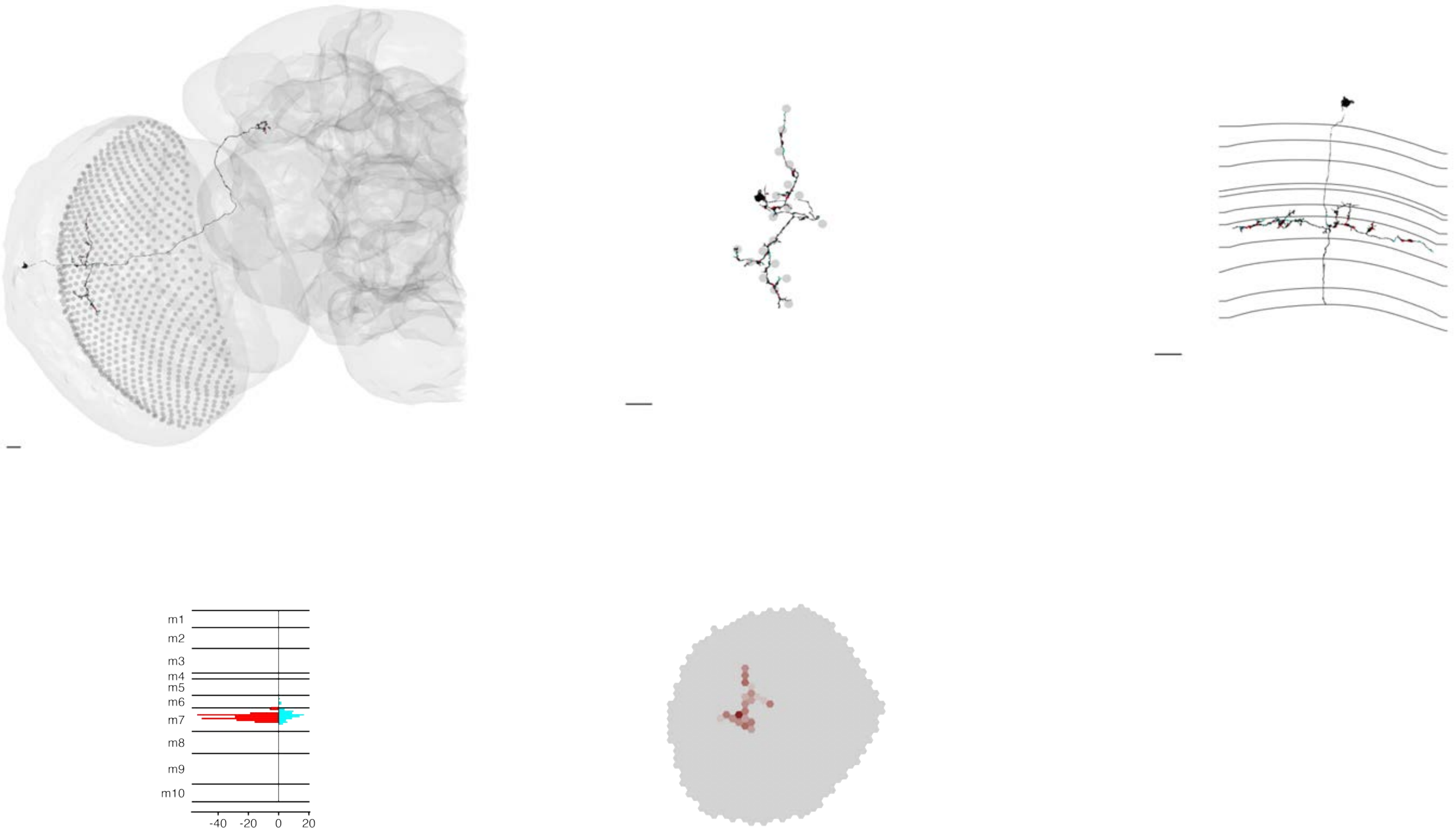

### MeTu3b_720575940633431972

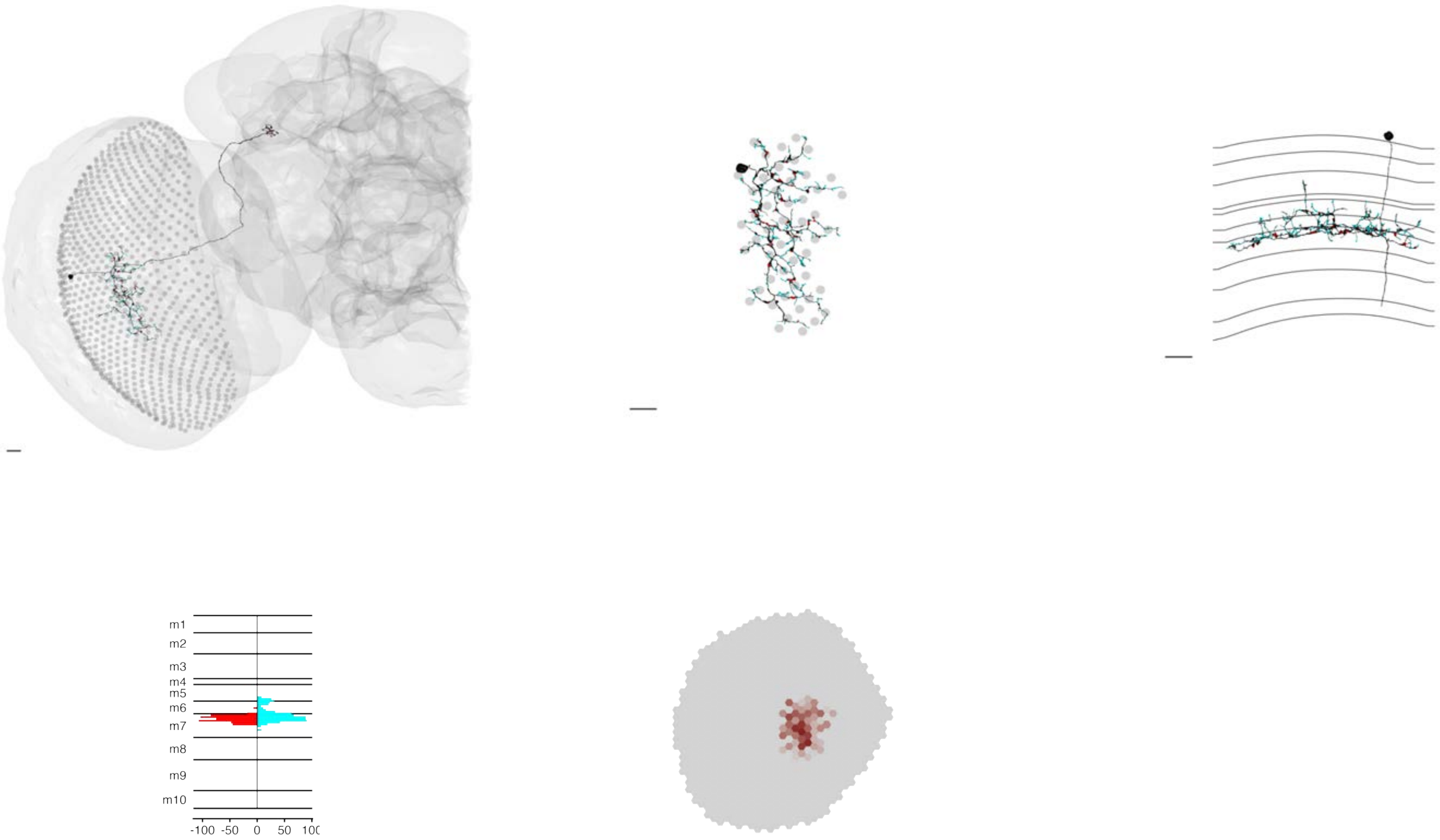

### MeTu3b_720575940633963872

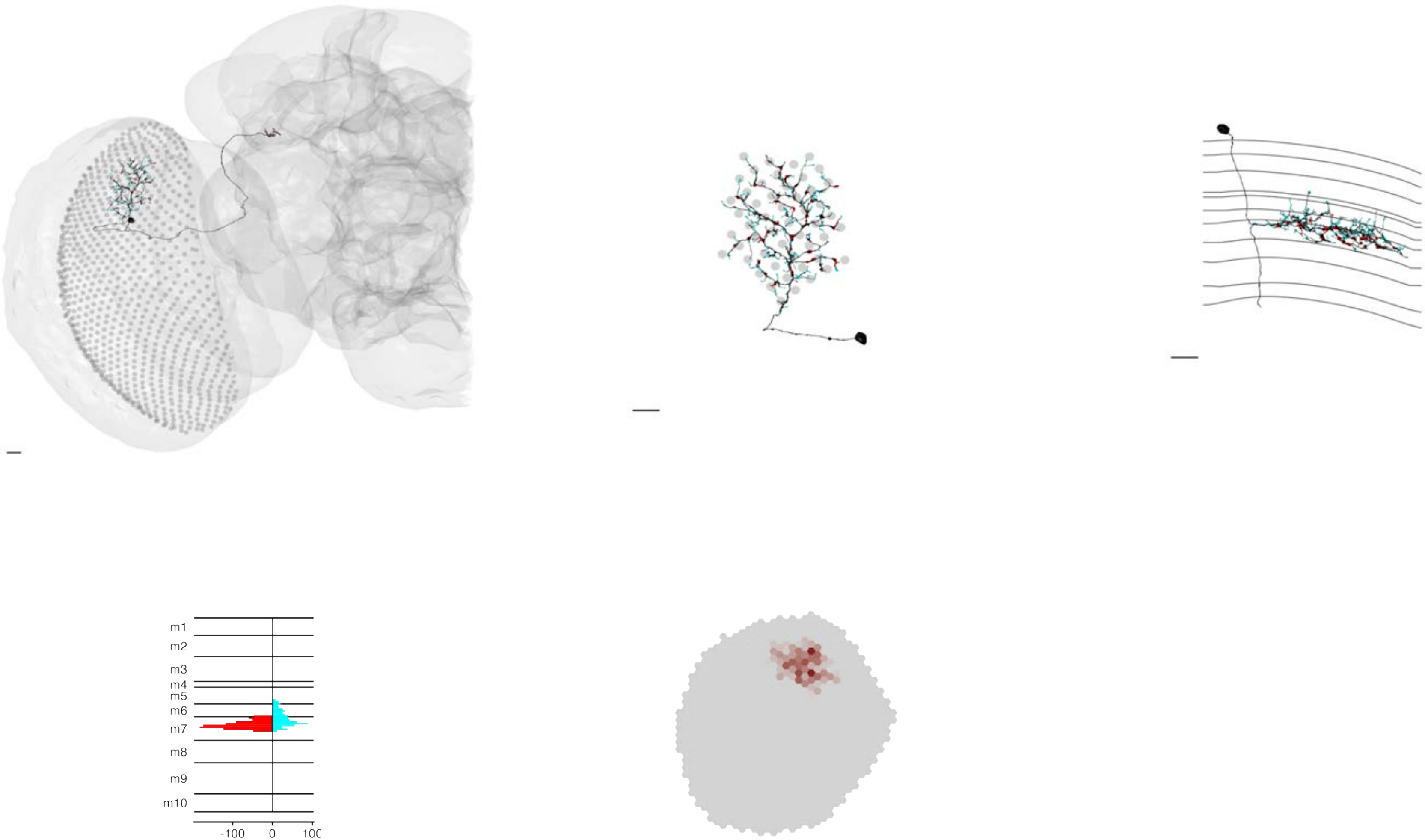

### MeTu3b_720575940634151348

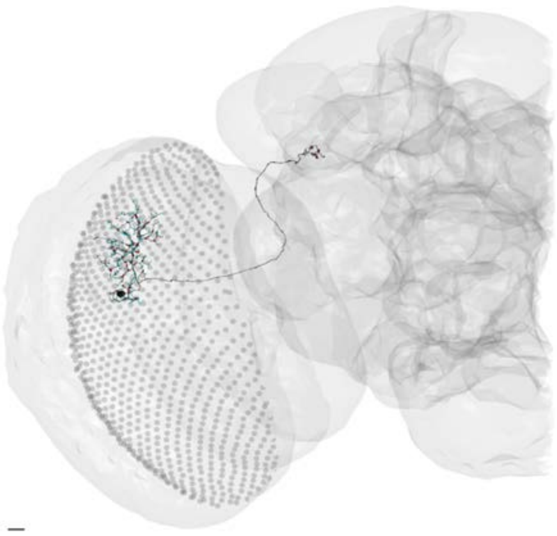

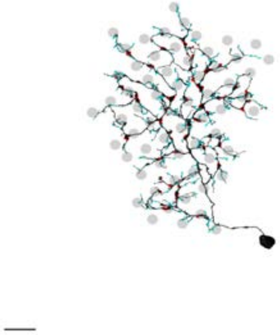

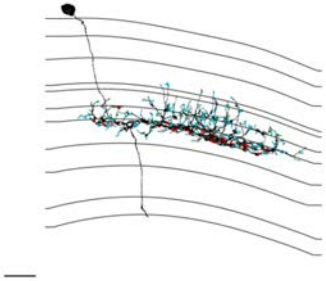

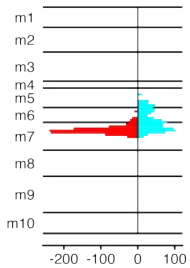


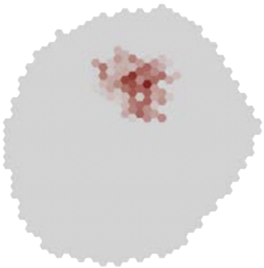

### MeTu3b_720575940634582623

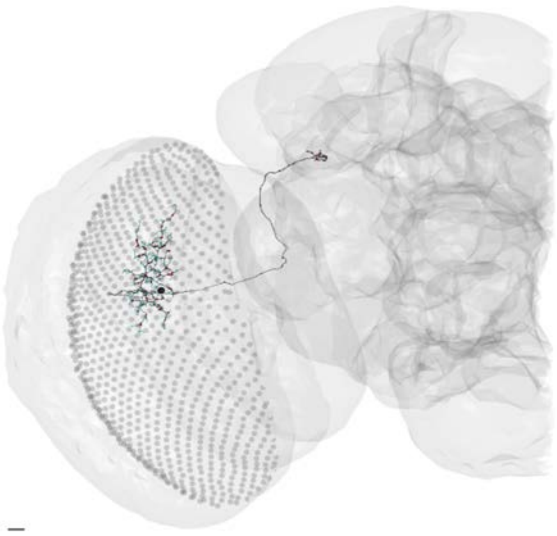

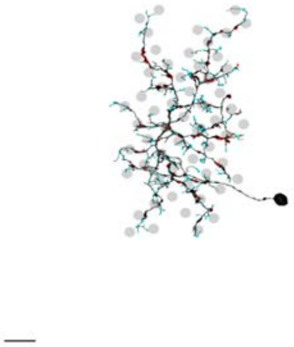

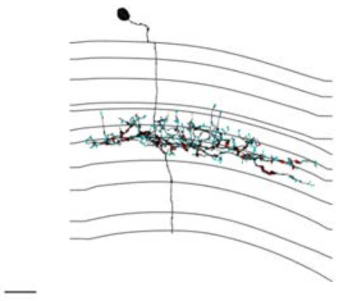

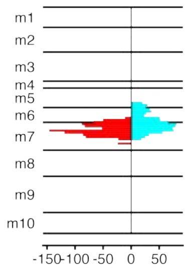


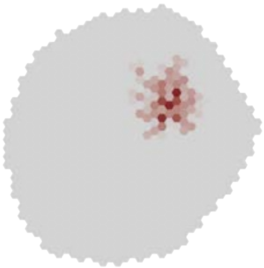

### MeTu3b_720575940635176032

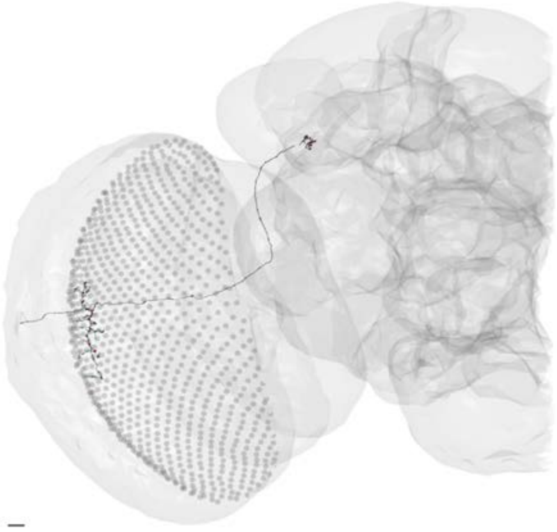

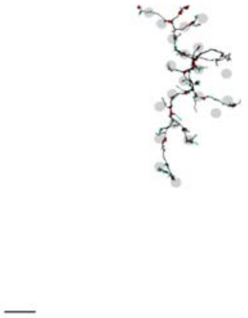

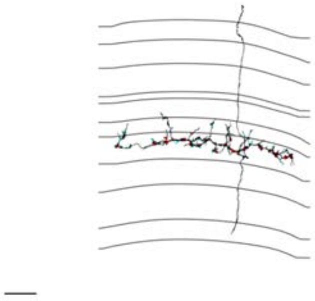

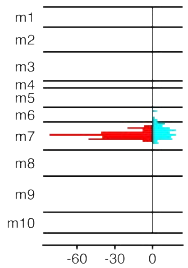


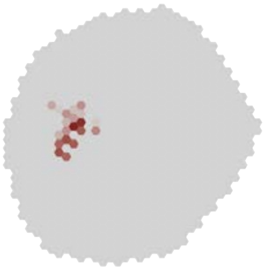

### MeTu3b_720575940637478234

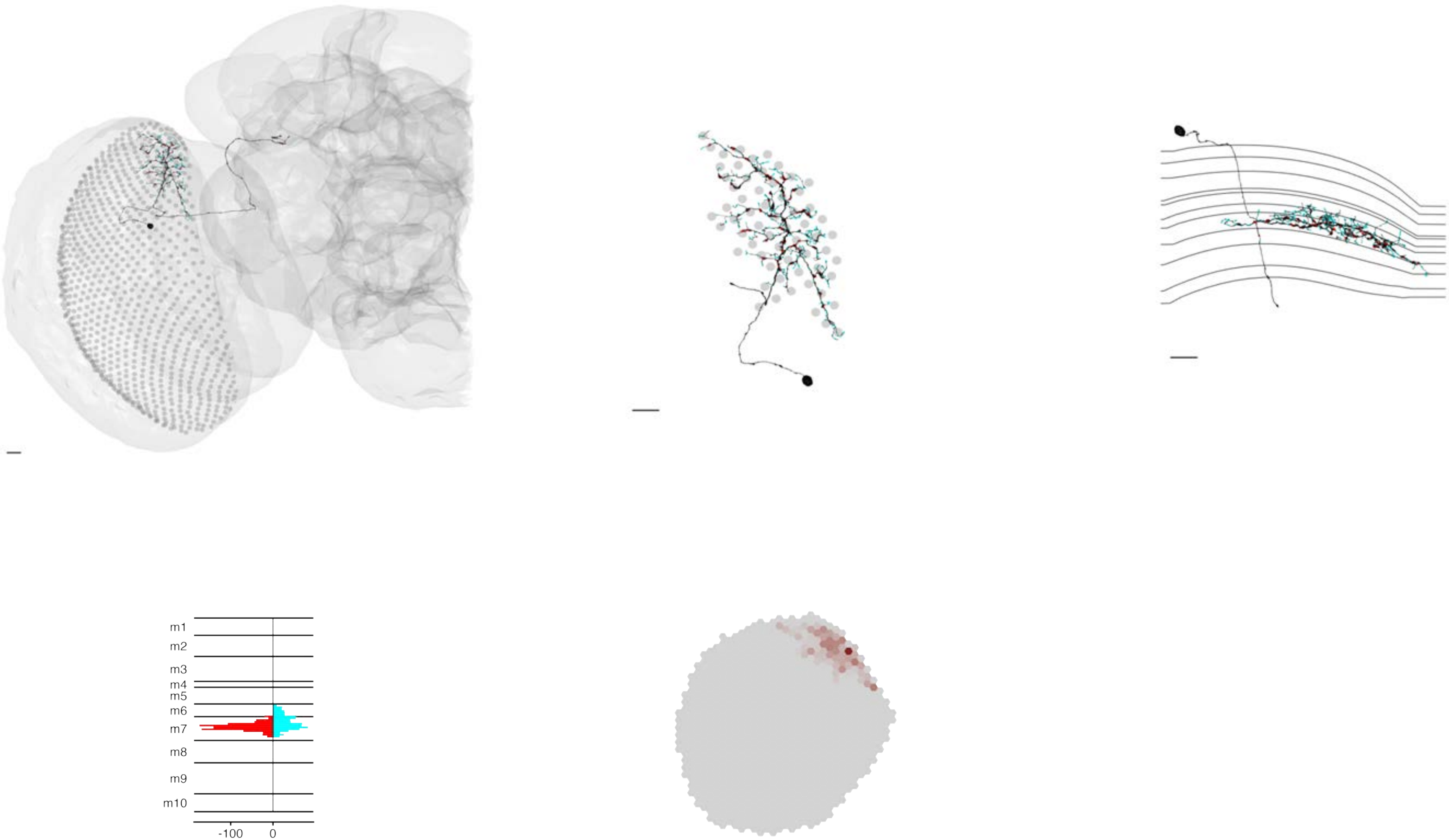

### MeTu3b_720575940637714803

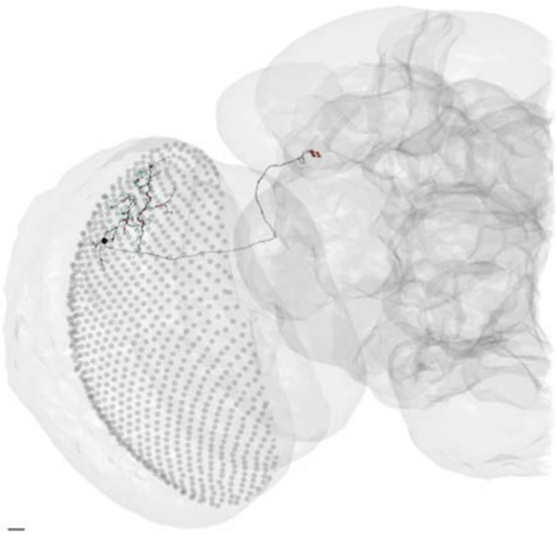

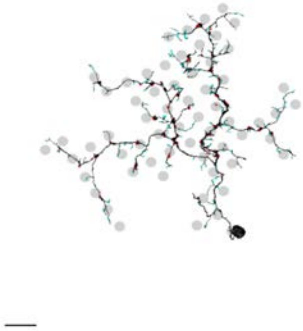

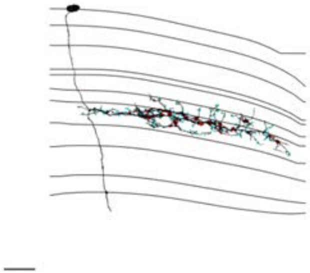

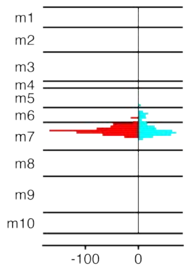


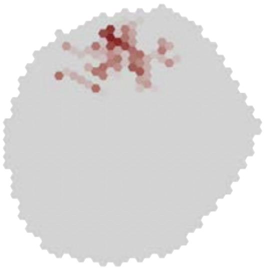

### MeTu3b_720575940638084174

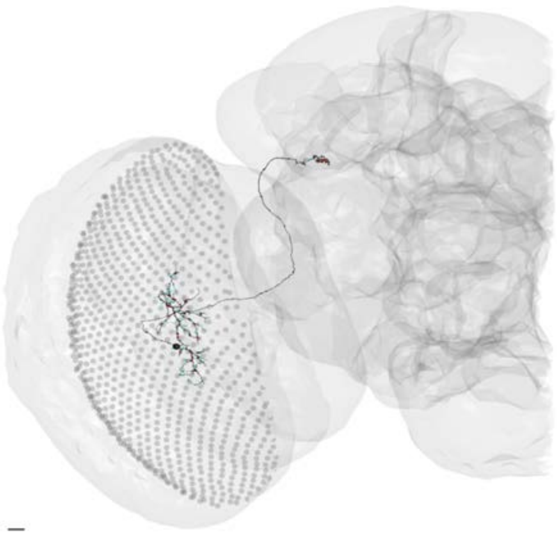

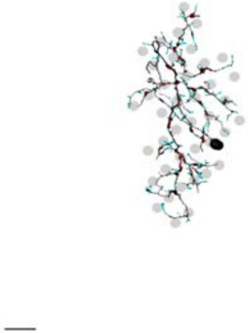

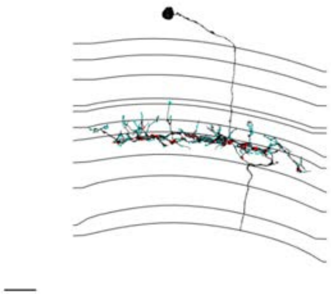

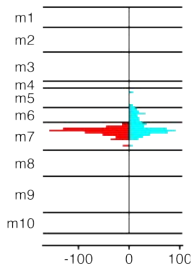


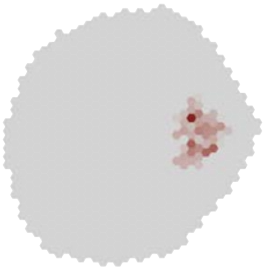

### MeTu3b_720575940638604221

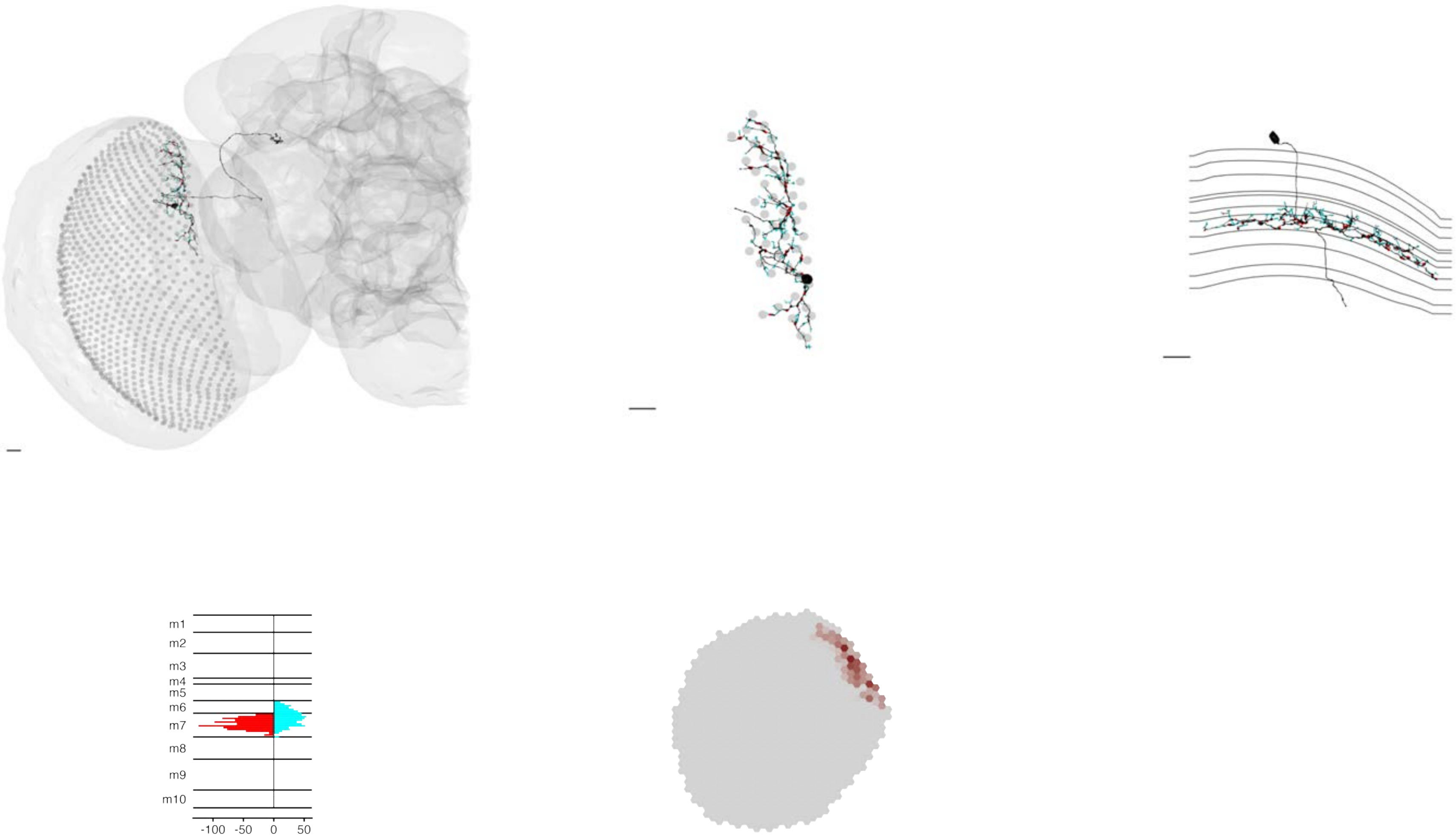

### MeTu3b_720575940642870925

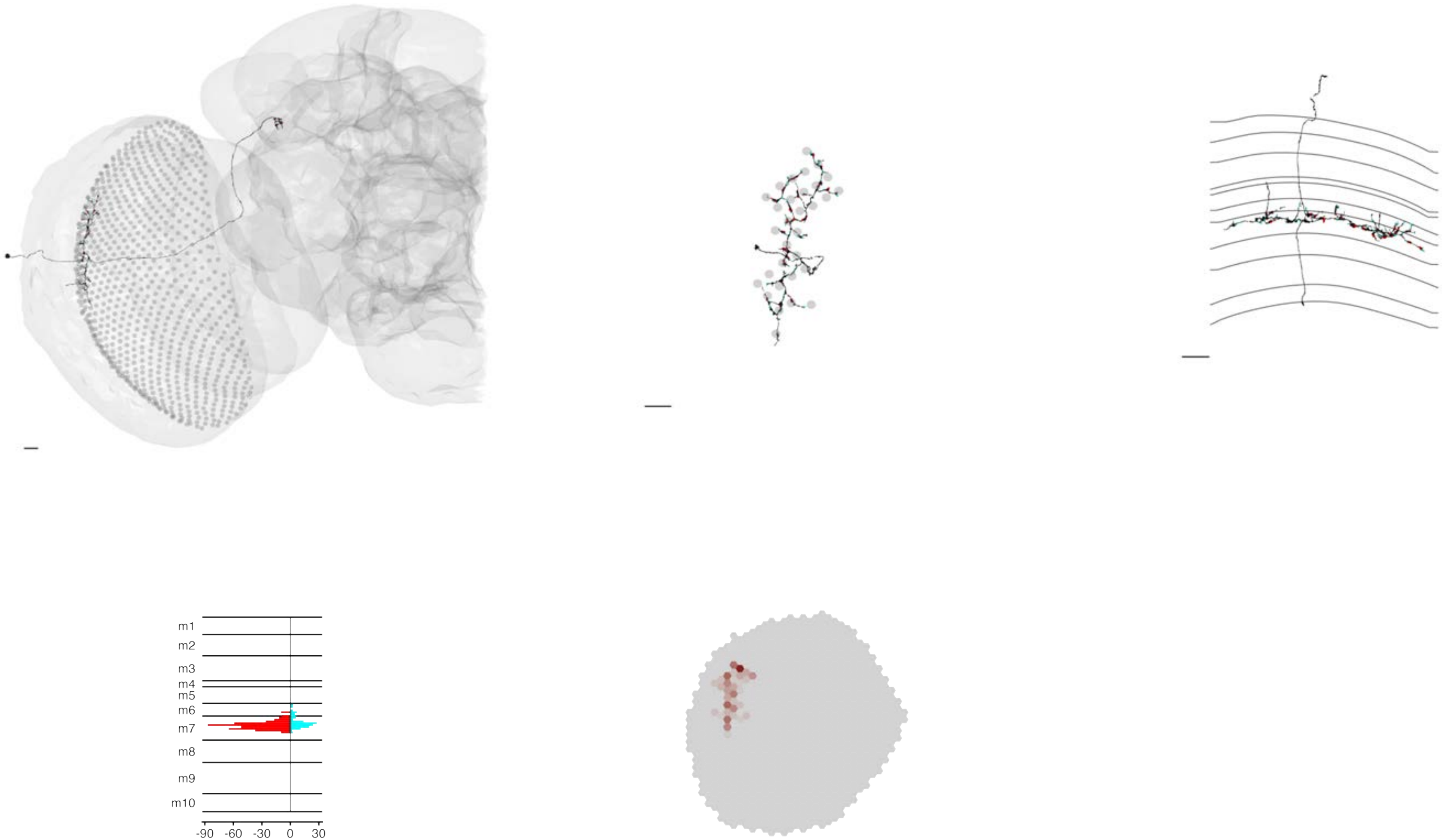

### MeTu3b_720575940645808407

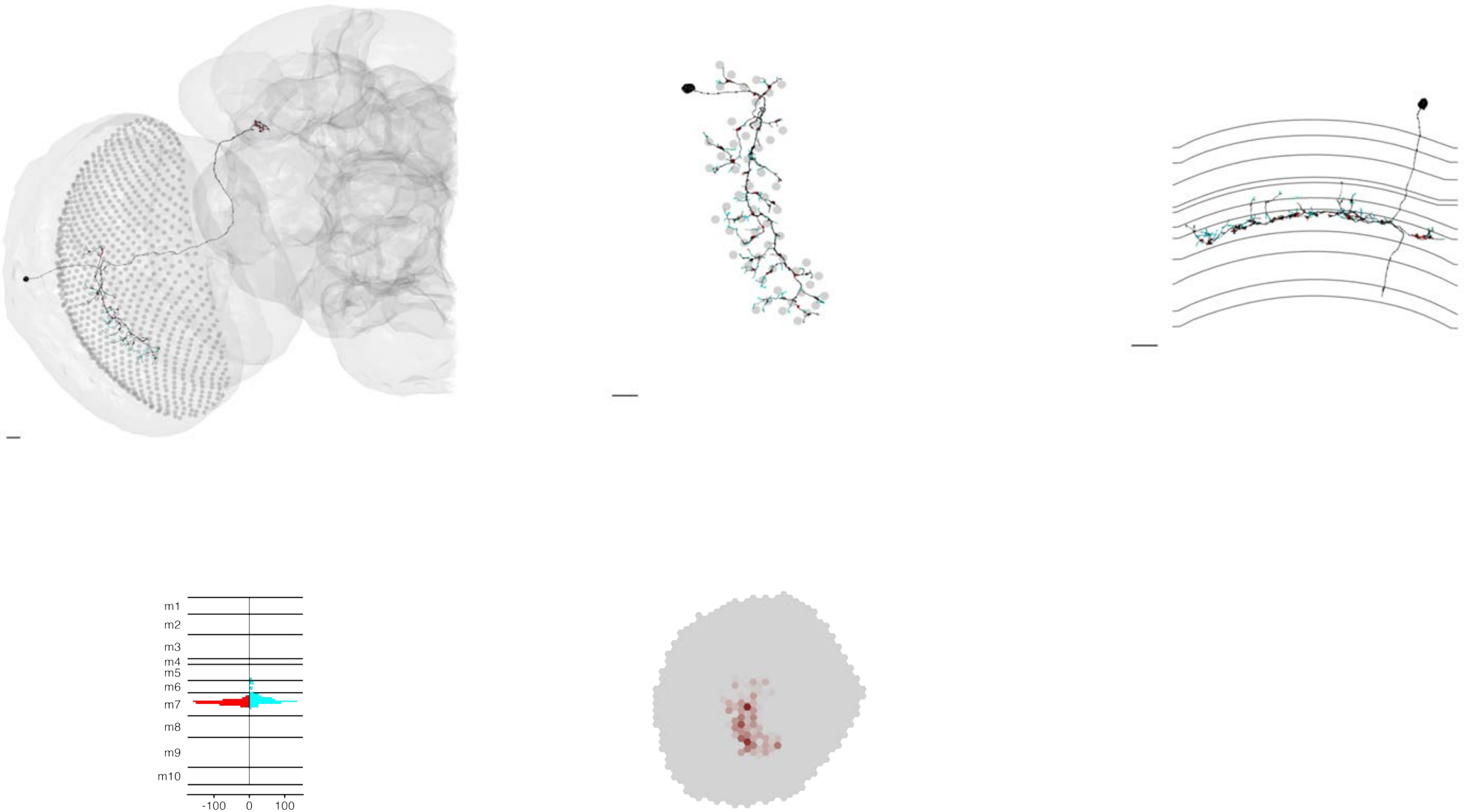

### MeTu3c_dorsal_720575940613764862

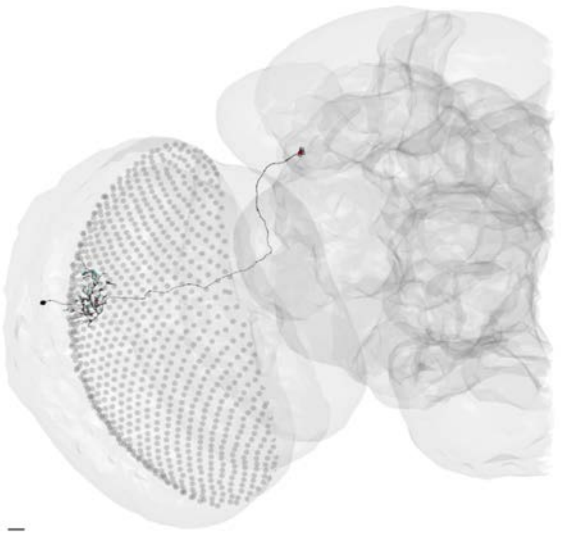

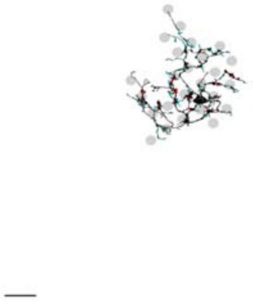

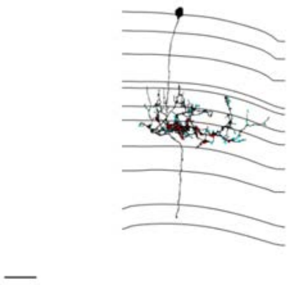

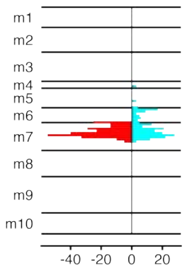


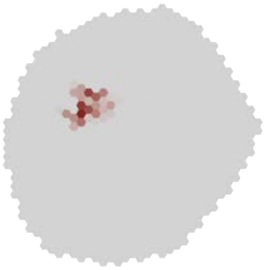

### MeTu3c_dorsal_720575940614930226

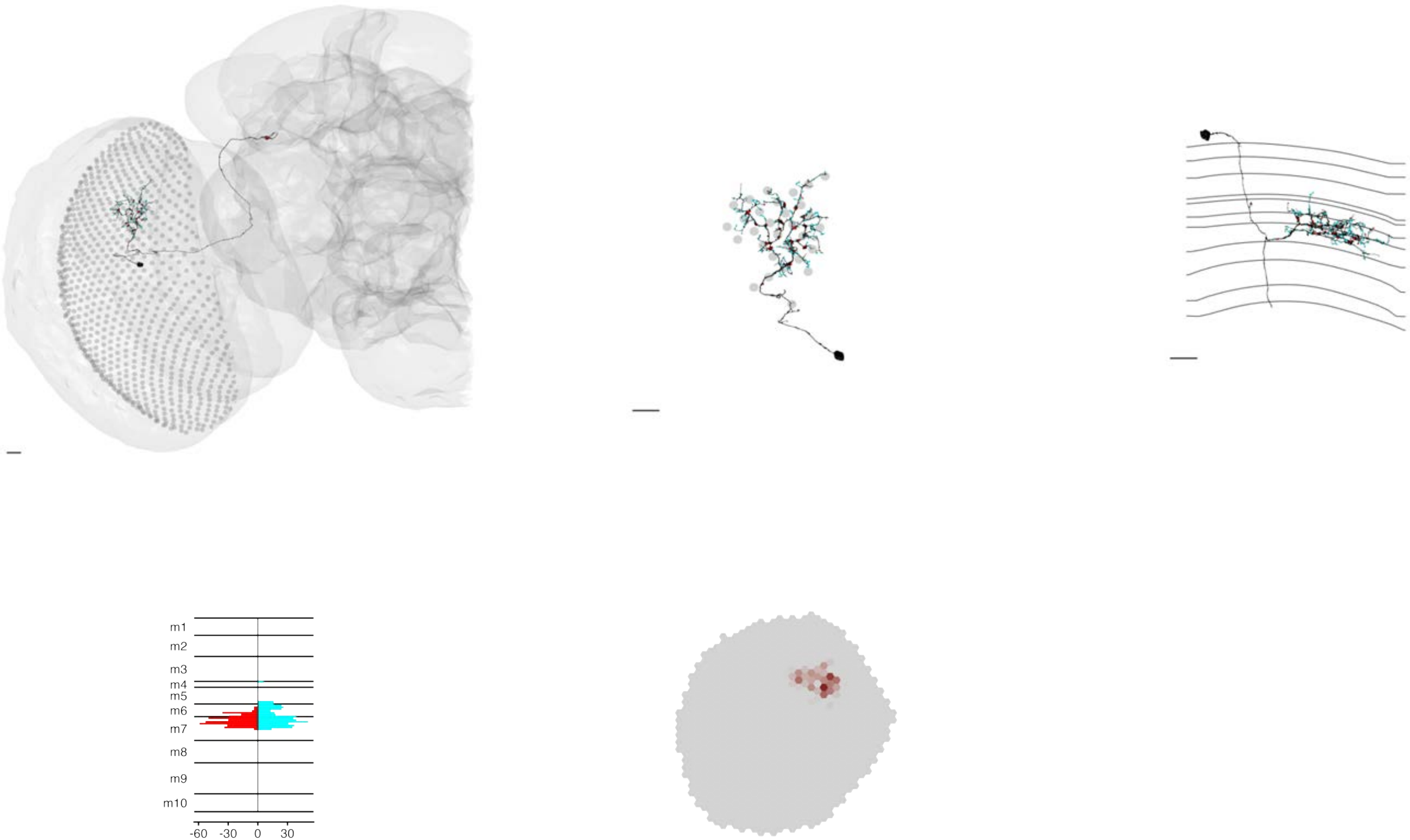

### MeTu3c_dorsal_720575940618767320

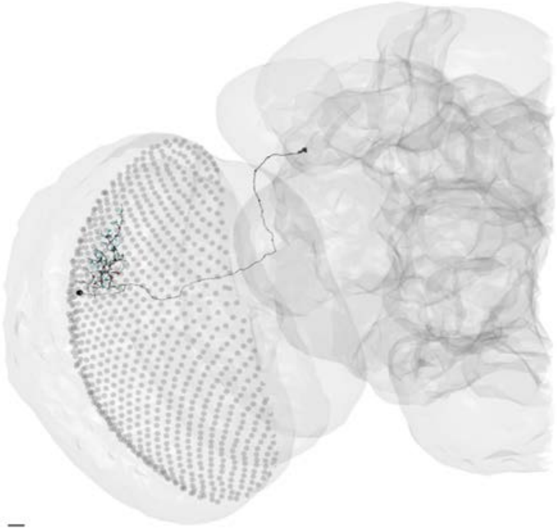

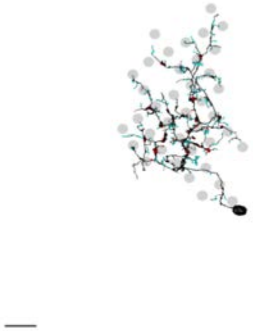

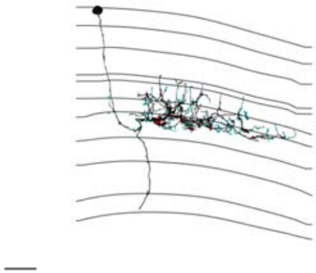

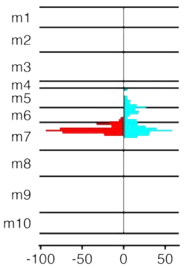


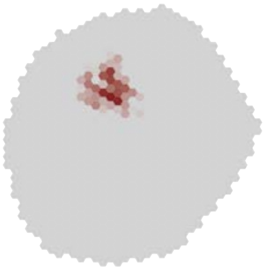

### MeTu3c_dorsal_720575940620048624

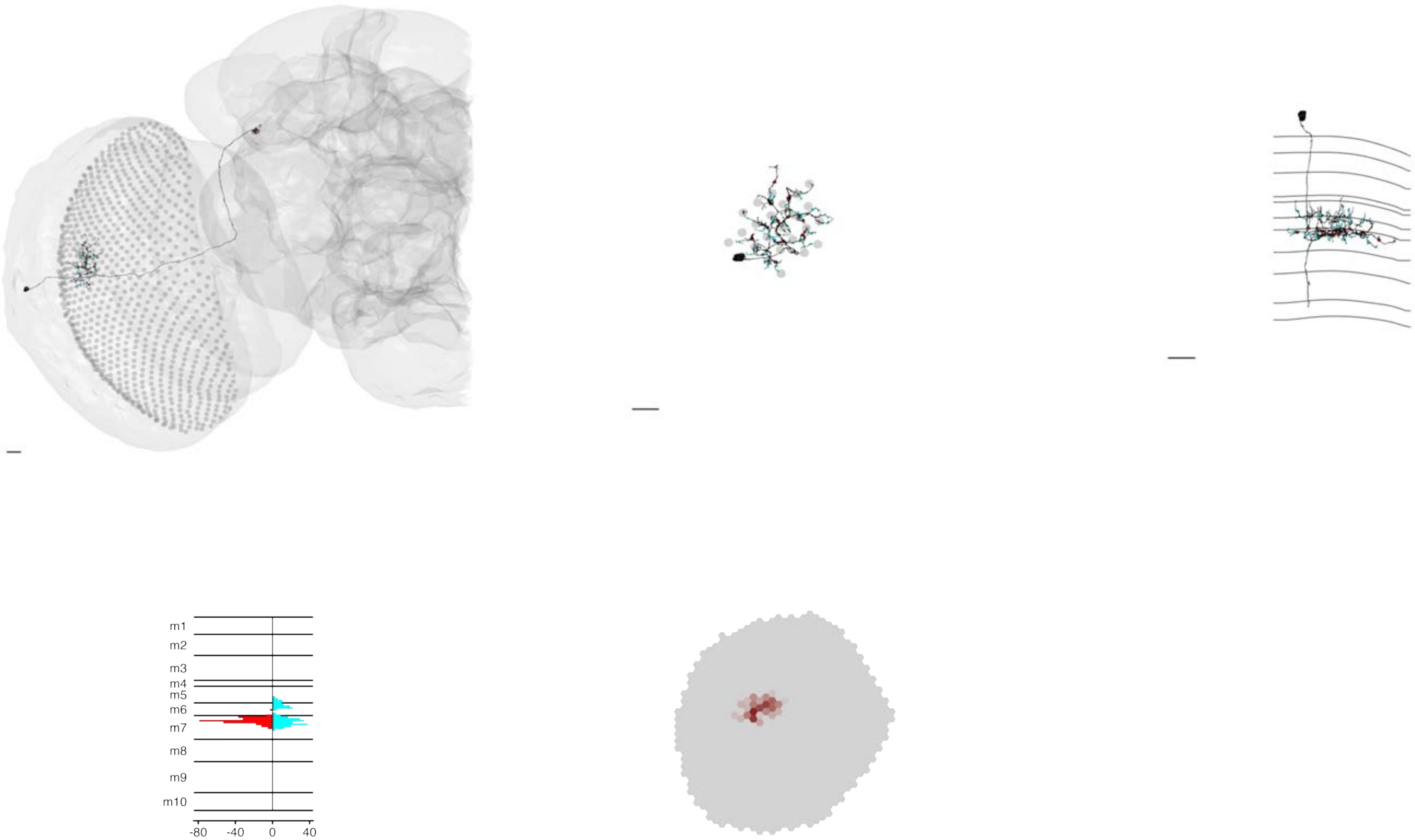

### MeTu3c_dorsal_720575940620826021

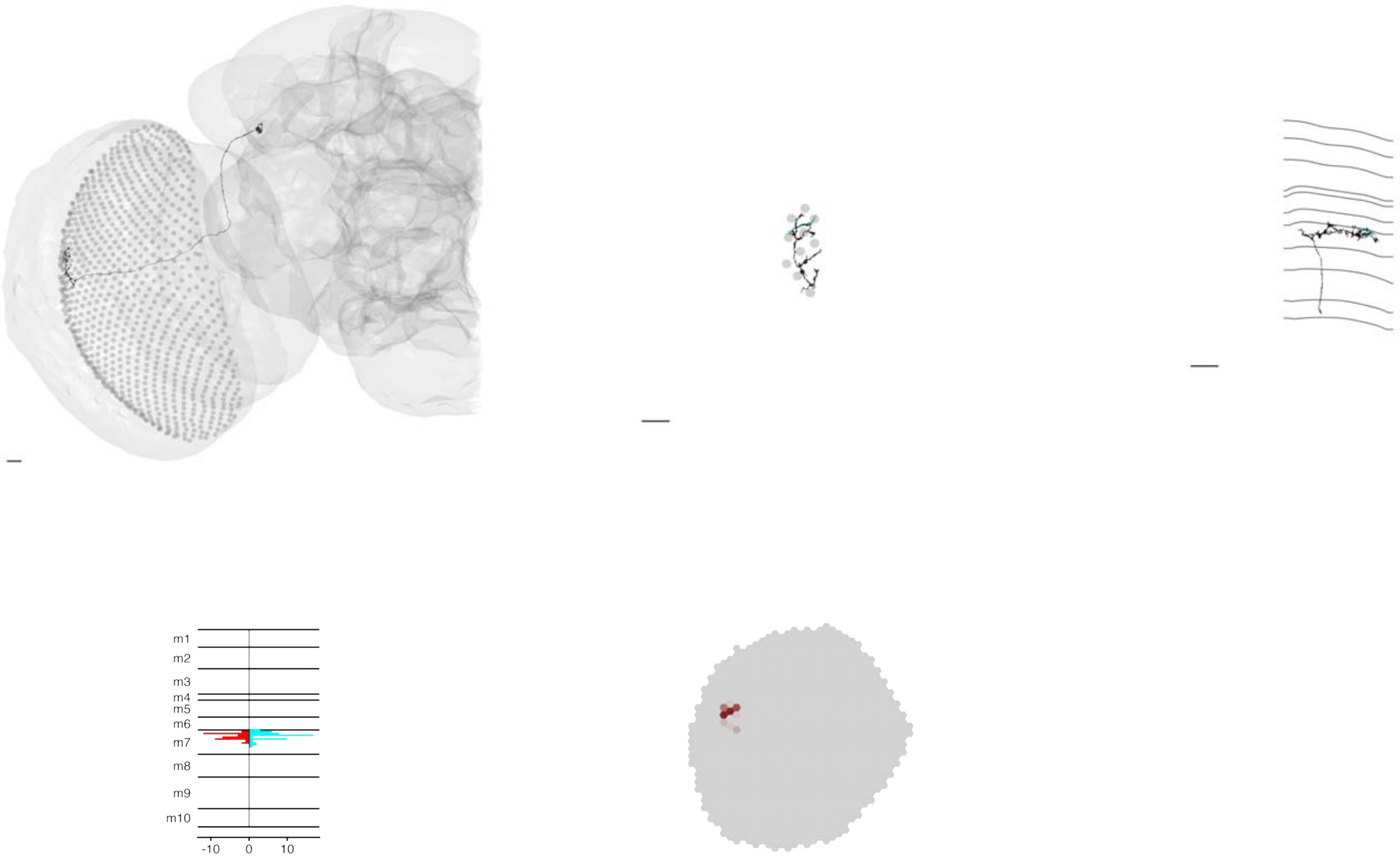

### MeTu3c_dorsal_720575940621023526

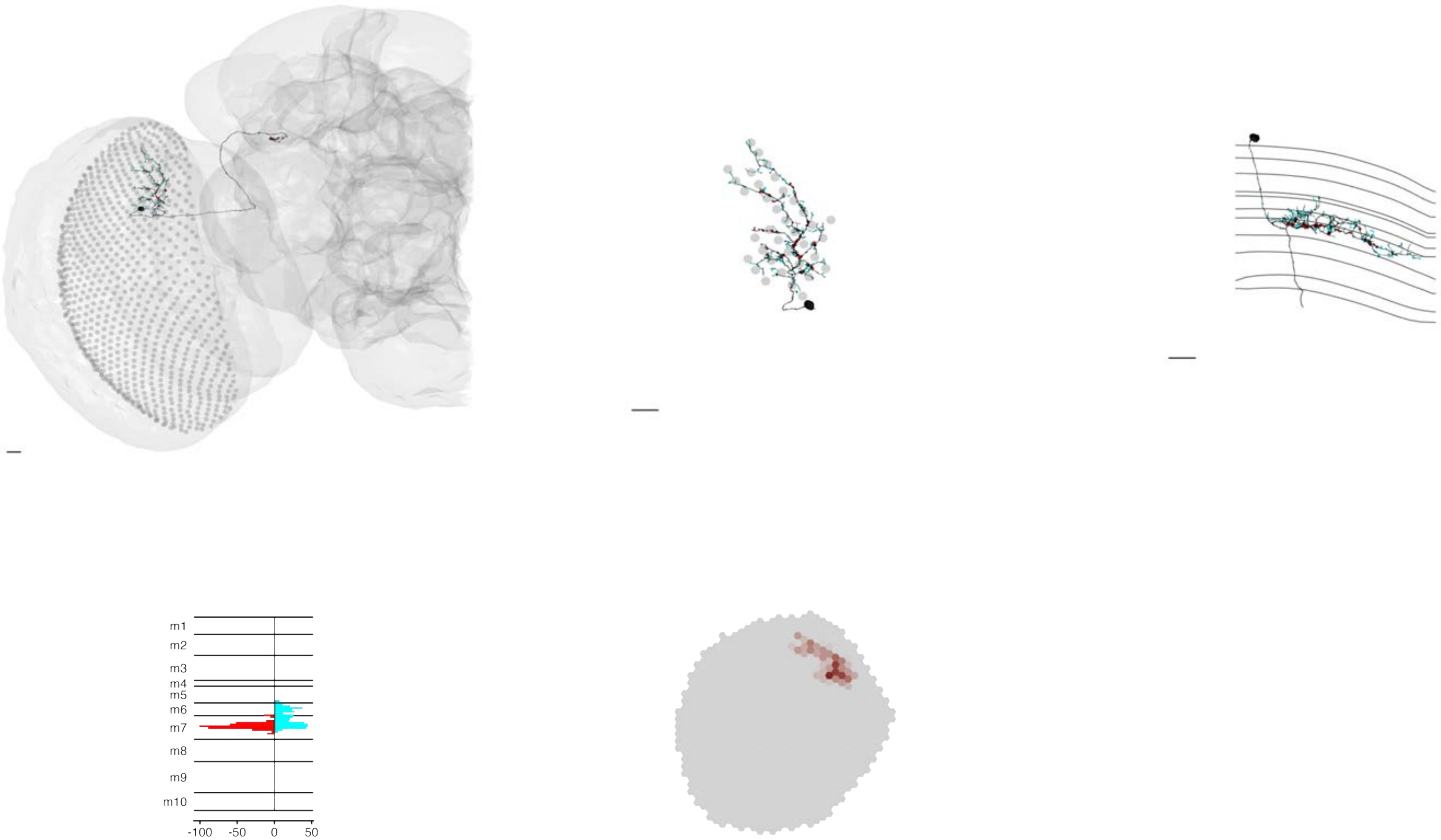

### MeTu3c_dorsal_720575940621487180

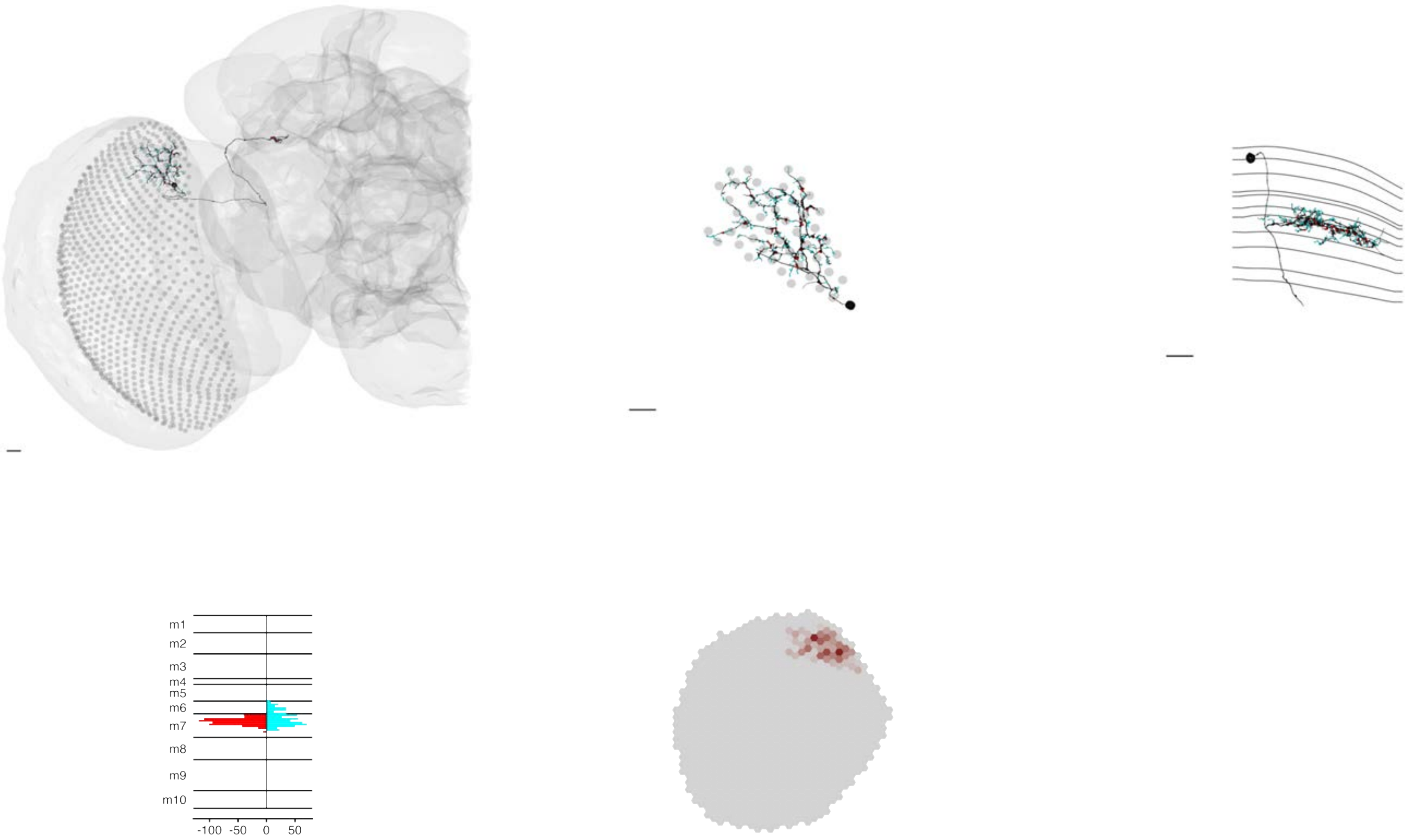

### MeTu3c_dorsal_720575940621529309

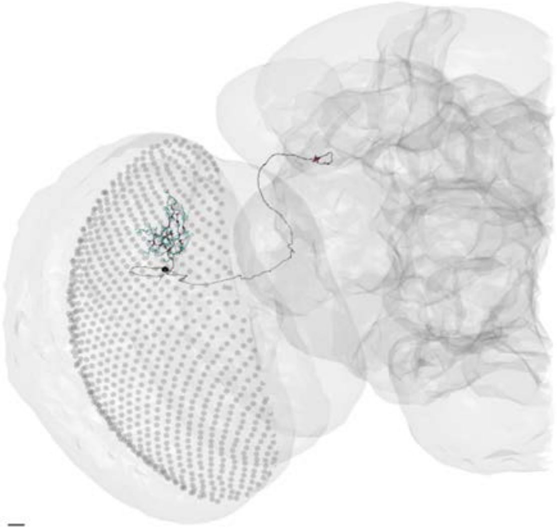

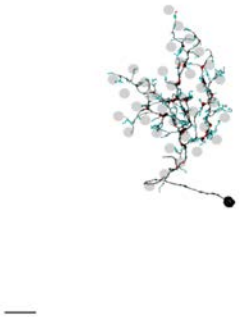

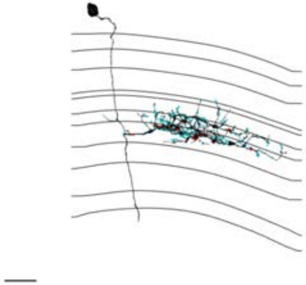

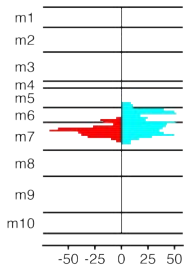


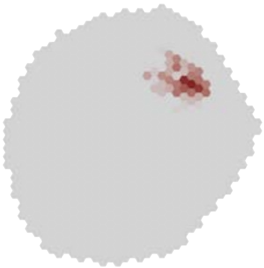

### MeTu3c_dorsal_720575940622137013

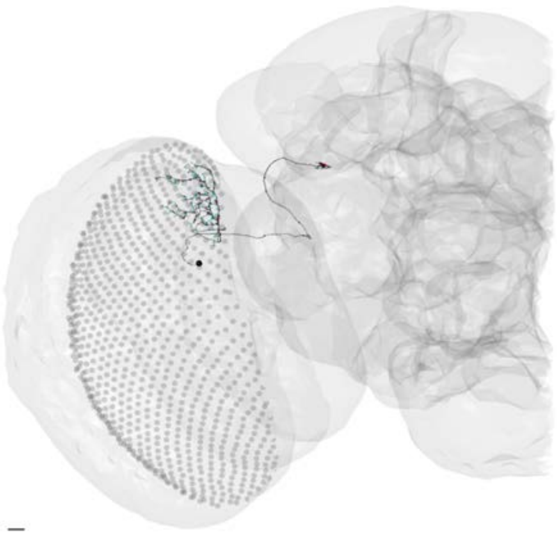

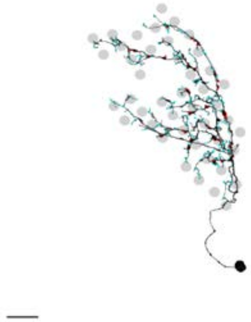

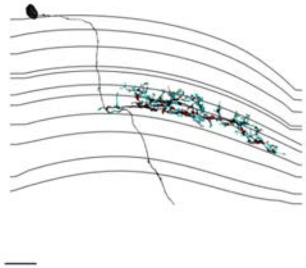

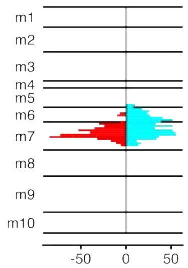


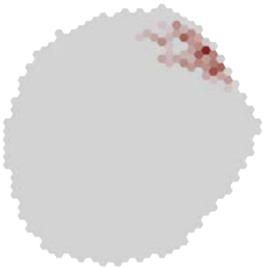

### MeTu3c_dorsal_720575940623010124

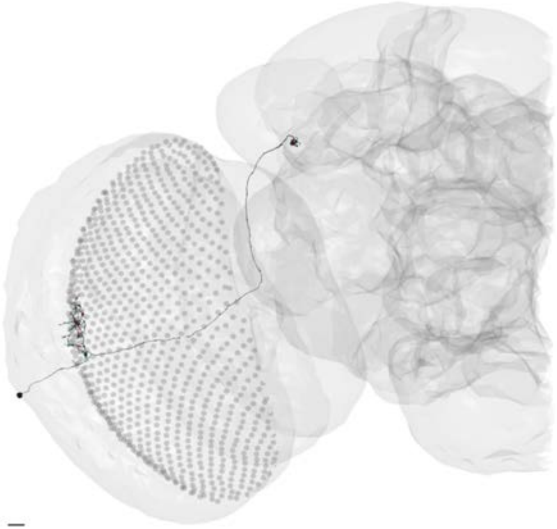

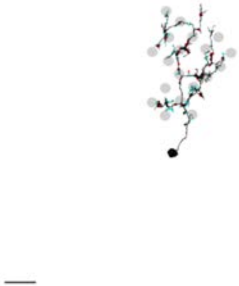

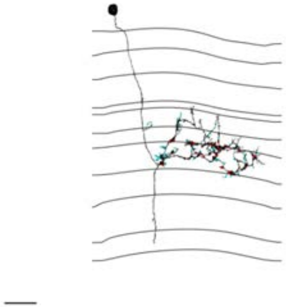

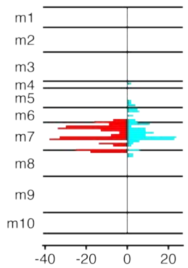


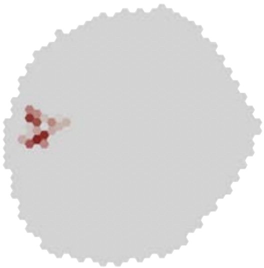

### MeTu3c_dorsal_720575940623380714

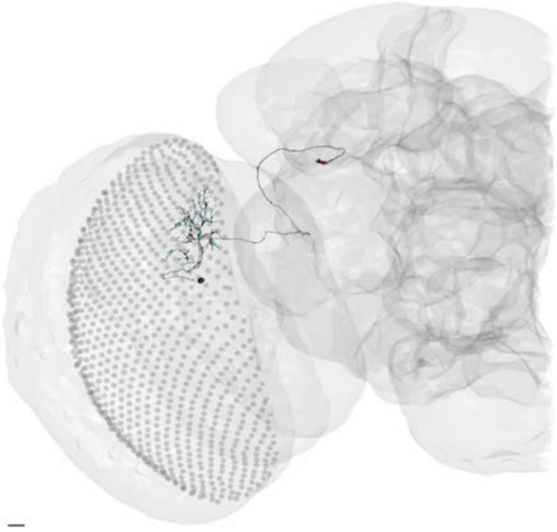

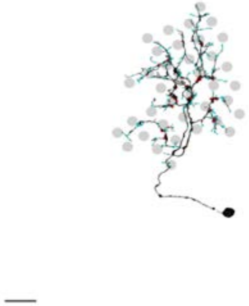

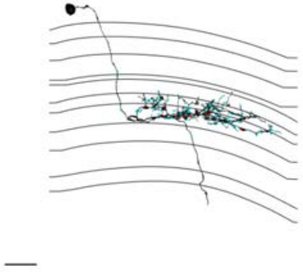

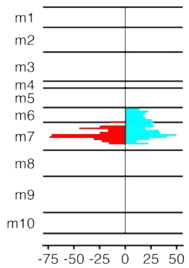


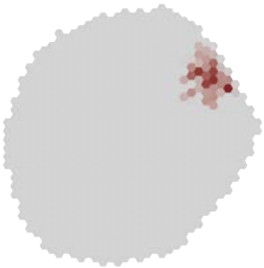

### MeTu3c_dorsal_720575940623886055

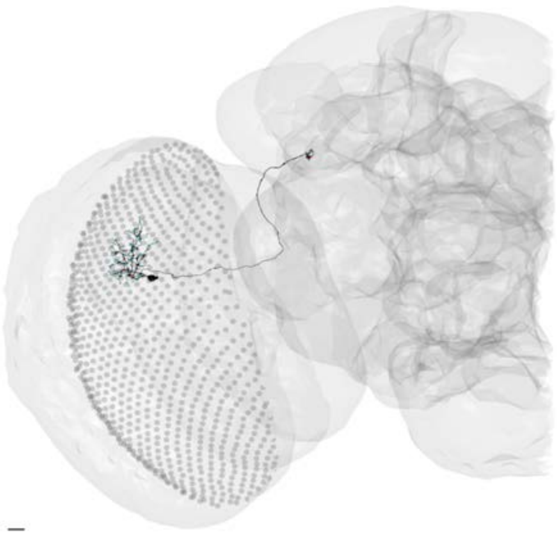

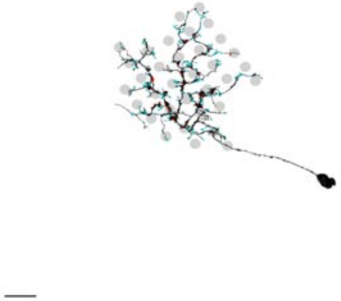

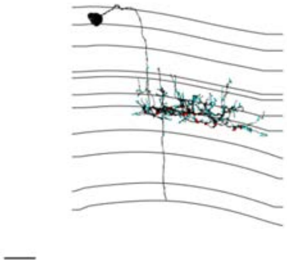

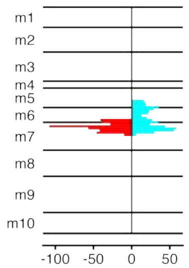


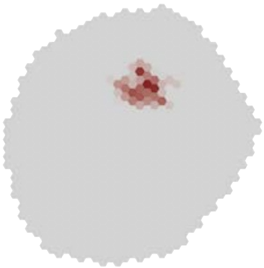

MeTu3c_dorsal_720575940624169653

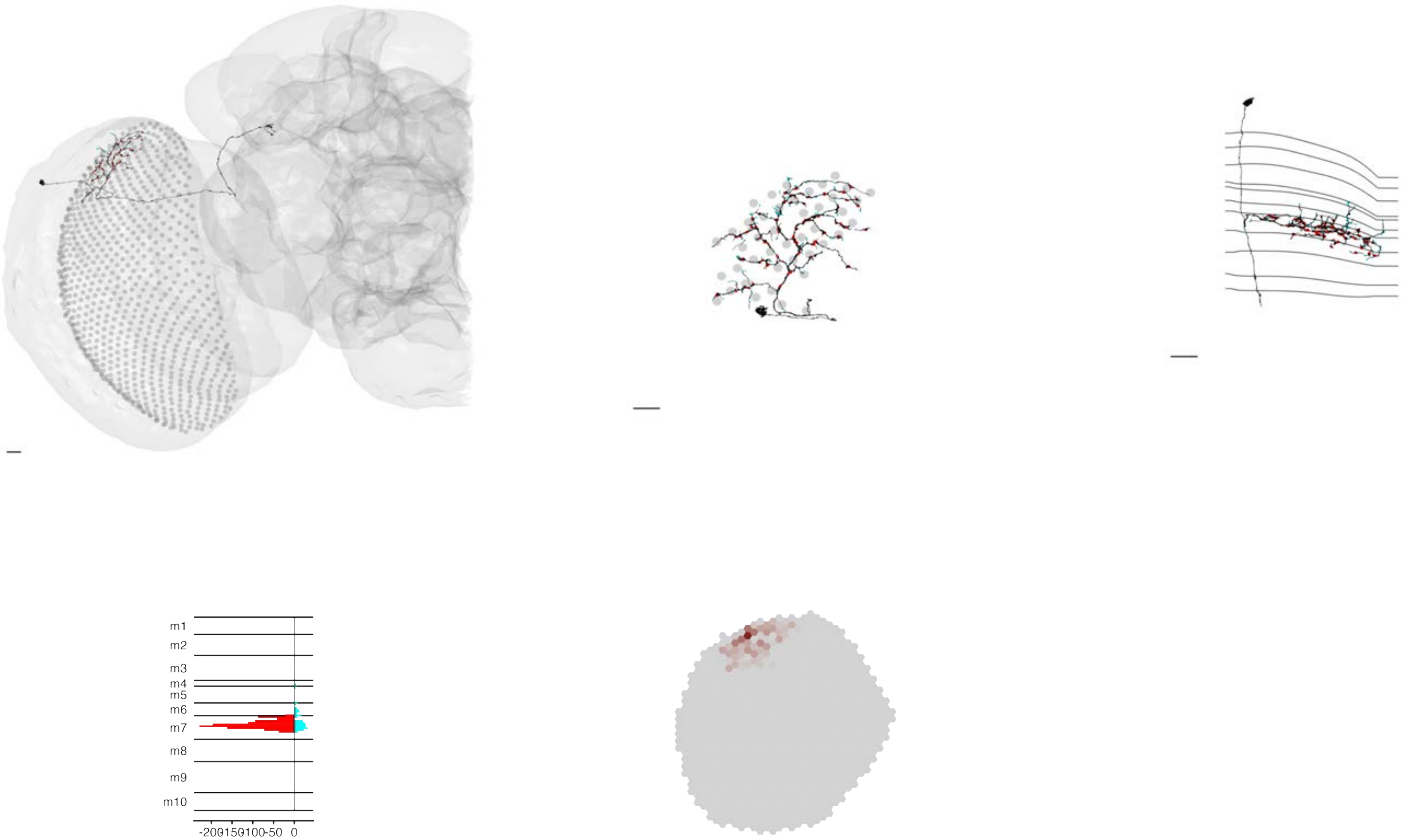

### MeTu3c_dorsal_720575940625838594

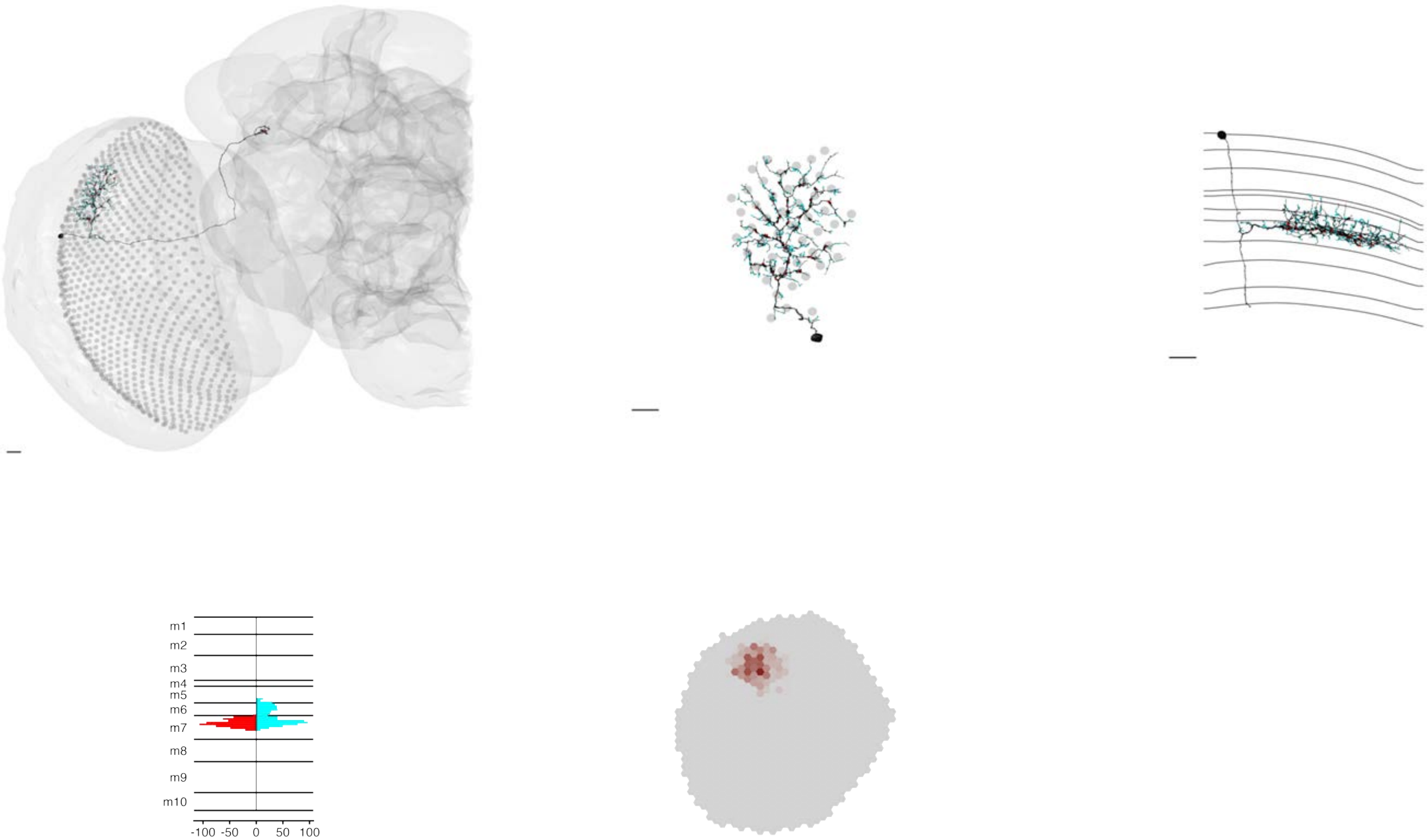

### MeTu3c_dorsal_720575940625877178

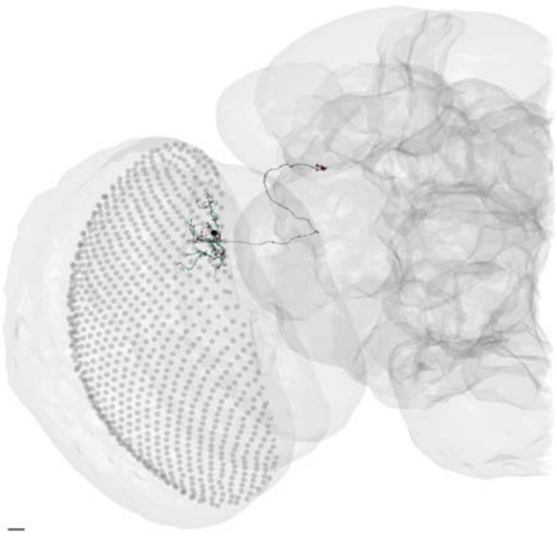

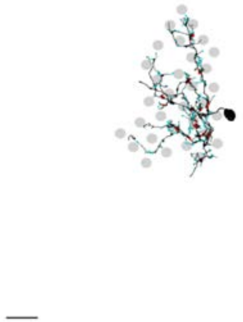

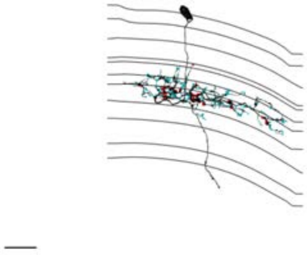

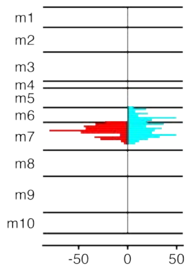


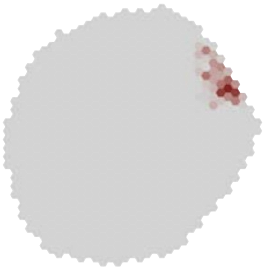

MeTu3c_dorsal_720575940626327112

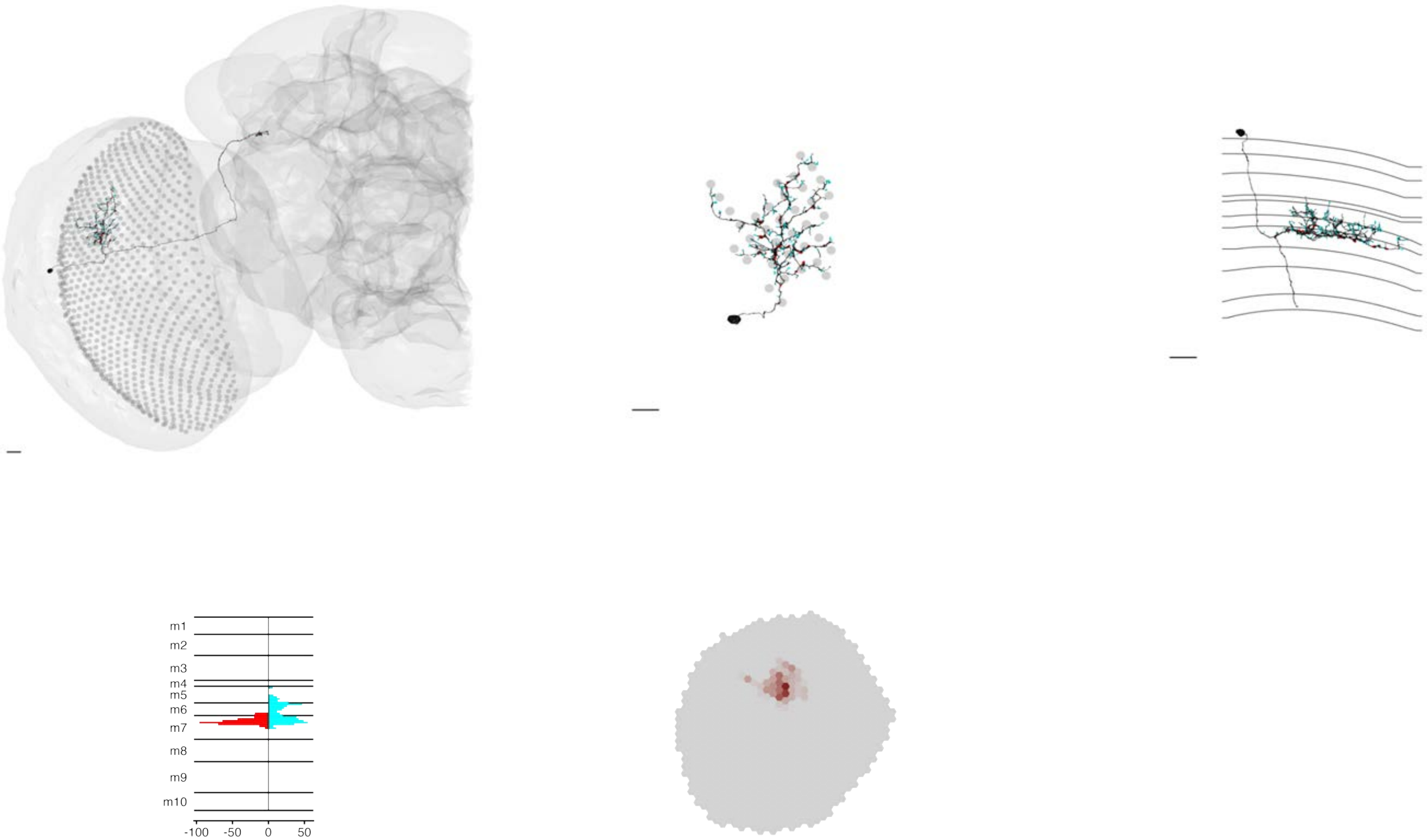

### MeTu3c_dorsal_720575940627933946

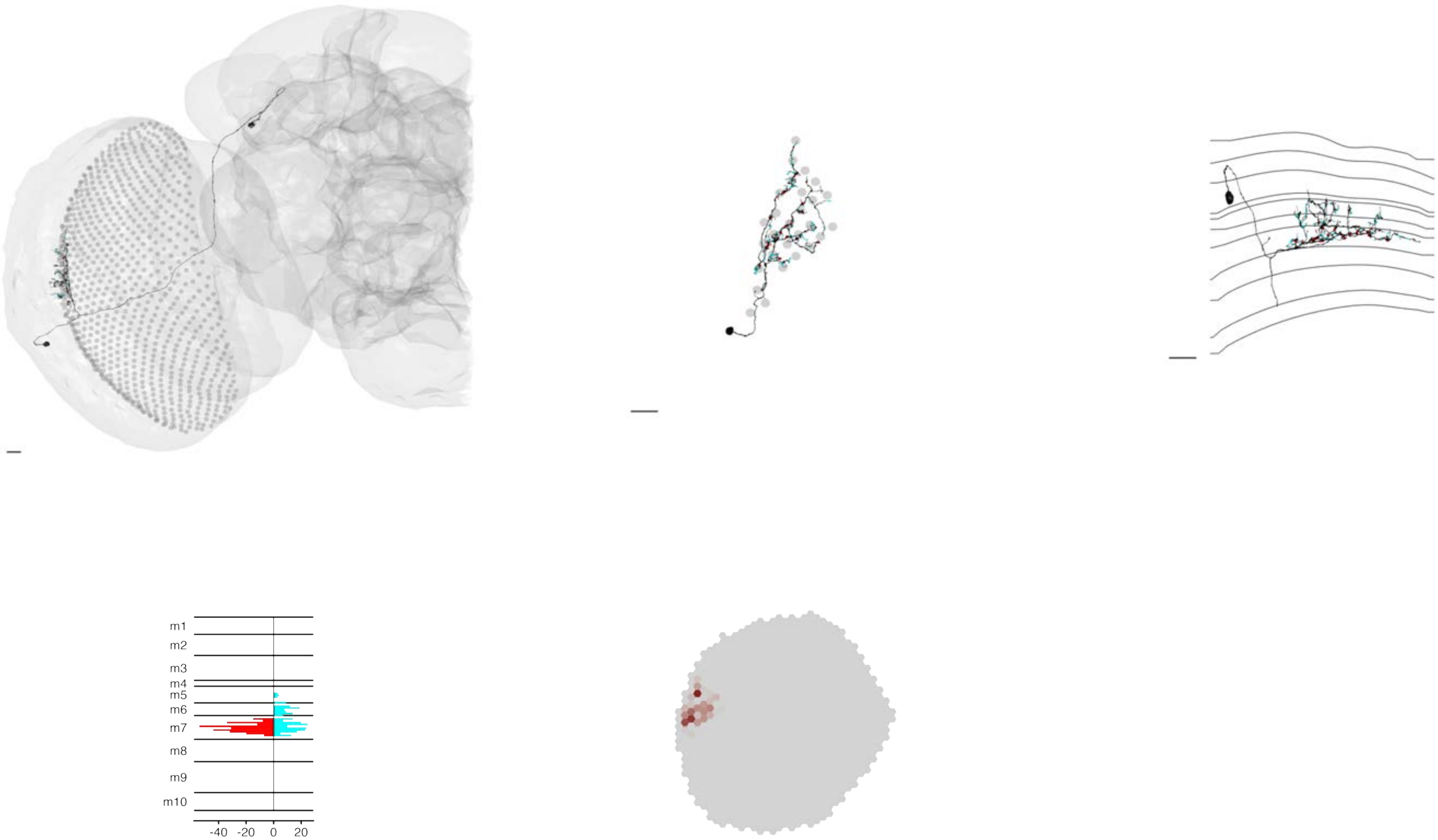

### MeTu3c_dorsal_720575940629254204

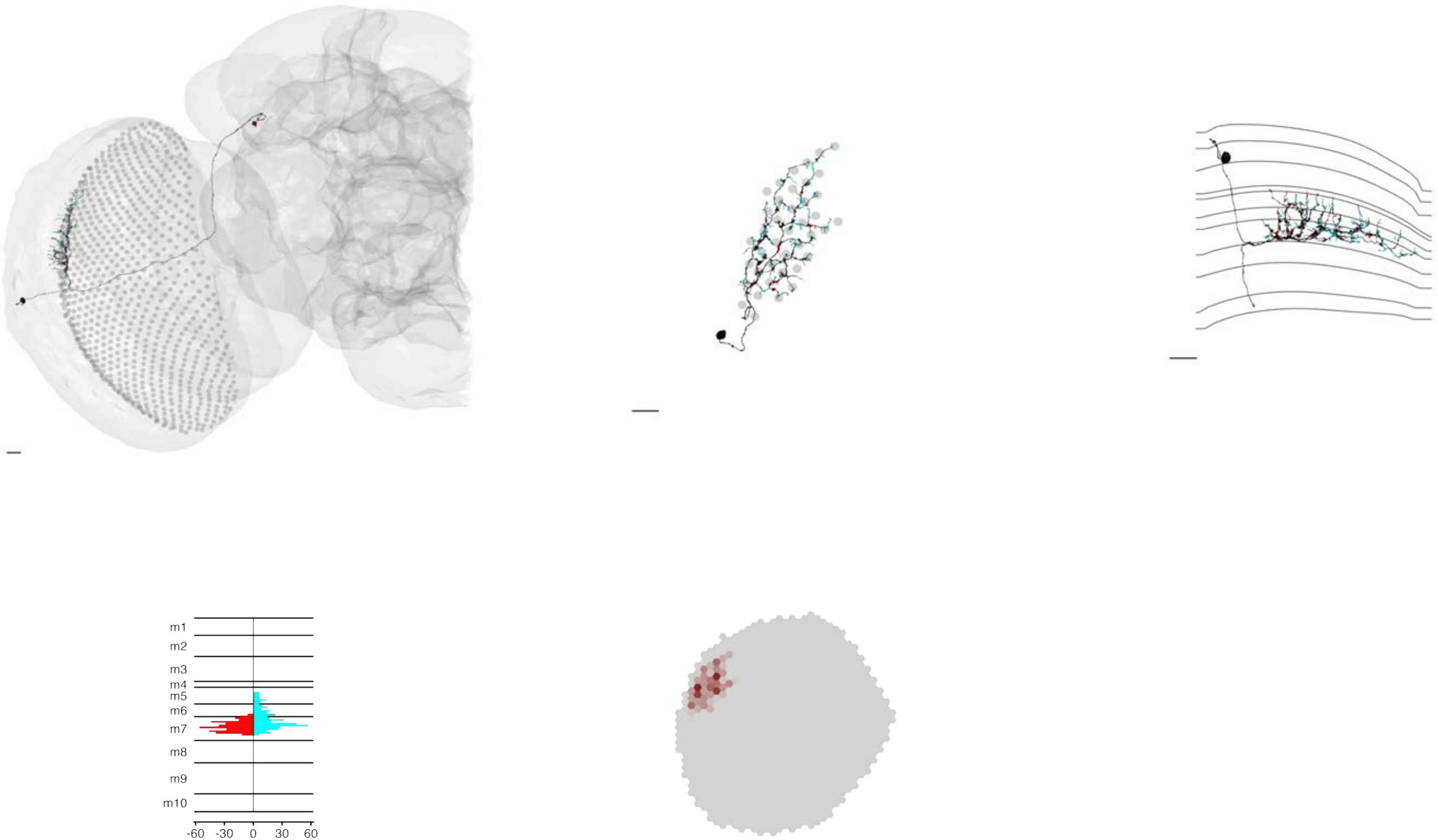

### MeTu3c_dorsal_720575940630603587

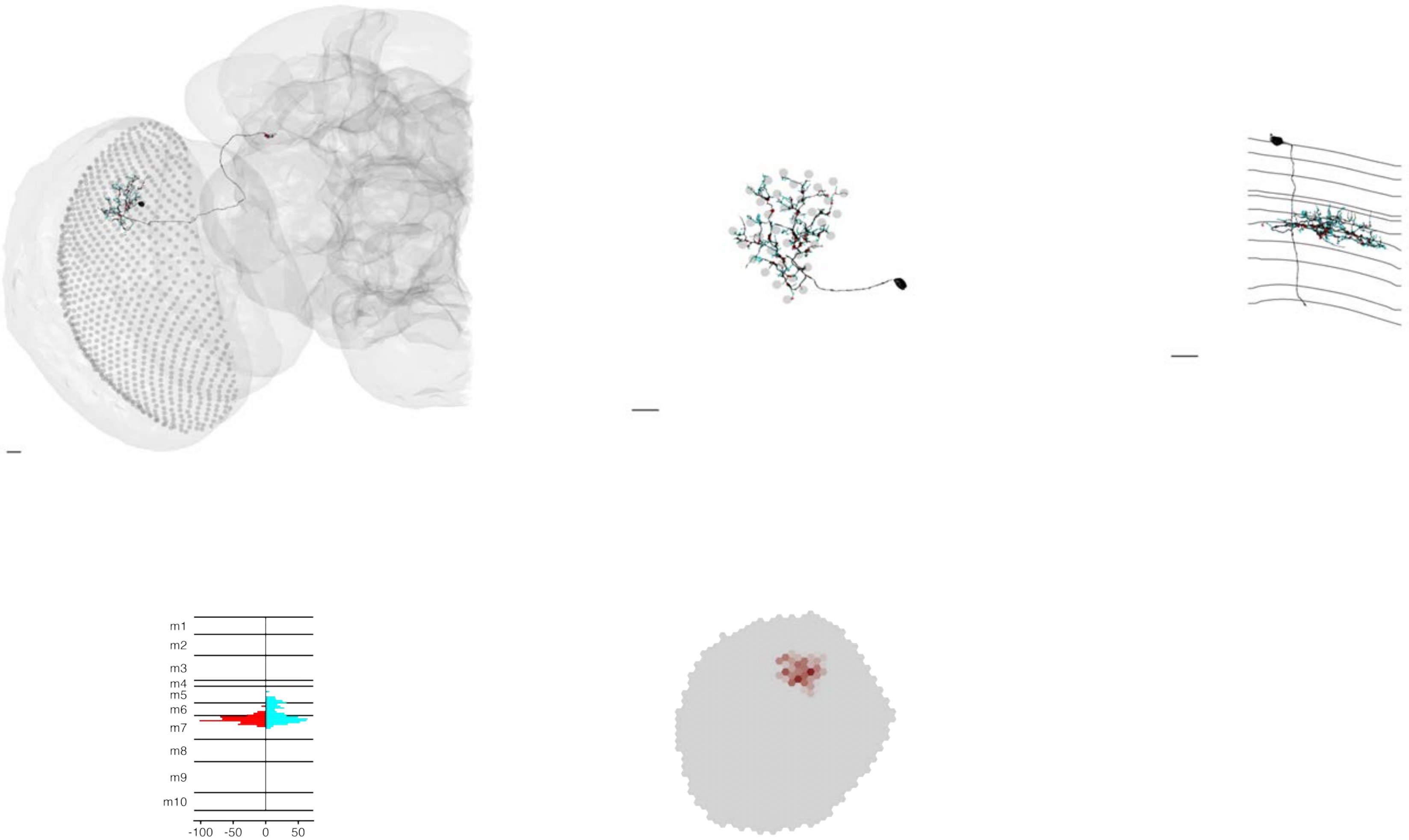

### MeTu3c_dorsal_720575940630888223

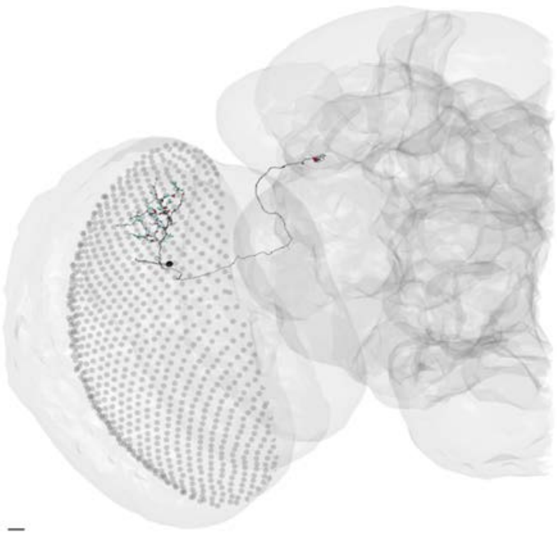

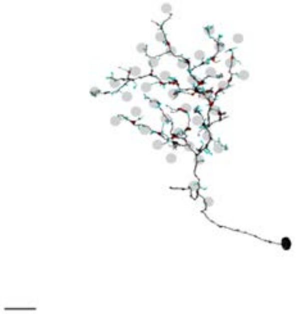

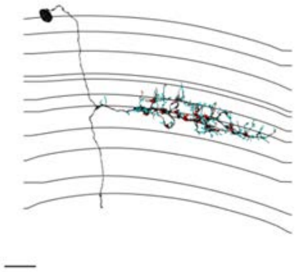

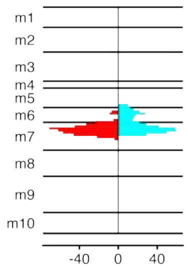


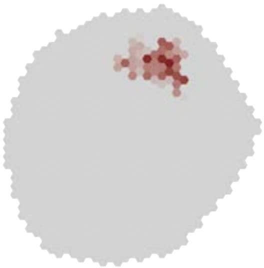

### MeTu3c_dorsal_720575940633871315

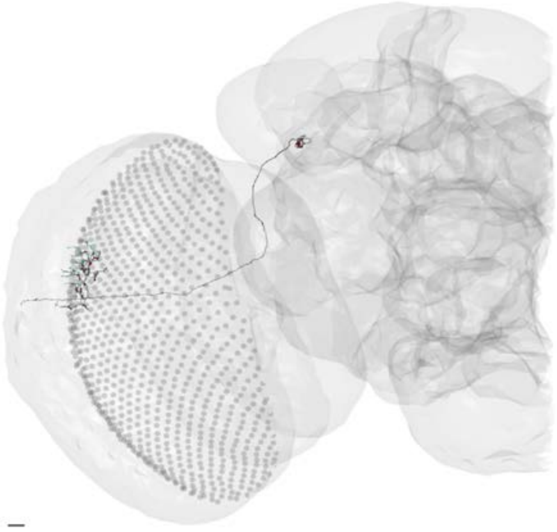

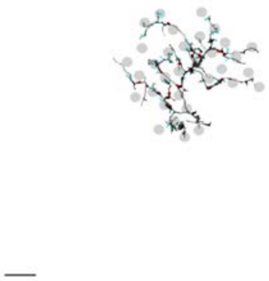

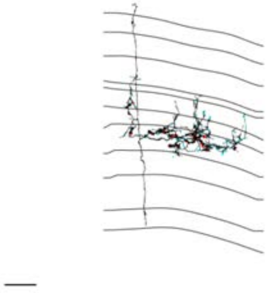

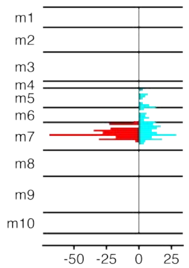


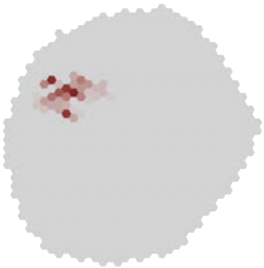

### MeTu3c_dorsal_720575940637486670

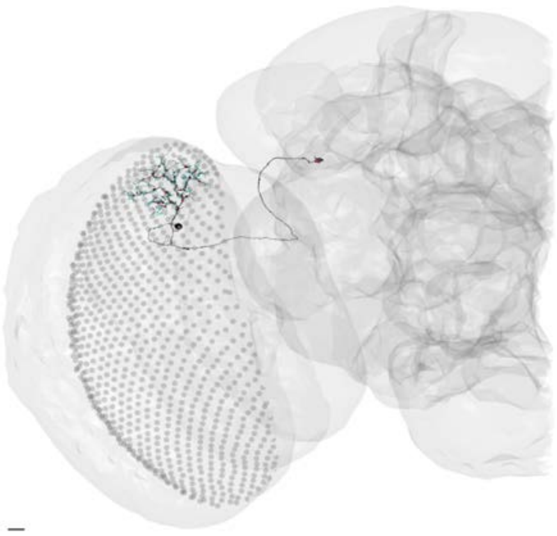

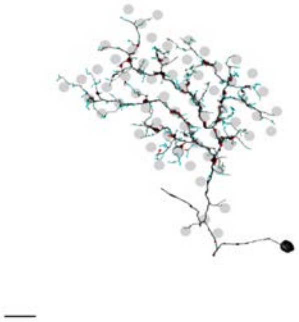

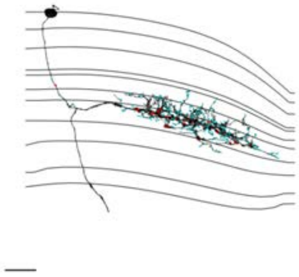

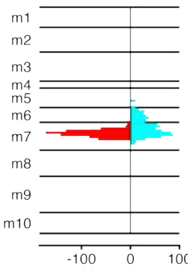


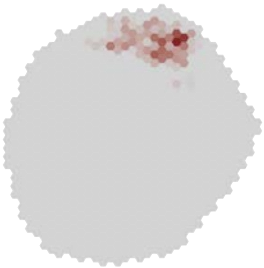

### MeTu3c_dorsal_720575940637959539

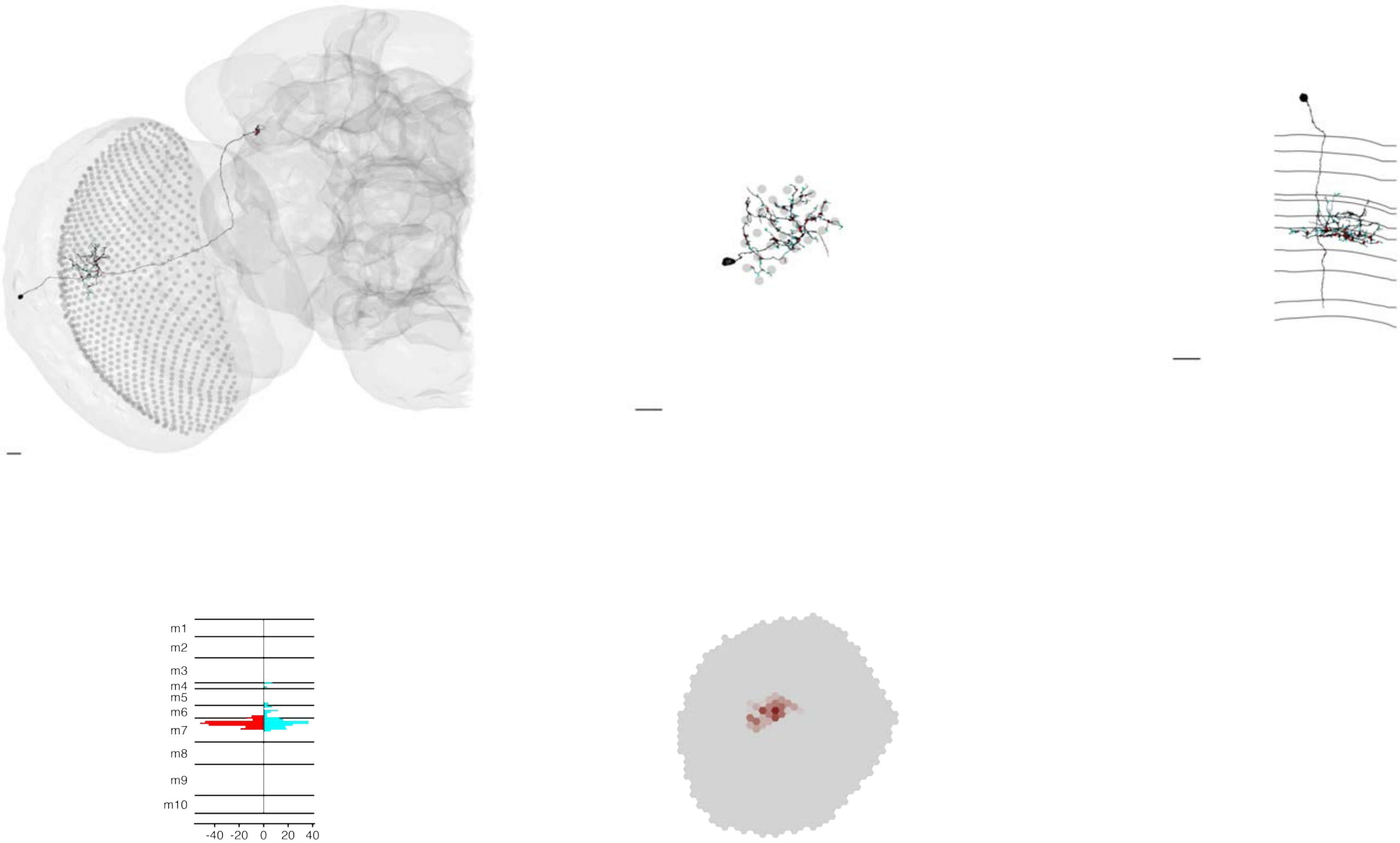

MeTu3c_dorsal_720575940641978101

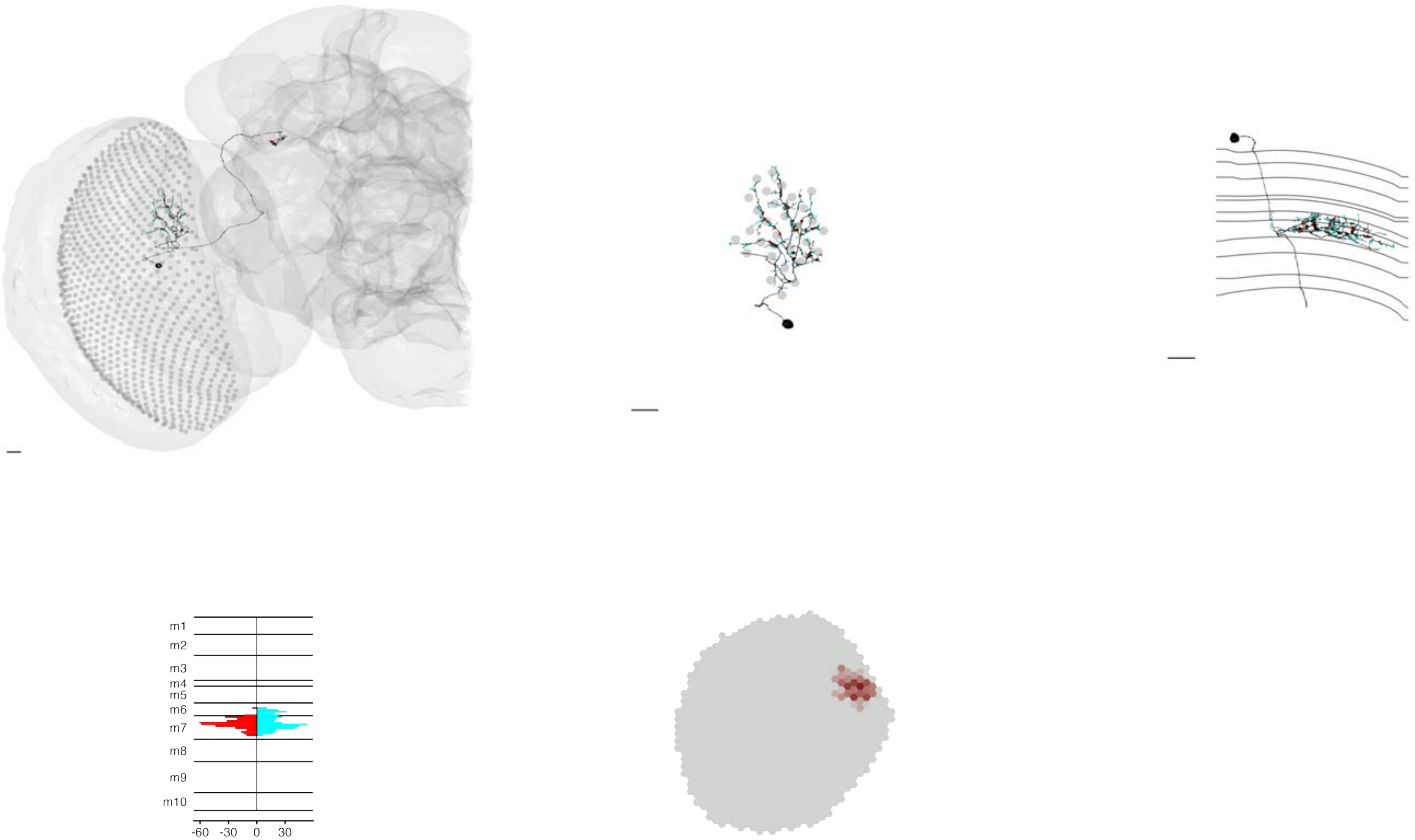

### MeTu3c_dorsal_720575940655446689

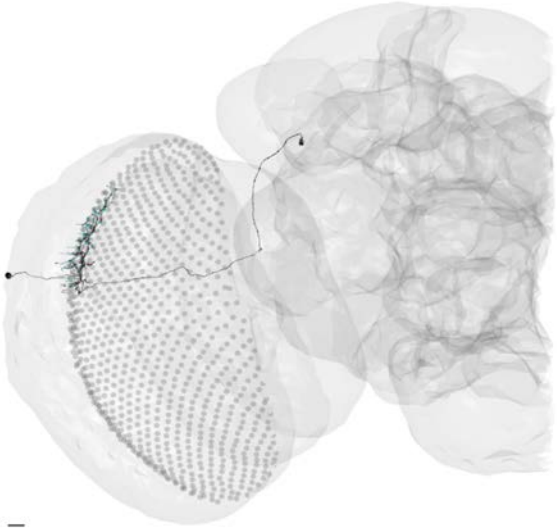

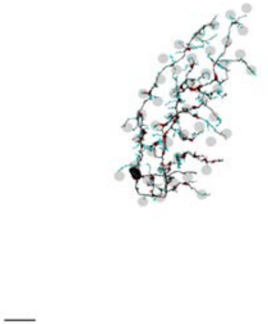

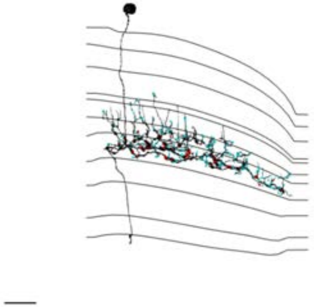

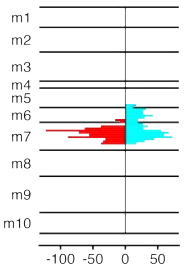


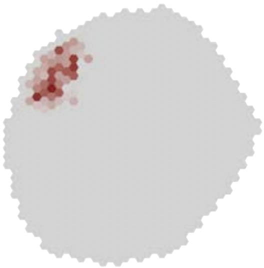

MeTu3c_ventral_720575940602996448

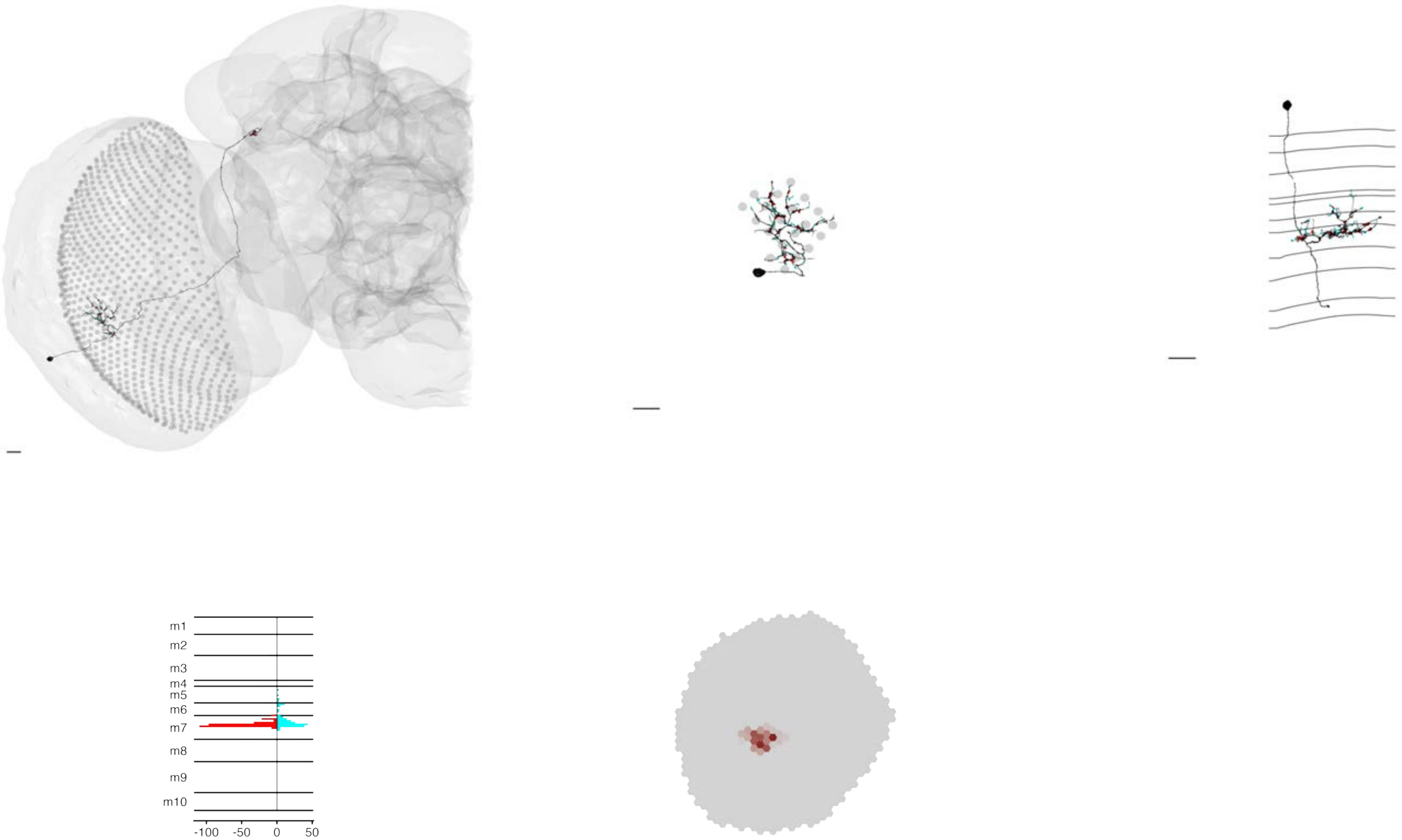

MeTu3c_ventral_720575940606044448

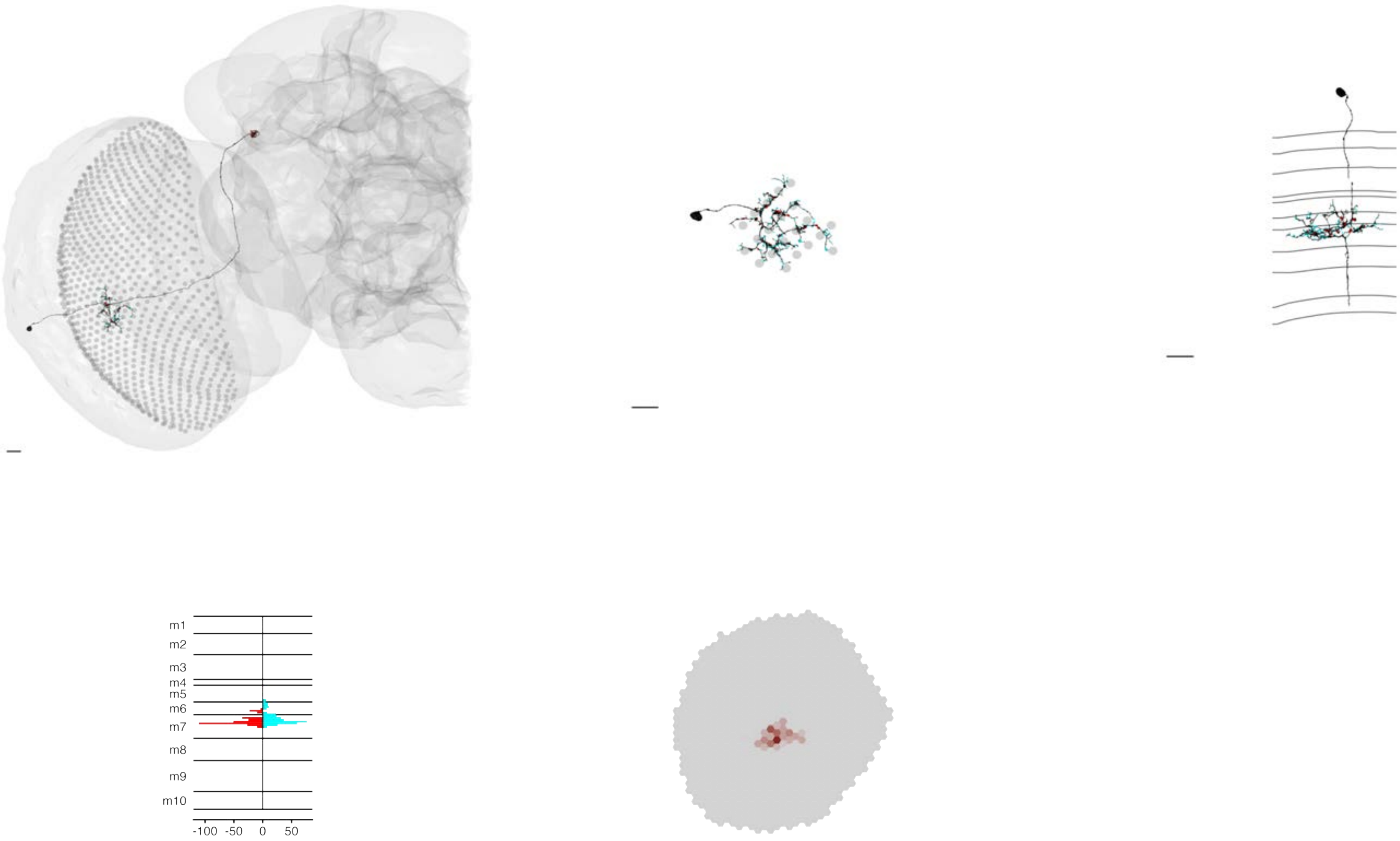

### MeTu3c_ventral_720575940609524946

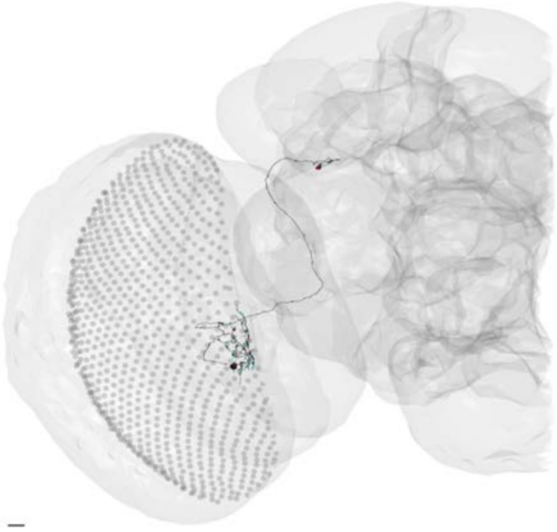

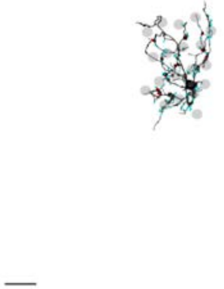

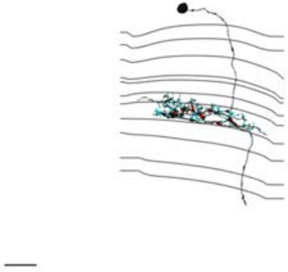

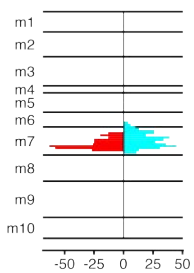


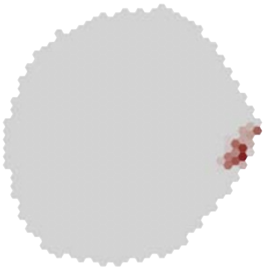

### MeTu3c_ventral_720575940610352590

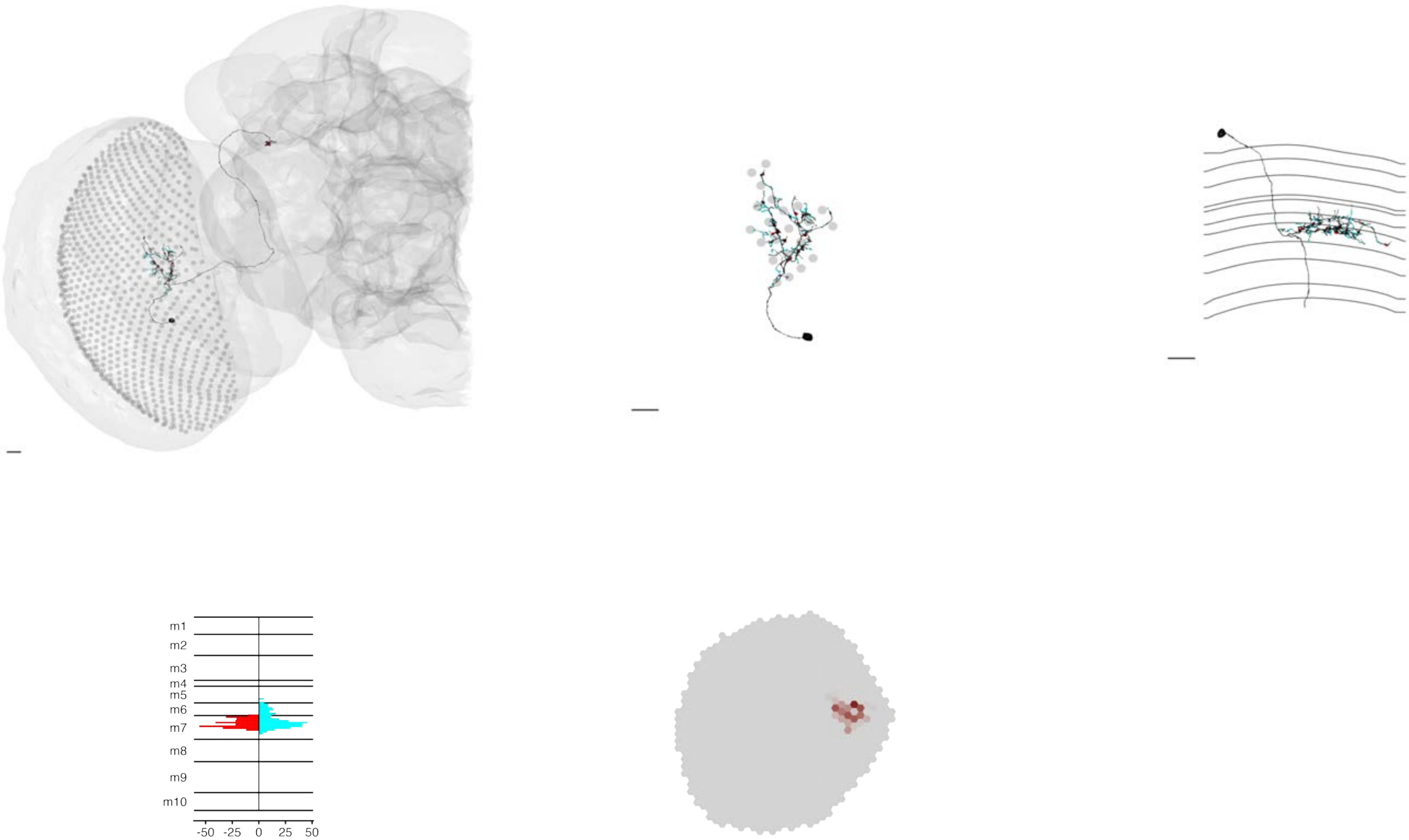

### MeTu3c_ventral_720575940611194390

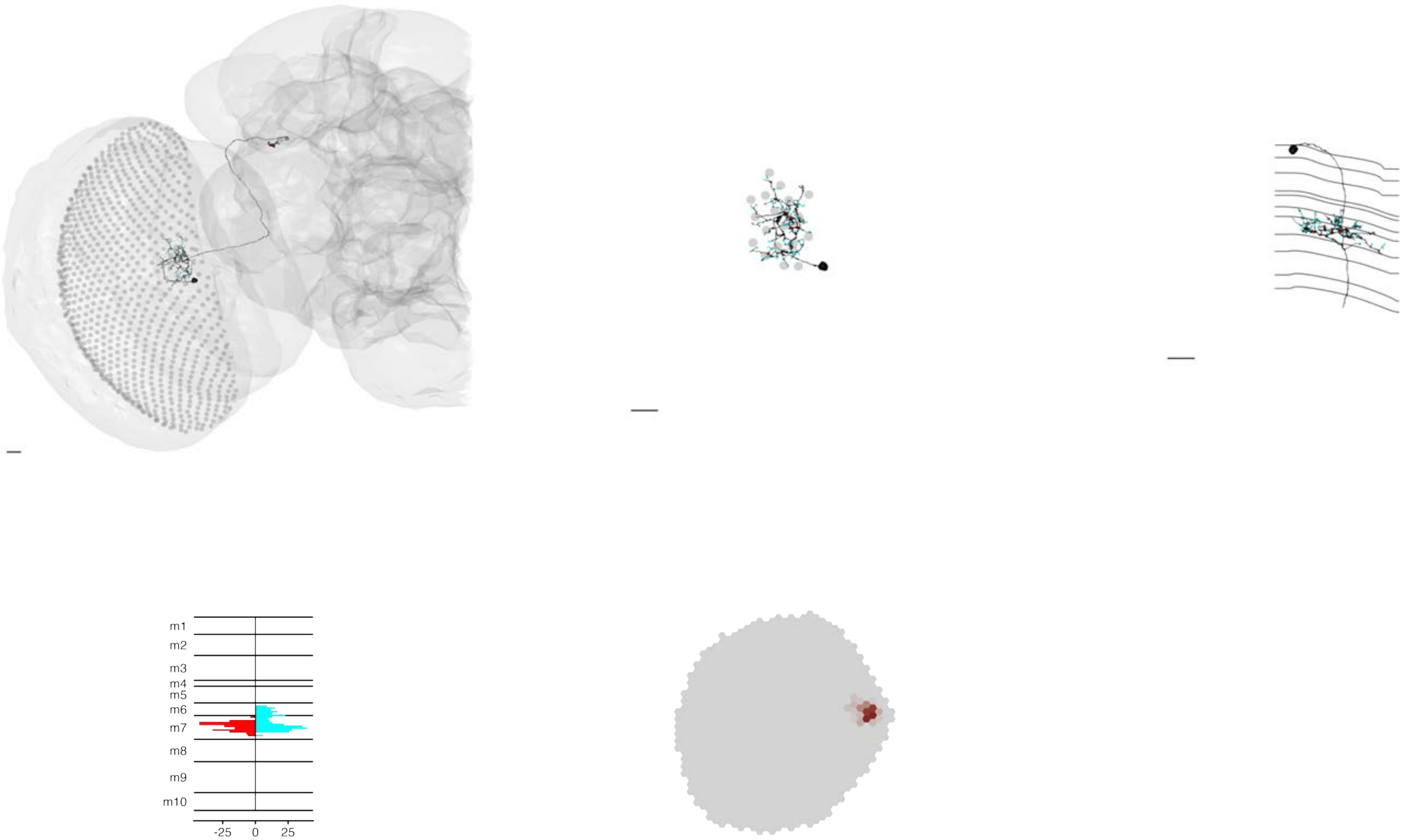

### MeTu3c_ventral_720575940612333462

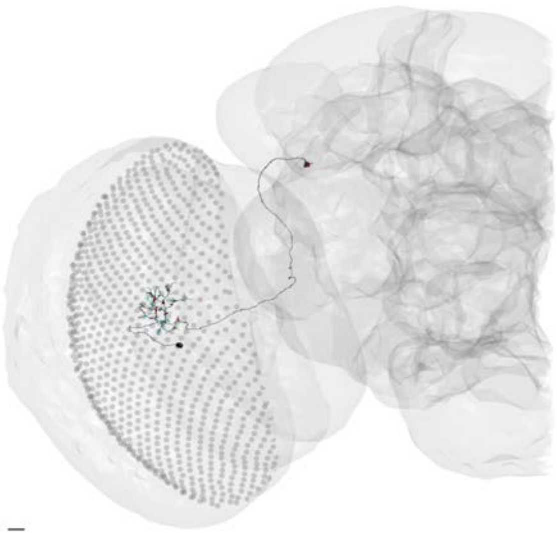

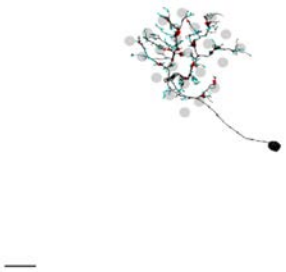

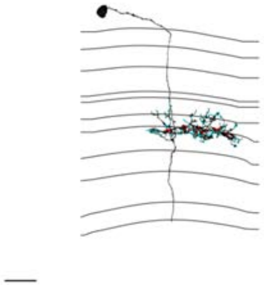

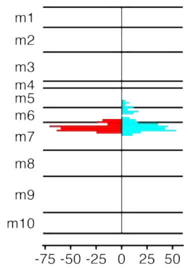


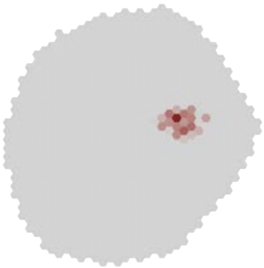

### MeTu3c_ventral_720575940612378152

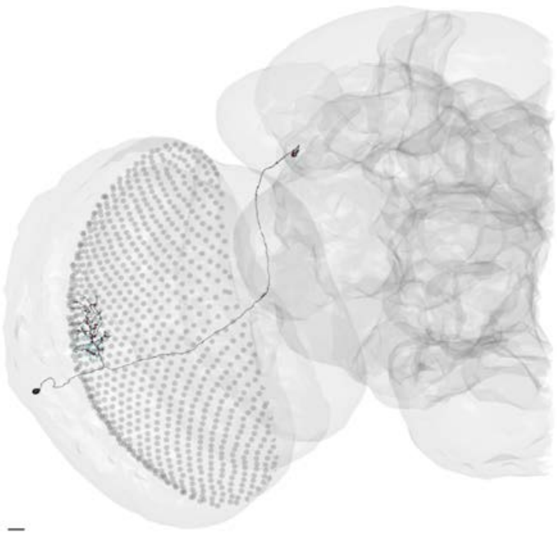

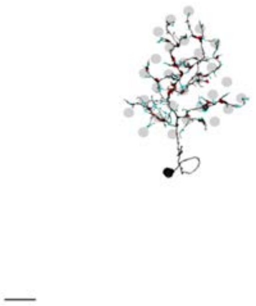

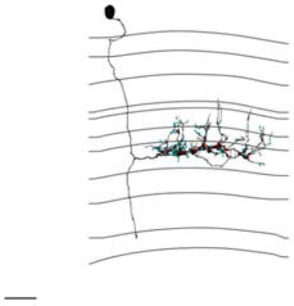

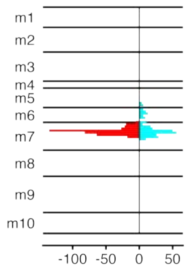


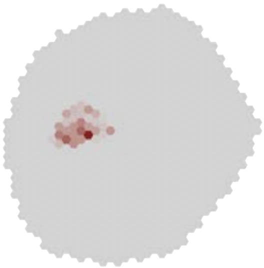

### MeTu3c_ventral_720575940613692414

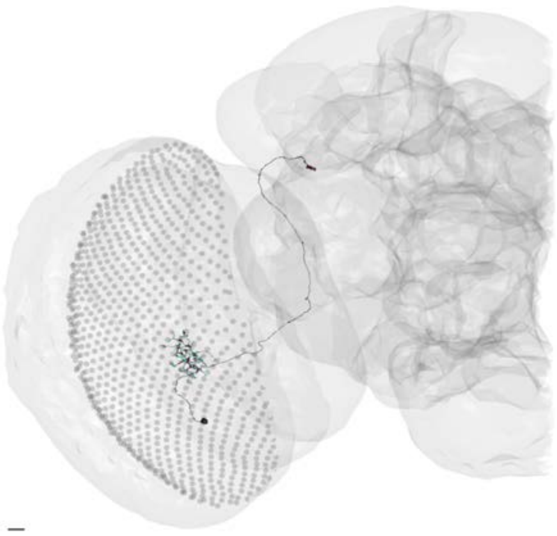

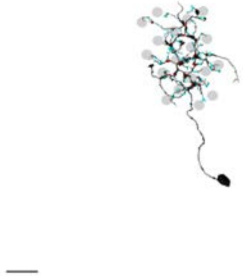

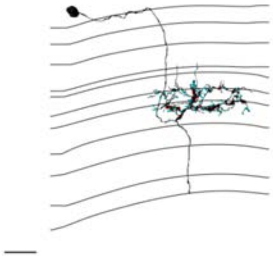

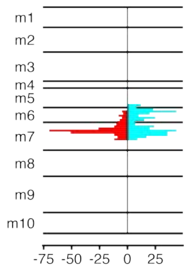


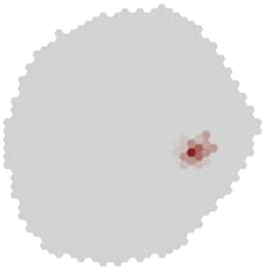

### MeTu3c_ventral_720575940615518870

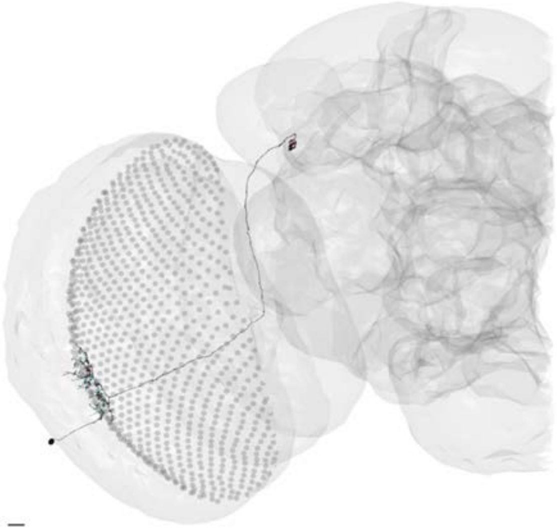

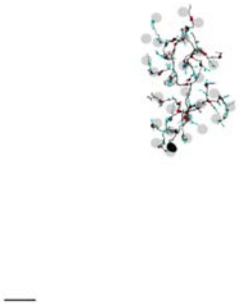

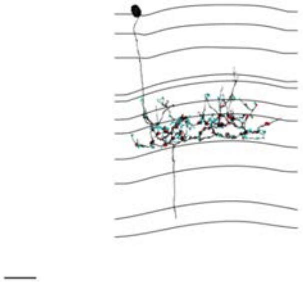

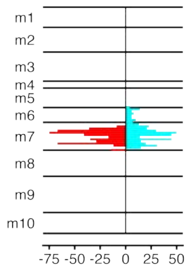


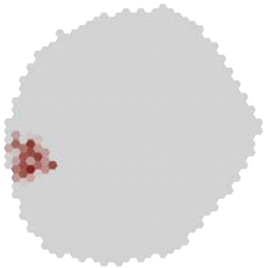

MeTu3c_ventral_720575940617087718

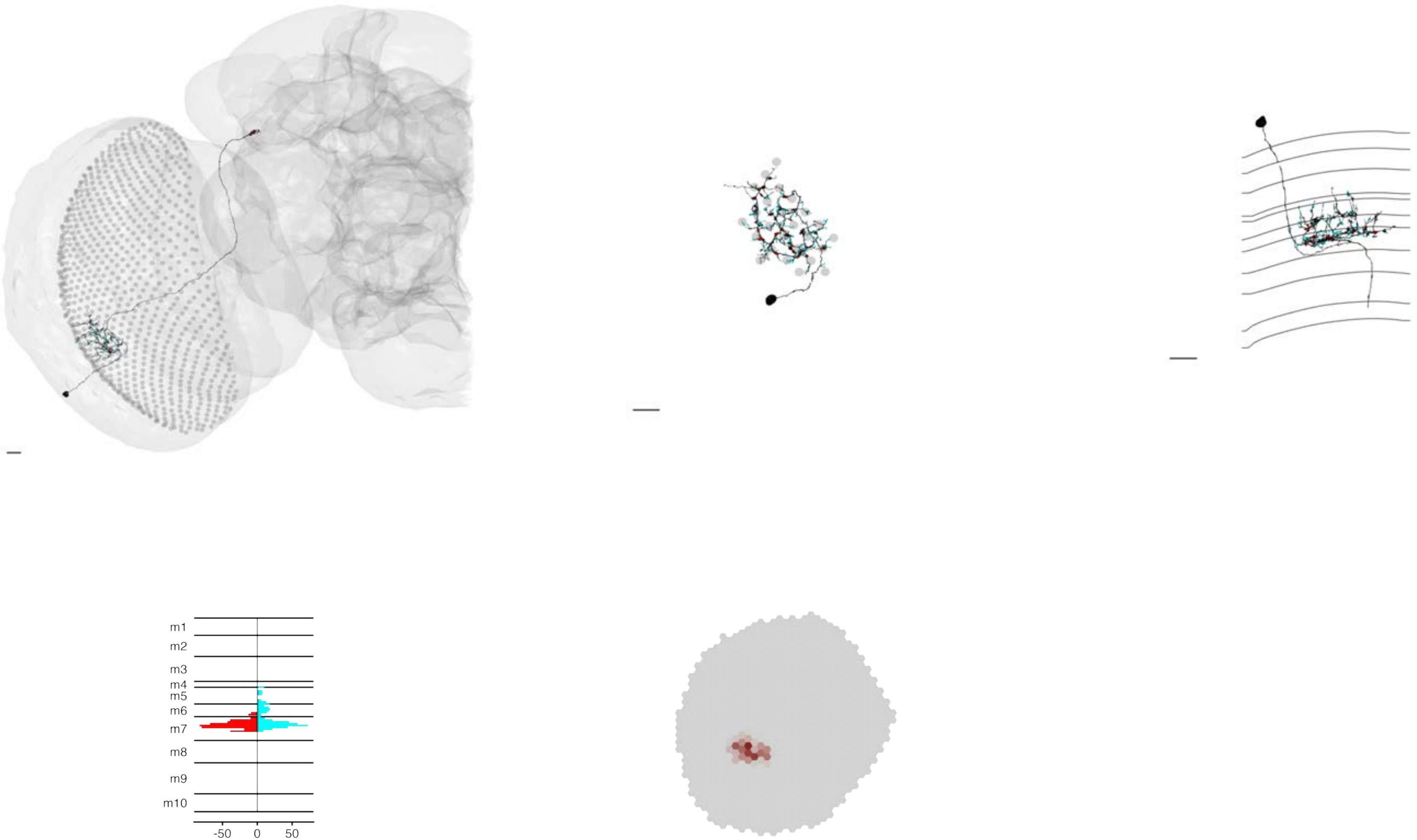

### MeTu3c_ventral_720575940617688866

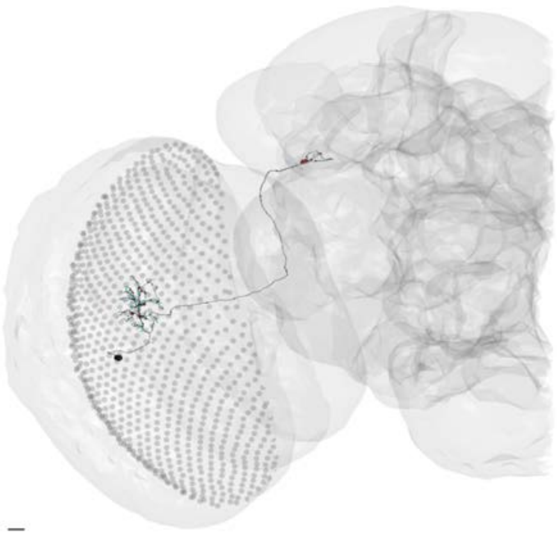

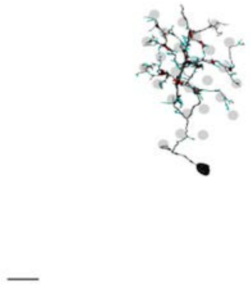

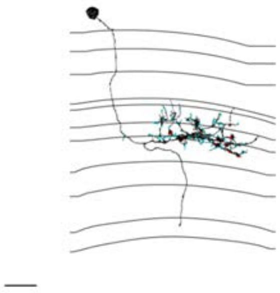

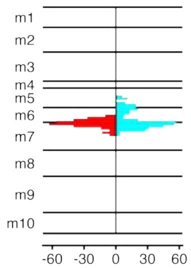


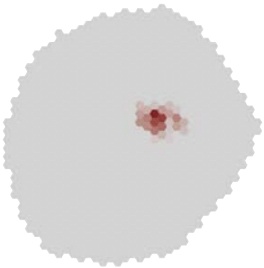

### MeTu3c_ventral_720575940618486181

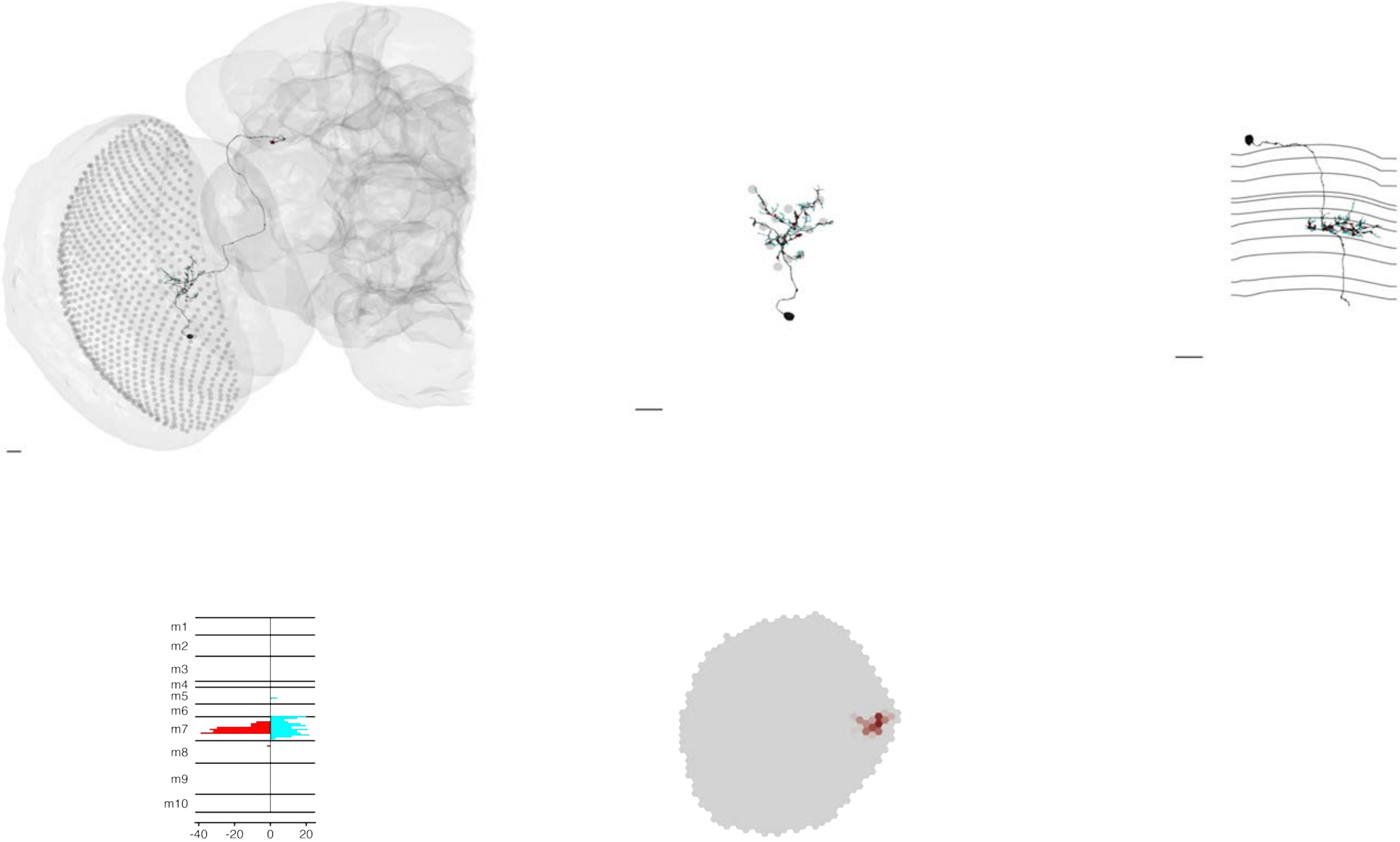

### MeTu3c_ventral_720575940618543445

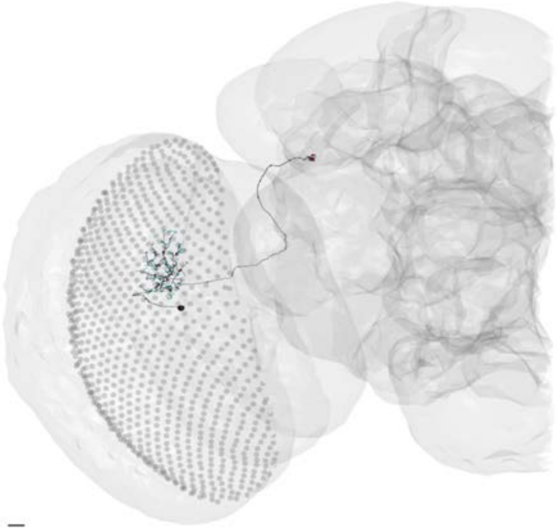

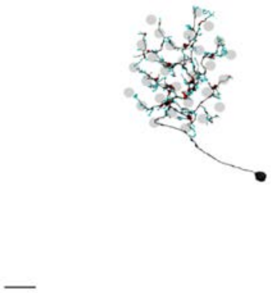

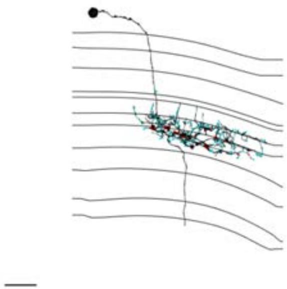

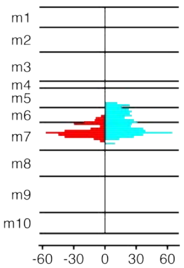


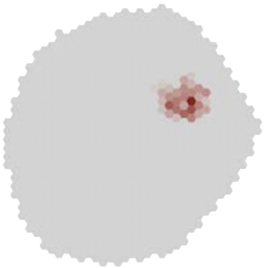

### MeTu3c_ventral_720575940618660145

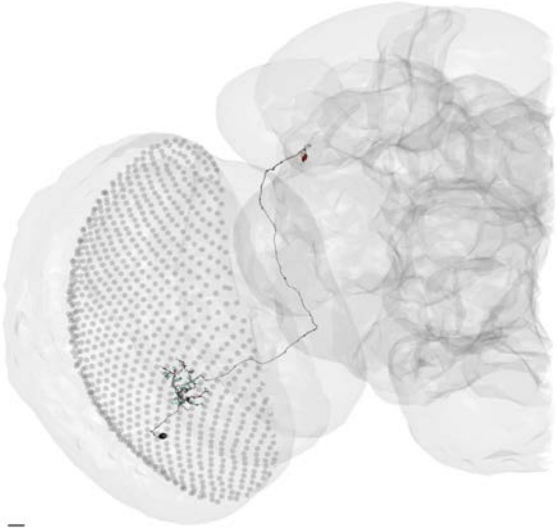

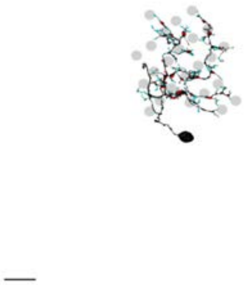

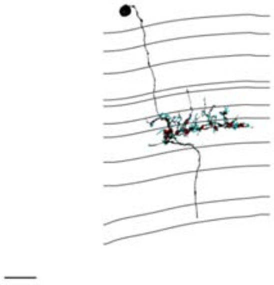

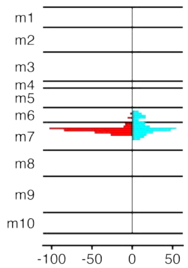


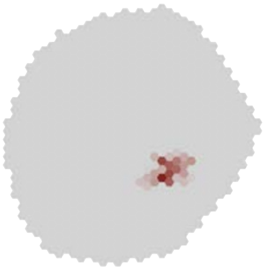

### MeTu3c_ventral_720575940619197319

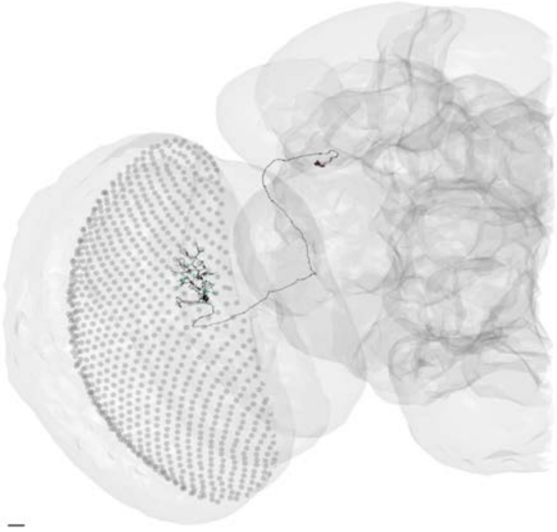

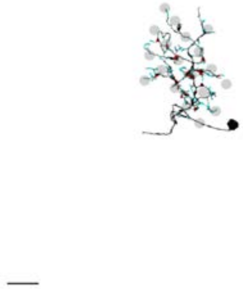

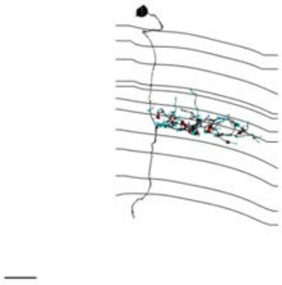

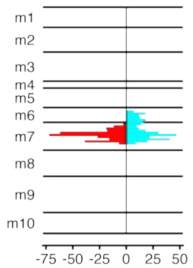


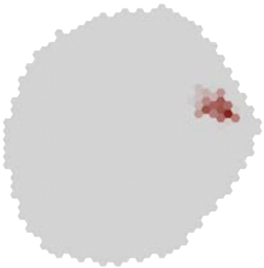

MeTu3c_ventral_720575940619768923

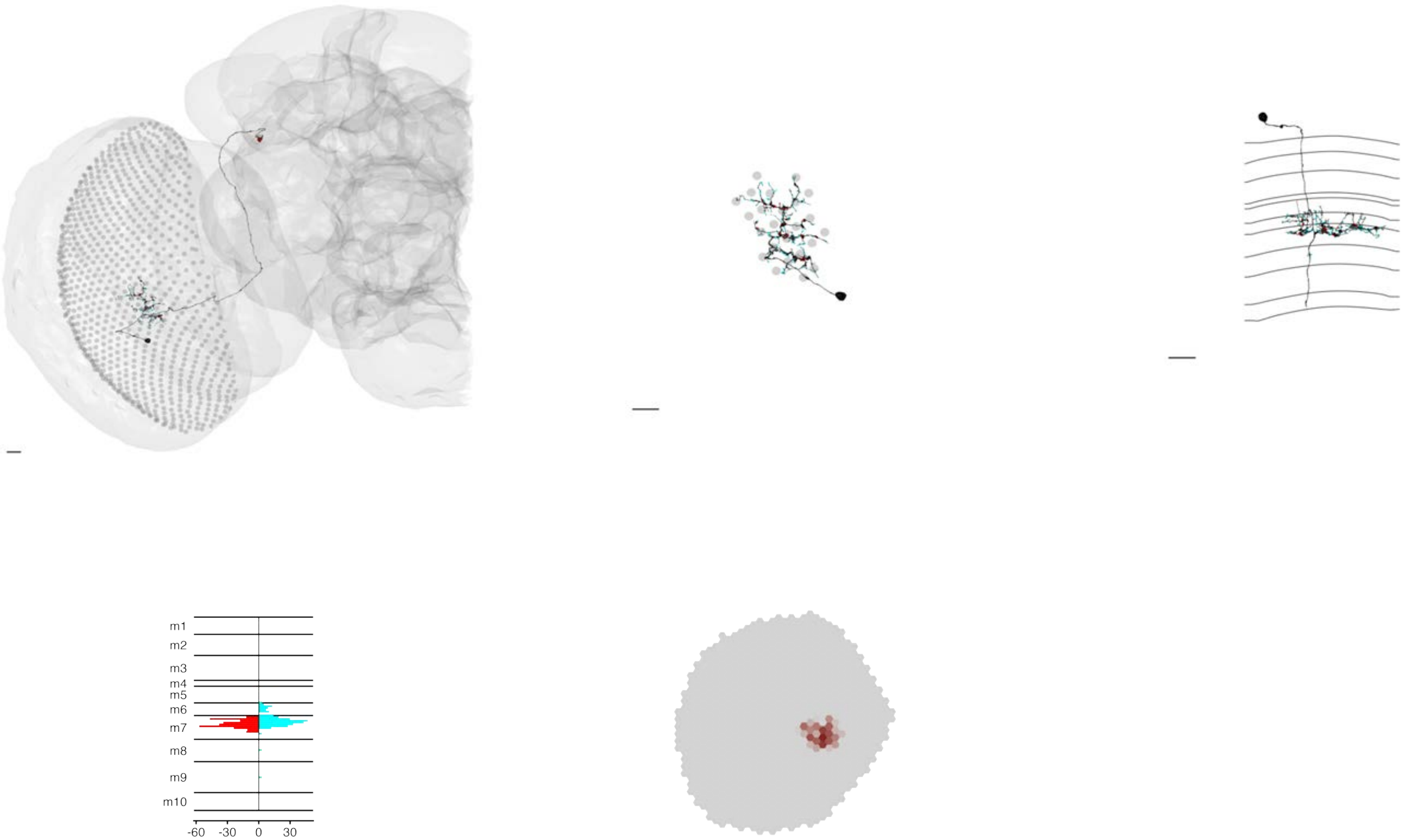

### MeTu3c_ventral_720575940620068317

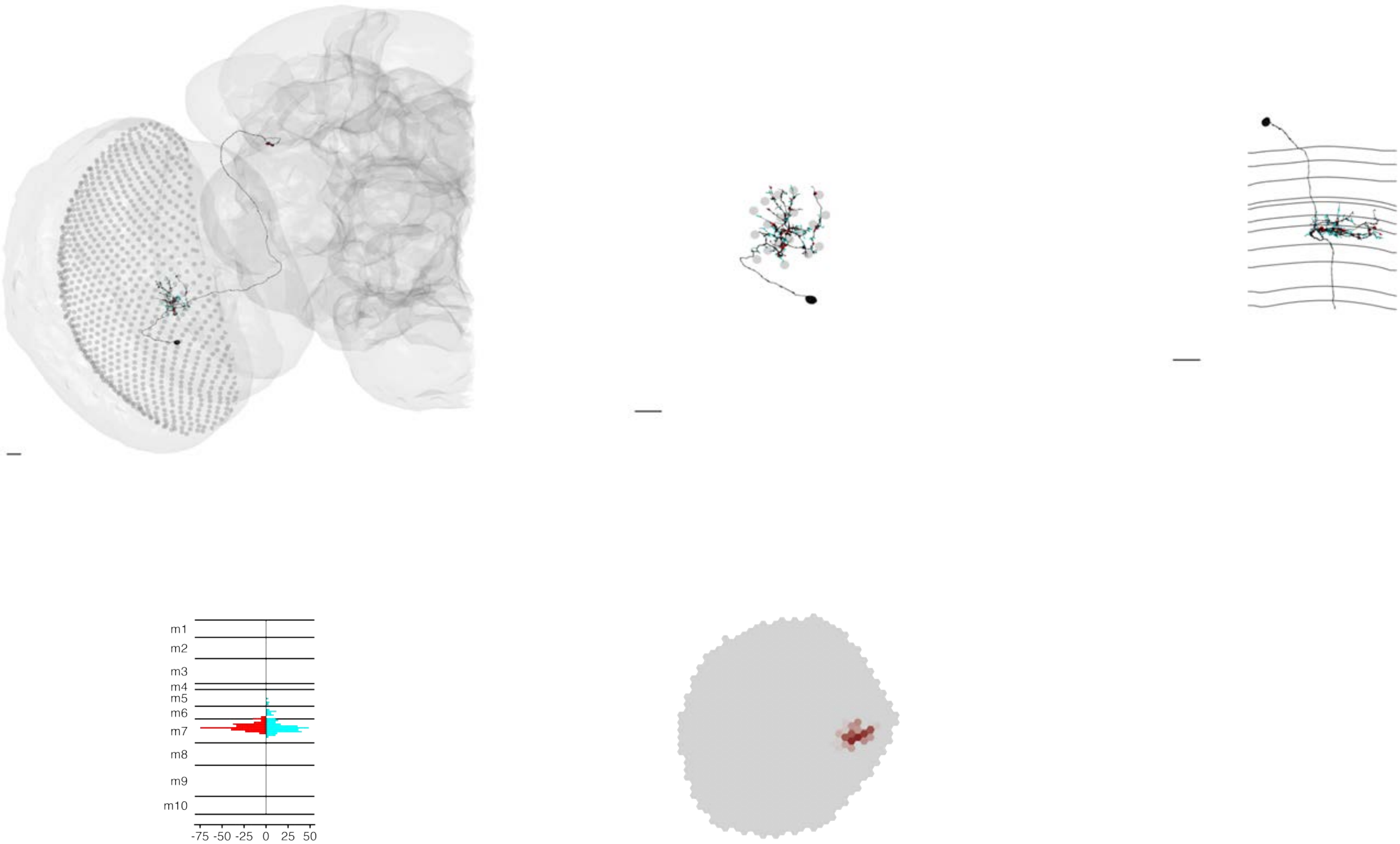

### MeTu3c_ventral_720575940621608406

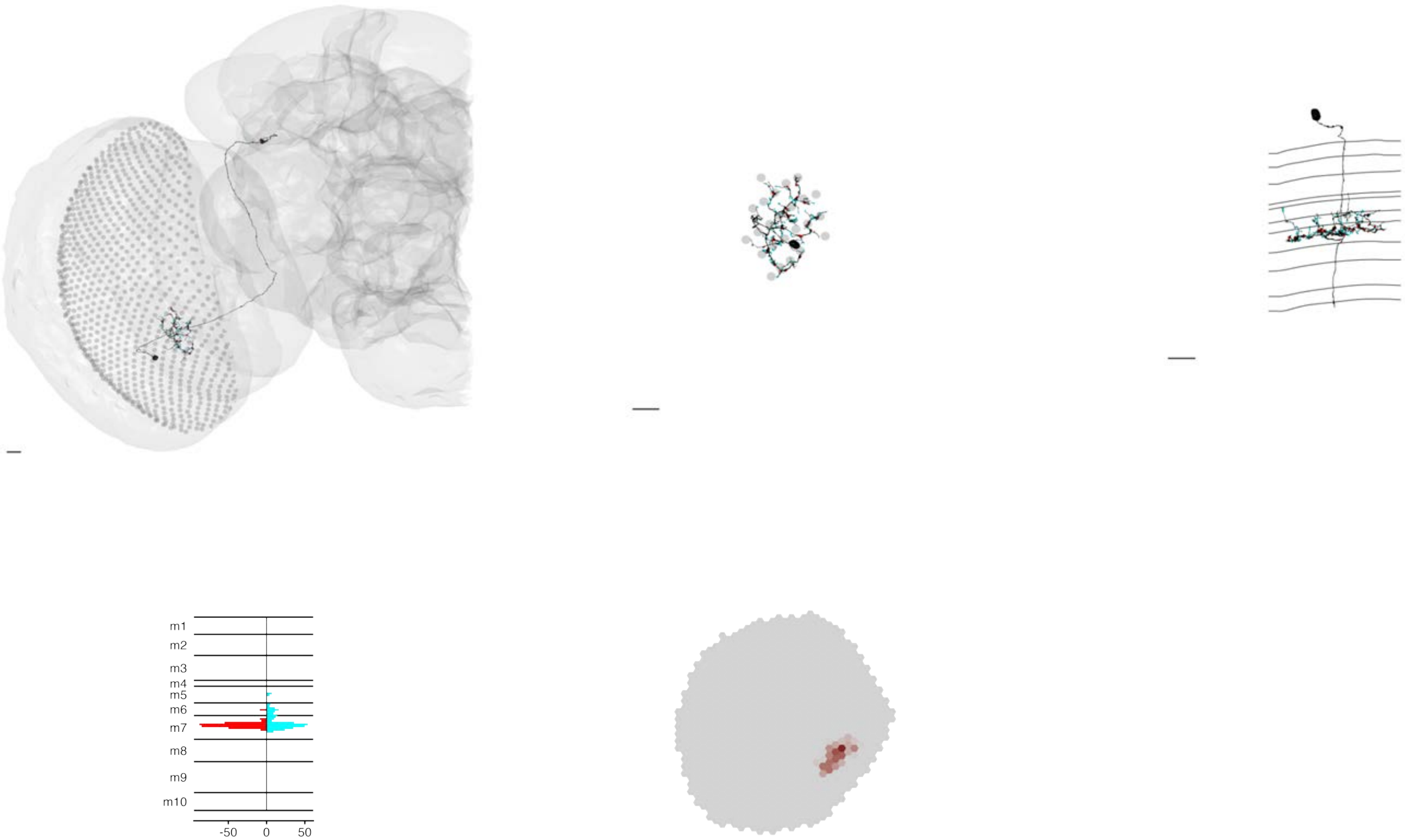

MeTu3c_ventral_720575940622028148

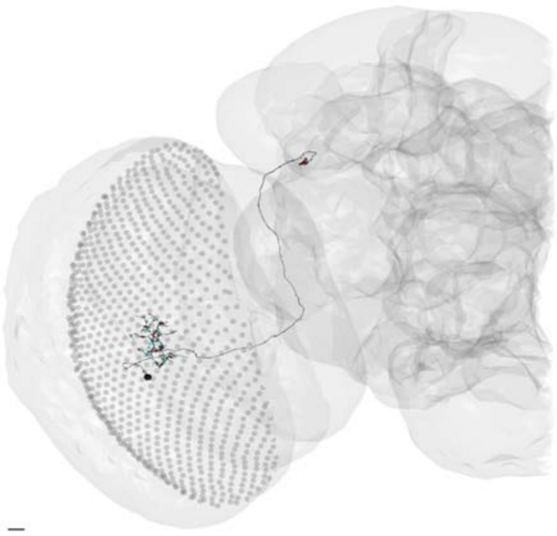

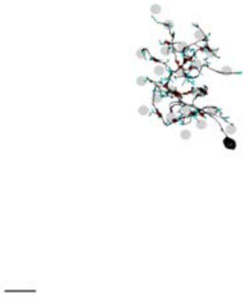

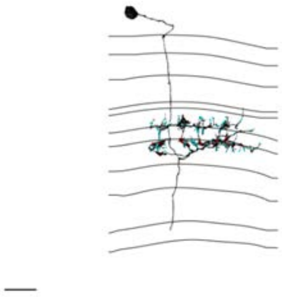

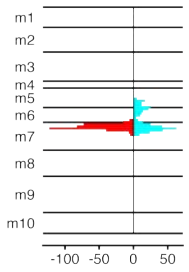


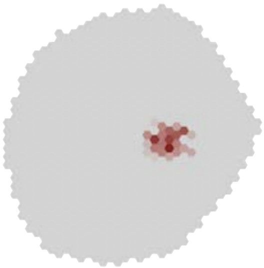

### MeTu3c_ventral_720575940622108679

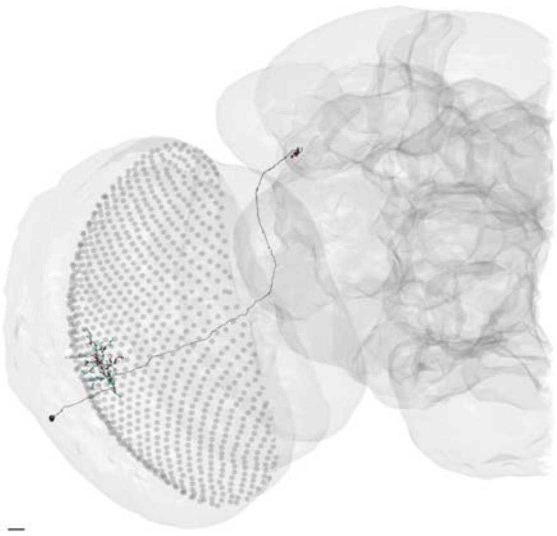

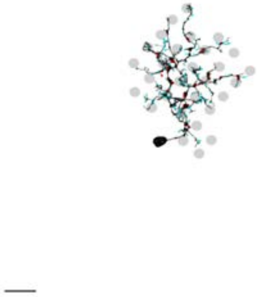

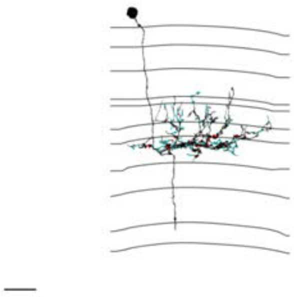

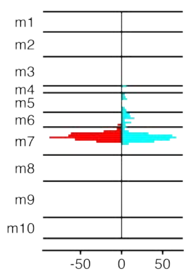


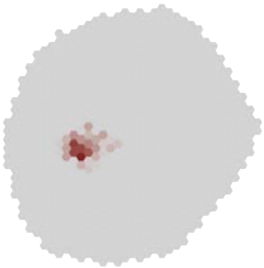

### MeTu3c_ventral_720575940622860314

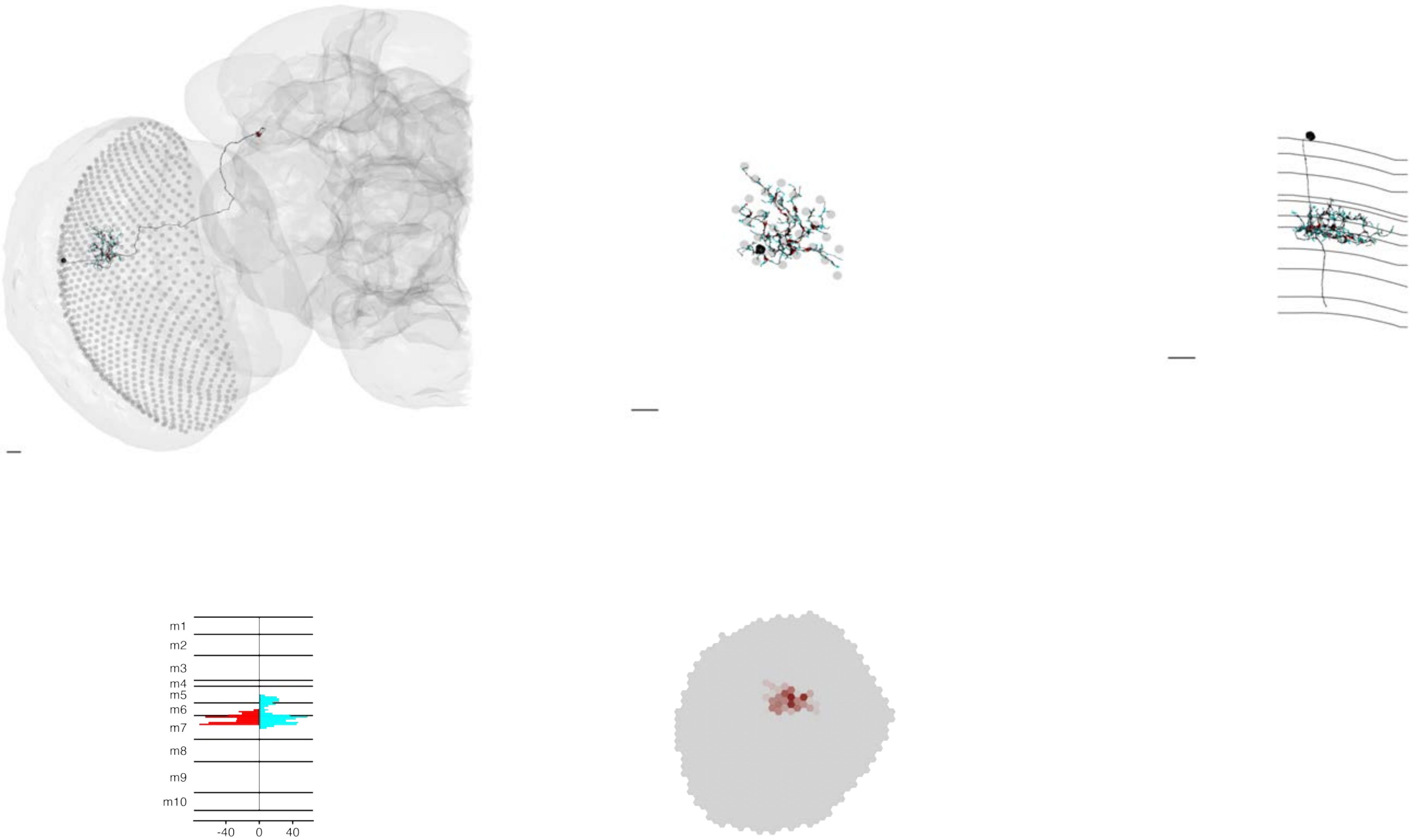

### MeTu3c_ventral_720575940622864154

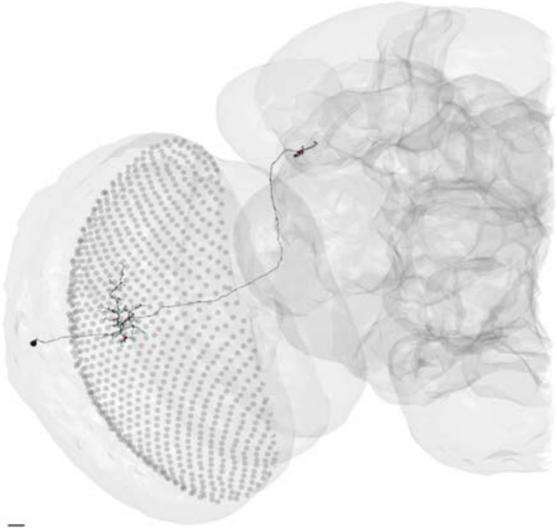

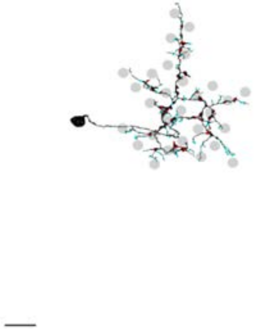

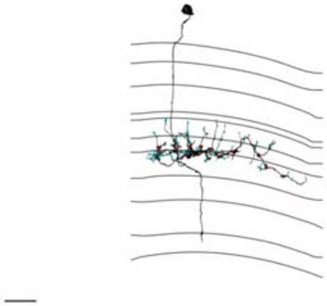

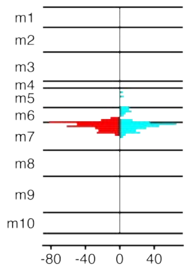


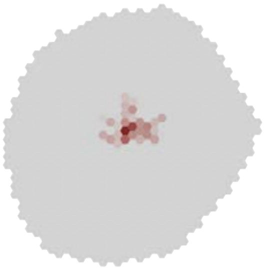

### MeTu3c_ventral_720575940623885861

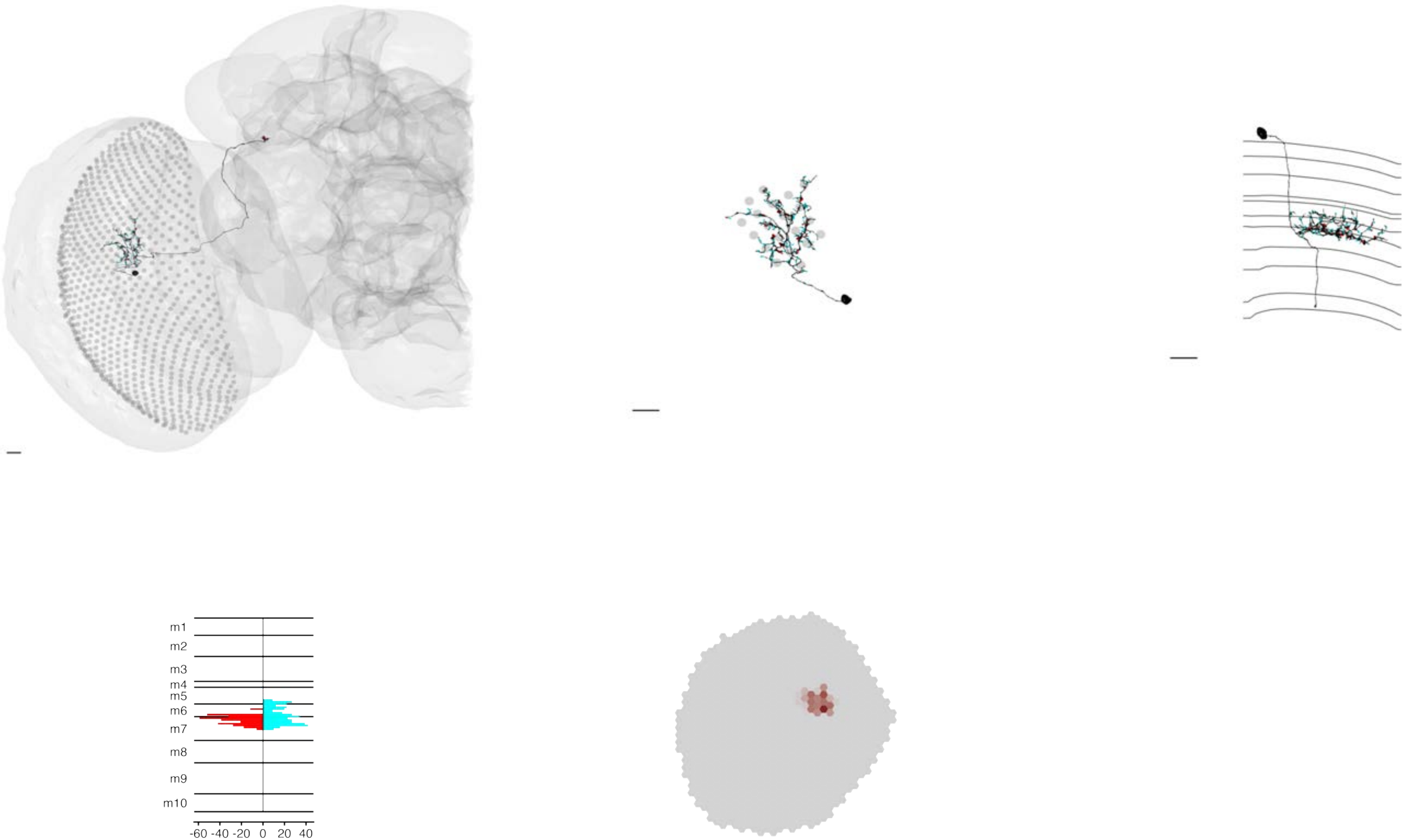

### MeTu3c_ventral_720575940624030990

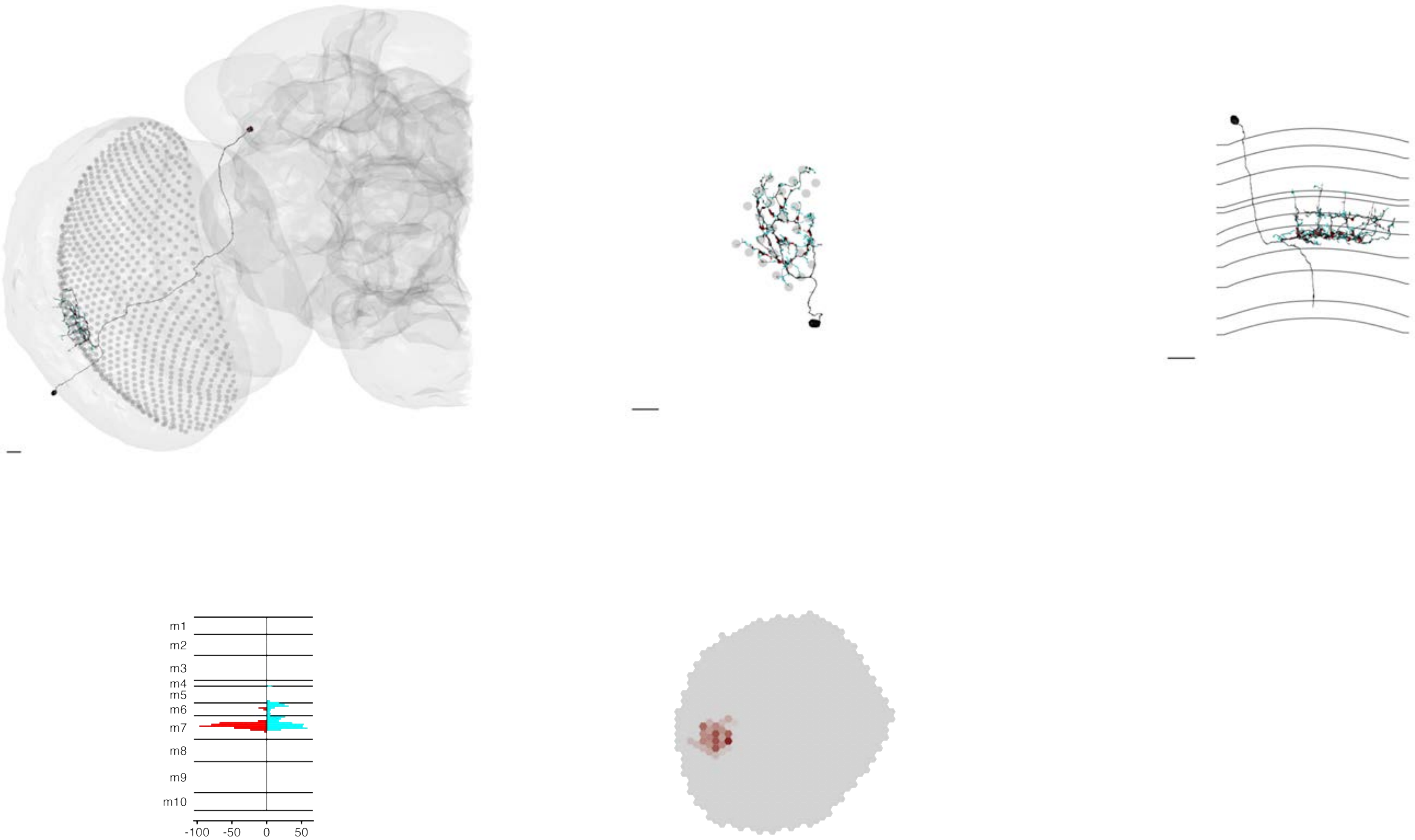

### MeTu3c_ventral_720575940625291907

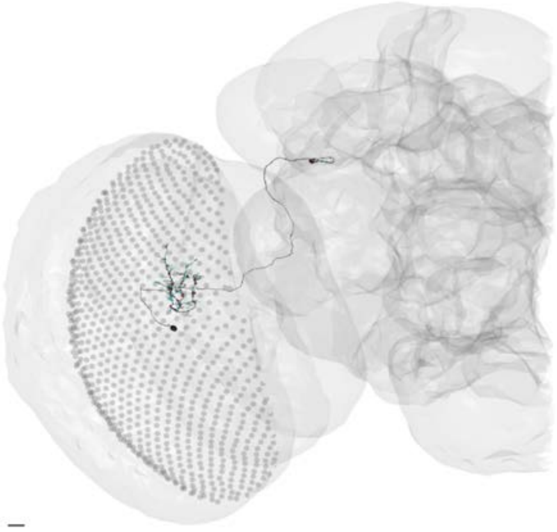

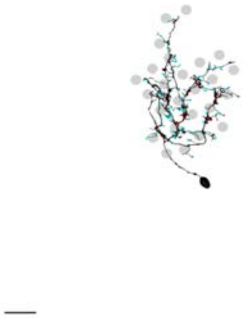

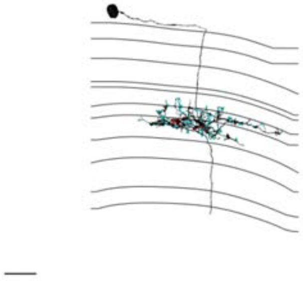

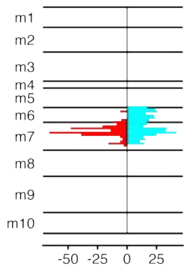


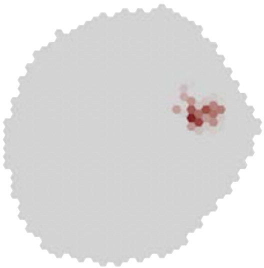

### MeTu3c_ventral_720575940625617418

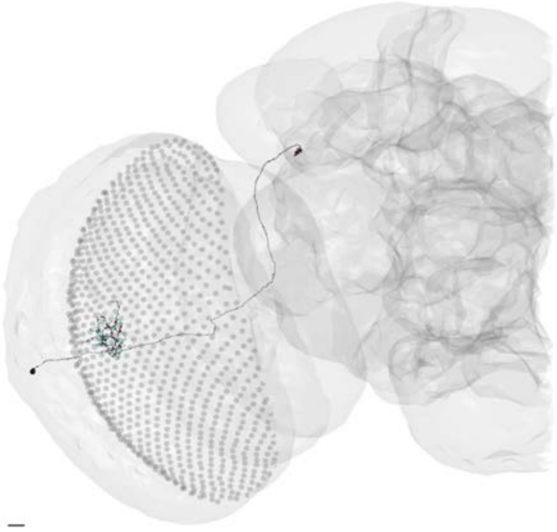

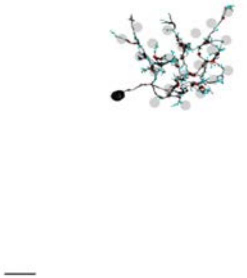

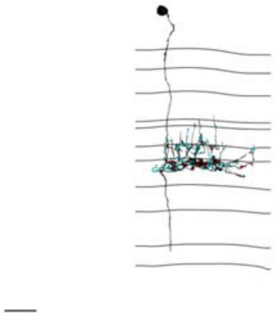

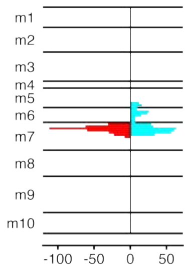


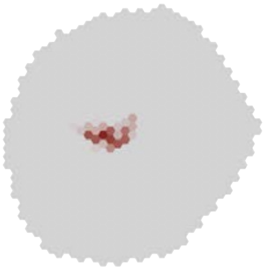

MeTu3c_ventral_720575940627419127

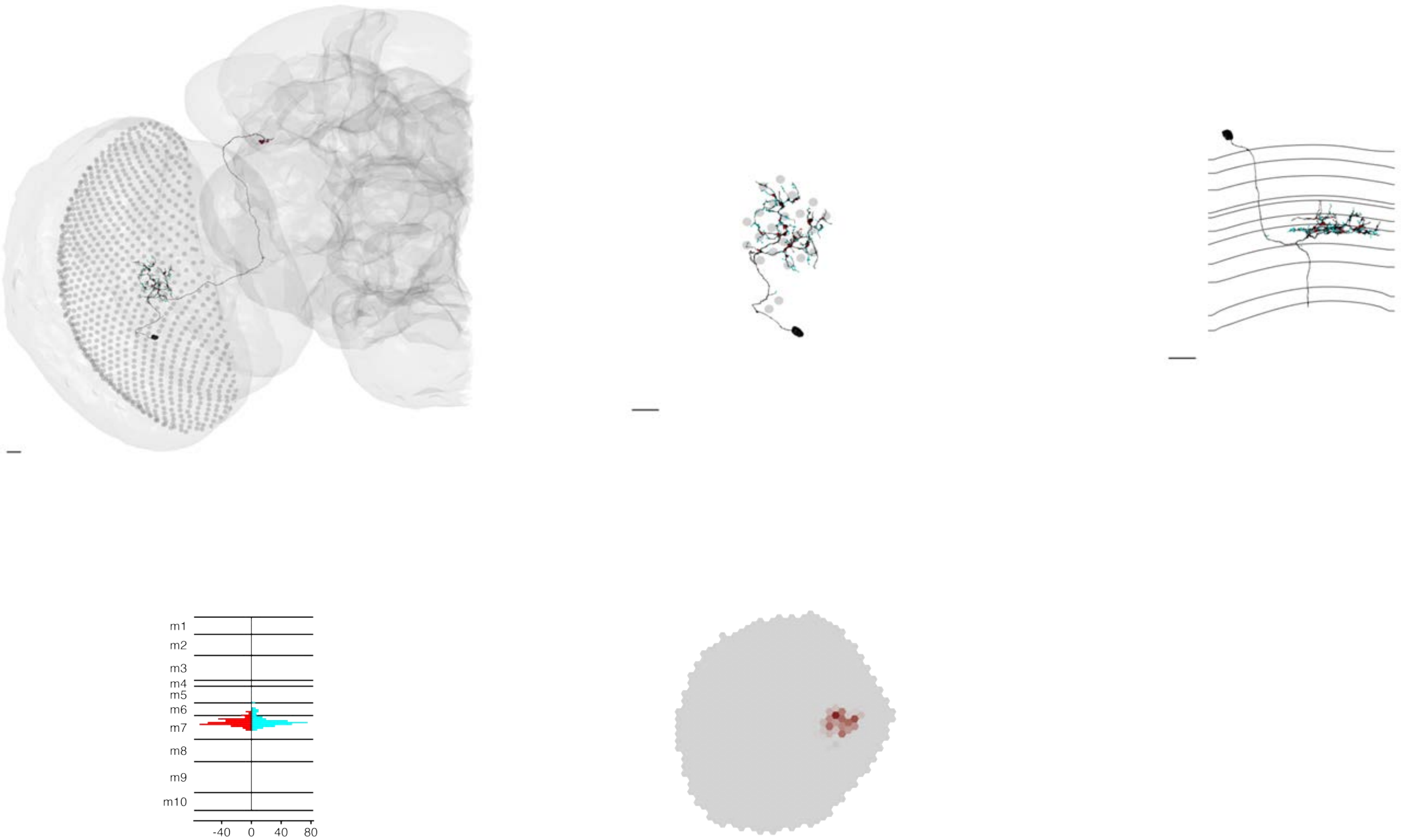

MeTu3c_ventral_720575940628495695

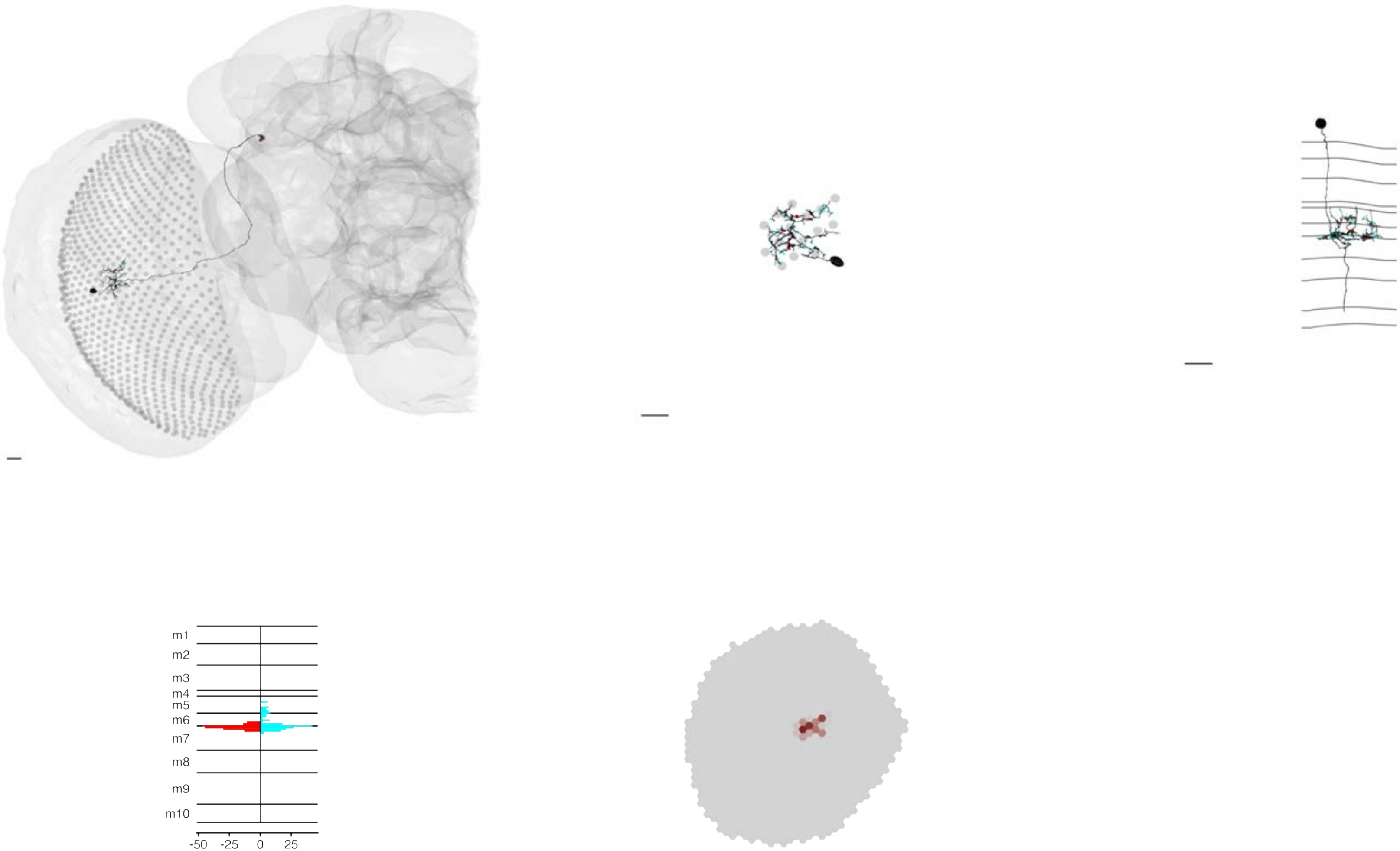

### MeTu3c_ventral_720575940628940670

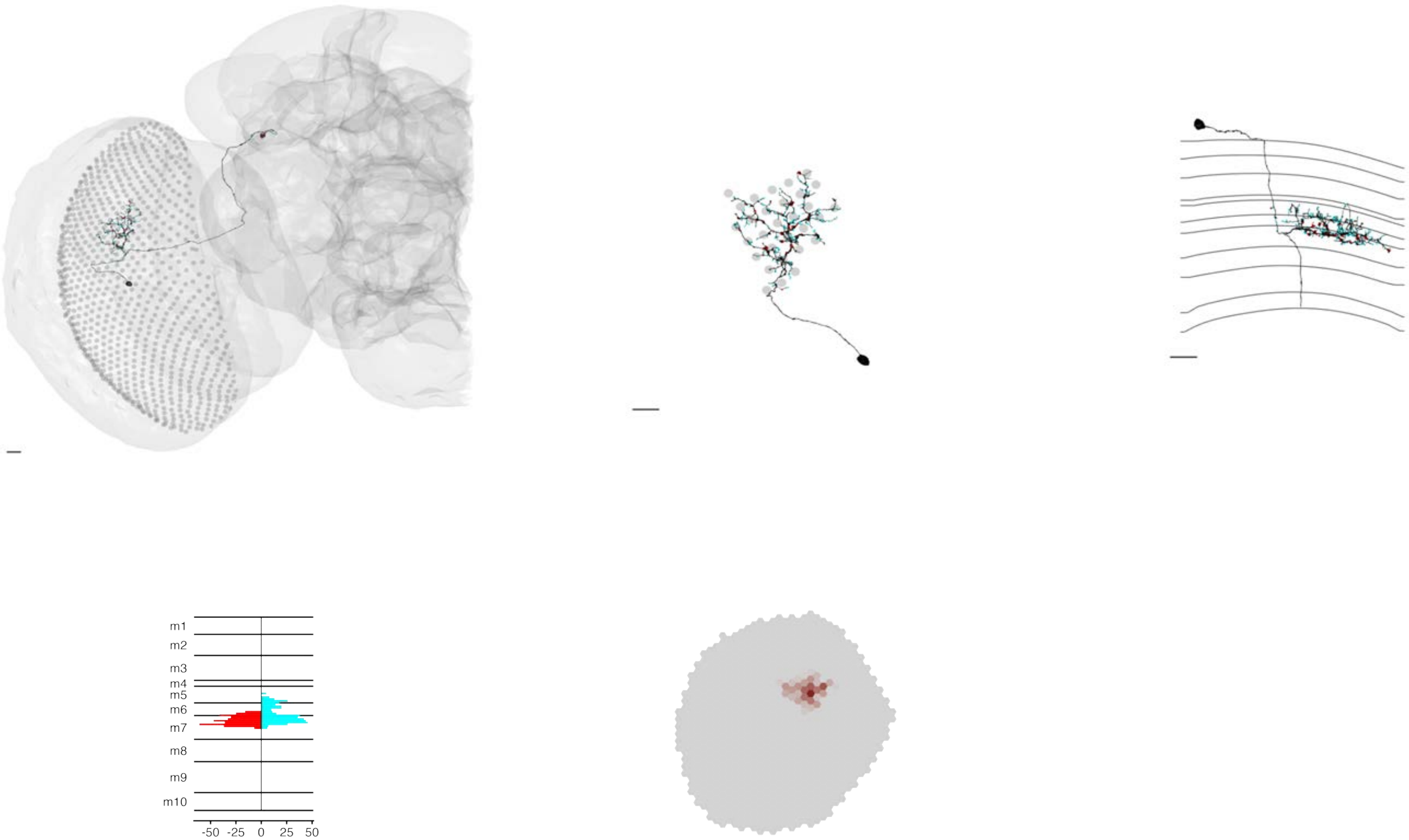

### MeTu3c_ventral_720575940629600271

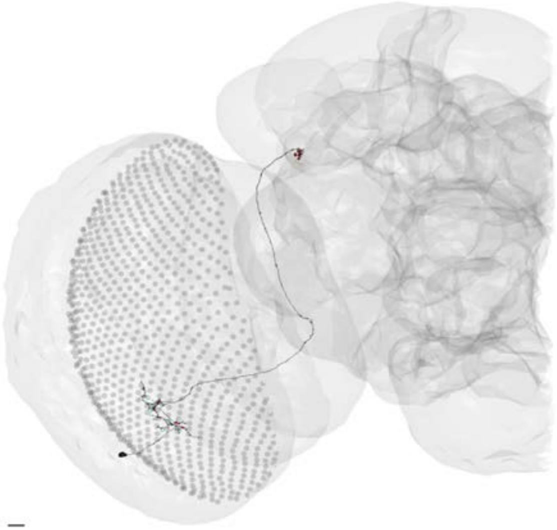

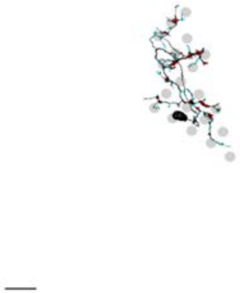

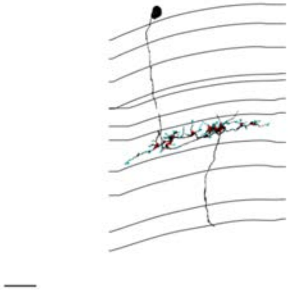

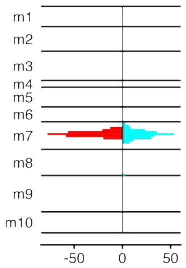


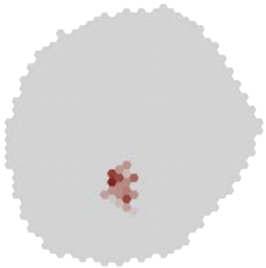

### MeTu3c_ventral_720575940630184044

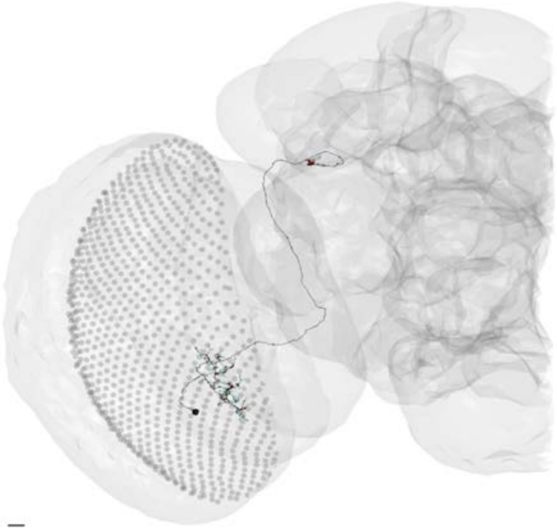

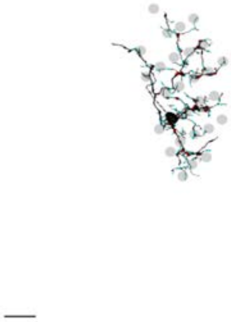

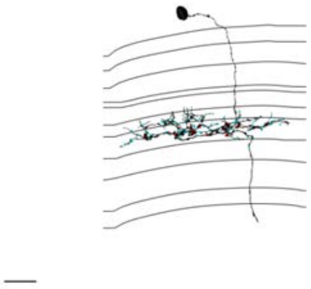

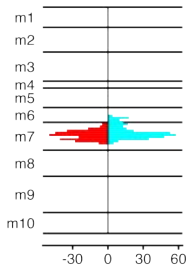


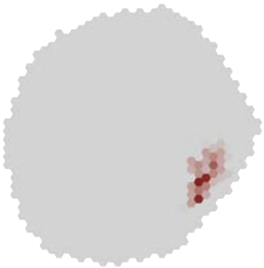

### MeTu3c_ventral_720575940630511663

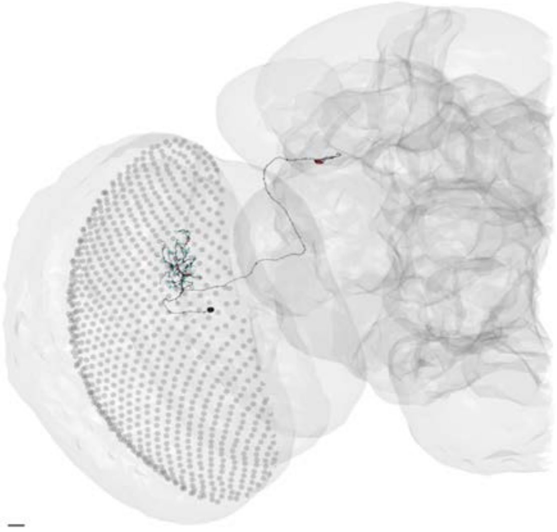

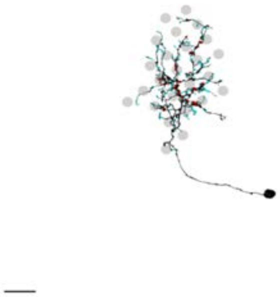

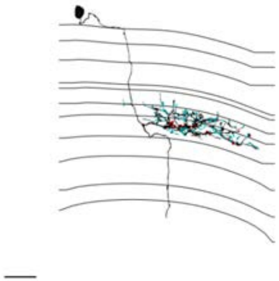

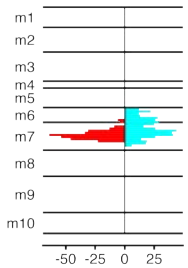


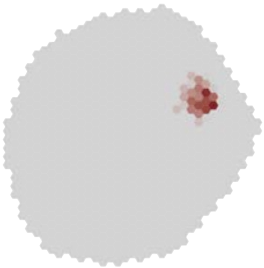

### MeTu3c_ventral_720575940631421231

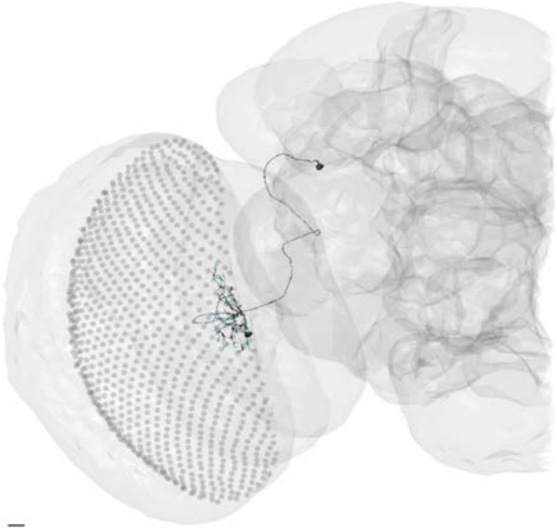

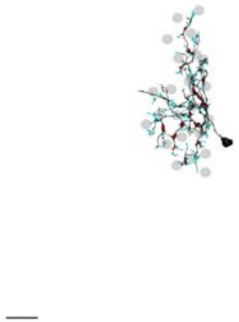

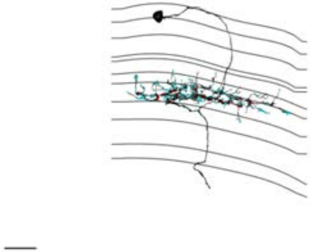

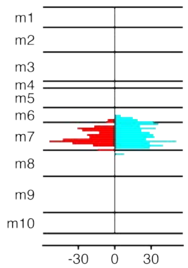


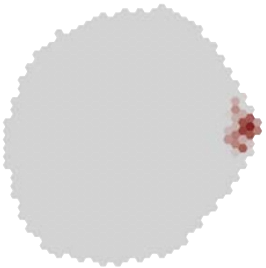

### MeTu3c_ventral_720575940632859729

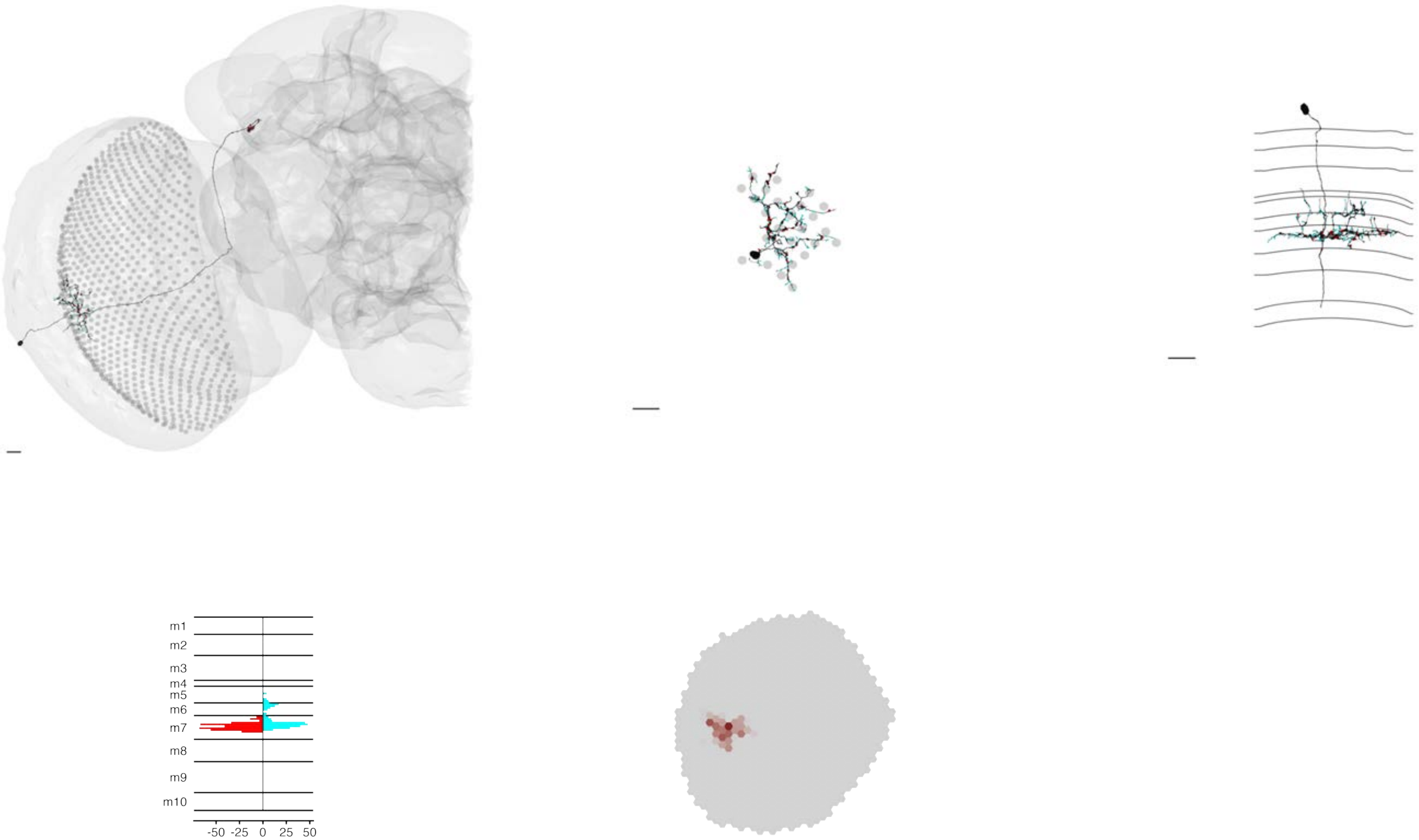

### MeTu3c_ventral_720575940633261267

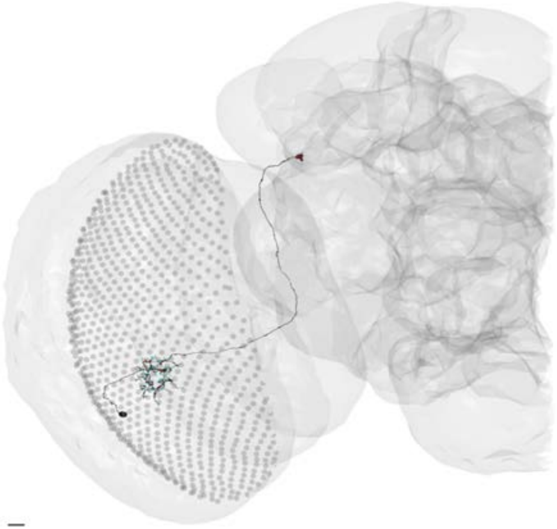

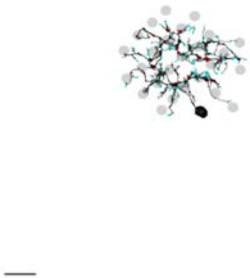

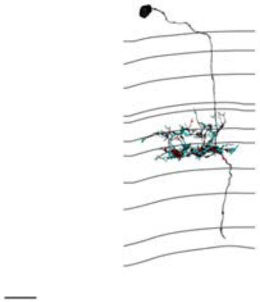

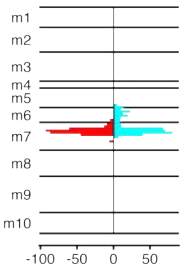


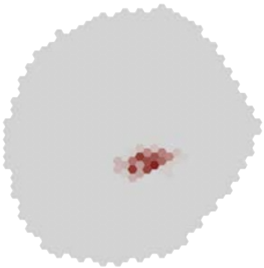

### MeTu3c_ventral_720575940634378351

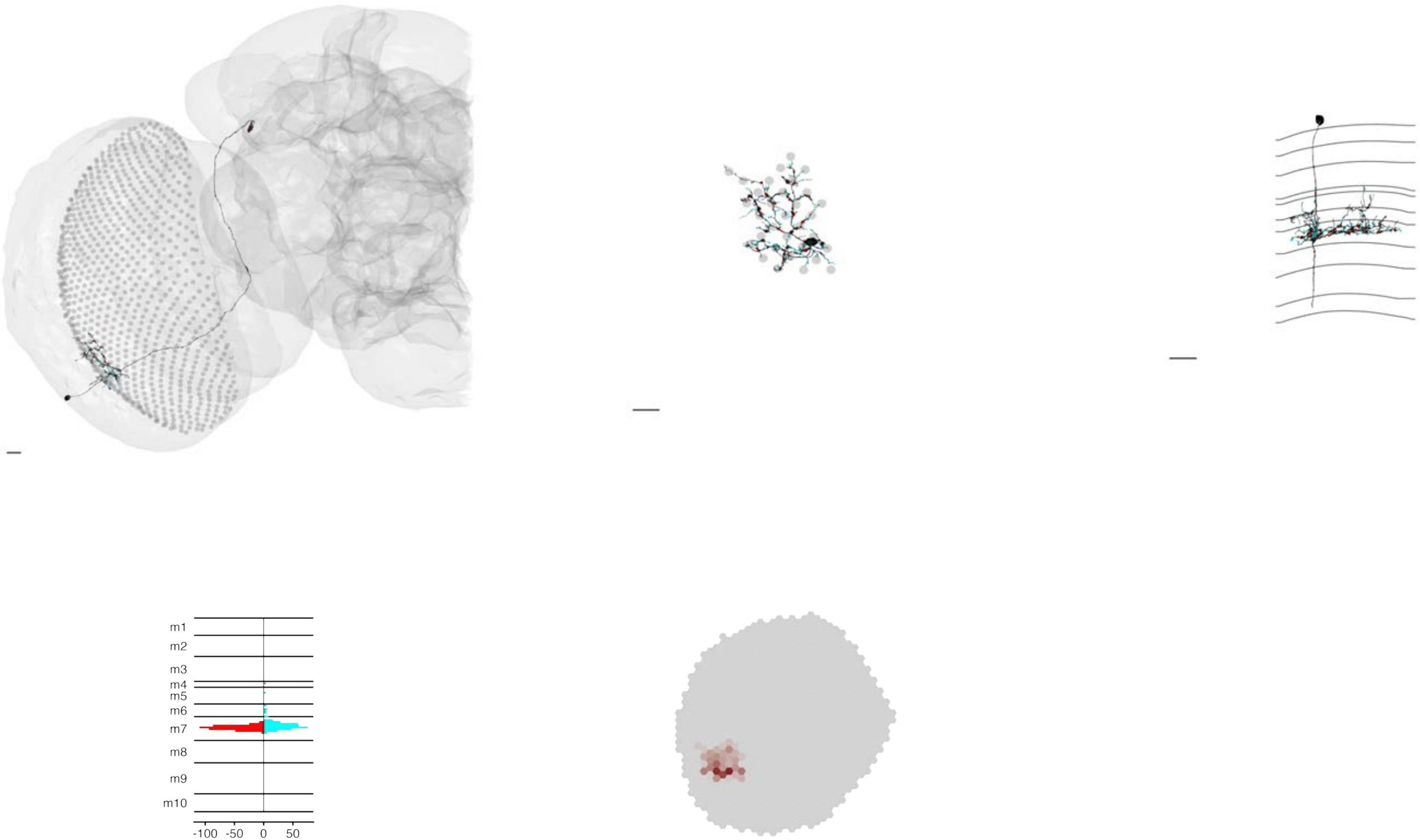

MeTu3c_ventral_720575940634649244

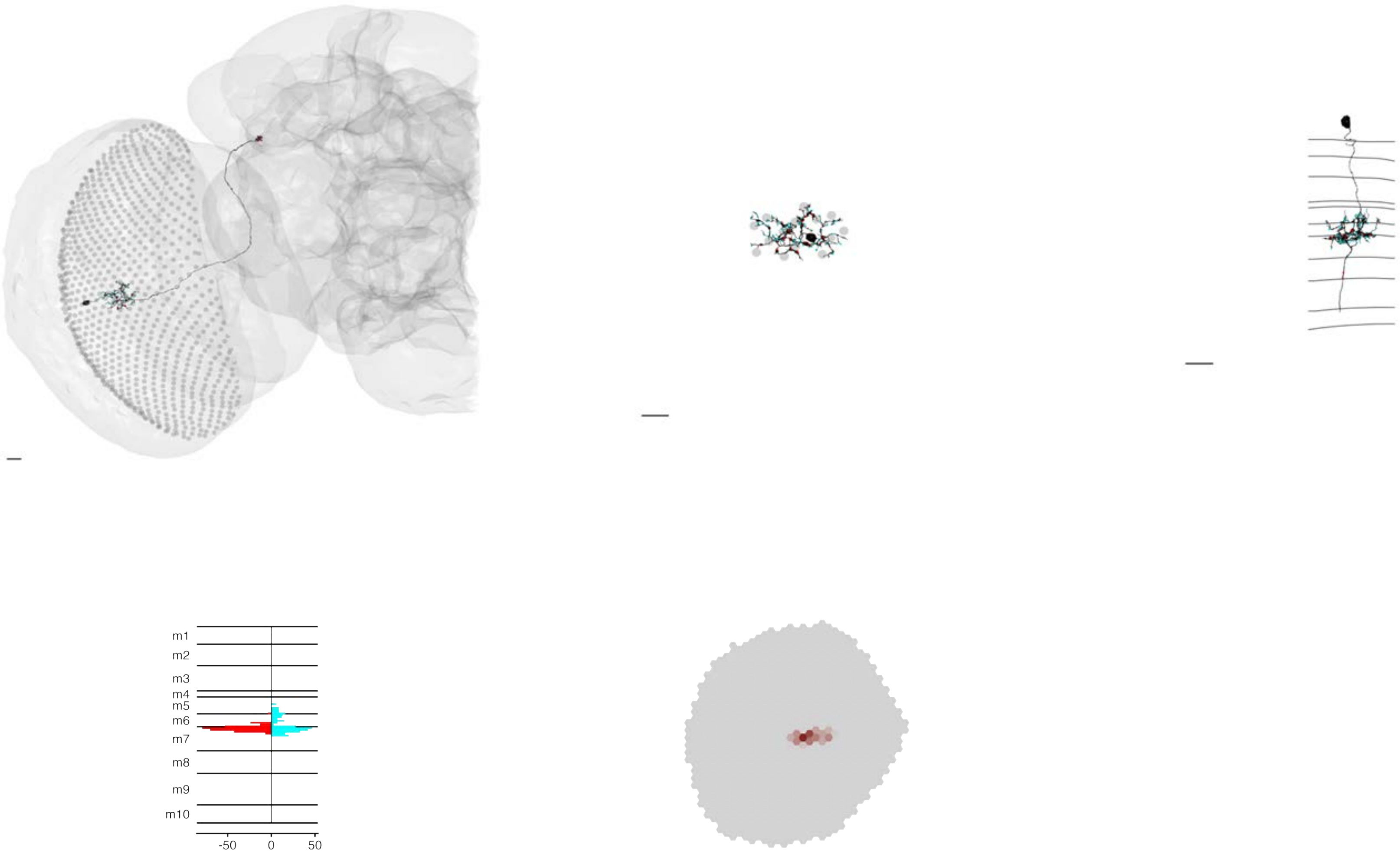

### MeTu3c_ventral_720575940639526901

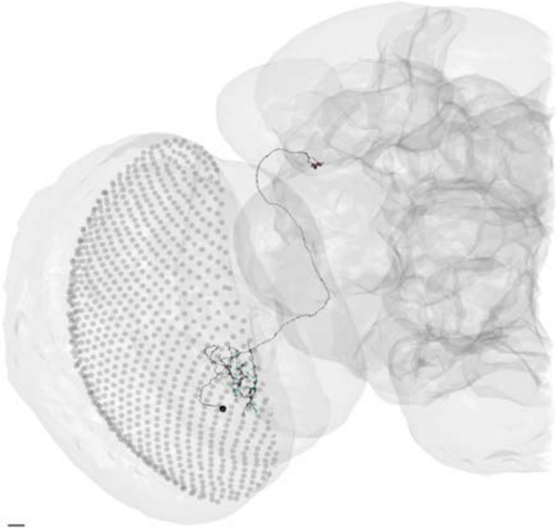

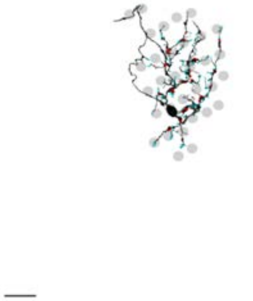

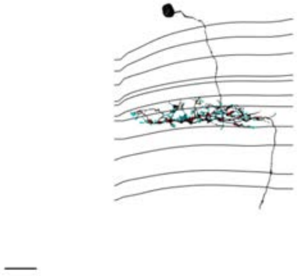

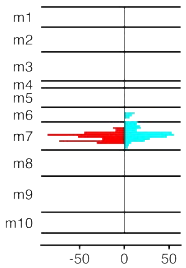


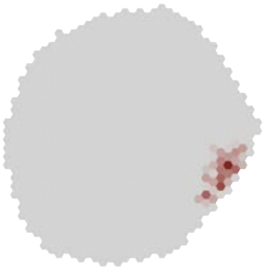

### MeTu3c_ventral_720575940652753825

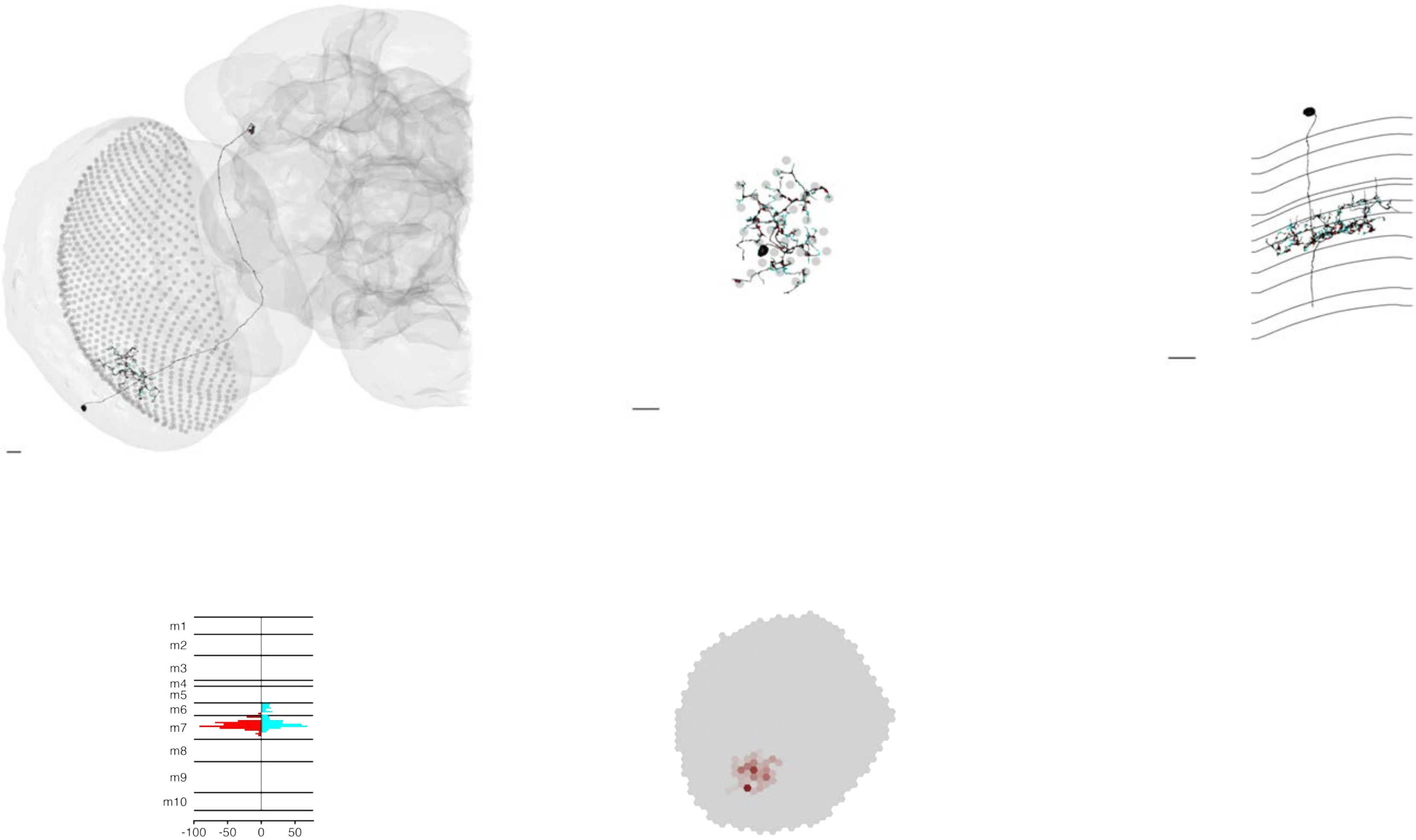

### MeTu4a_720575940605476994

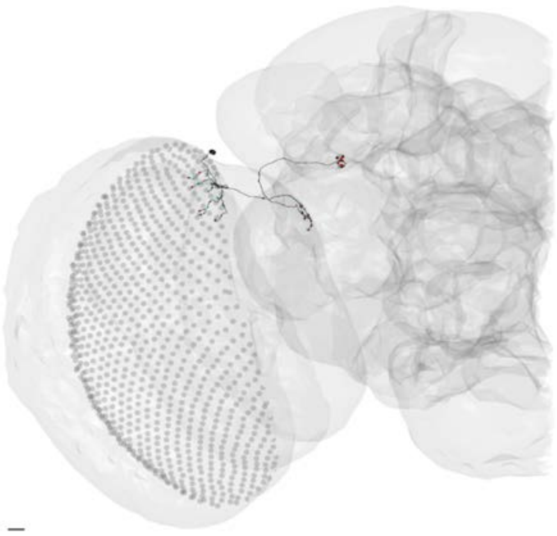

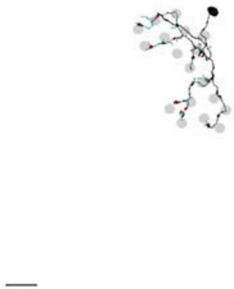

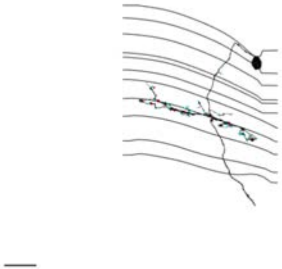

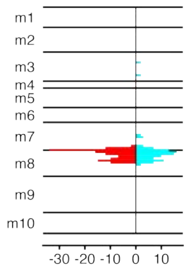


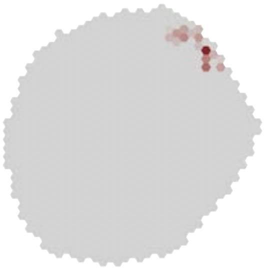

### MeTu4a_720575940606768049

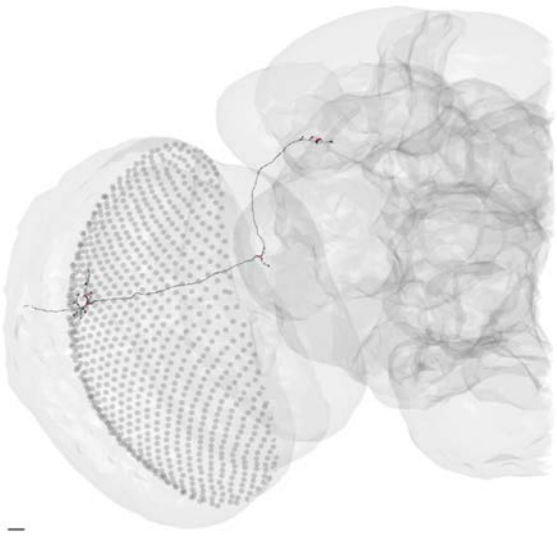

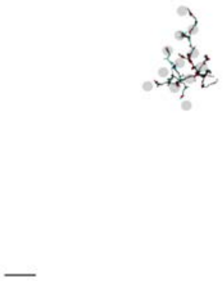

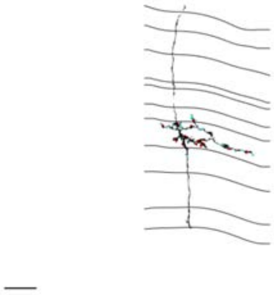

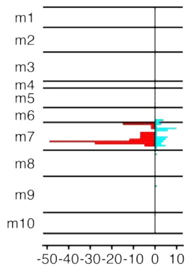


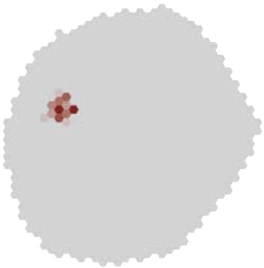

### MeTu4a_720575940607630172

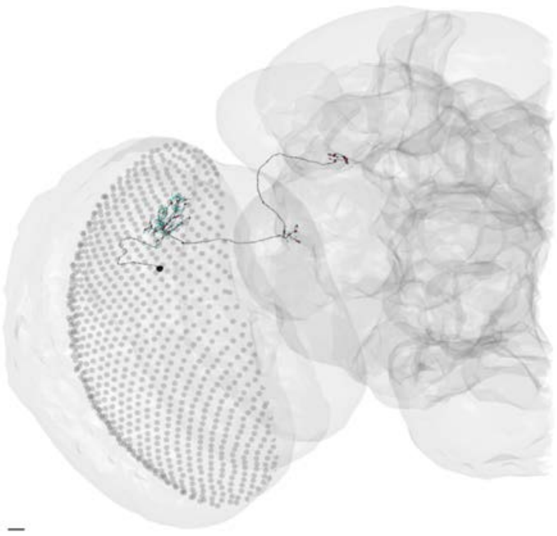

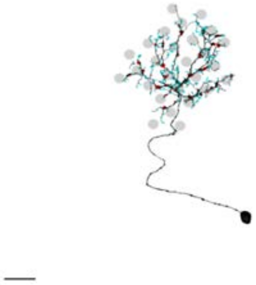

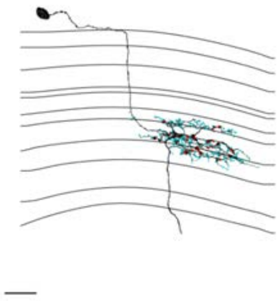

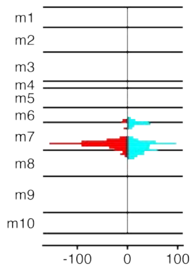


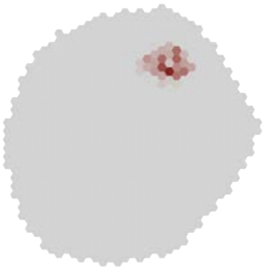

### MeTu4a_720575940608756600

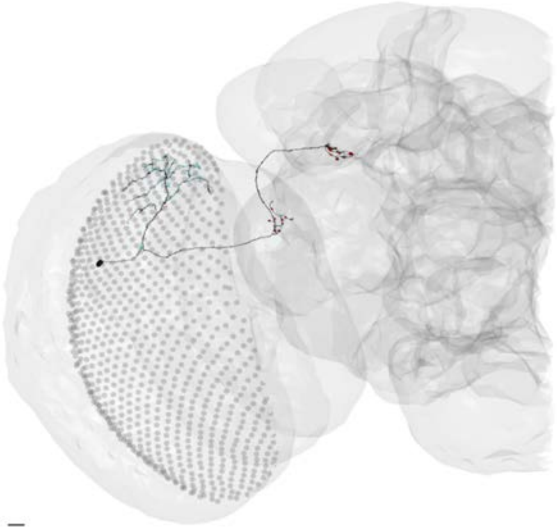

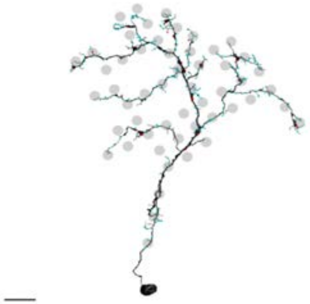

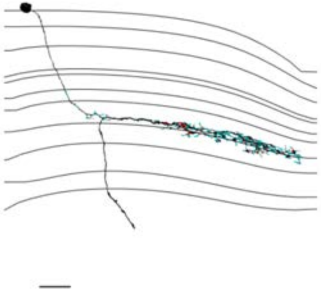

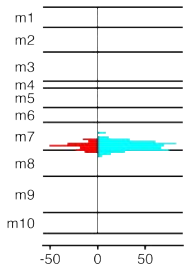


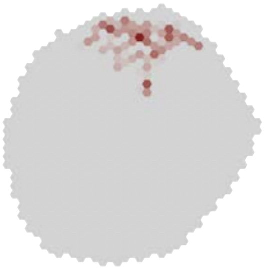

MeTu4a_720575940610238418

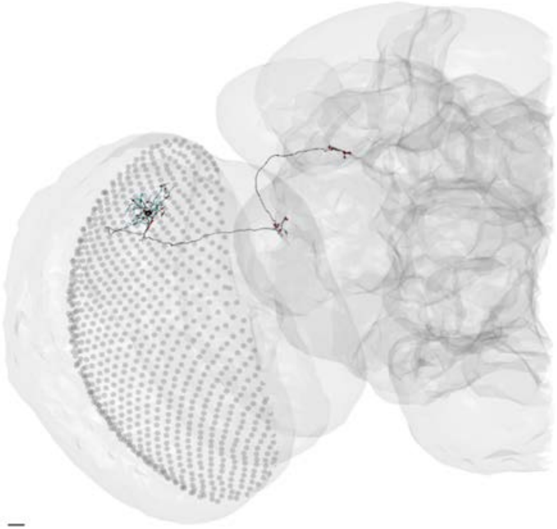

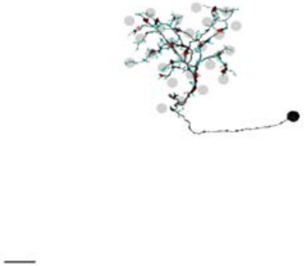

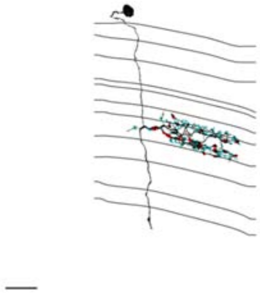

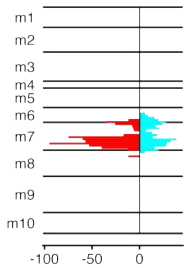


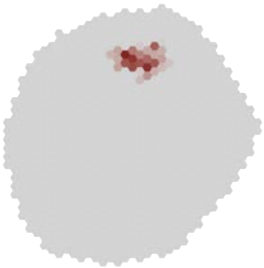

### MeTu4a_720575940610765380

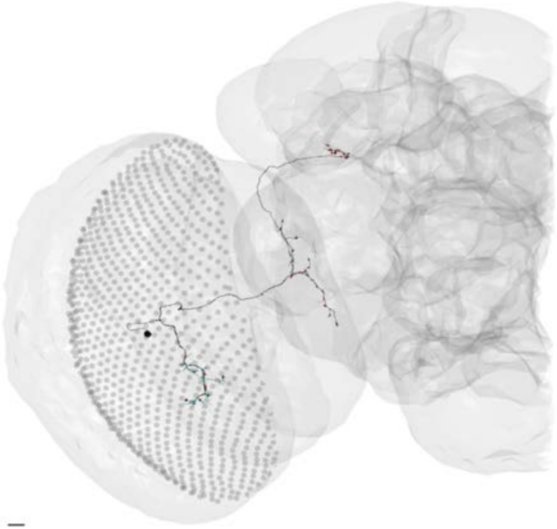

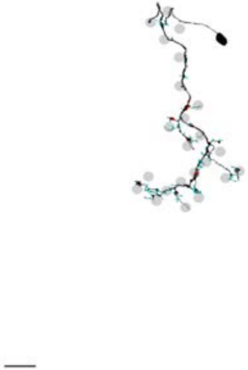

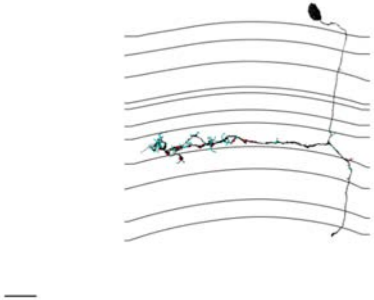

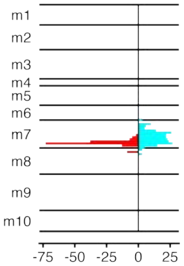


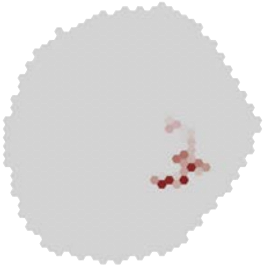

### MeTu4a_720575940610804453

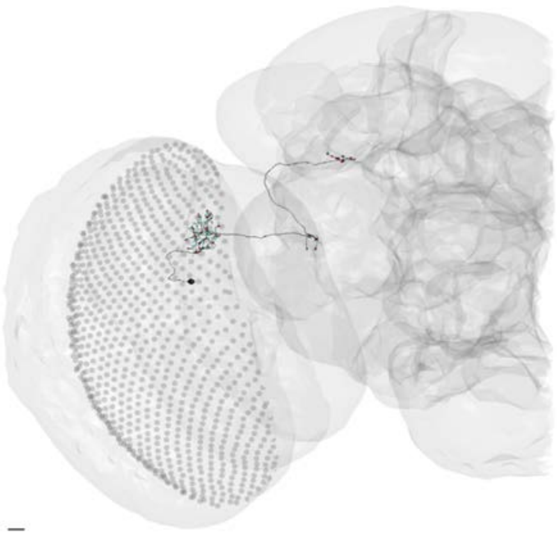

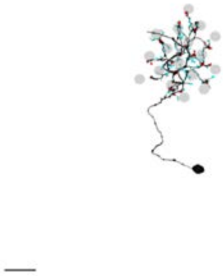

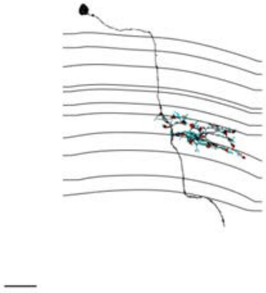

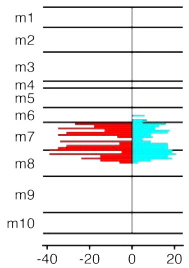


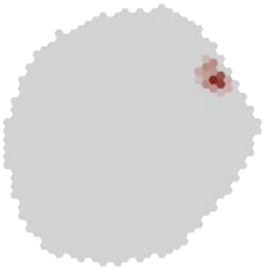

### MeTu4a_720575940610809061

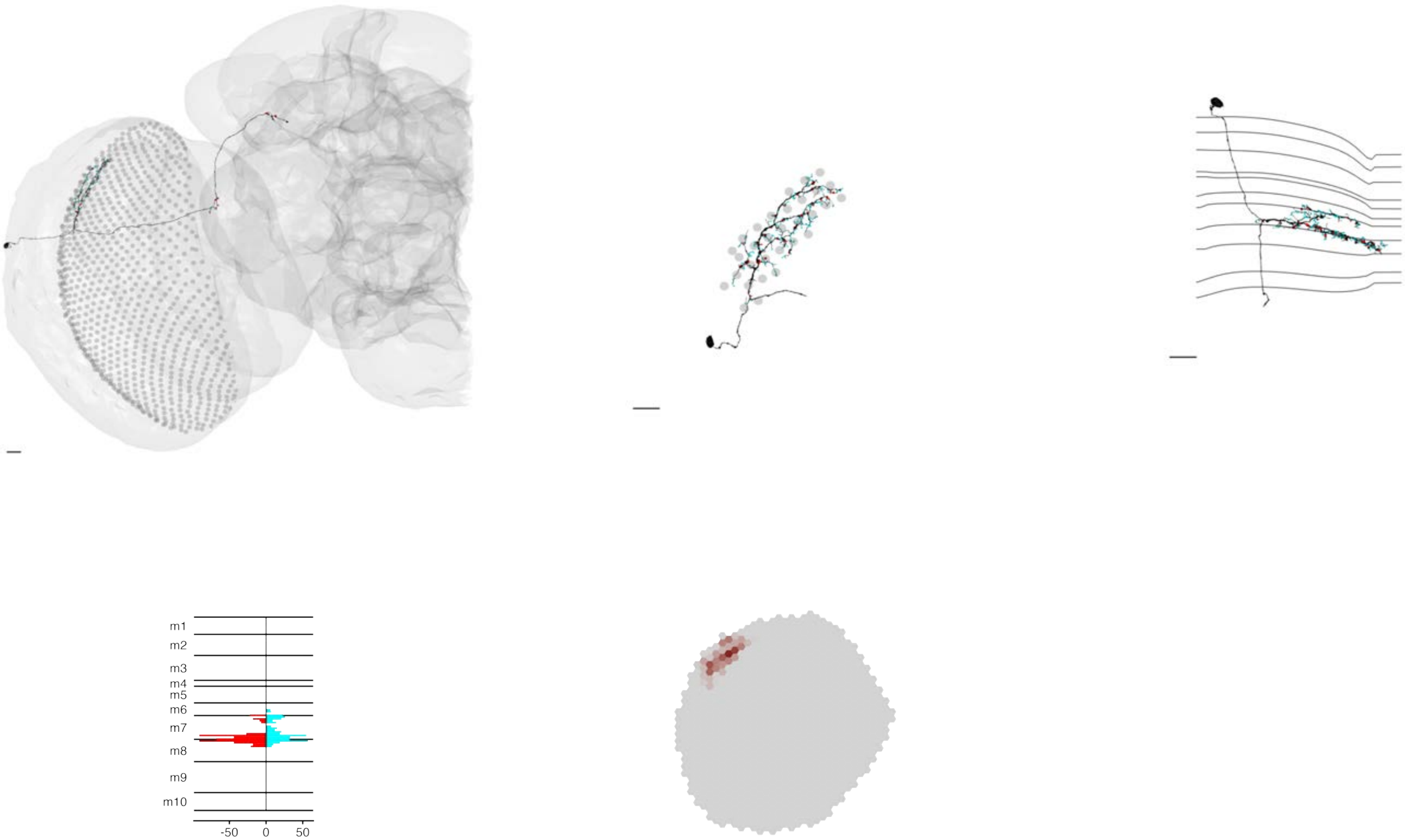

### MeTu4a_720575940613072382

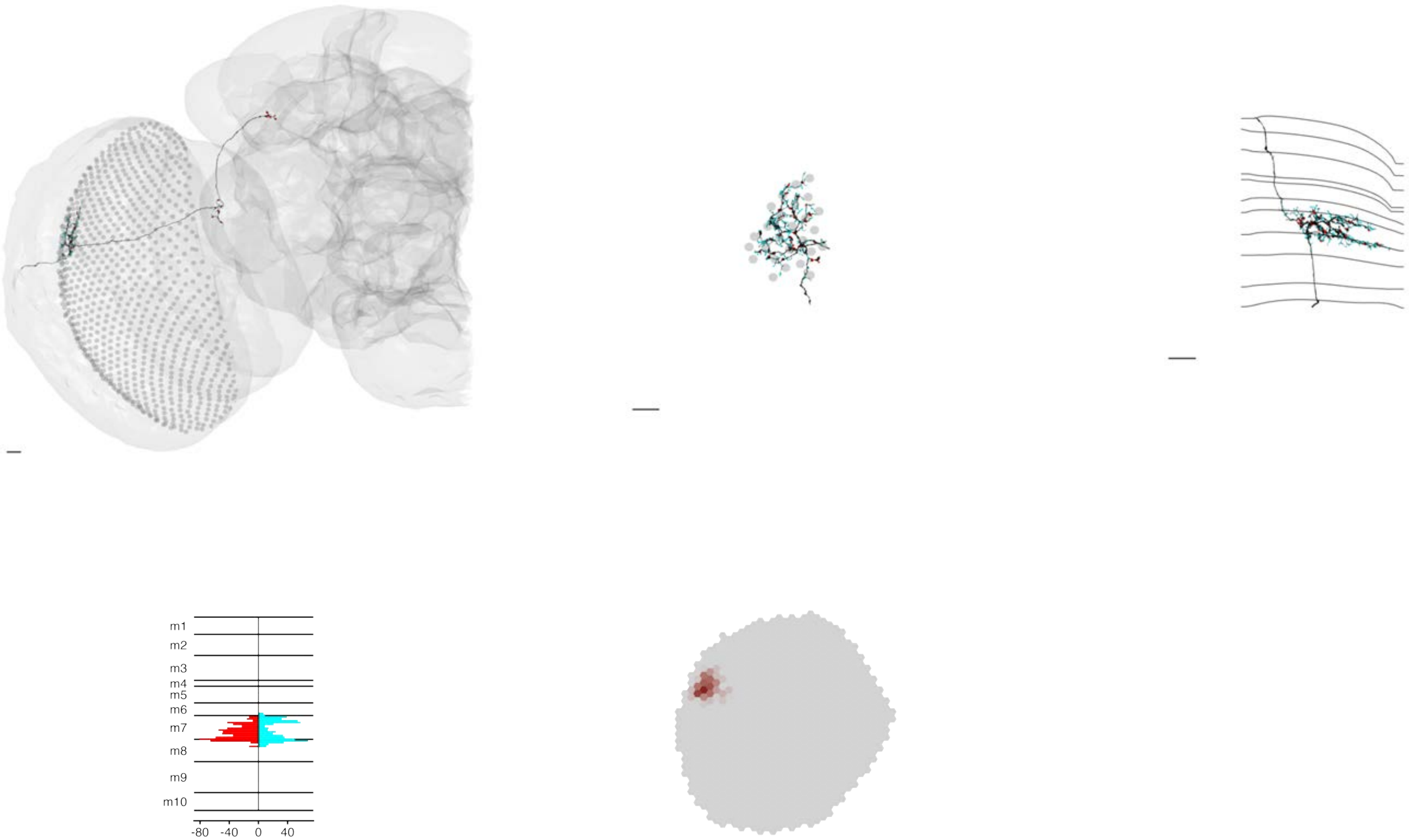

### MeTu4a_720575940613681662

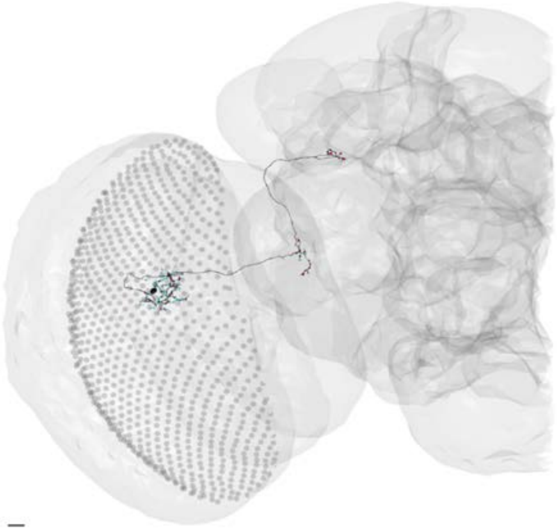

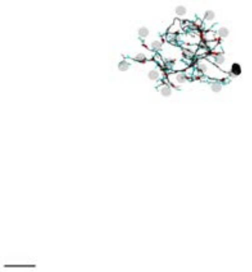

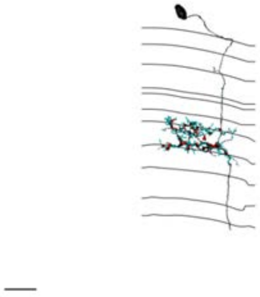

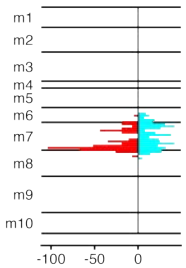


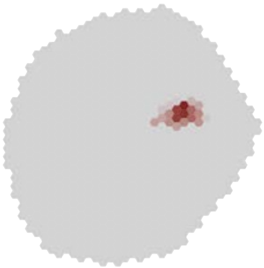

### MeTu4a_720575940613977503

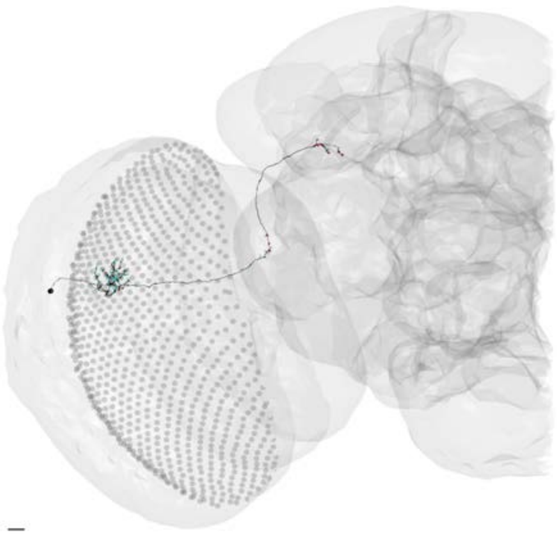

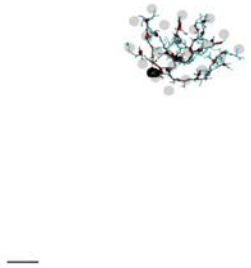

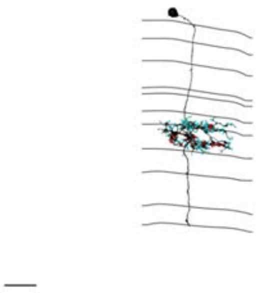

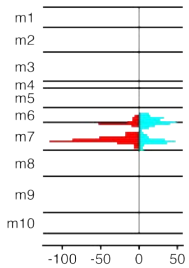


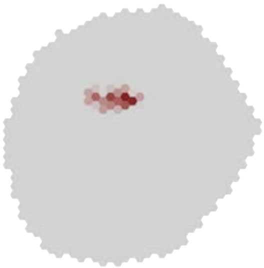

### MeTu4a_720575940614421471

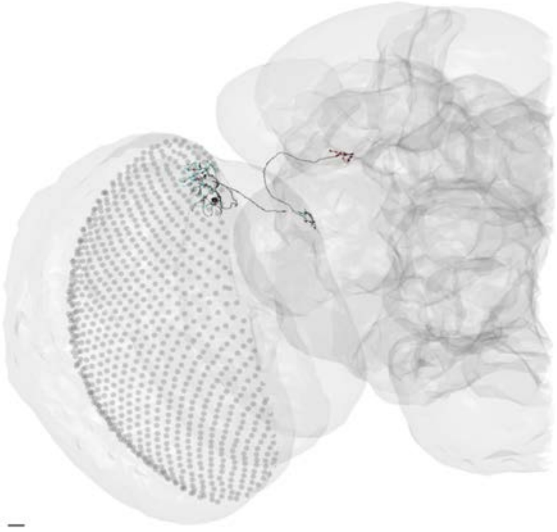

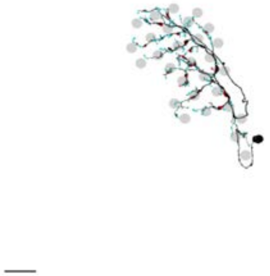

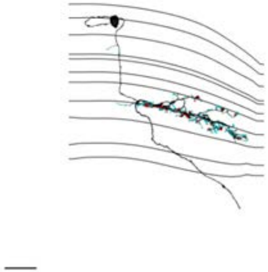

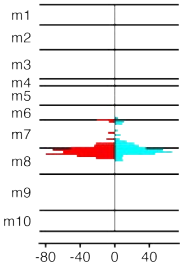


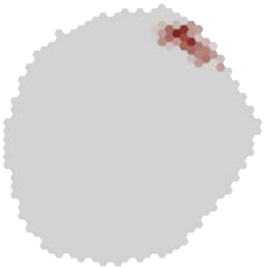

### MeTu4a_720575940616290004

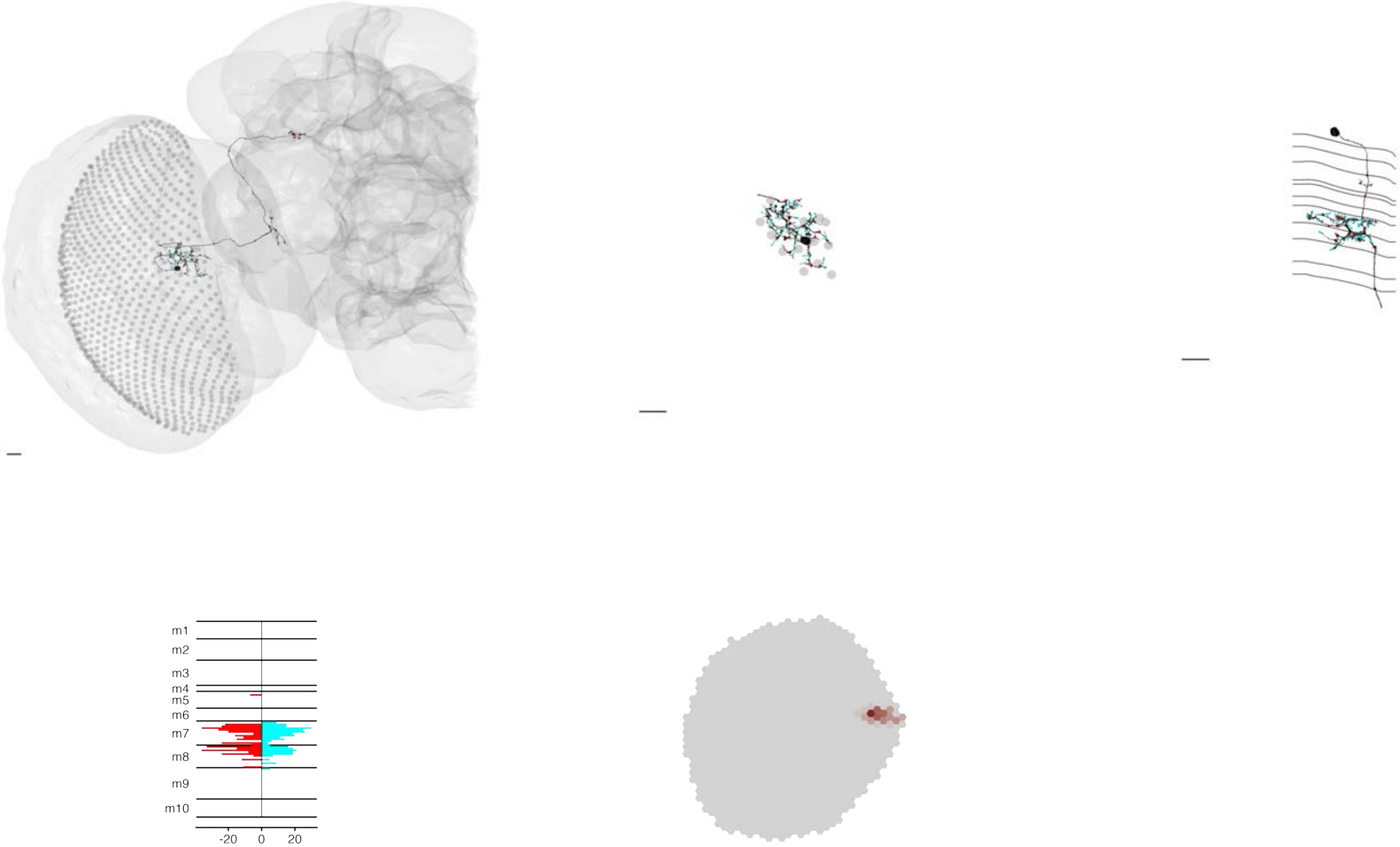

### MeTu4a_720575940617311574

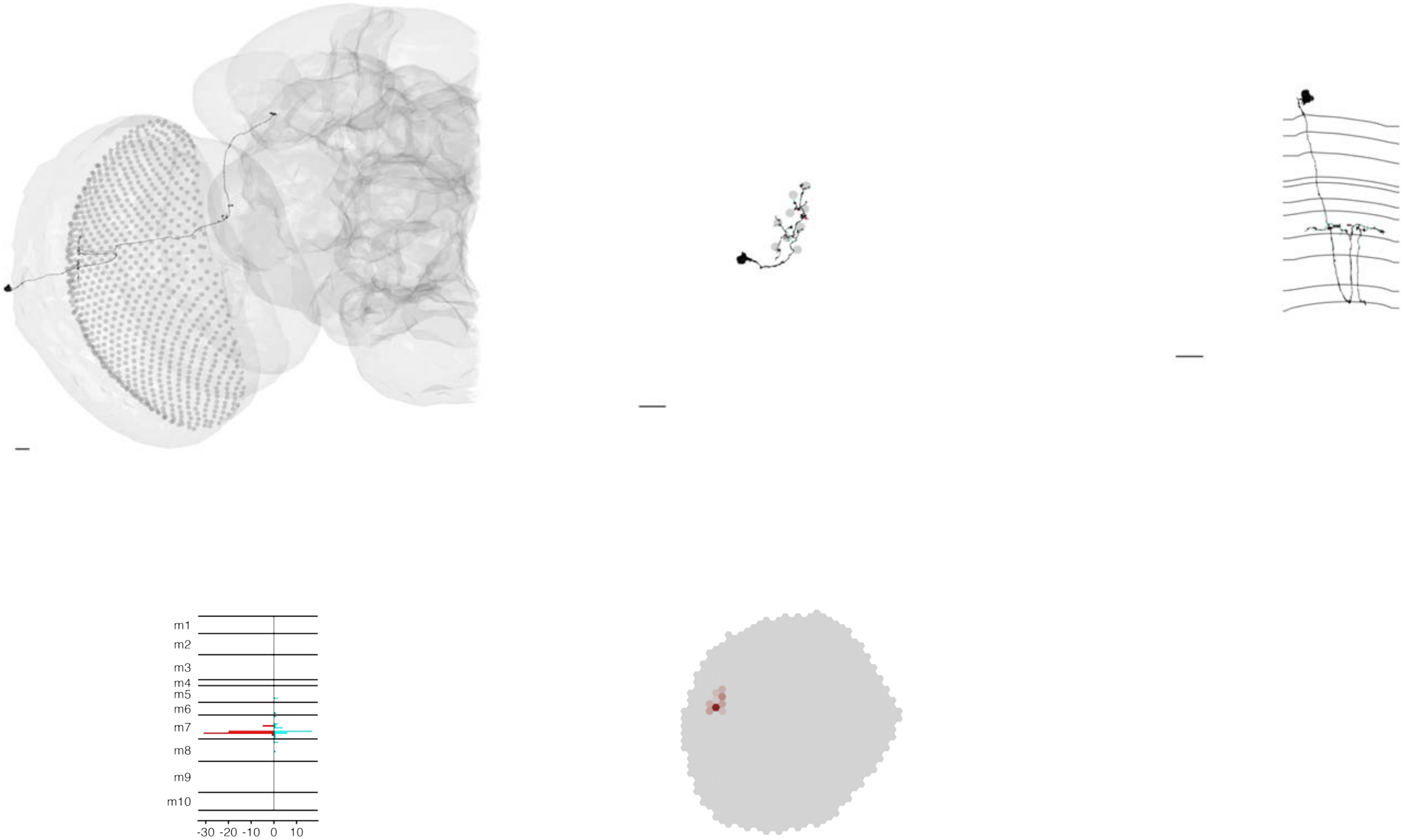

### MeTu4a_720575940618761044

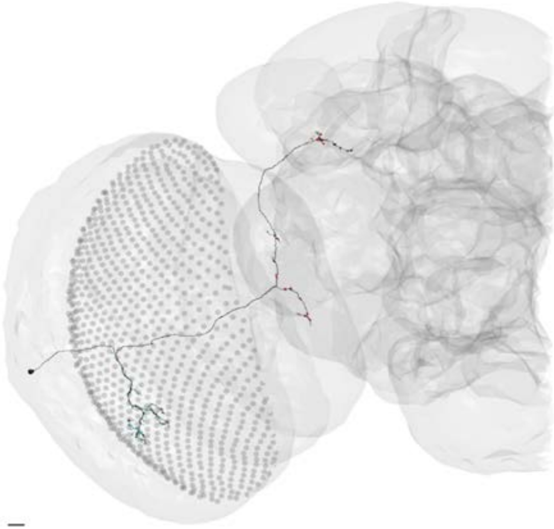

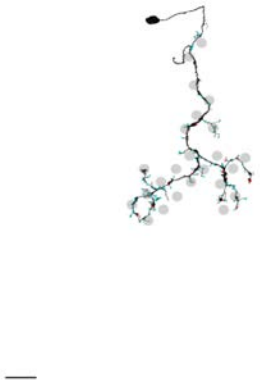

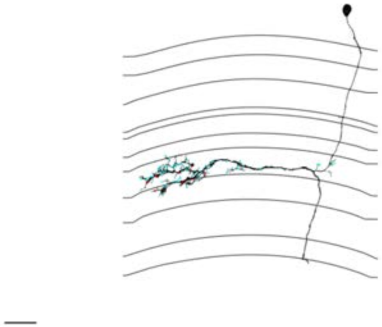

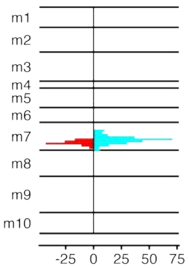


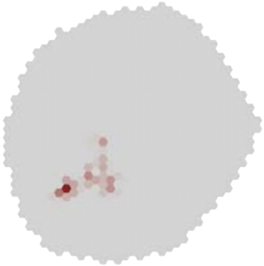

### MeTu4a_720575940619279064

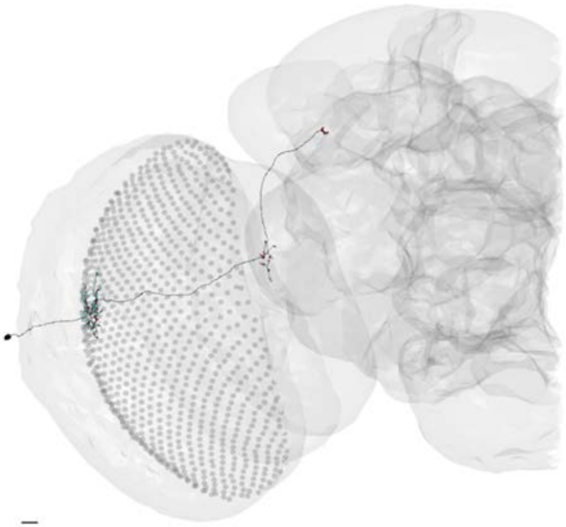

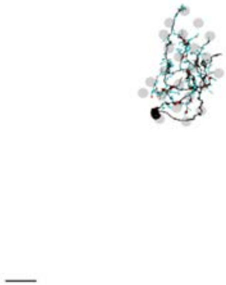

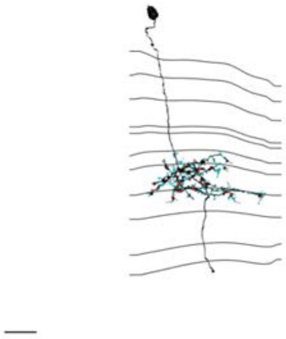

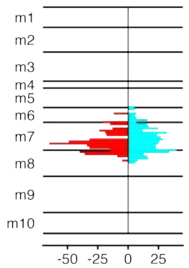


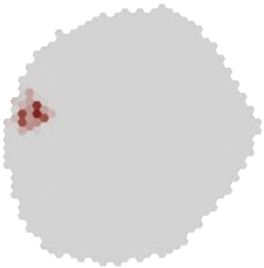

MeTu4a_720575940619324069

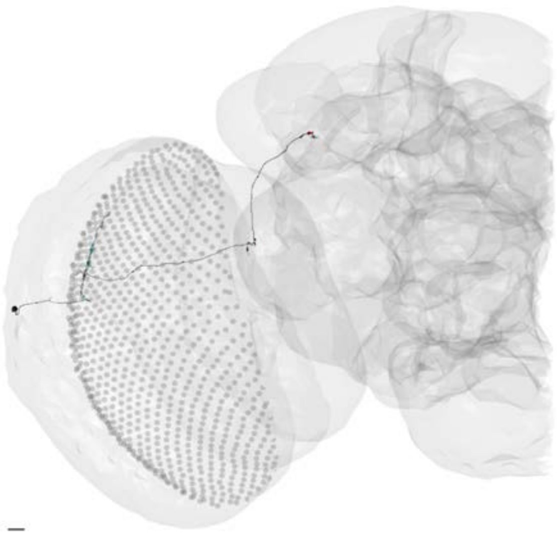

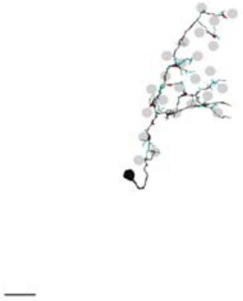

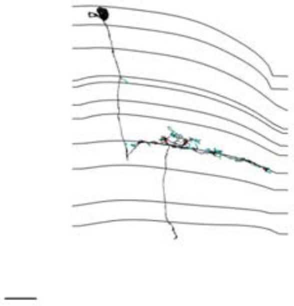

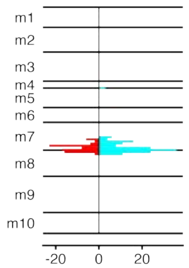


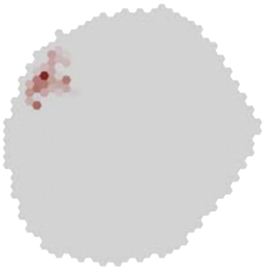

### MeTu4a_720575940620577974

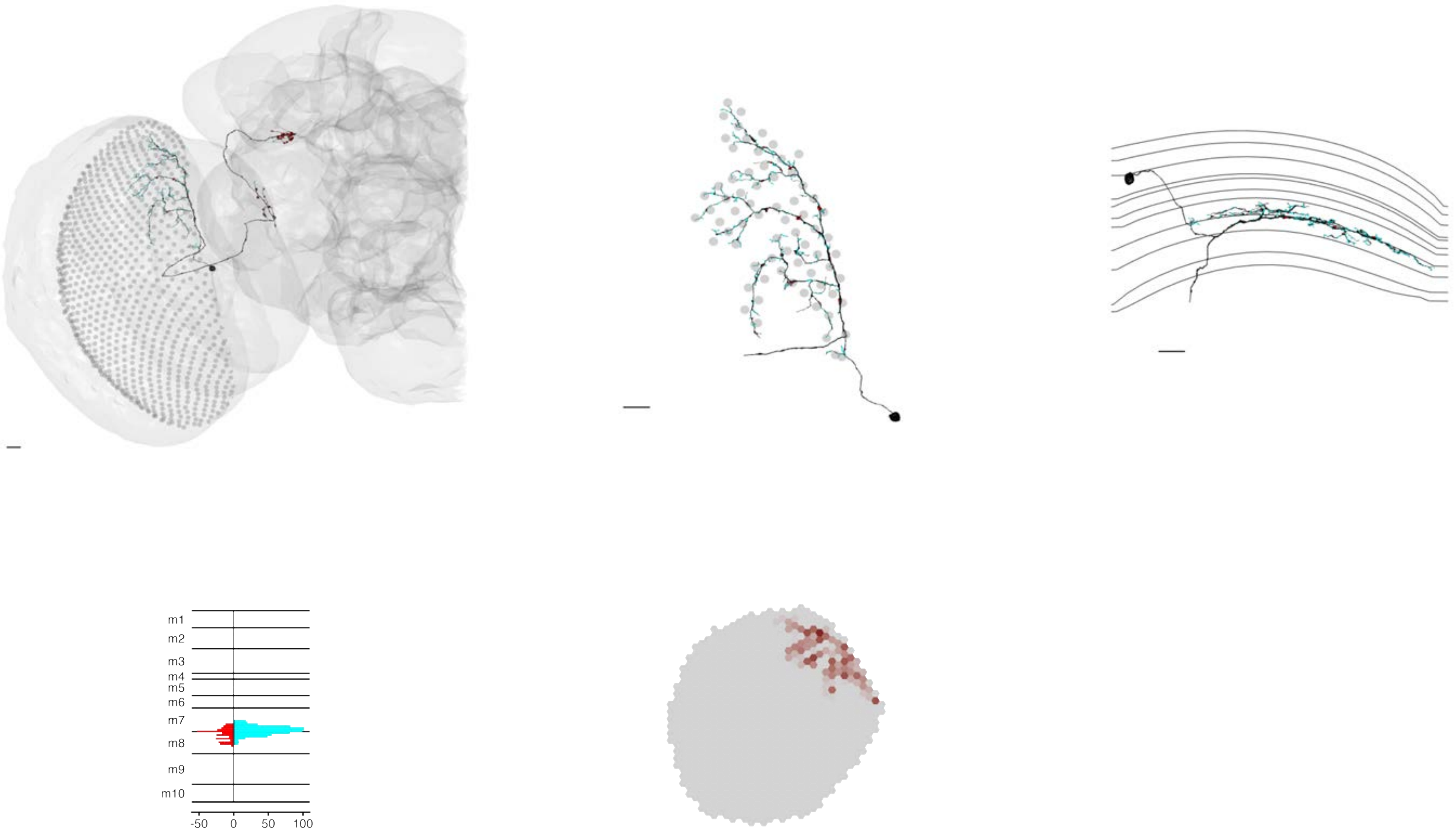

### MeTu4a_720575940620927472

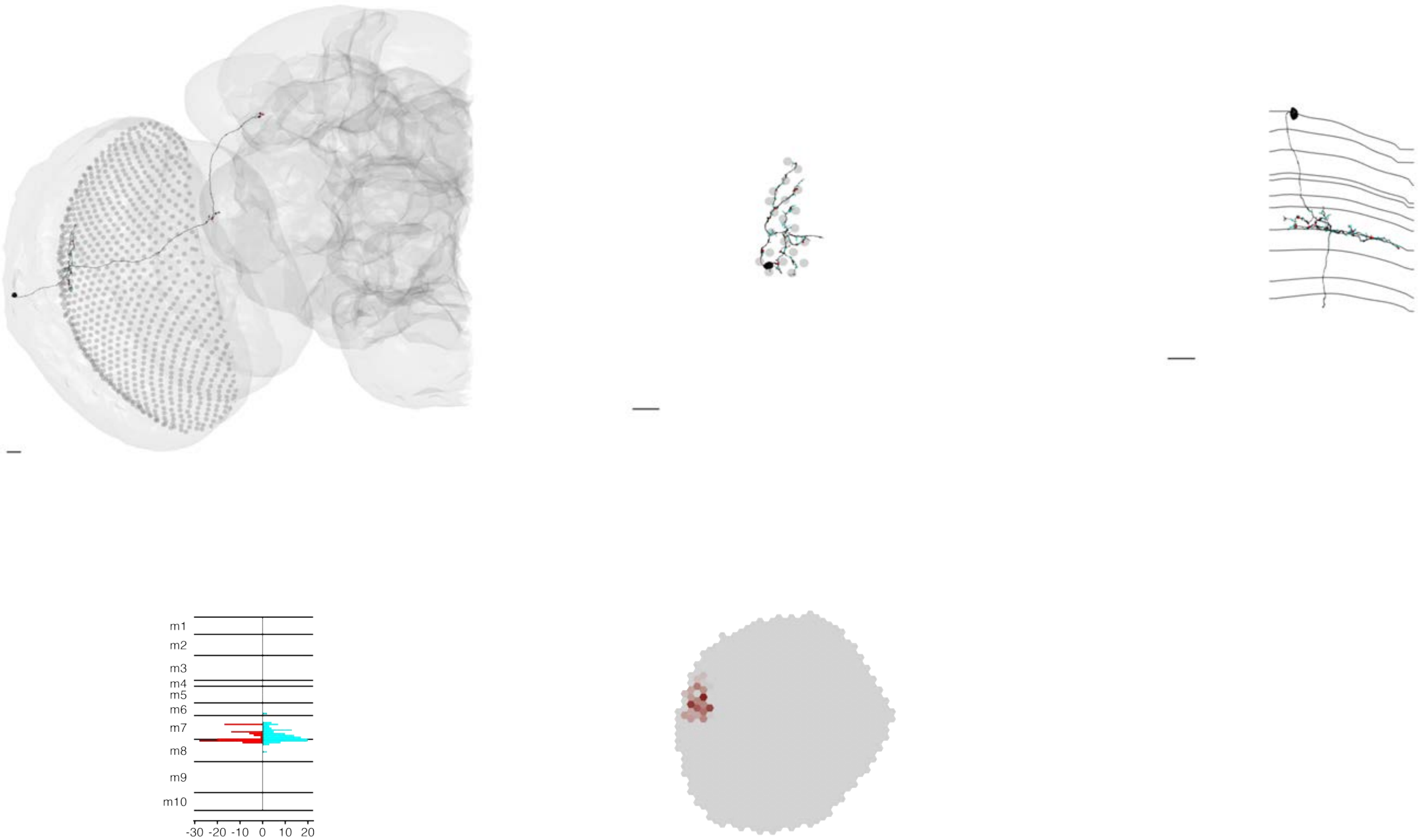

### MeTu4a_720575940622112183

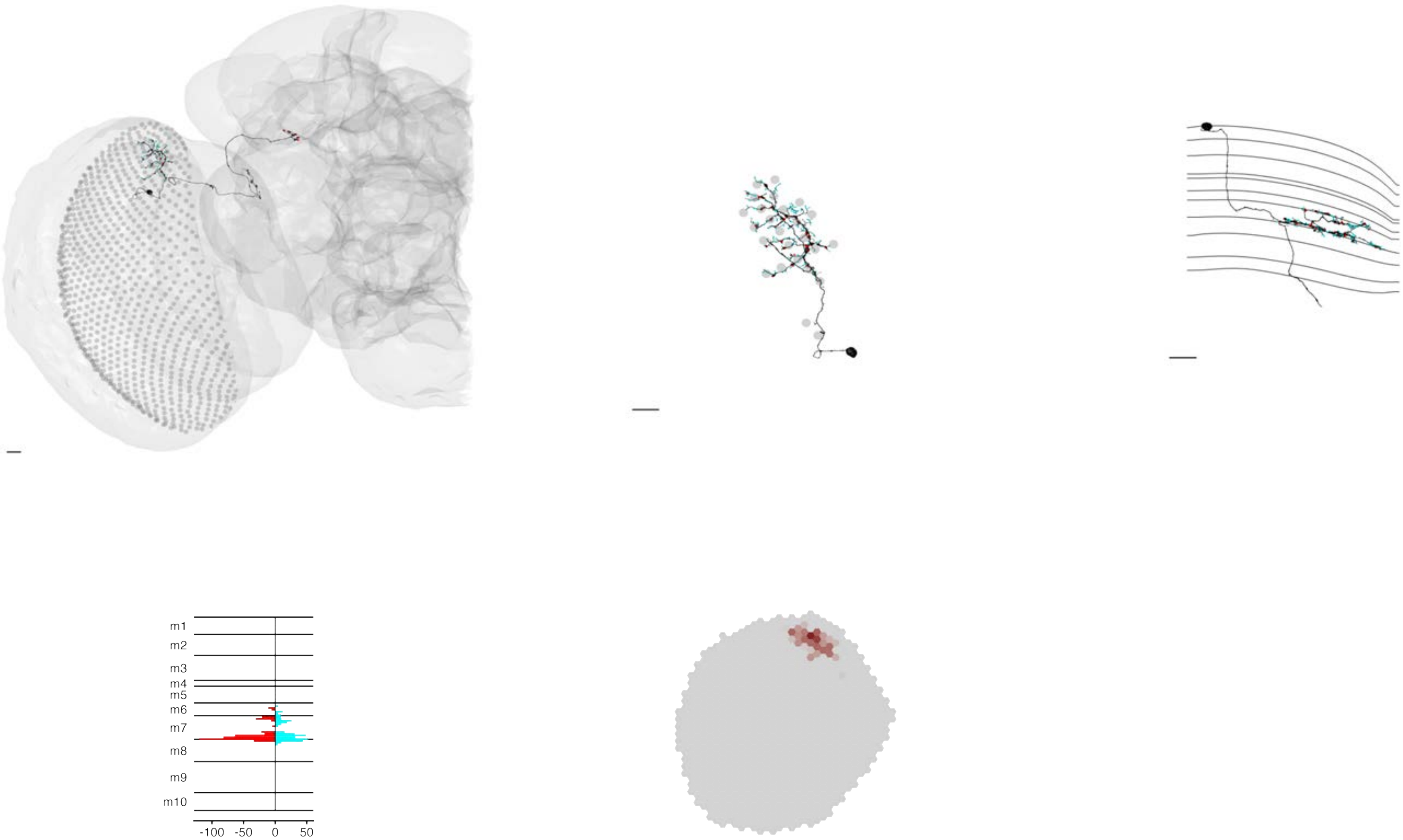

MeTu4a_720575940622781865

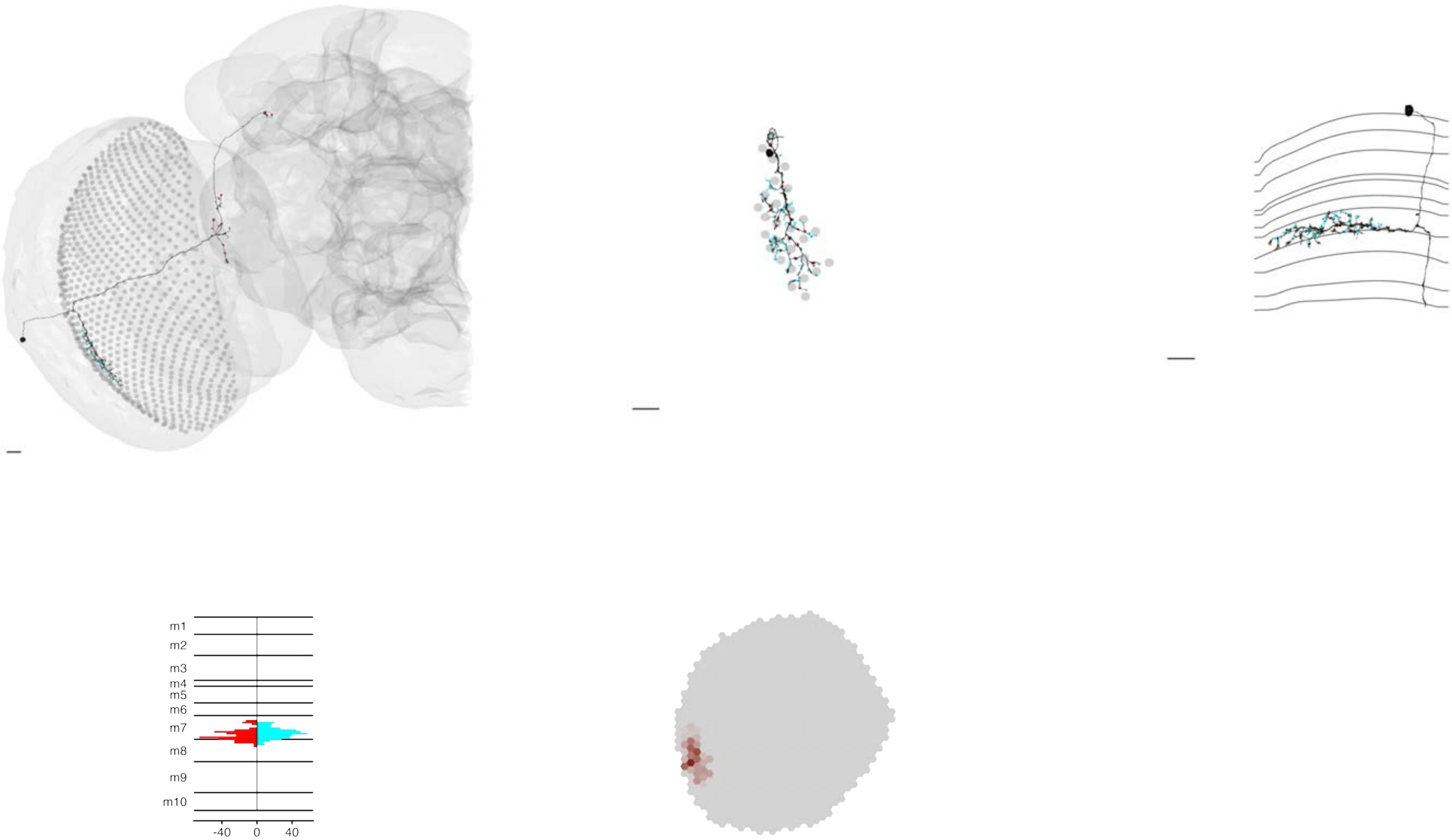

### MeTu4a_720575940623175182

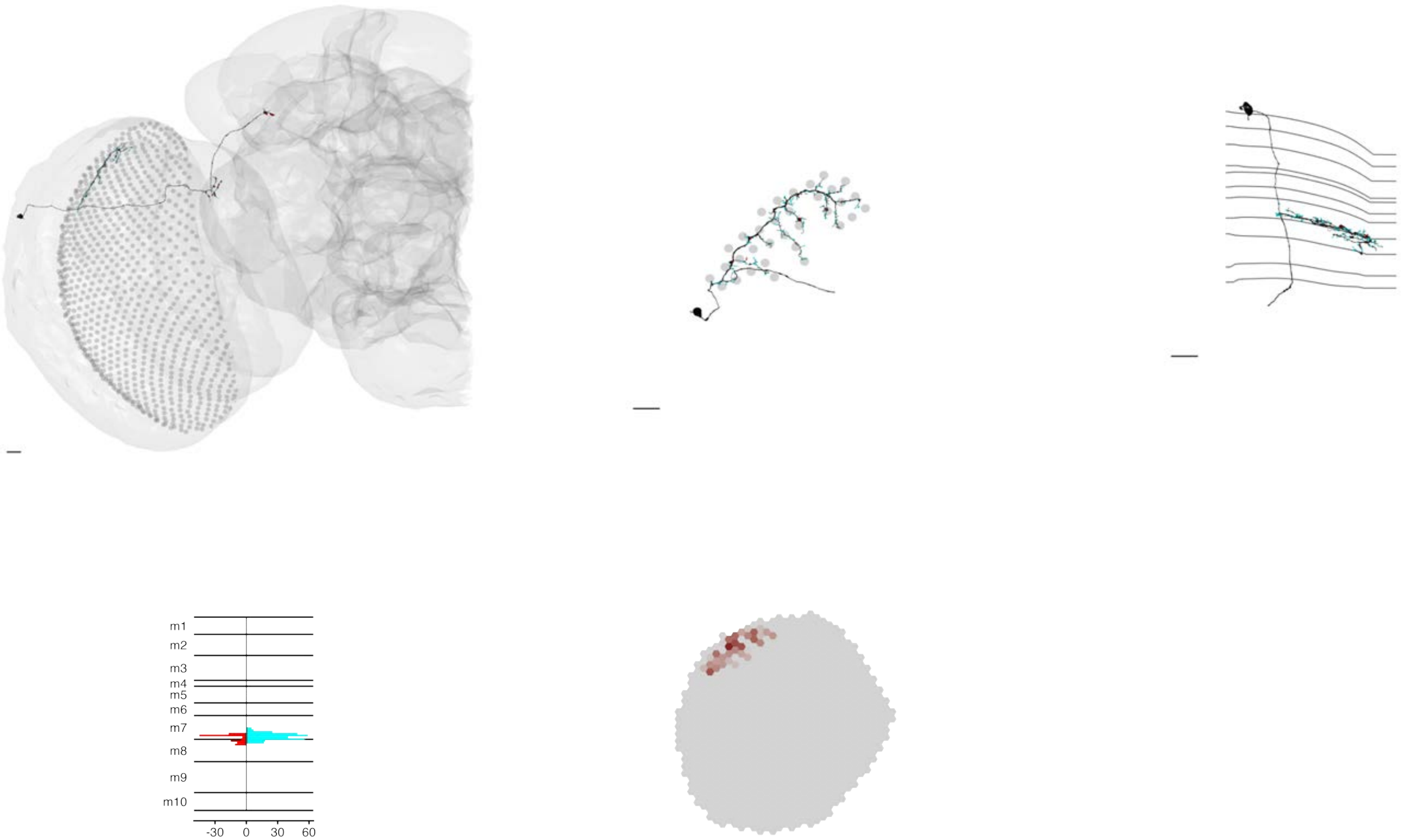

### MeTu4a_720575940623251455

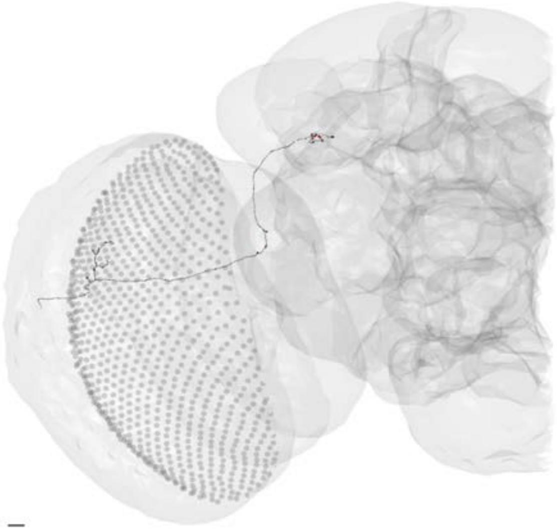

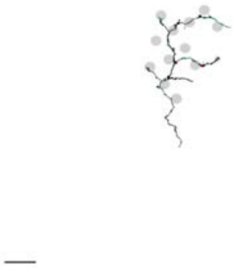

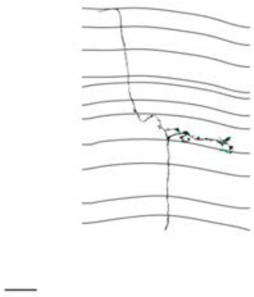

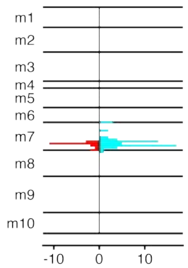


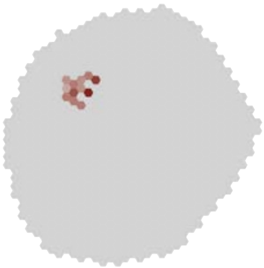

### MeTu4a_720575940623343613

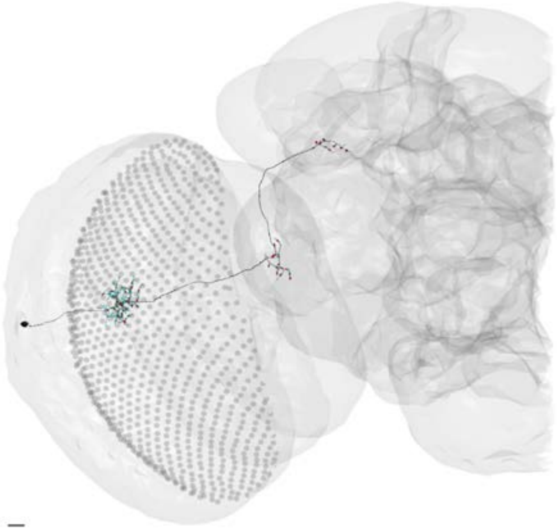

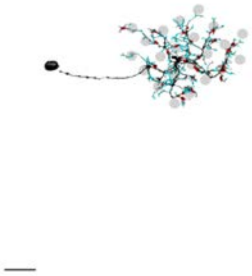

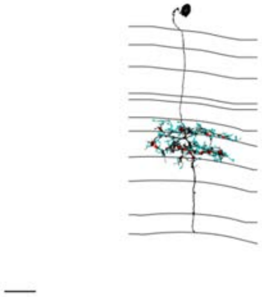

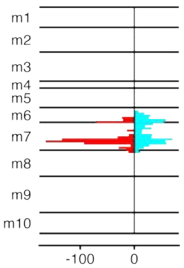


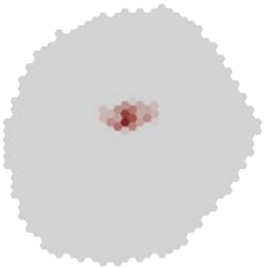

### MeTu4a_720575940623425543

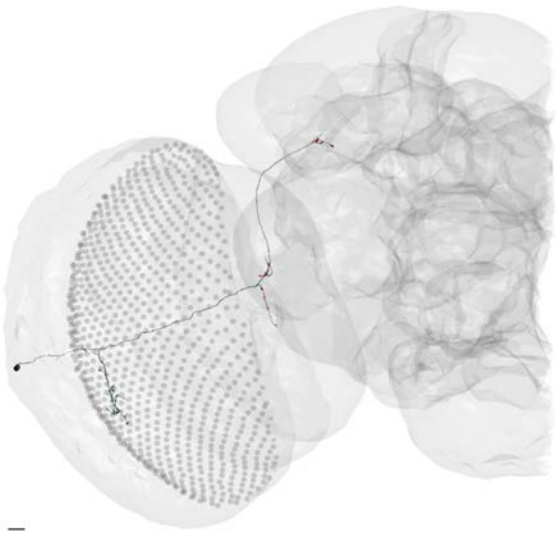

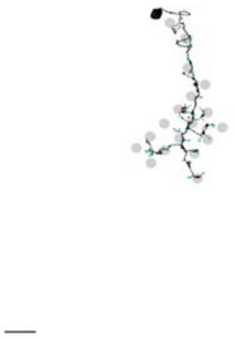

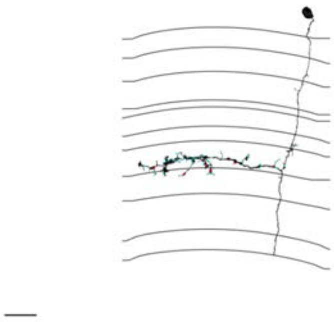

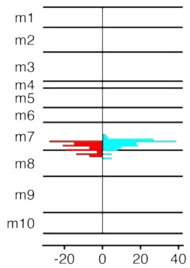


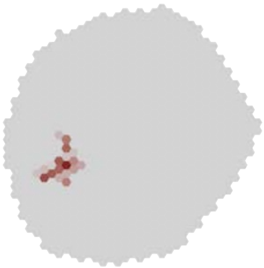

MeTu4a_720575940623987214

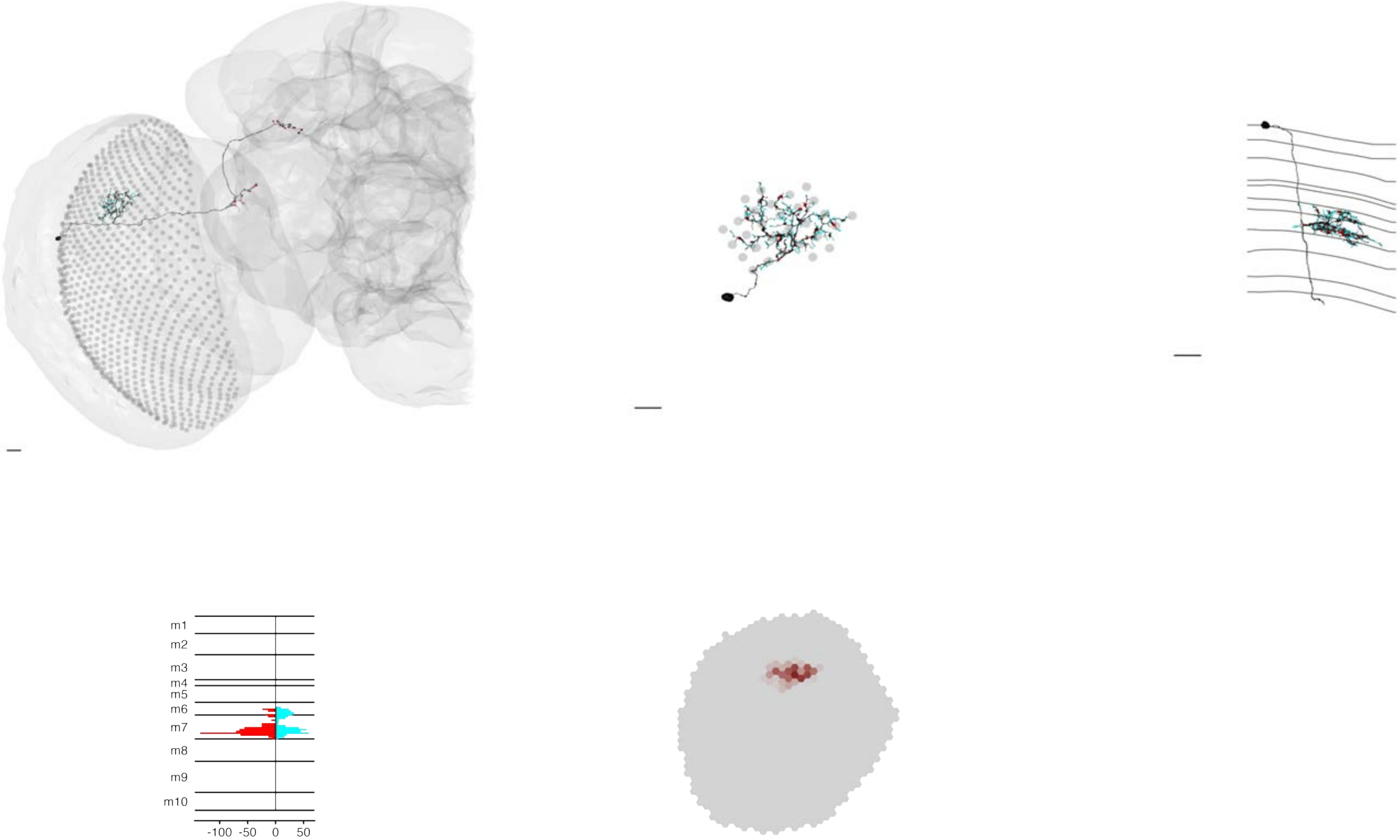

### MeTu4a_720575940624370765

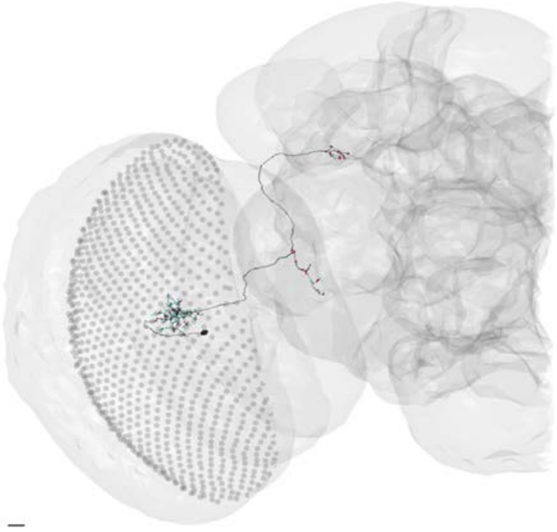

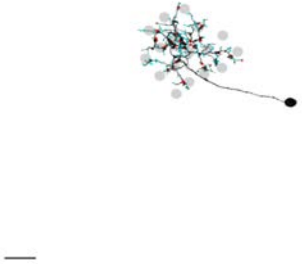

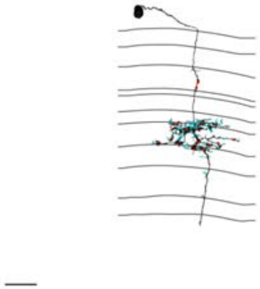

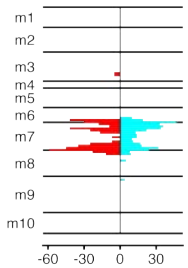


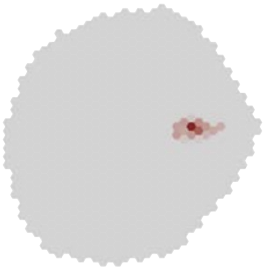

### MeTu4a_720575940624585725

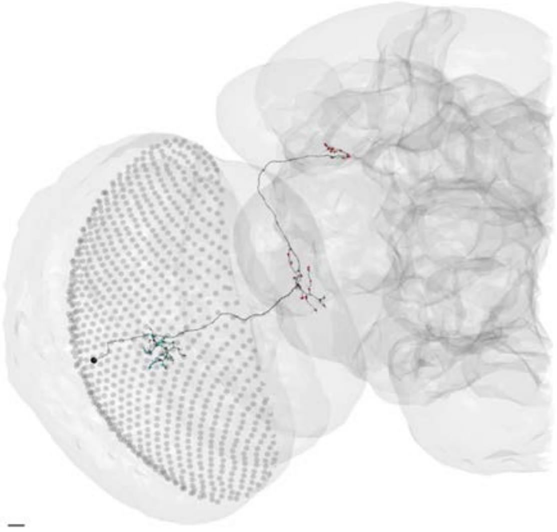

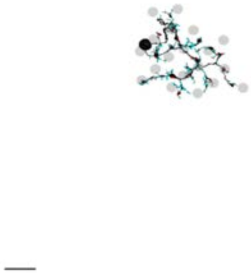

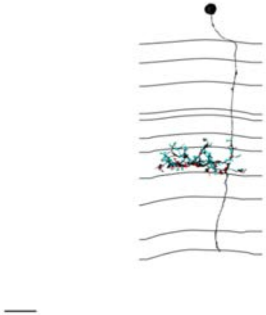

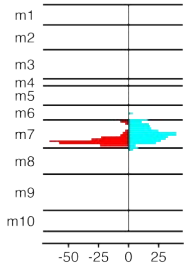


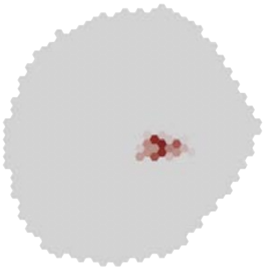

### MeTu4a_720575940624728551

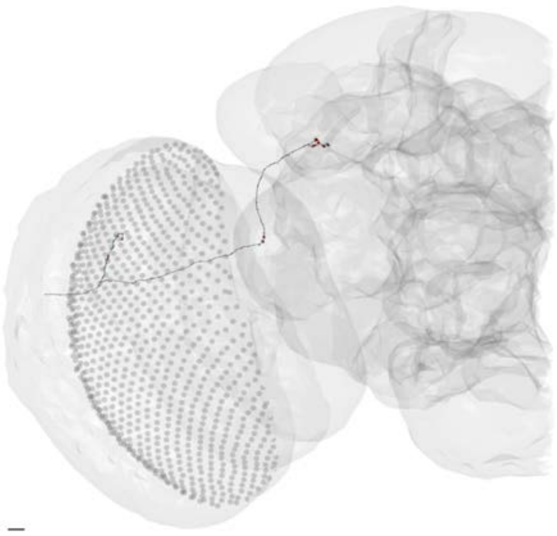

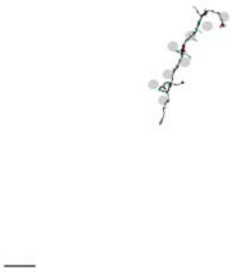

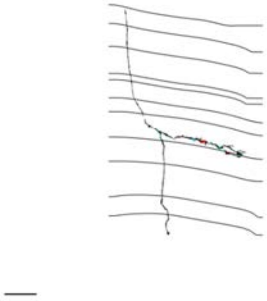

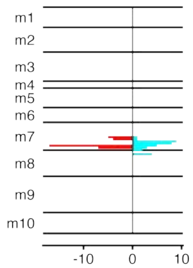


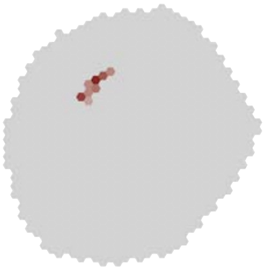

### MeTu4a_720575940625067312

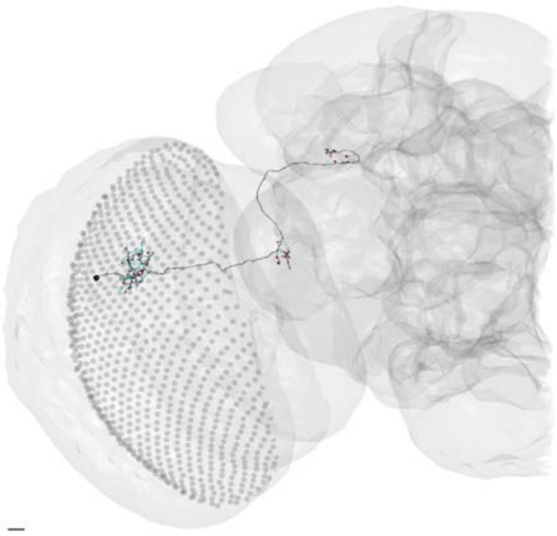

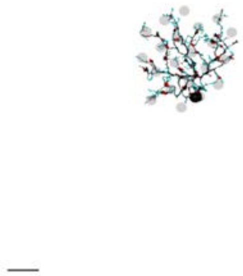

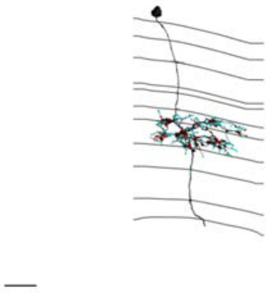

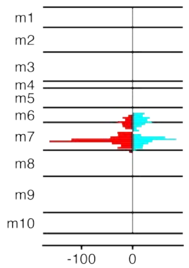


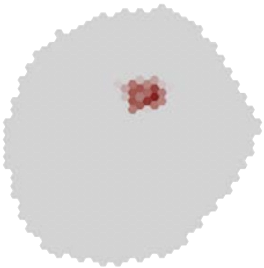

### MeTu4a_720575940625230323

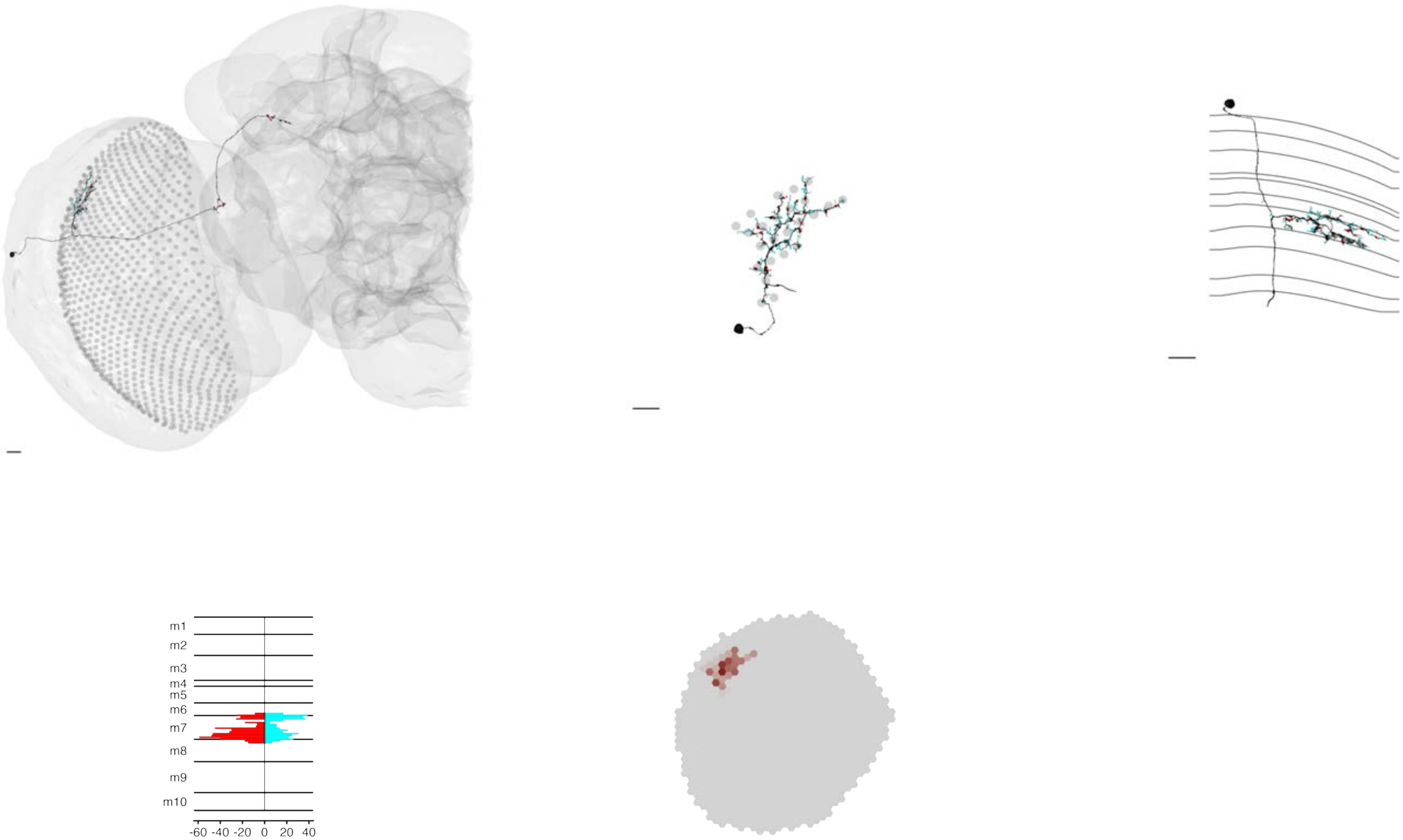

### MeTu4a_720575940625237396

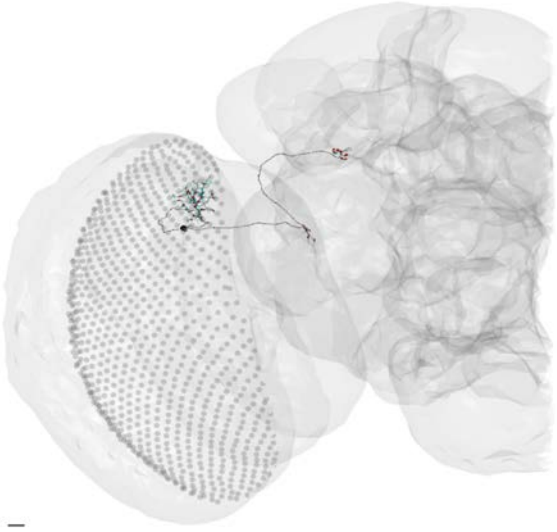

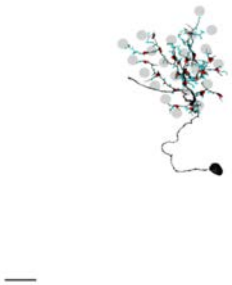

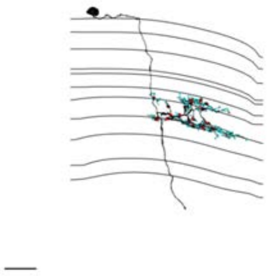

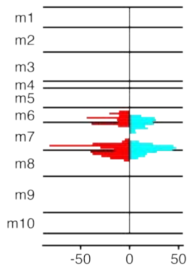


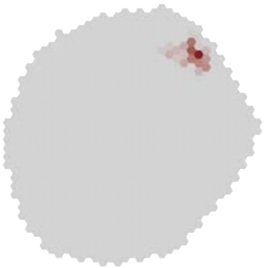

### MeTu4a_720575940625622018

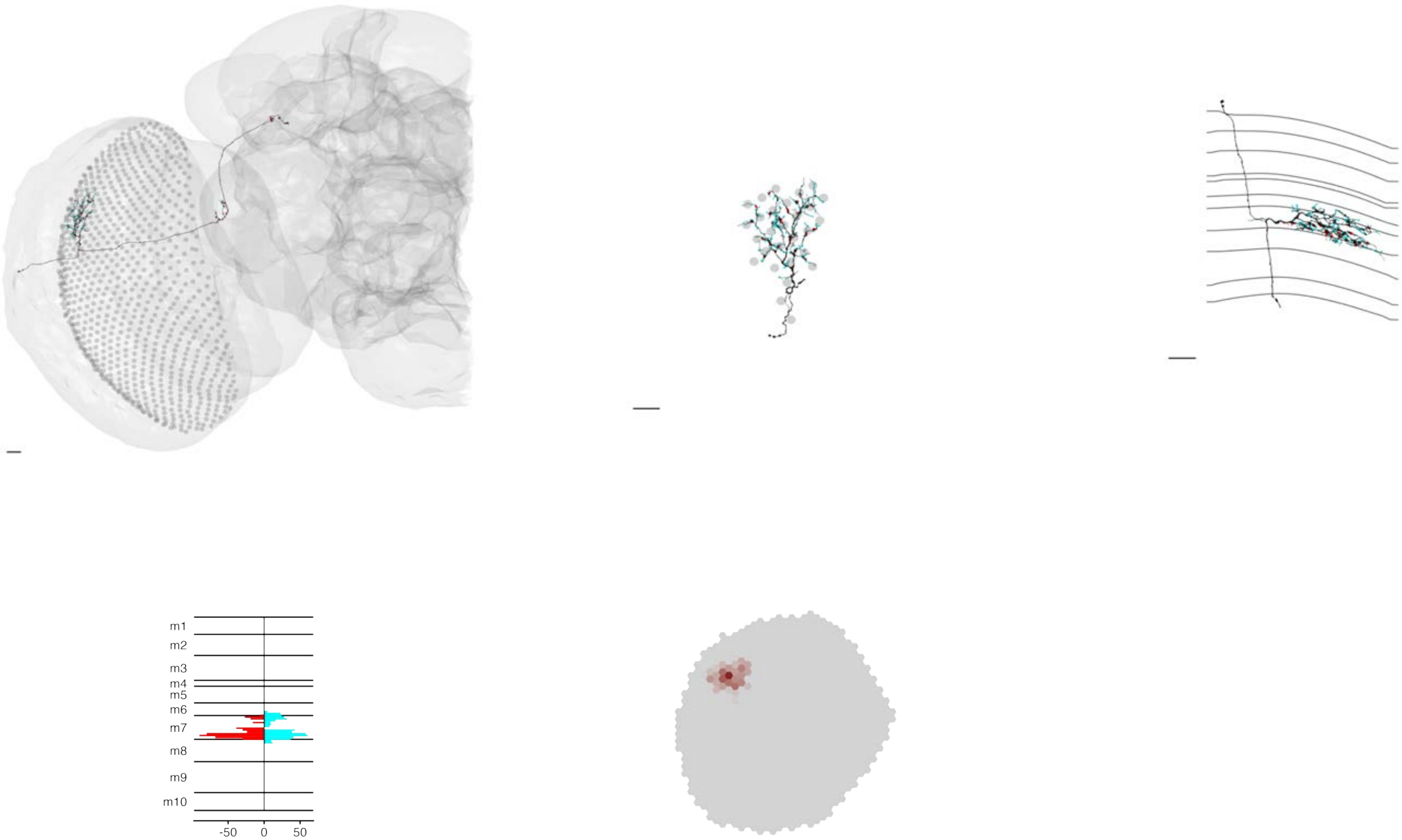

### MeTu4a_720575940626797841

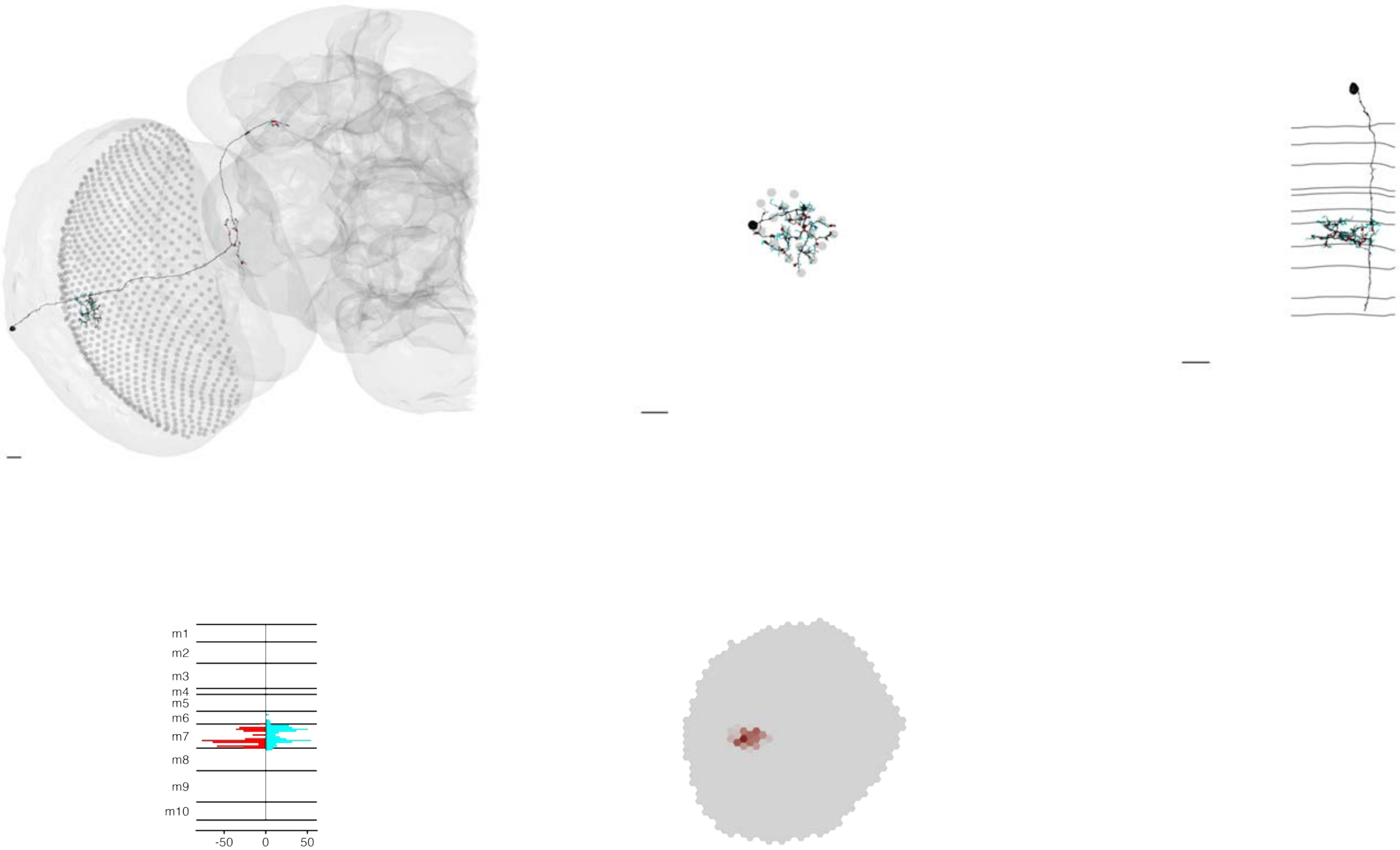

### MeTu4a_720575940627305212

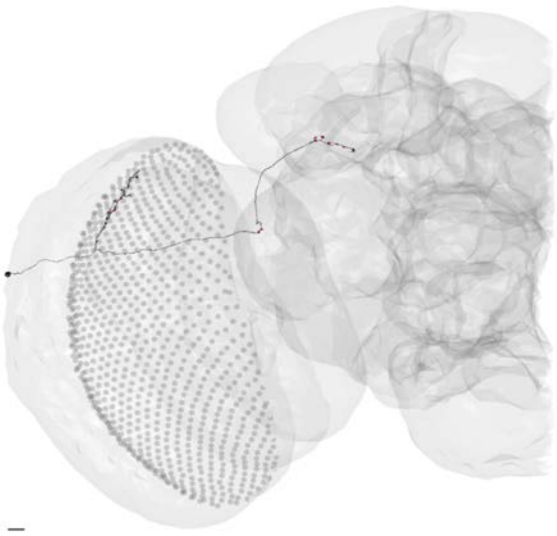

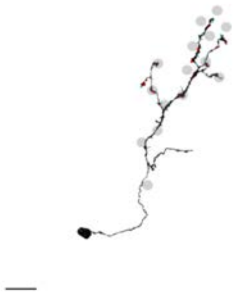

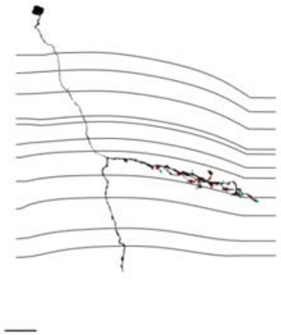

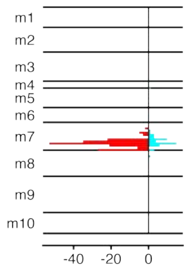


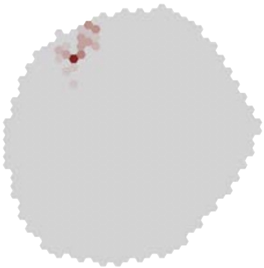

### MeTu4a_720575940627892234

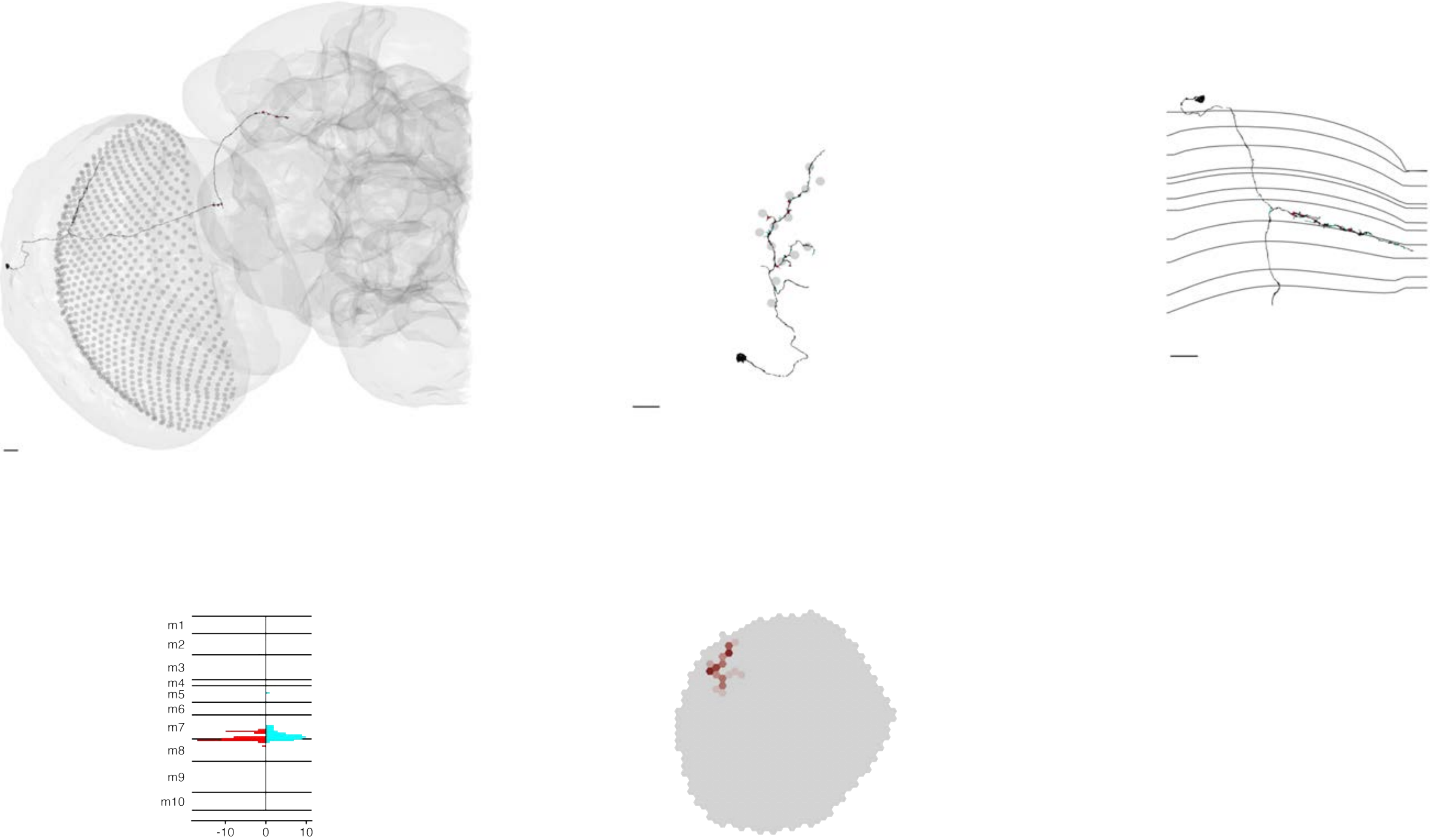

### MeTu4a_720575940628100395

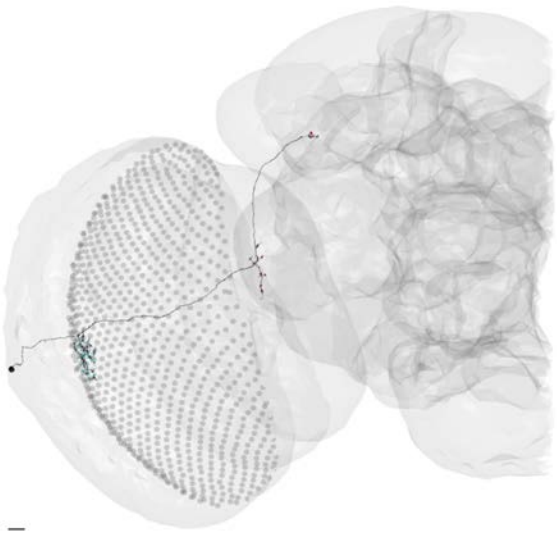

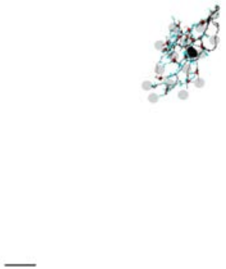

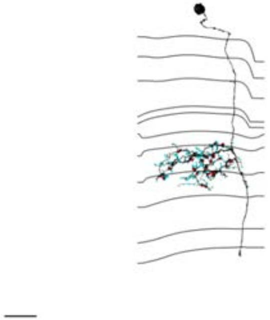

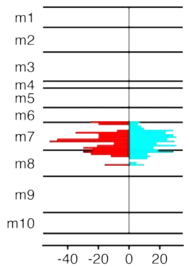


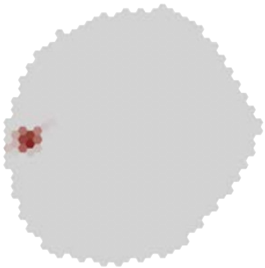

### MeTu4a_720575940628450503

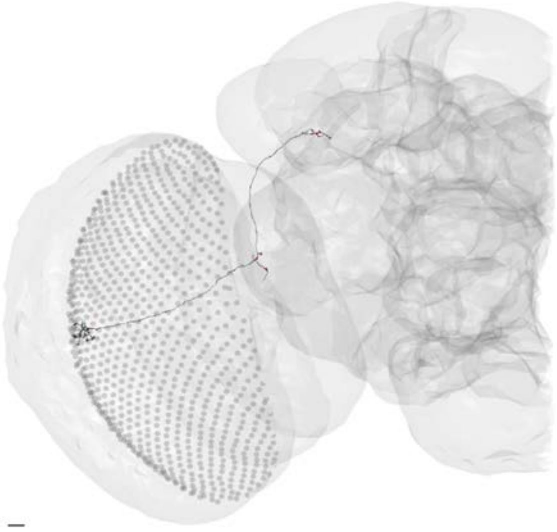

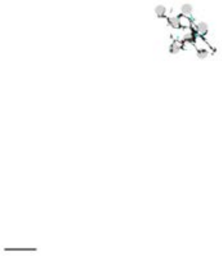

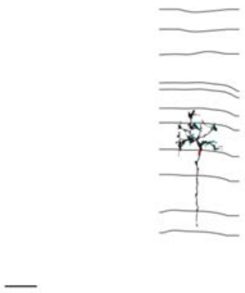

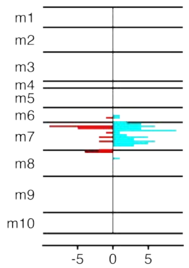


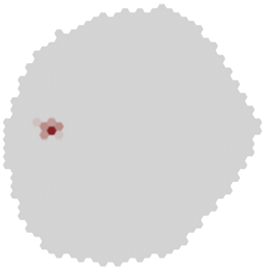

### MeTu4a_720575940628801900

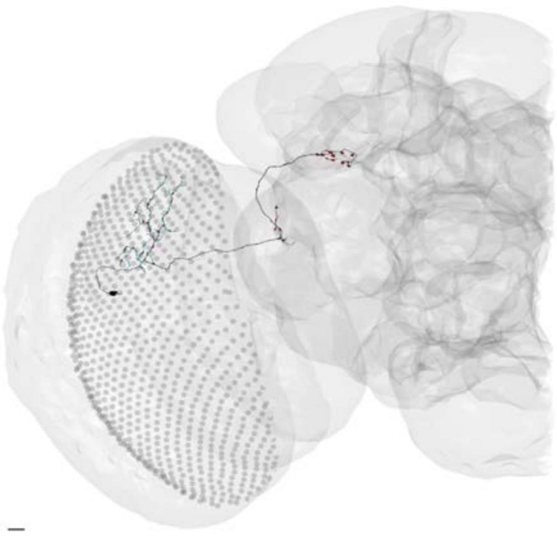

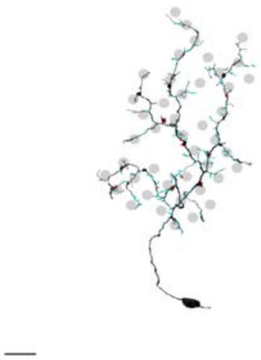

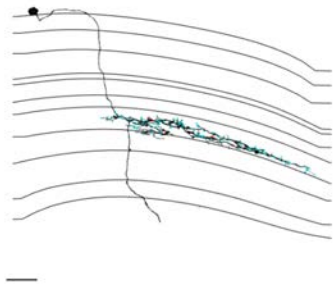

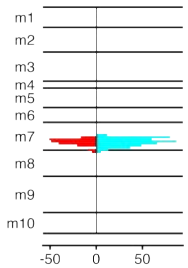


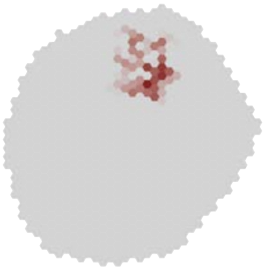

### MeTu4a_720575940630418682

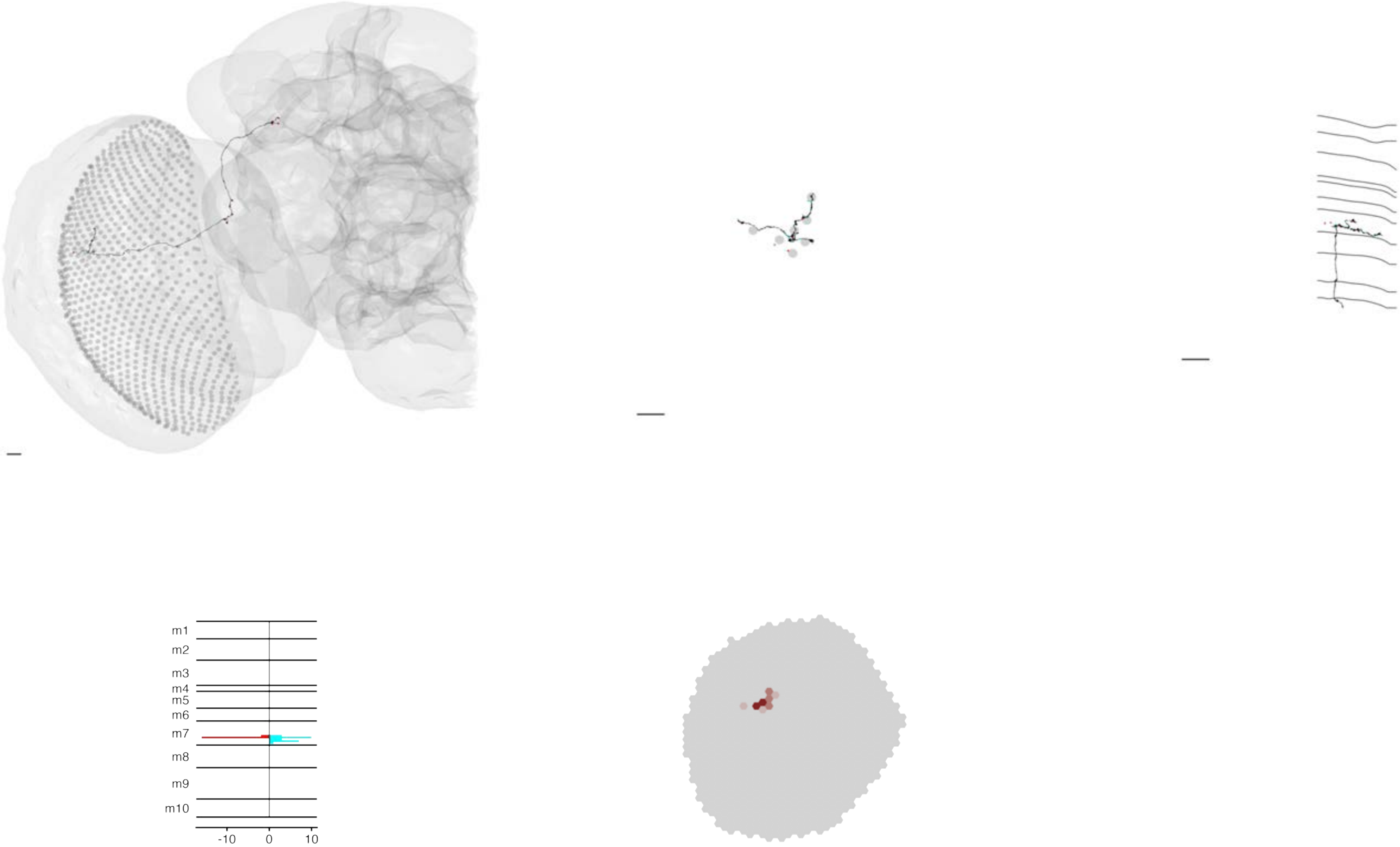

### MeTu4a_720575940630483564

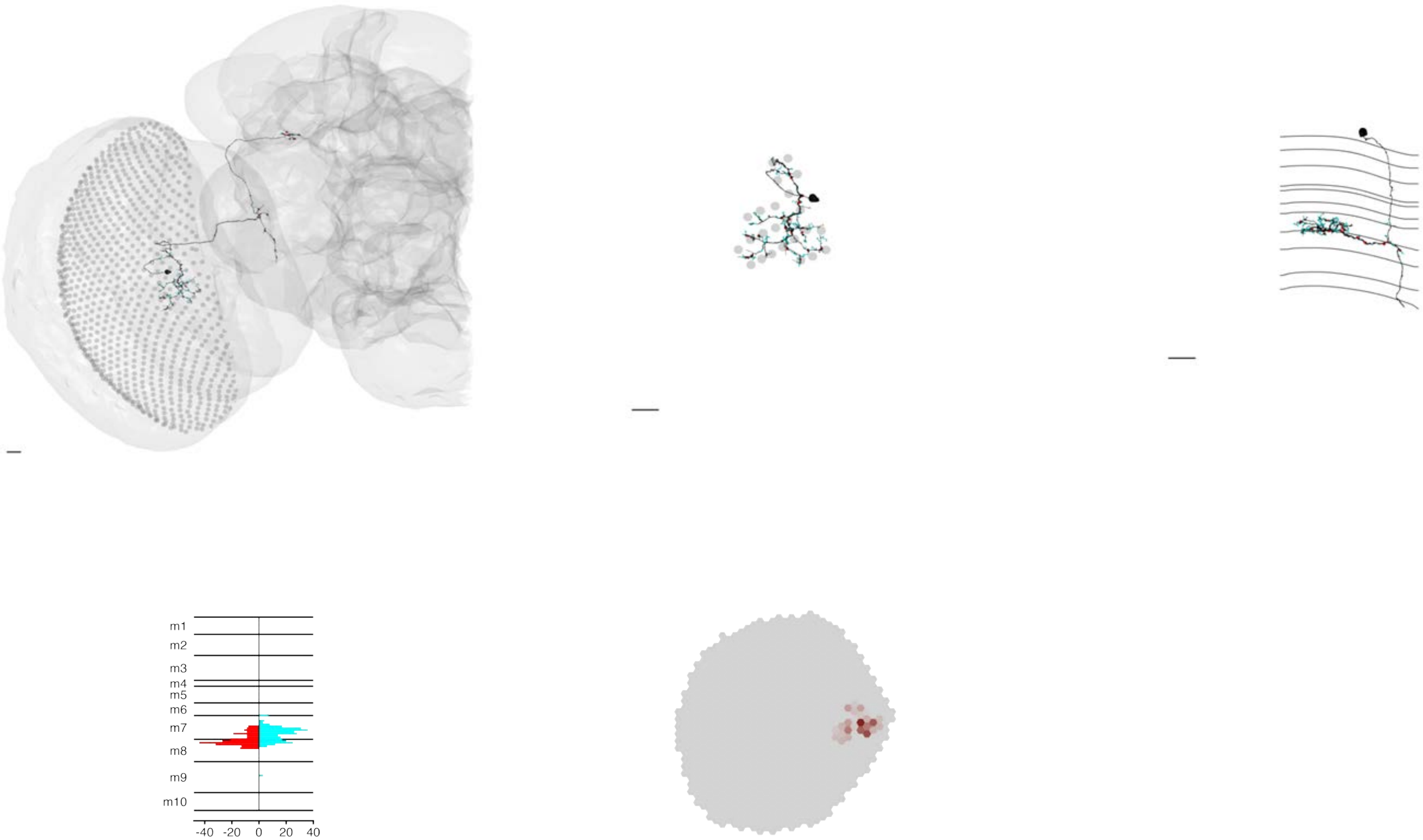

MeTu4a_720575940630619985

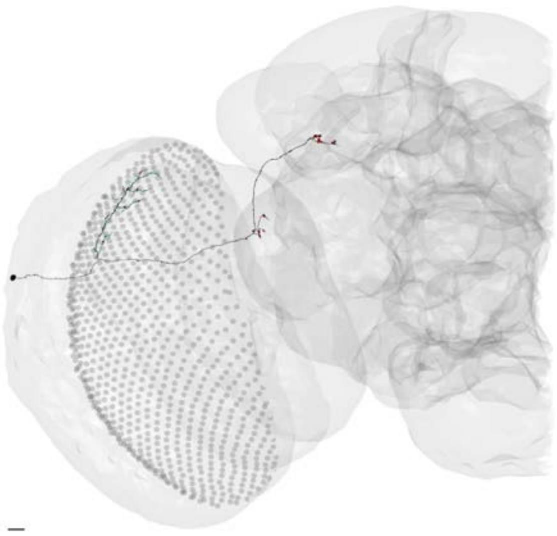

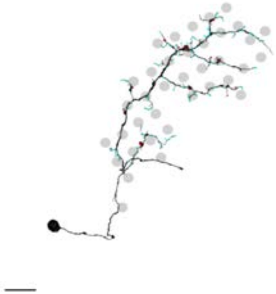

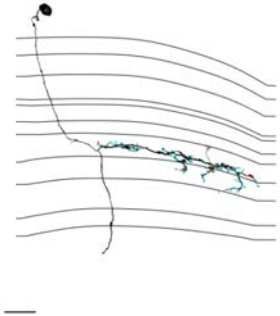

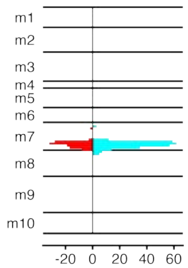


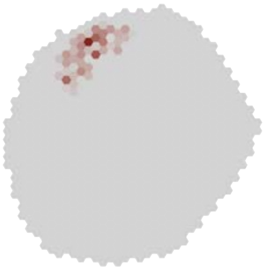

MeTu4a_720575940630692412

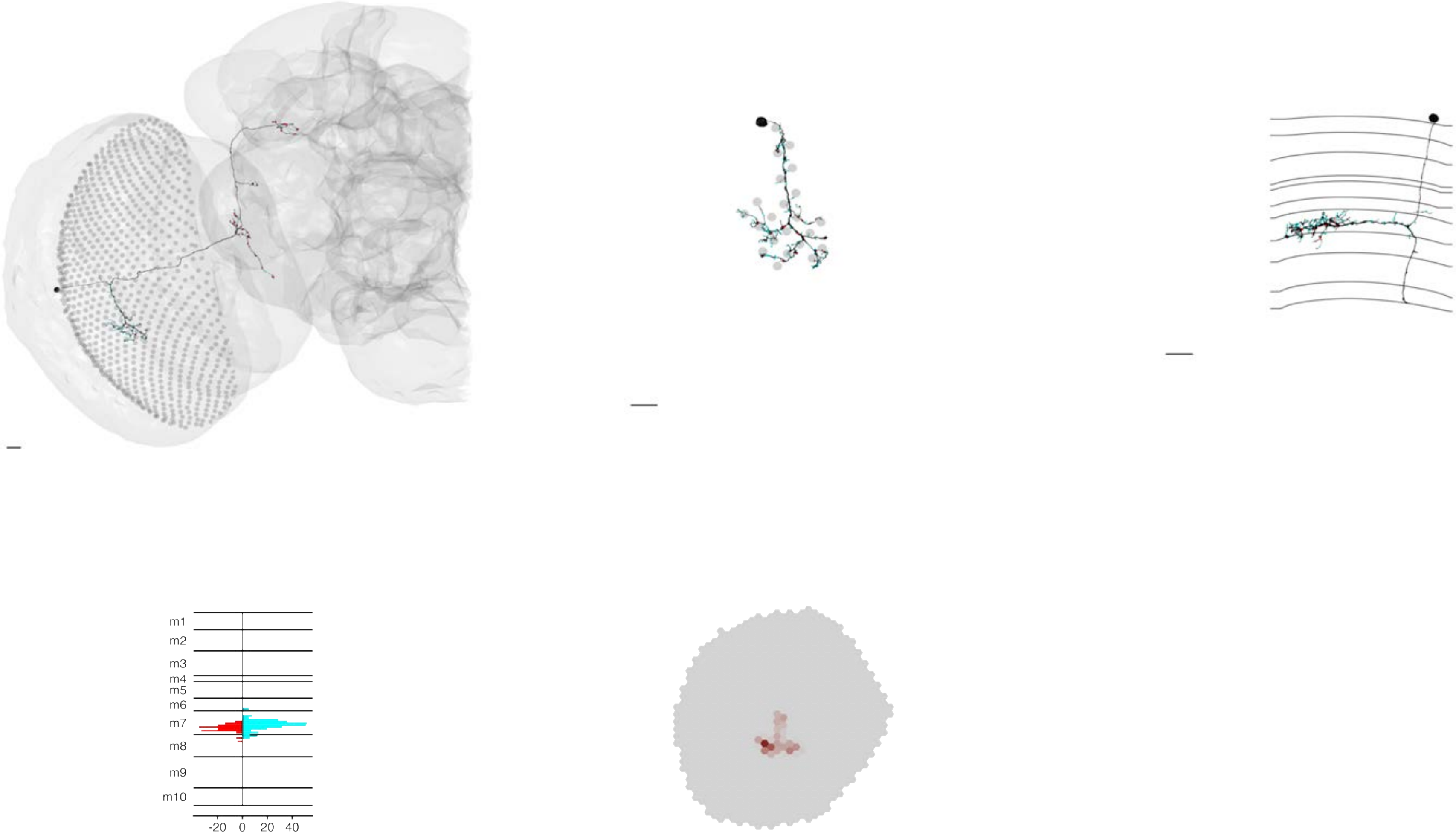

MeTu4a_720575940631170809

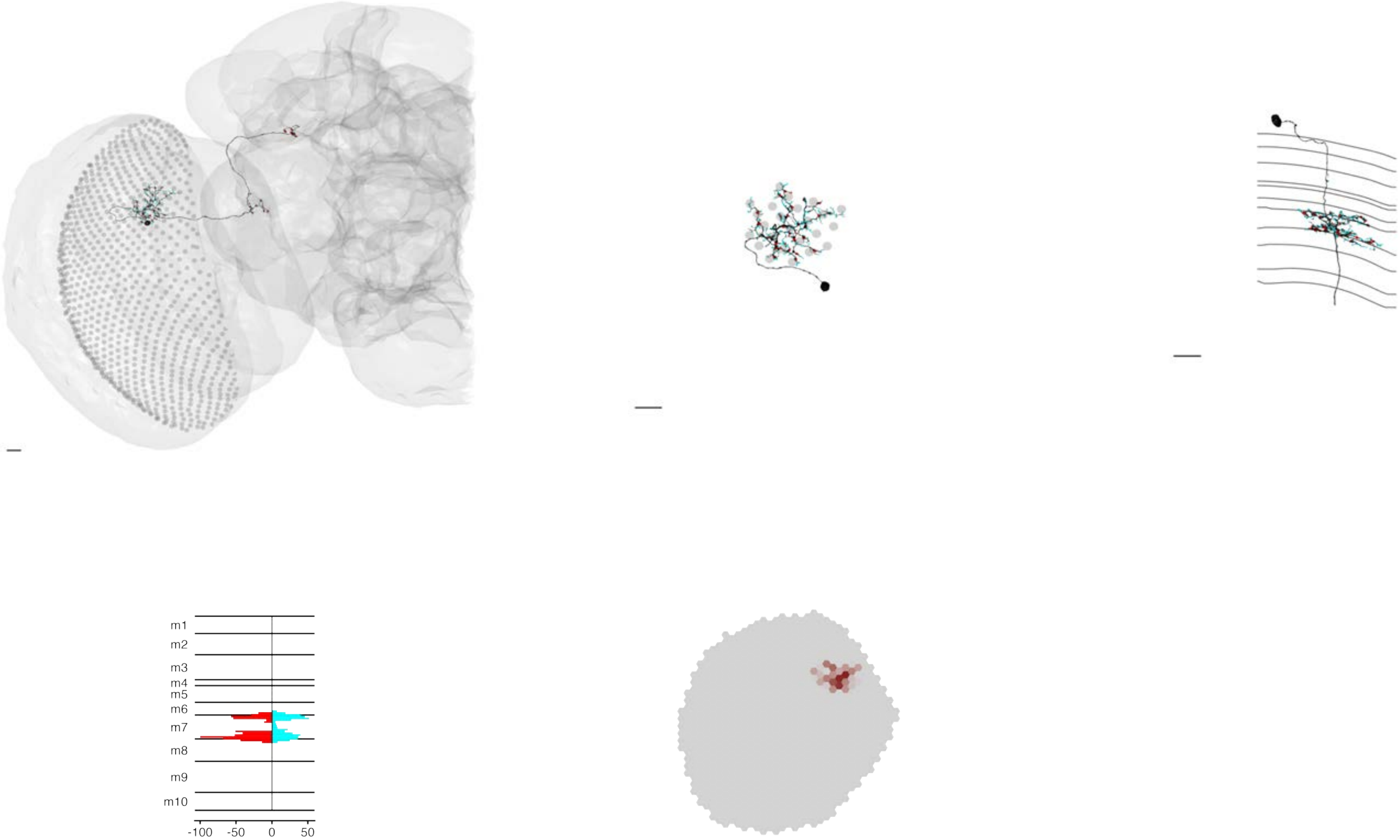

### MeTu4a_720575940631232249

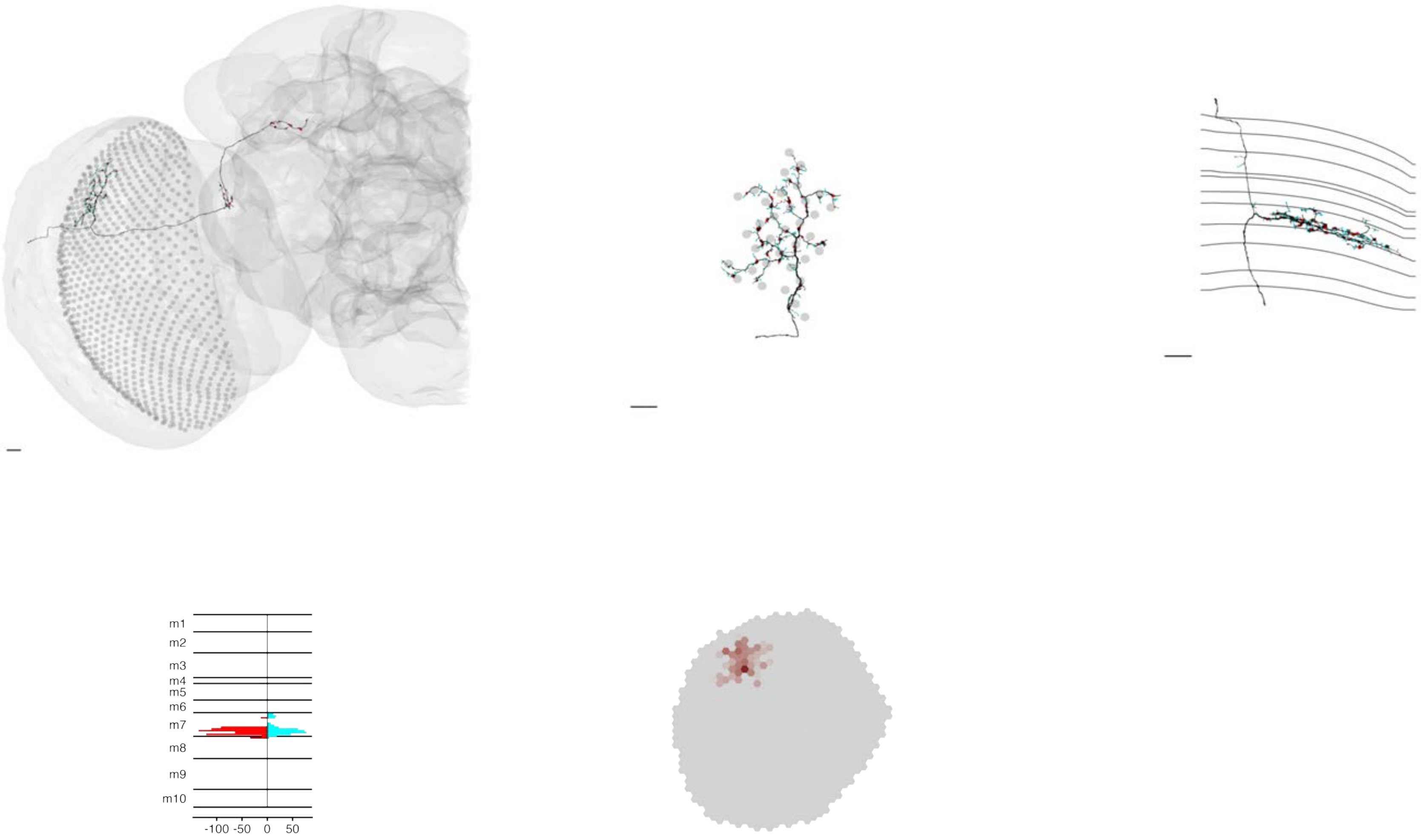

MeTu4a_720575940631346895

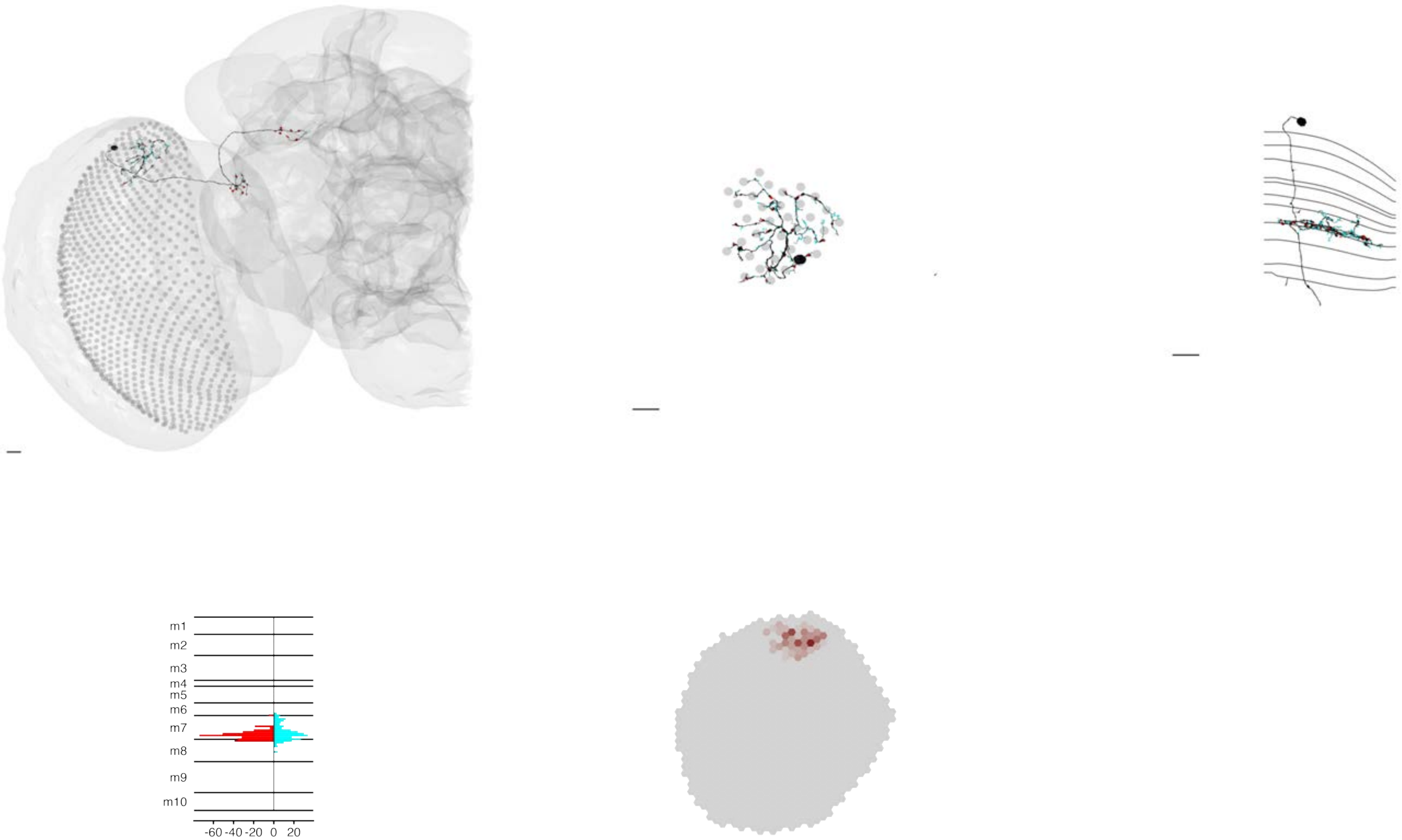

MeTu4a_720575940631366351

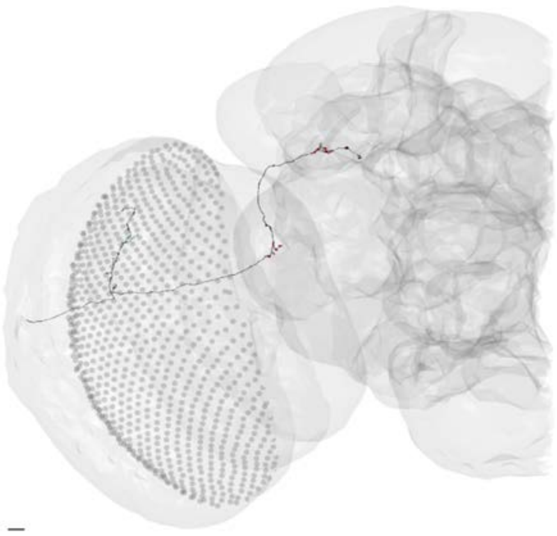

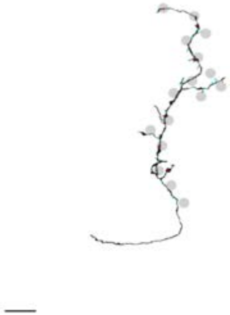

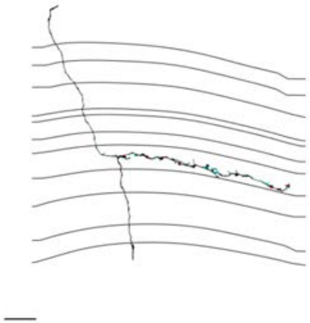

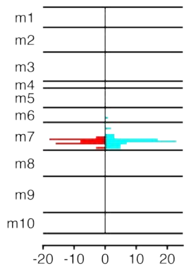


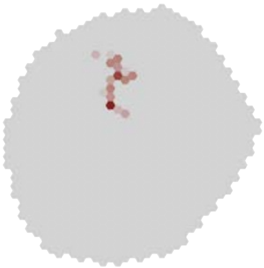

### MeTu4a_720575940631634975

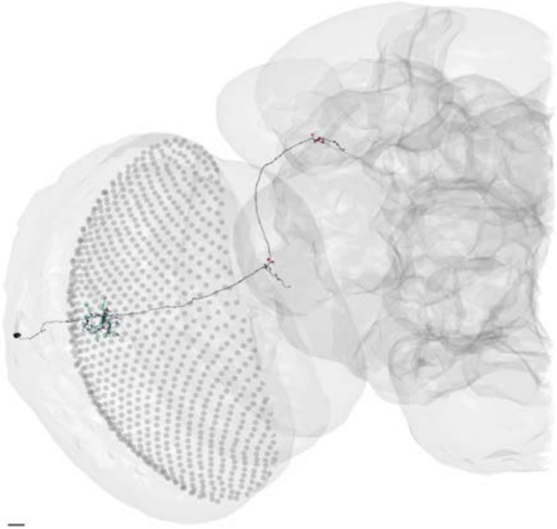

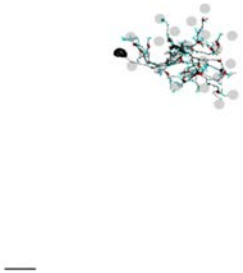

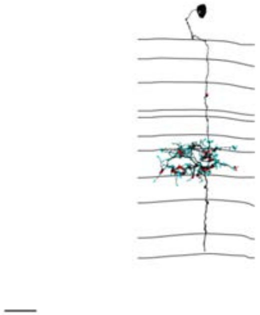

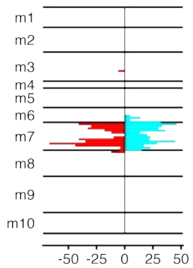


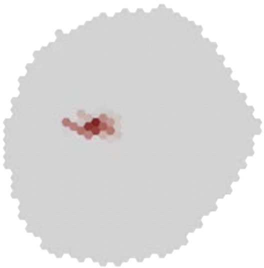

### MeTu4a_720575940632035882

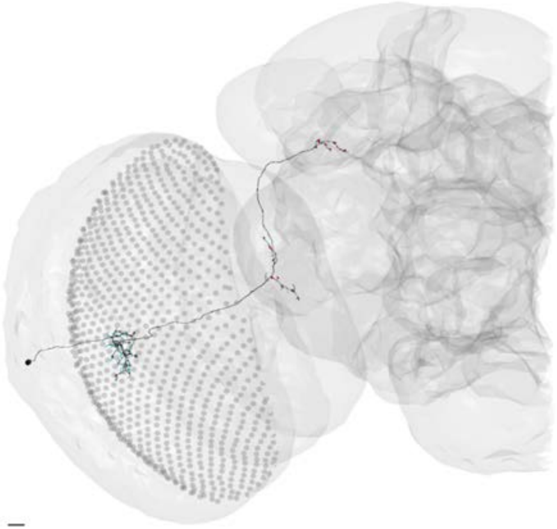

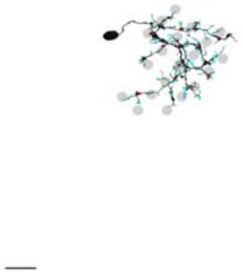

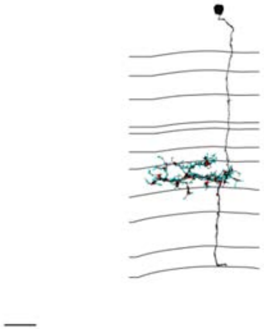

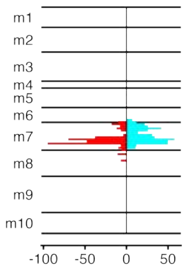


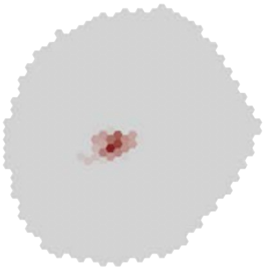

### MeTu4a_720575940632085900

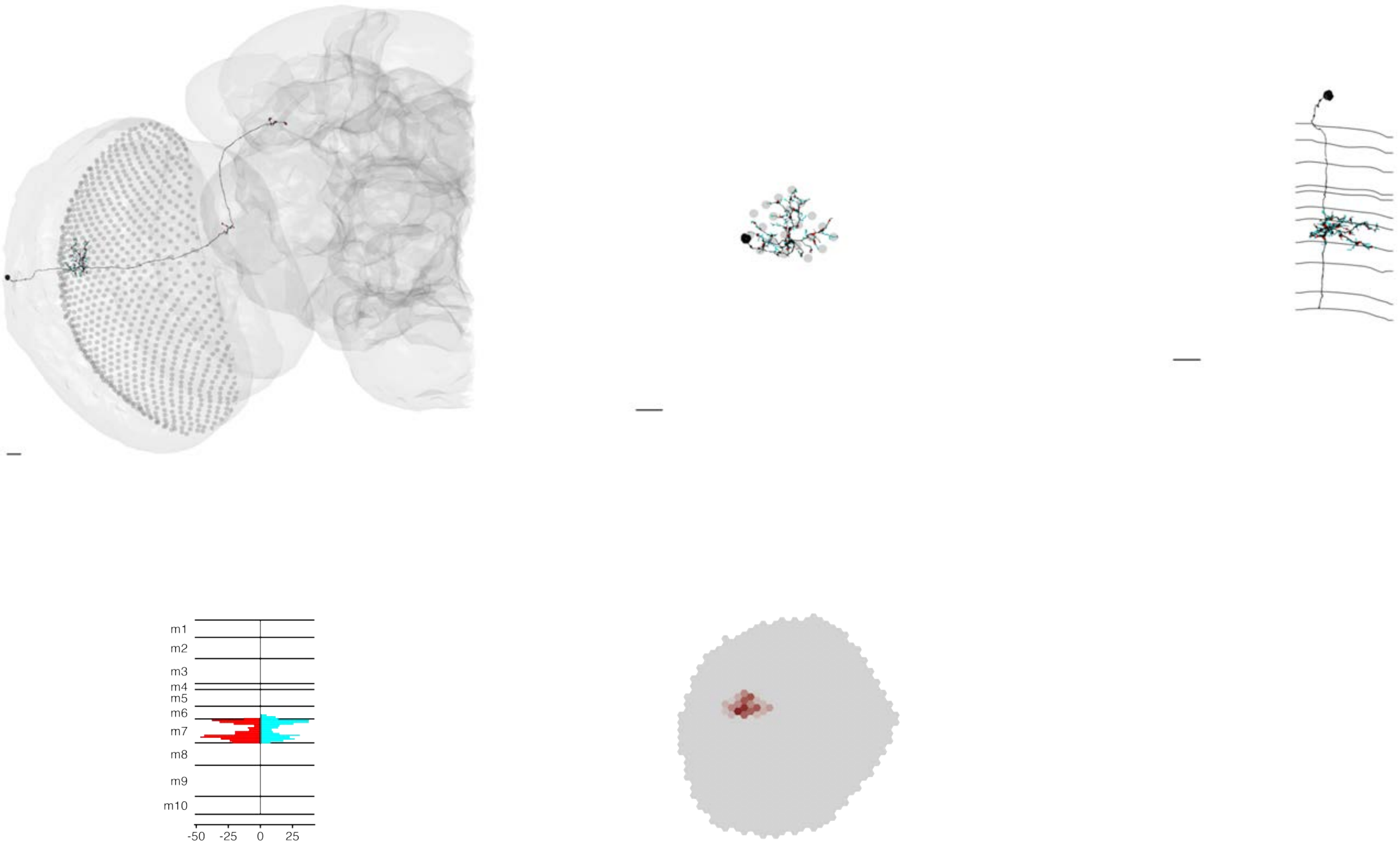

### MeTu4a_720575940632293262

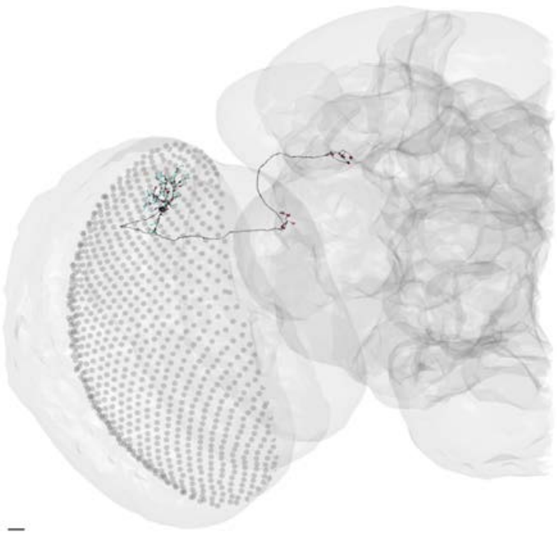

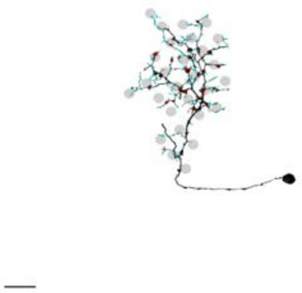

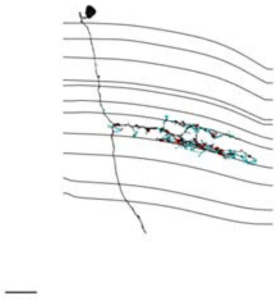

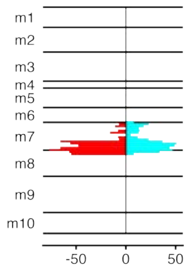


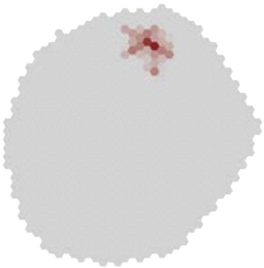

### MeTu4a_720575940632669666

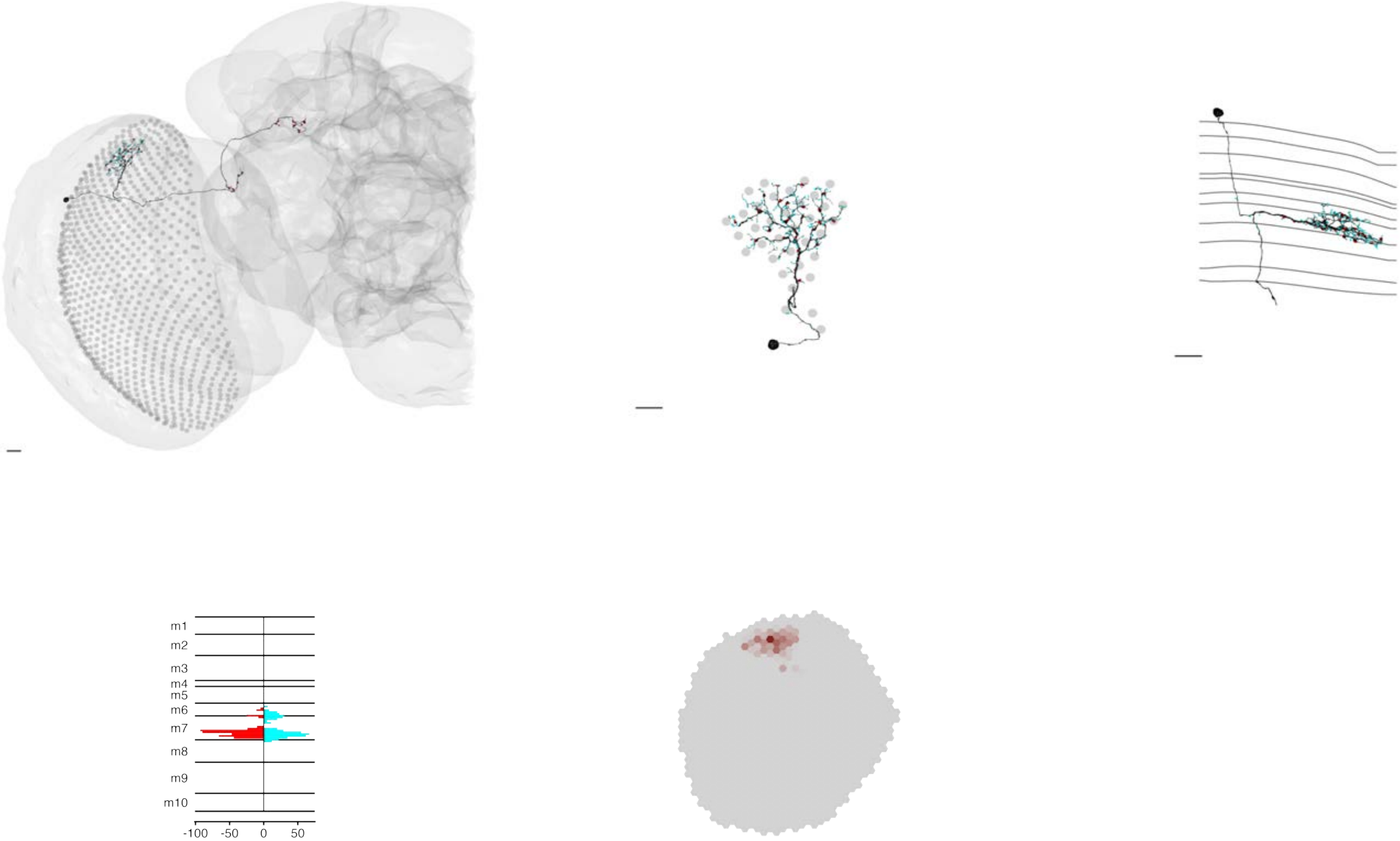

### MeTu4a_720575940632798177

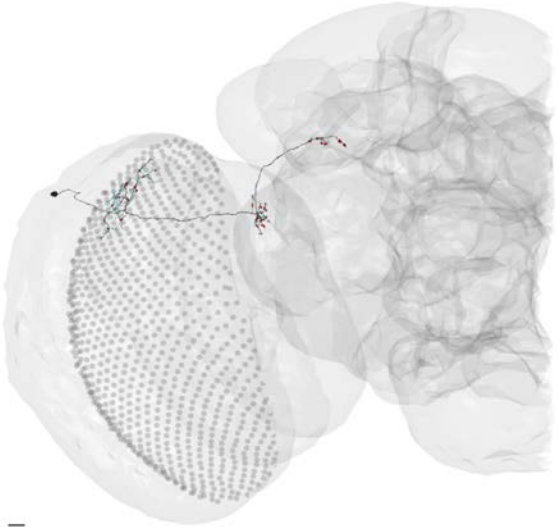

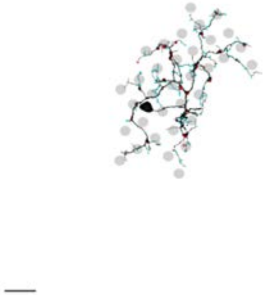

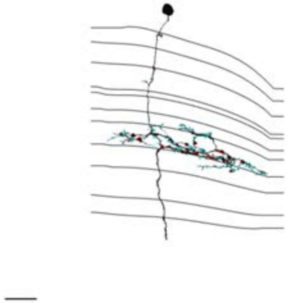

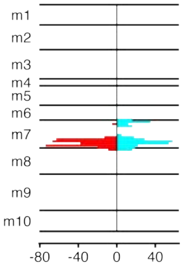


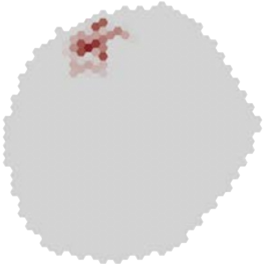

### MeTu4a_720575940632952986

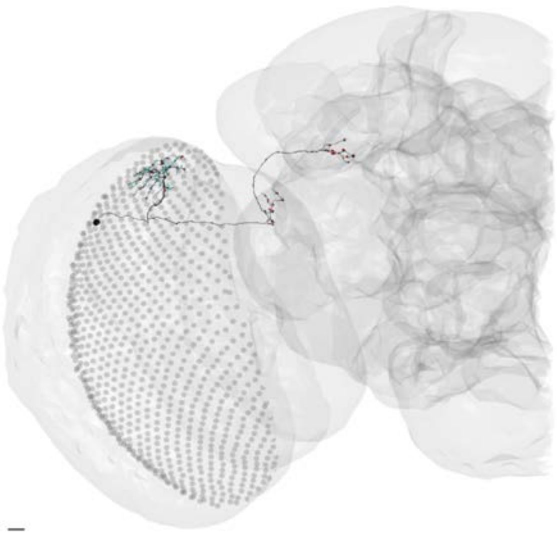

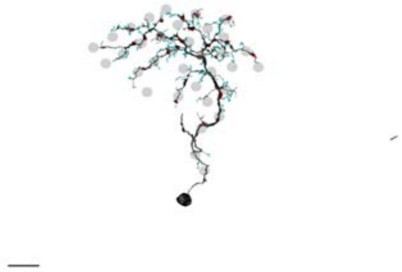

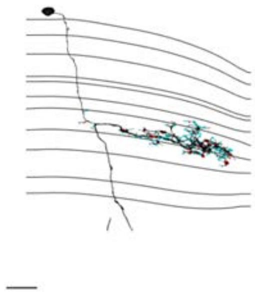

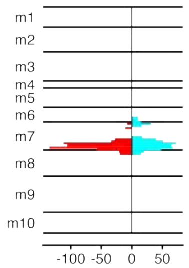


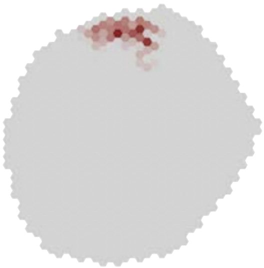

### MeTu4a_720575940633369939

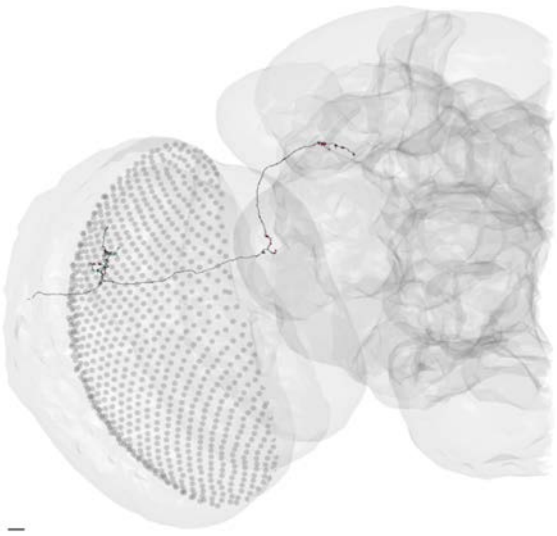

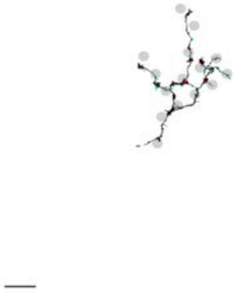

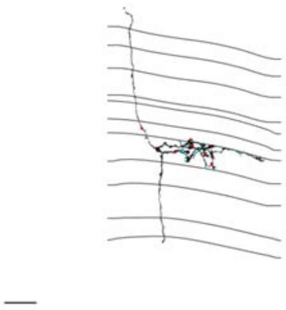

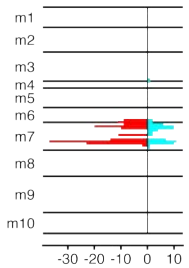


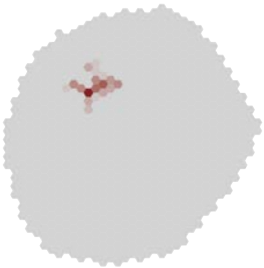

### MeTu4a_720575940633929908

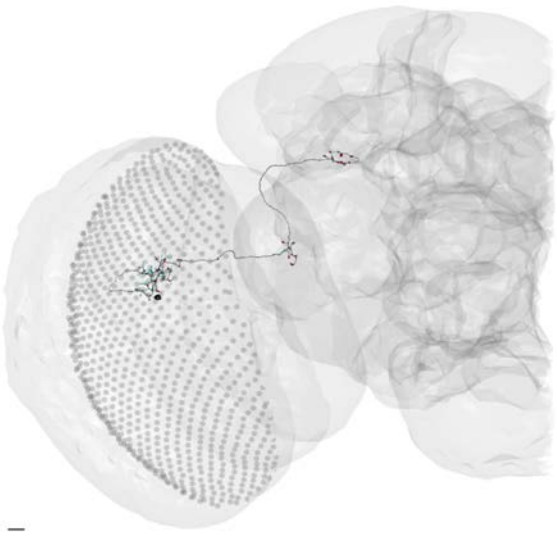

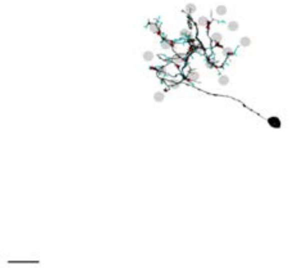

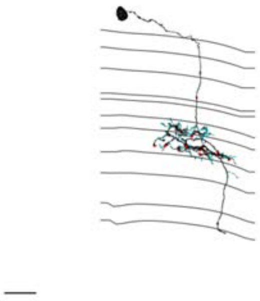

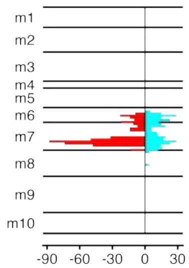


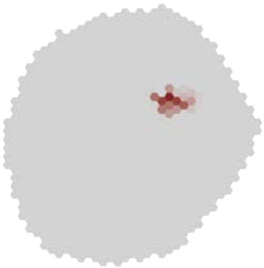

### MeTu4a_720575940639904565

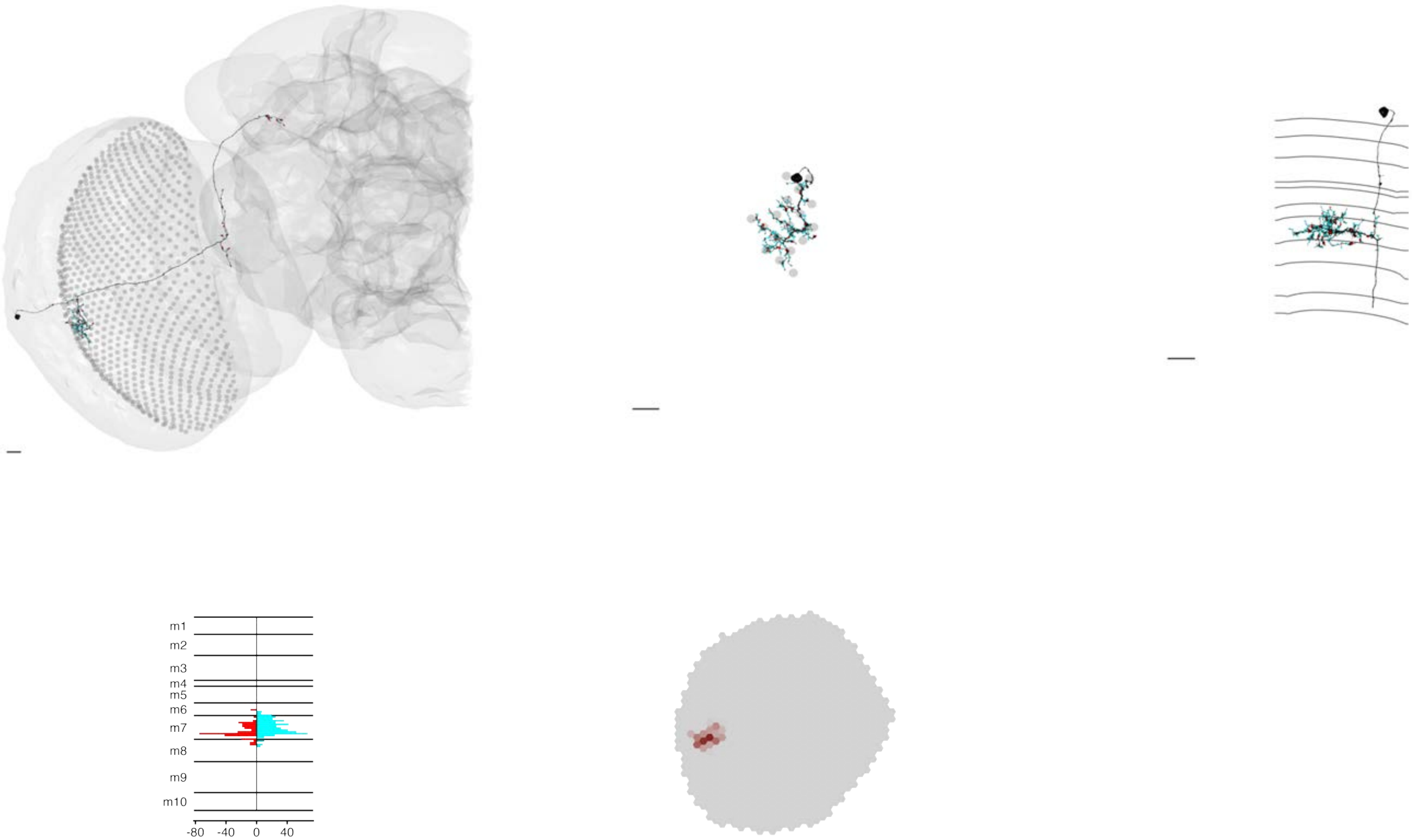

### MeTu4a_720575940643890888

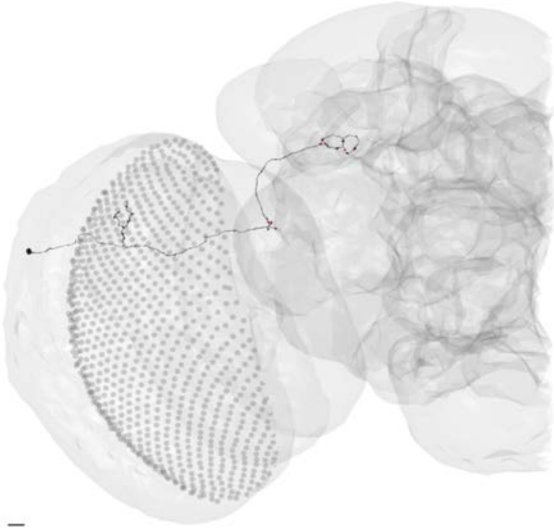

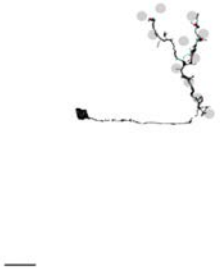

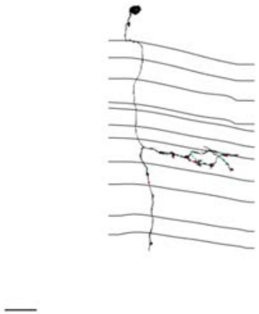

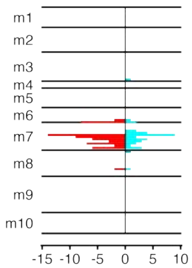


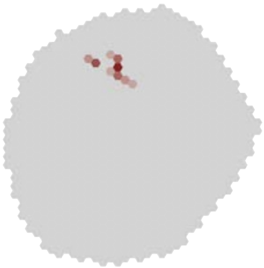

MeTu4a_720575940648817017

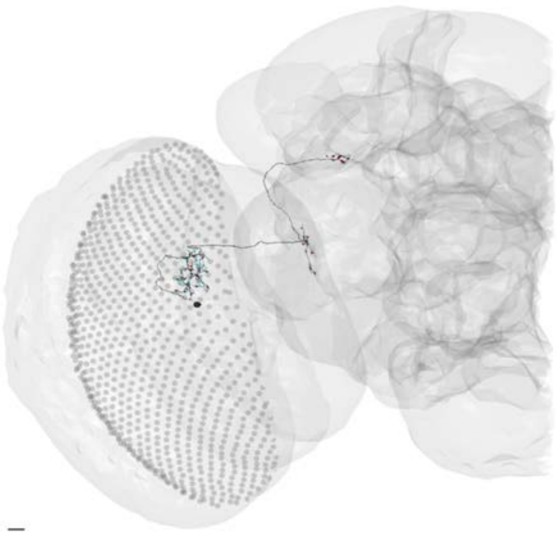

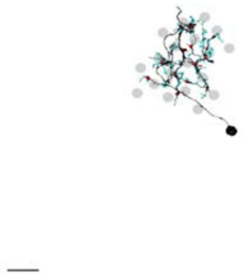

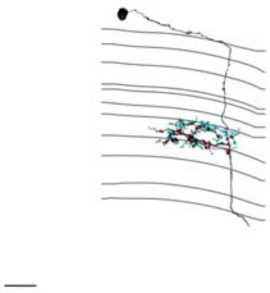

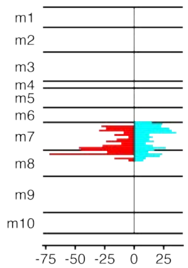


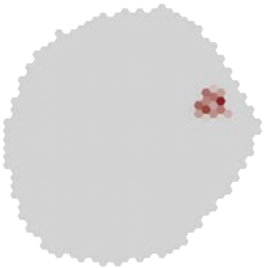

### MeTu4a_720575940650581625

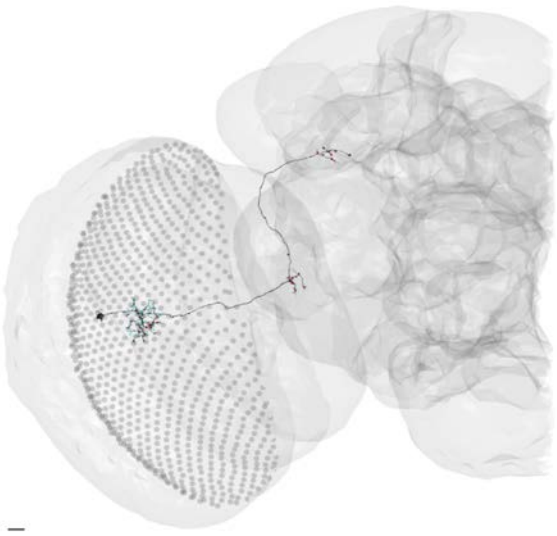

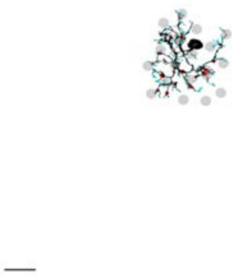

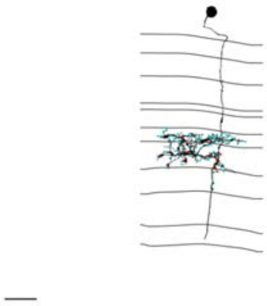

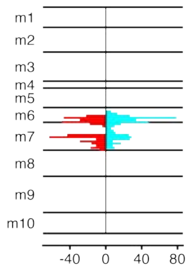


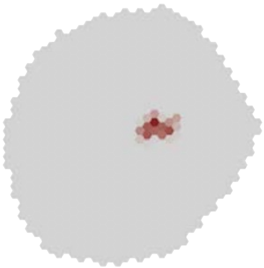

### MeTu4b_720575940609956547

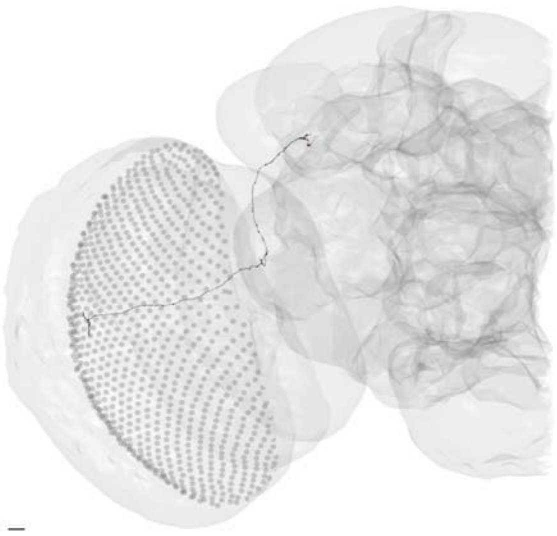

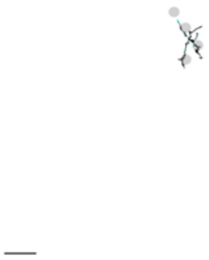

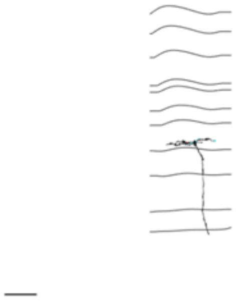

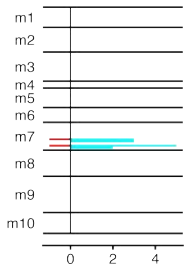


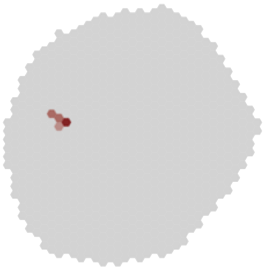

### MeTu4b_720575940610350961

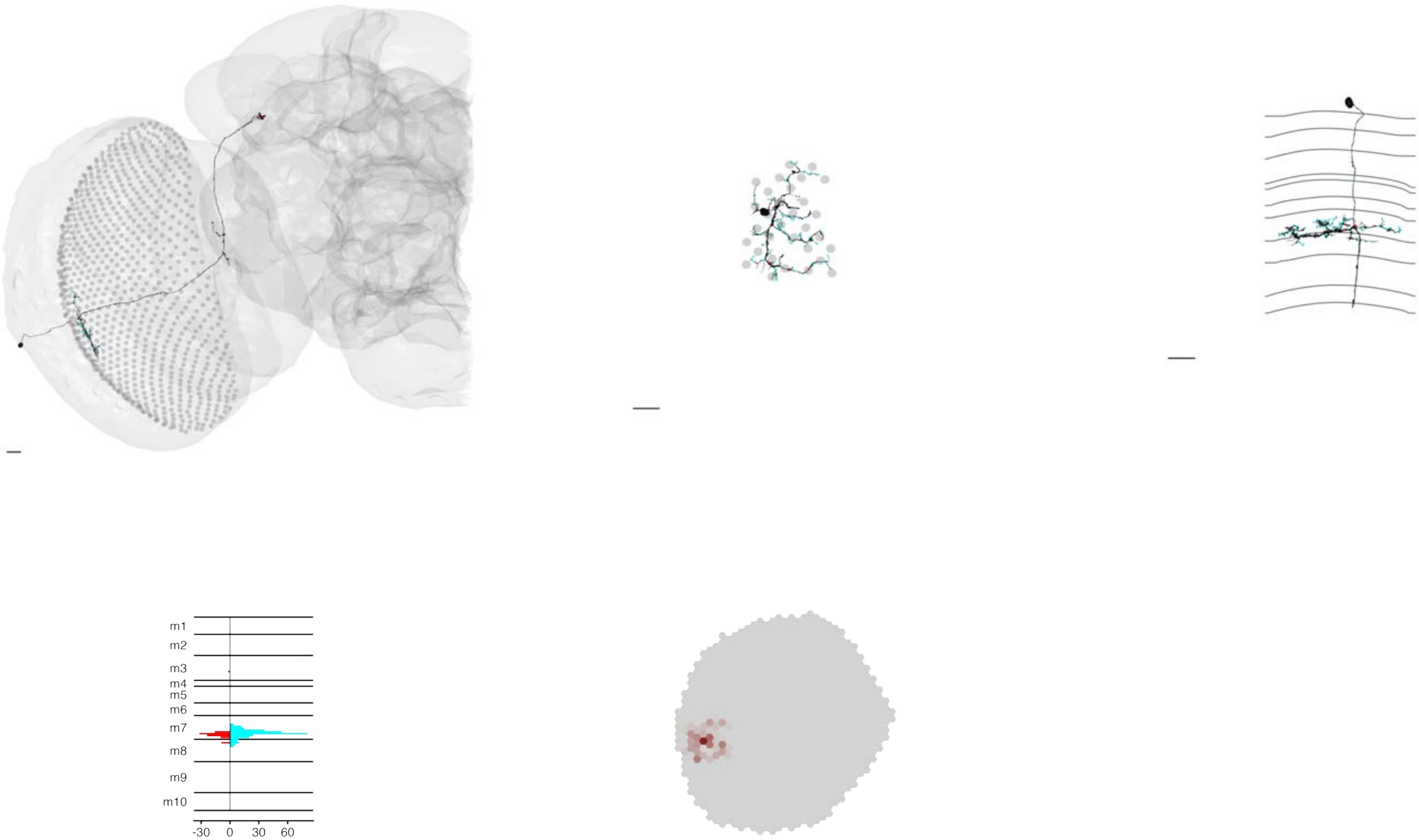

### MeTu4b_720575940613735934

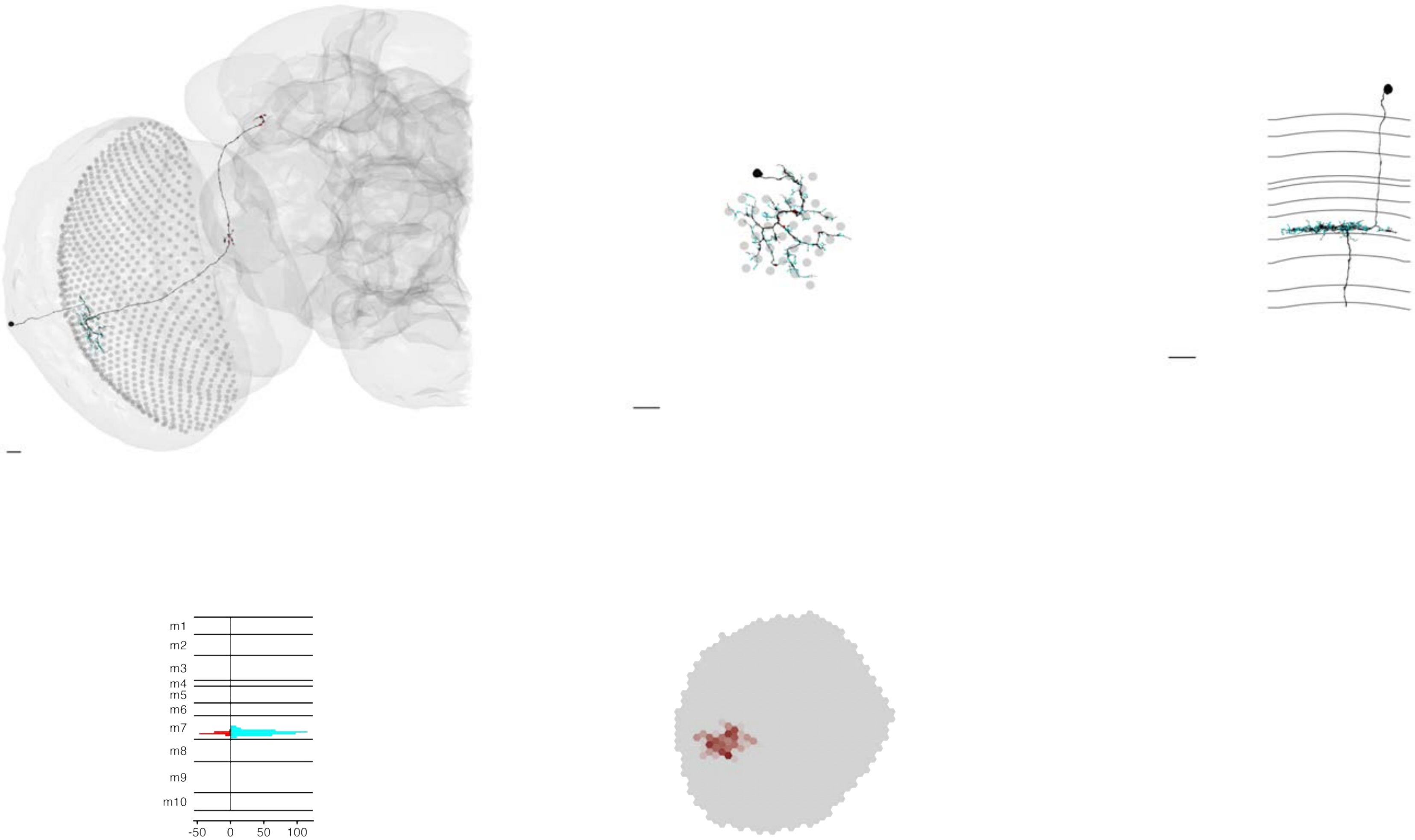

### MeTu4b_720575940613803747

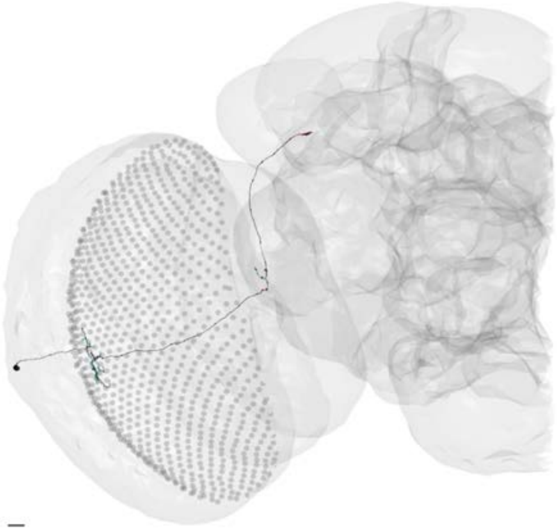

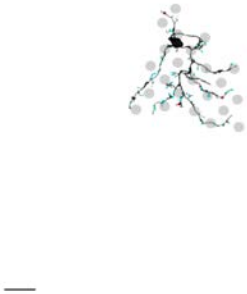

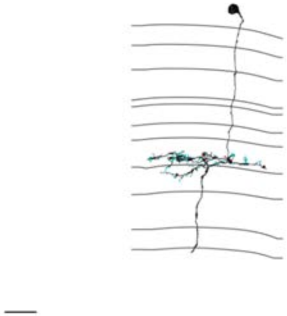

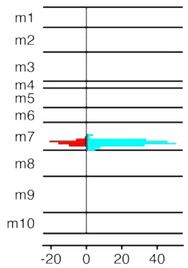


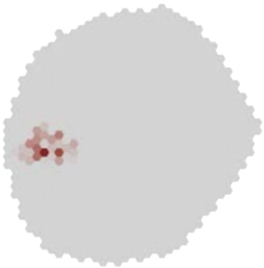

### MeTu4b_720575940617195934

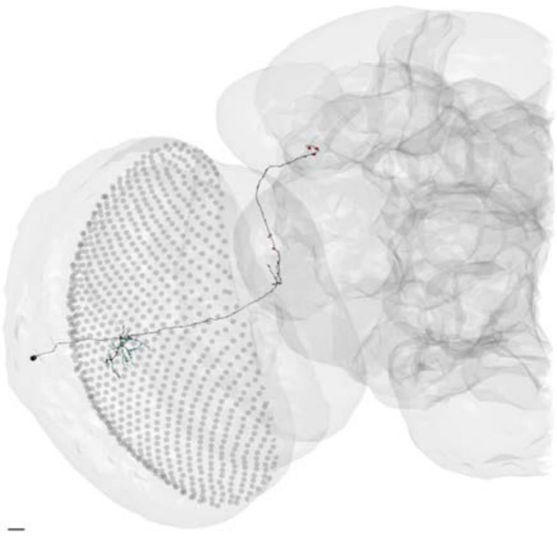

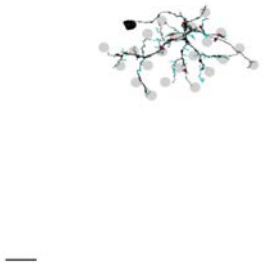

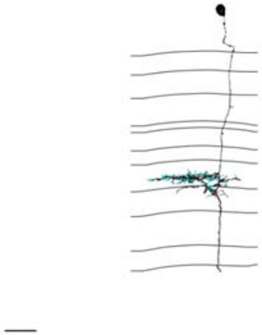

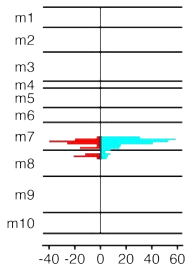


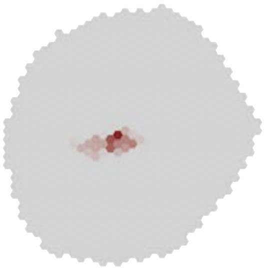

### MeTu4b_720575940618751508

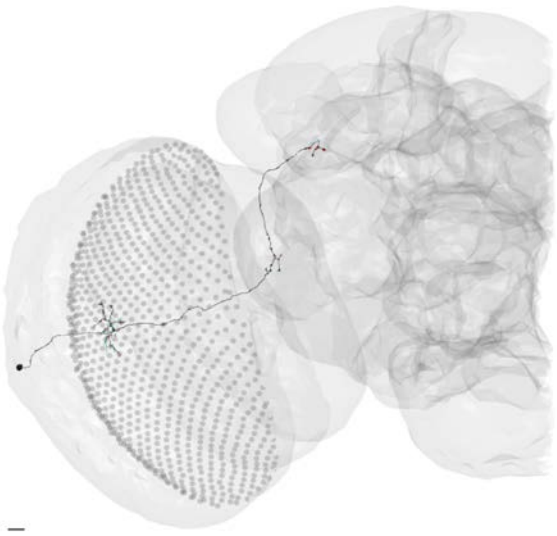

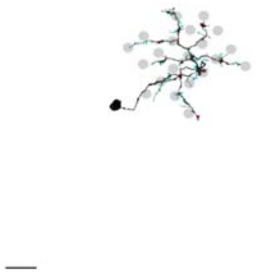

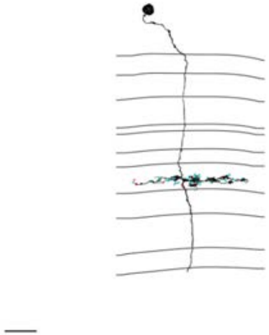

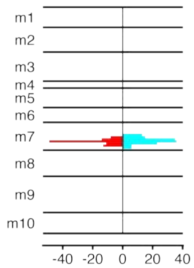


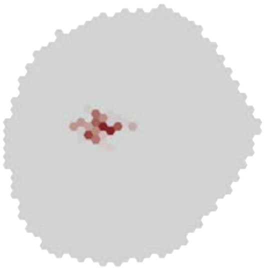

MeTu4b_720575940620151873

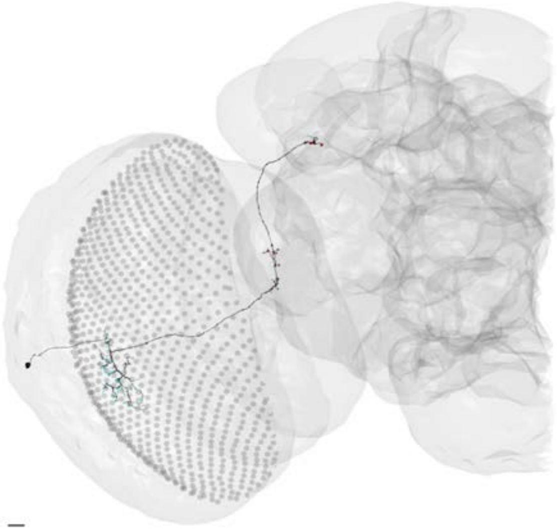

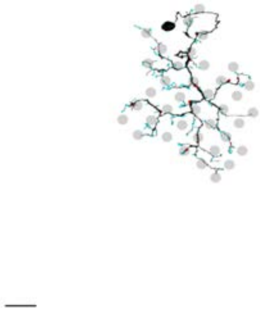

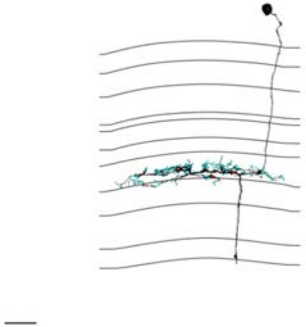

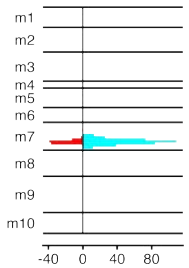


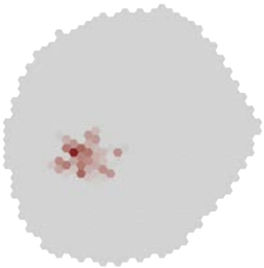

### MeTu4b_720575940620465524

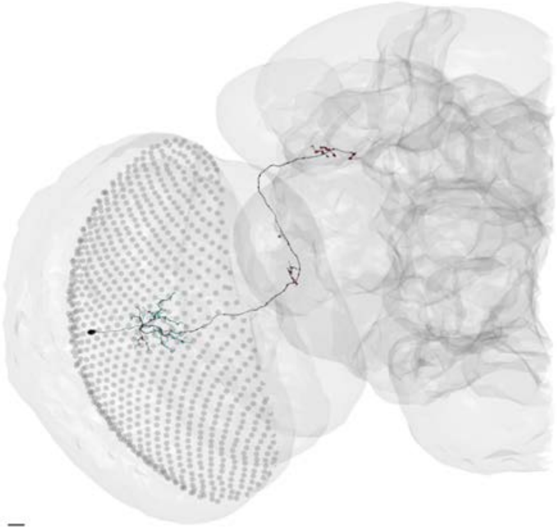

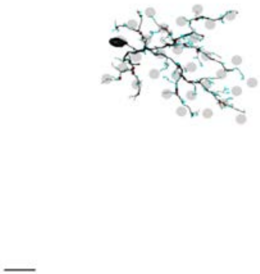

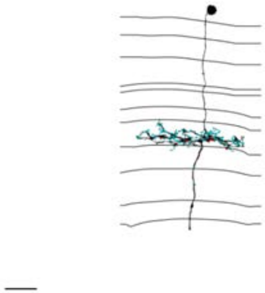

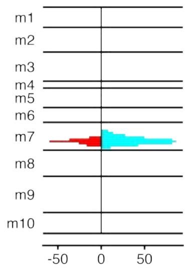


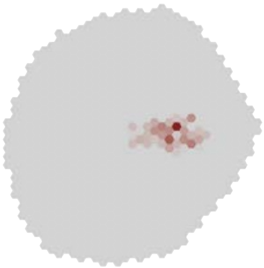

### MeTu4b_720575940621165261

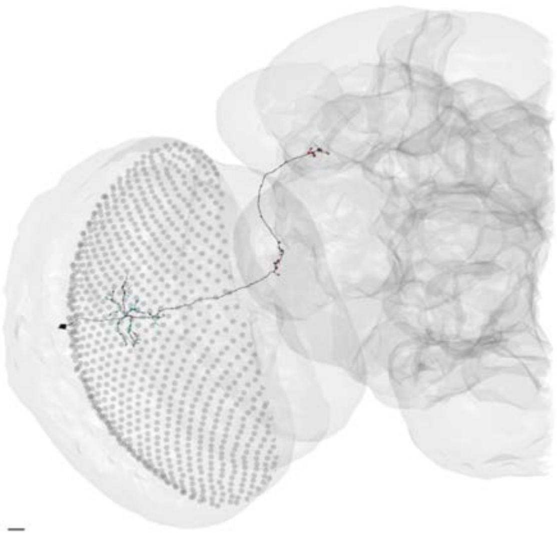

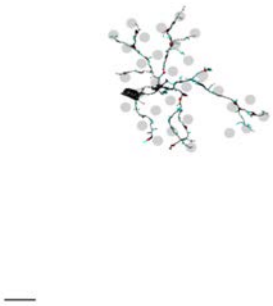

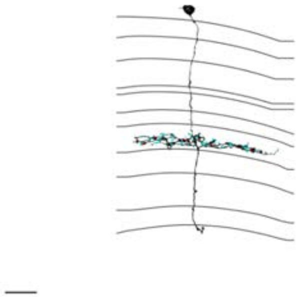

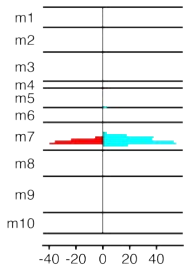


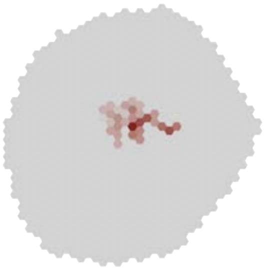

### MeTu4b_720575940624529229

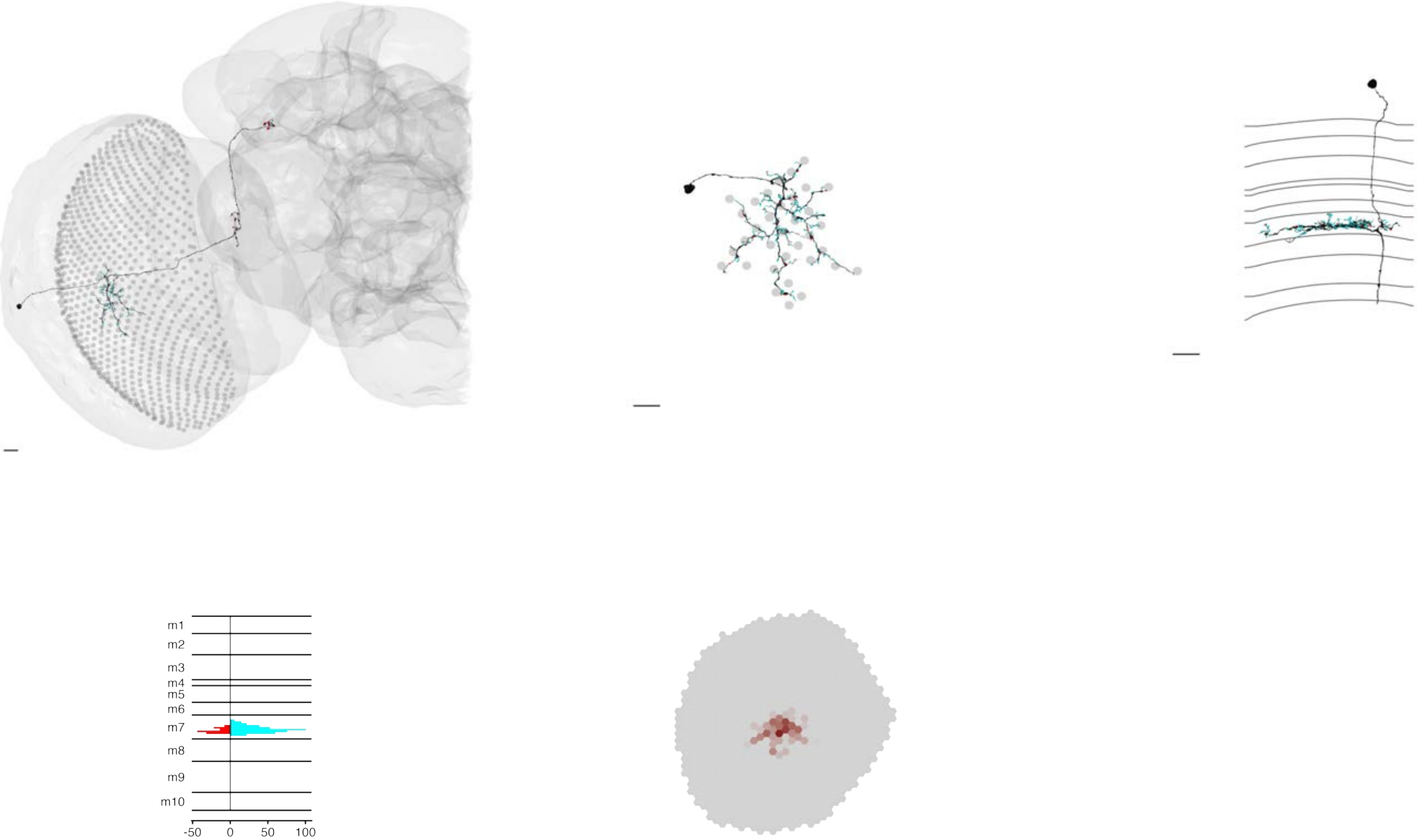

### MeTu4b_720575940625227160

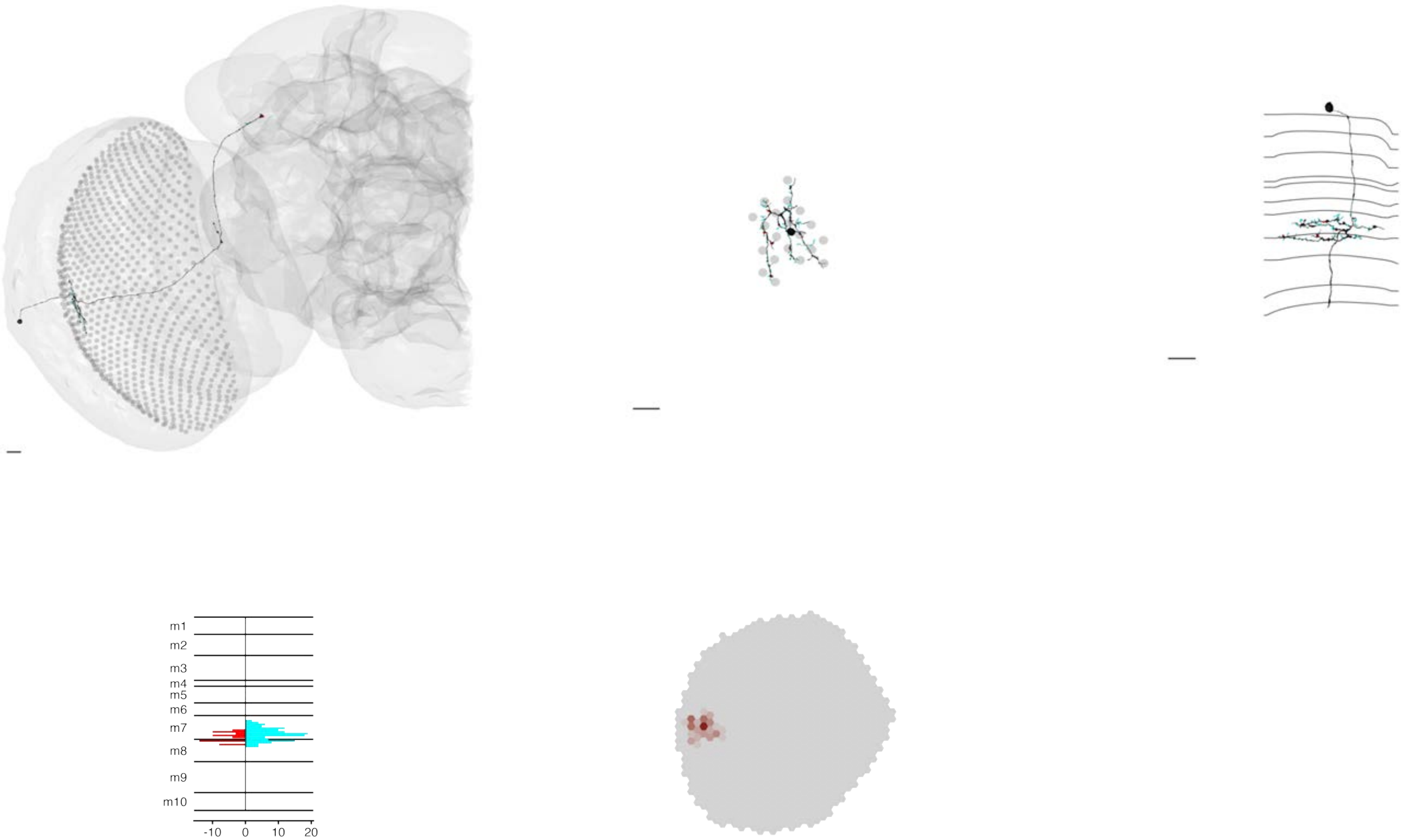

### MeTu4b_720575940626194718

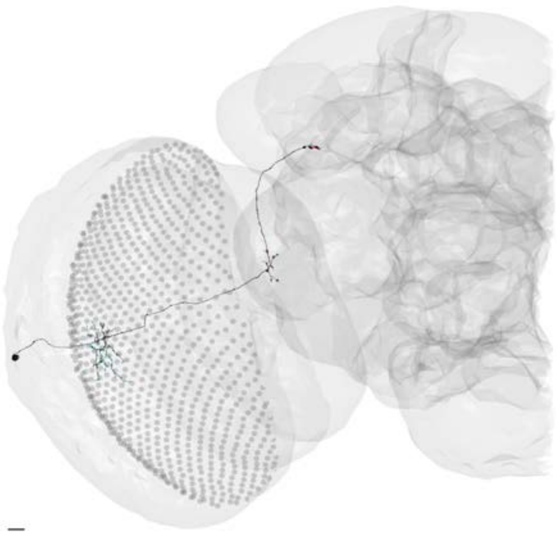

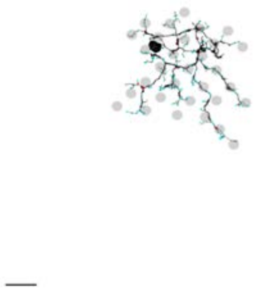

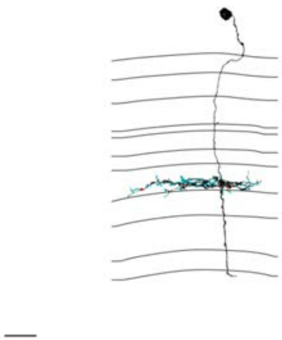

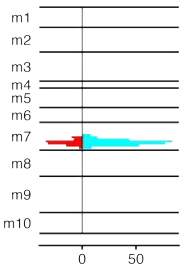


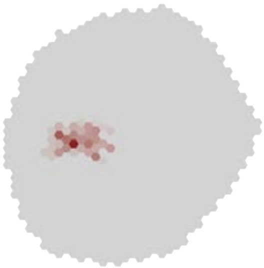

### MeTu4c_720575940604007392

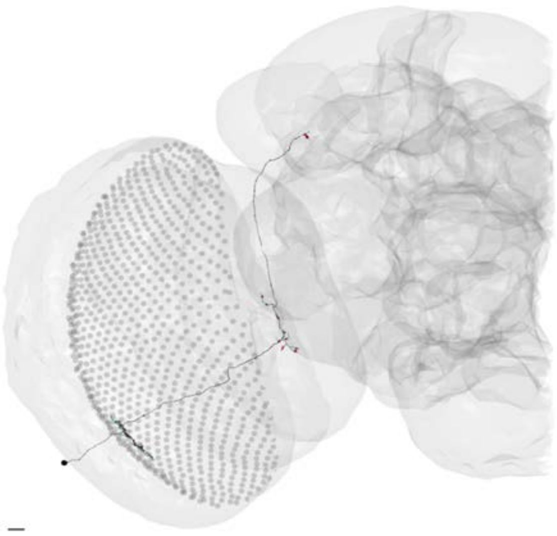

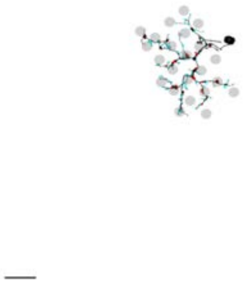

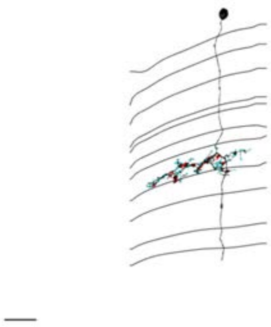

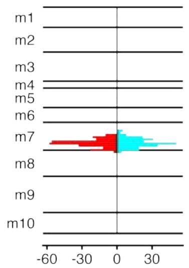


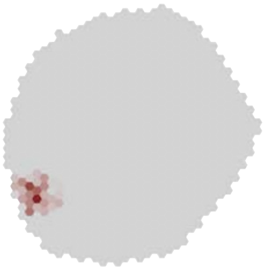

### MeTu4c_720575940605791422

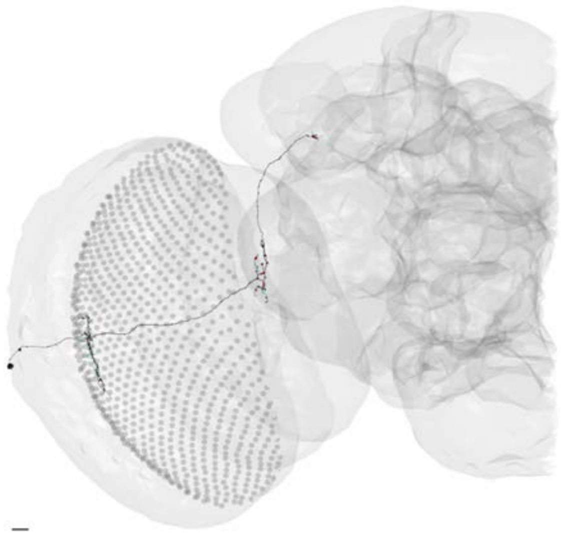

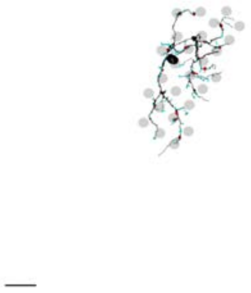

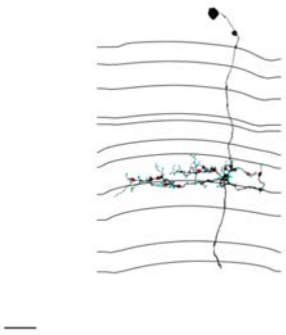

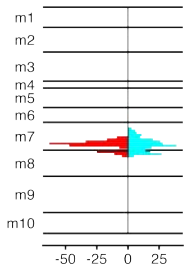


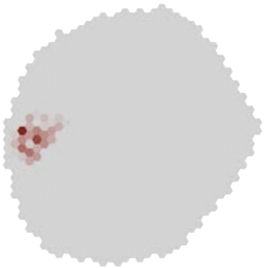

### MeTu4c_720575940607603548

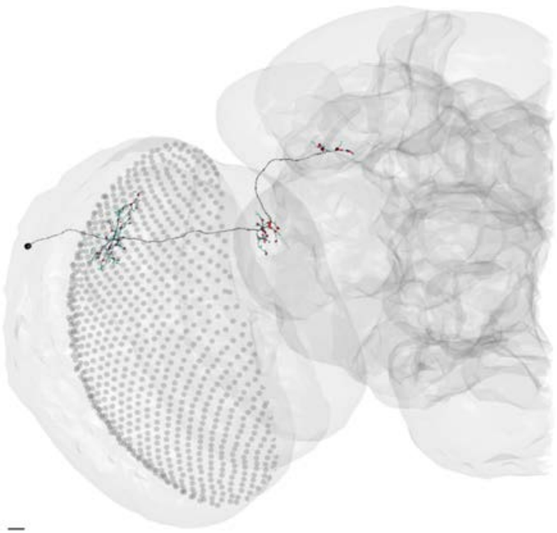

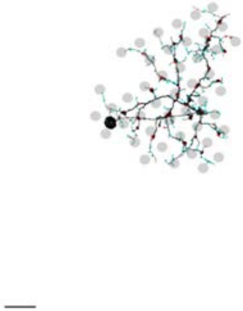

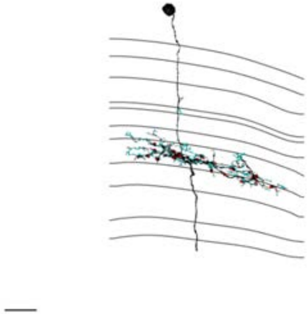

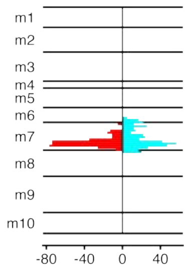


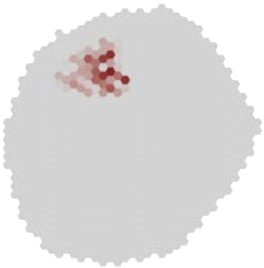

MeTu4c_720575940607671241

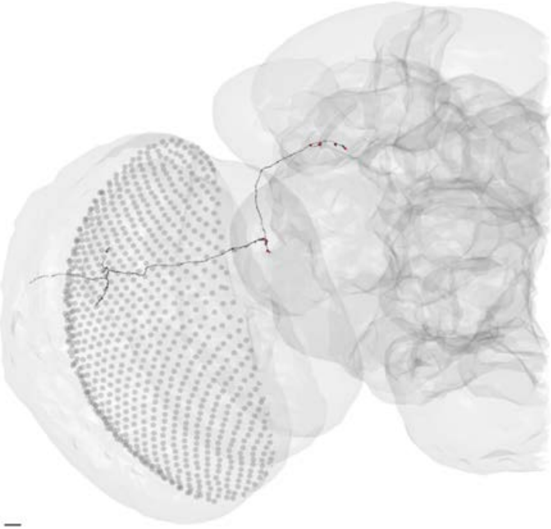

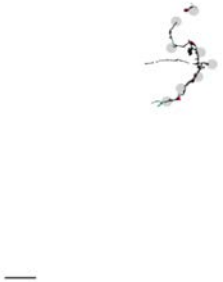

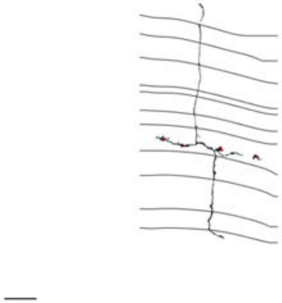

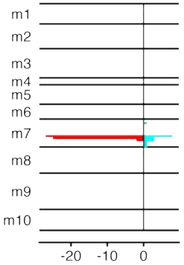


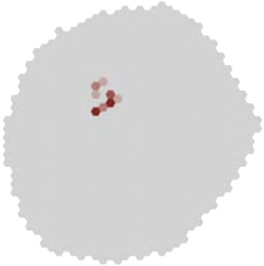

### MeTu4c_720575940616021899

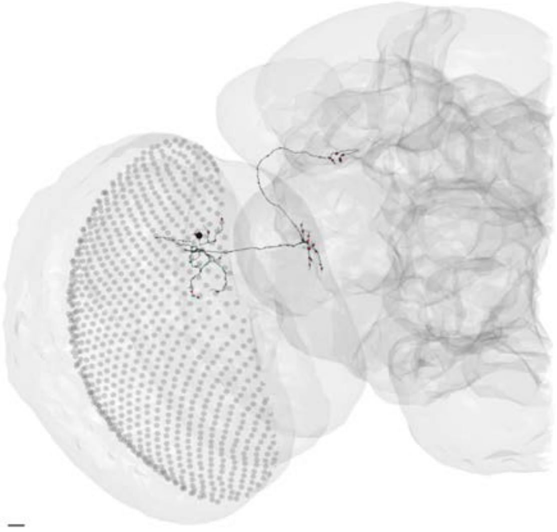

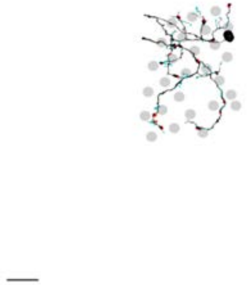

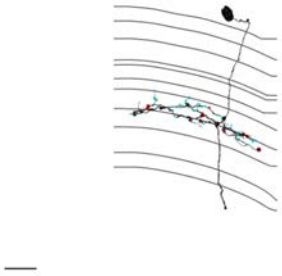

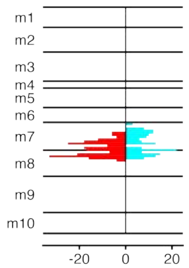


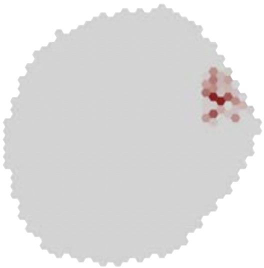

### MeTu4c_720575940616445638

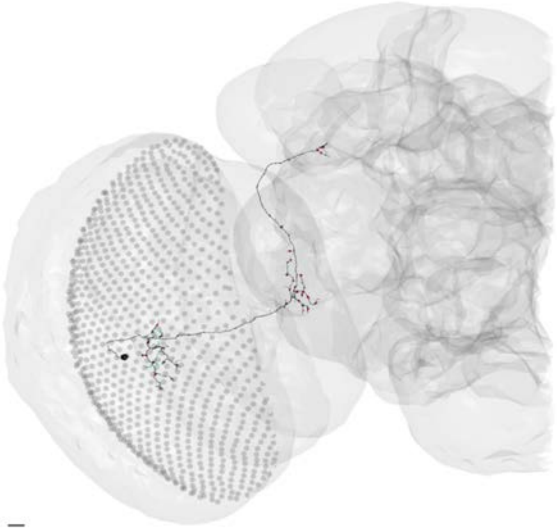

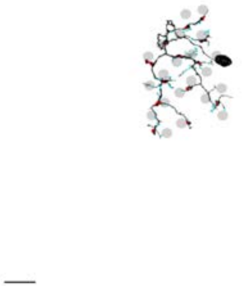

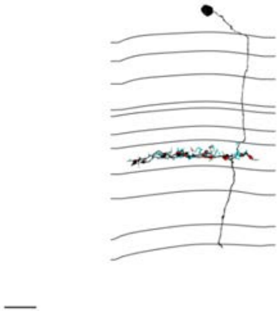

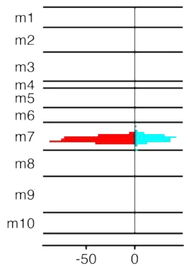


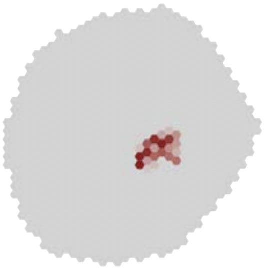

### MeTu4c_720575940617979682

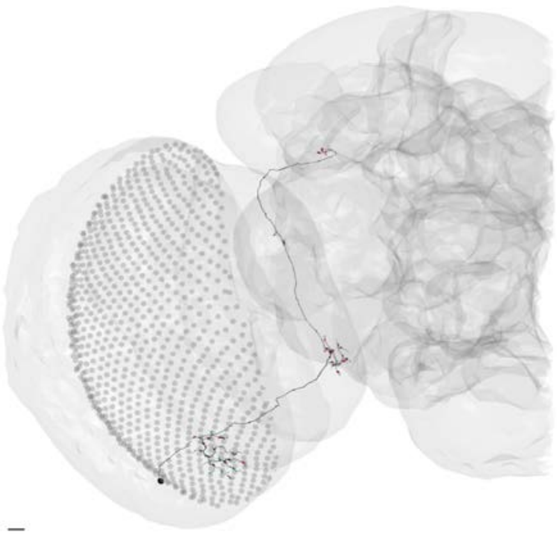

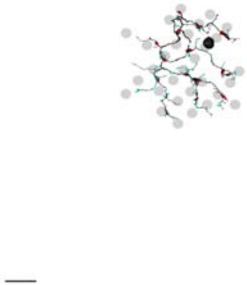

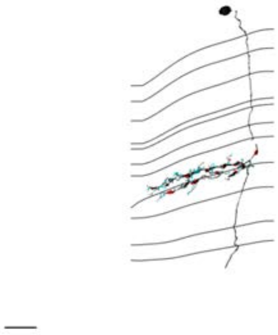

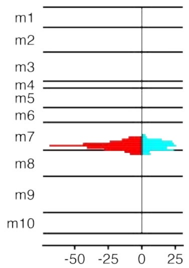


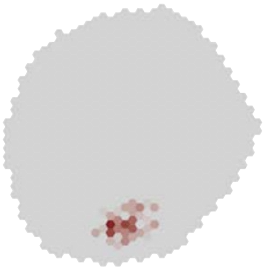

### MeTu4c_720575940618758592

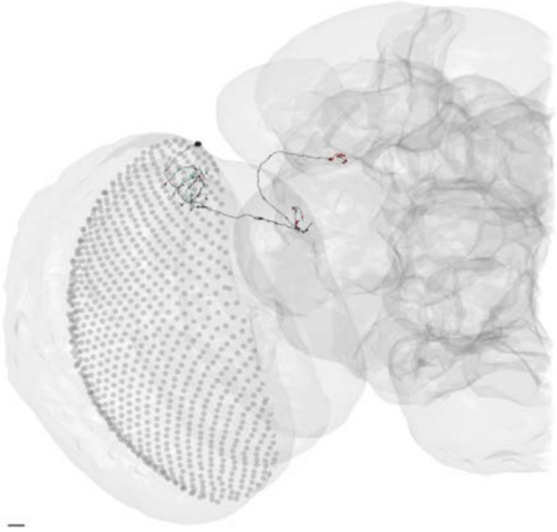

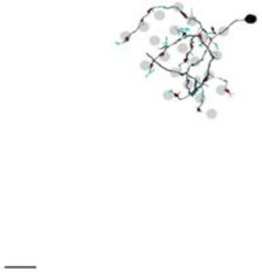

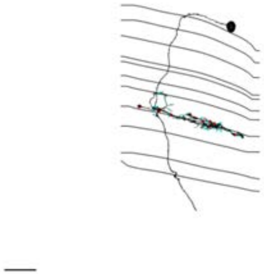

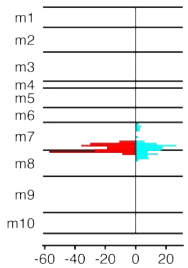


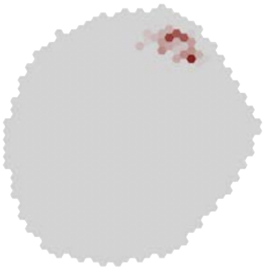

### MeTu4c_720575940618964672

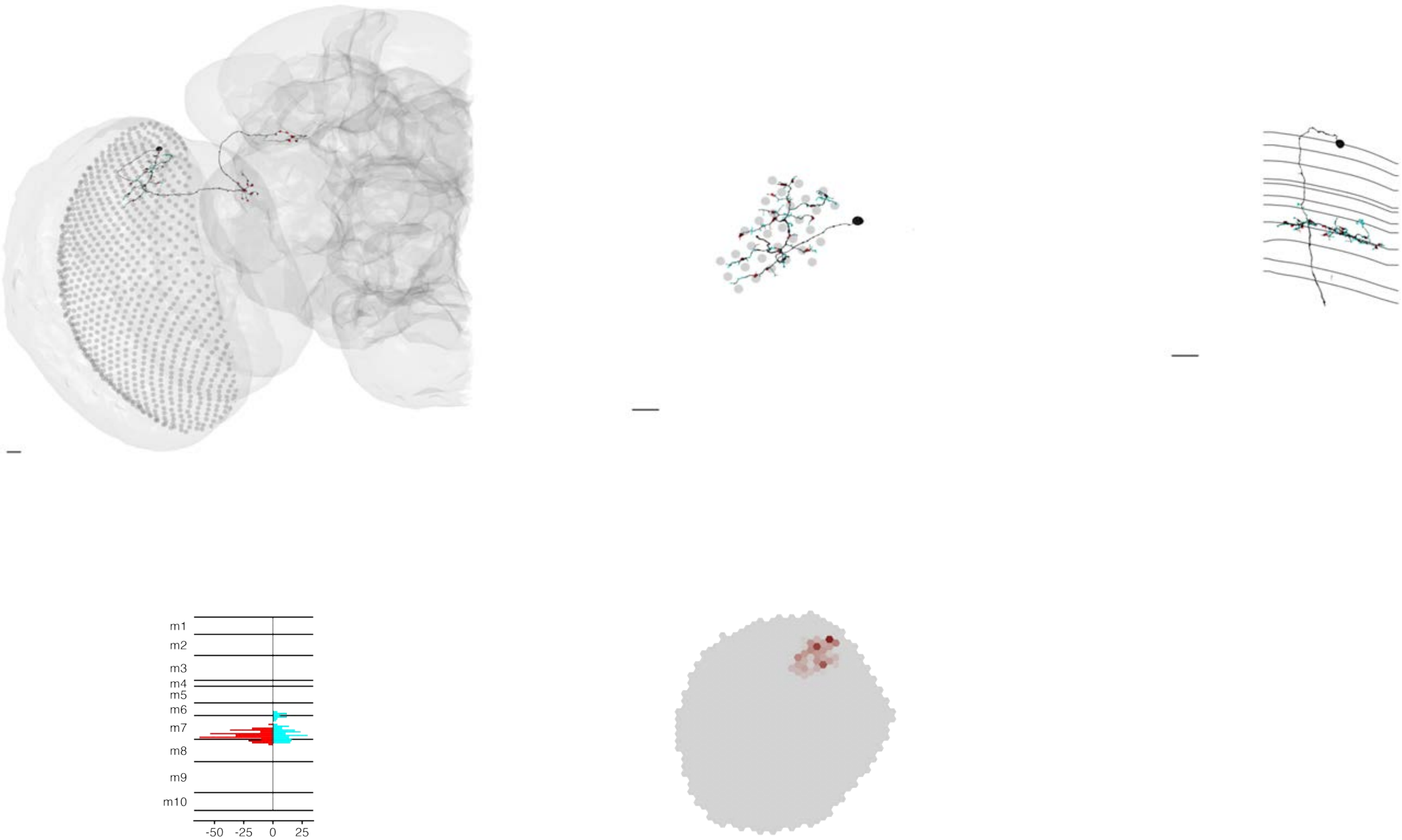

### MeTu4c_720575940619072592

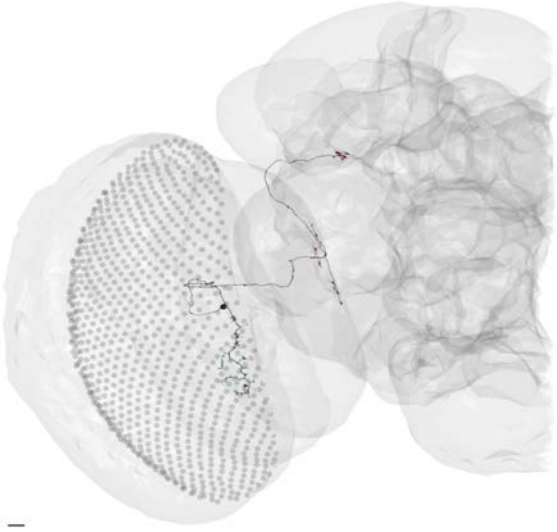

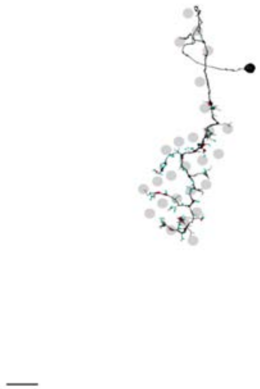

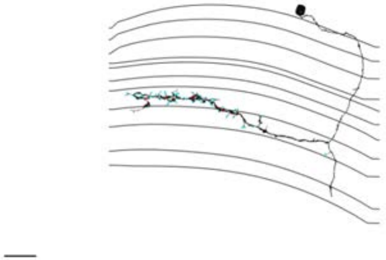

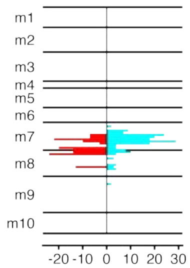


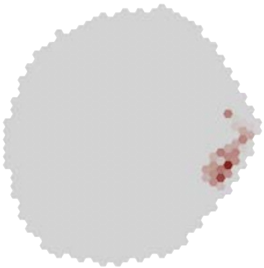

### MeTu4c_720575940619442113

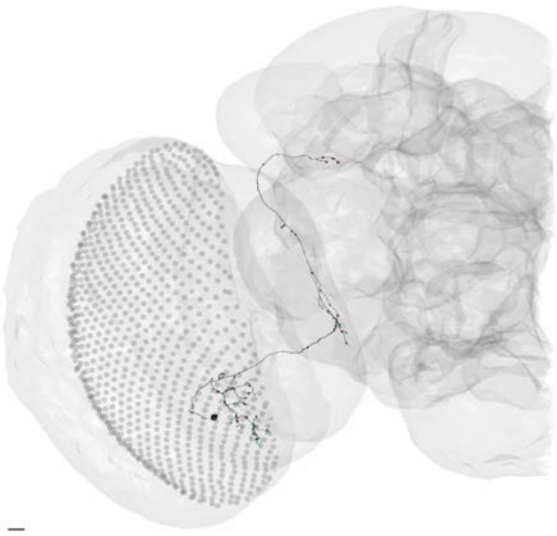

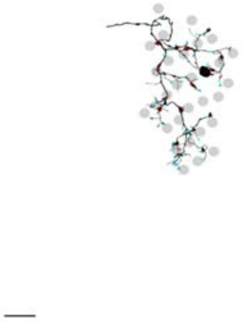

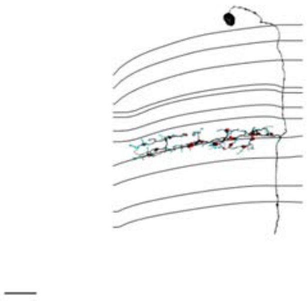

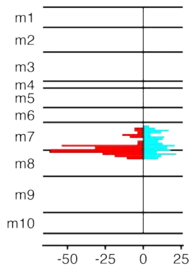


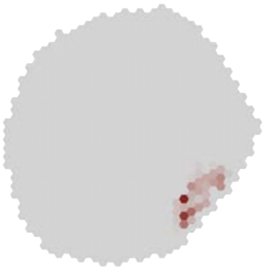

### MeTu4c_720575940619742075

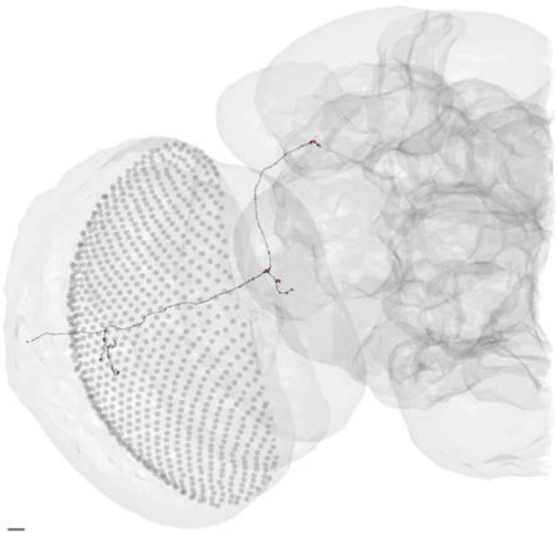

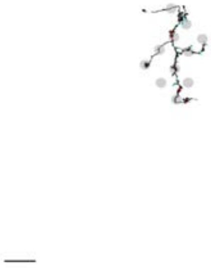

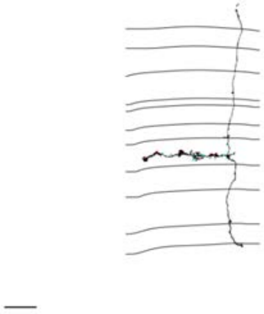

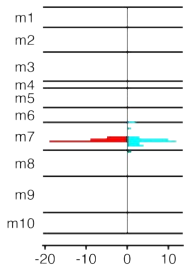


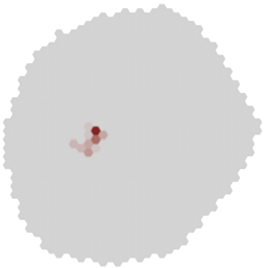

MeTu4c_720575940620422485

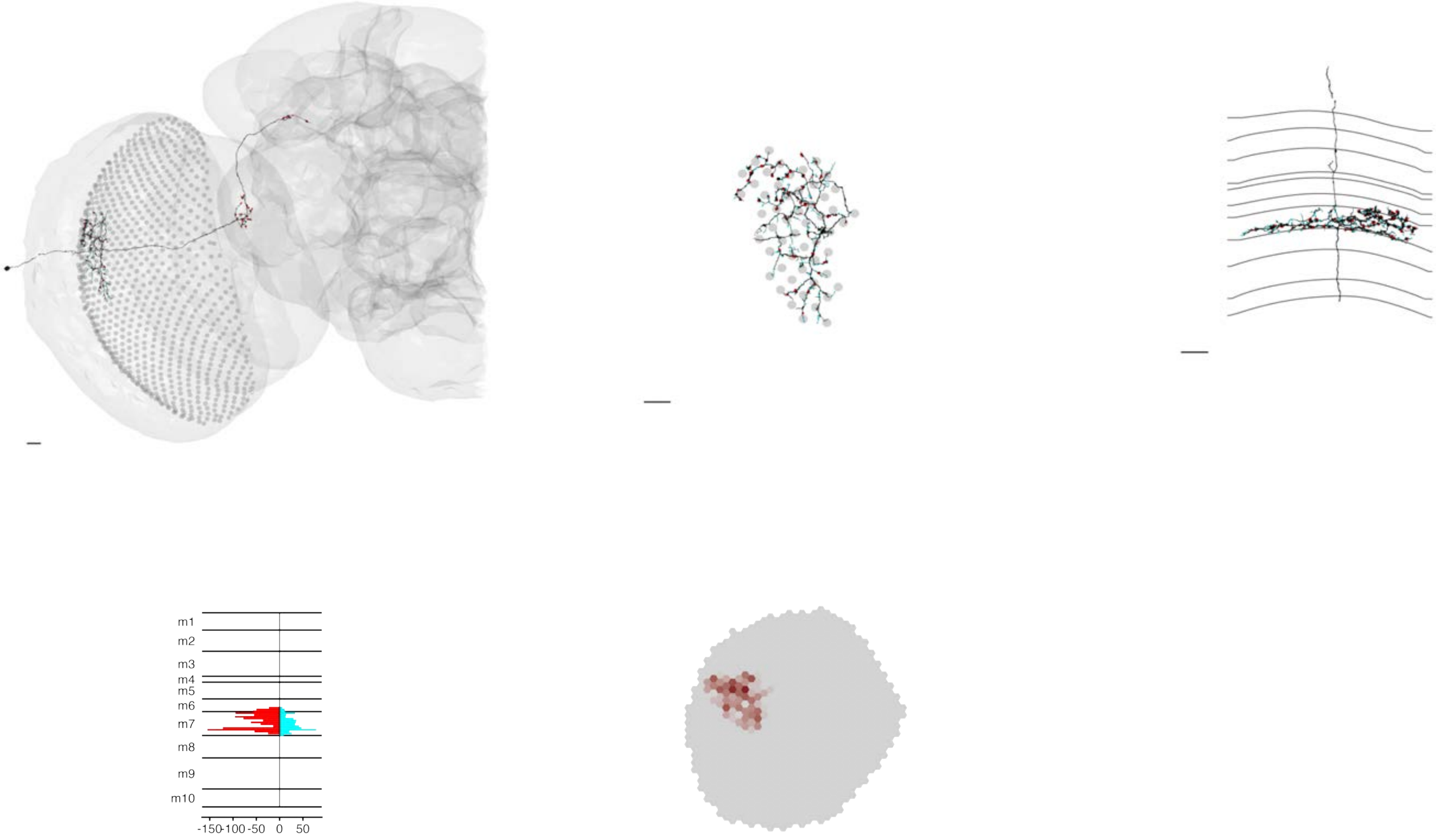

### MeTu4c_720575940620534710

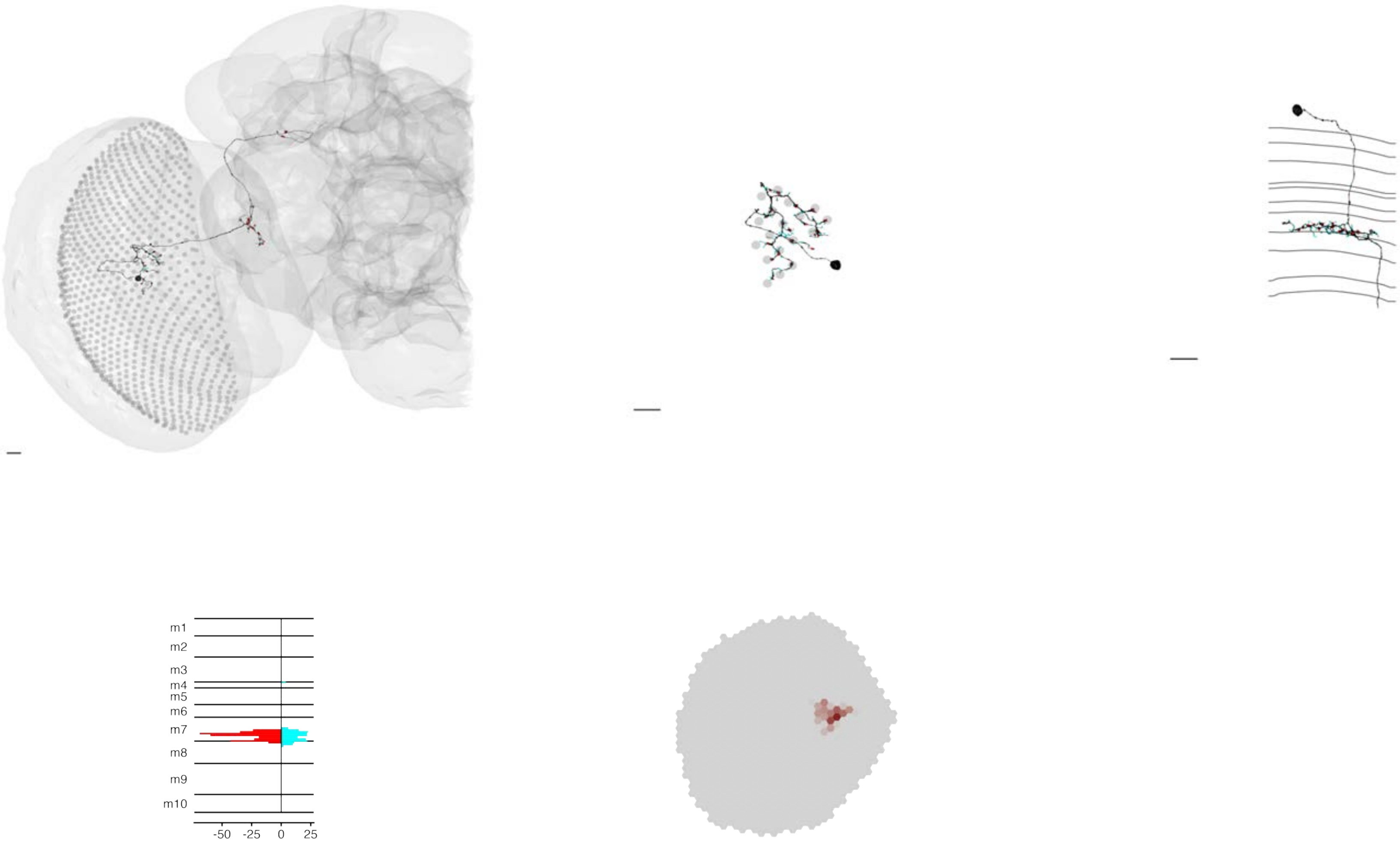

### MeTu4c_720575940620658928

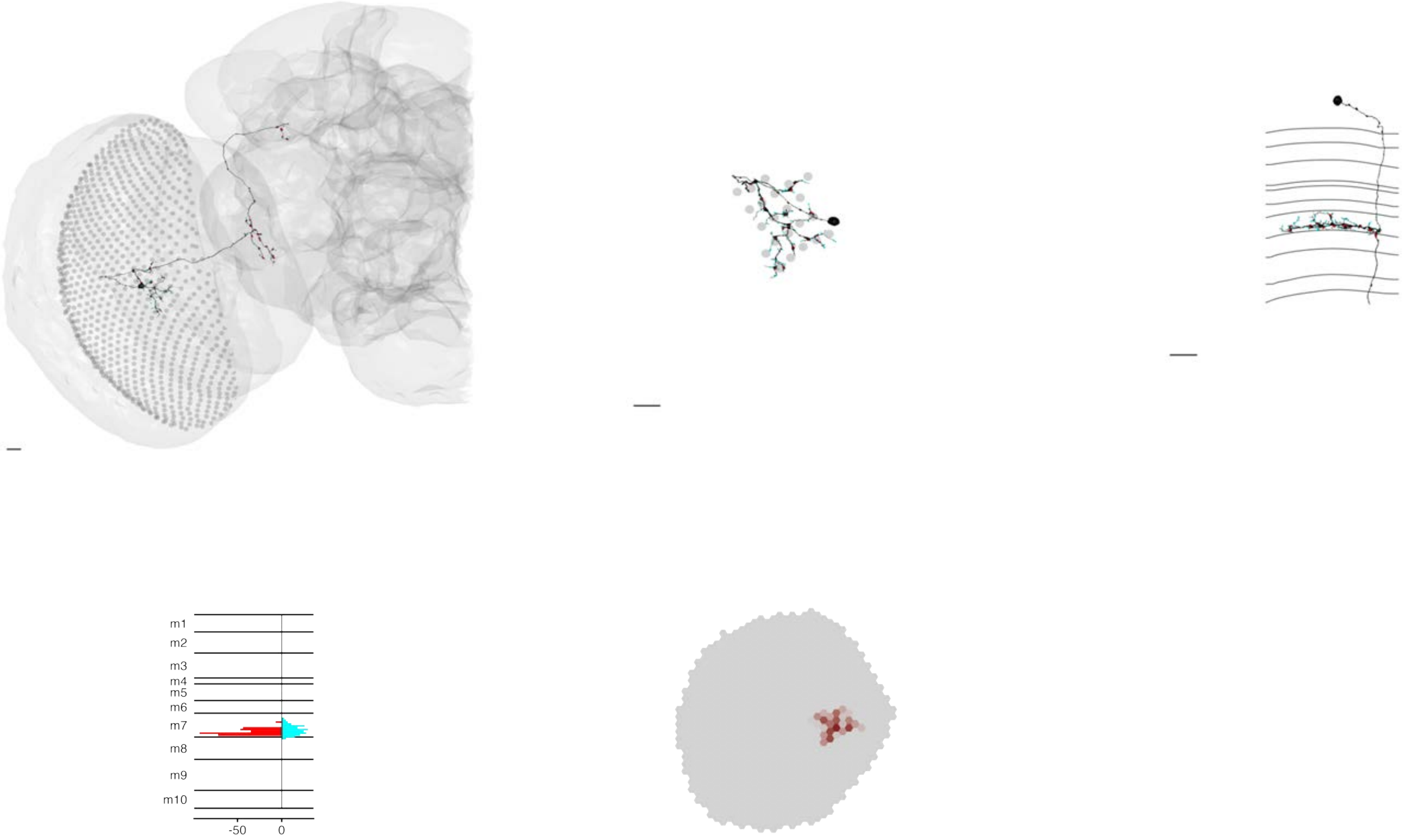

### MeTu4c_720575940620660901

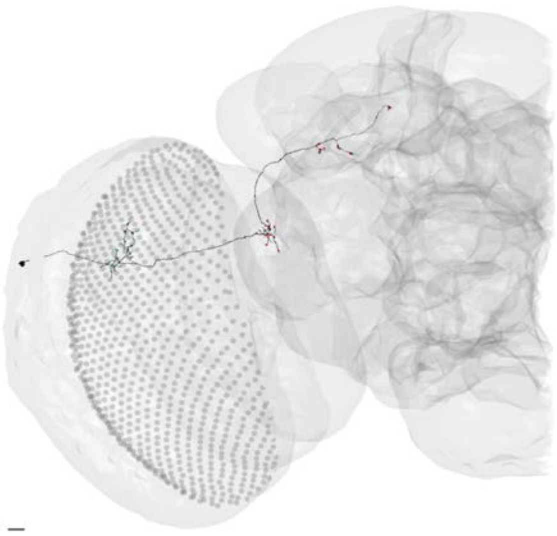

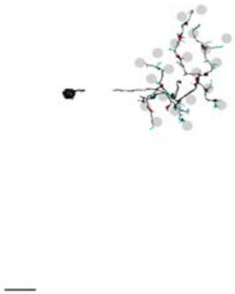

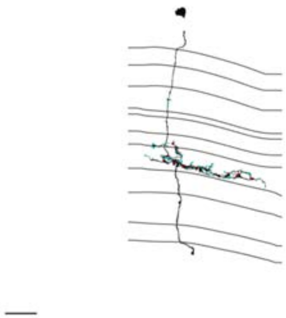

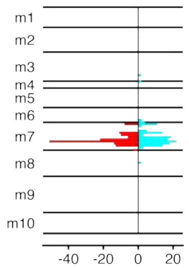


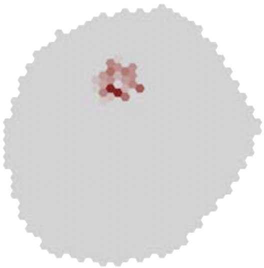

### MeTu4c_720575940620989862

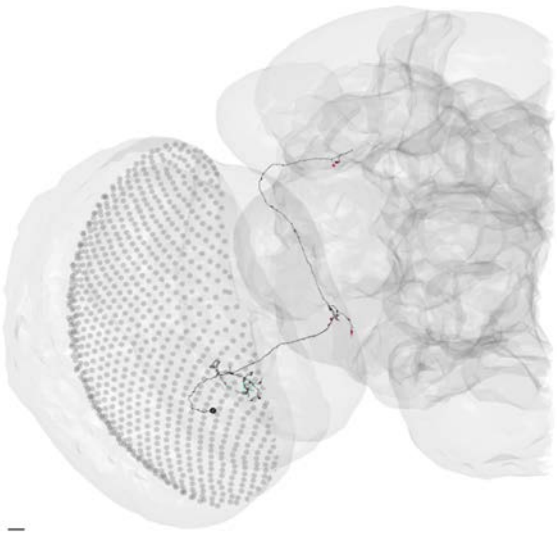

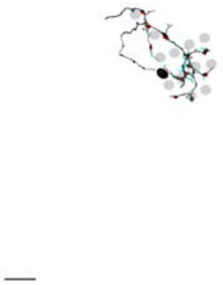

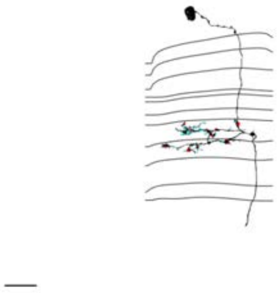

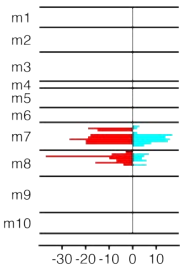


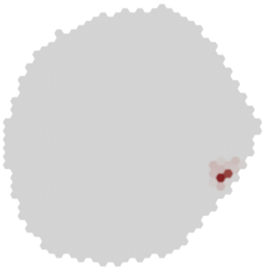

### MeTu4c_720575940621182043

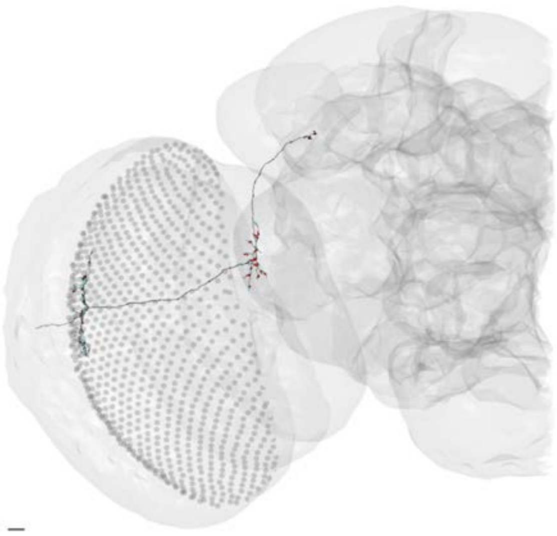

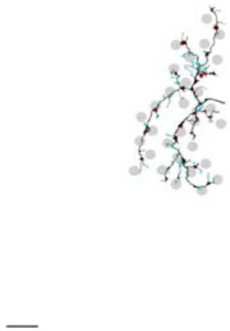

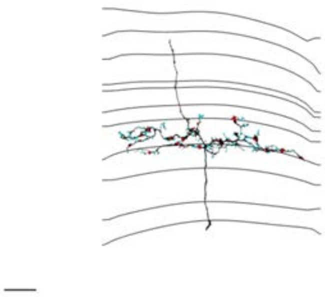

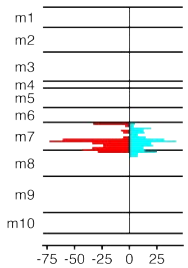


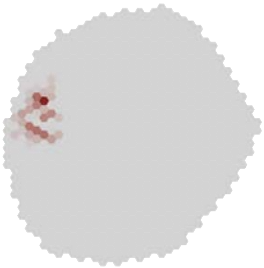

### MeTu4c_720575940621412193

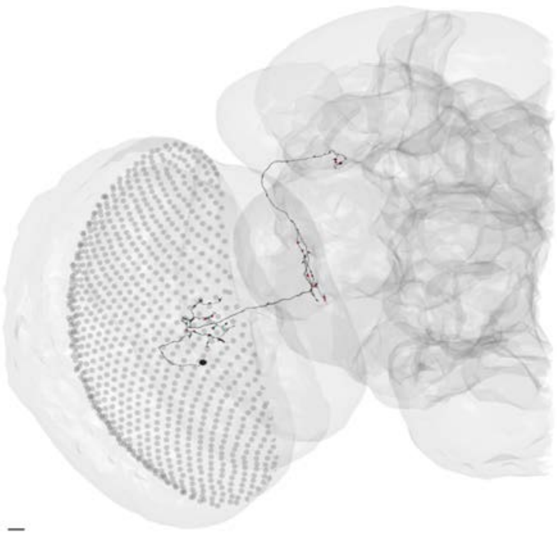

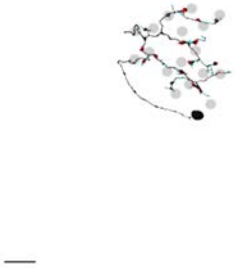

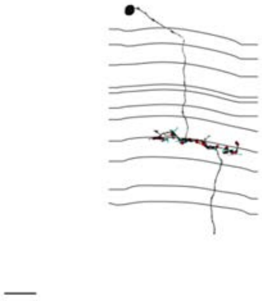

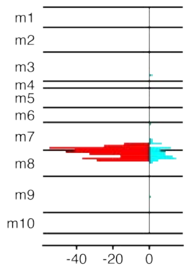


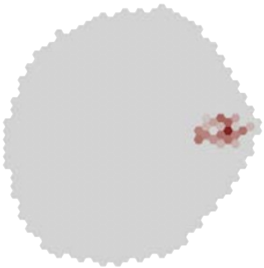

### MeTu4c_720575940622182732

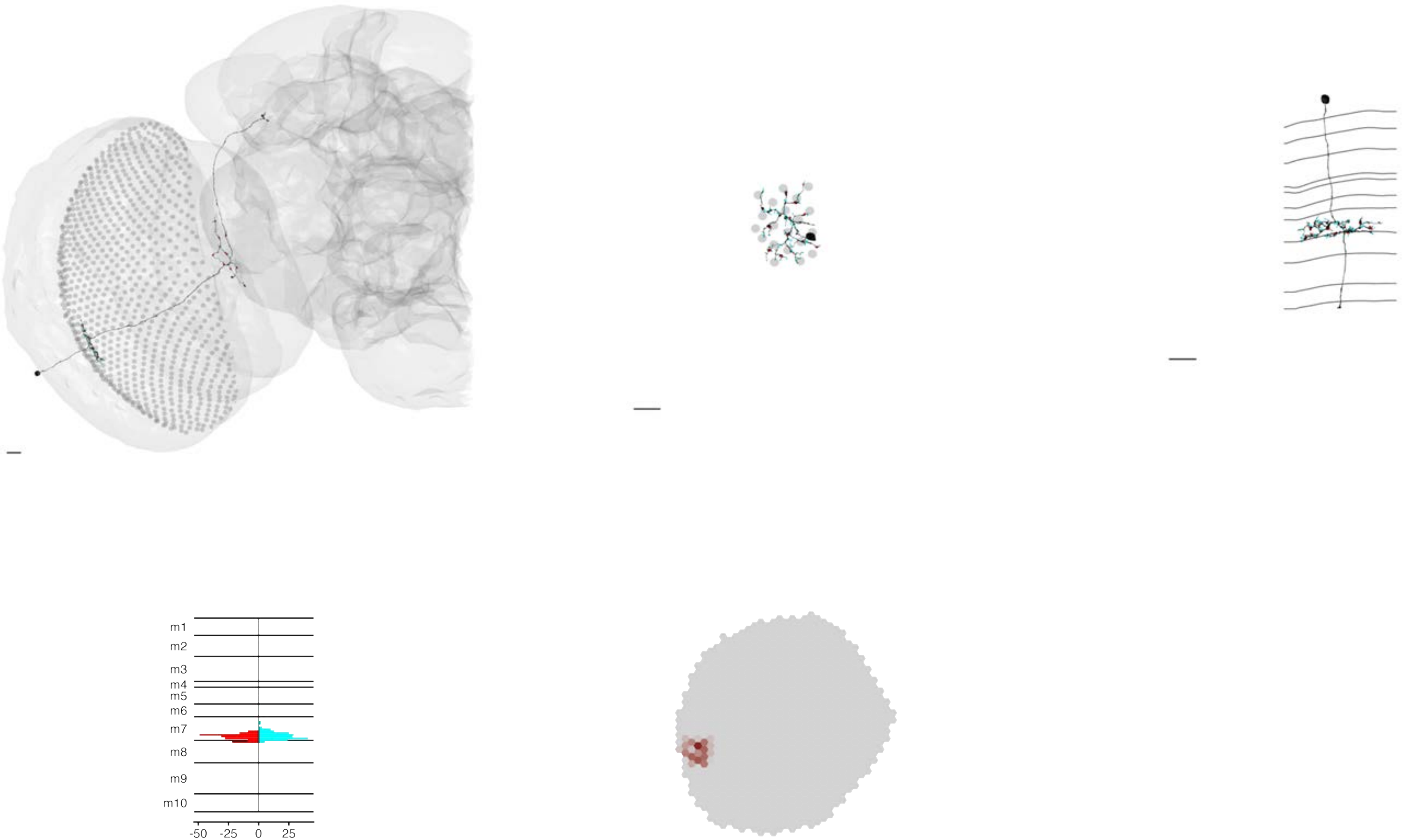

### MeTu4c_720575940622516971

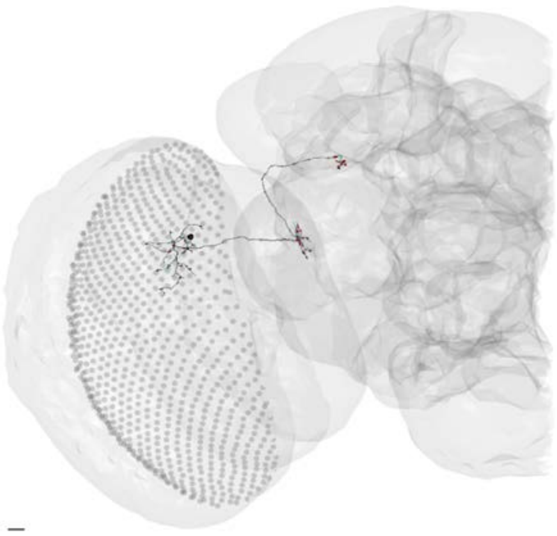

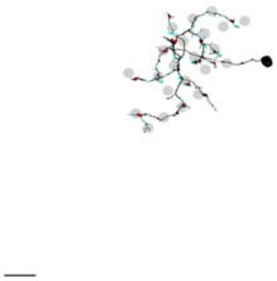

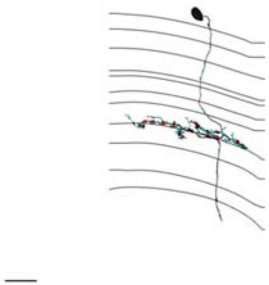

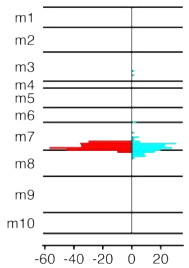


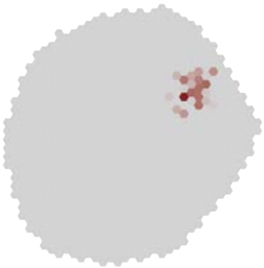

### MeTu4c_720575940622541252

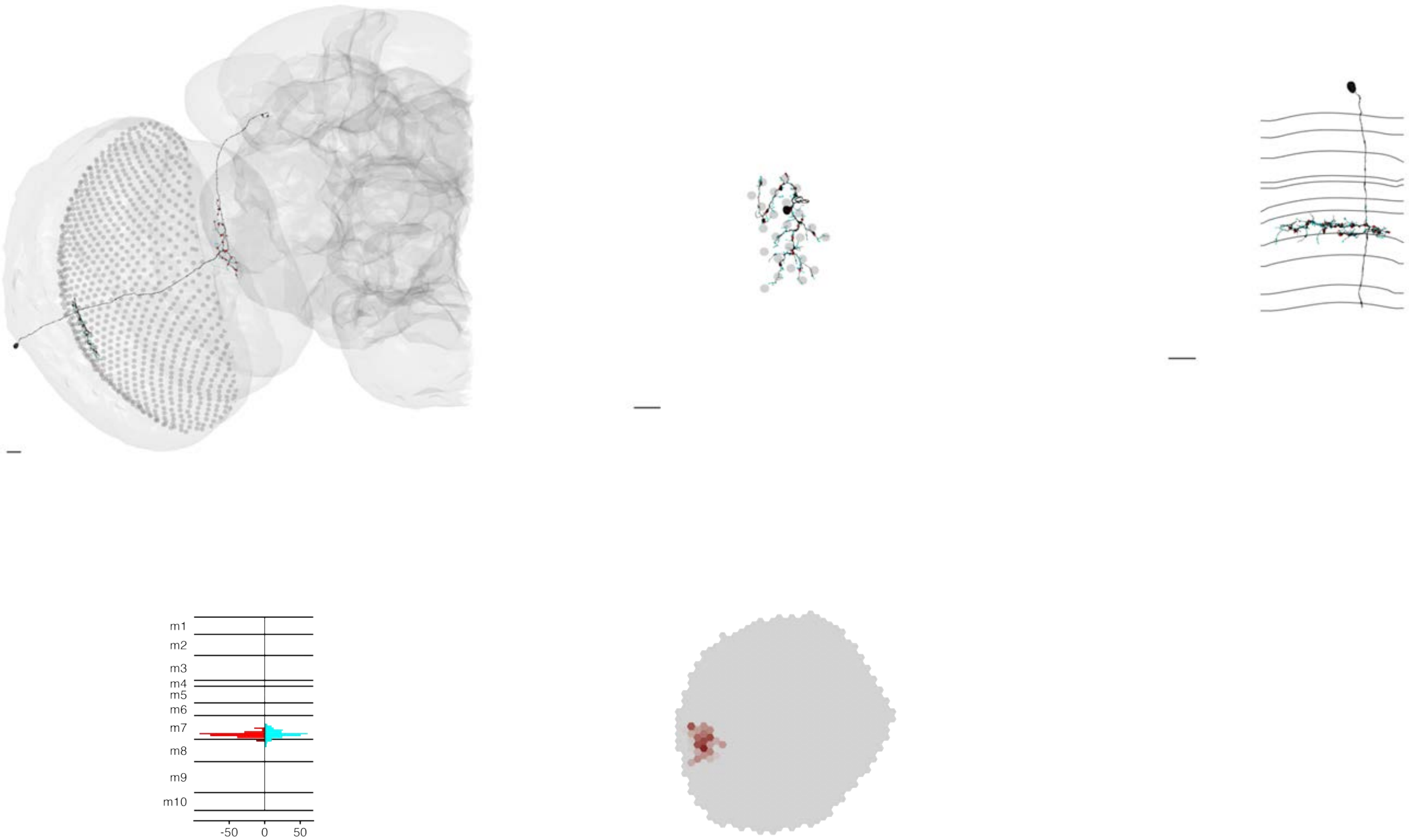

### MeTu4c_720575940622760717

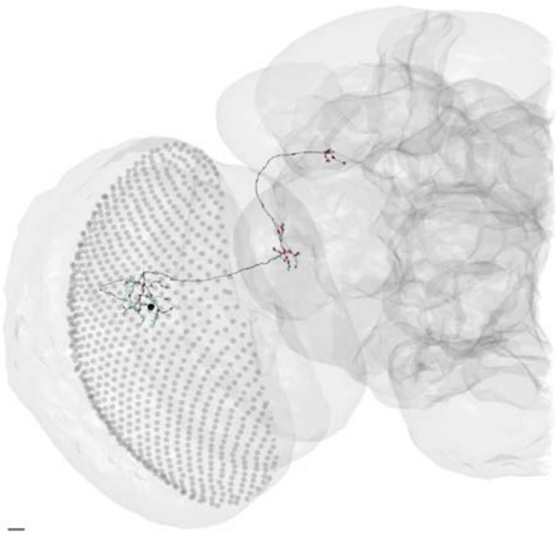

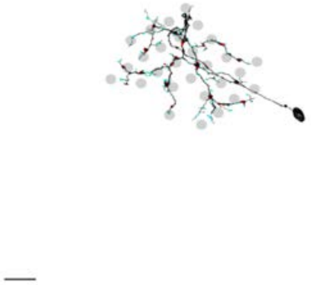

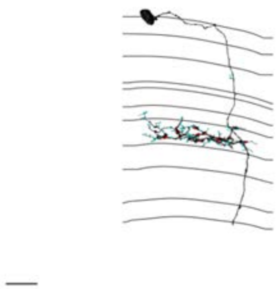

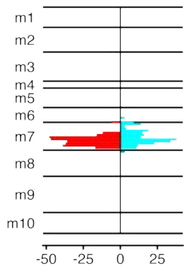


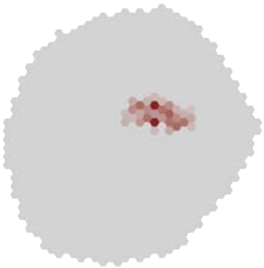

### MeTu4c_720575940622814588

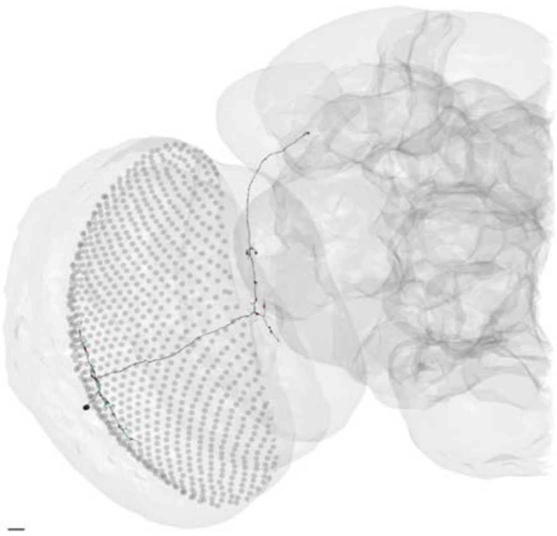

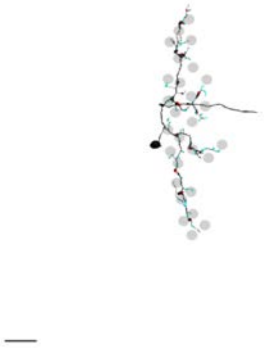

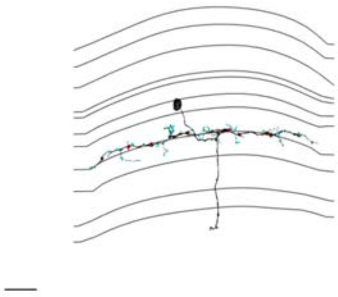

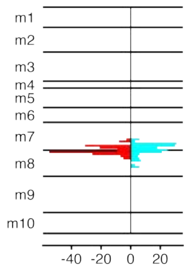


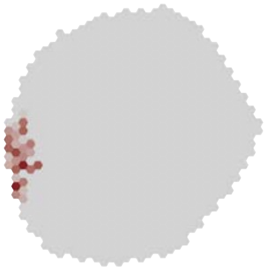

### MeTu4c_720575940623275593

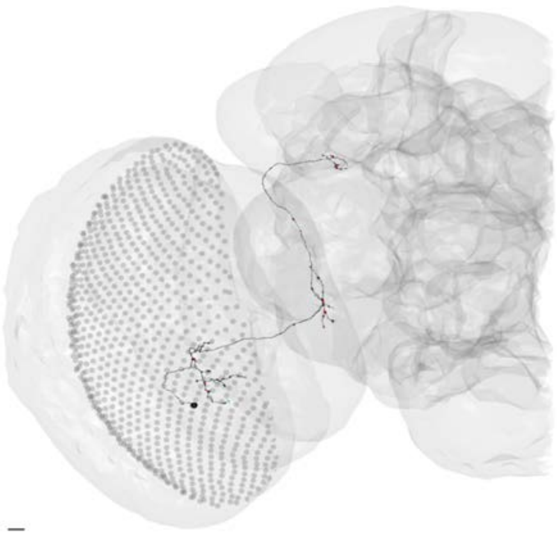

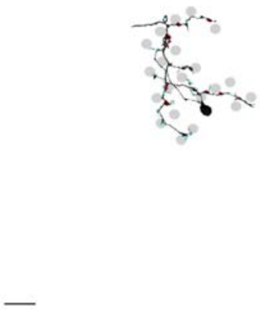

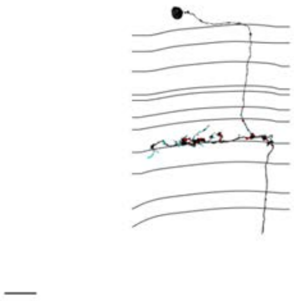

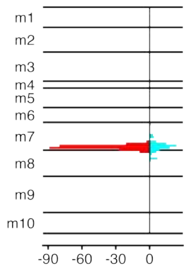


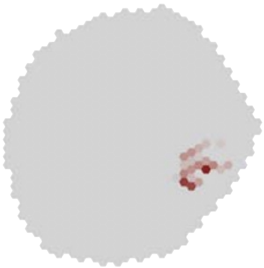

### MeTu4c_720575940624369642

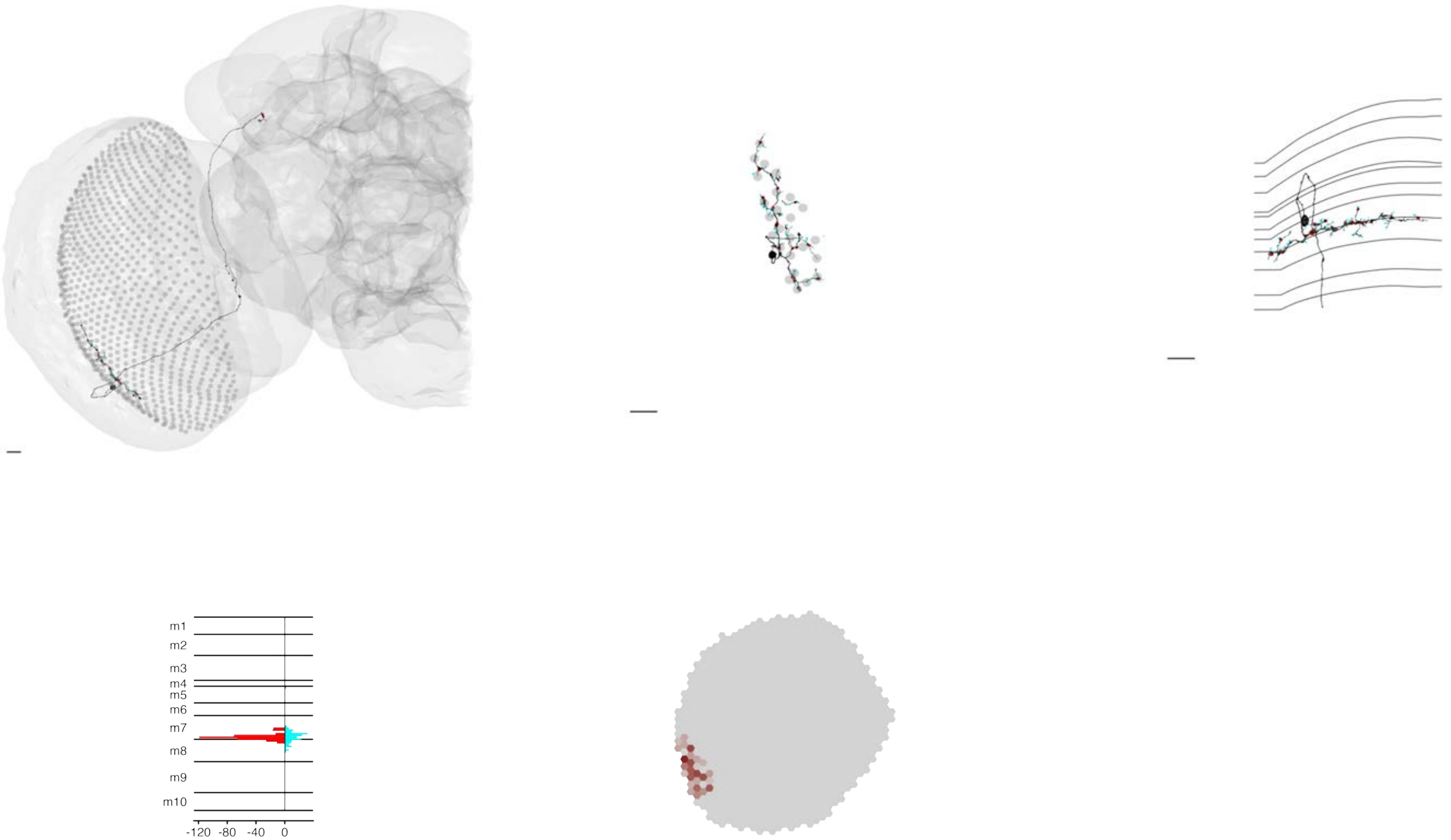

### MeTu4c_720575940626452220

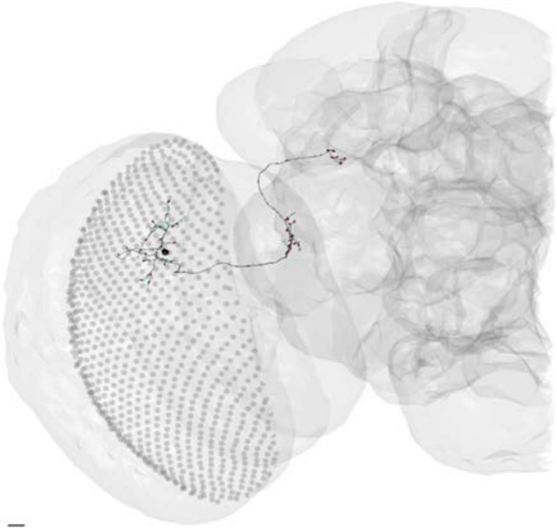

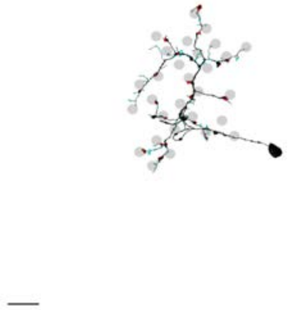

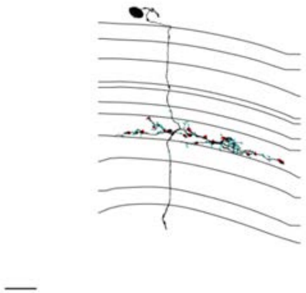

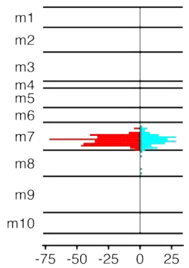


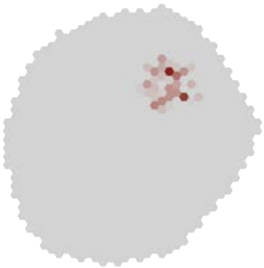

### MeTu4c_720575940627051536

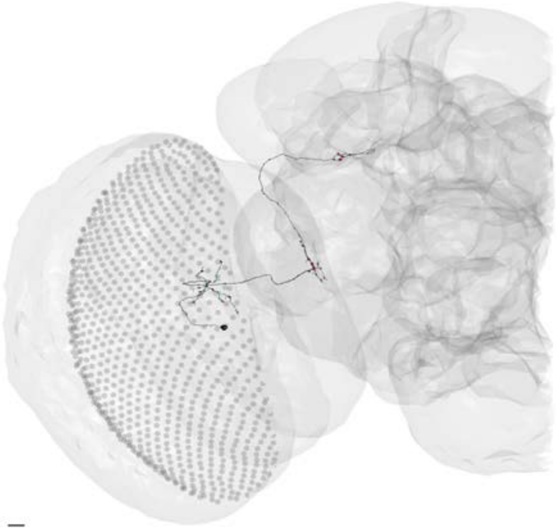

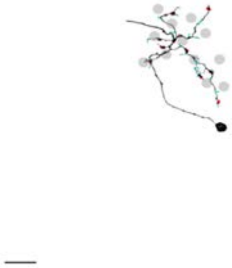

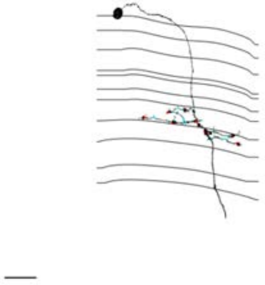

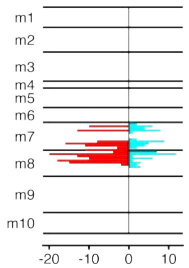


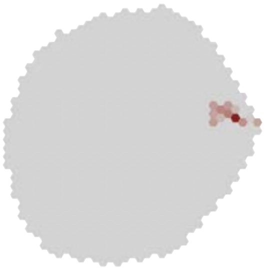

### MeTu4c_720575940627322627

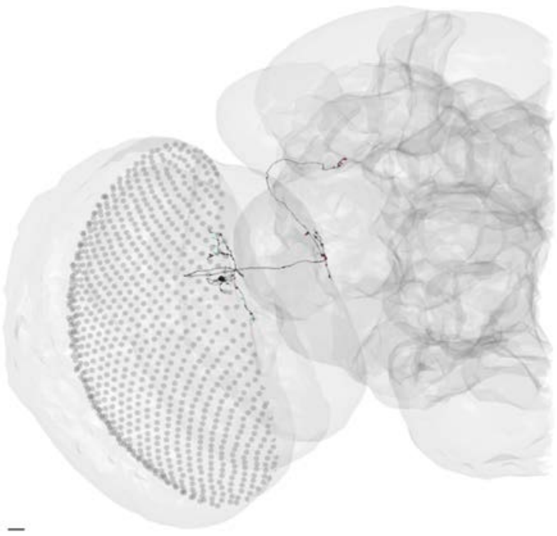

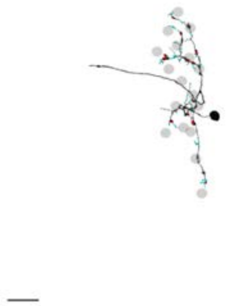

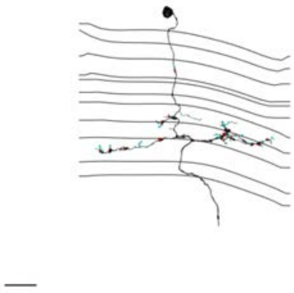

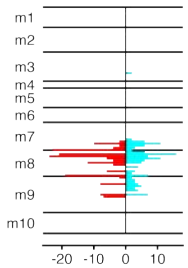


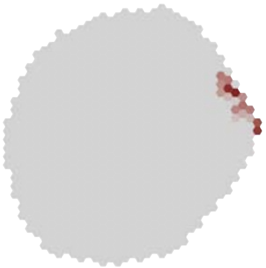

### MeTu4c_720575940627619687

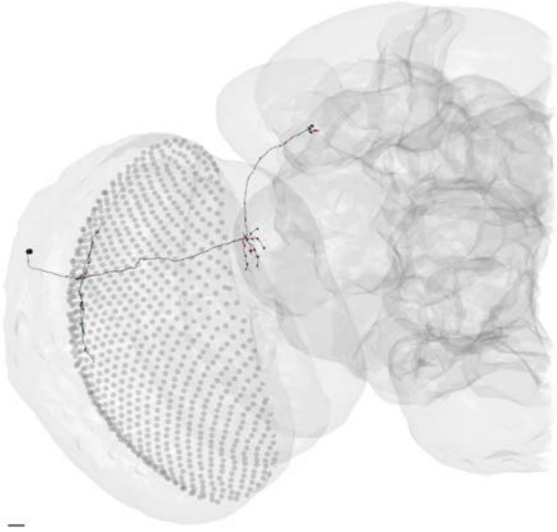

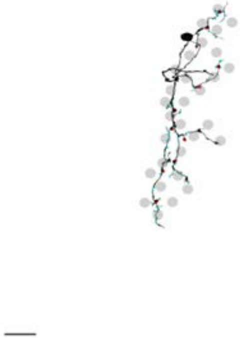

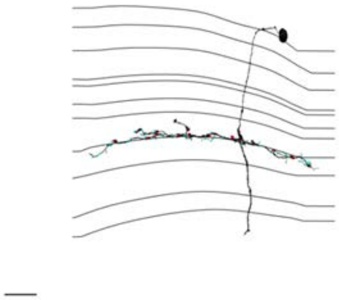

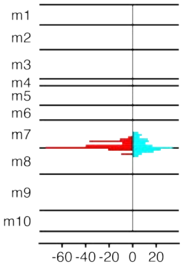


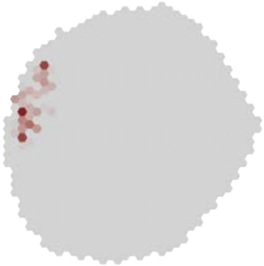

### MeTu4c_720575940627911273

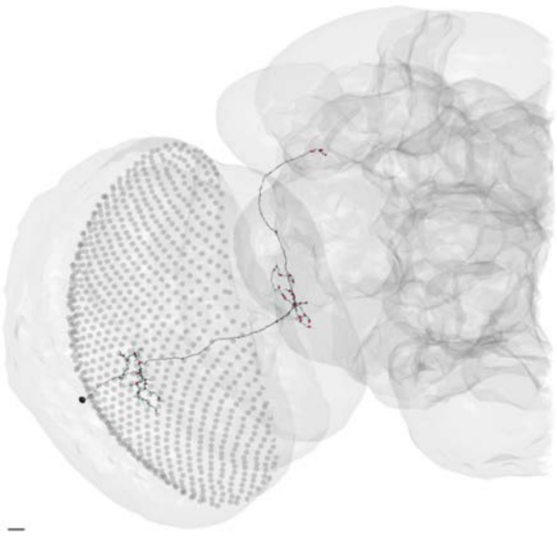

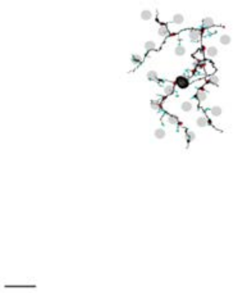

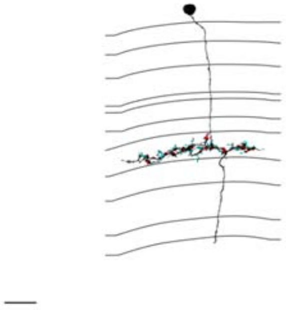

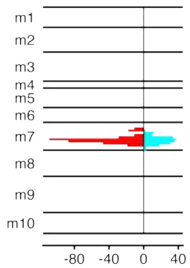


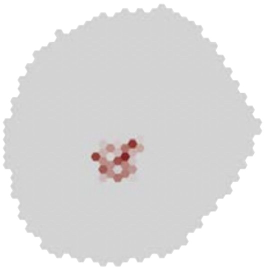

### MeTu4c_720575940627920719

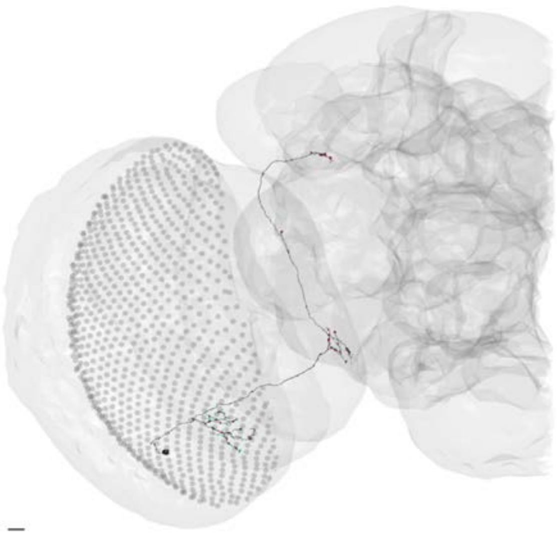

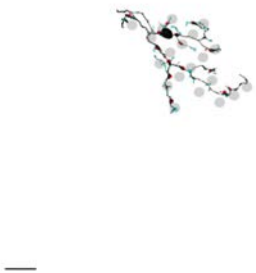

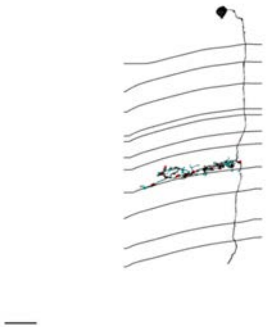

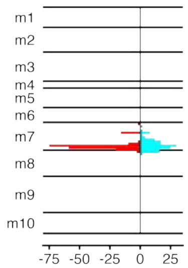


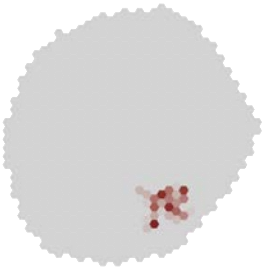

### MeTu4c_720575940628321284

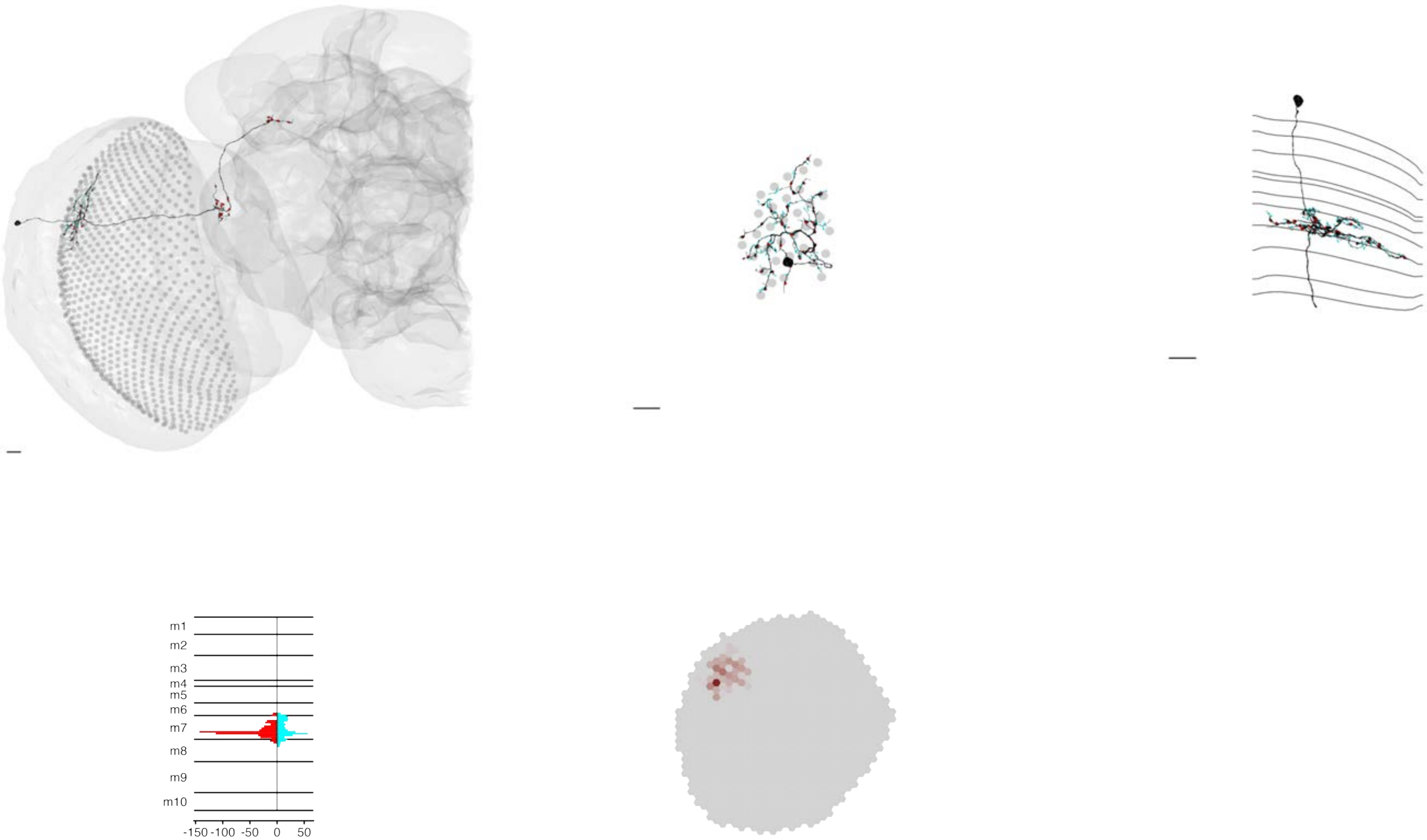

### MeTu4c_720575940628416772

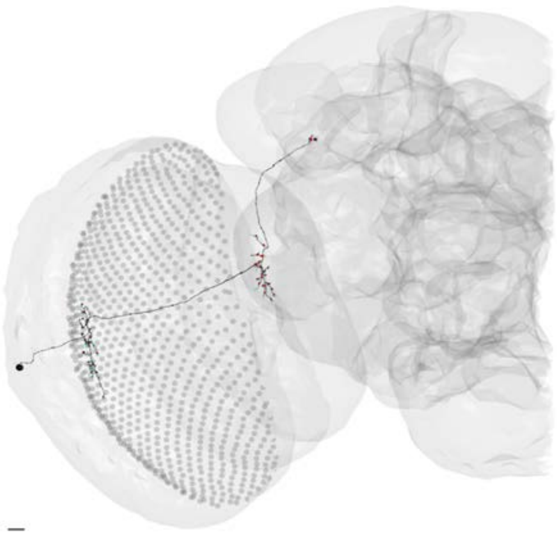

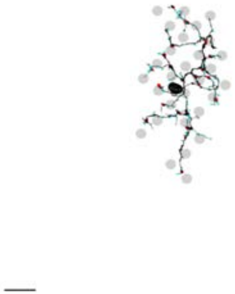

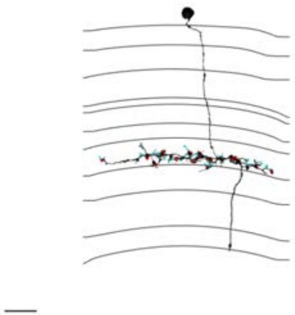

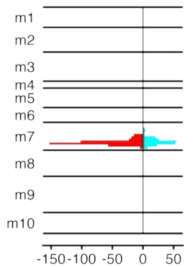


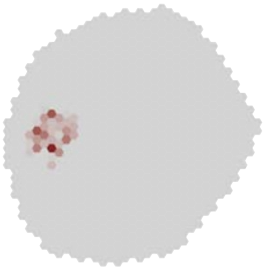

### MeTu4c_720575940628565320

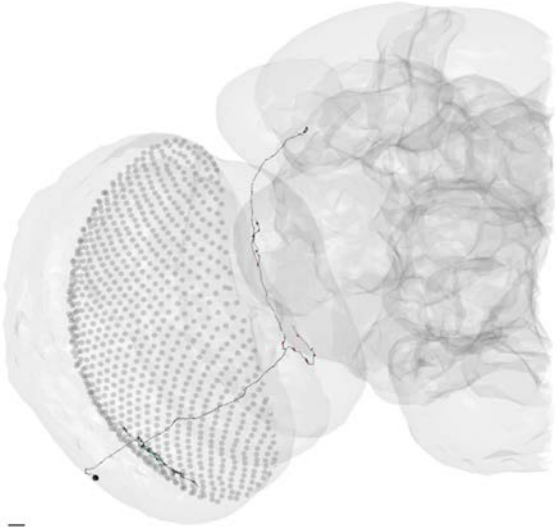

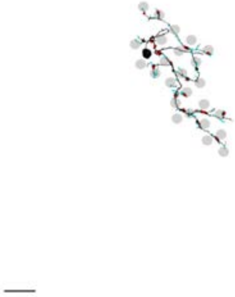

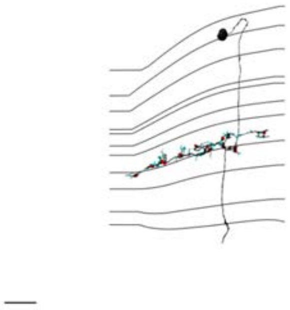

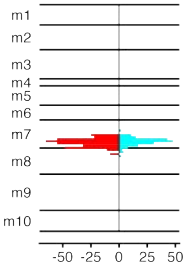


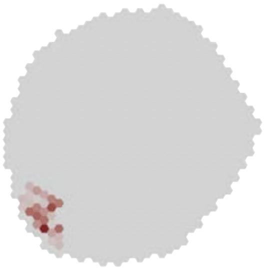

### MeTu4c_720575940628787815

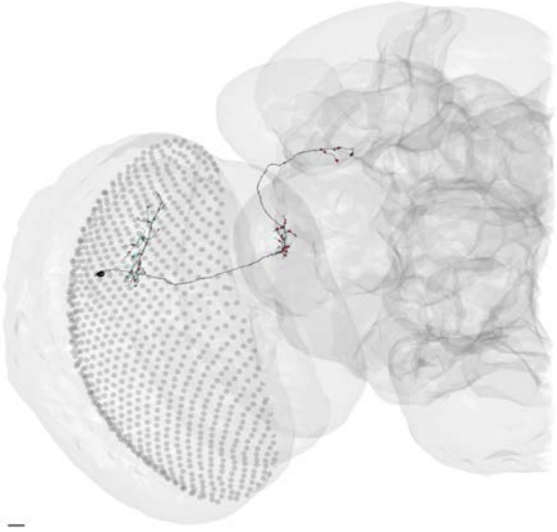

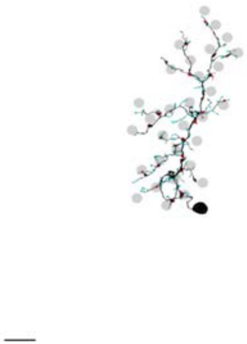

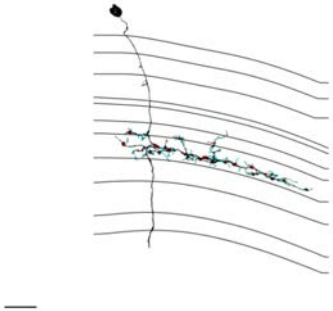

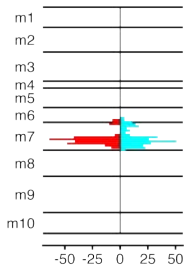


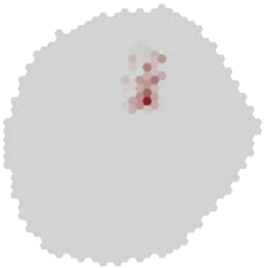

### MeTu4c_720575940629083643

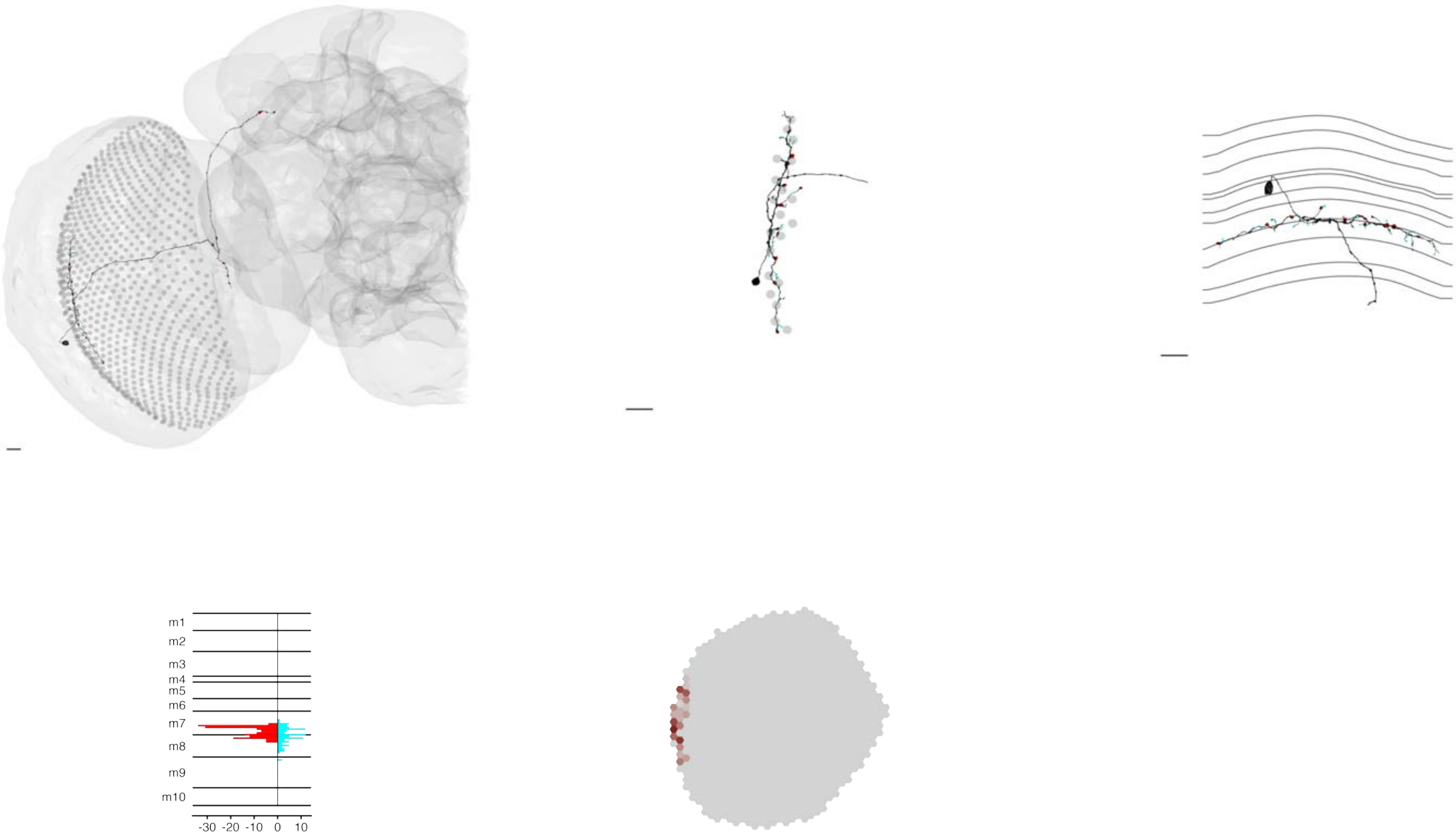

### MeTu4c_720575940629756524

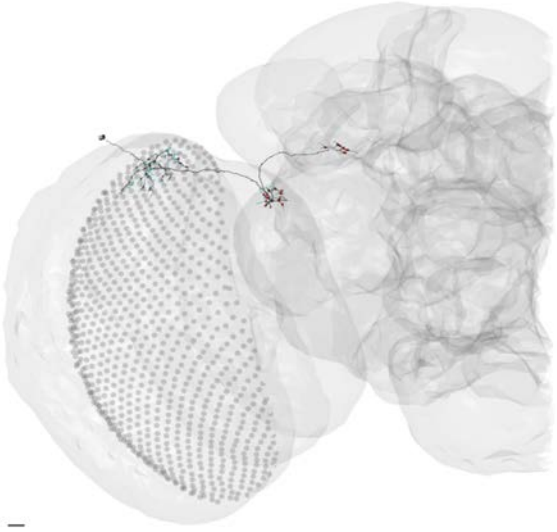

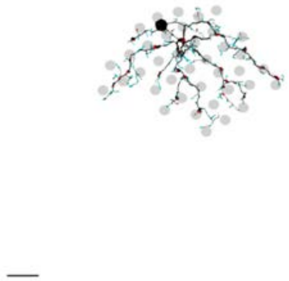

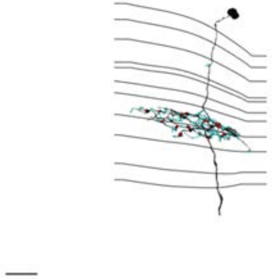

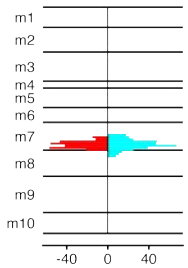


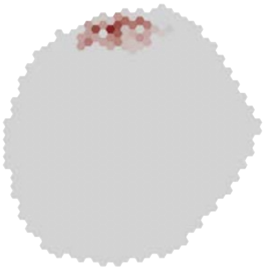

### MeTu4c_720575940631479460

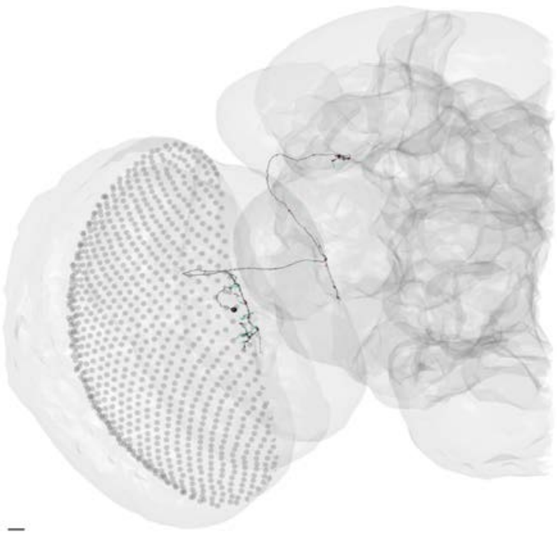

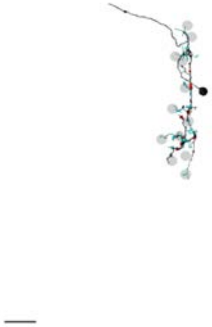

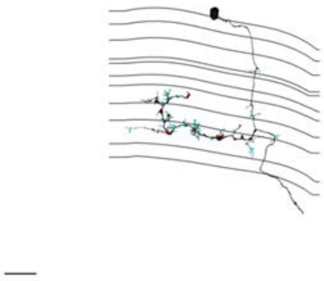

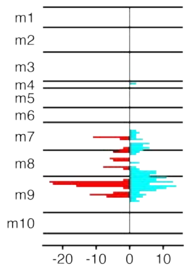


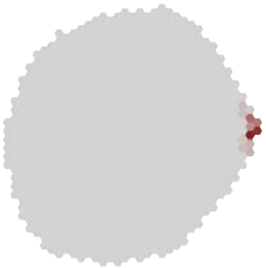

### MeTu4c_720575940631709900

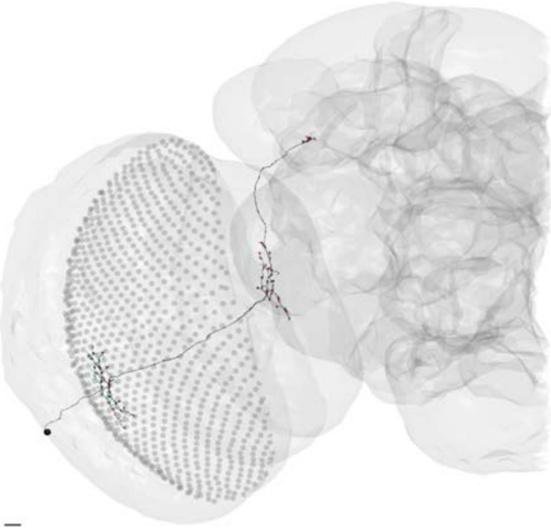

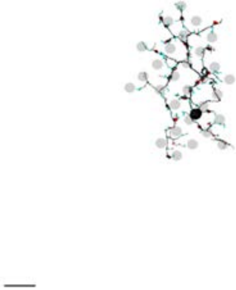

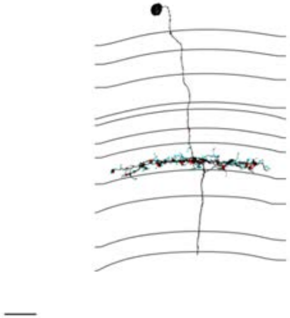

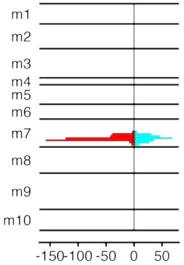


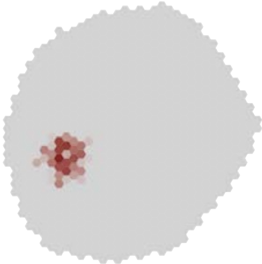

### MeTu4c_720575940633837779

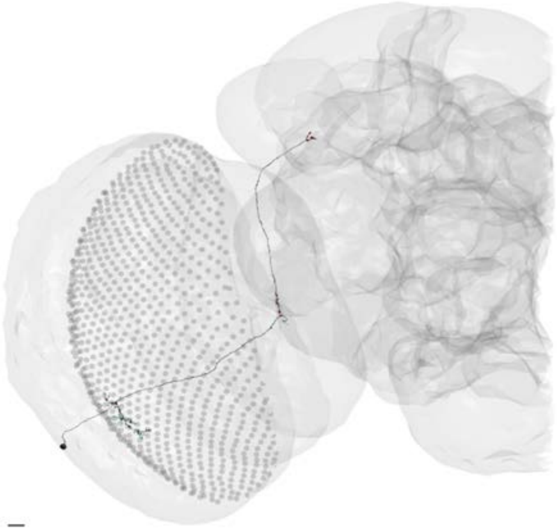

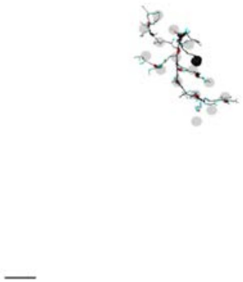

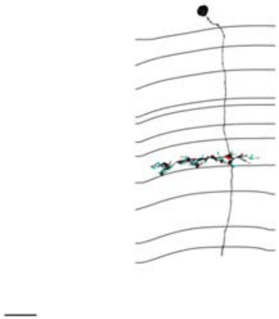

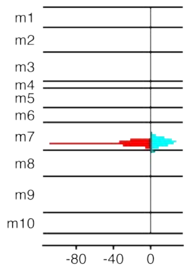


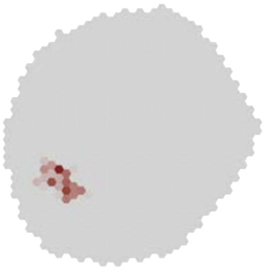

### MeTu4c_720575940634827188

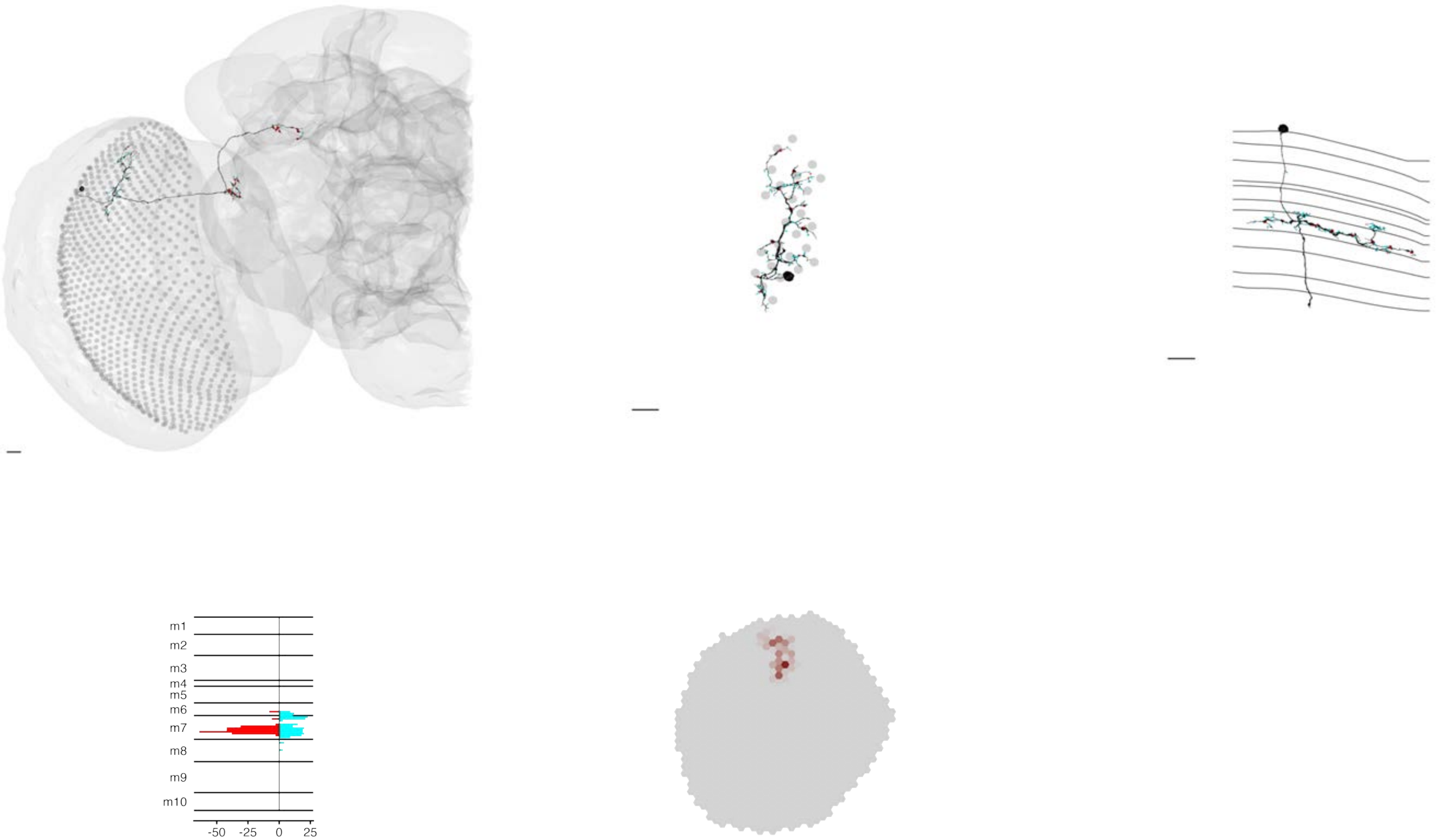

### MeTu4c_720575940635647630

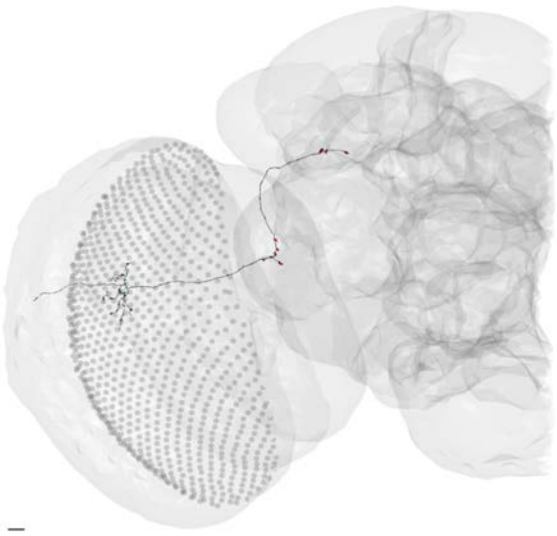

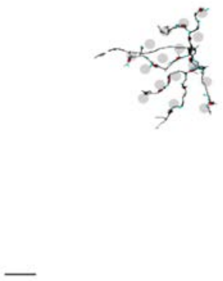

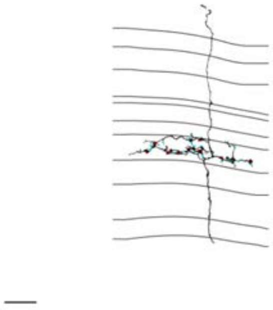

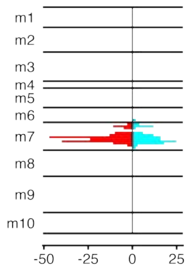


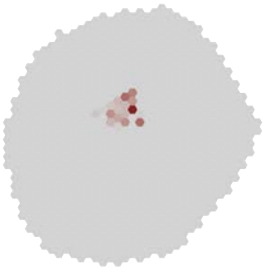

### MeTu4c_720575940638341198

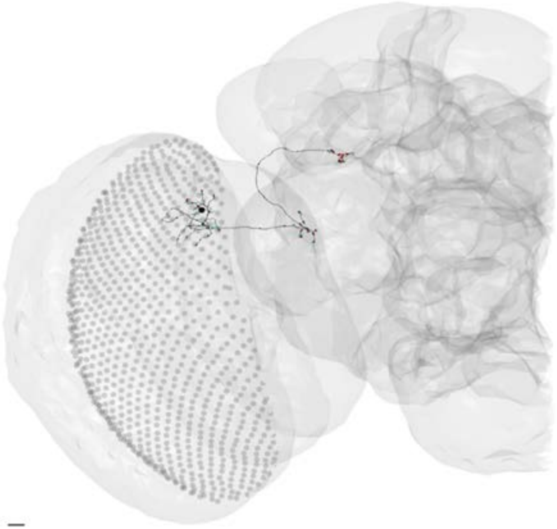

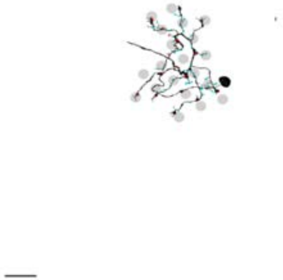

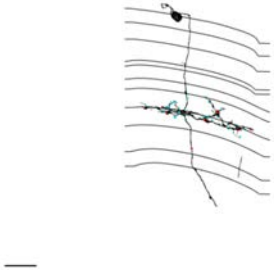

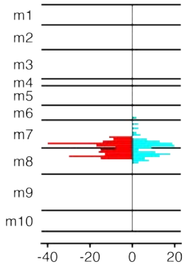


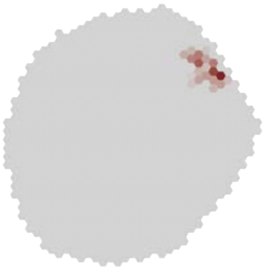

### MeTu4c_720575940639039184

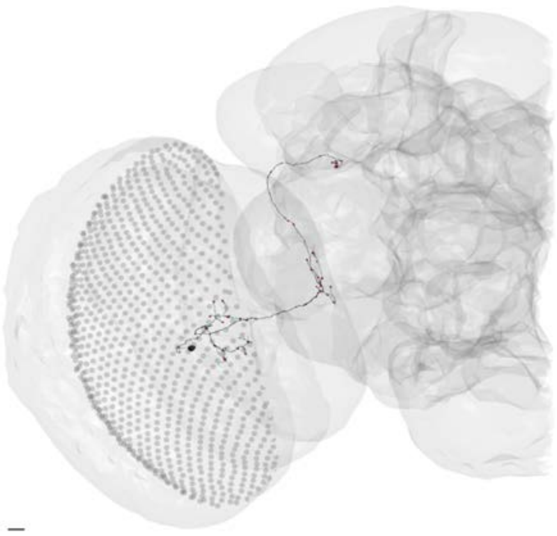

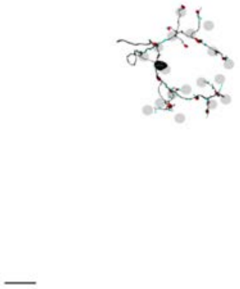

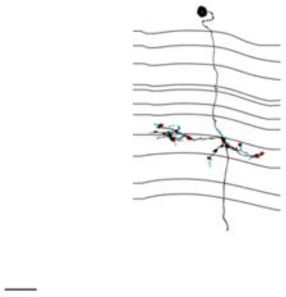

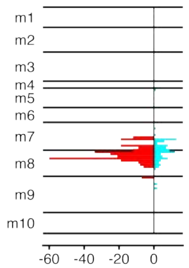


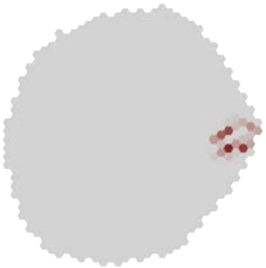

### MeTu4c_720575940640040832

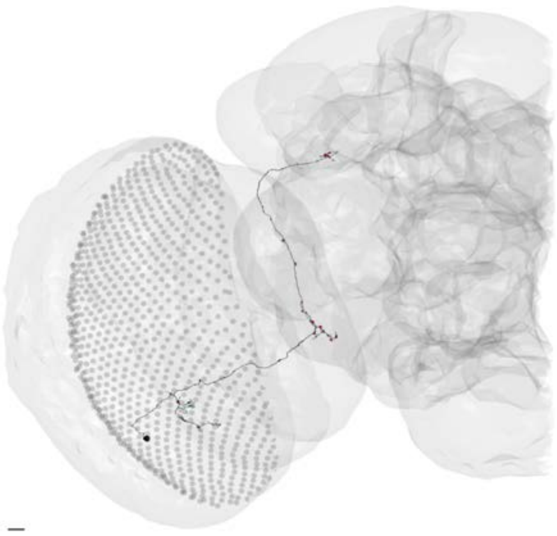

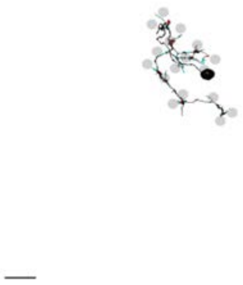

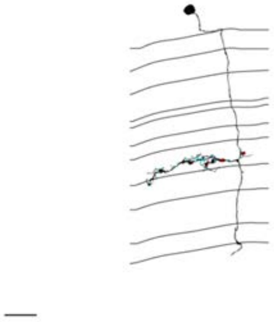

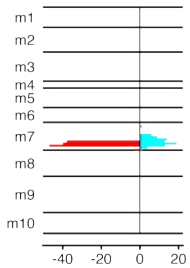


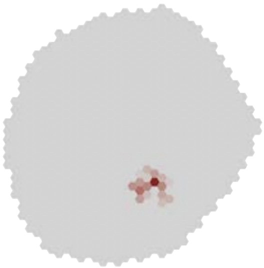

### MeTu4c_720575940643051405

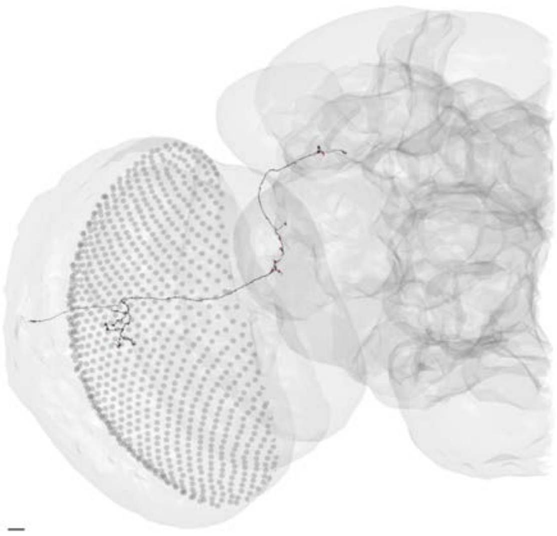

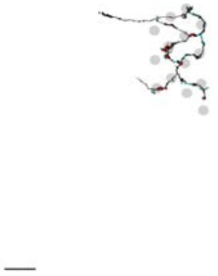

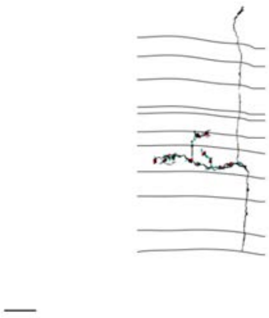

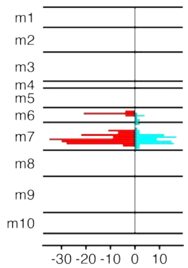


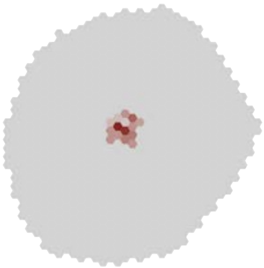

### MeTu4c_720575940644157600

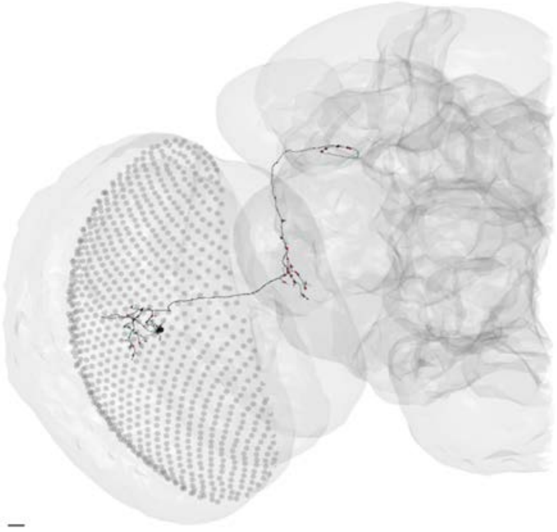

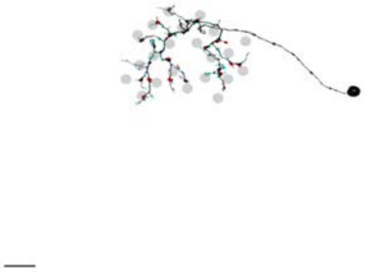

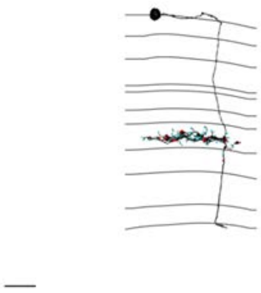

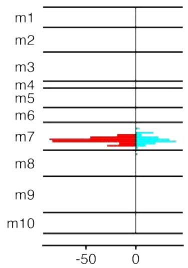


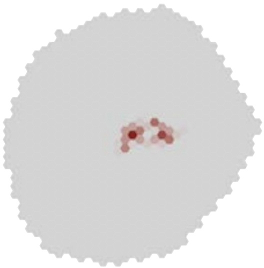

### MeTu4d_720575940609307147

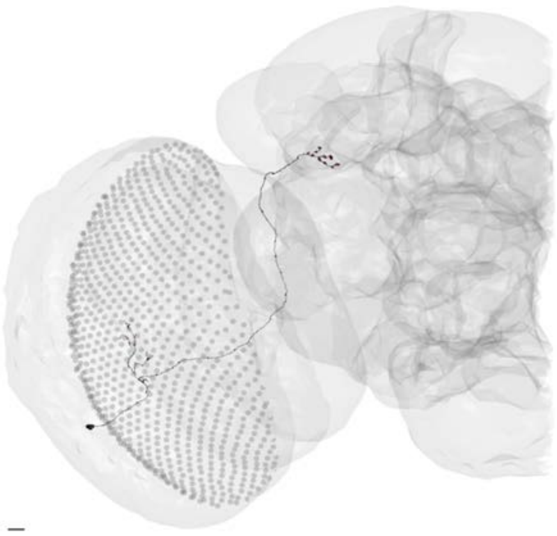

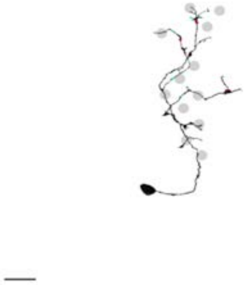

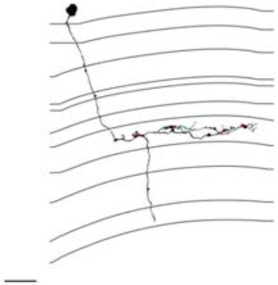

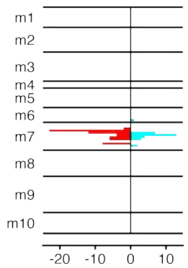


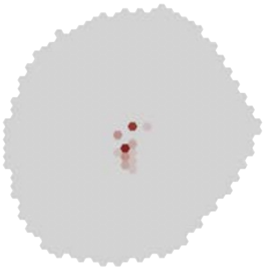

### MeTu4d_720575940612778649

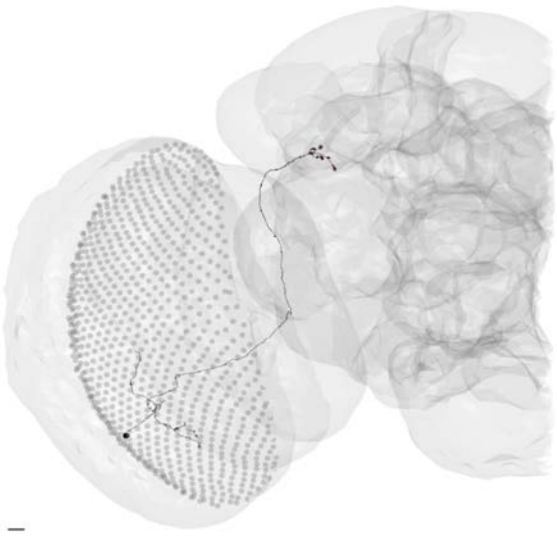

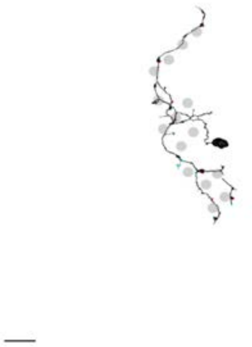

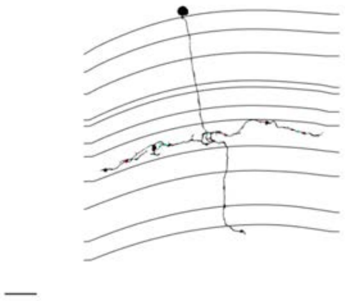

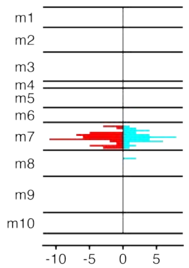


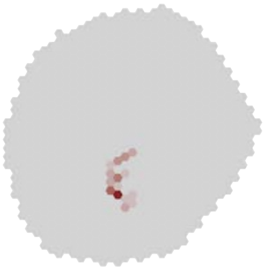

### MeTu4d_720575940614282705

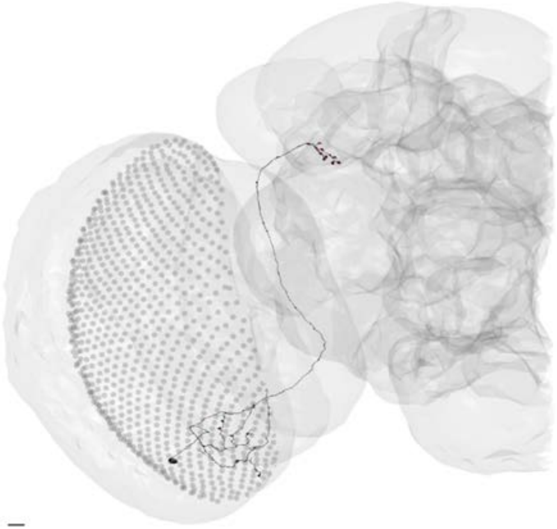

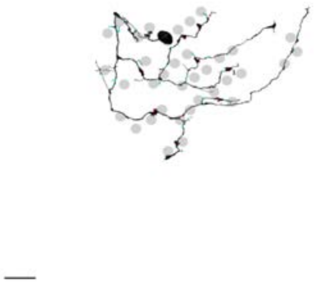

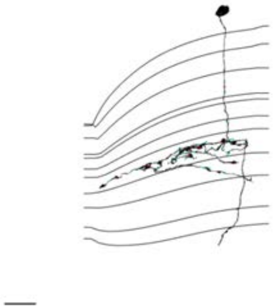

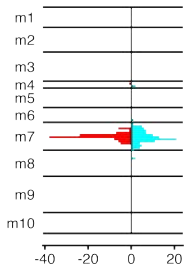


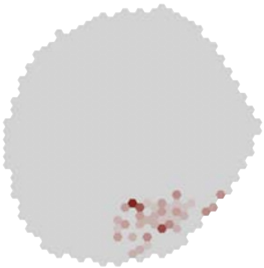

### MeTu4d_720575940619183014

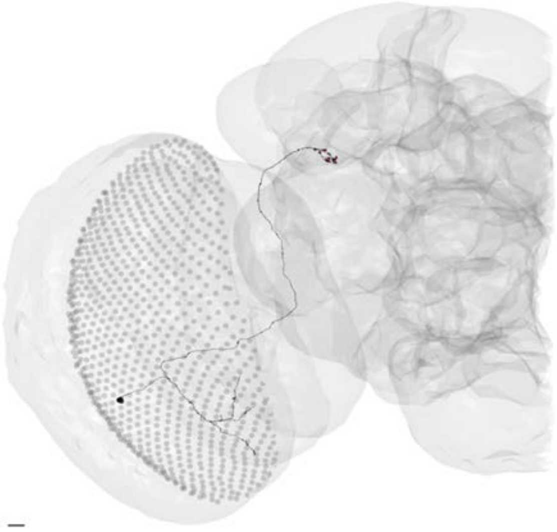

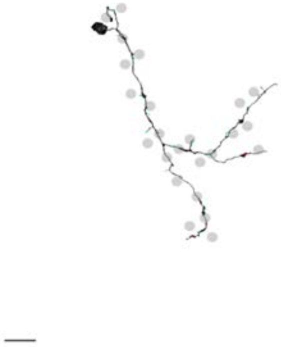

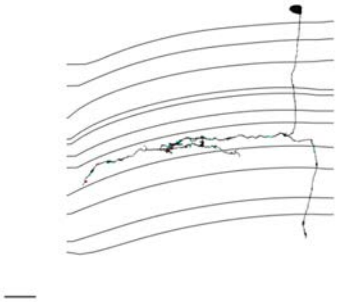

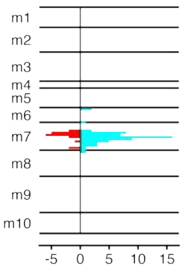


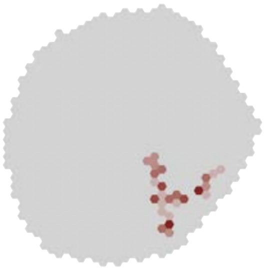

### MeTu4d_720575940621250651

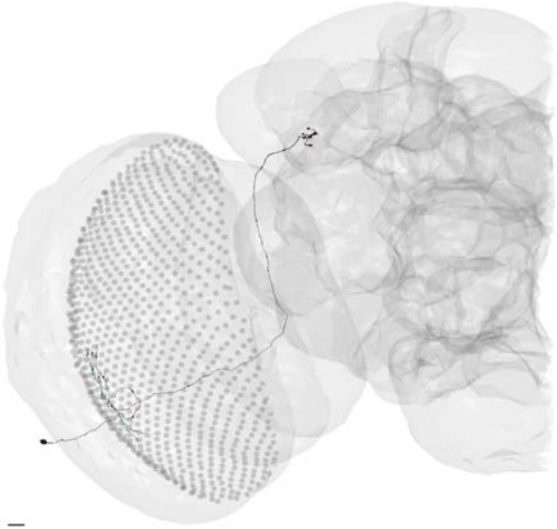

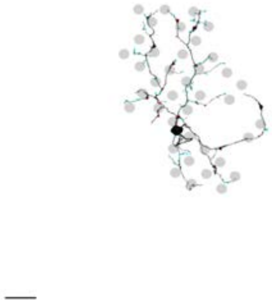

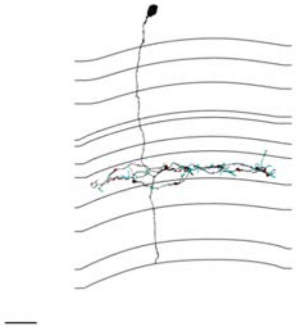

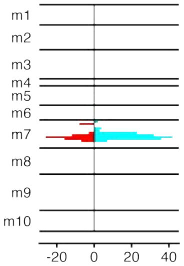


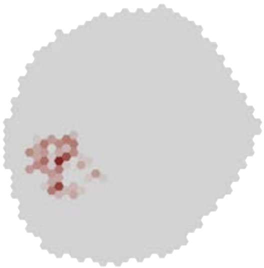

### MeTu4d_720575940621609430

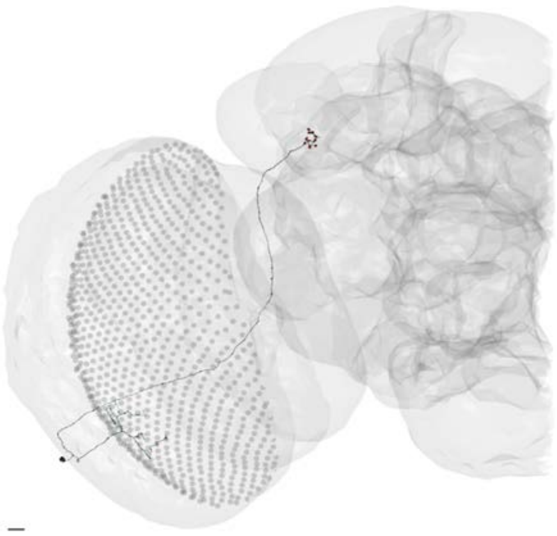

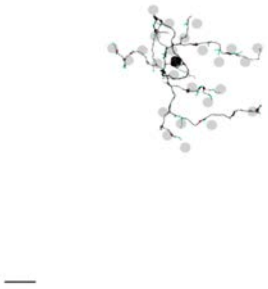

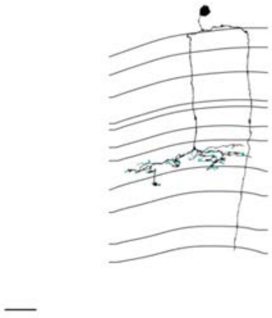

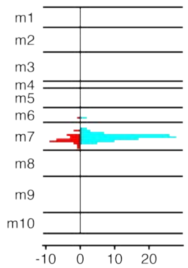


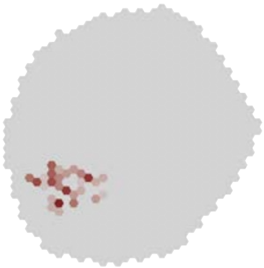

### MeTu4d_720575940621648993

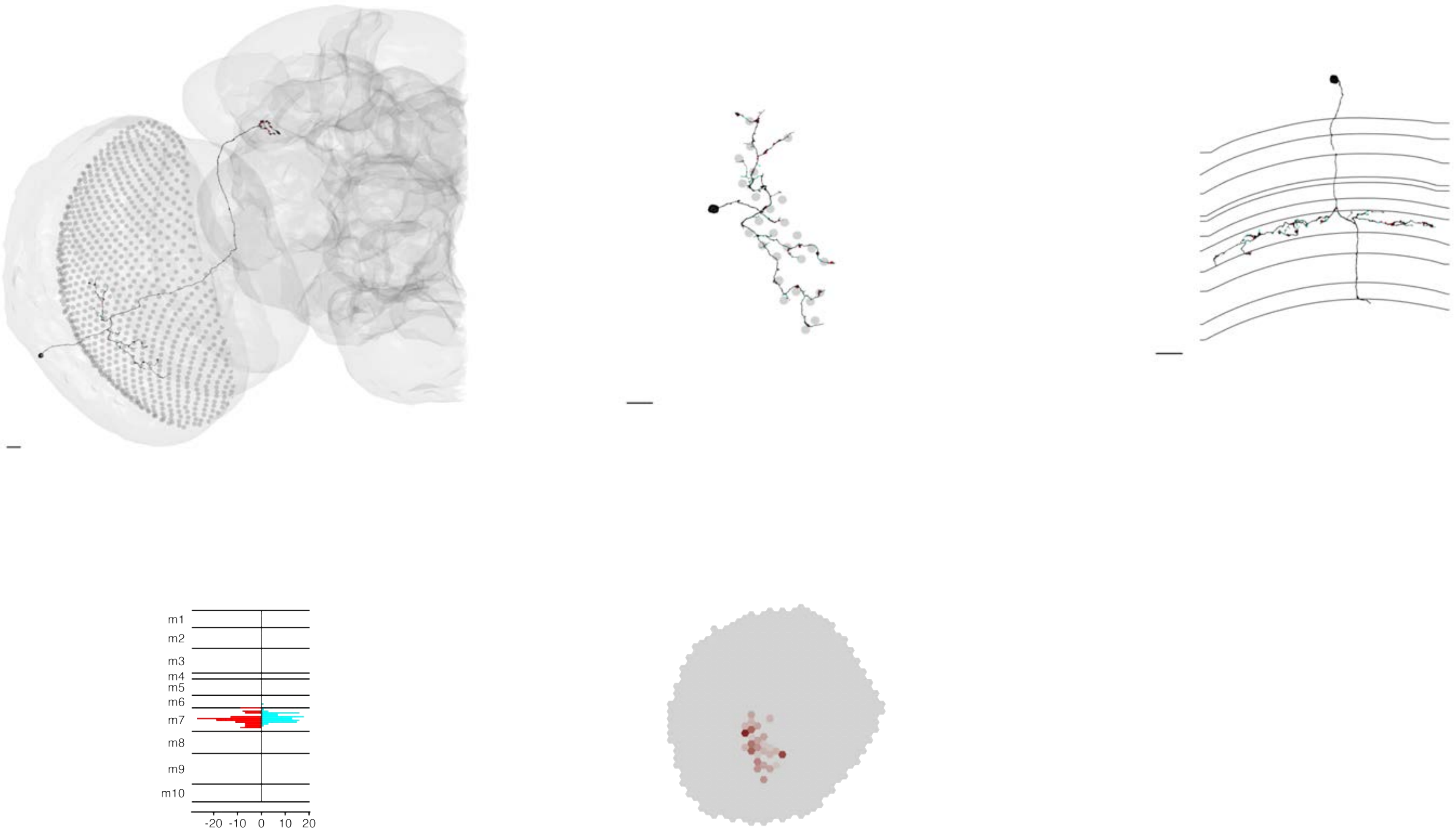

### MeTu4d_720575940622383198

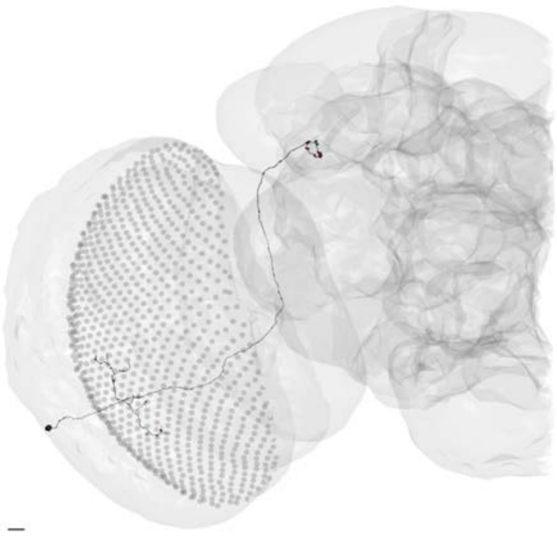

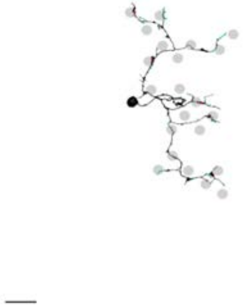

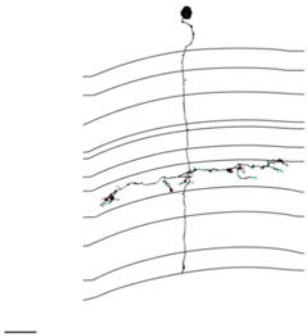

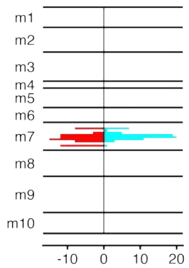


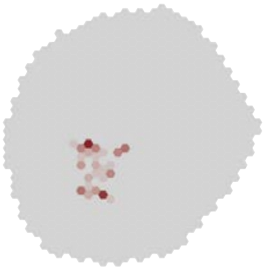

### MeTu4d_720575940622807646

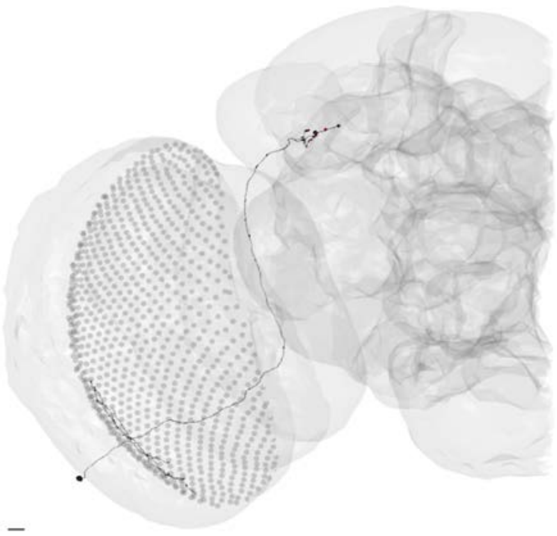

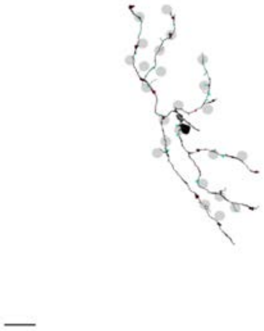

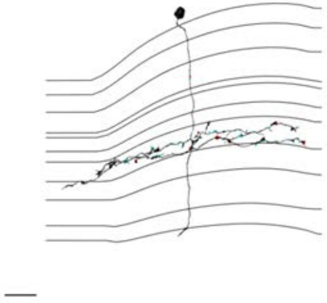

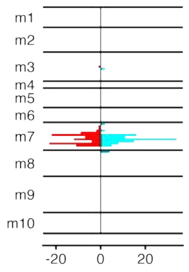


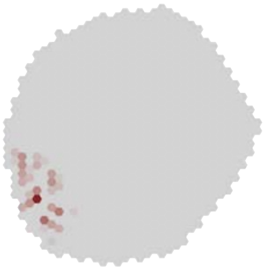

### MeTu4d_720575940625879696

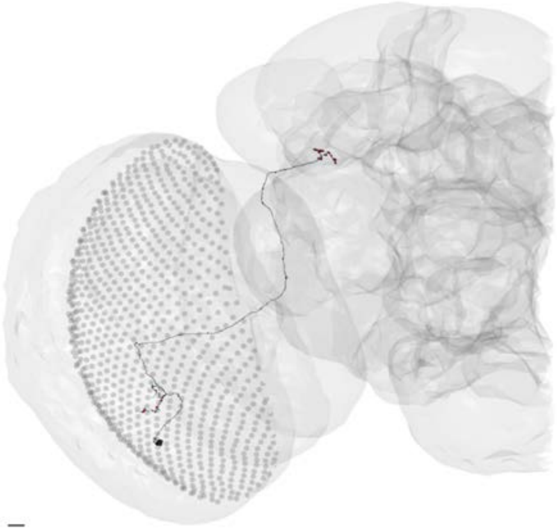

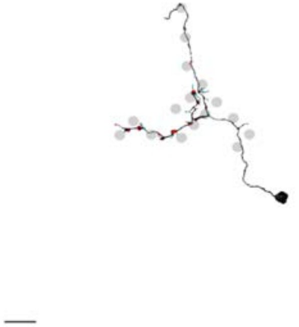

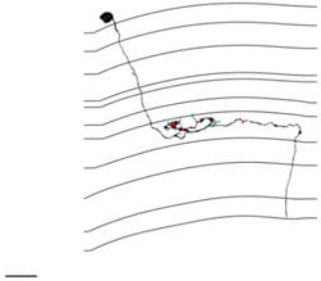

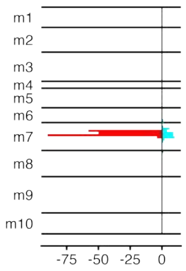


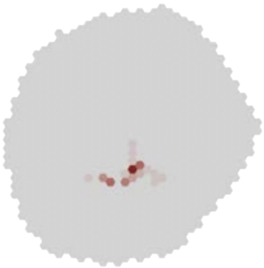

MeTu4d_720575940626588106

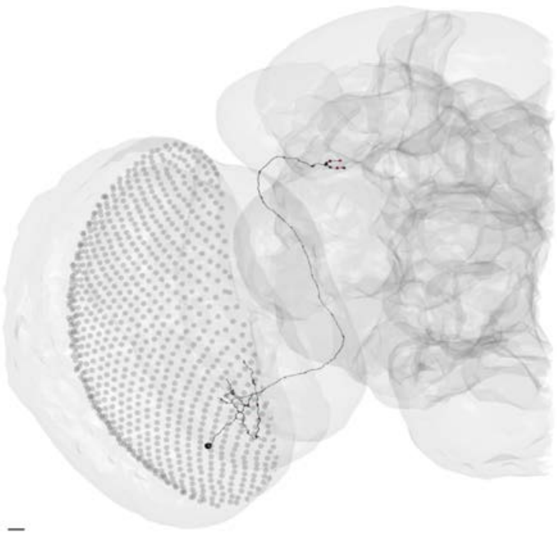

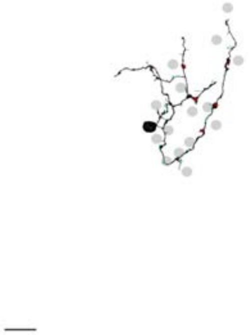

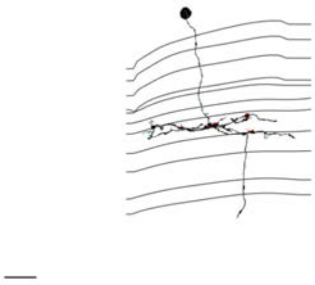

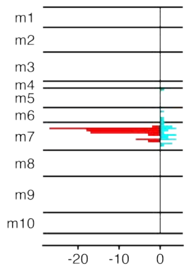


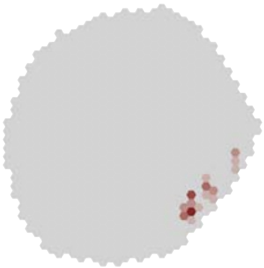

### MeTu4d_720575940629819855

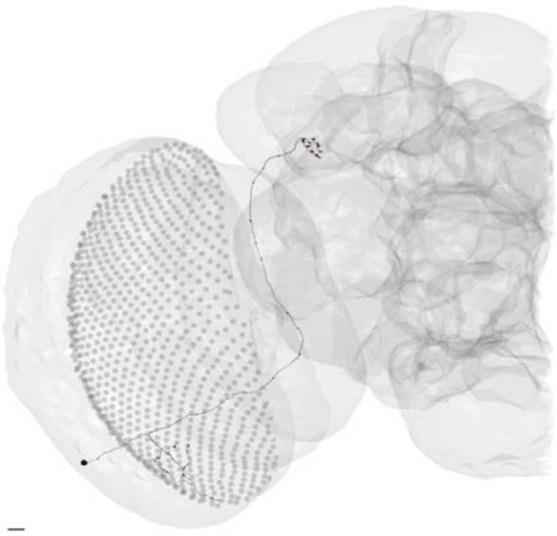

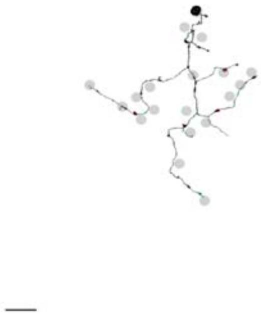

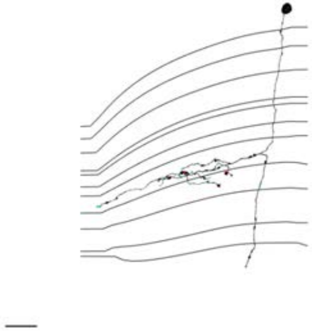

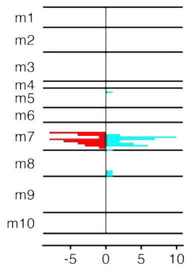


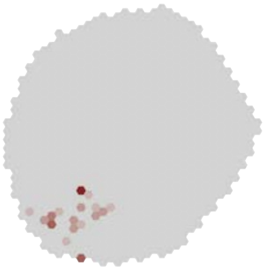

### MeTu4d_720575940630662442

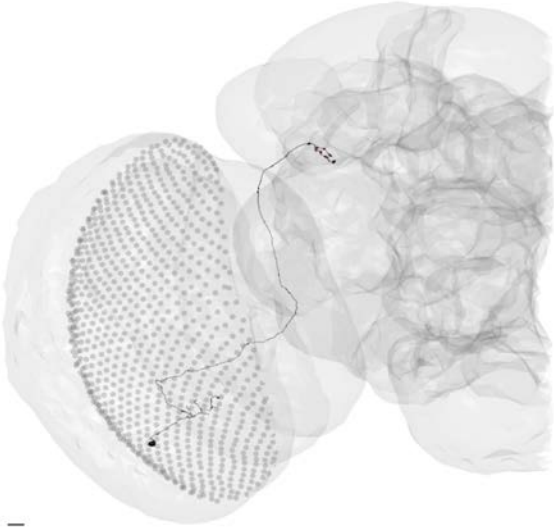

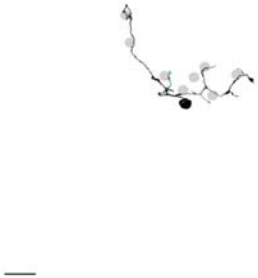

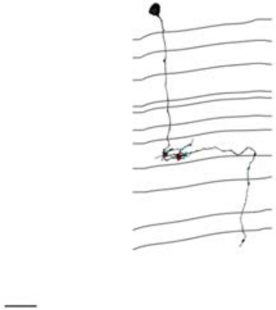

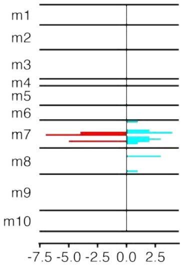


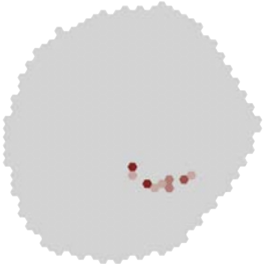

### MeTu4d_720575940631077480

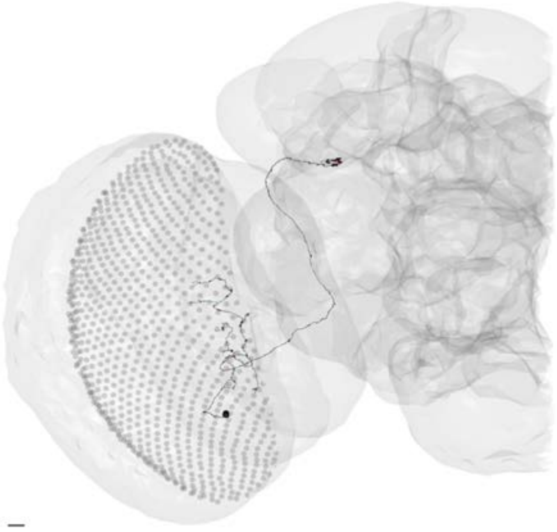

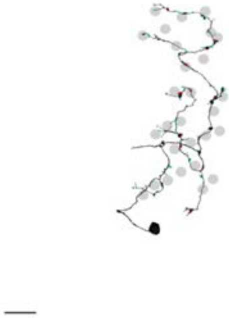

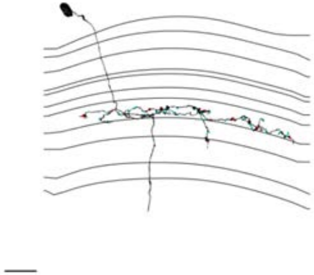

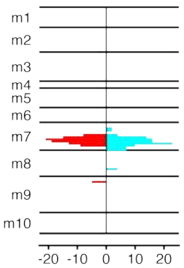


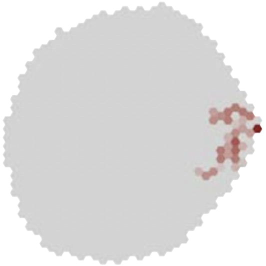

### MeTu4d_720575940633295967

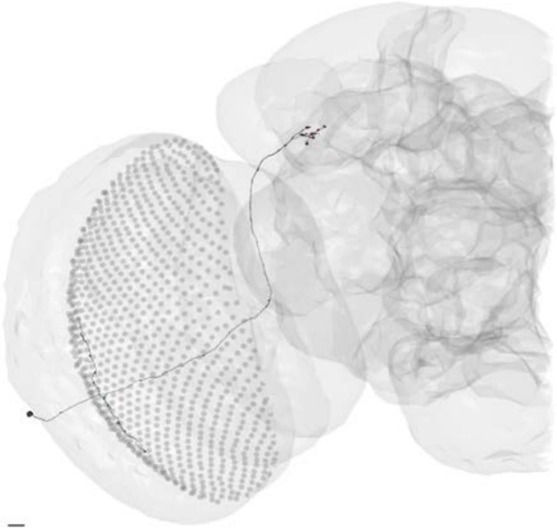

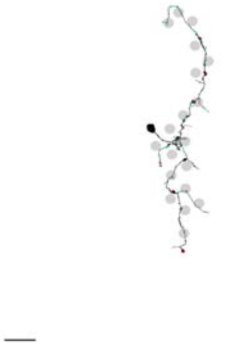

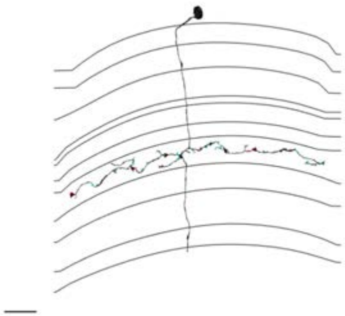

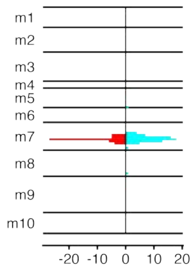


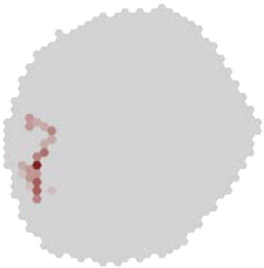

### MeTu4d_720575940633372499

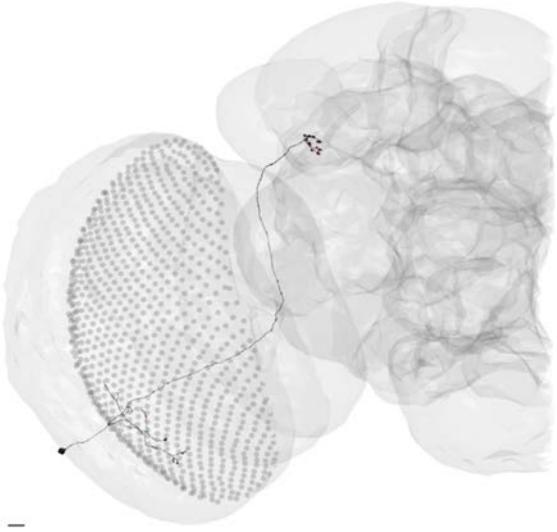

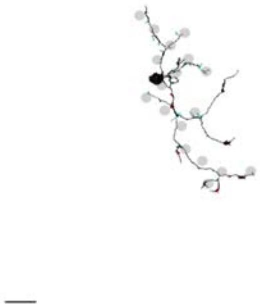

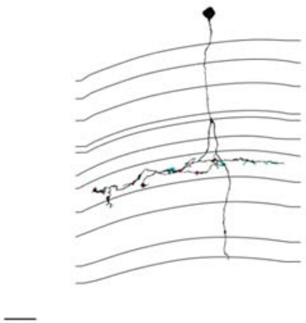

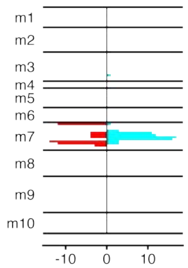


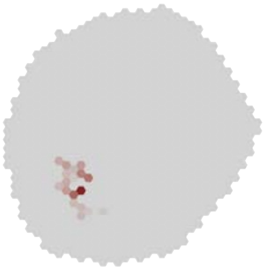

### MeTu4d_720575940636083380

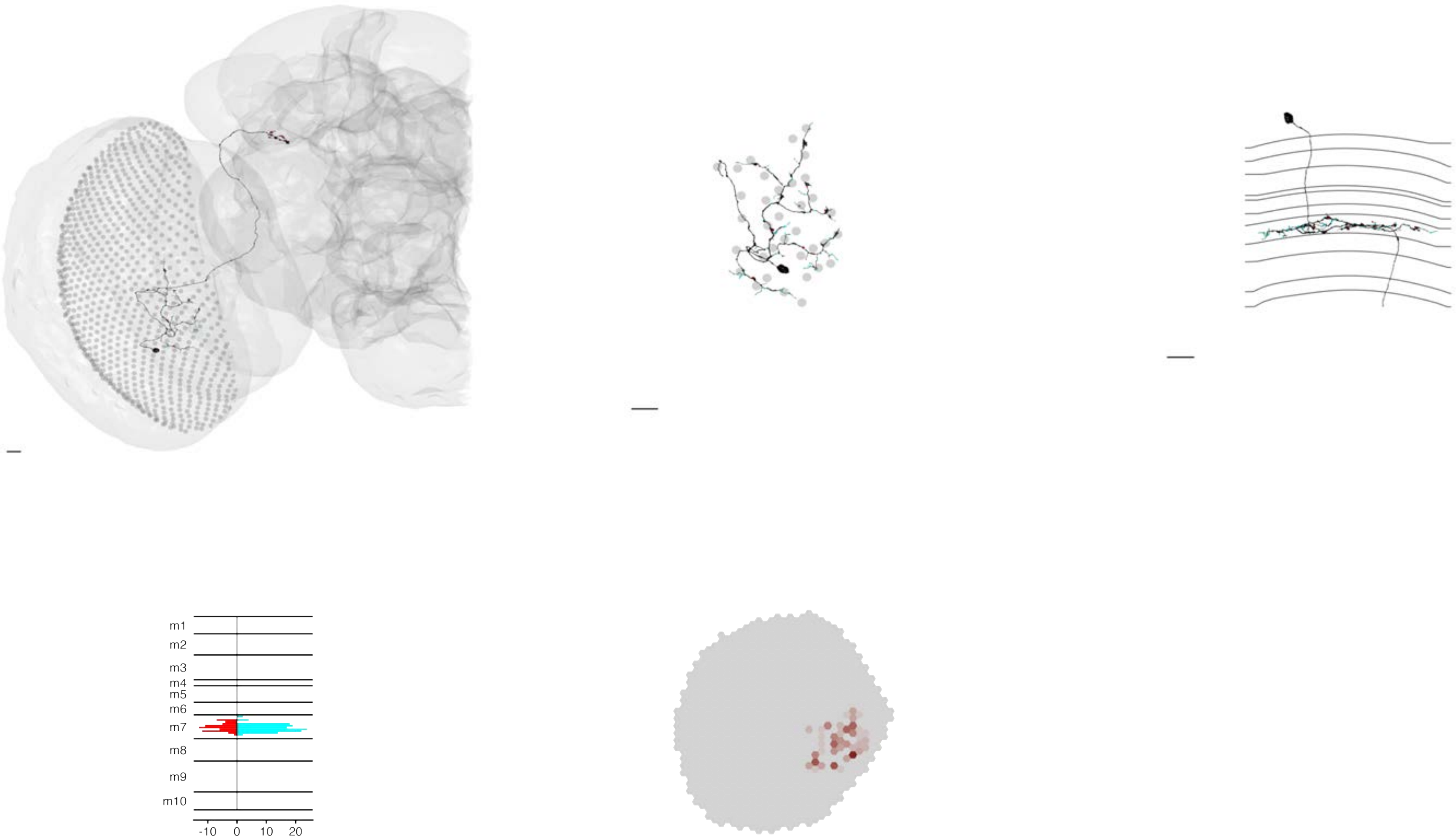

### MeTu4d_720575940647352196

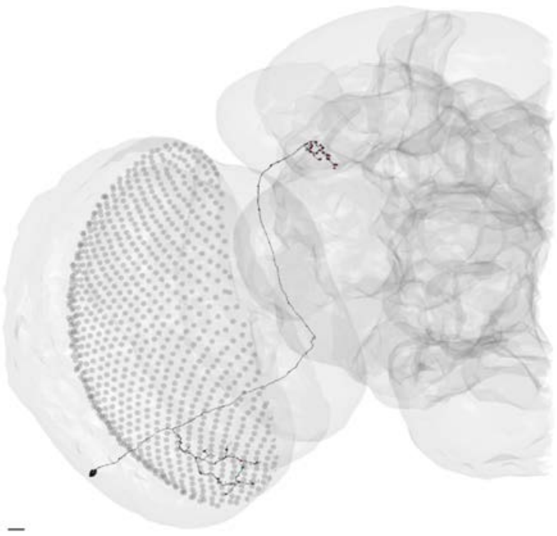

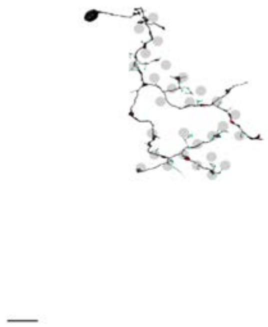

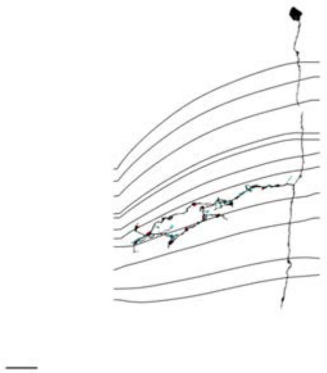

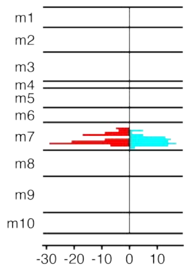


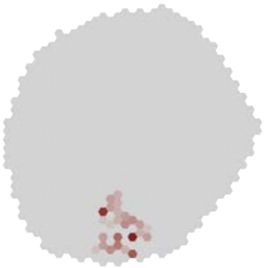
