## Supplementary material for "Connectomic reconstruction predicts the functional organization of visual inputs to the navigation center of the *Drosophila* brain": Data set 2

## ER2_ad 720575940628733292

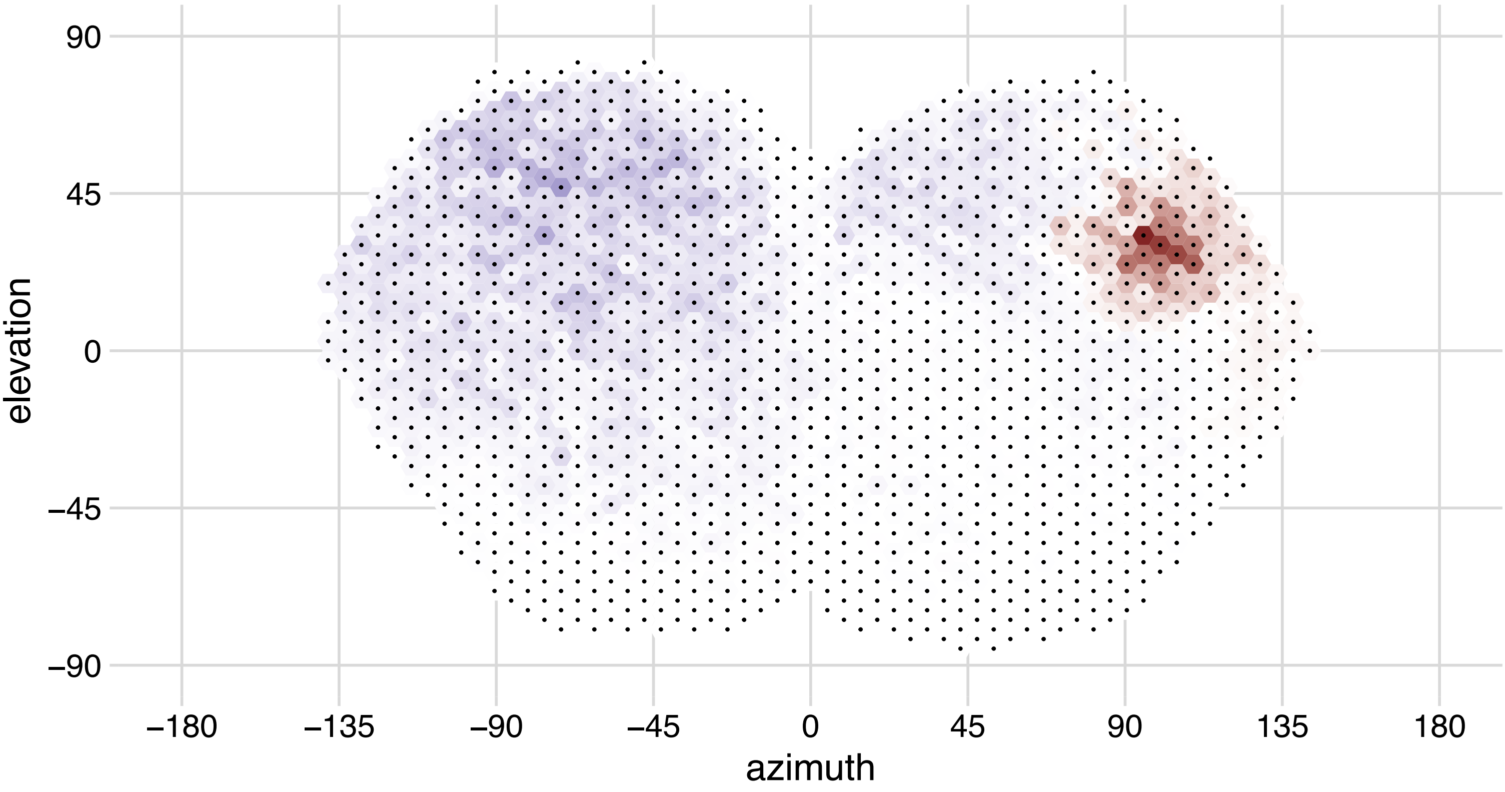

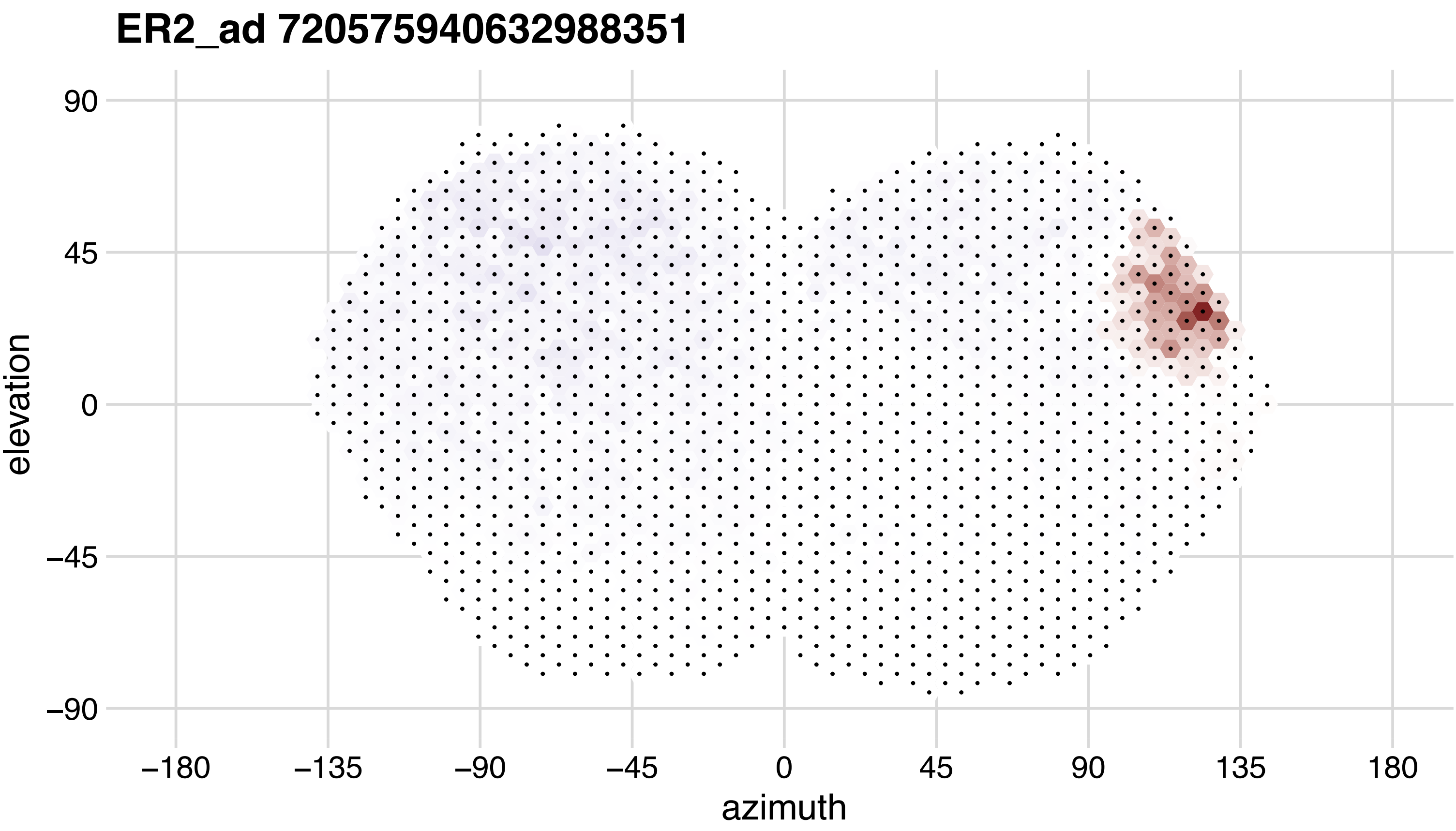
ER2_ad 720575940632988351
elevation
azimuth
90
45
0
−45
−90
−180
−135
−90
−45
0
45
90
135
180

## ER2_ad 720575940635373935

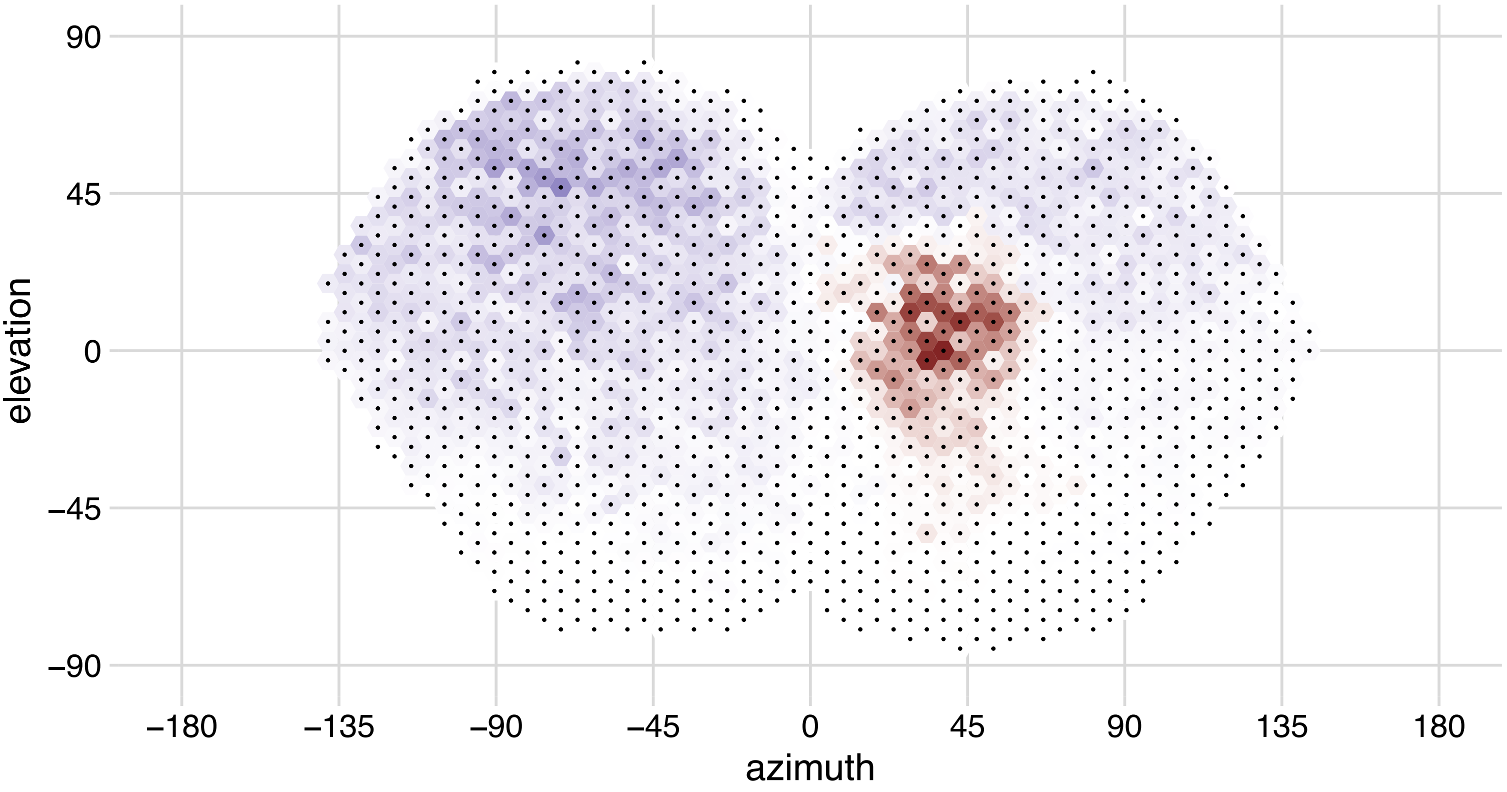

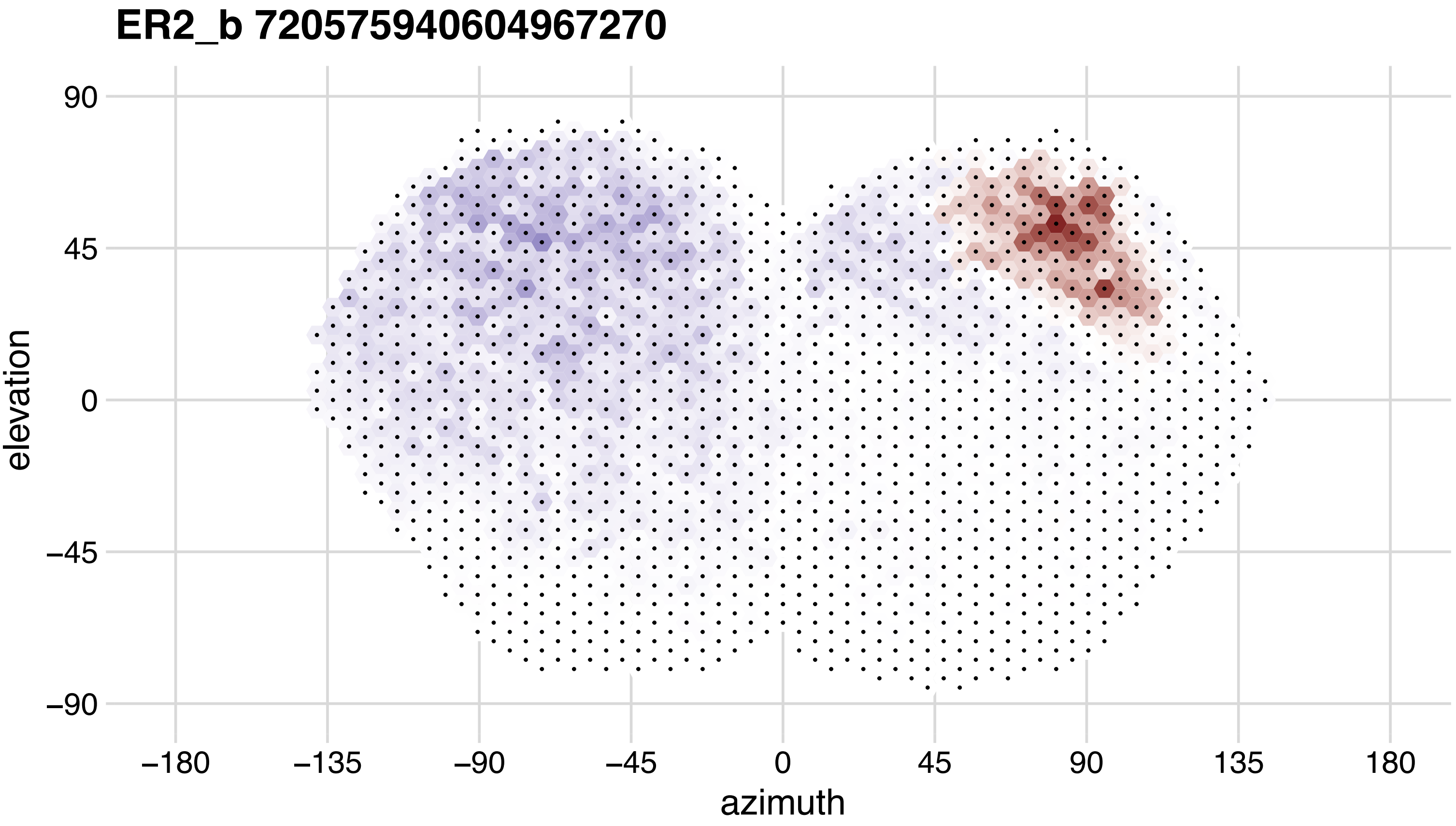
ER2_b 720575940604967270
90
45
0
−45
−90
elevation
−180
−135
−90
−45
0
45
90
135
180
azimuth

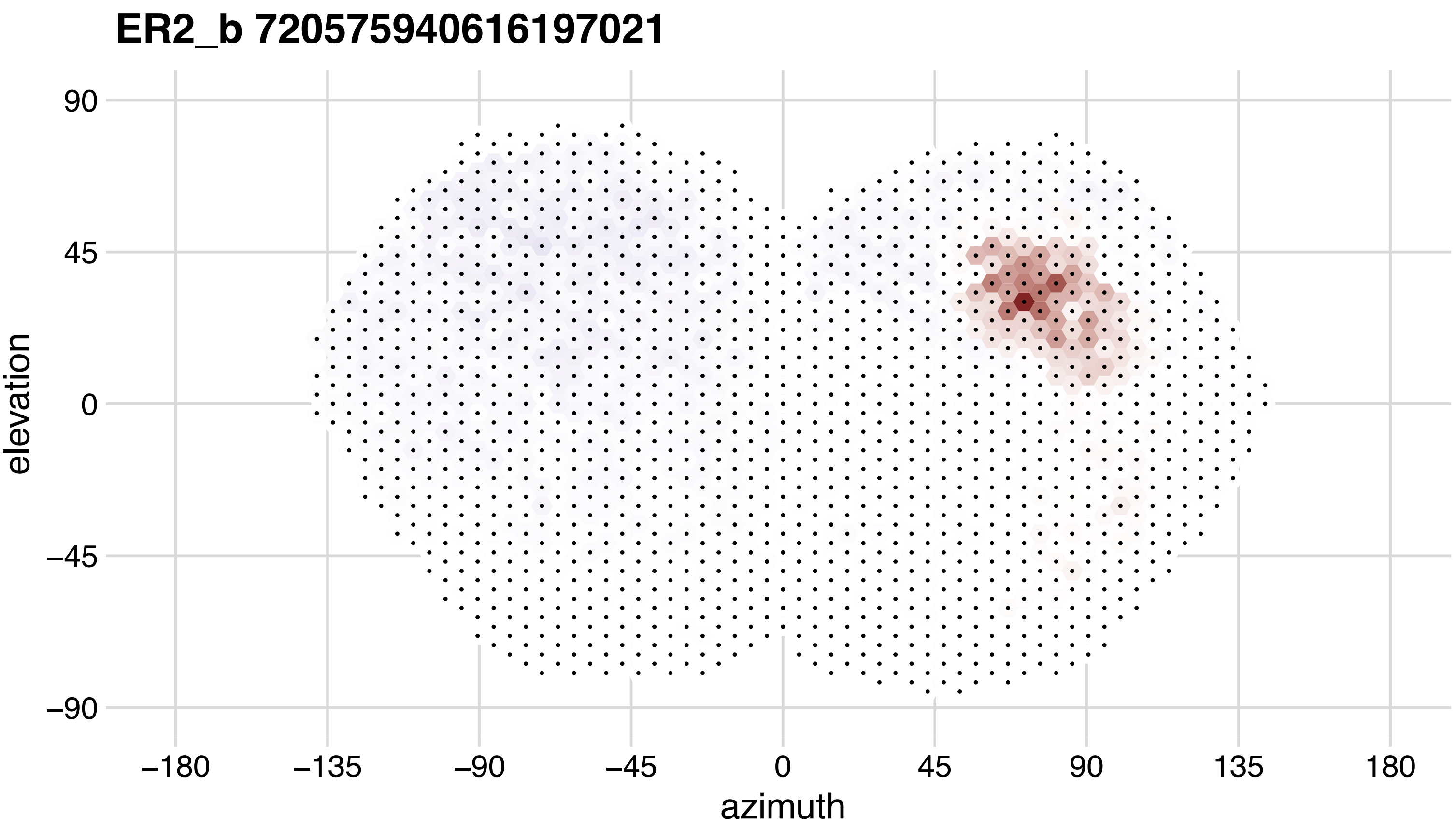
ER2_b 720575940616197021
90
45
0
−45
−90
elevation
−180
−135
−90
−45
0
45
90
135
180
azimuth

## ER2_b 720575940625366846

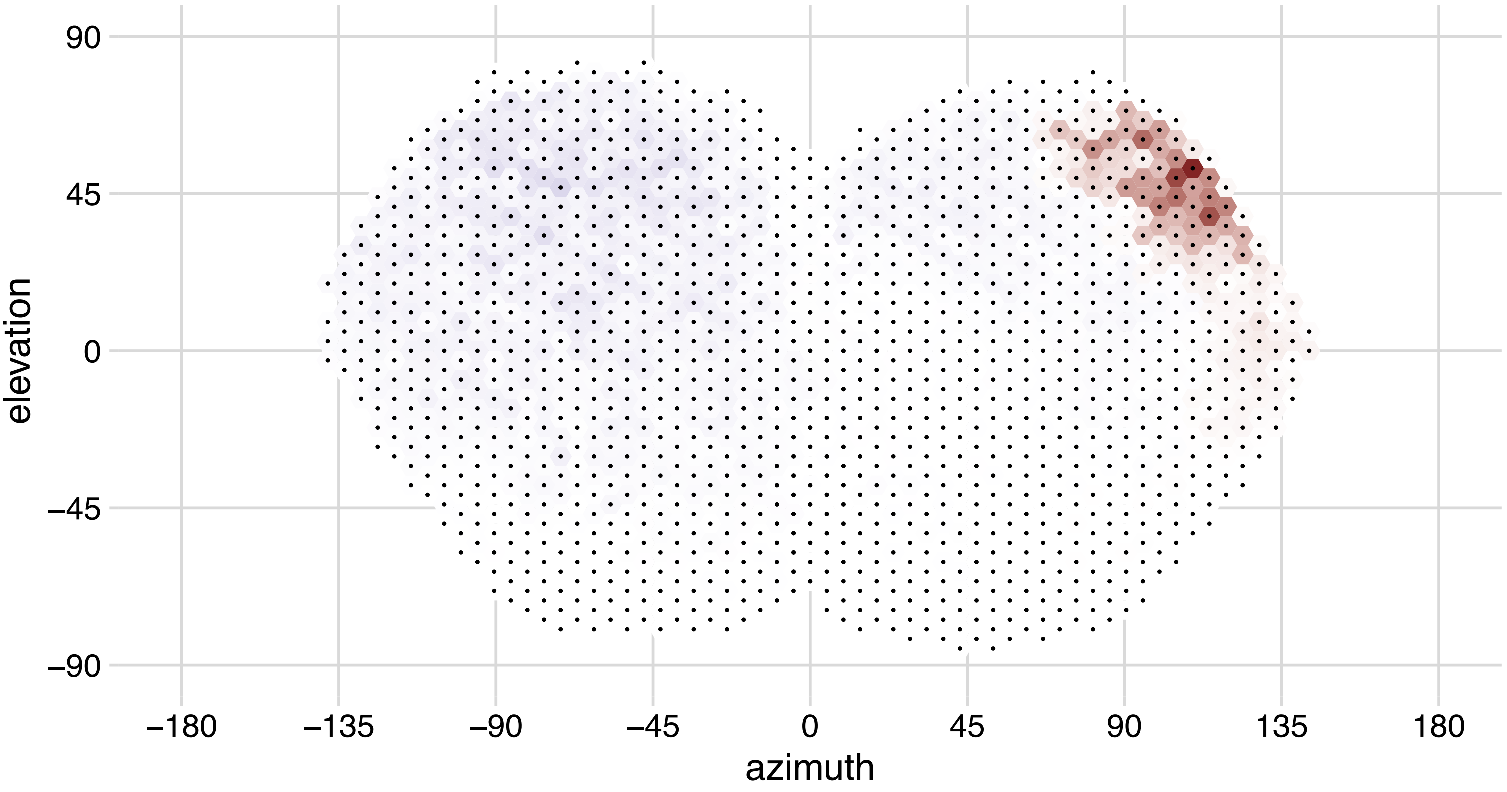

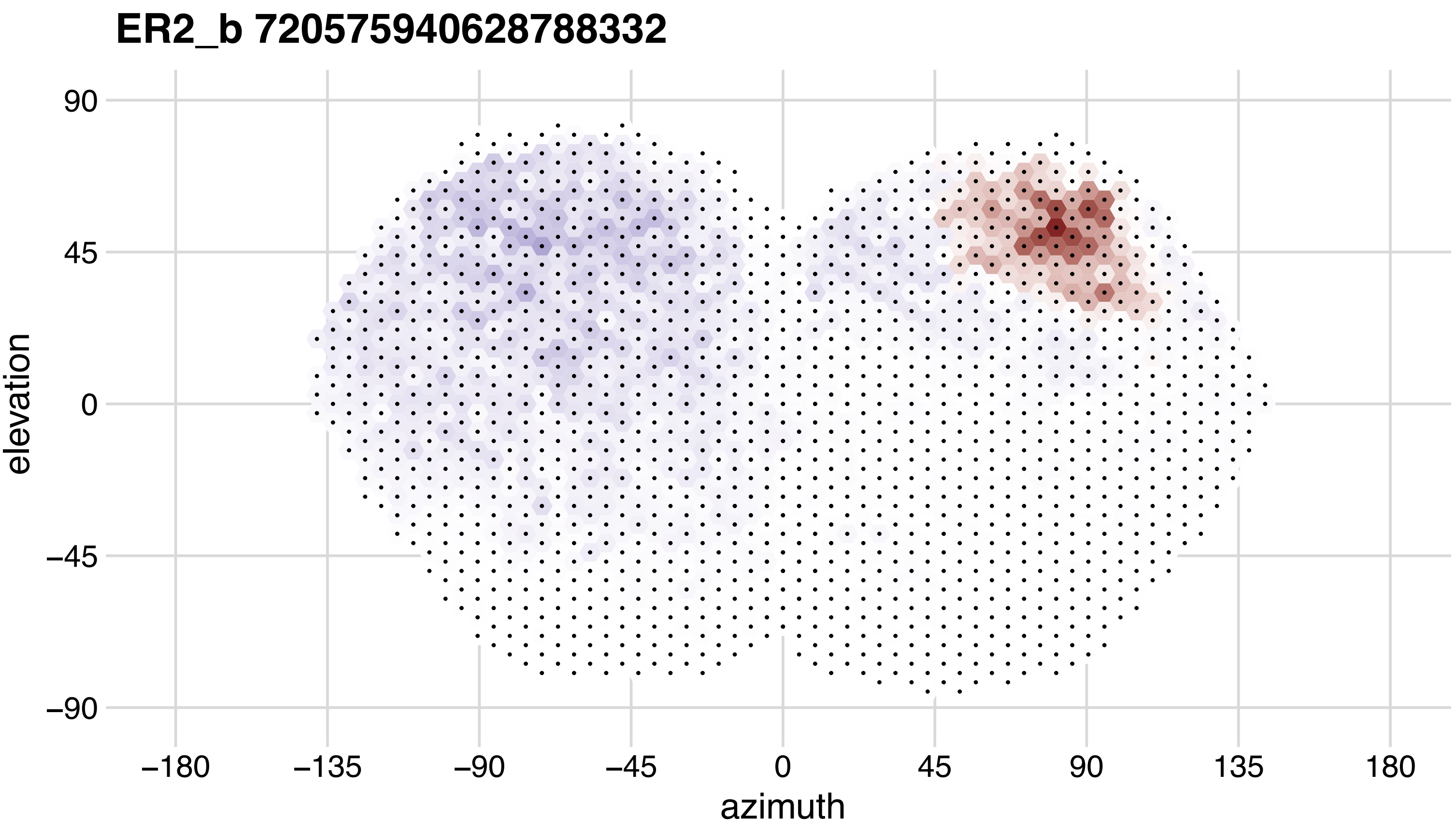
ER2_b 720575940628788332
90
45
0
−45
−90
elevation
−180
−135
−90
−45
0
45
90
135
180
azimuth

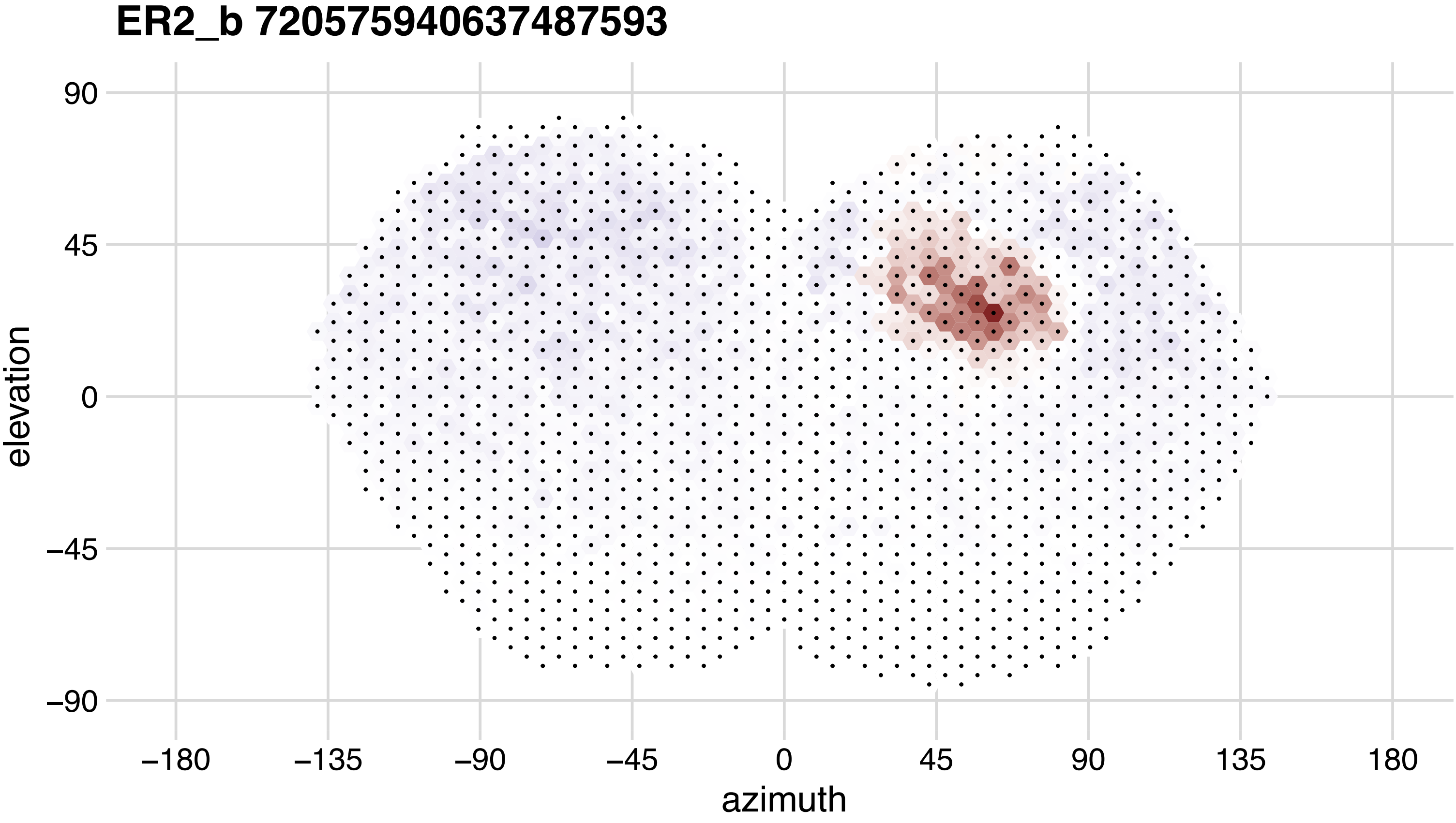
ER2_b 720575940637487593
90
45
0
−45
−90
elevation
−180
−135
−90
−45
0
45
90
135
180
azimuth

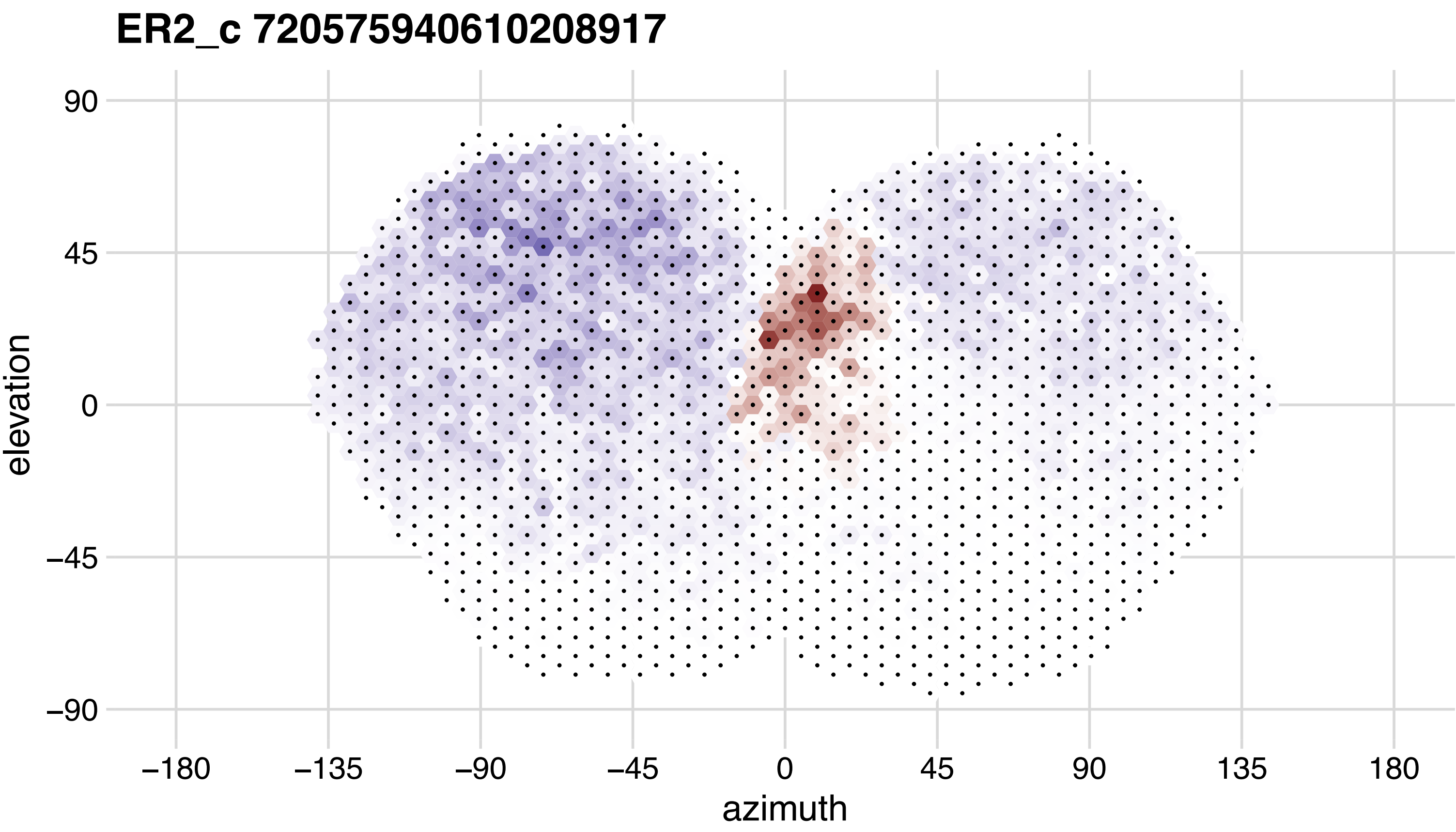
ER2_c 720575940610208917
90
45
0
−45
−90
elevation
−180
−135
−90
−45
0
45
90
135
180
azimuth

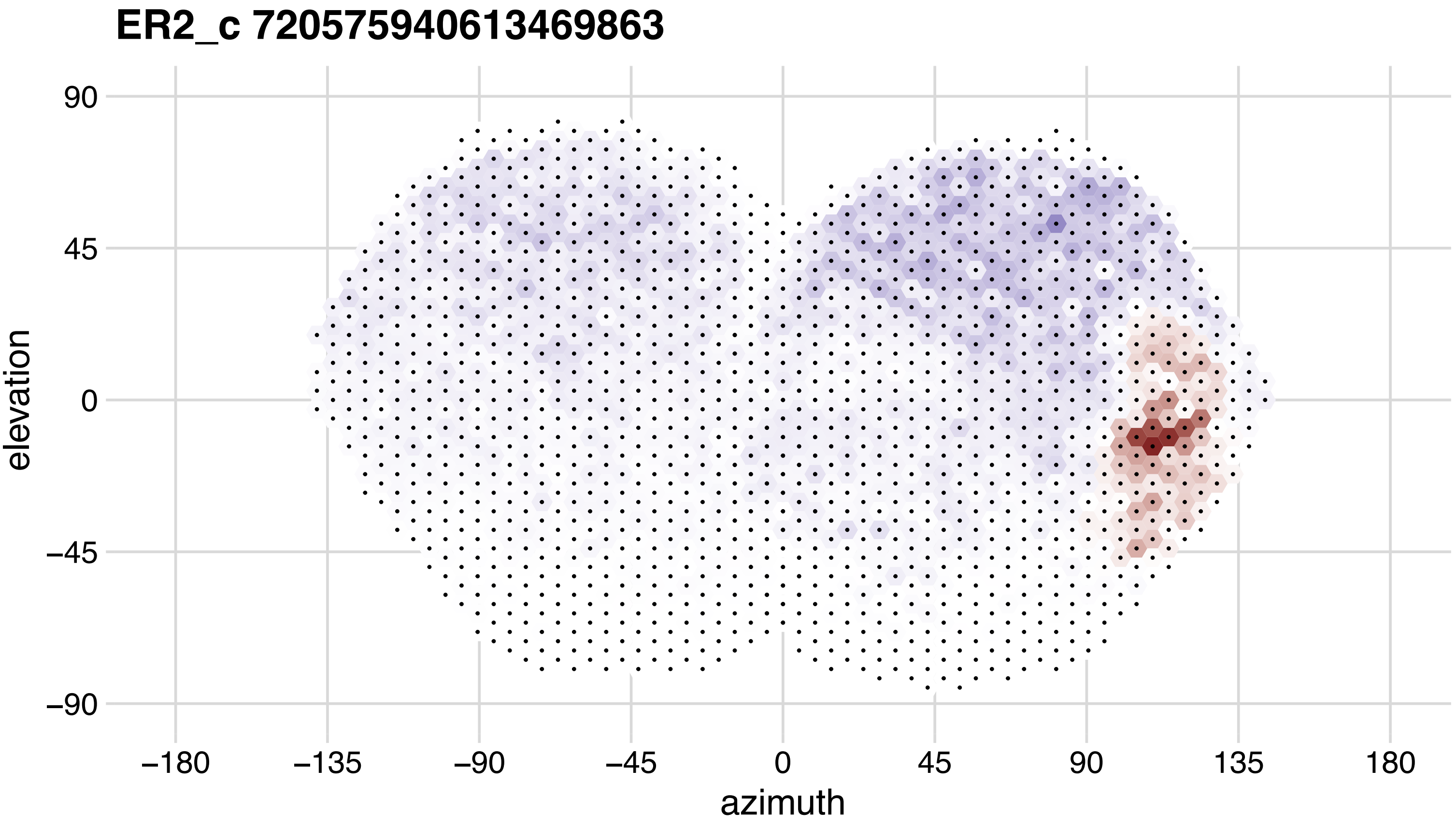
ER2_c 720575940613469863
elevation
azimuth
90
45
0
−45
−90
−180
−135
−90
−45
0
45
90
135
180

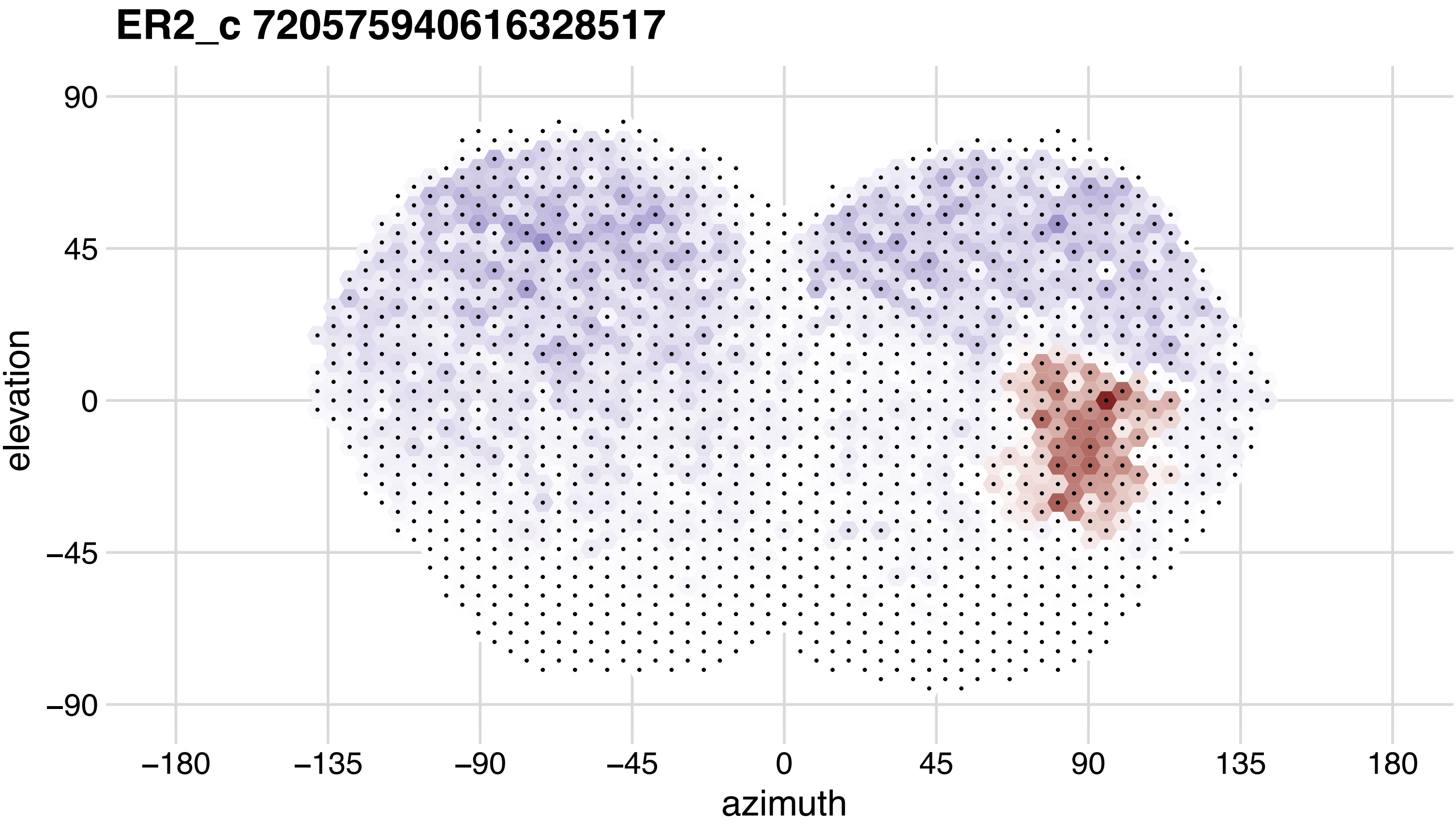
ER2_c 720575940616328517
90
45
0
−45
−90
elevation
−180
−135
−90
−45
0
45
90
135
180
azimuth

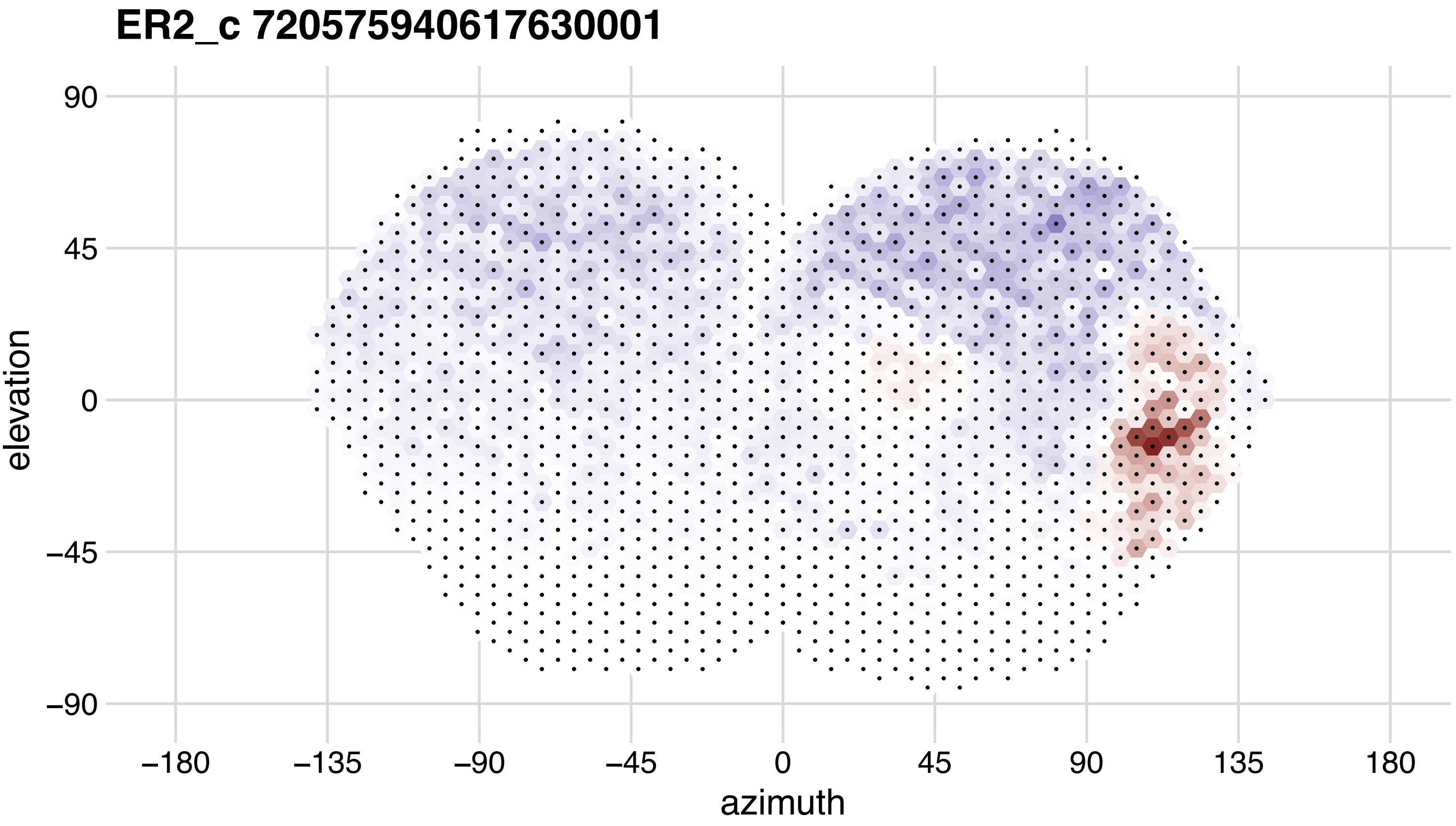

ER2_c 720575940617630001
90
45
0
−45
−90
elevation
−180
−135
−90
−45
0
45
90
135
180
azimuth

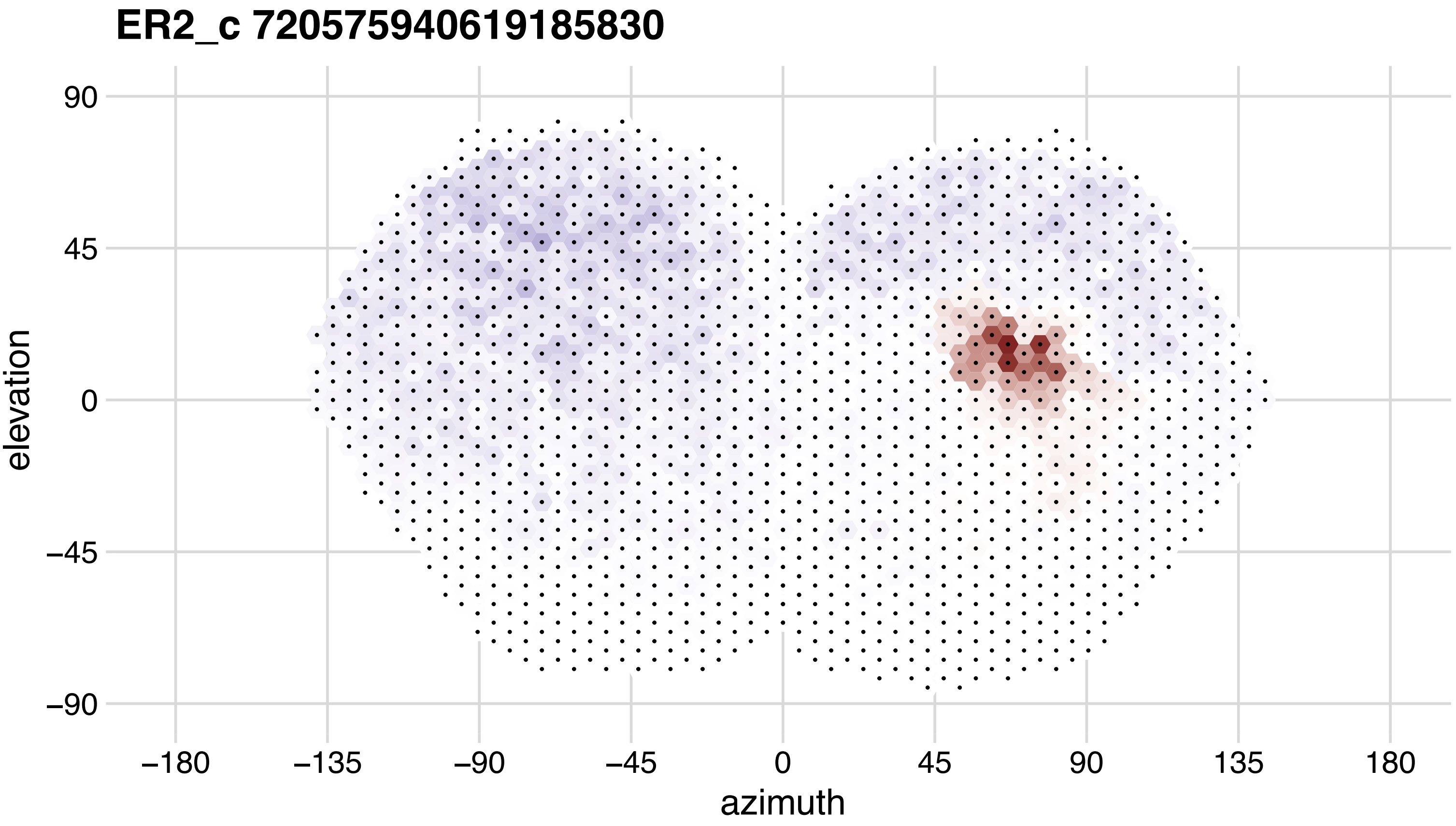

ER2_c 720575940619185830
90
45
0
−45
−90
elevation
−180
−135
−90
−45
0
45
90
135
180
azimuth

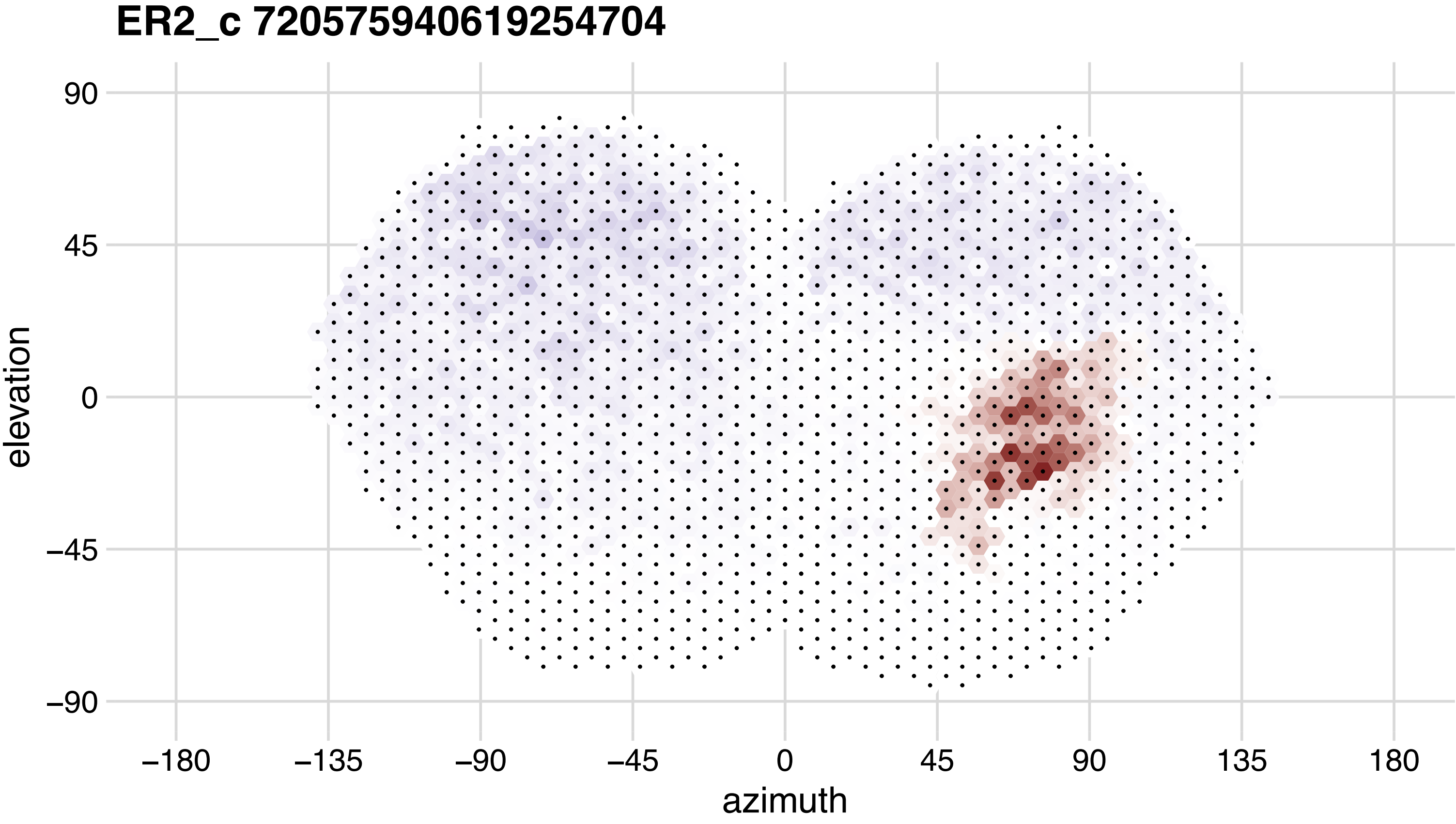
ER2_c 720575940619254704
90
45
0
−45
−90
elevation
−180
−135
−90
−45
0
45
90
135
180
azimuth

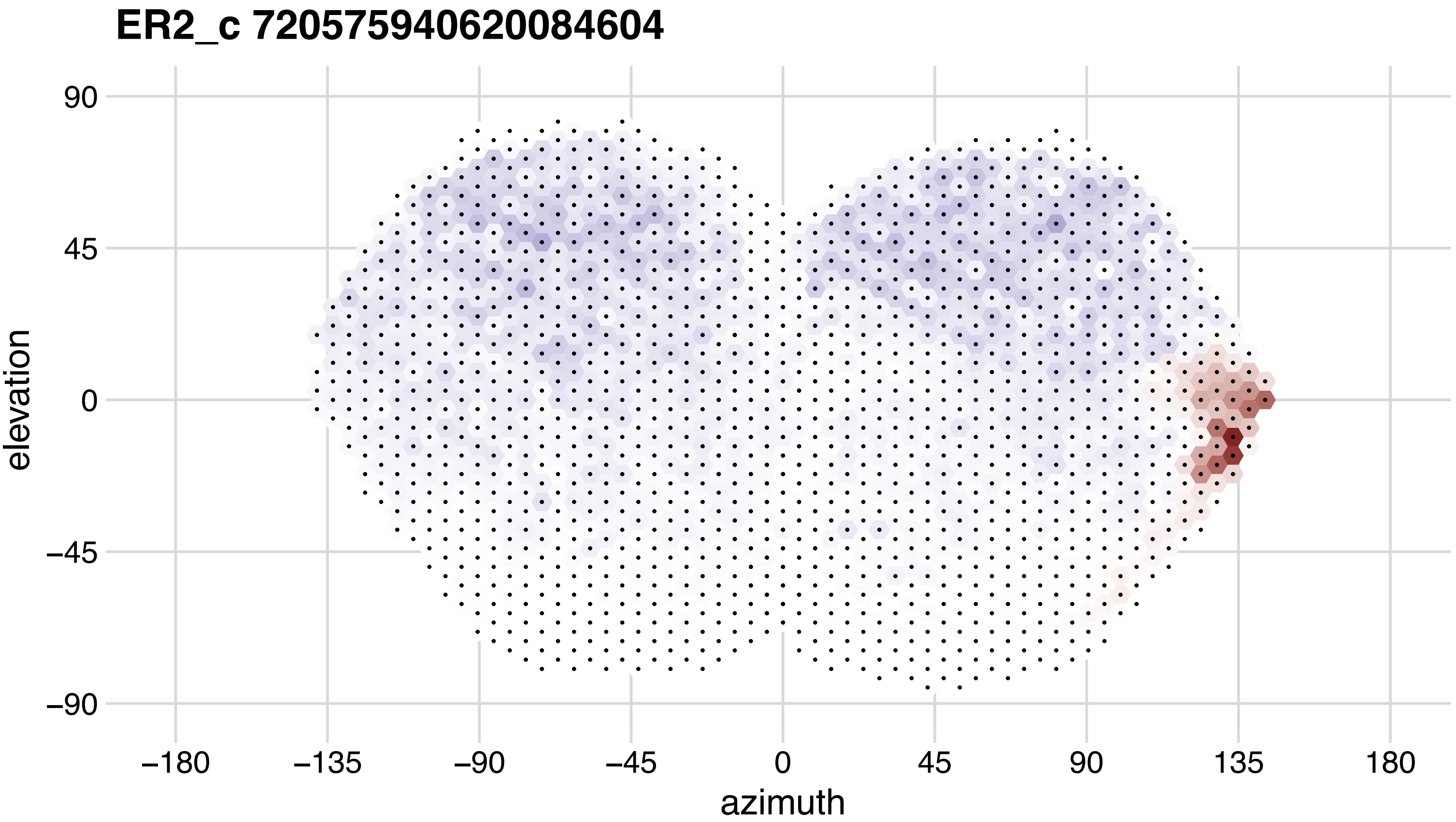
ER2_c 720575940620084604
90
45
0
−45
−90
elevation
−180
−135
−90
−45
0
45
90
135
180
azimuth

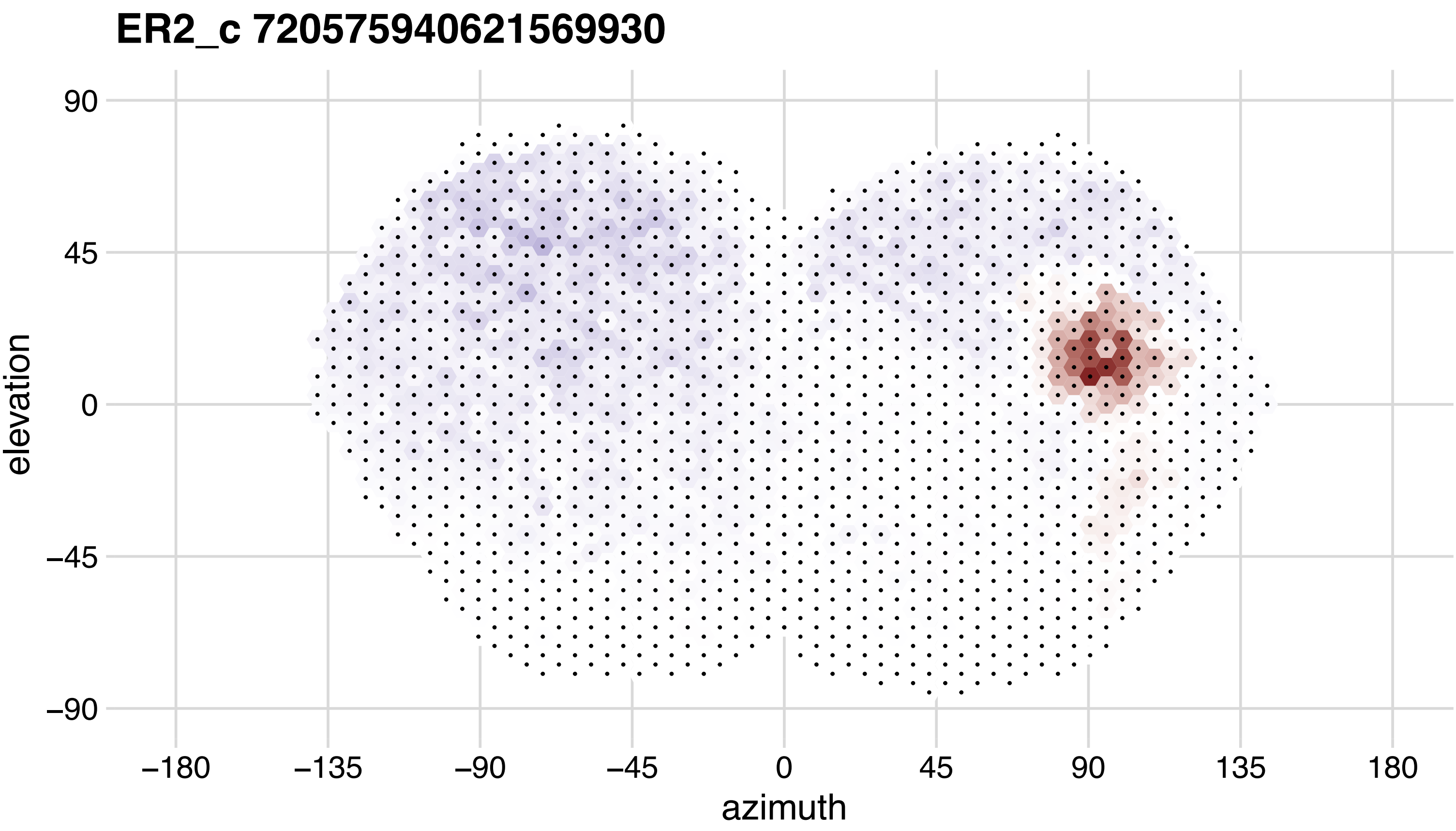
ER2_c 720575940621569930
90
45
0
−45
−90
elevation
−180
−135
−90
−45
0
45
90
135
180
azimuth

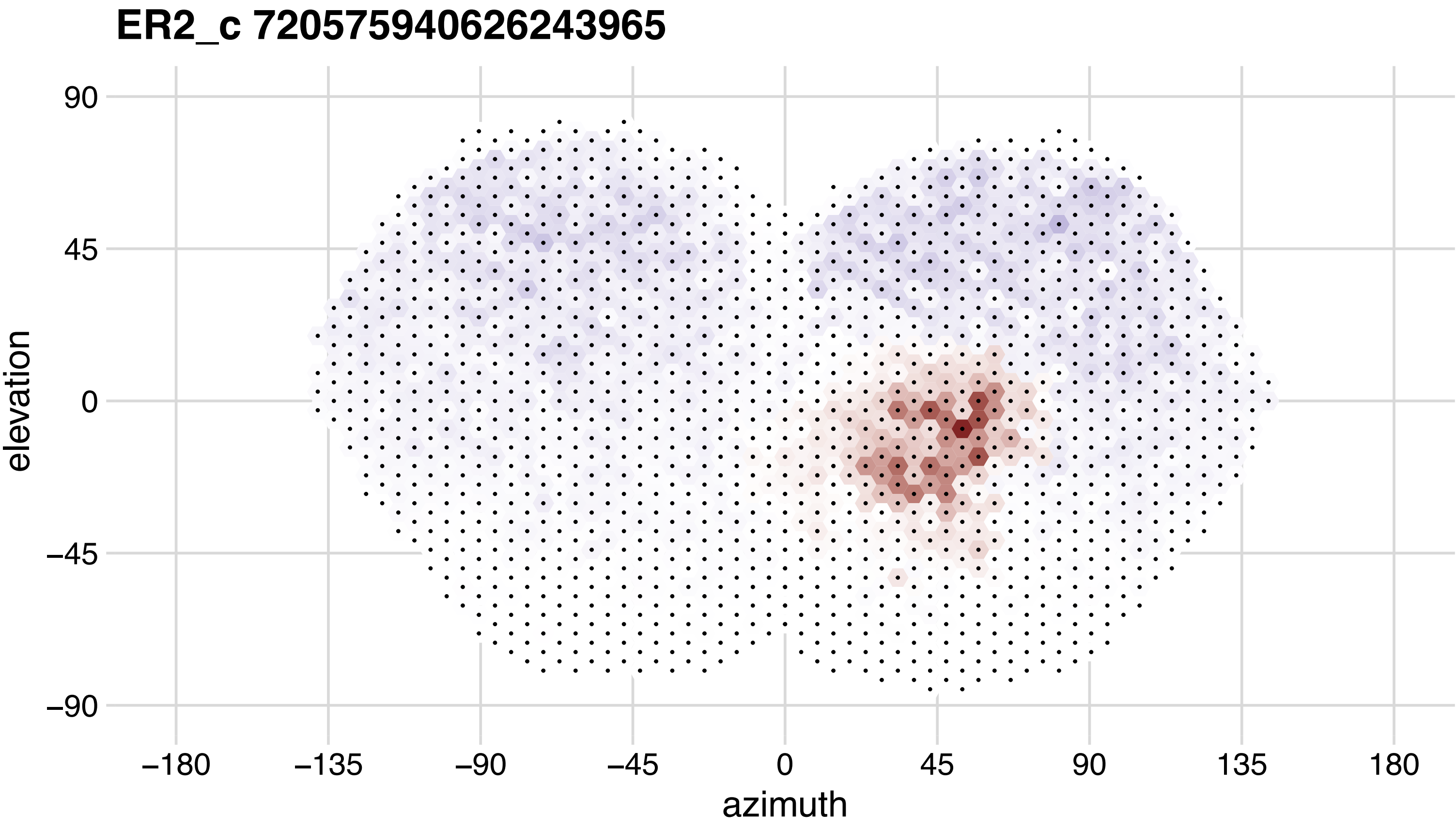
ER2_c 720575940626243965
elevation
90
45
0
−45
−90
azimuth
−180
−135
−90
−45
0
45
90
135
180

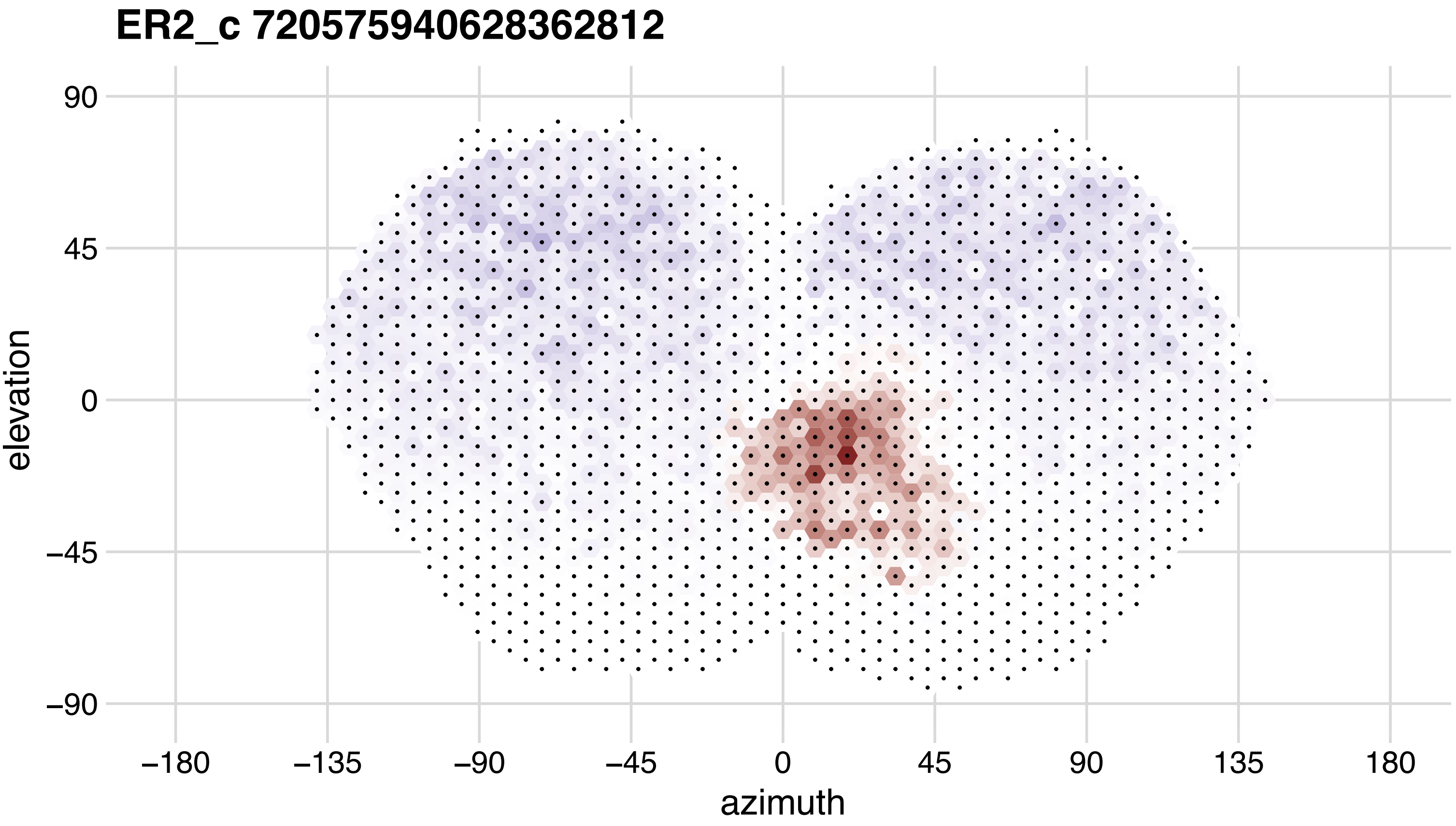
ER2_c 720575940628362812
90
45
0
−45
−90
elevation
−180
−135
−90
−45
0
45
90
135
180
azimuth

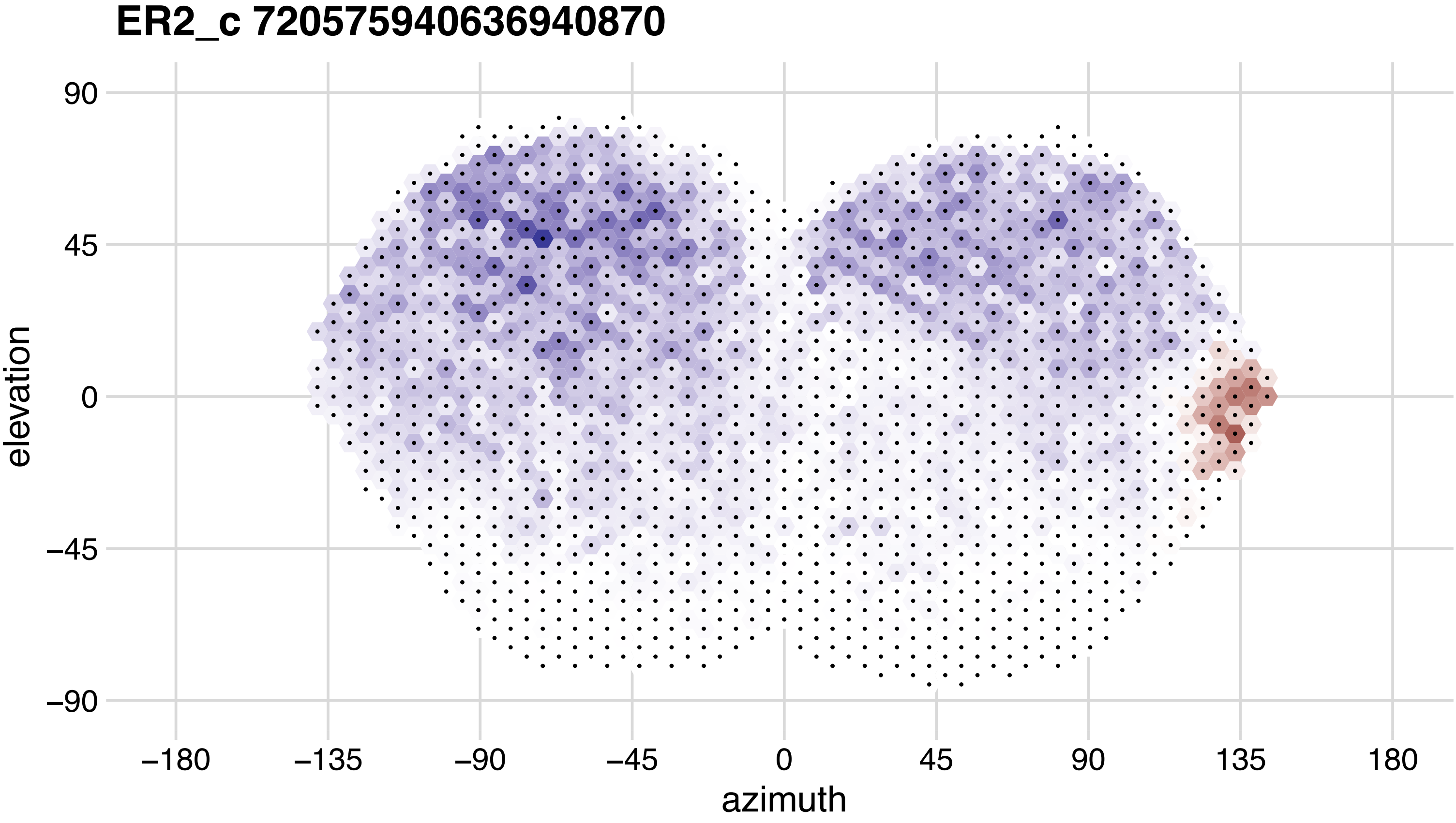
ER2_c 720575940636940870
90
45
0
−45
−90
elevation
−180
−135
−90
−45
0
45
90
135
180
azimuth

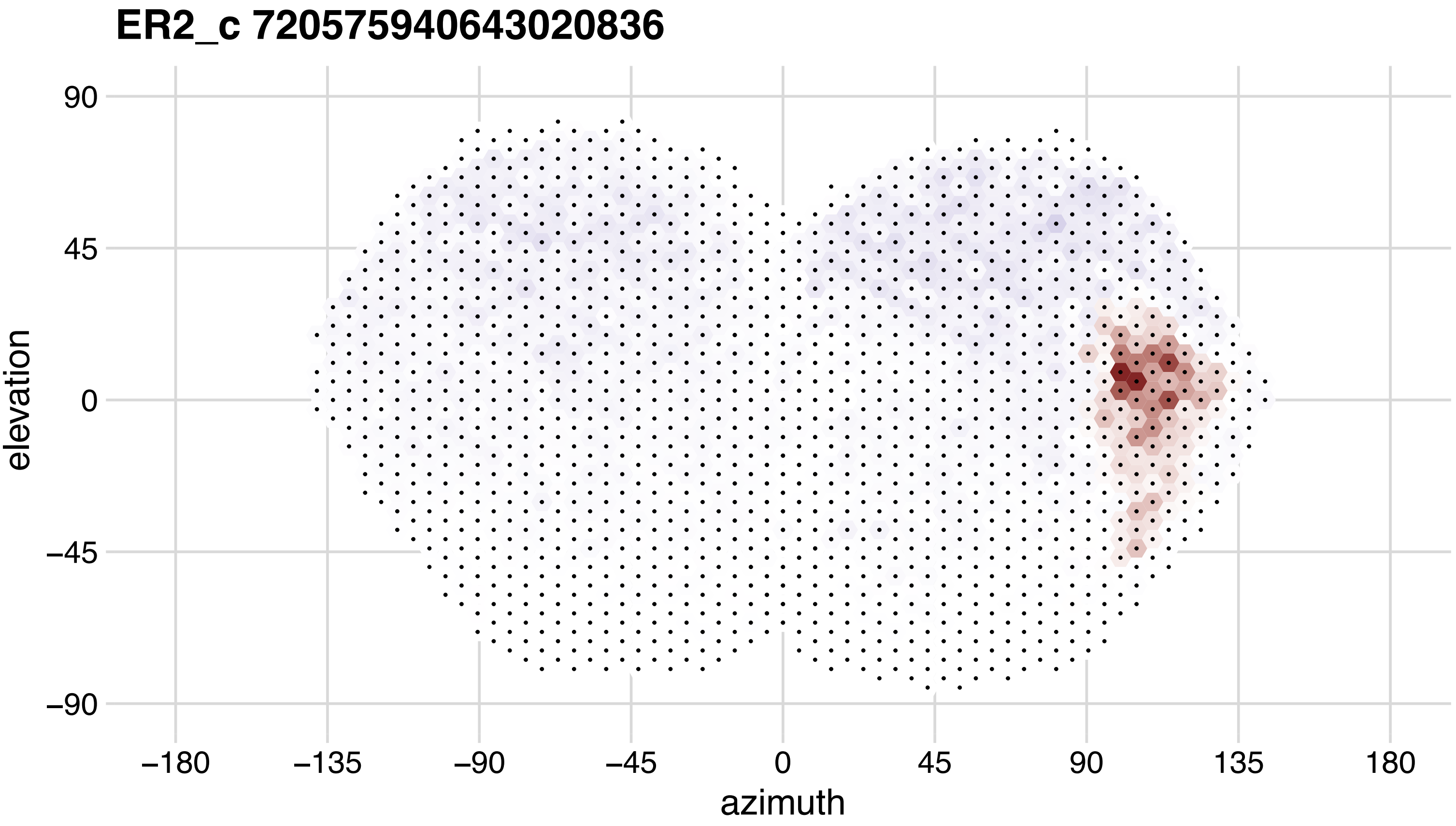

ER2_c 720575940643020836
90
45
0
−45
−90
elevation
−180
−135
−90
−45
0
45
90
135
180
azimuth

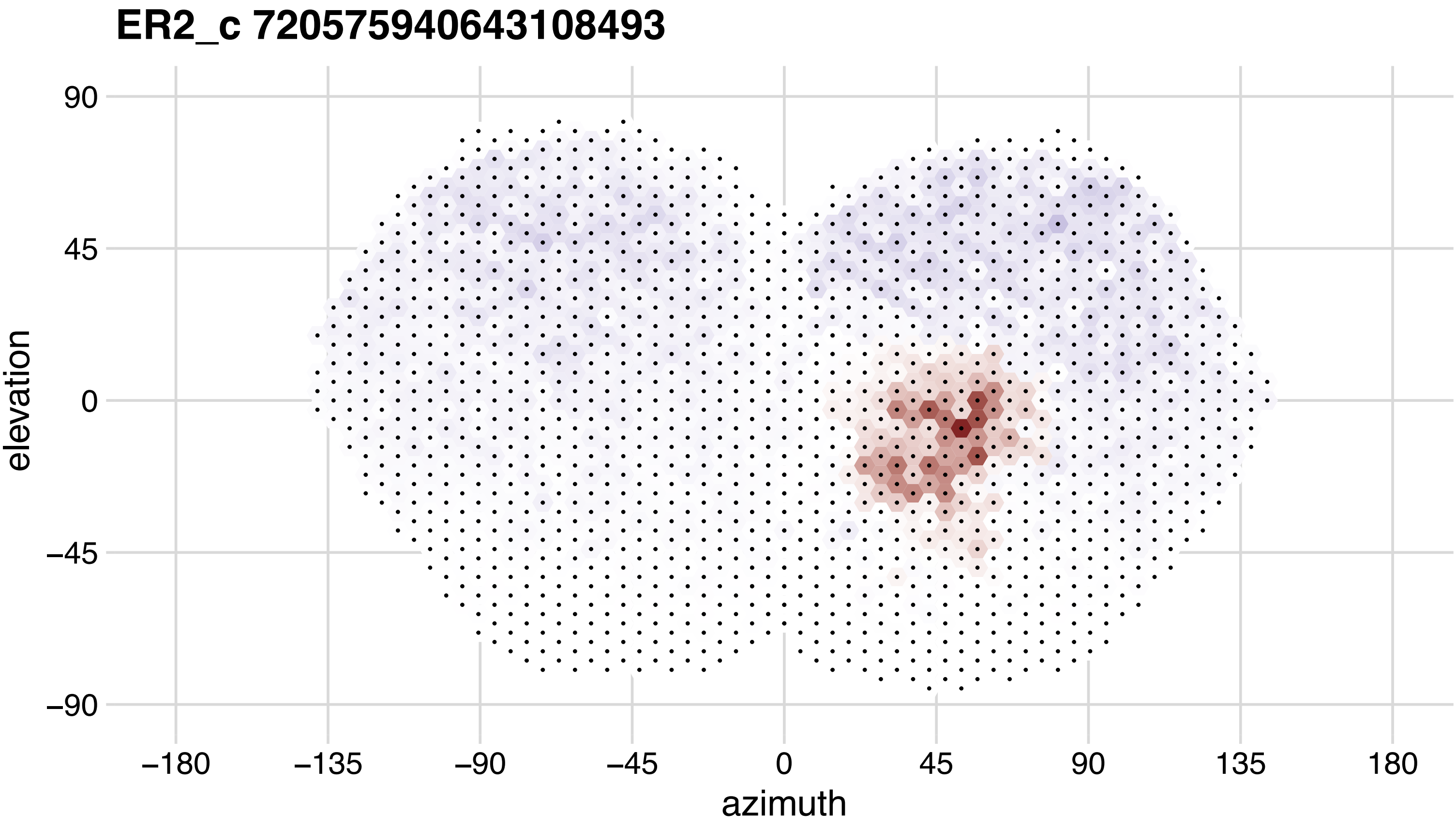
ER2_c 720575940643108493
90
45
0
−45
−90
elevation
−180
−135
−90
−45
0
45
90
135
180
azimuth

## ER3a_ad 720575940607621059

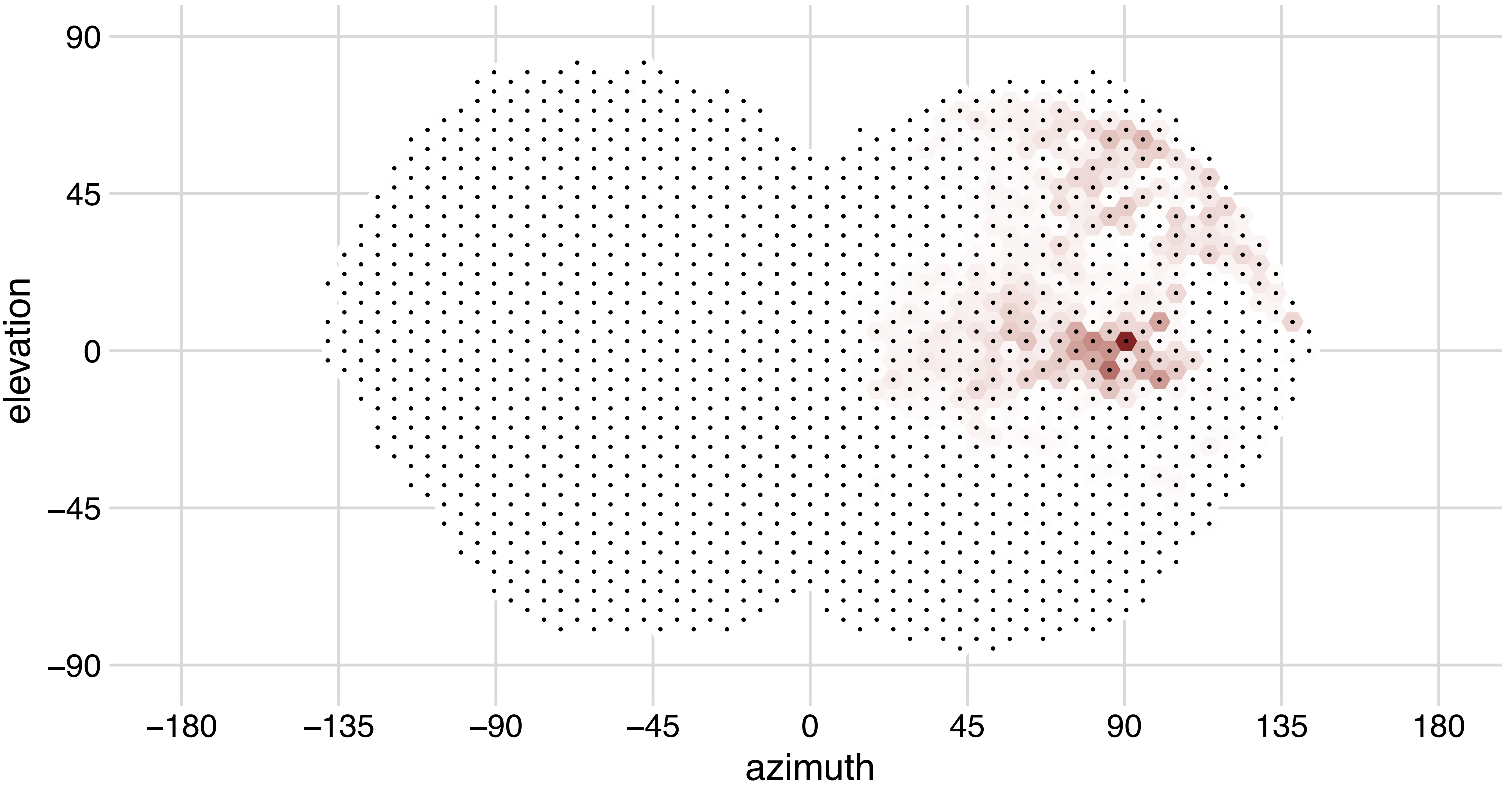

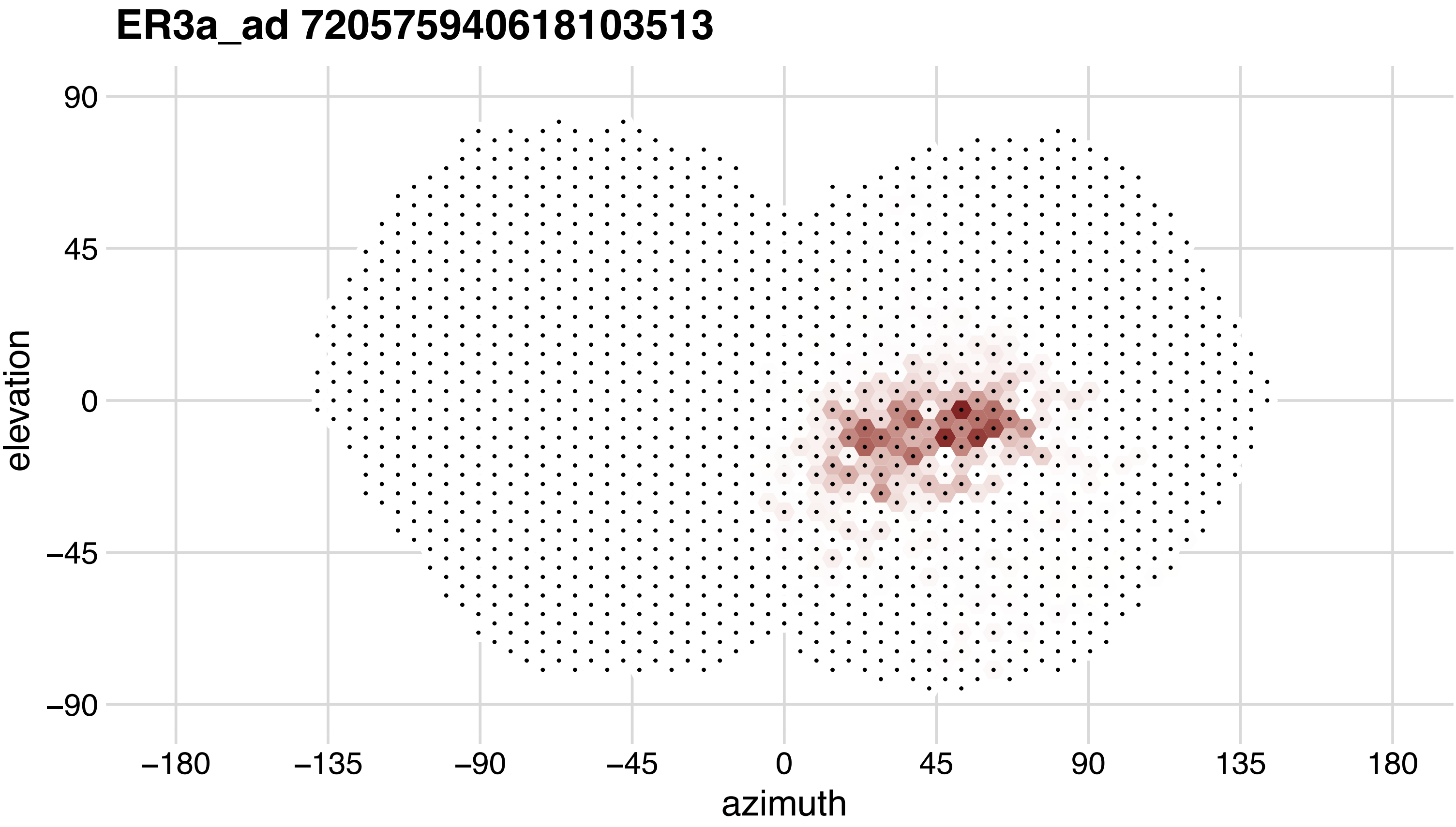
ER3a_ad 720575940618103513
90
45
0
−45
−90
elevation
−180
−135
−90
−45
0
45
90
135
180
azimuth

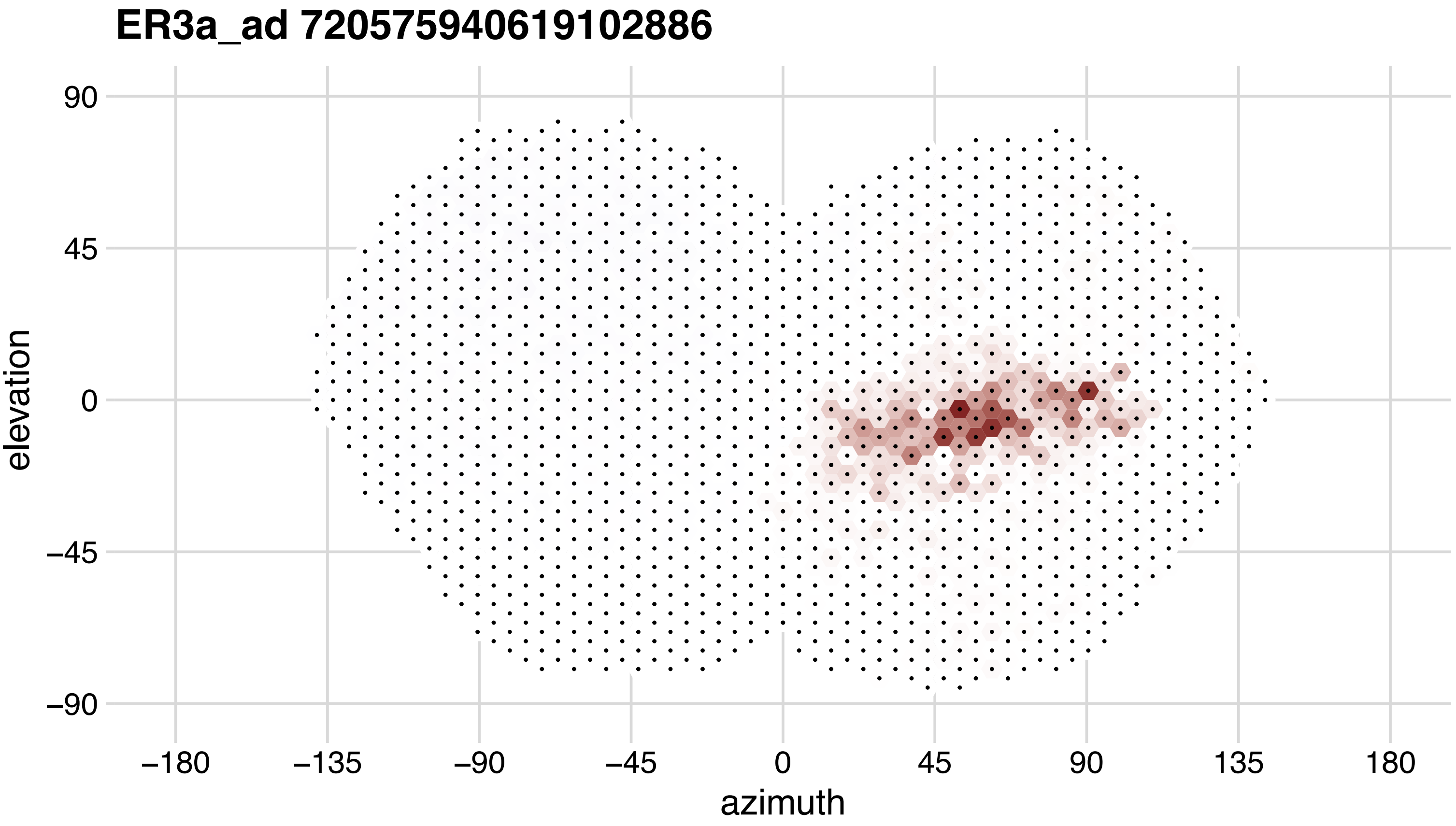
ER3a_ad 720575940619102886
elevation
90
45
0
−45
−90
azimuth
−180
−135
−90
−45
0
45
90
135
180

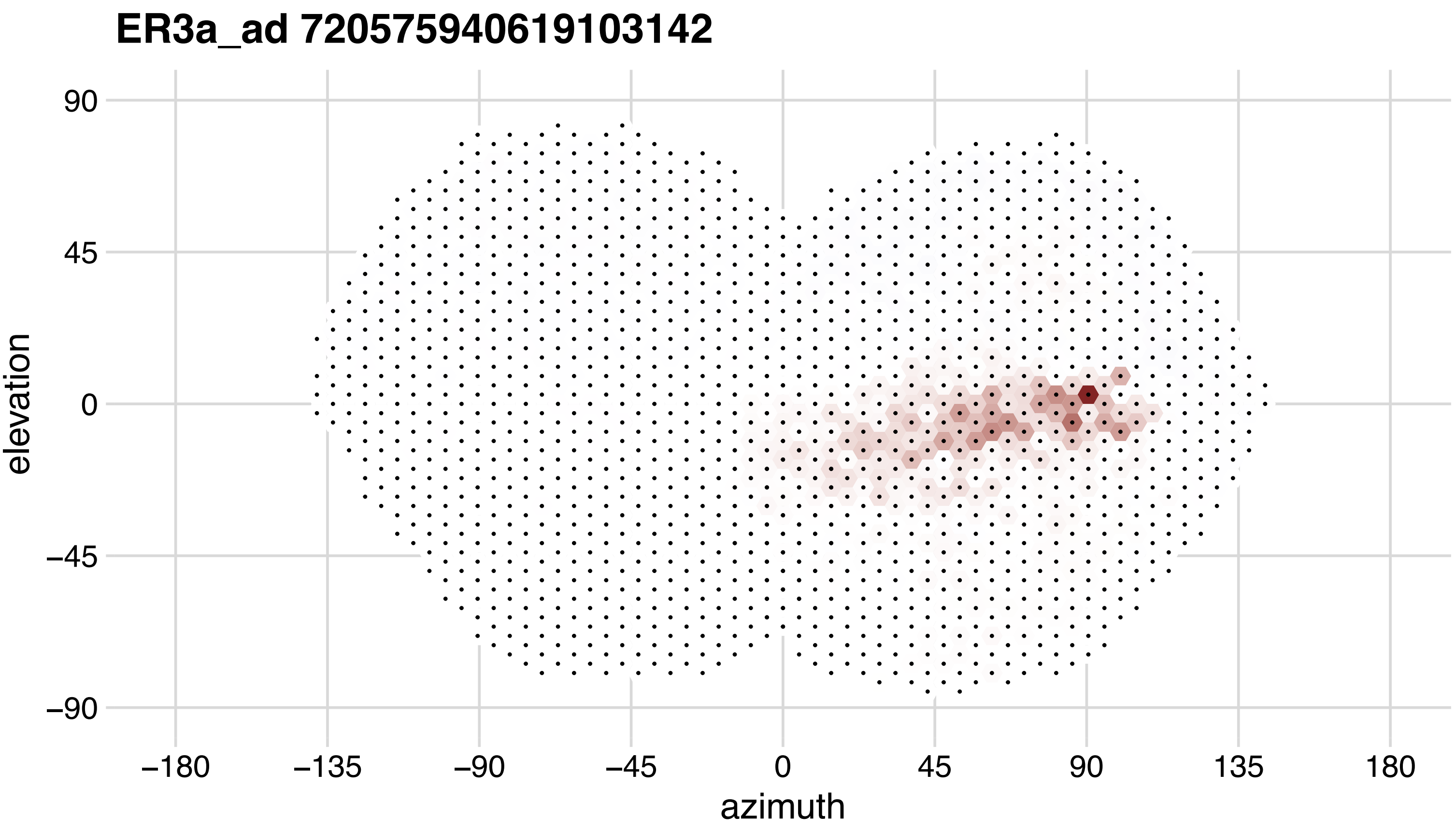

ER3a_ad 720575940619103142
90
45
0
−45
−90
elevation
−180
−135
−90
−45
0
45
90
135
180
azimuth

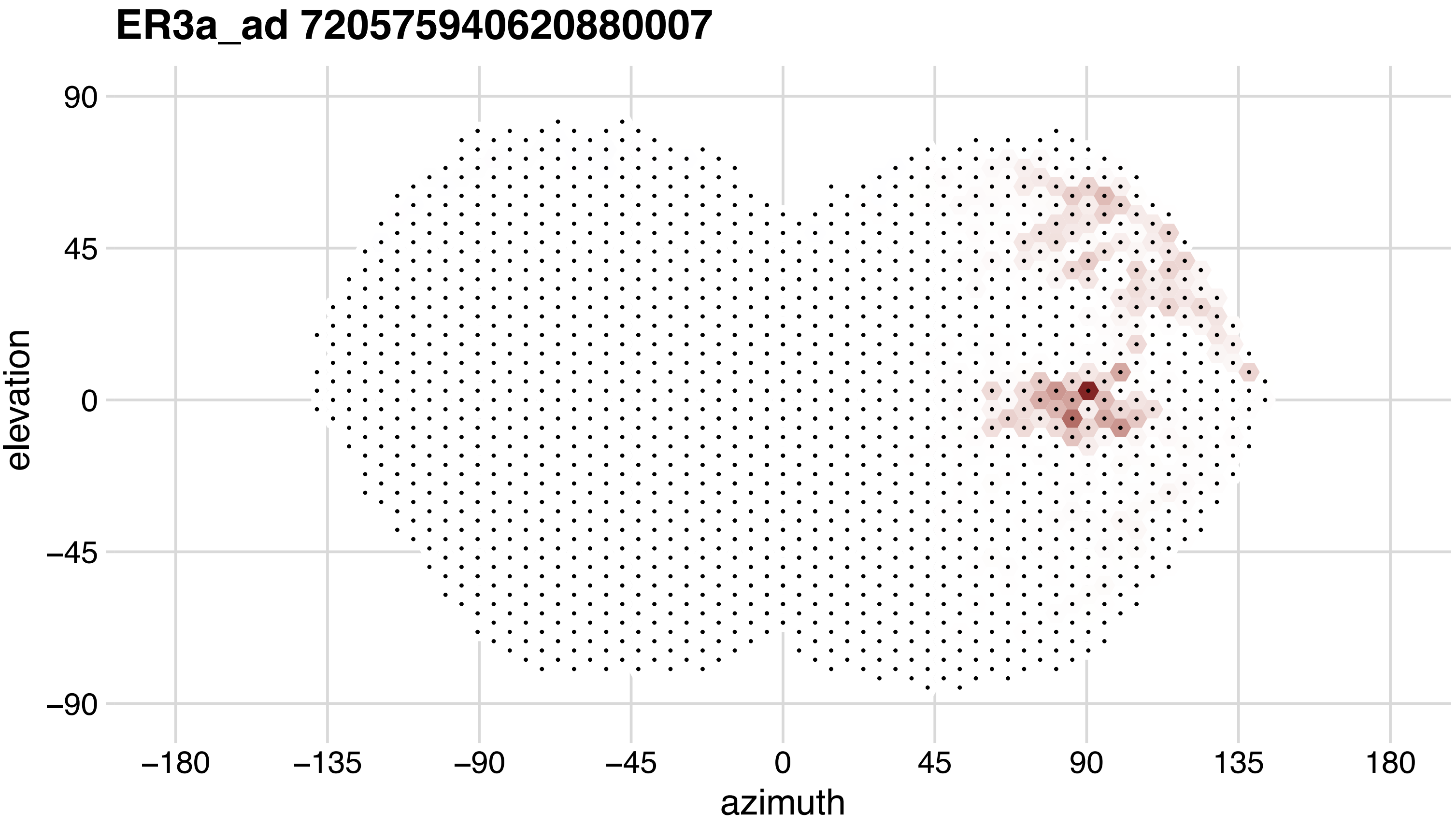
ER3a_ad 720575940620880007
90
45
0
−45
−90
elevation
−180
−135
−90
−45
0
45
90
135
180
azimuth

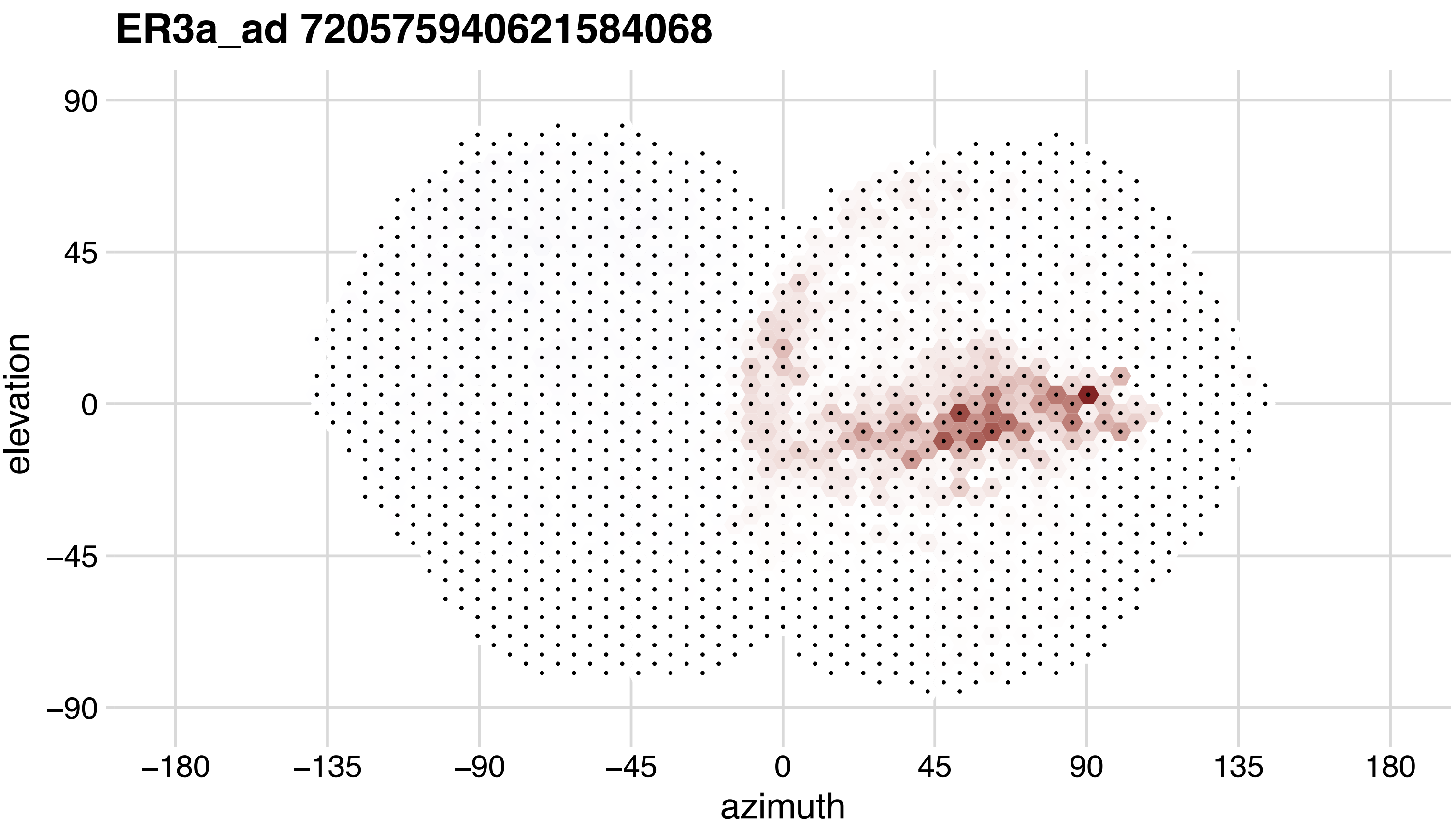

ER3a_ad 720575940621584068
elevation
90
45
0
−45
−90
−180
−135
−90
−45
0
45
90
135
180
azimuth

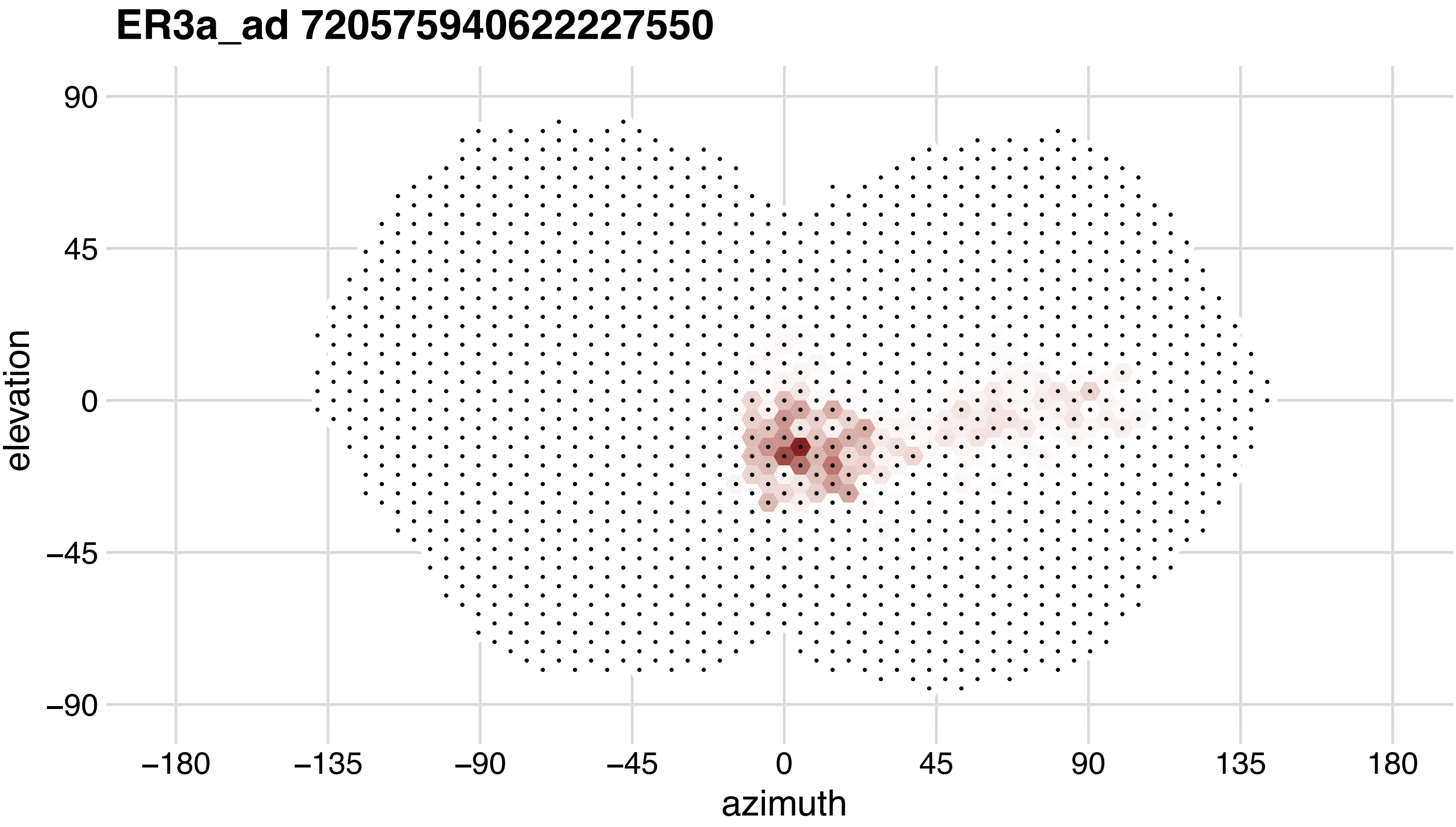
ER3a_ad 720575940622227550
90
45
0
−45
−90
elevation
−180
−135
−90
−45
0
45
90
135
180
azimuth

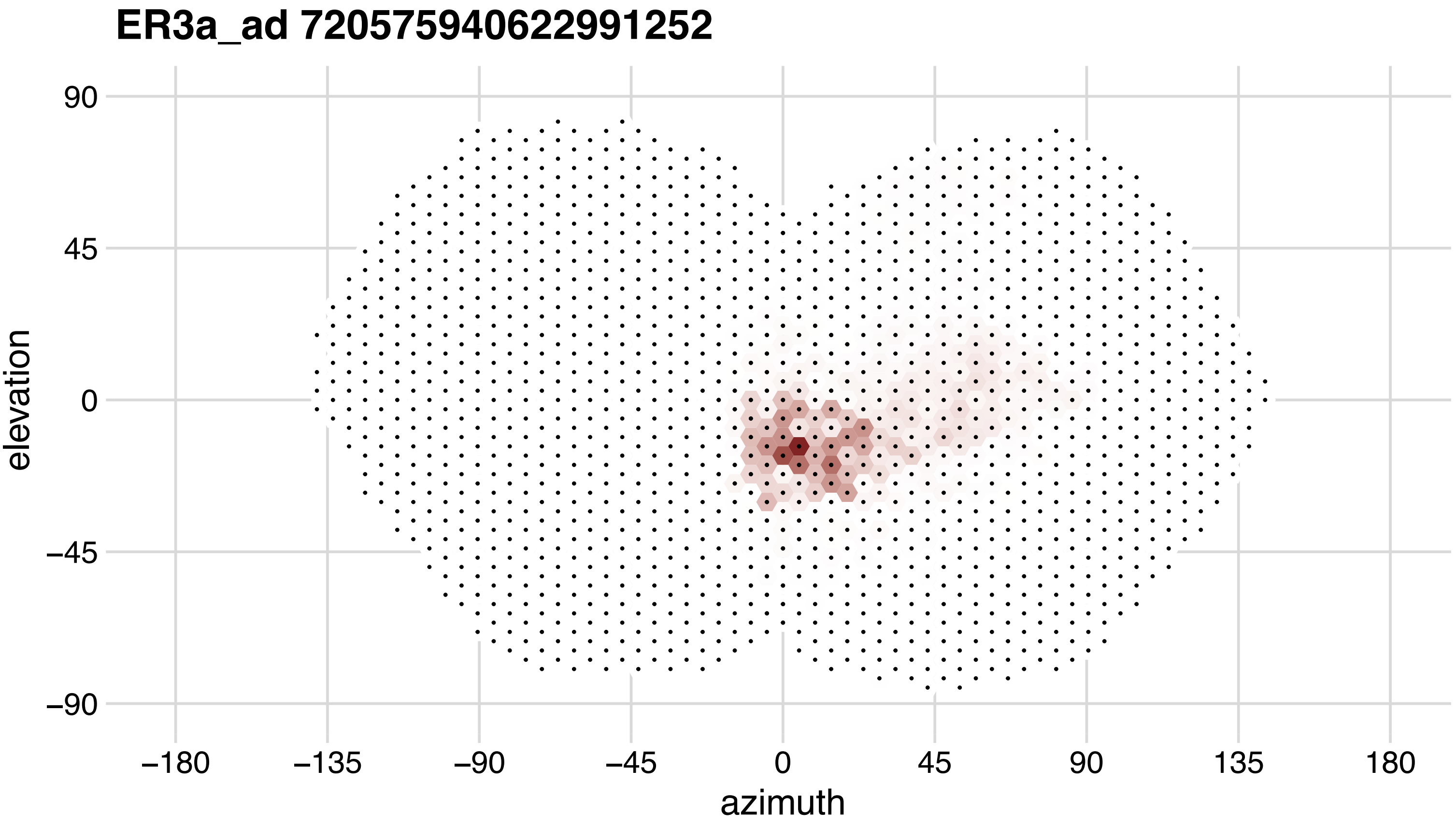
ER3a_ad 720575940622991252
90
45
0
−45
−90
elevation
−180
−135
−90
−45
0
45
90
135
180
azimuth

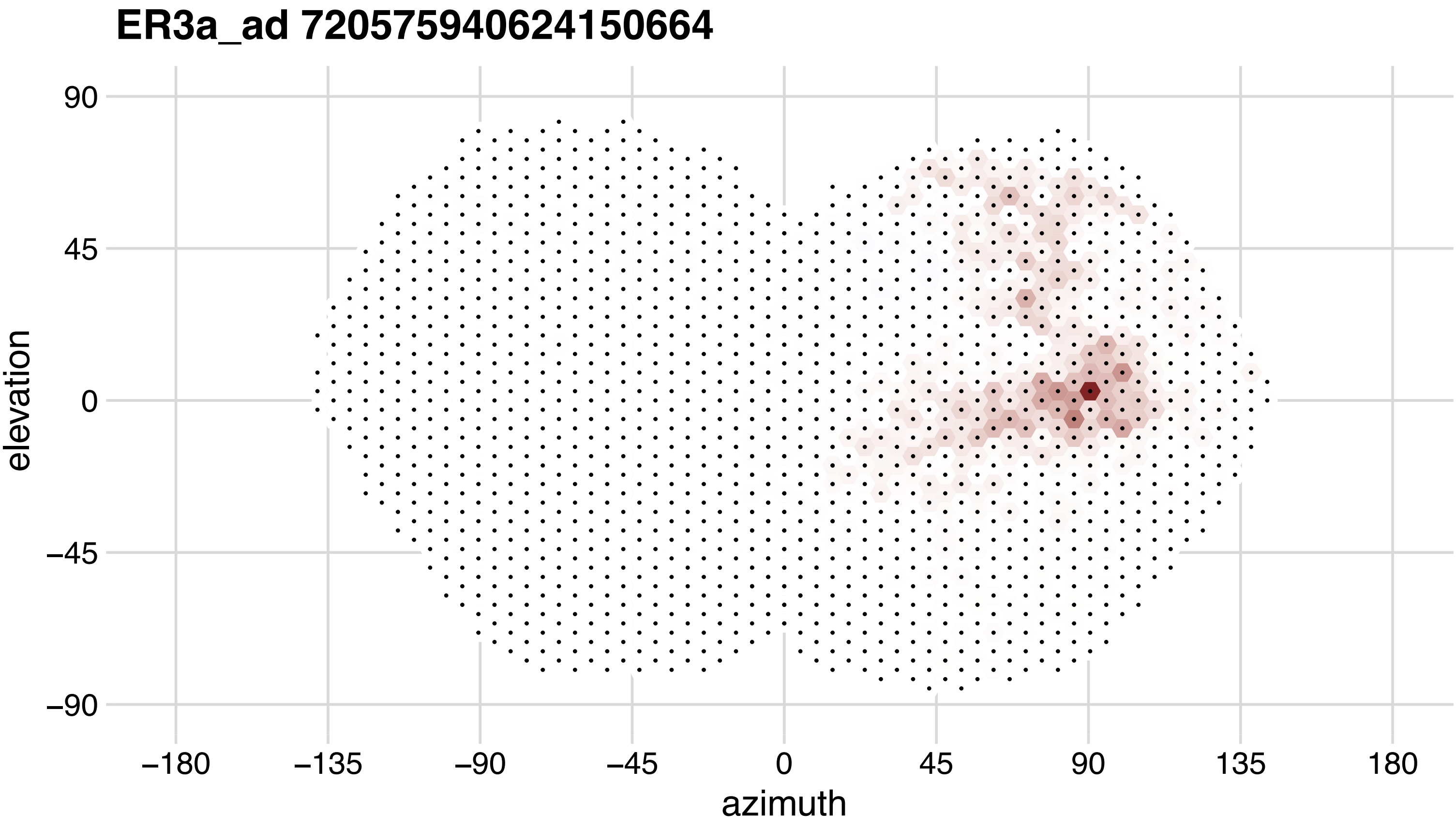
ER3a_ad 720575940624150664
90
45
0
−45
−90
elevation
−180
−135
−90
−45
0
45
90
135
180
azimuth

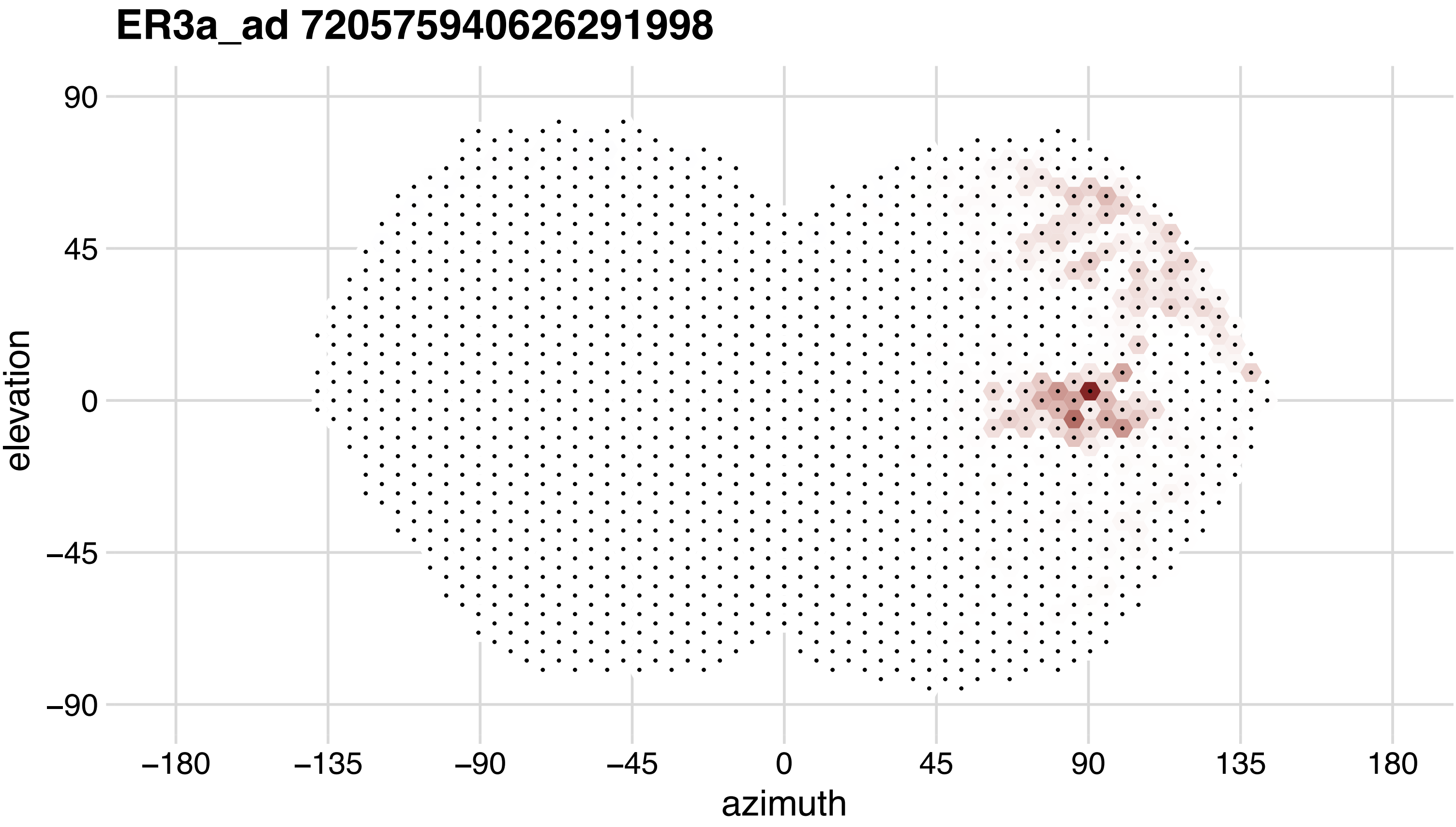
ER3a_ad 720575940626291998
90
45
0
−45
−90
elevation
−180
−135
−90
−45
0
45
90
135
180
azimuth

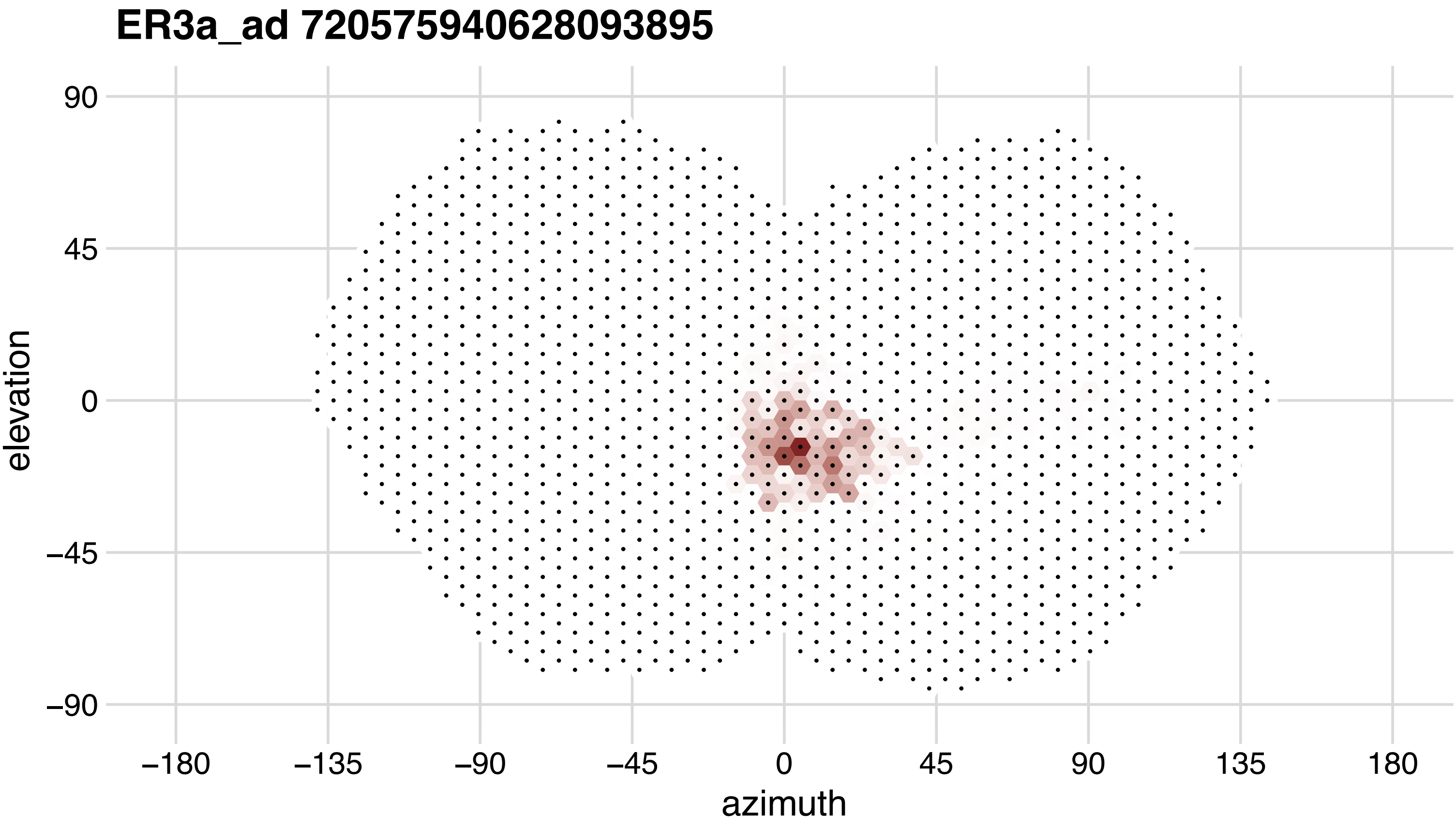
ER3a_ad 720575940628093895
90
45
0
−45
−90
elevation
−180
−135
−90
−45
0
45
90
135
180
azimuth

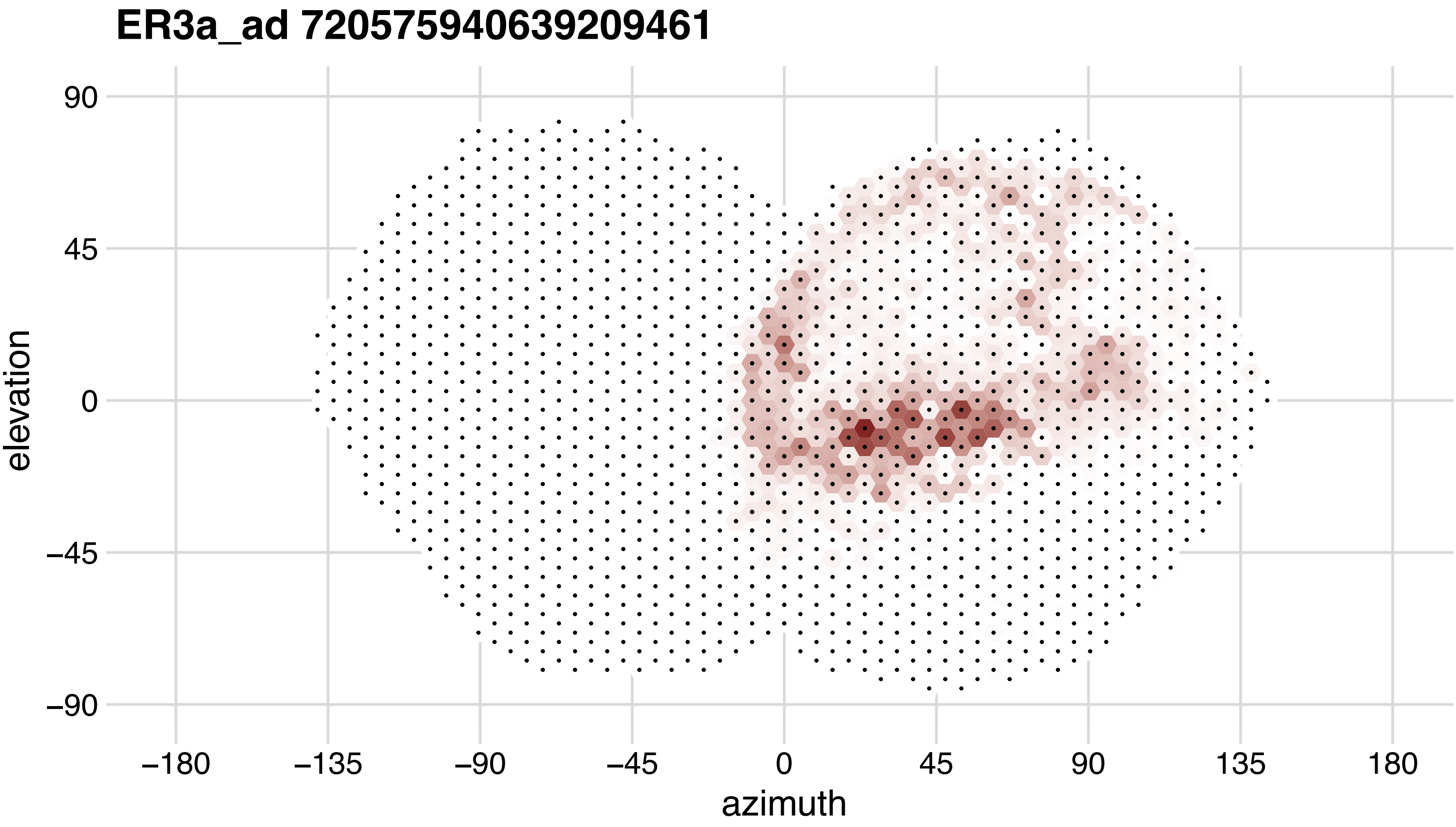
ER3a_ad 720575940639209461
90
45
0
−45
−90
elevation
−180
−135
−90
−45
0
45
90
135
180
azimuth

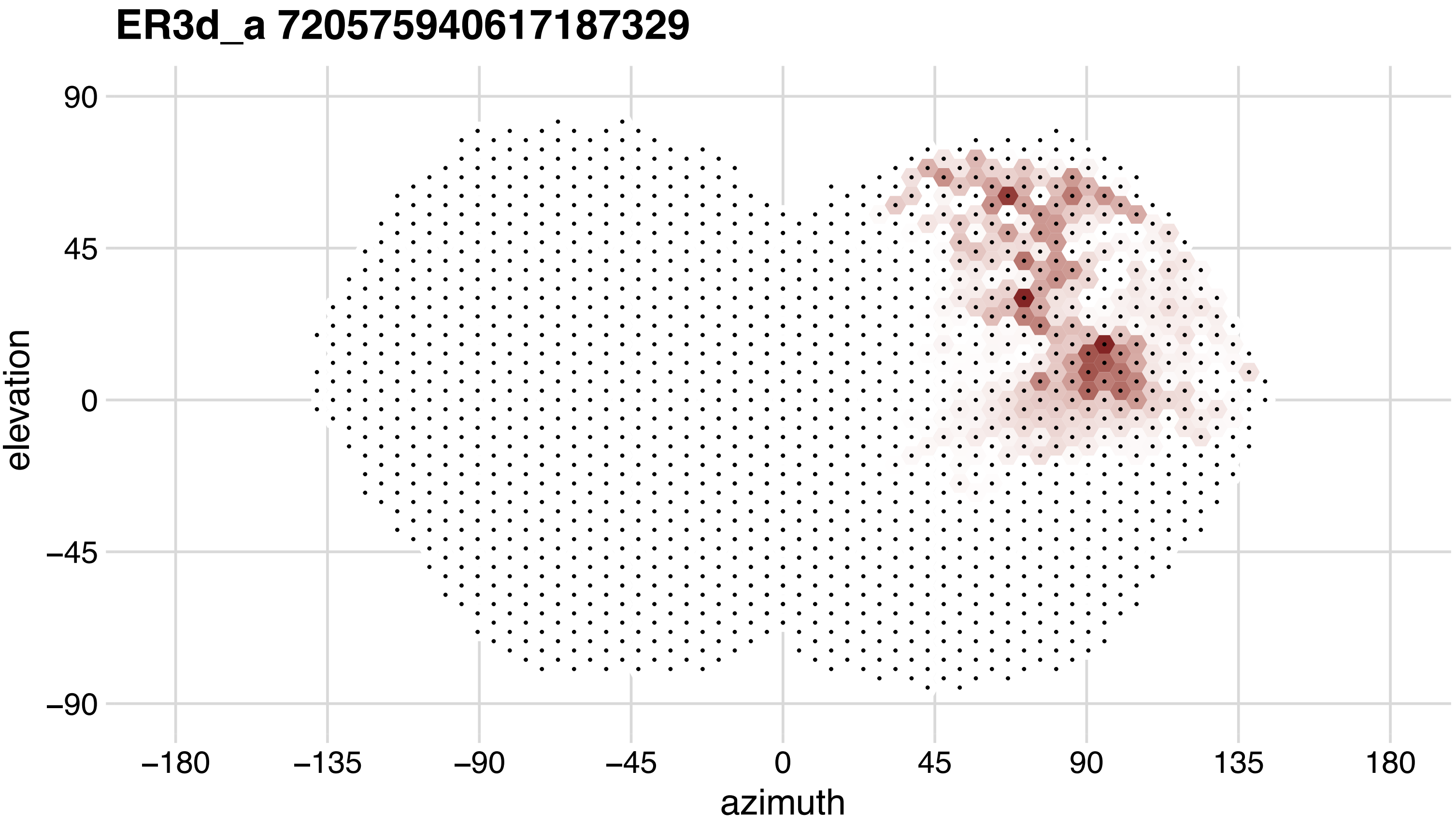
ER3d_a 720575940617187329
90
45
0
−45
−90
elevation
−180
−135
−90
−45
0
45
90
135
180
azimuth

# ER3d_a 720575940619767744

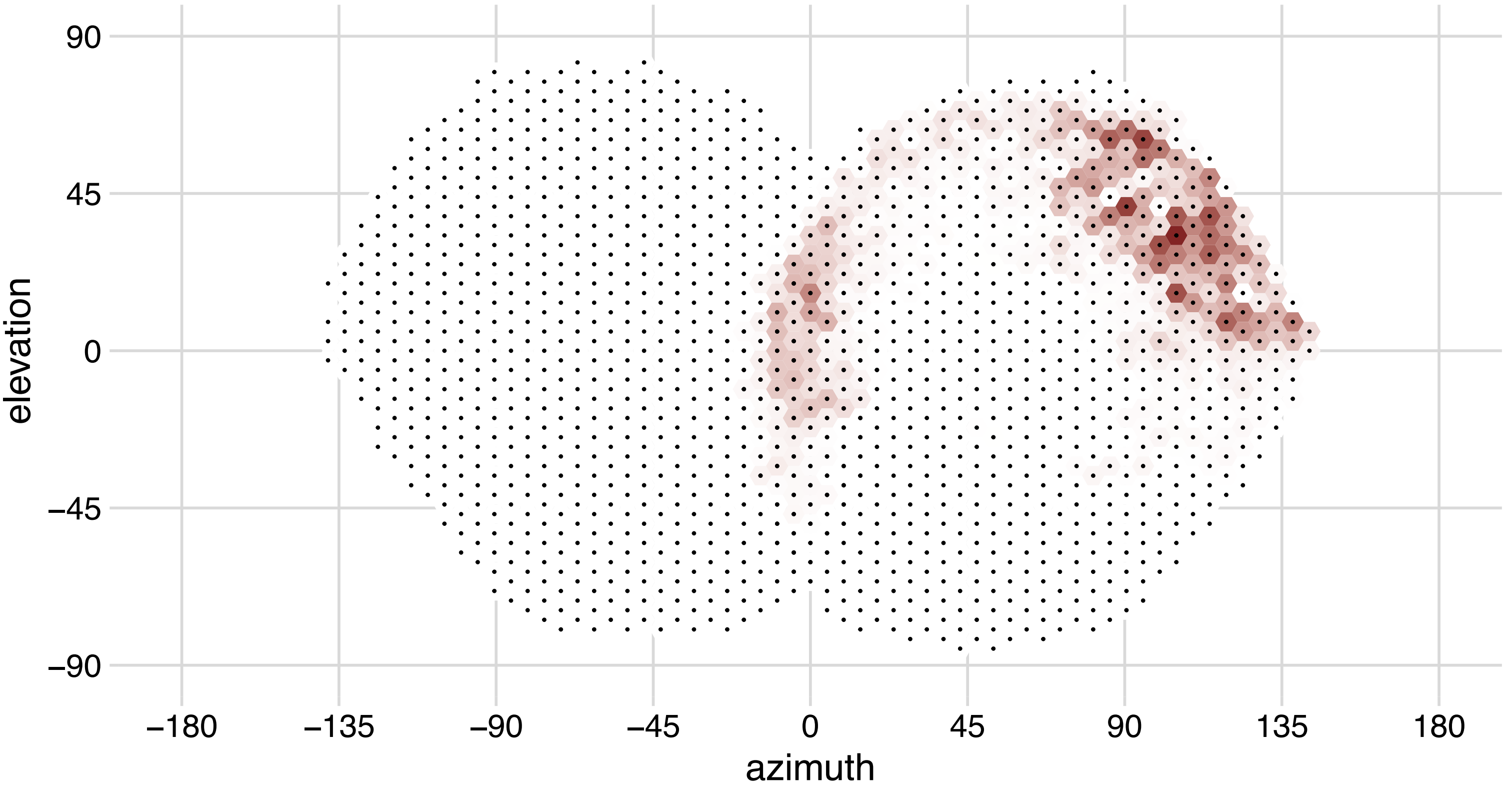

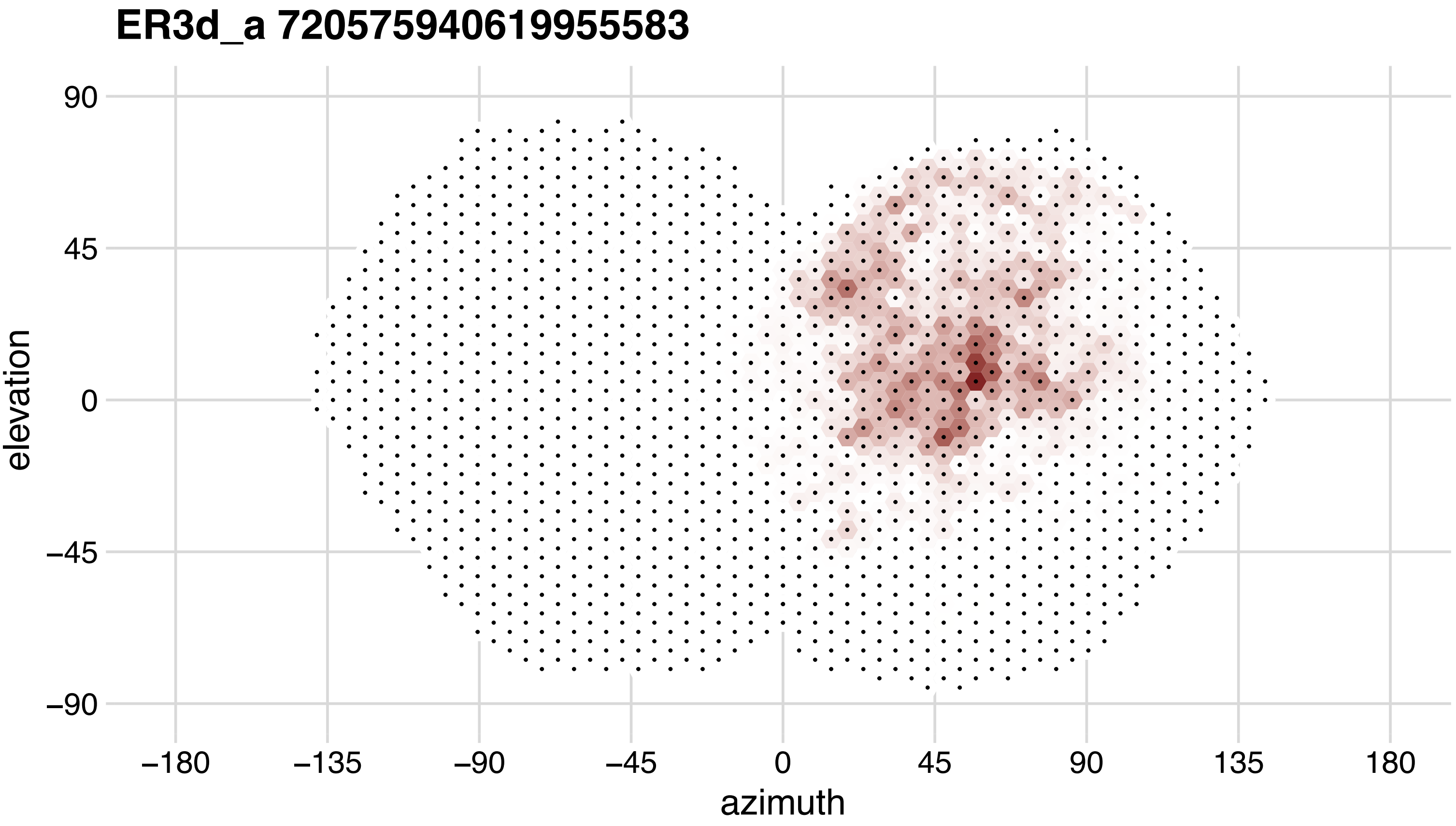
ER3d_a 720575940619955583
90
45
0
−45
−90
elevation
−180
−135
−90
−45
0
45
90
135
180
azimuth

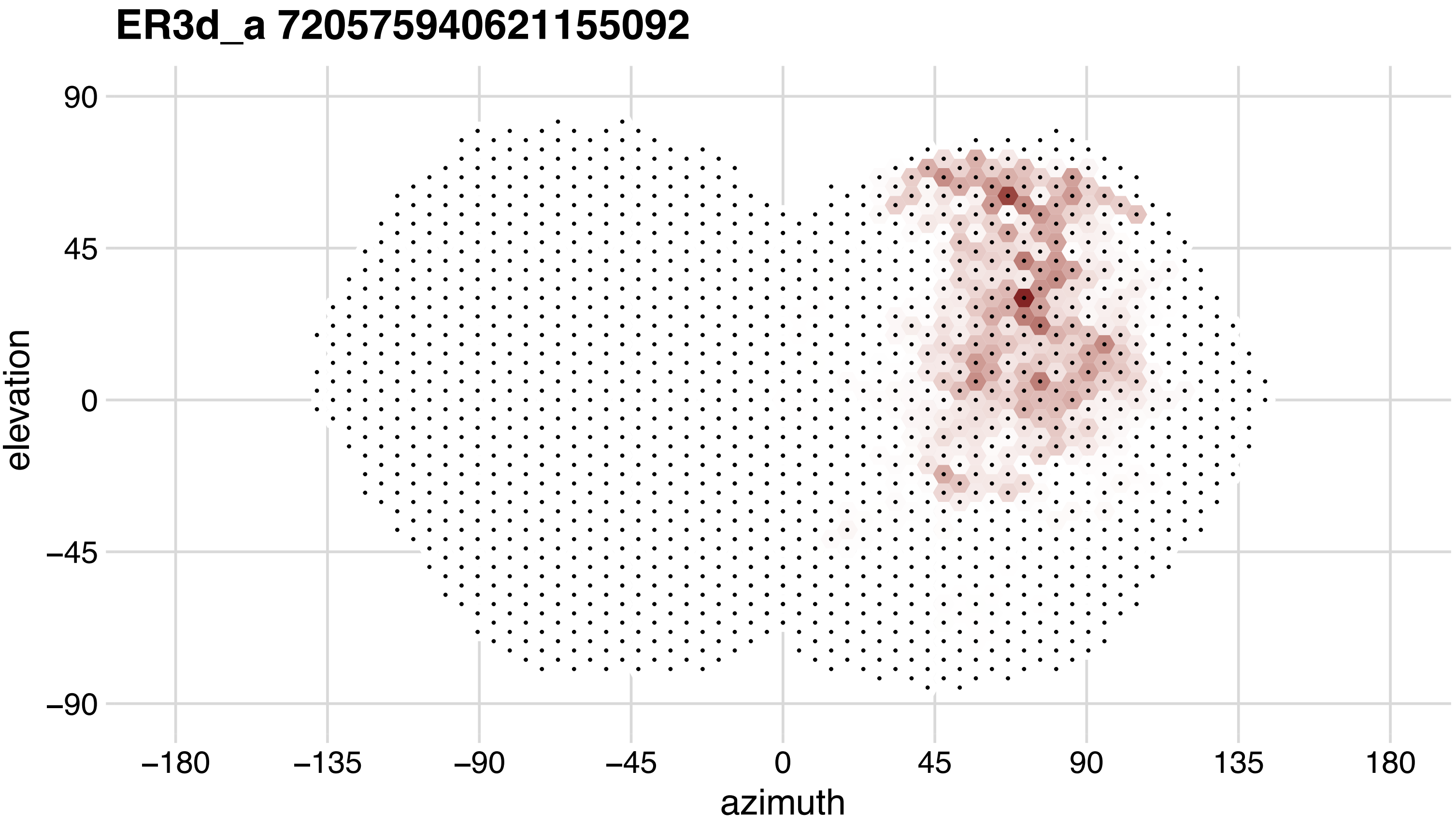

ER3d_a 720575940621155092
90
45
0
−45
−90
elevation
−180
−135
−90
−45
0
45
90
135
180
azimuth

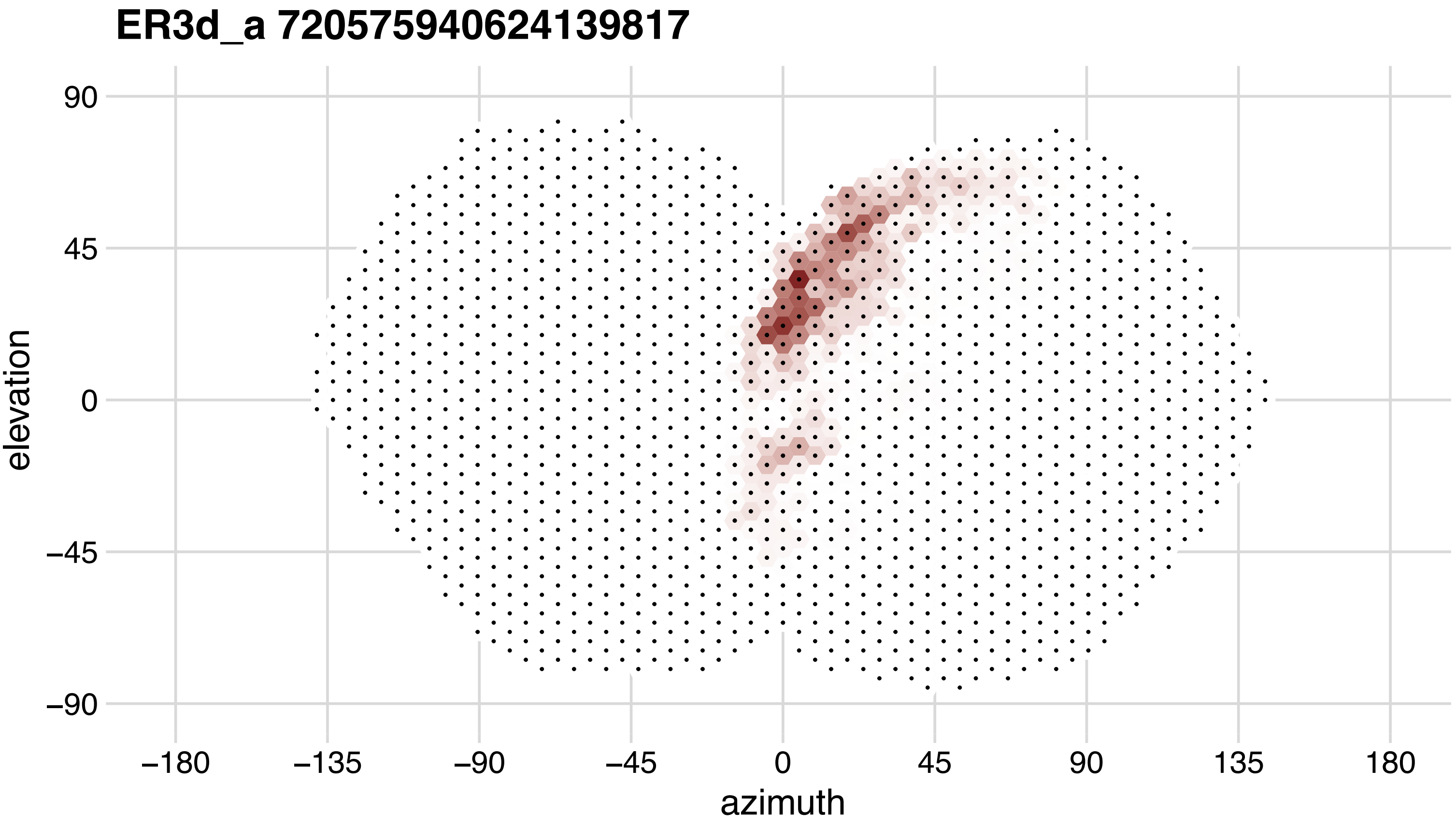
ER3d_a 720575940624139817
90
45
0
−45
−90
elevation
−180
−135
−90
−45
0
45
90
135
180
azimuth

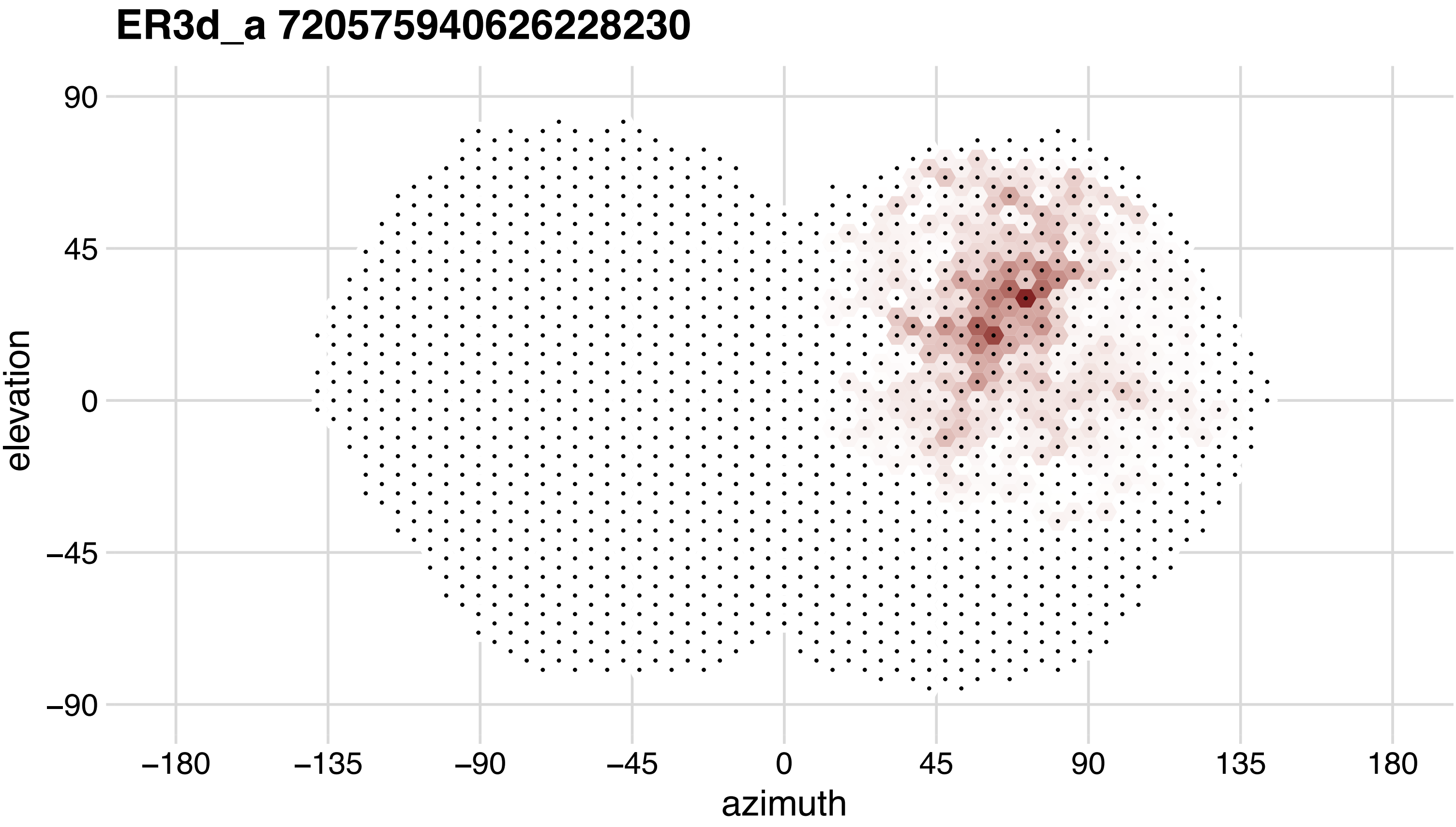
ER3d_a 720575940626228230
90
45
0
−45
−90
elevation
−180
−135
−90
−45
0
45
90
135
180
azimuth

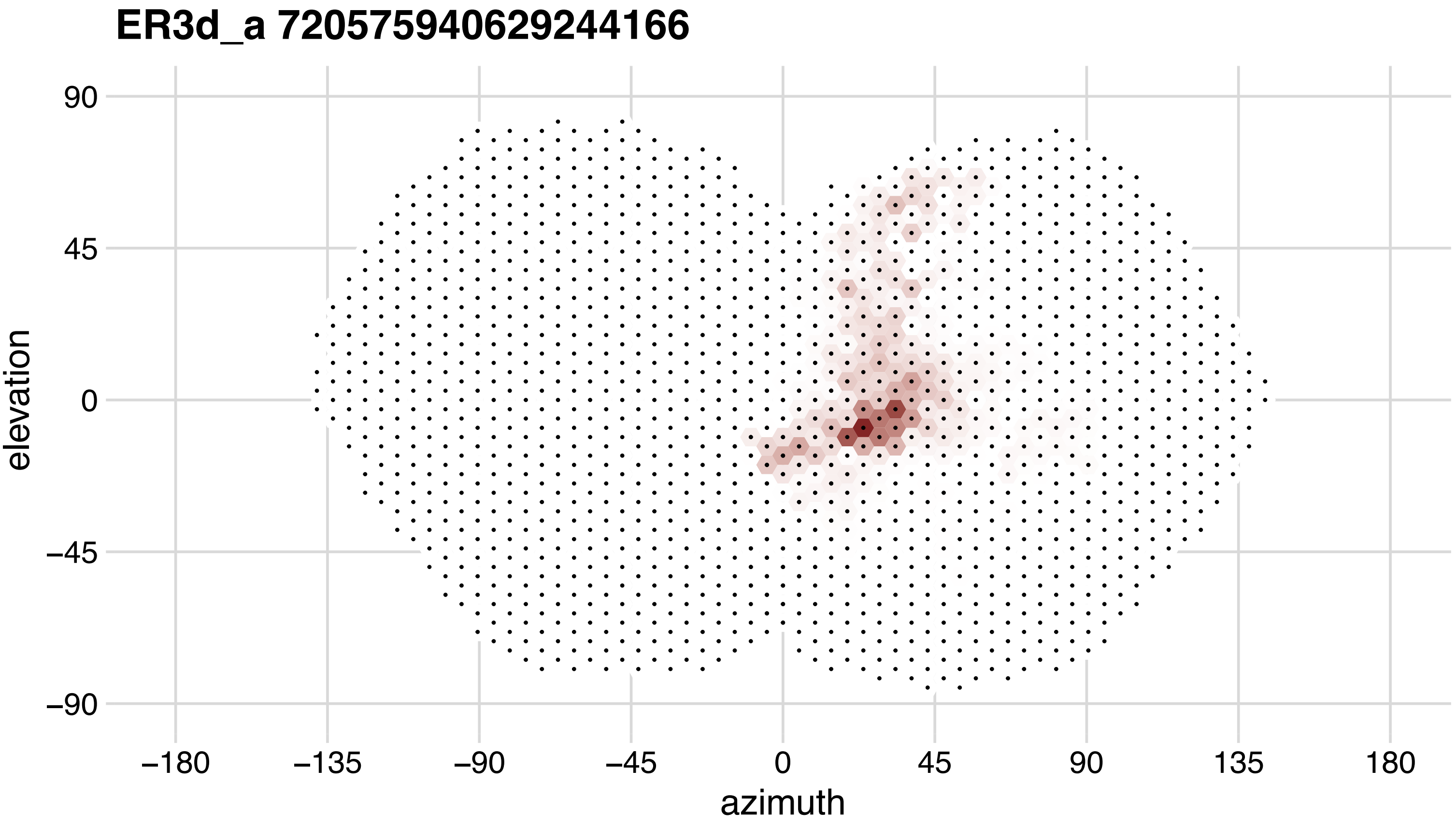
ER3d_a 720575940629244166
elevation
azimuth
90
45
0
−45
−90
−180
−135
−90
−45
0
45
90
135
180

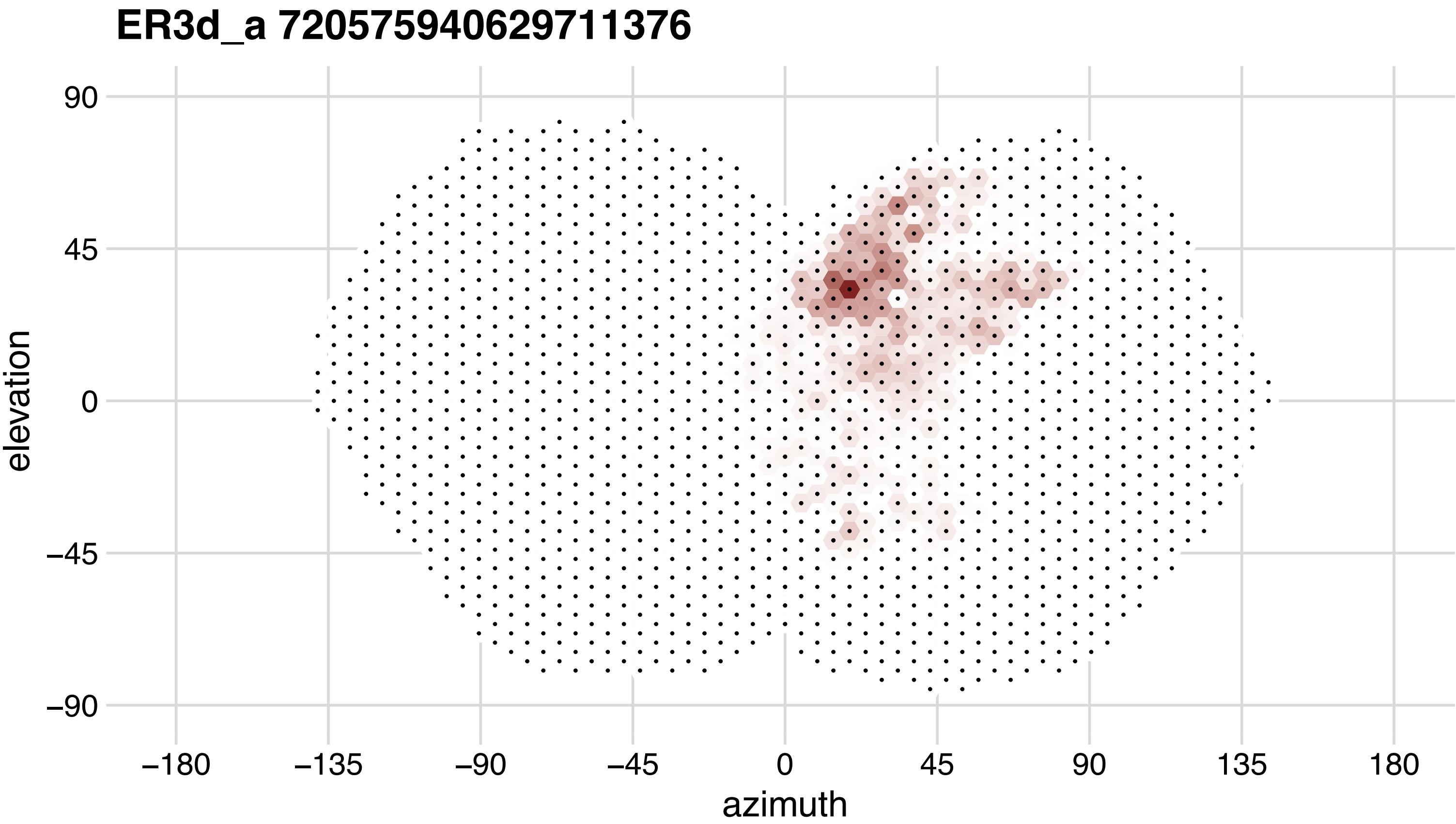
ER3d_a 720575940629711376
90
45
0
−45
−90
elevation
−180
−135
−90
−45
0
45
90
135
180
azimuth

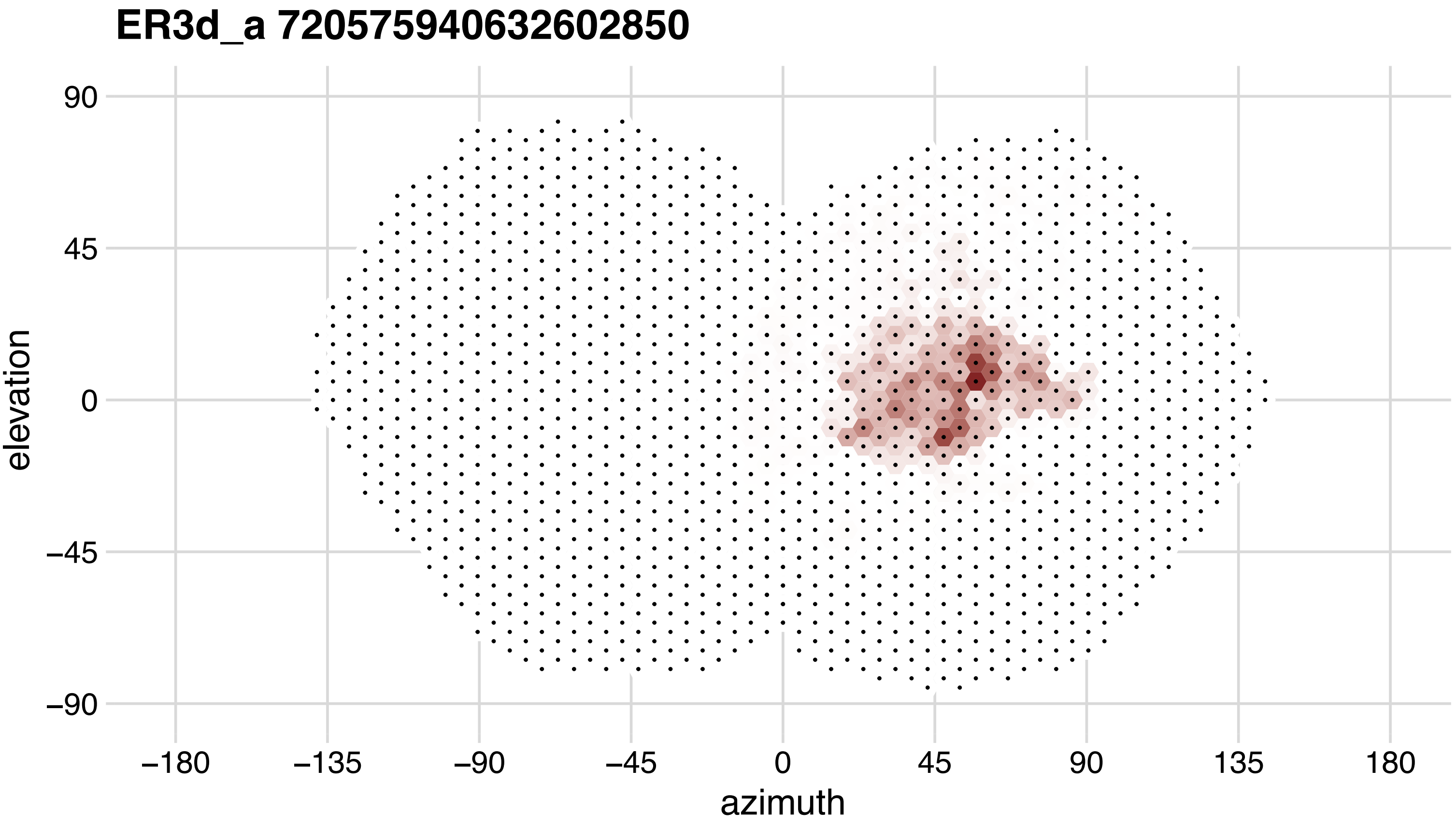

ER3d_a 720575940632602850
90
45
0
−45
−90
elevation
−180
−135
−90
−45
0
45
90
135
180
azimuth

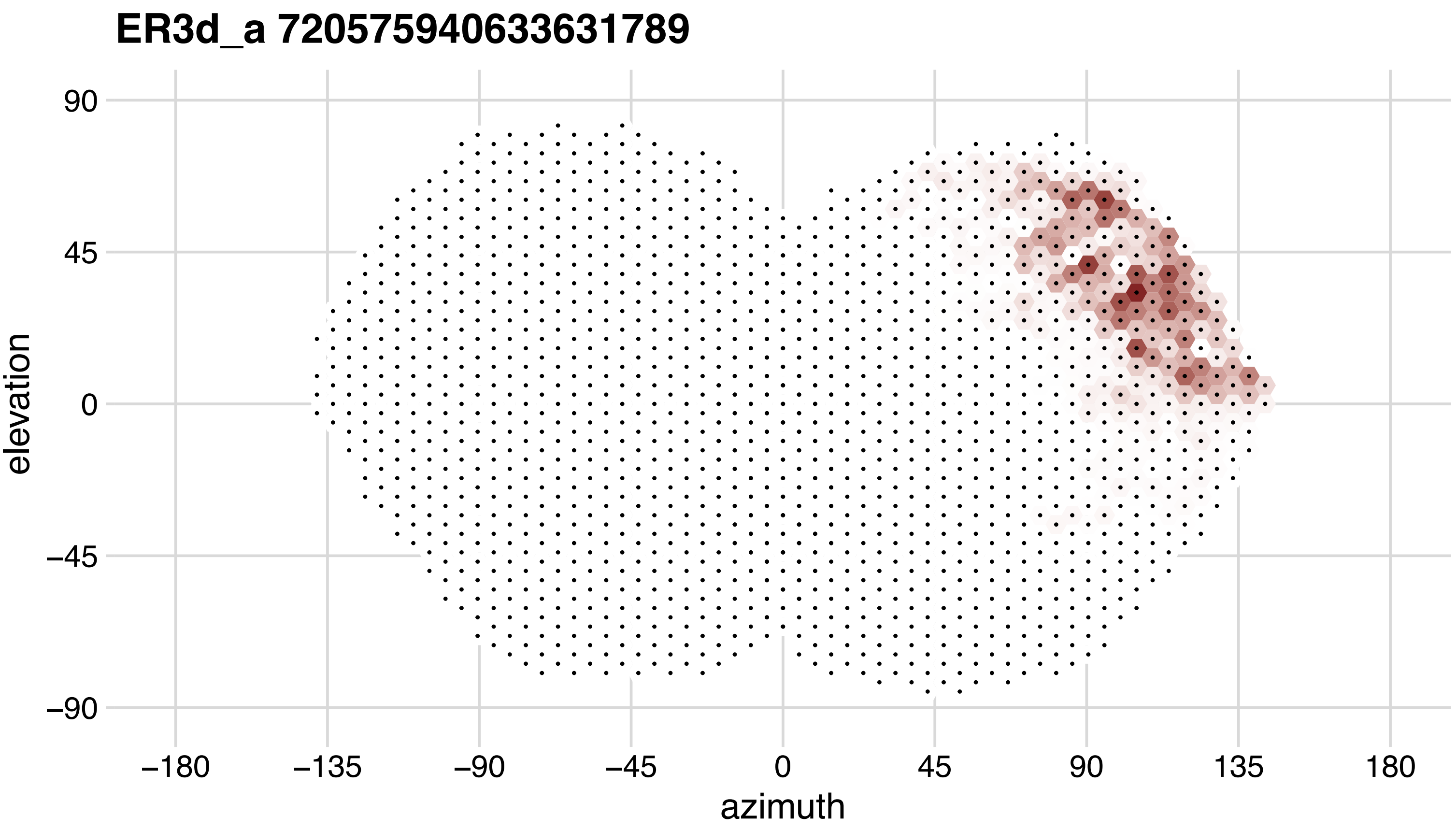
ER3d_a 720575940633631789
90
45
0
−45
−90
elevation
−180
−135
−90
−45
0
45
90
135
180
azimuth

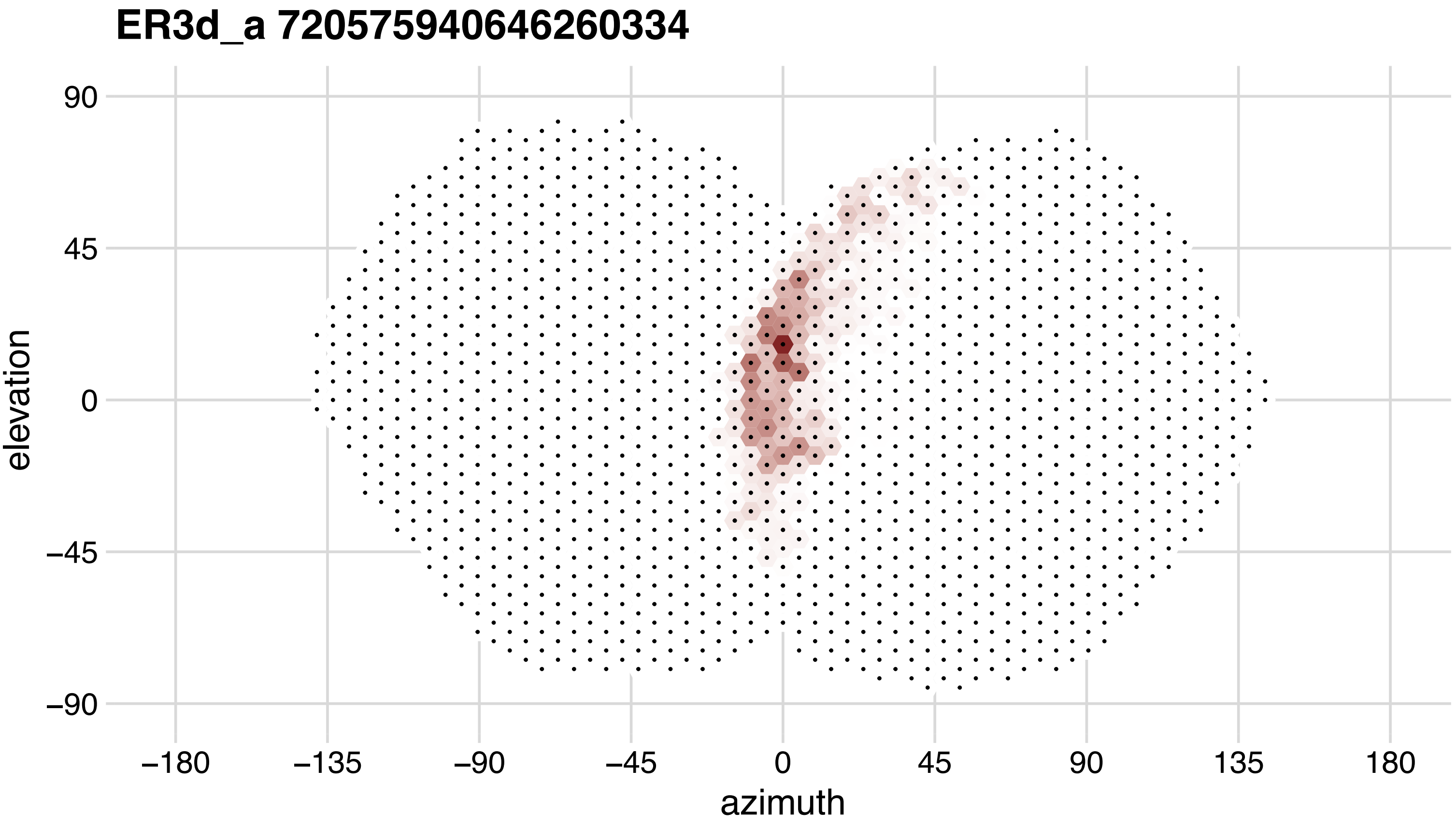
ER3d_a 720575940646260334
90
45
0
−45
−90
elevation
−180
−135
−90
−45
0
45
90
135
180
azimuth

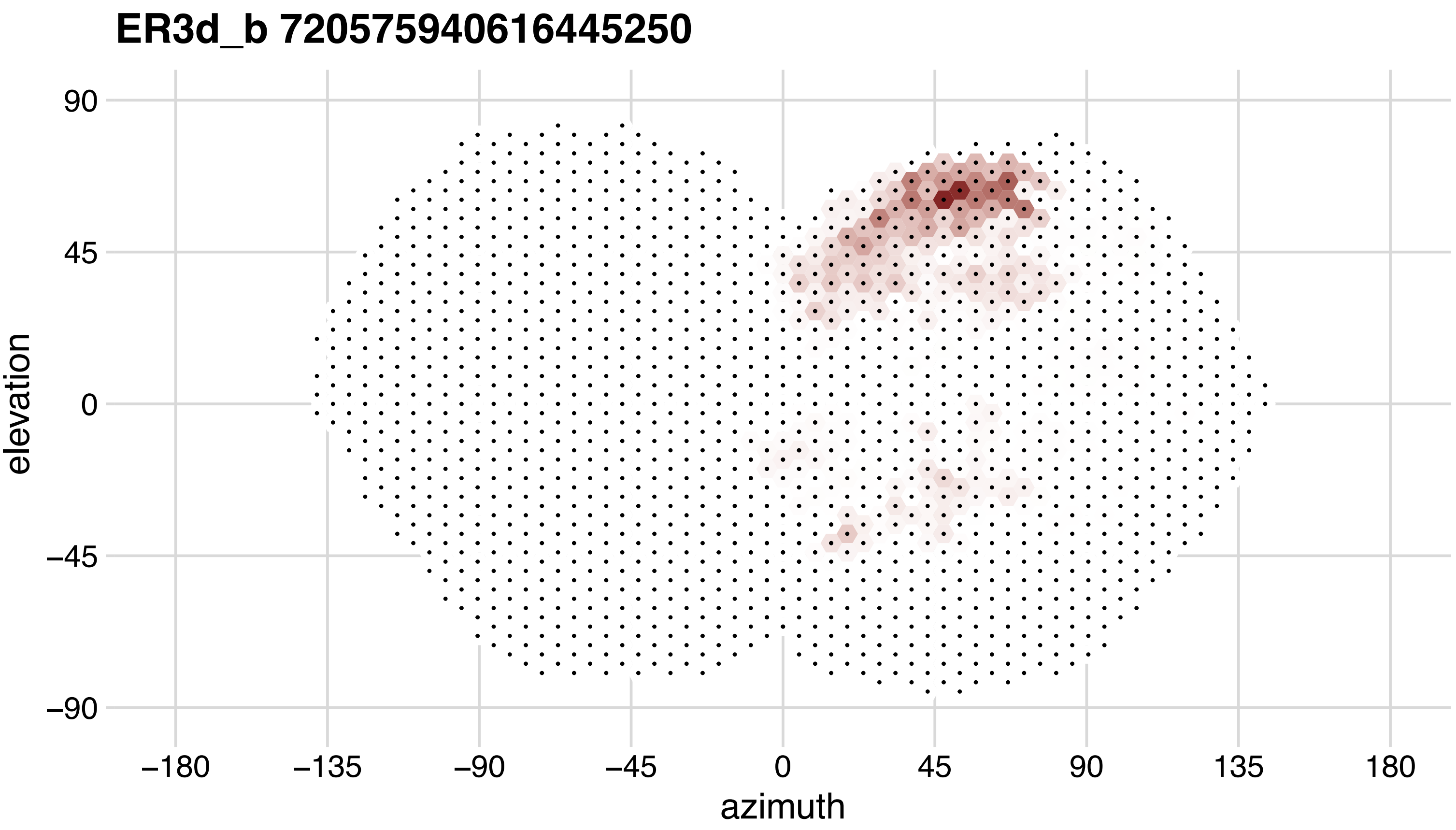
ER3d_b 720575940616445250
90
45
0
−45
−90
elevation
−180
−135
−90
−45
0
45
90
135
180
azimuth

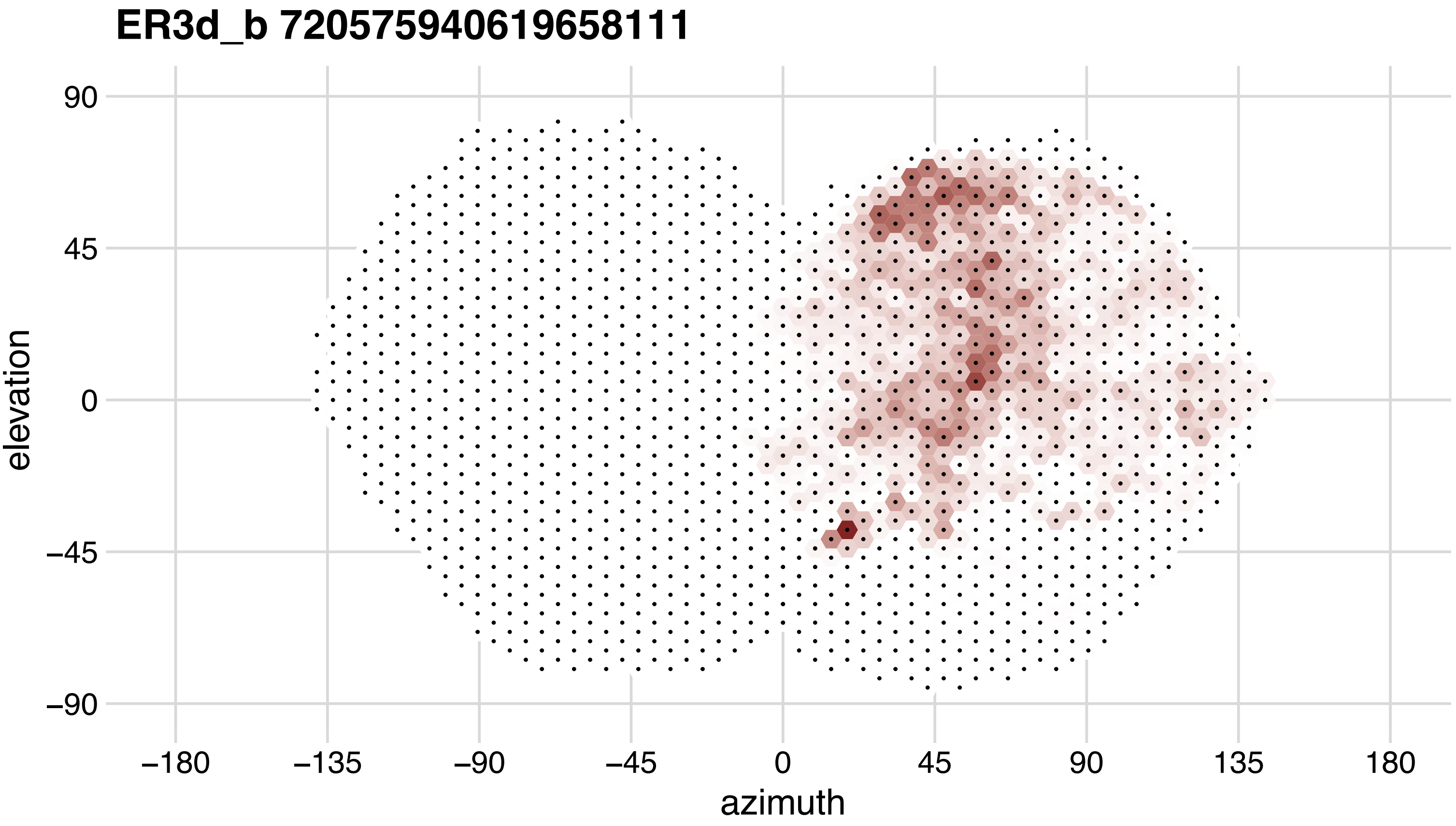

ER3d_b 720575940619658111
90
45
0
−45
−90
elevation
−180
−135
−90
−45
0
45
90
135
180
azimuth

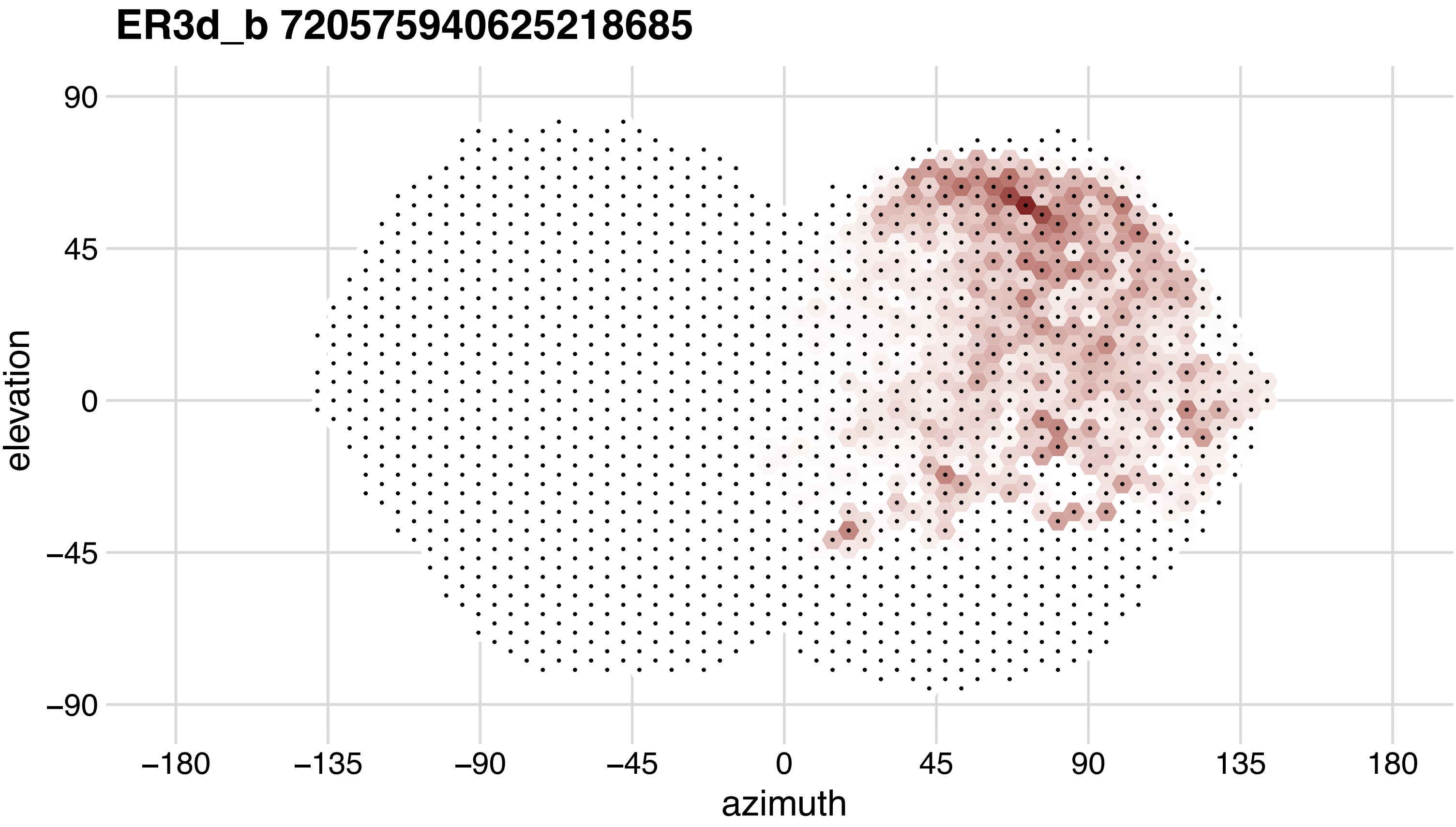
ER3d_b 720575940625218685
90
45
0
−45
−90
elevation
−180
−135
−90
−45
0
45
90
135
180
azimuth

## ER3d_b 720575940626827024

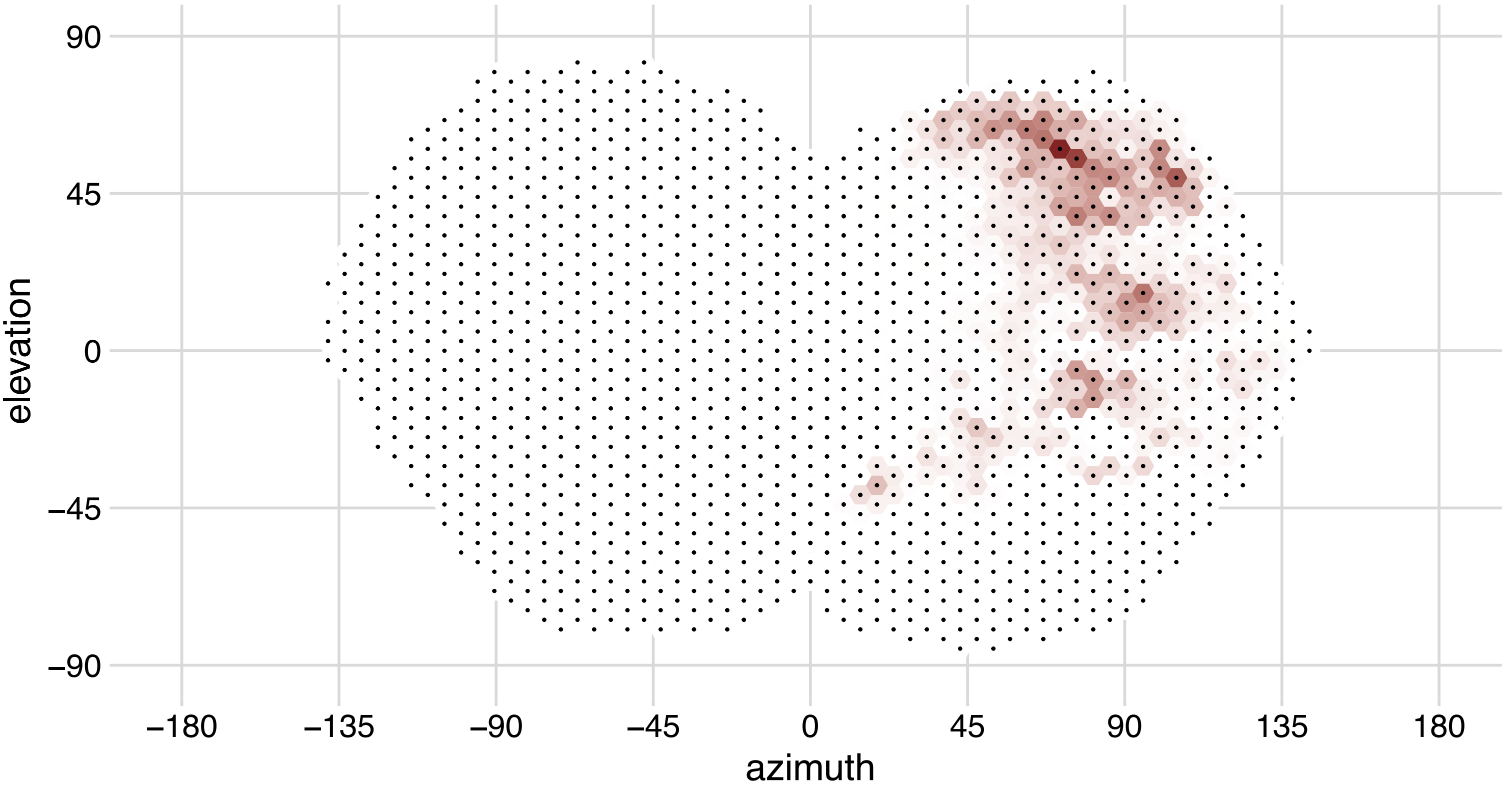

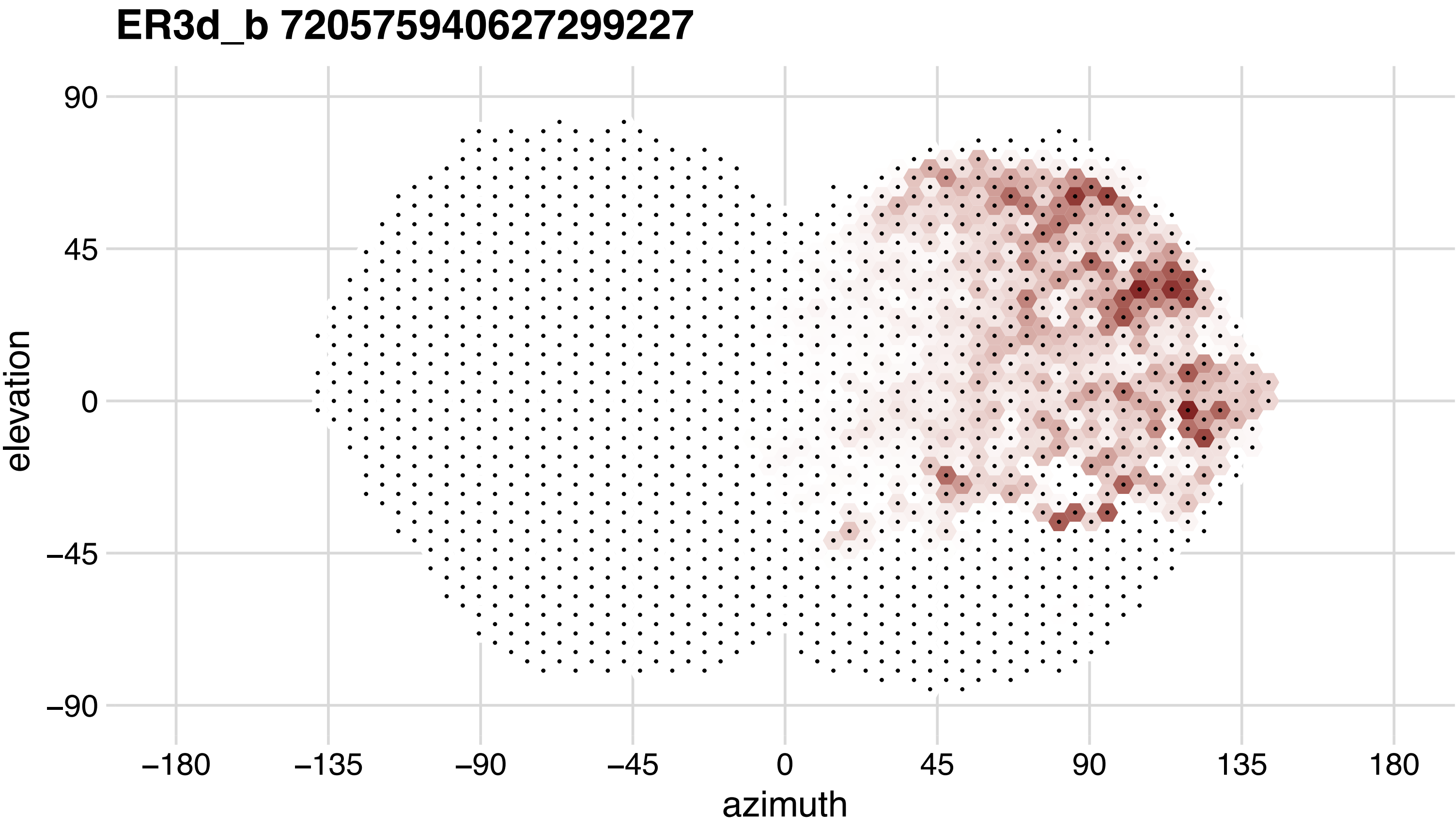

ER3d_b 720575940627299227
90
45
0
−45
−90
elevation
−180
−135
−90
−45
0
45
90
135
180
azimuth

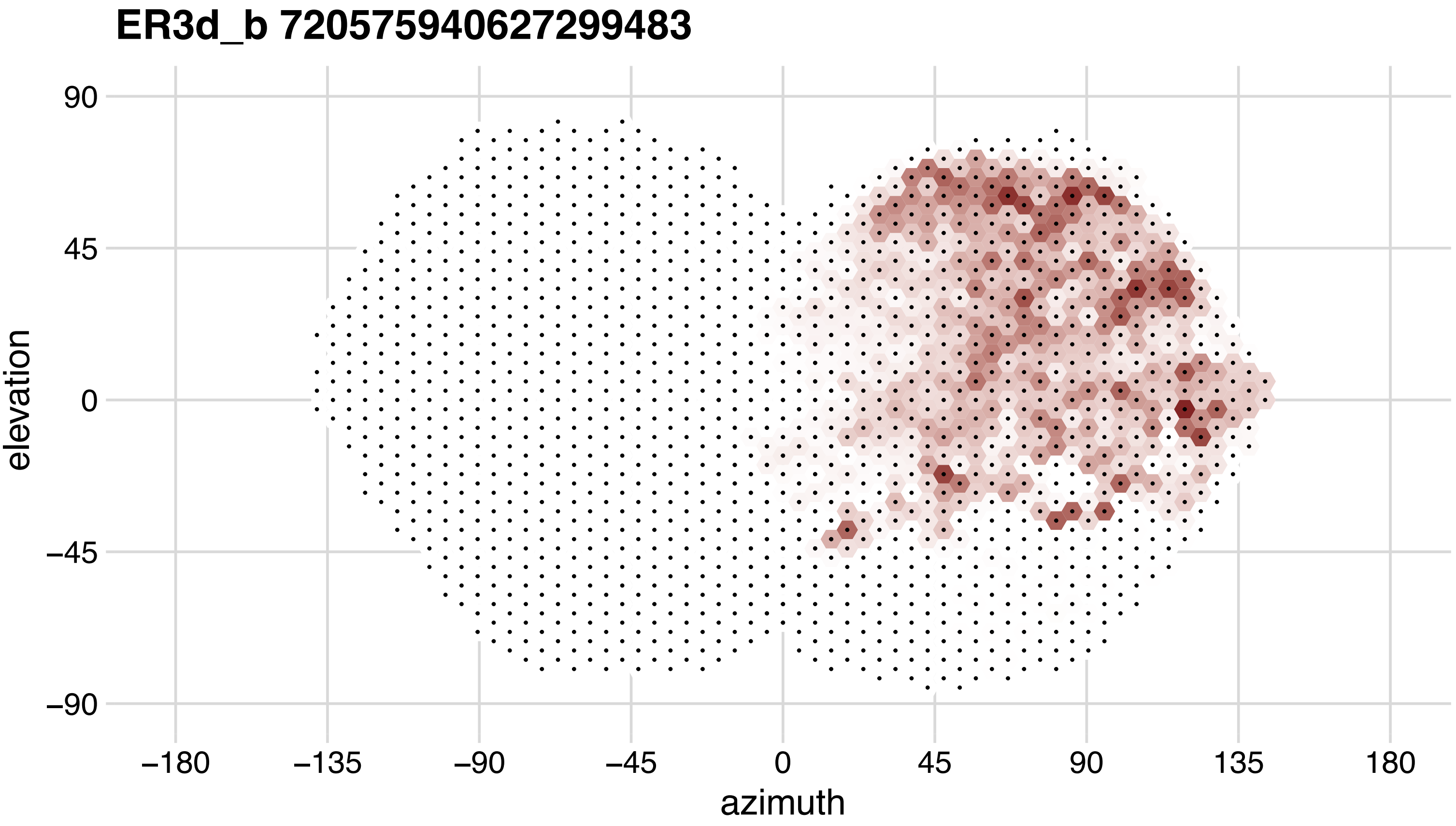
ER3d_b 720575940627299483
90
45
0
−45
−90
elevation
−180
−135
−90
−45
0
45
90
135
180
azimuth

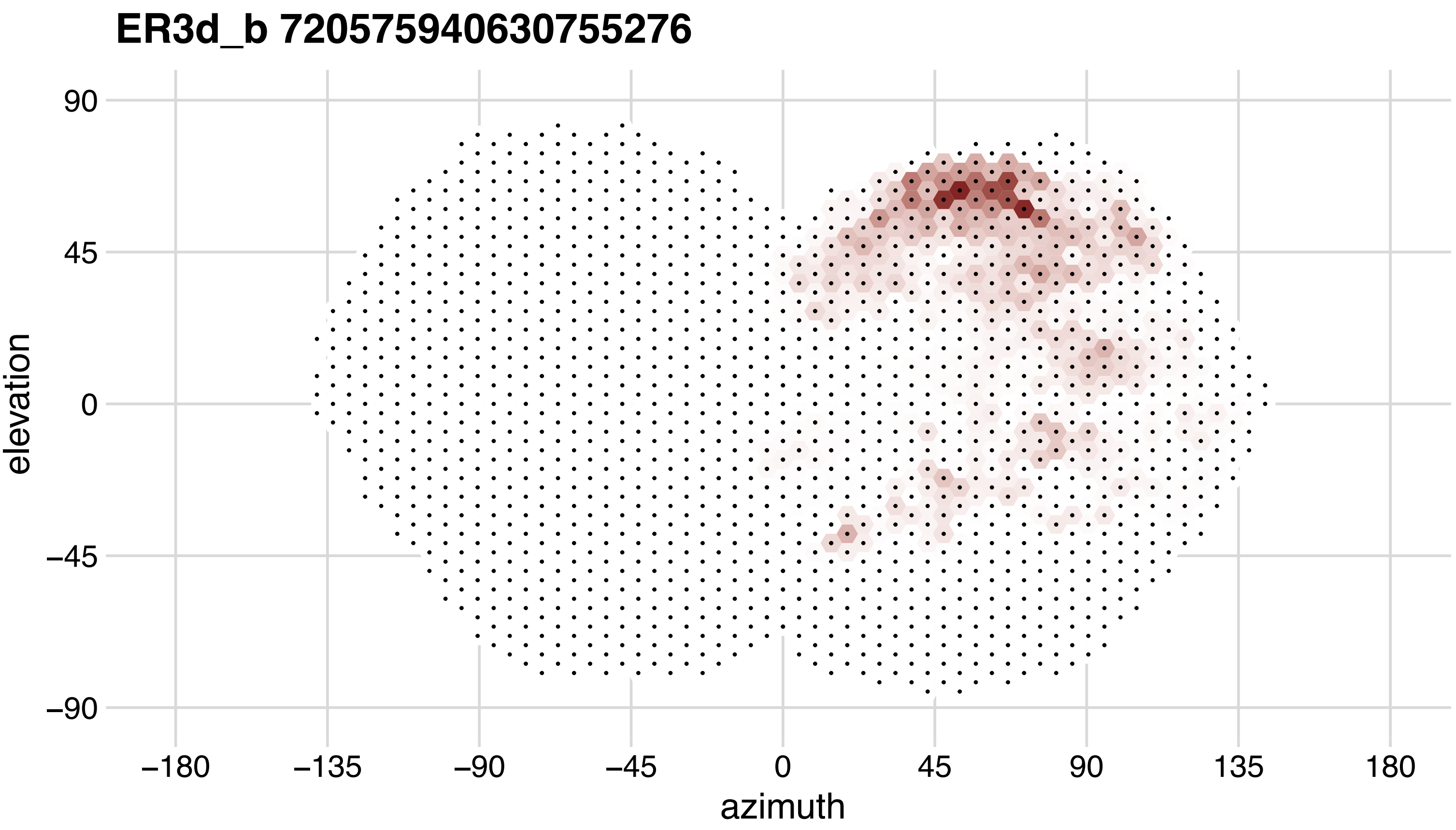
ER3d_b 720575940630755276
90
45
0
−45
−90
elevation
−180
−135
−90
−45
0
45
90
135
180
azimuth

## ER3d_c 720575940616205083

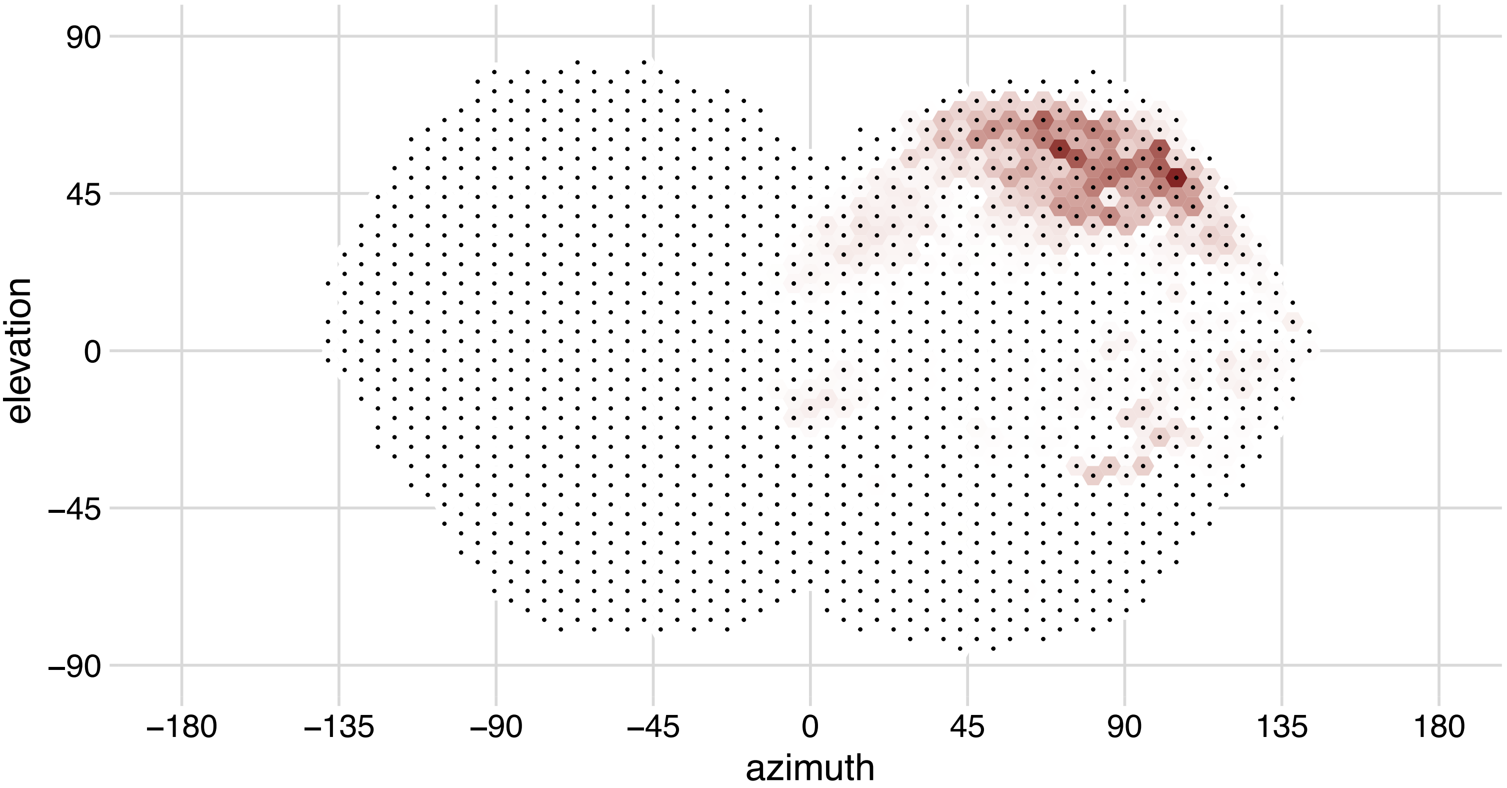

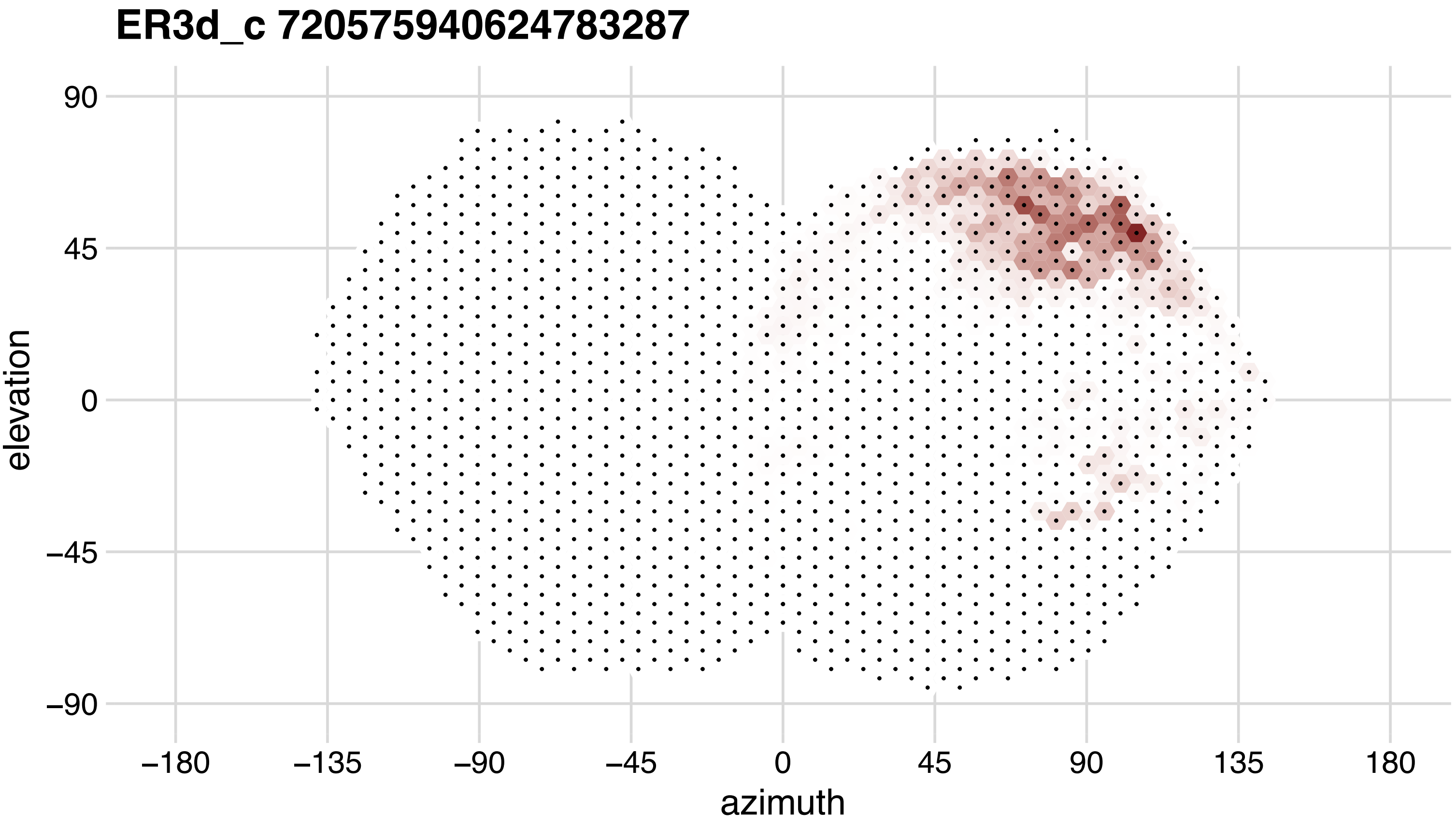
ER3d_c 720575940624783287
90
45
0
−45
−90
elevation
−180
−135
−90
−45
0
45
90
135
180
azimuth

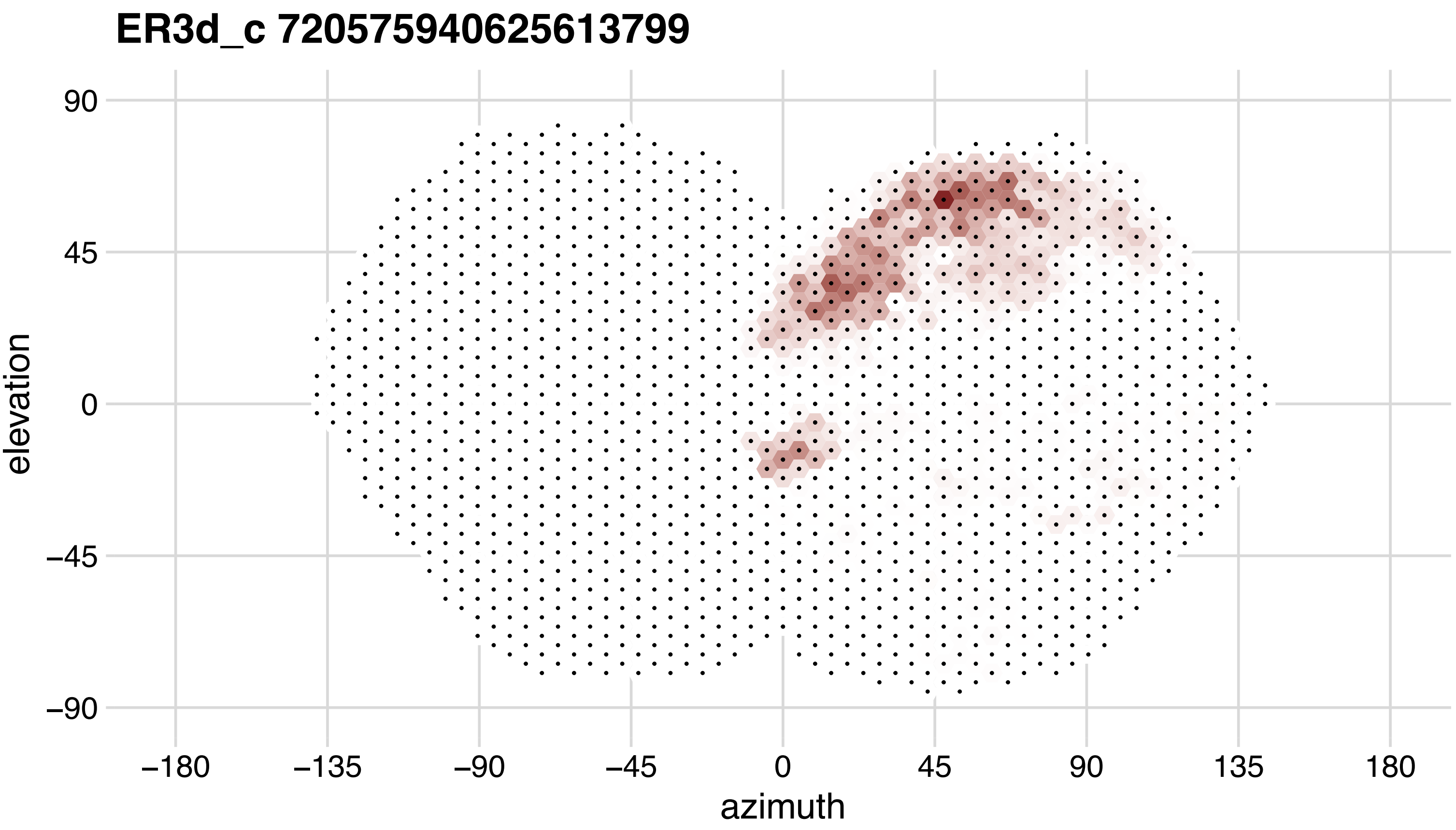
ER3d_c 720575940625613799
90
45
0
−45
−90
elevation
−180
−135
−90
−45
0
45
90
135
180
azimuth

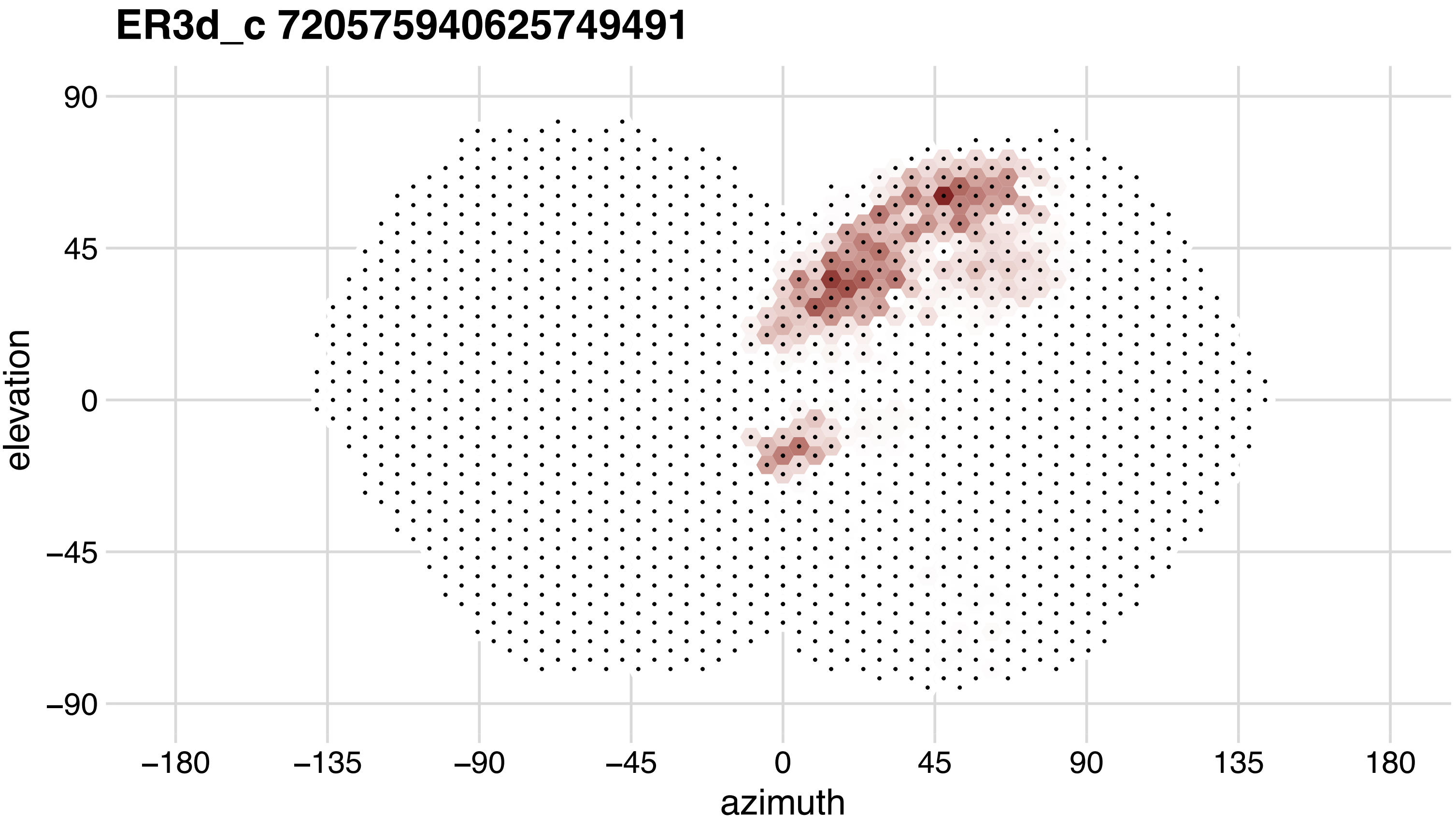
ER3d_c 720575940625749491
90
45
0
−45
−90
elevation
−180
−135
−90
−45
0
45
90
135
180
azimuth

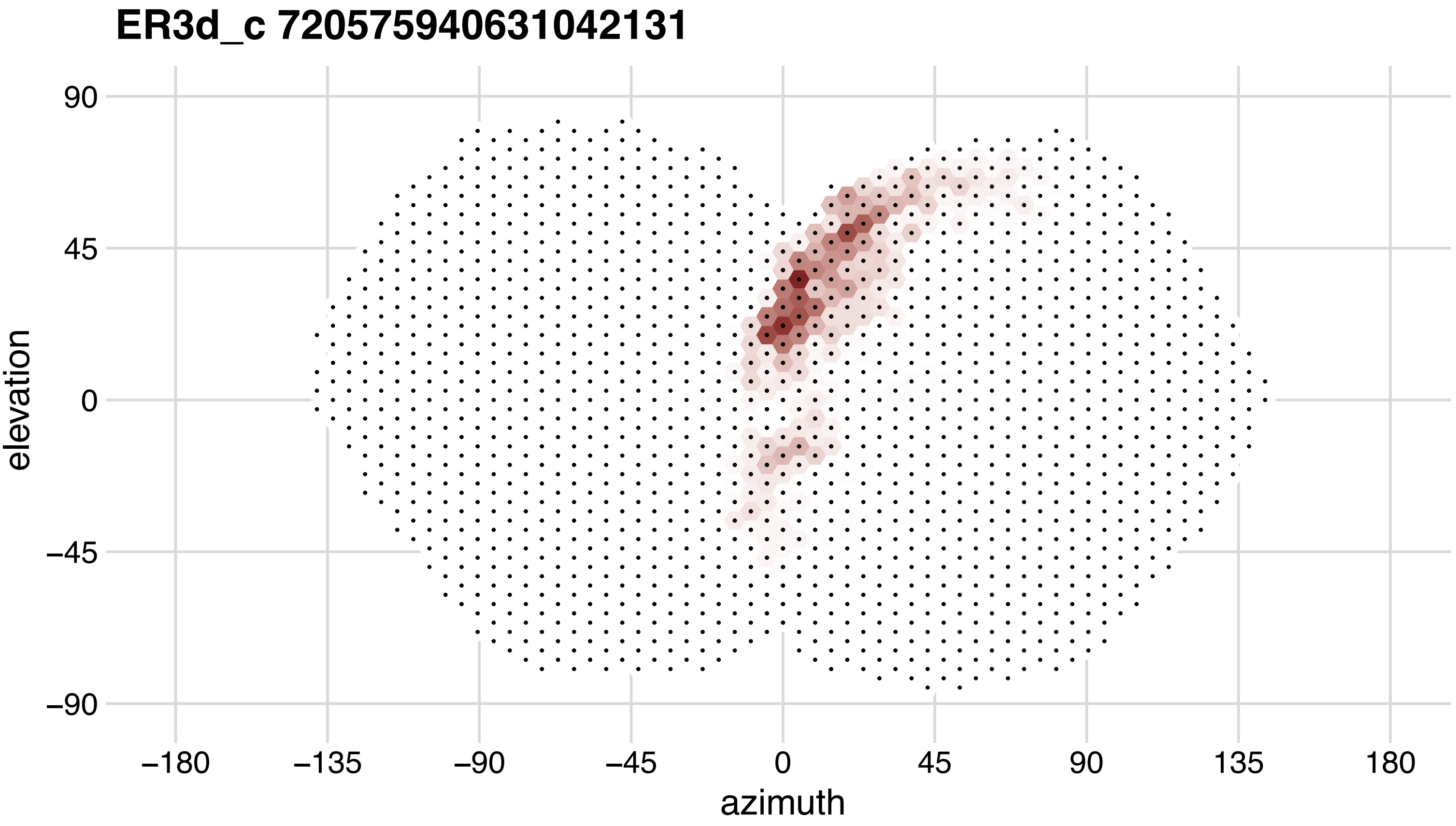

ER3d_c 720575940631042131
90
45
0
−45
−90
elevation
−180
−135
−90
−45
0
45
90
135
180
azimuth

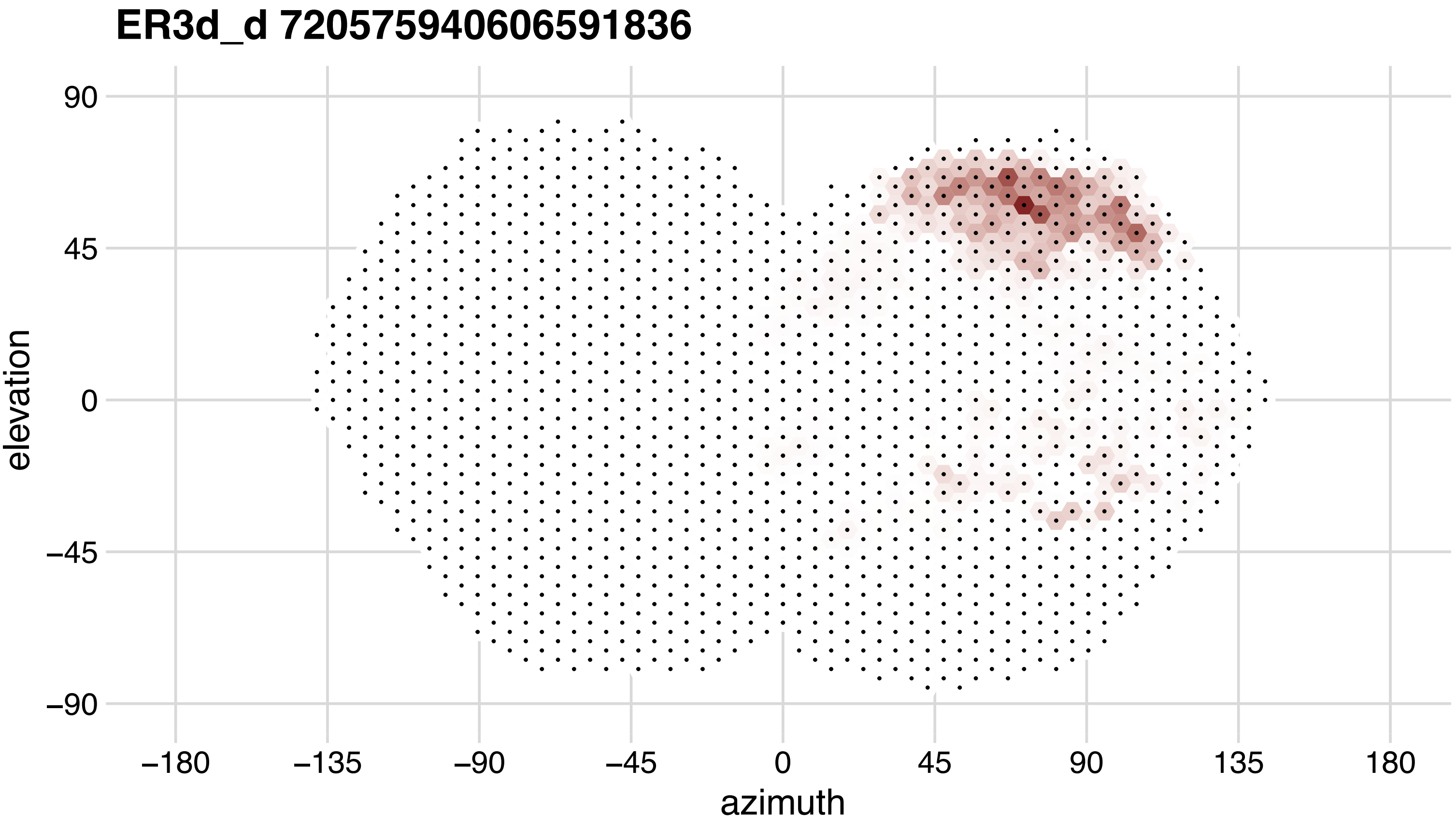
ER3d_d 720575940606591836
elevation
90
45
0
−45
−90
−180
−135
−90
−45
0
45
90
135
180
azimuth

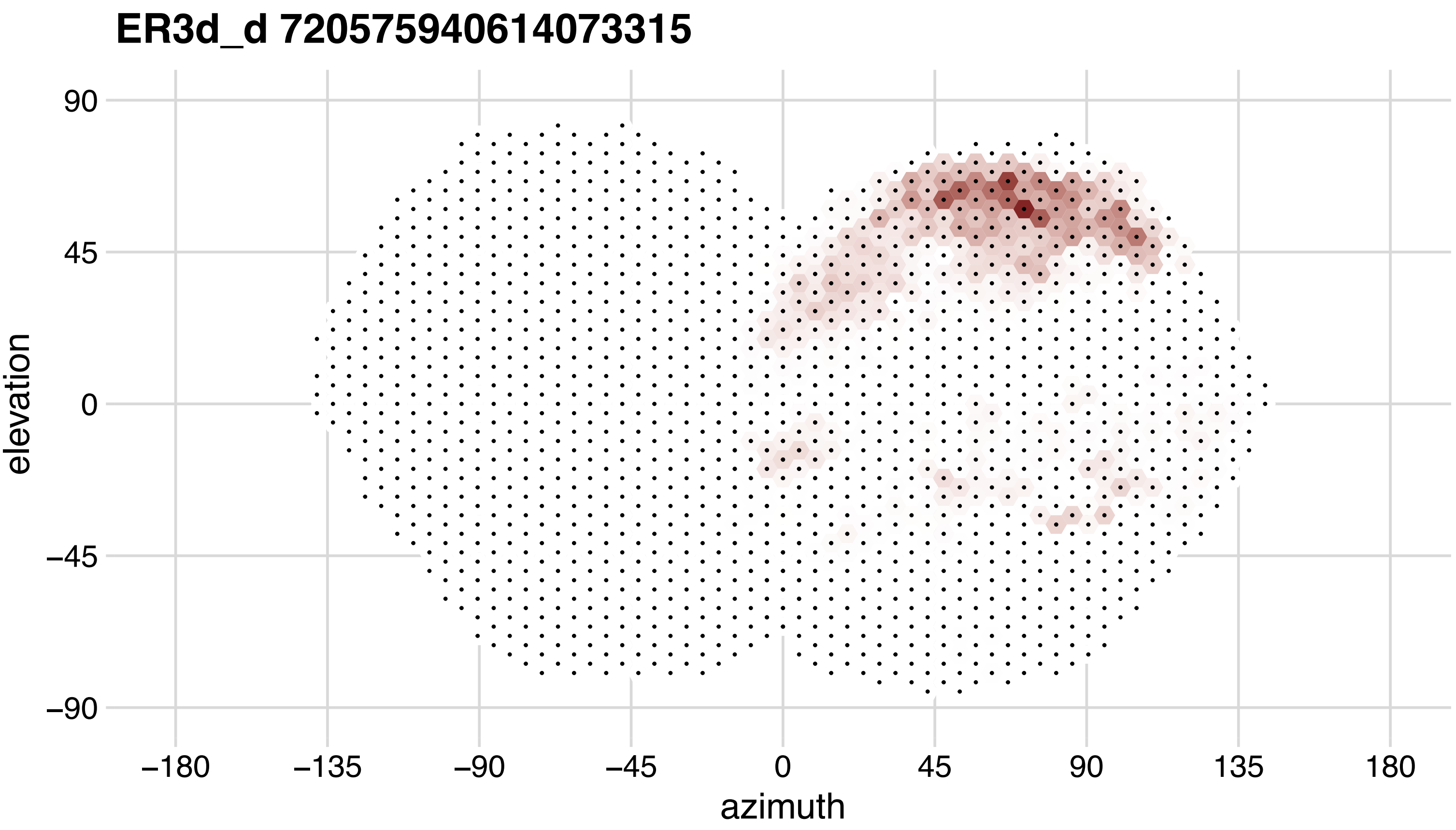
ER3d_d 720575940614073315
elevation
90
45
0
−45
−90
azimuth
−180
−135
−90
−45
0
45
90
135
180

## ER3d_d 720575940619829765

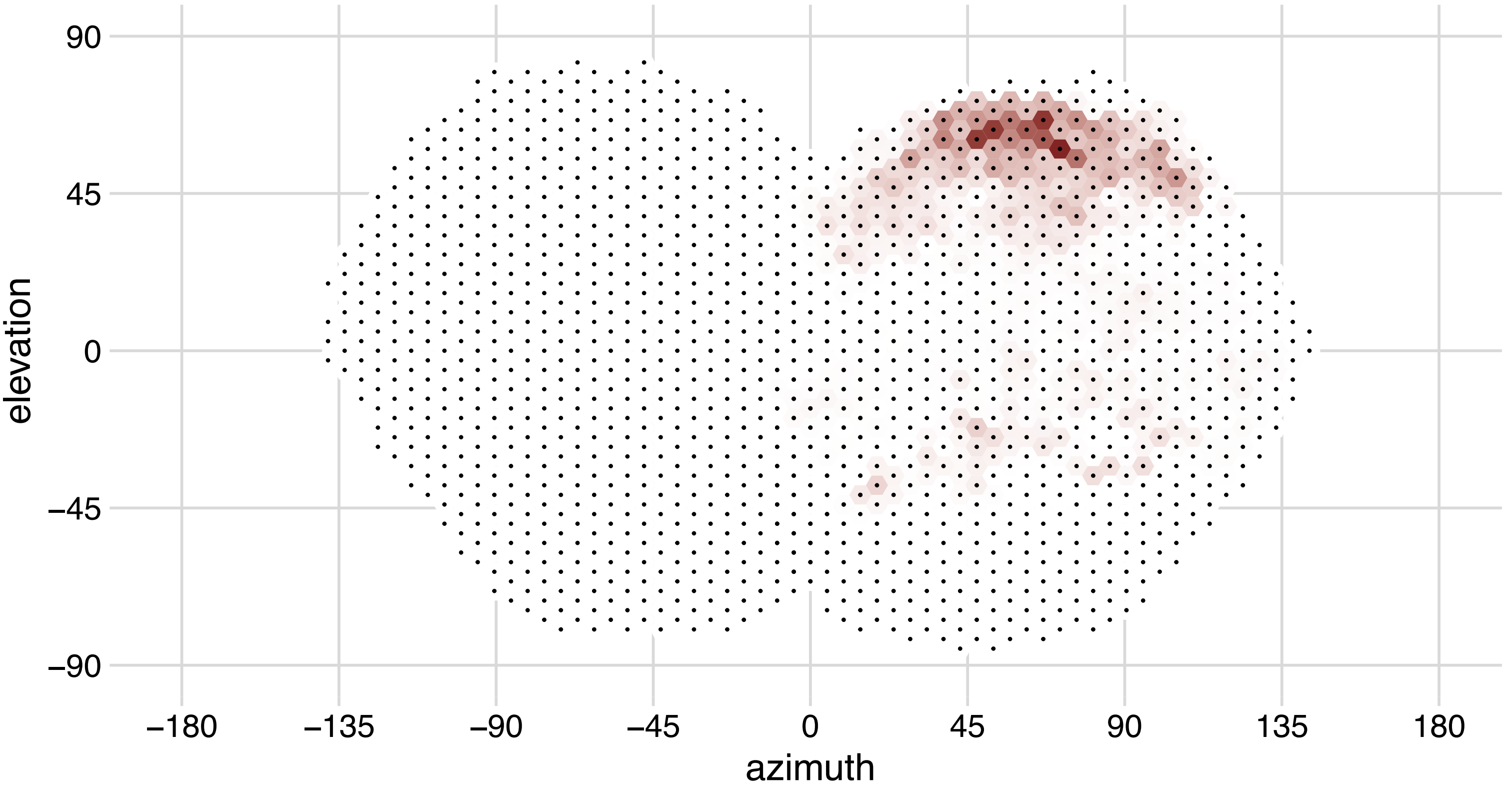

## ER3m 720575940609621003

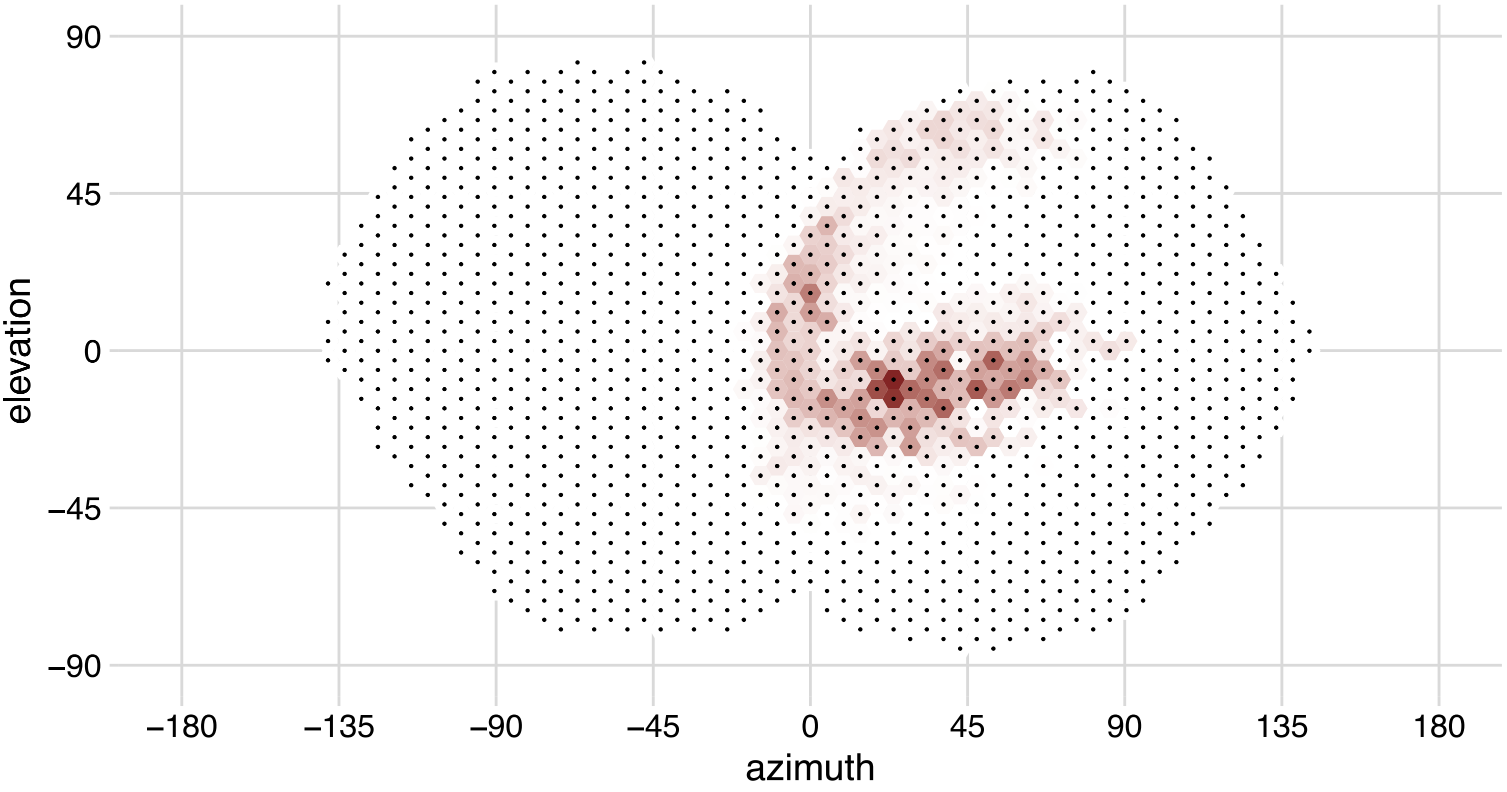

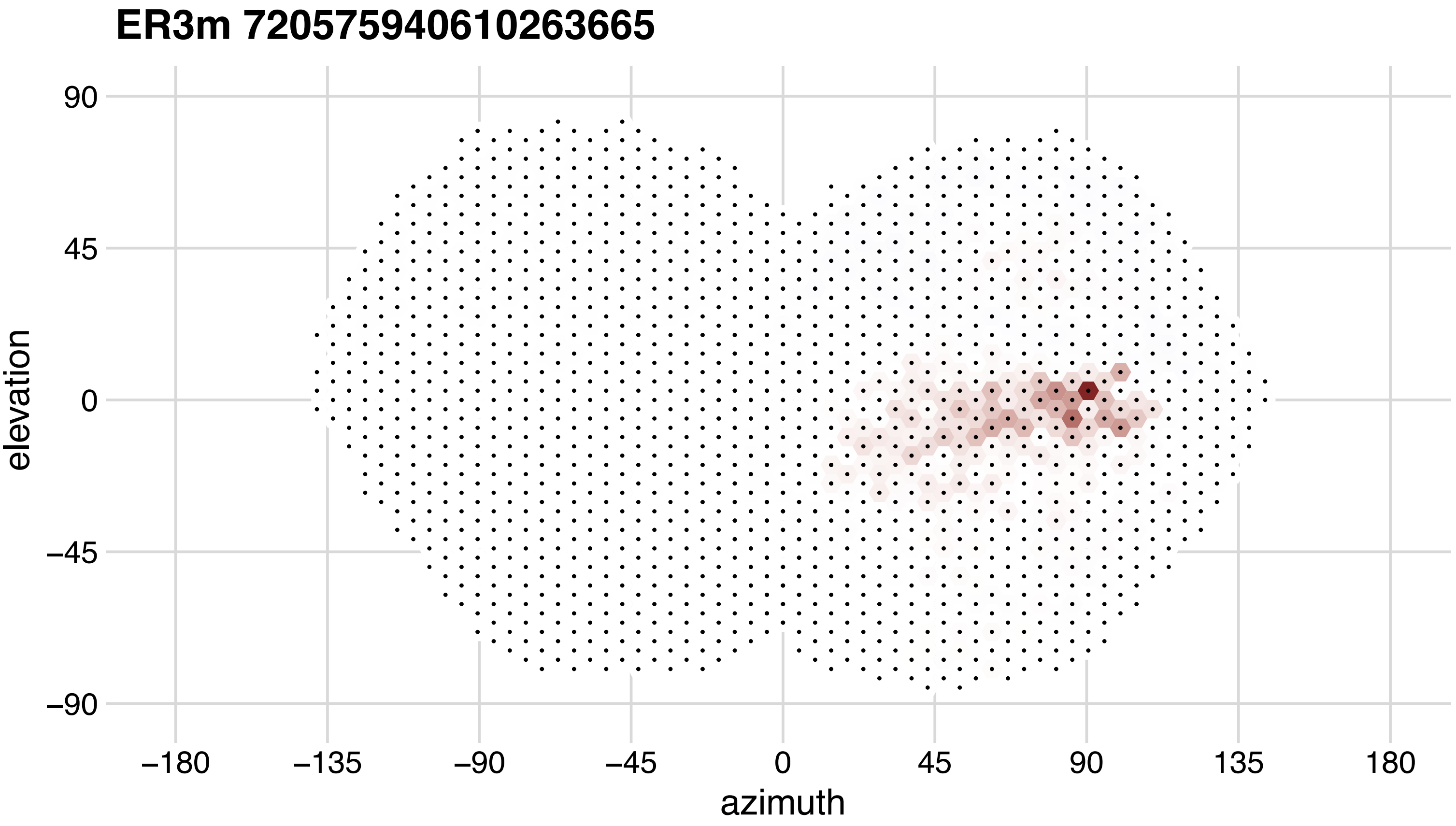
ER3m 720575940610263665
90
45
0
−45
−90
elevation
−180
−135
−90
−45
0
45
90
135
180
azimuth

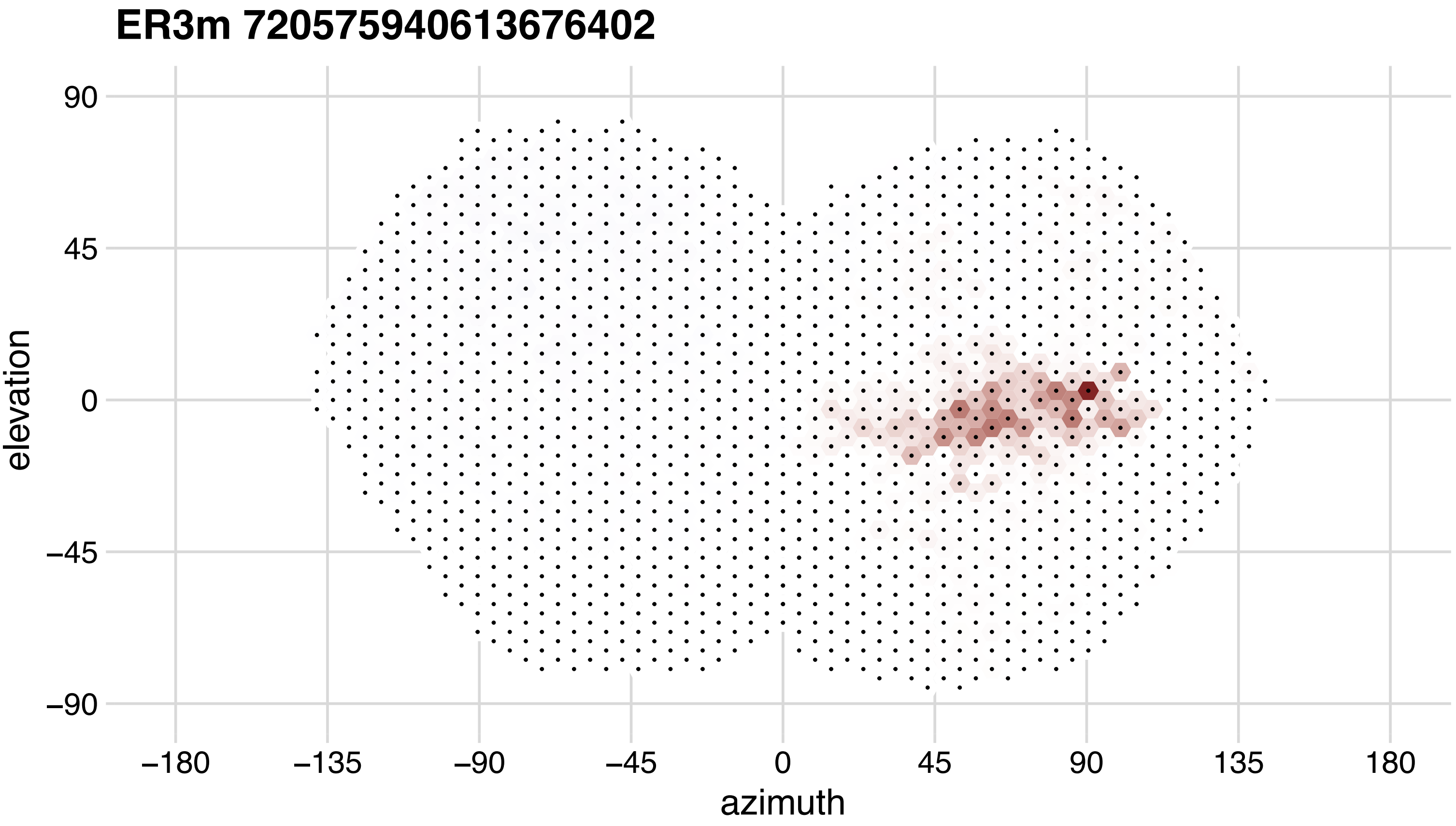

ER3m 720575940613676402
90
45
0
−45
−90
elevation
−180
−135
−90
−45
0
45
90
135
180
azimuth

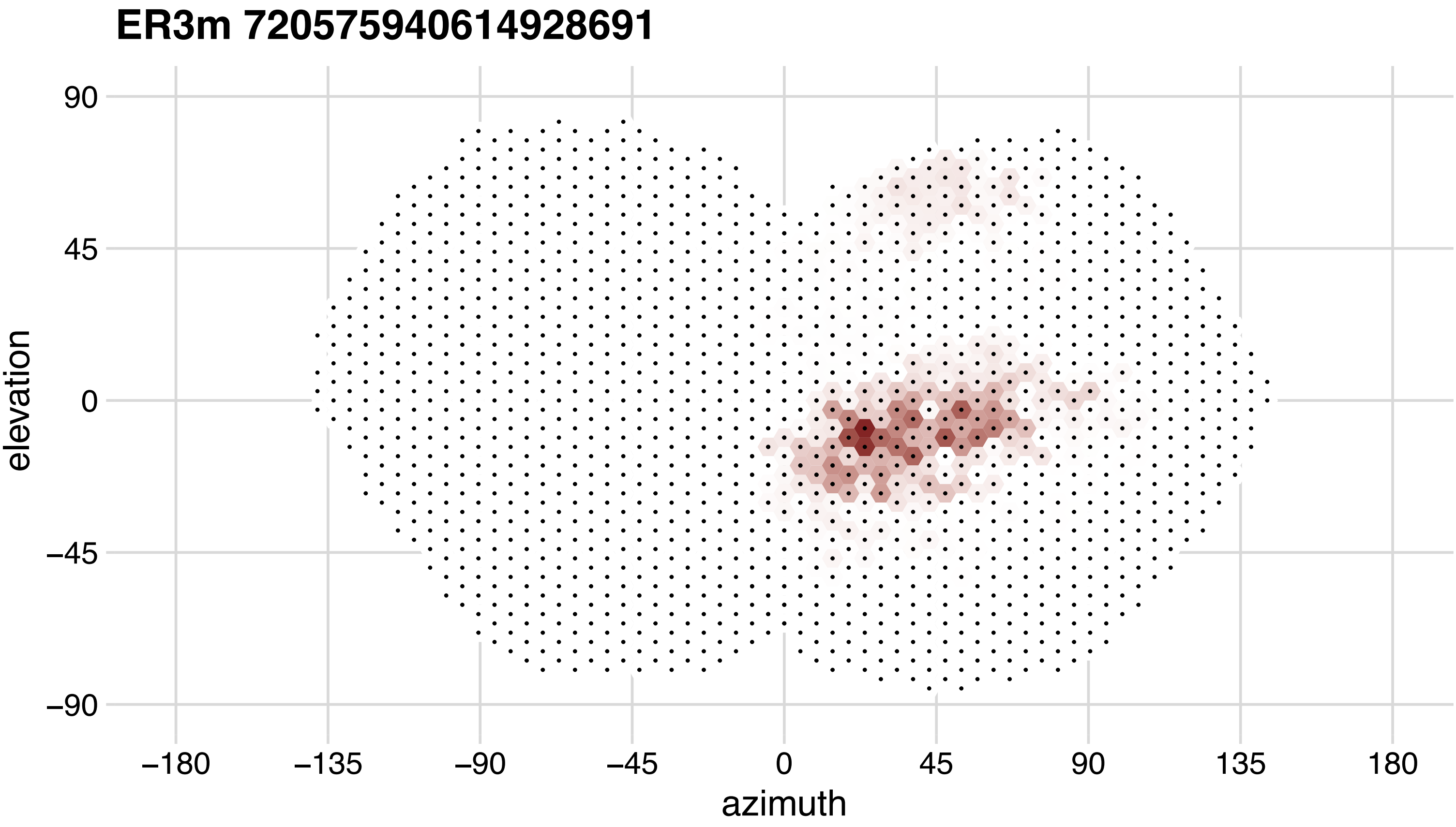
ER3m 720575940614928691
90
45
0
−45
−90
elevation
−180
−135
−90
−45
0
45
90
135
180
azimuth

## ER3m 720575940616441410

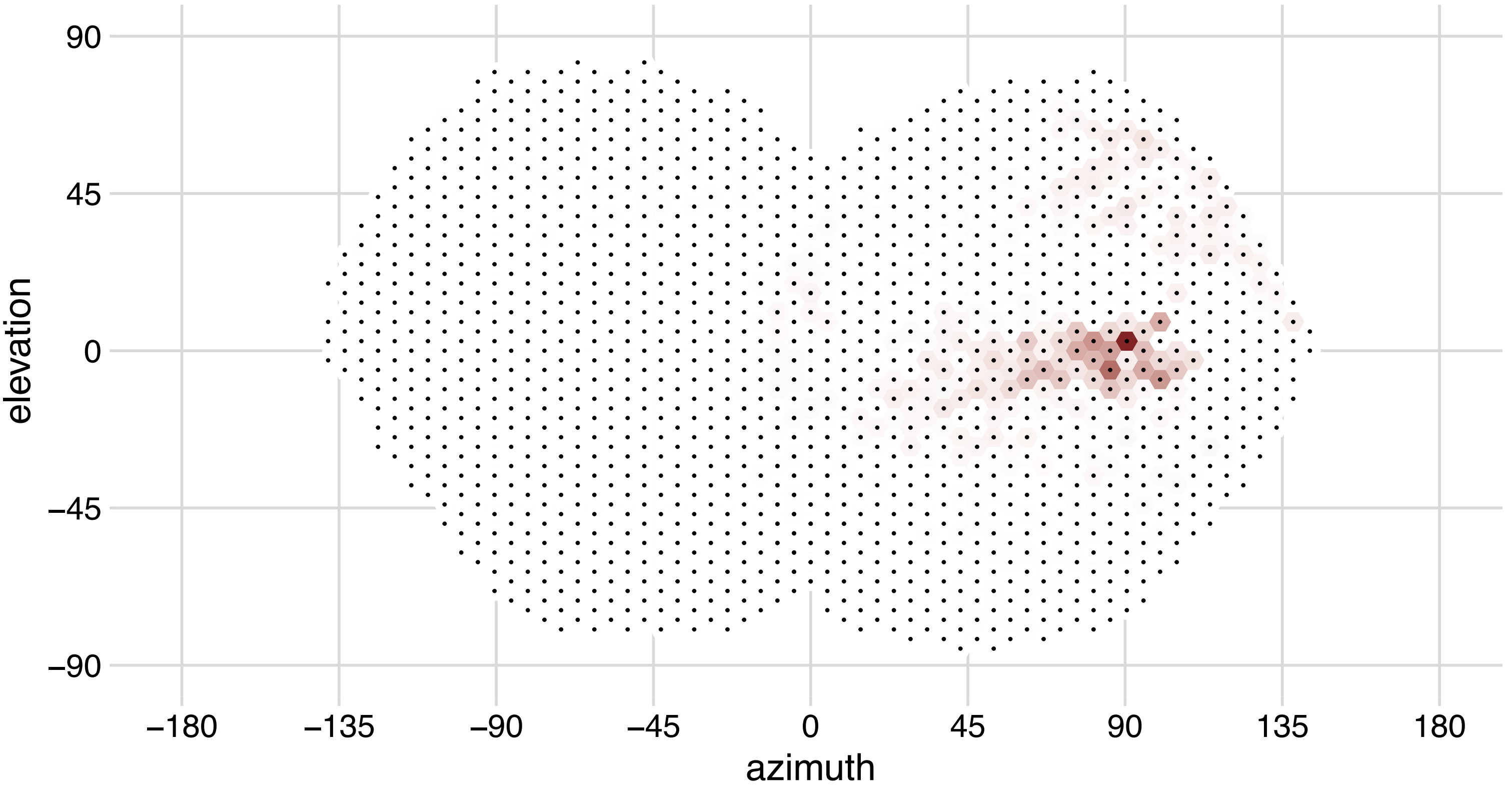

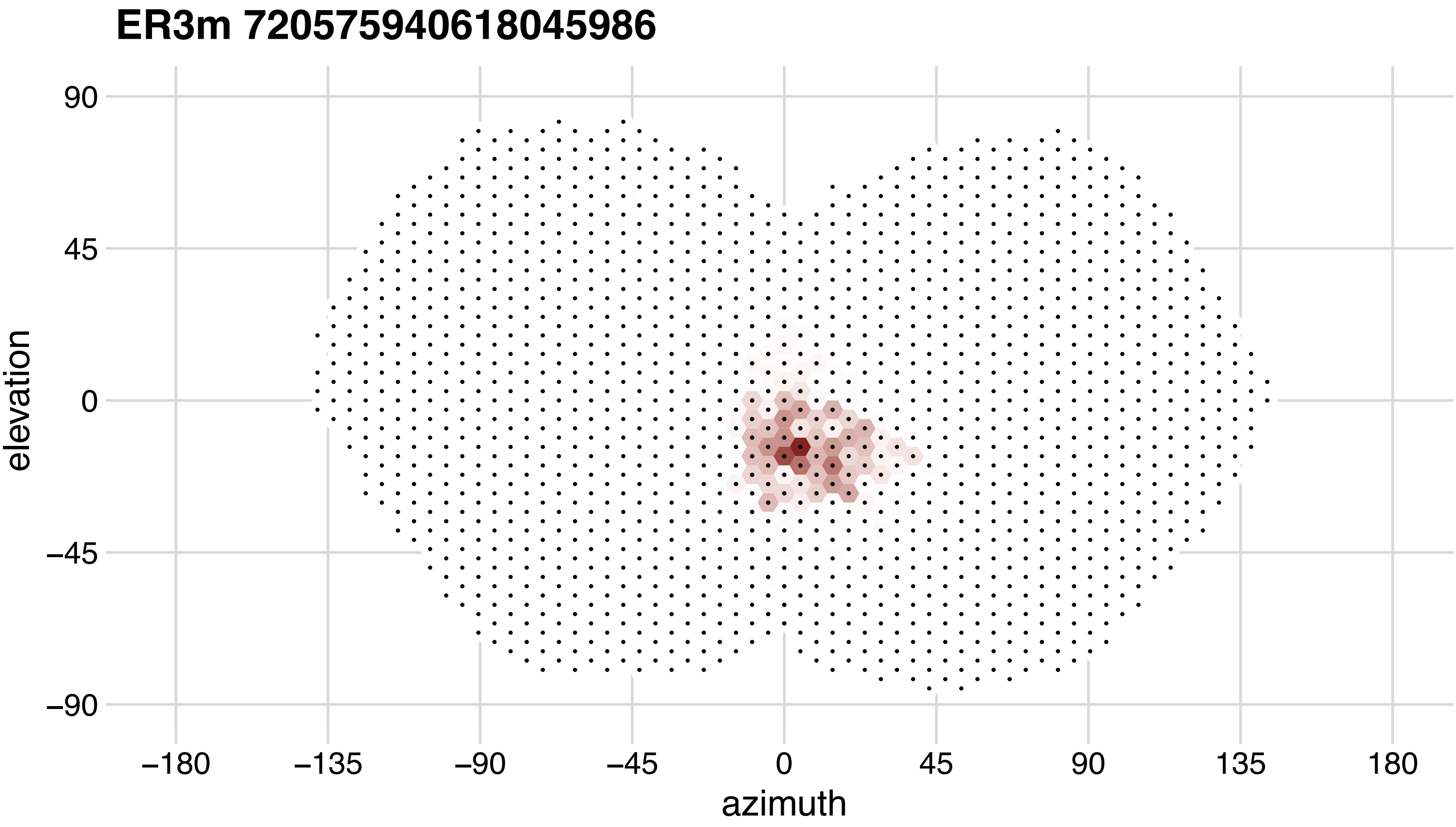
ER3m 720575940618045986
elevation
azimuth
90
45
0
−45
−90
−180
−135
−90
−45
0
45
90
135
180

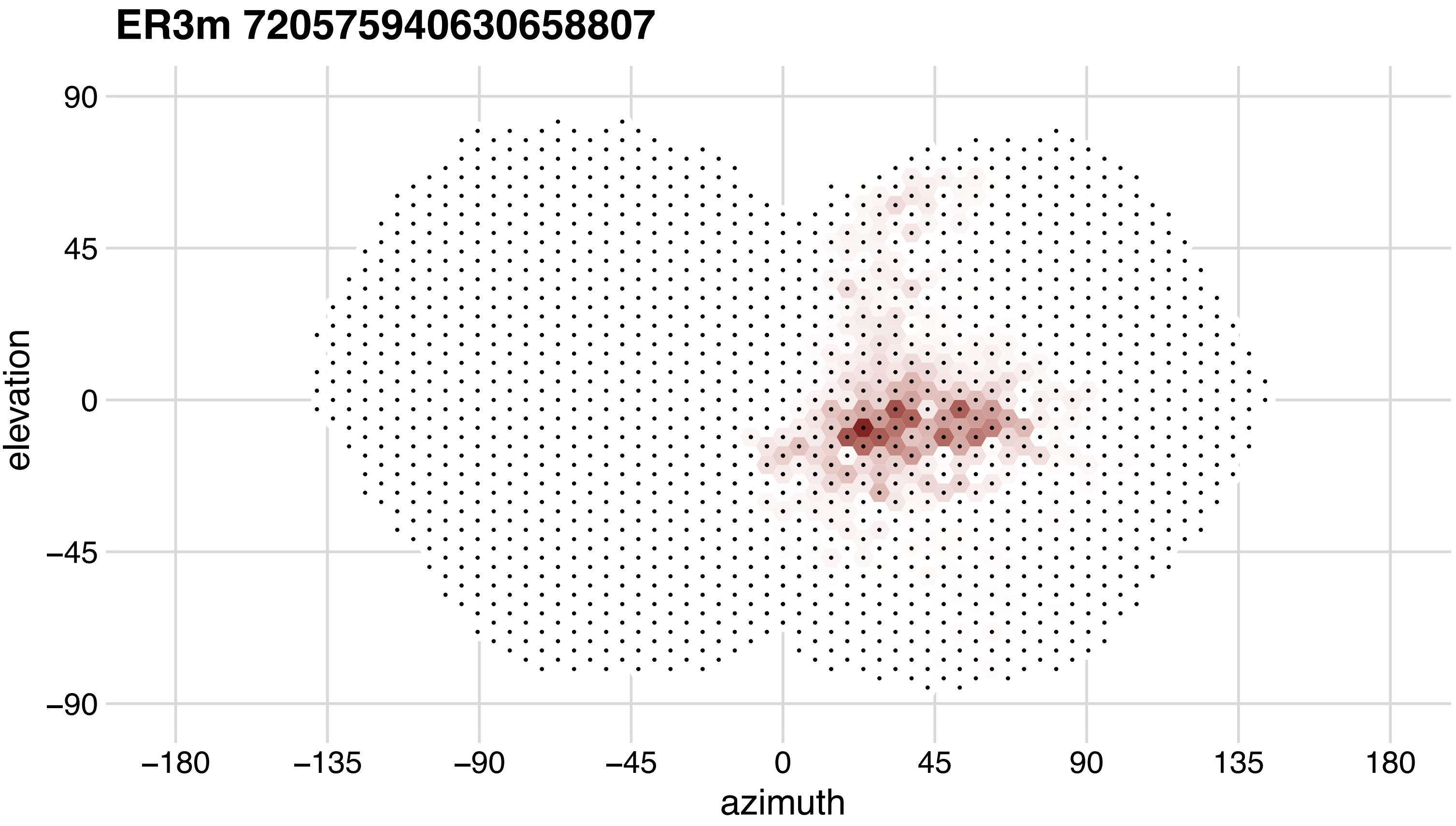
ER3m 720575940630658807
90
45
0
−45
−90
elevation
−180
−135
−90
−45
0
45
90
135
180
azimuth

## ER3p_ab 720575940613700674

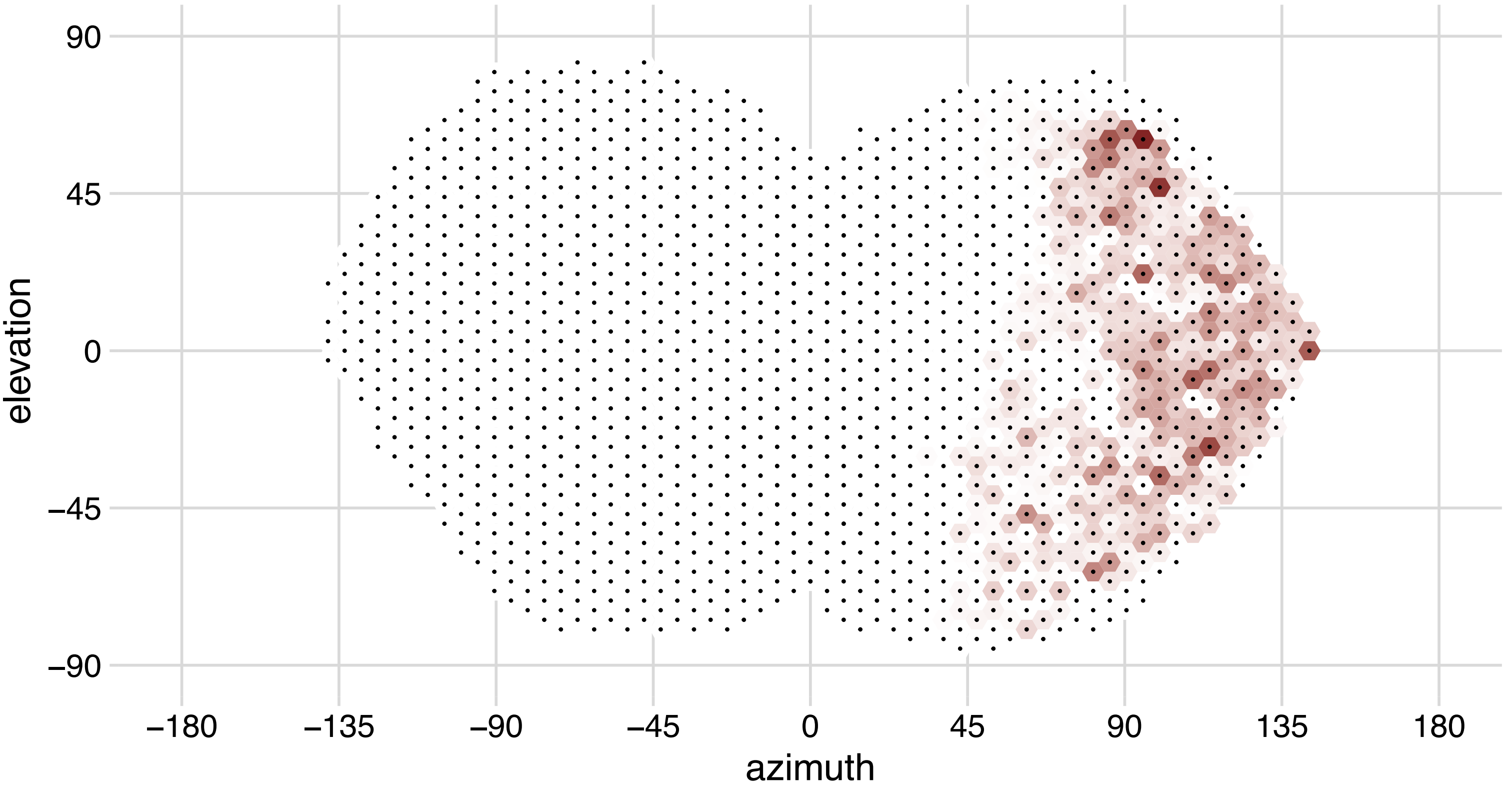

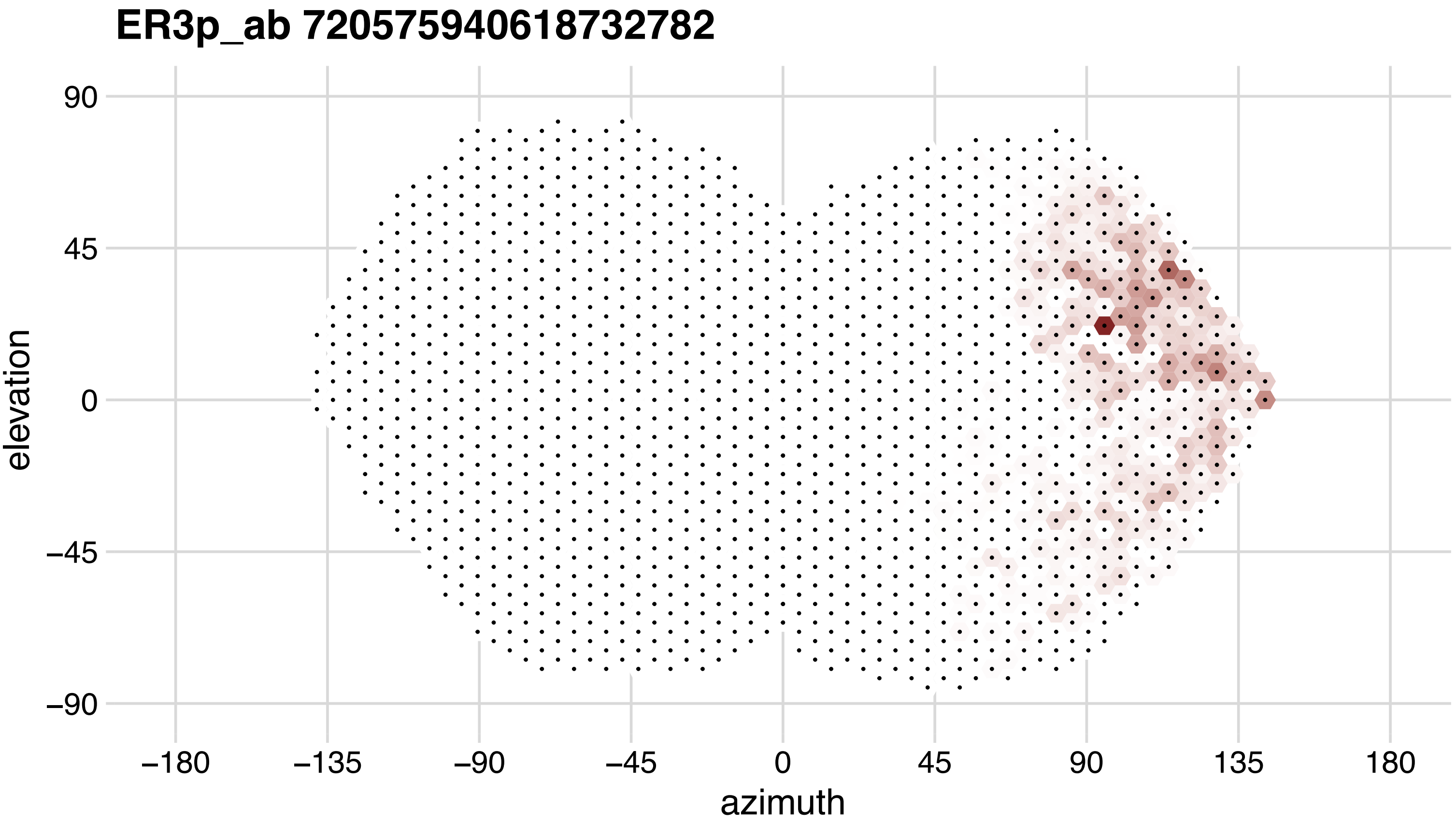
ER3p_ab 720575940618732782
90
45
0
−45
−90
elevation
−180
−135
−90
−45
0
45
90
135
180
azimuth

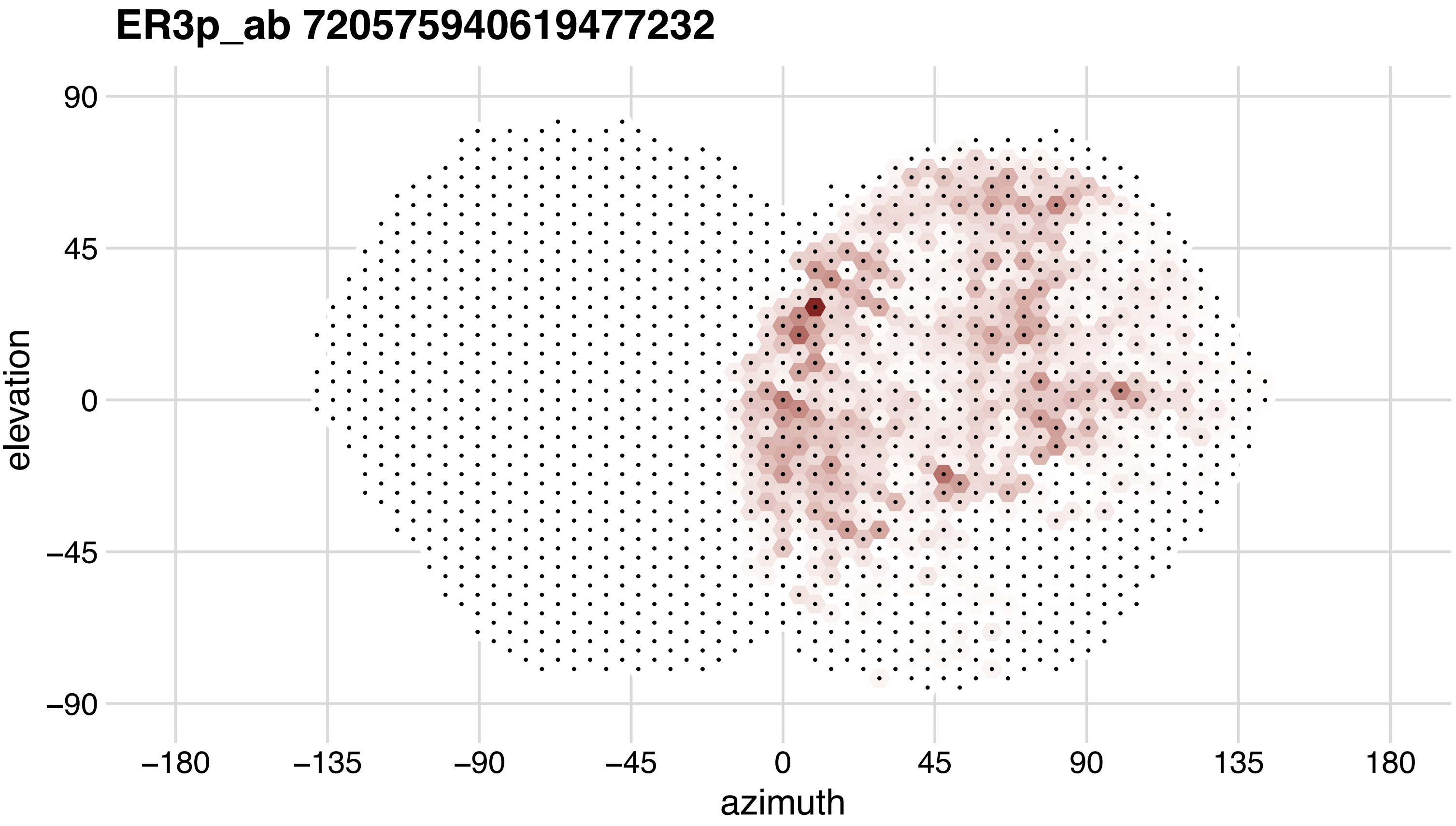
ER3p_ab 720575940619477232
90
45
0
−45
−90
elevation
−180
−135
−90
−45
0
45
90
135
180
azimuth

## ER3p_ab 720575940619957786

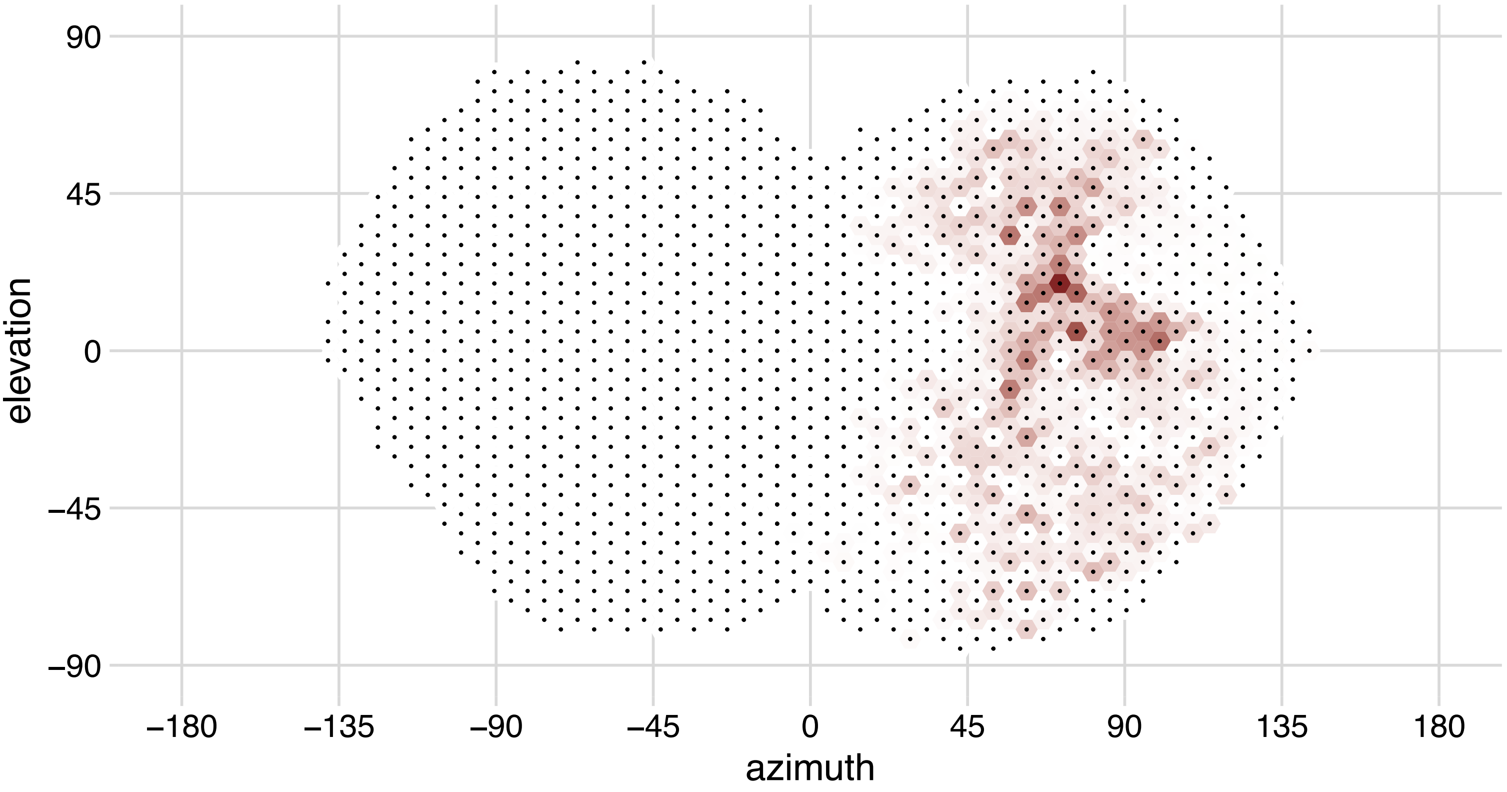

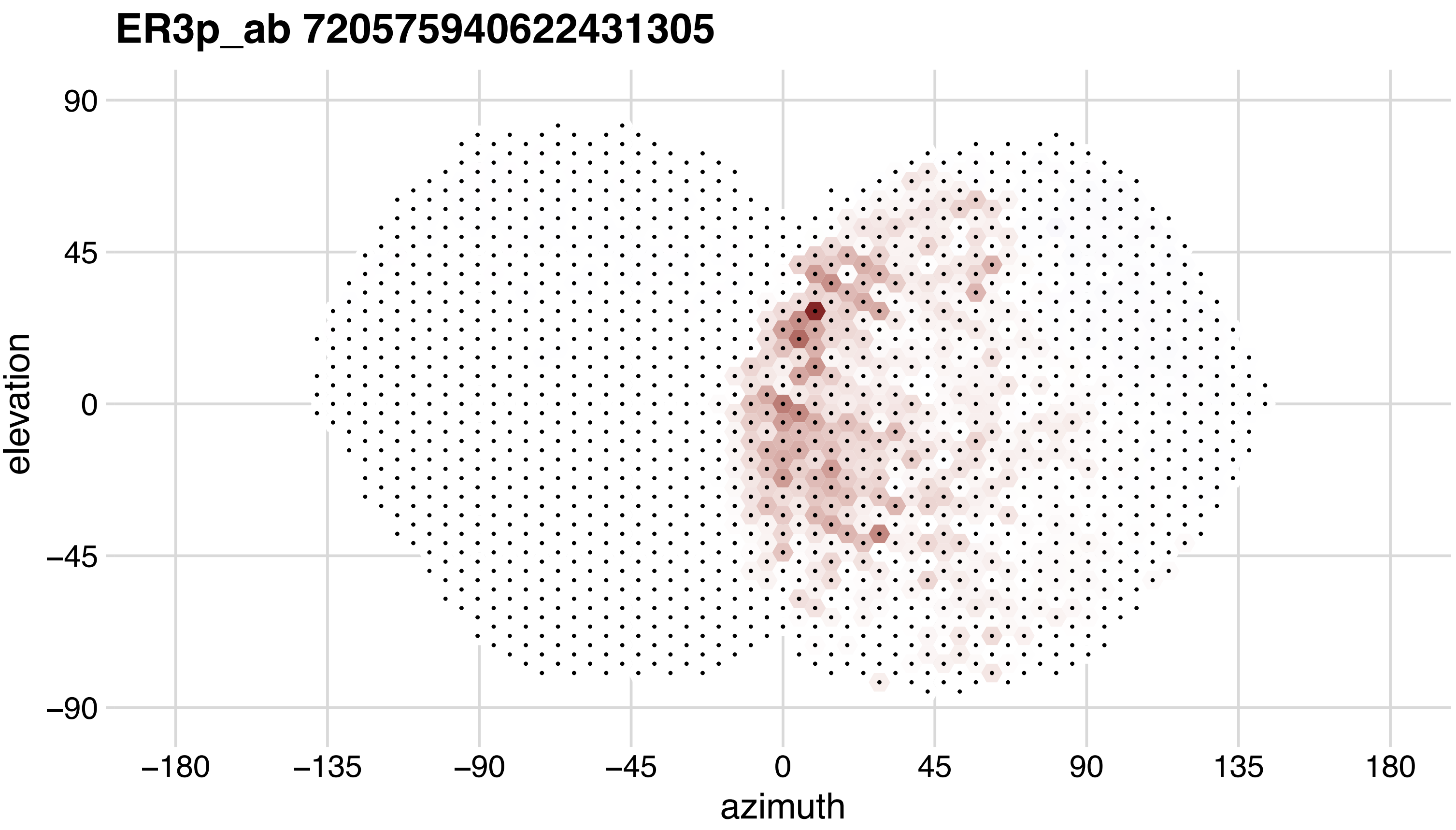
ER3p_ab 720575940622431305
90
45
0
−45
−90
elevation
−180
−135
−90
−45
0
45
90
135
180
azimuth

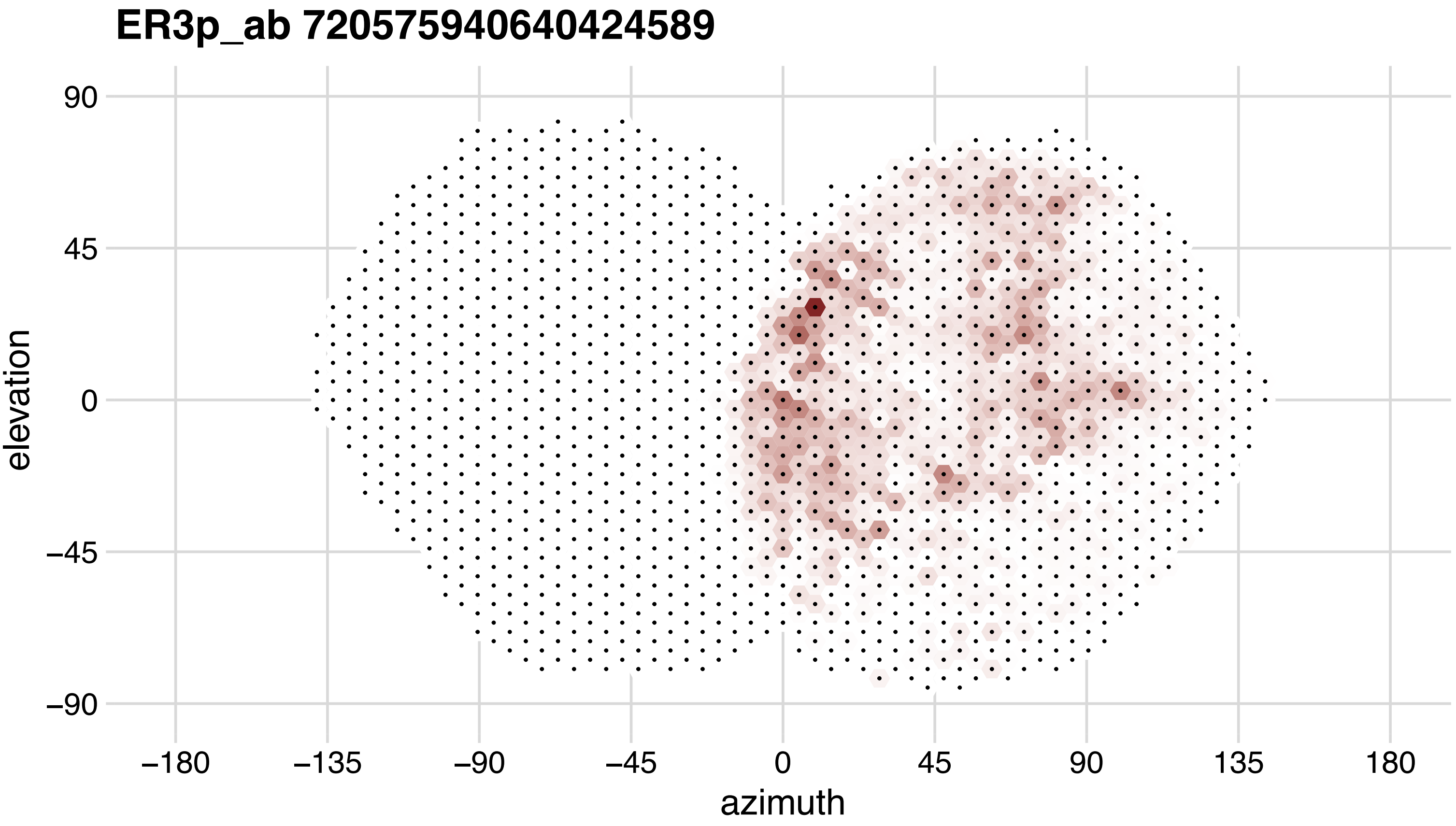
ER3p_ab 720575940640424589
elevation
90
45
0
−45
−90
azimuth
−180
−135
−90
−45
0
45
90
135
180

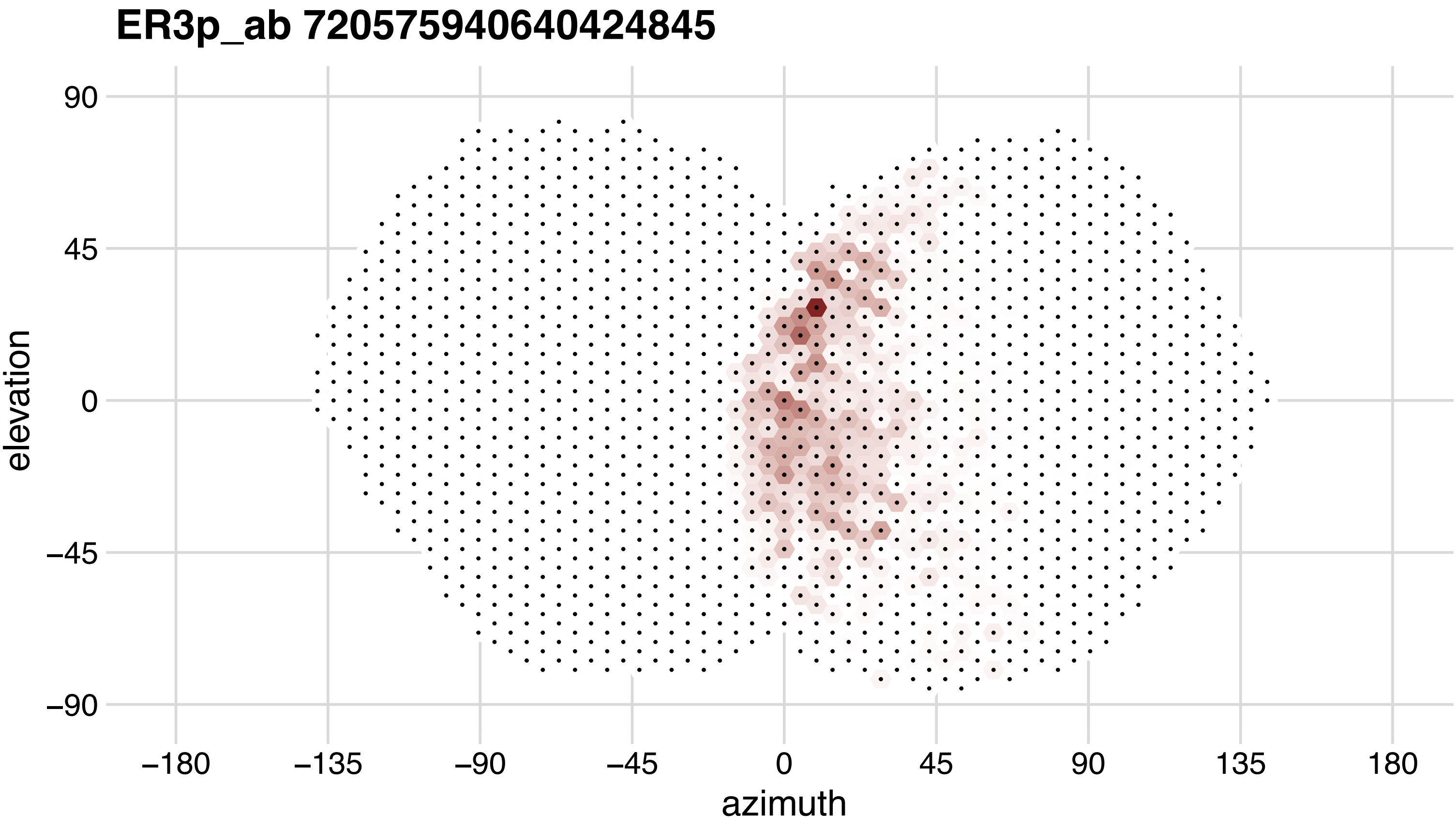
ER3p_ab 720575940640424845
90
45
0
−45
−90
elevation
−180
−135
−90
−45
0
45
90
135
180
azimuth

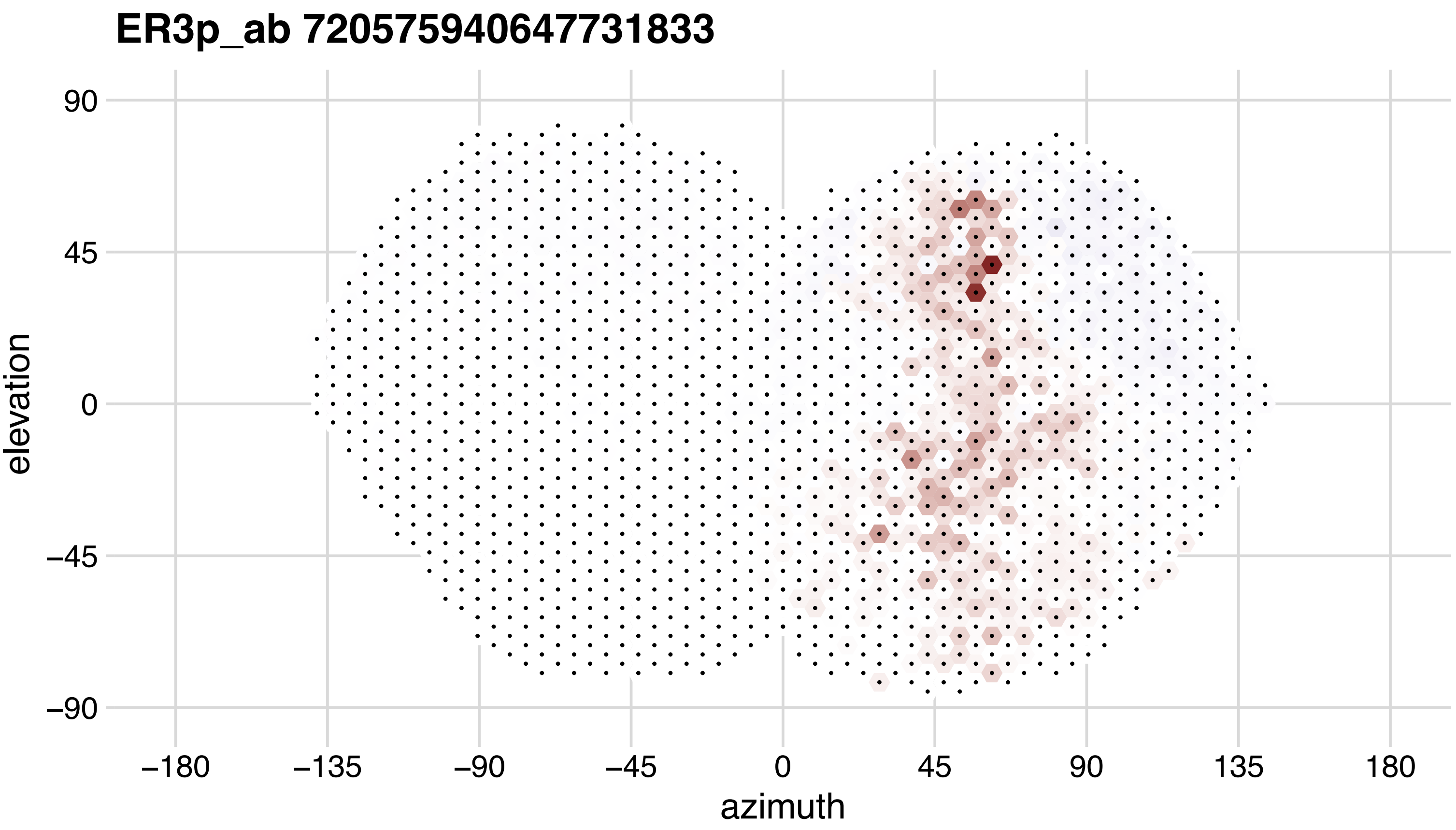
ER3p_ab 720575940647731833
90
45
0
−45
−90
elevation
−180
−135
−90
−45
0
45
90
135
180
azimuth

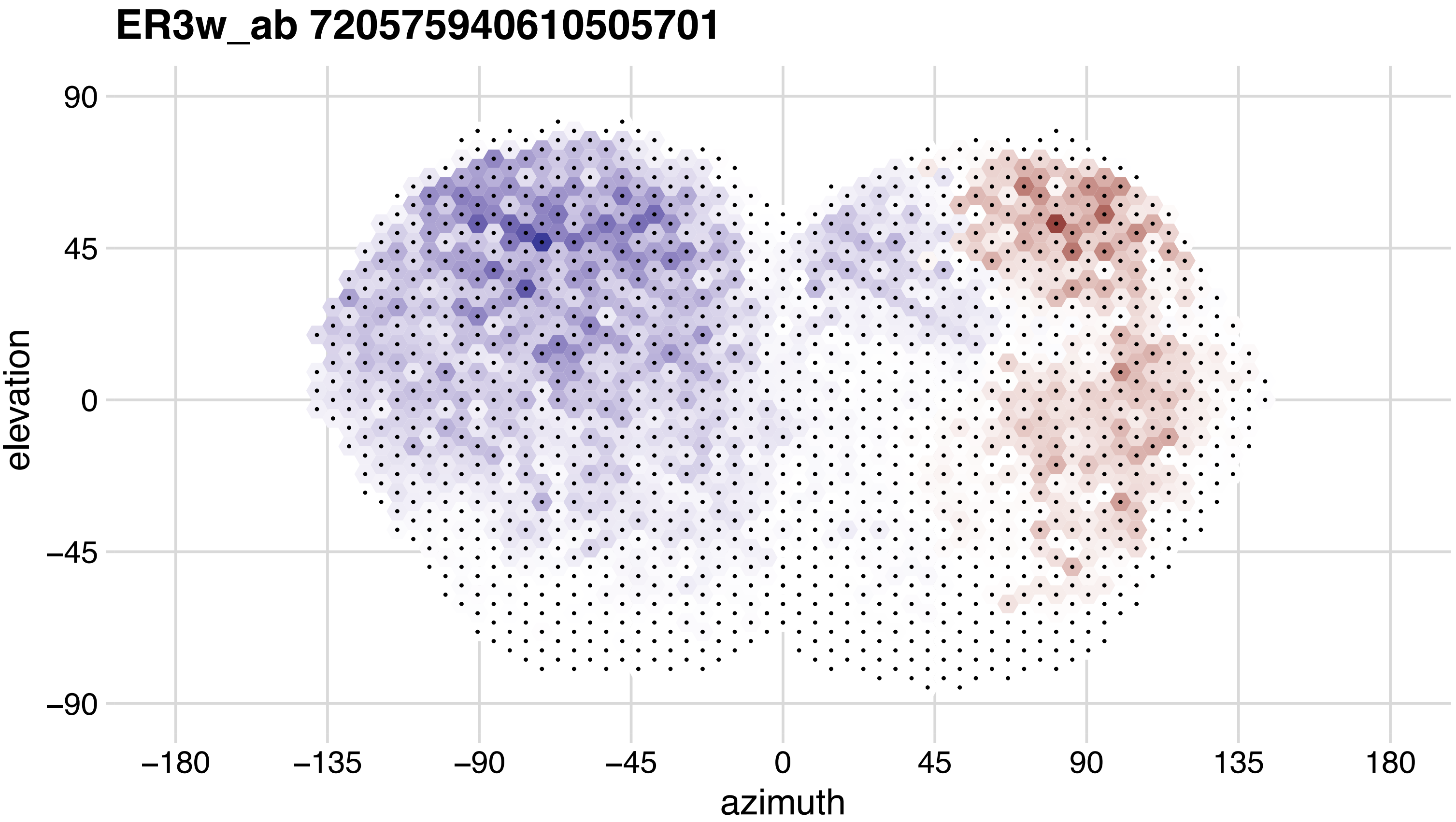
ER3w_ab 720575940610505701
90
45
0
−45
−90
elevation
−180
−135
−90
−45
0
45
90
135
180
azimuth

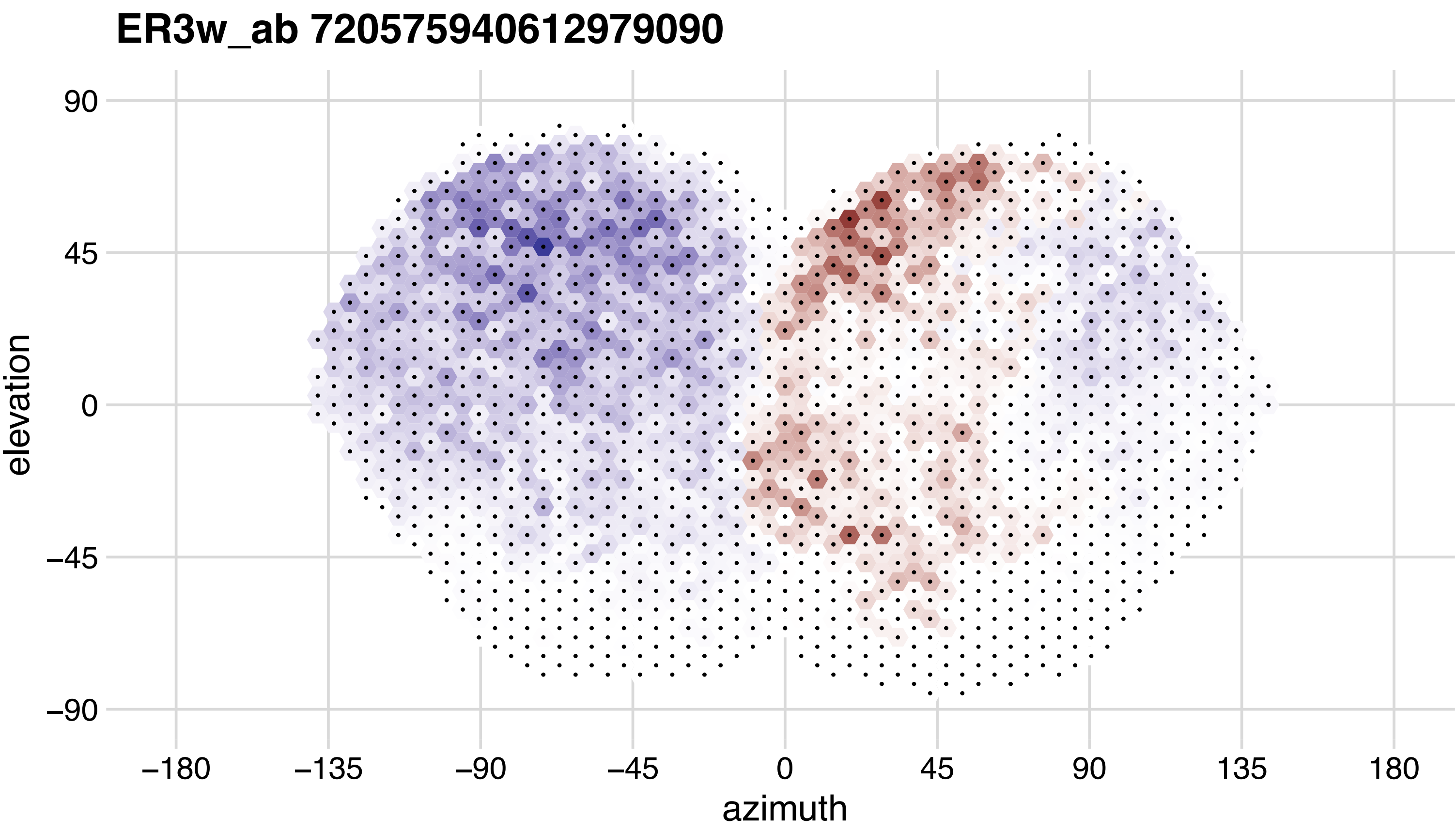
ER3w_ab 720575940612979090
90
45
0
−45
−90
elevation
−180
−135
−90
−45
0
45
90
135
180
azimuth

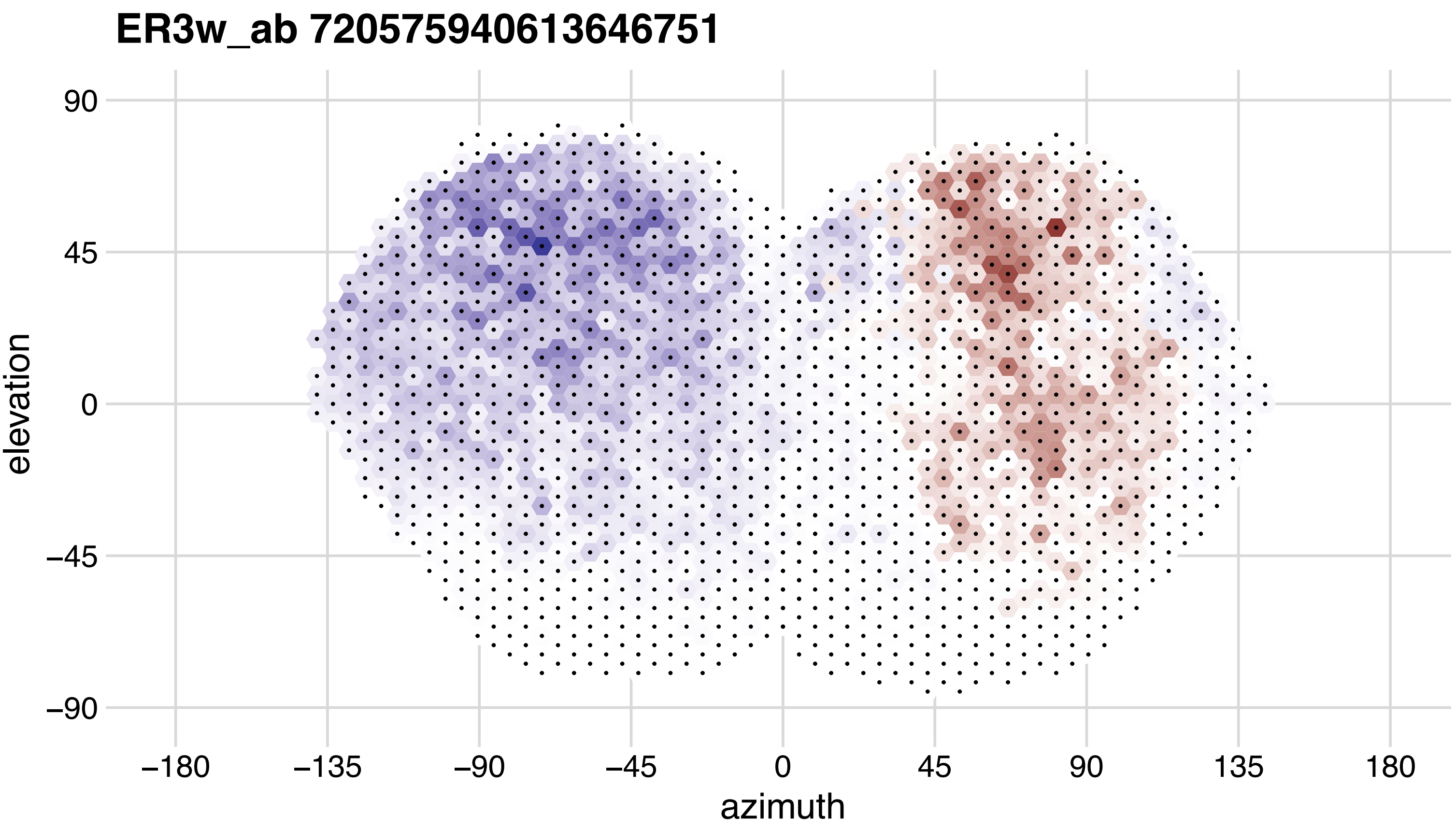
ER3w_ab 720575940613646751
90
45
0
−45
−90
elevation
−180
−135
−90
−45
0
45
90
135
180
azimuth

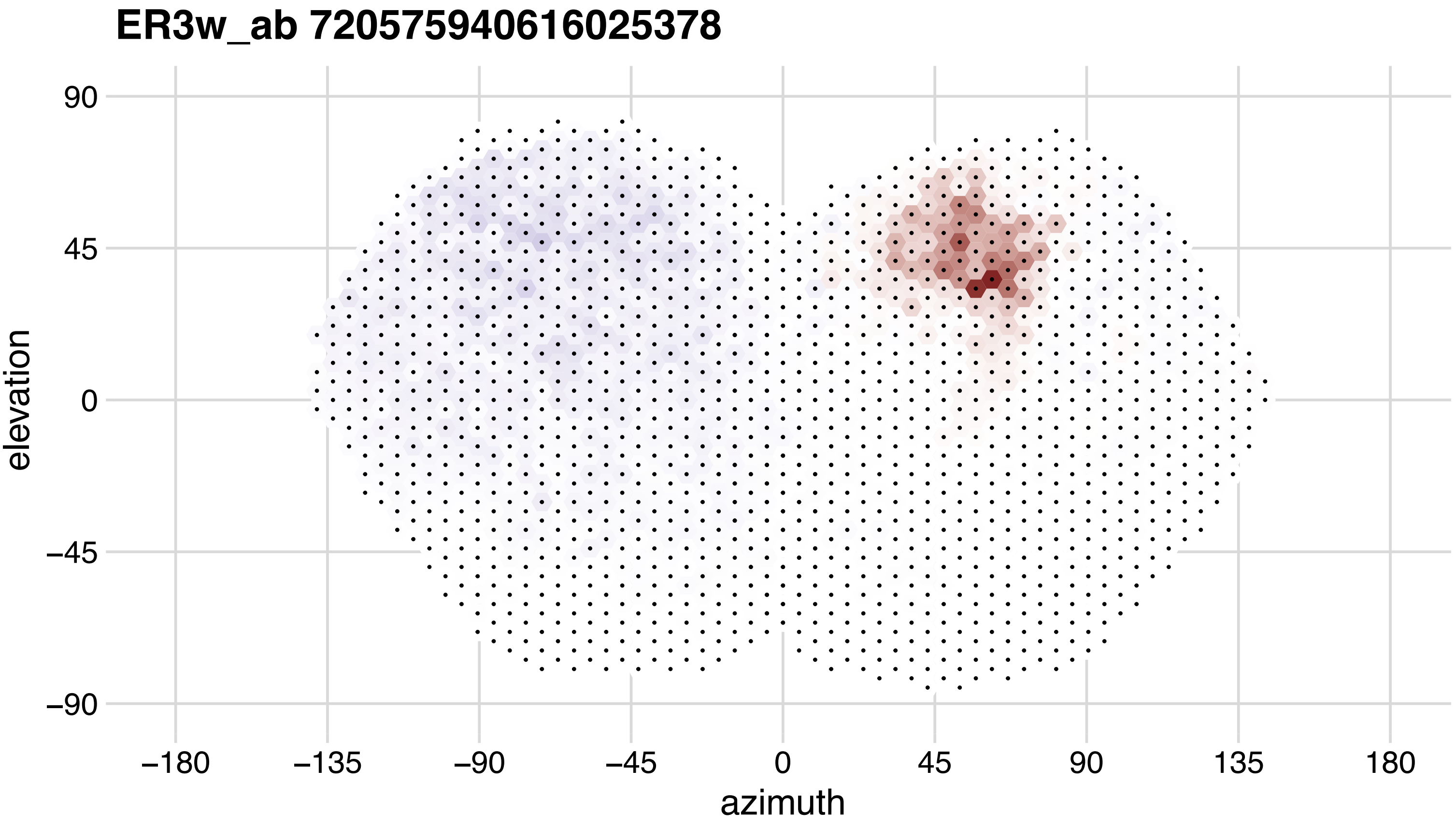
ER3w_ab 720575940616025378
elevation
azimuth
90
45
0
−45
−90
−180
−135
−90
−45
0
45
90
135
180

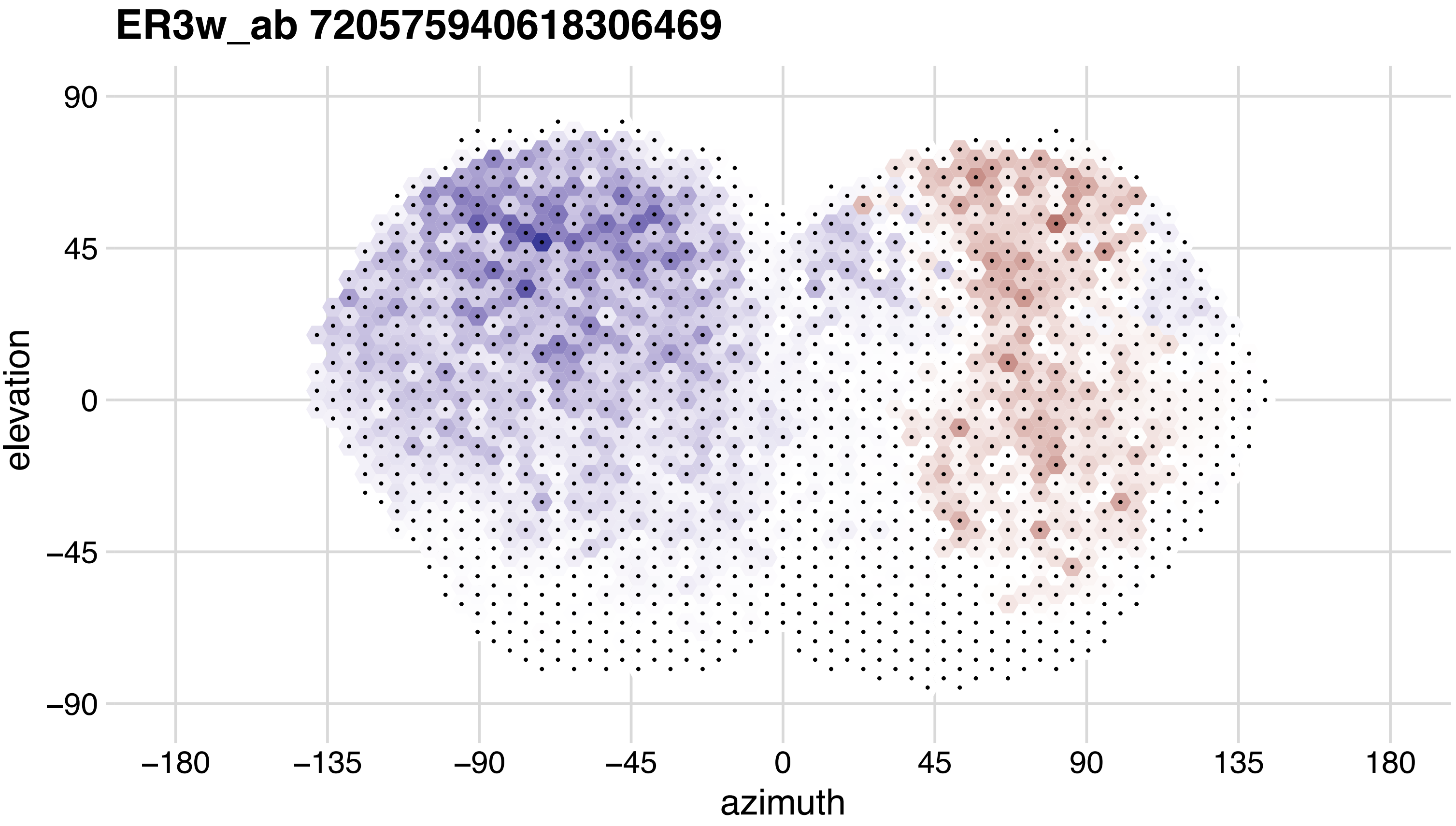
ER3w_ab 720575940618306469
90
45
0
−45
−90
elevation
−180
−135
−90
−45
0
45
90
135
180
azimuth

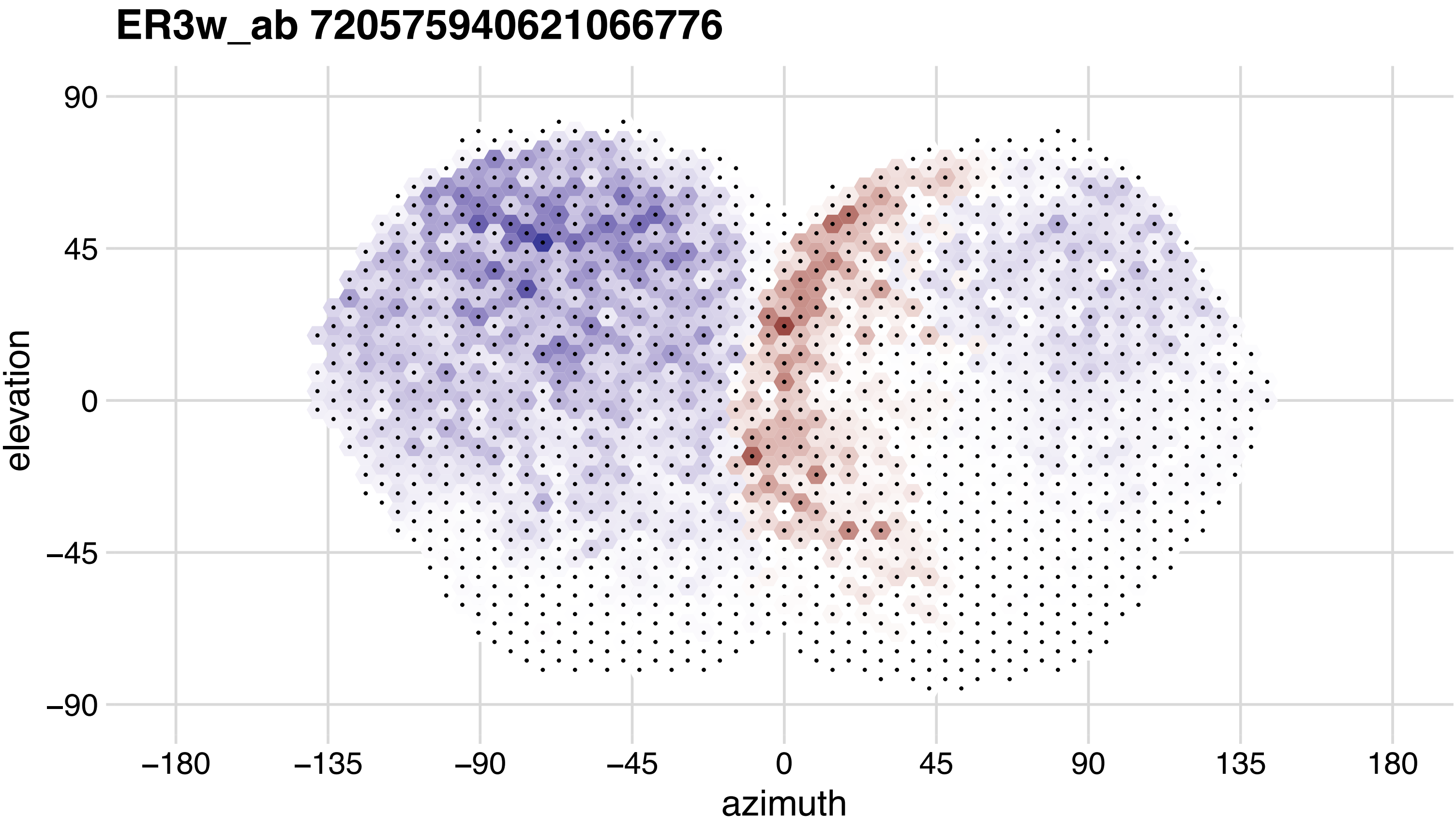
ER3w_ab 720575940621066776
90
45
0
−45
−90
elevation
−180
−135
−90
−45
0
45
90
135
180
azimuth

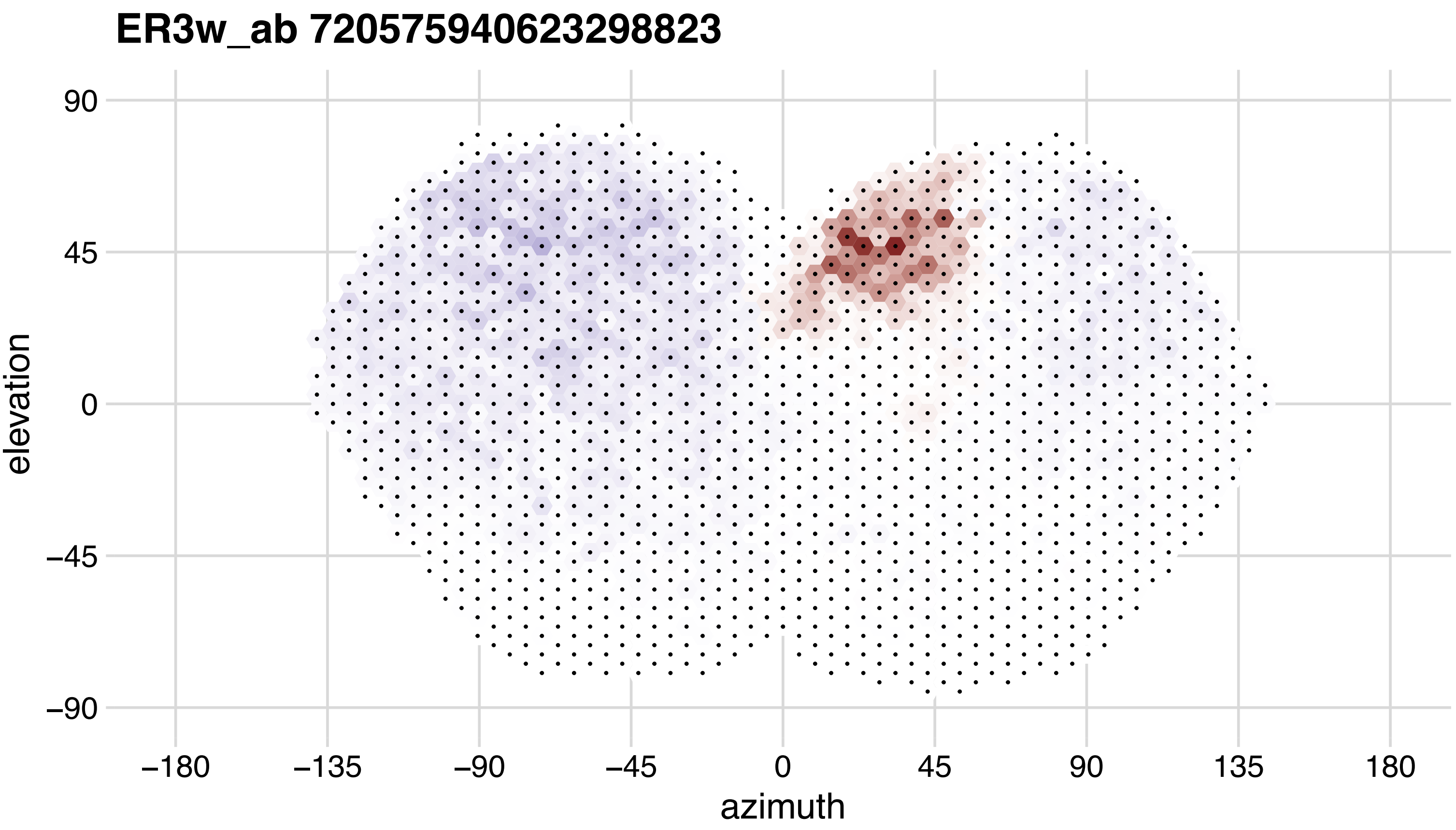
ER3w_ab 720575940623298823
elevation
azimuth
90
45
0
−45
−90
−180
−135
−90
−45
0
45
90
135
180

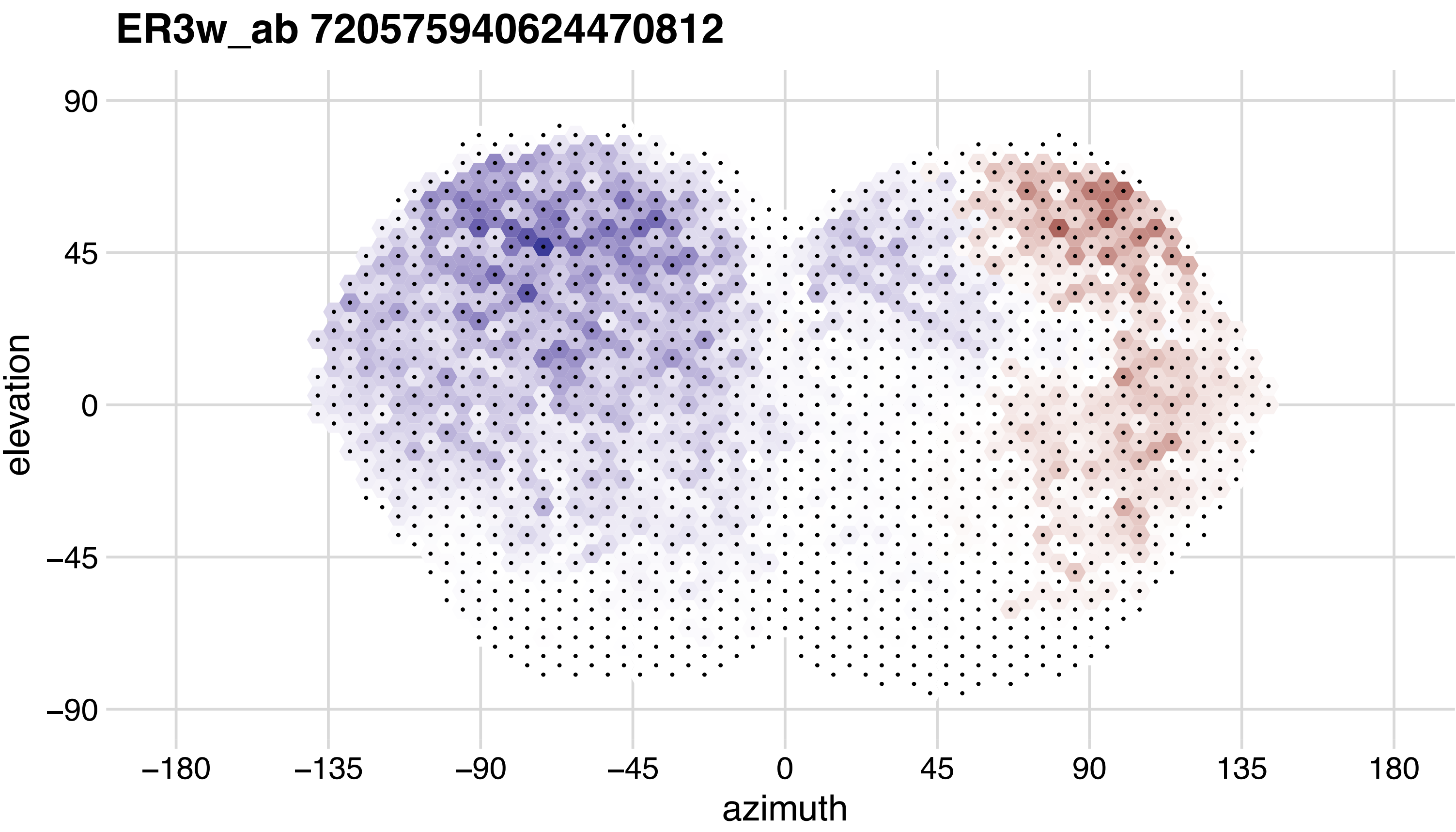

ER3w_ab 720575940624470812
90
45
0
−45
−90
elevation
−180
−135
−90
−45
0
45
90
135
180
azimuth

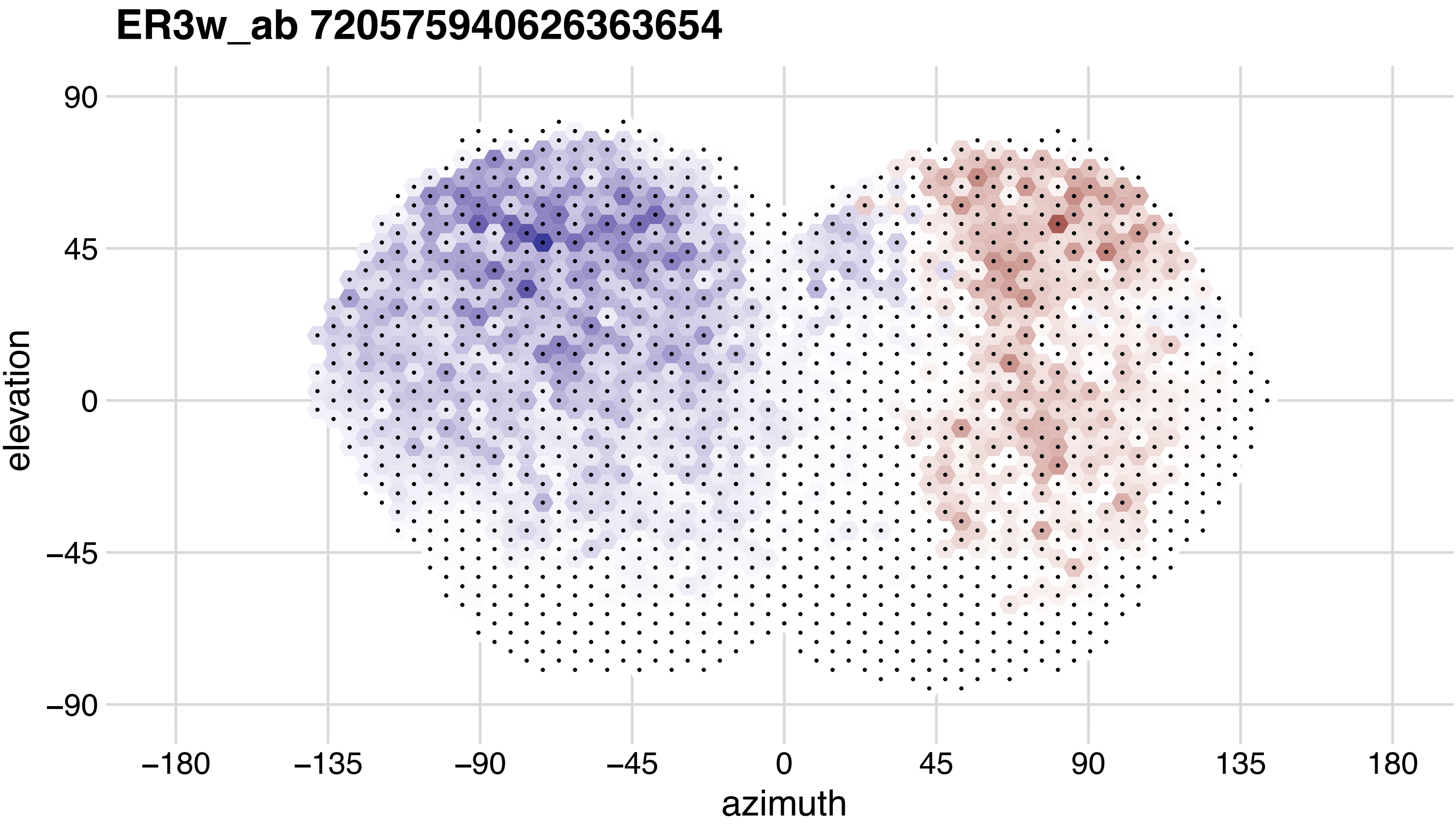
ER3w_ab 720575940626363654
90
45
0
−45
−90
elevation
−180
−135
−90
−45
0
45
90
135
180
azimuth

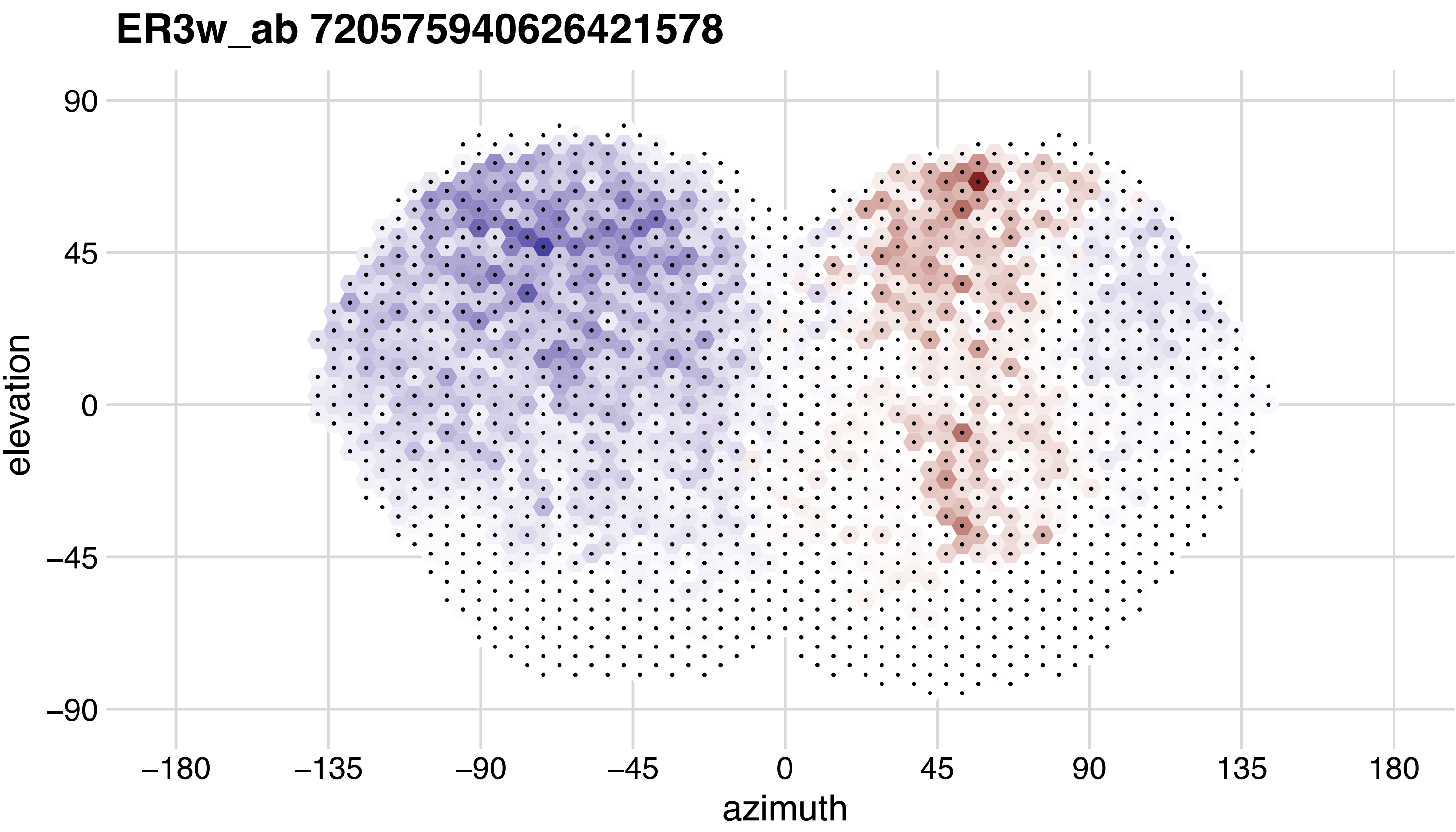

ER3w_ab 720575940626421578
90
45
0
−45
−90
elevation
−180
−135
−90
−45
0
45
90
135
180
azimuth

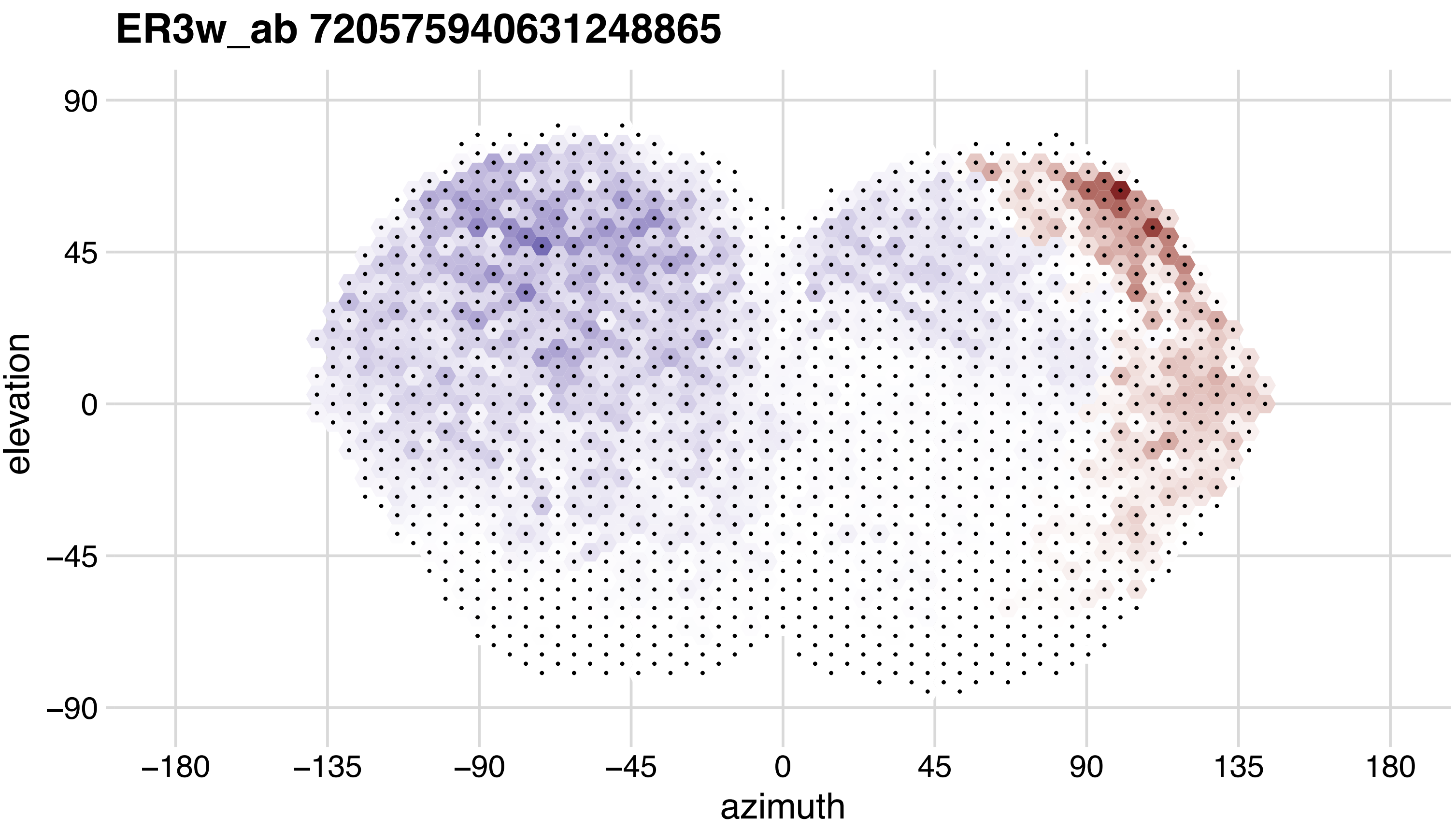
ER3w_ab 720575940631248865
90
45
0
−45
−90
elevation
−180
−135
−90
−45
0
45
90
135
180
azimuth

## ER3w_ab 720575940637681267

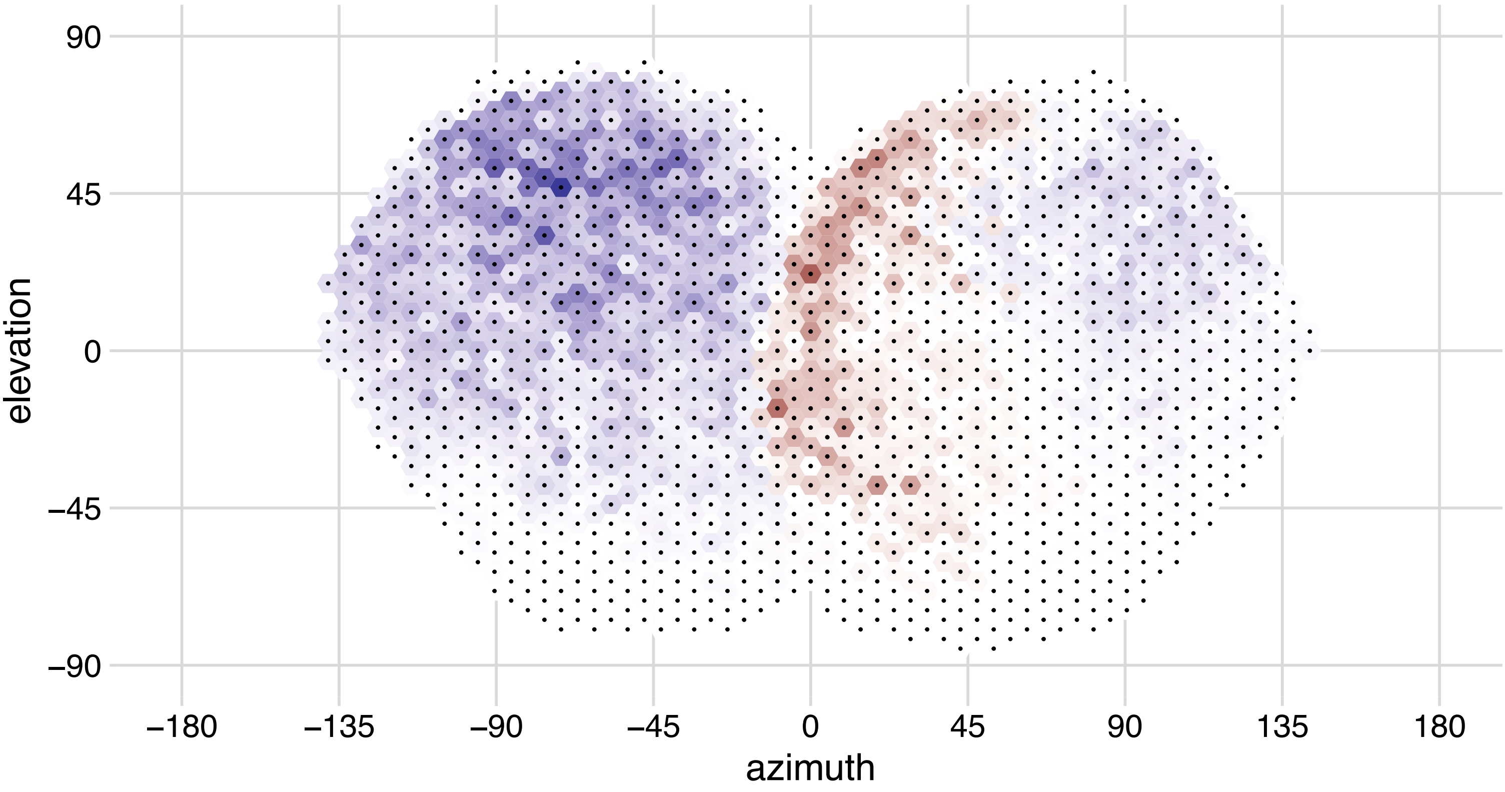

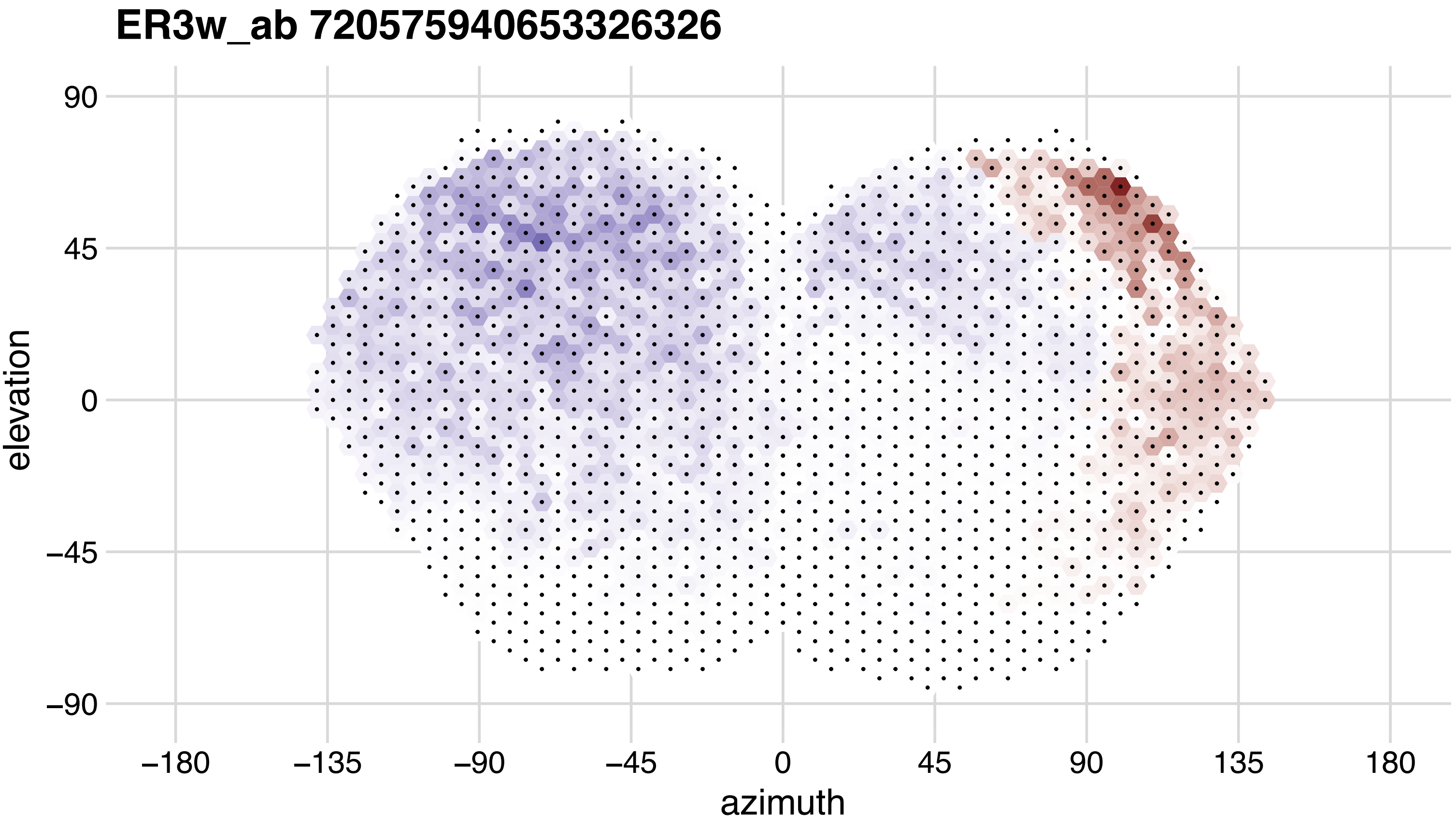
ER3w_ab 720575940653326326
90
45
0
−45
−90
elevation
−180
−135
−90
−45
0
45
90
135
180
azimuth

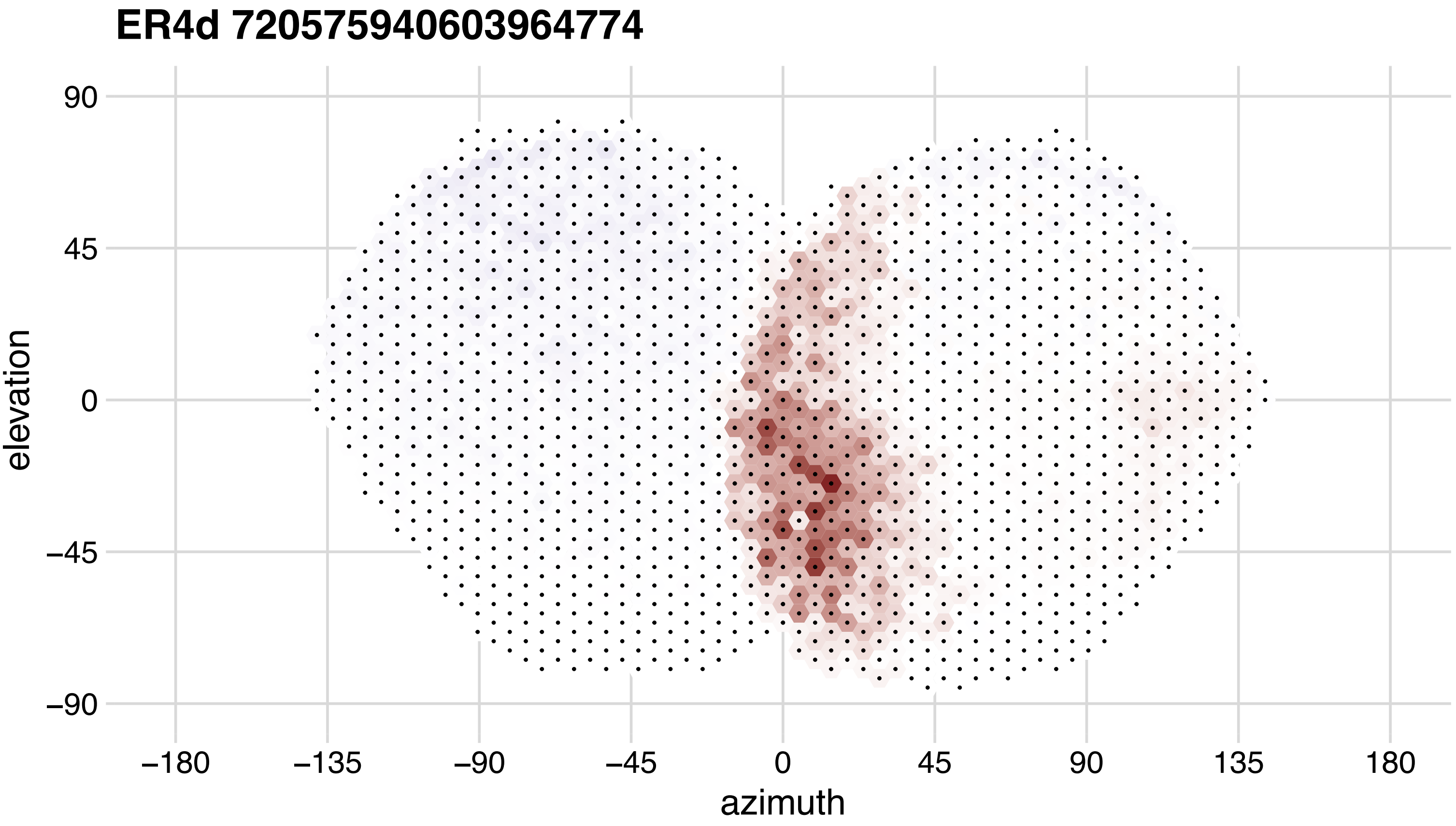
ER4d 720575940603964774
elevation
azimuth
90
45
0
−45
−90
−180
−135
−90
−45
0
45
90
135
180

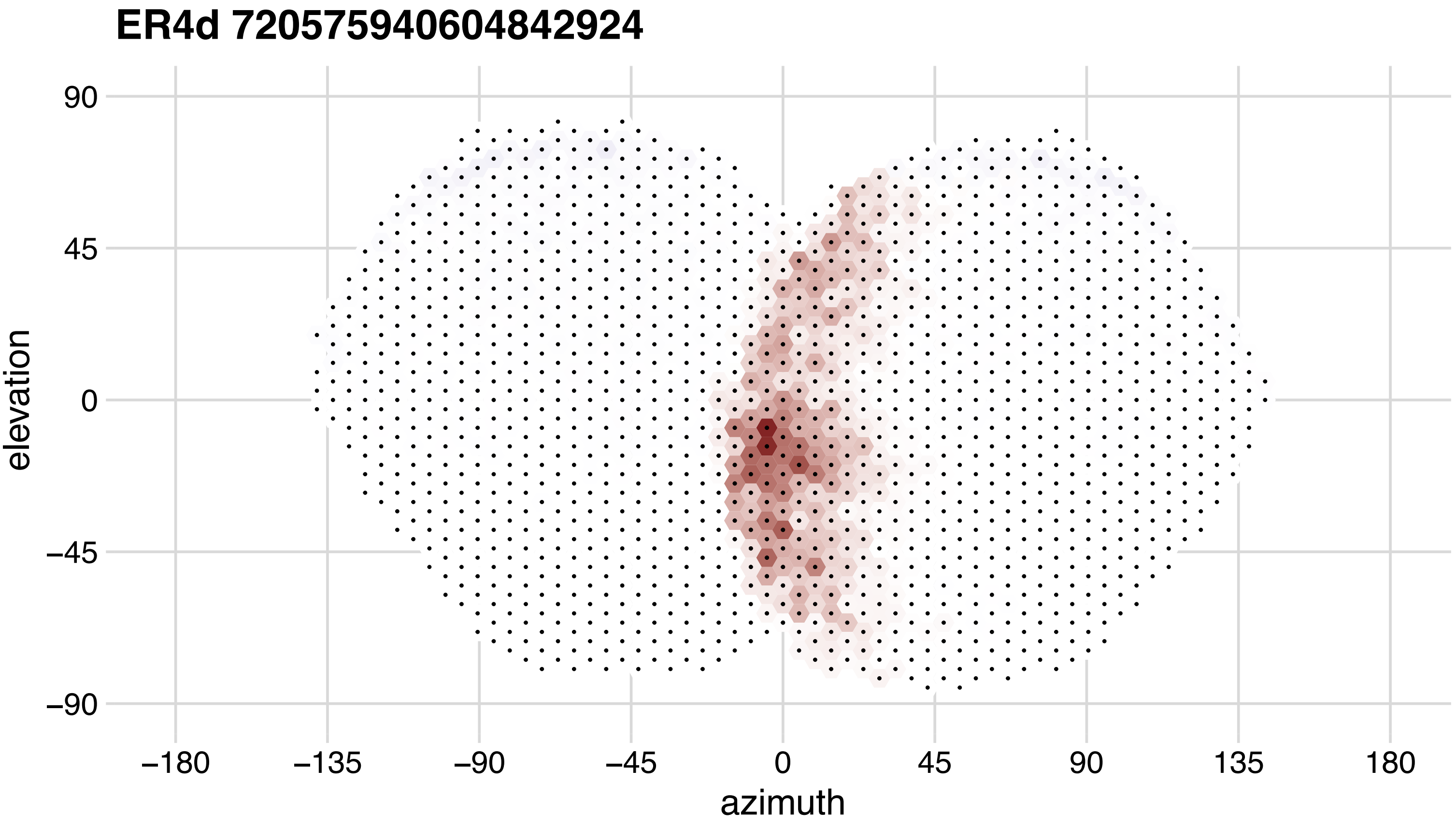

ER4d 720575940604842924
90
45
0
−45
−90
elevation
−180
−135
−90
−45
0
45
90
135
180
azimuth

# ER4d 720575940606102729

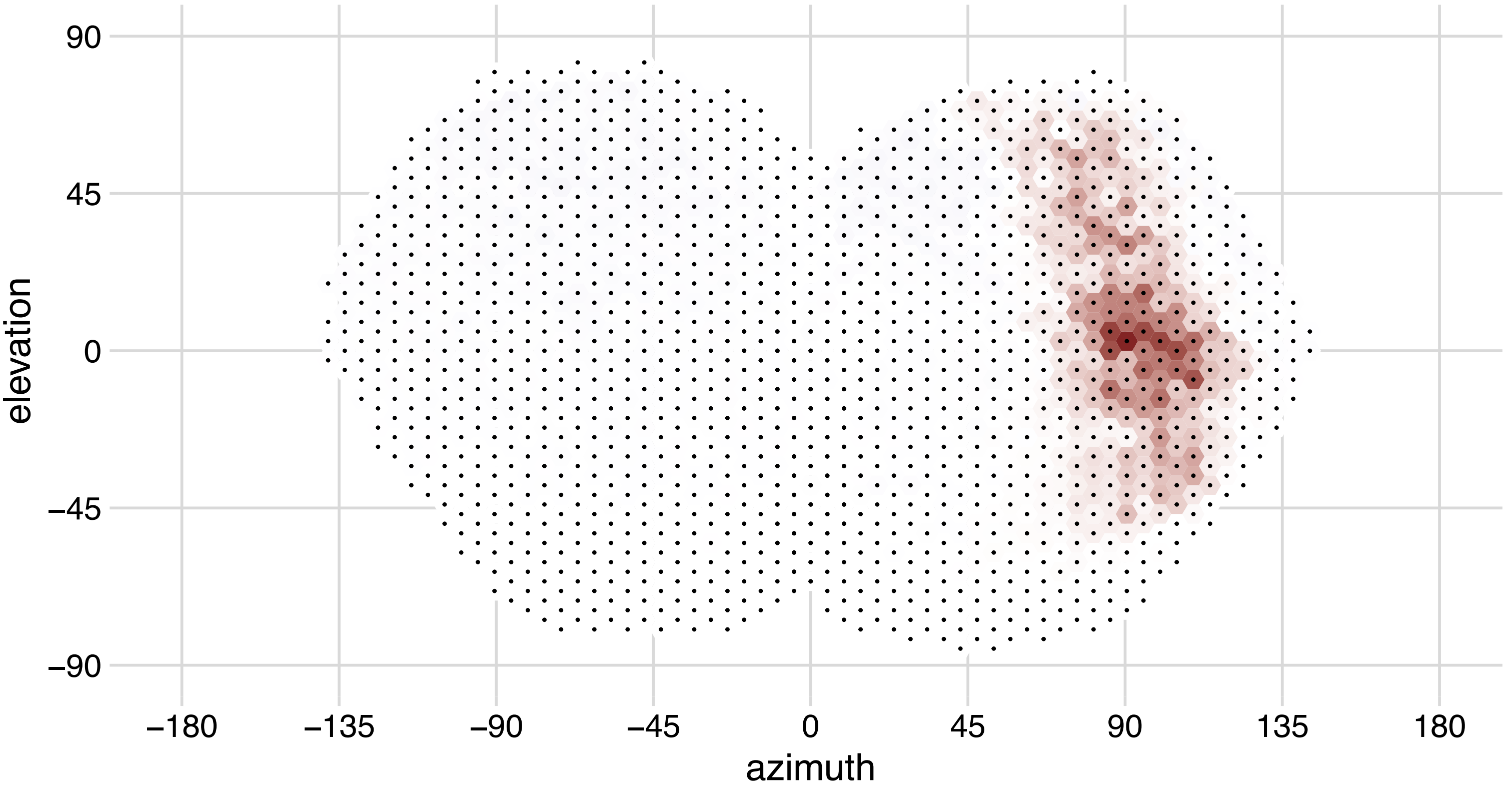

## ER4d 720575940619062357

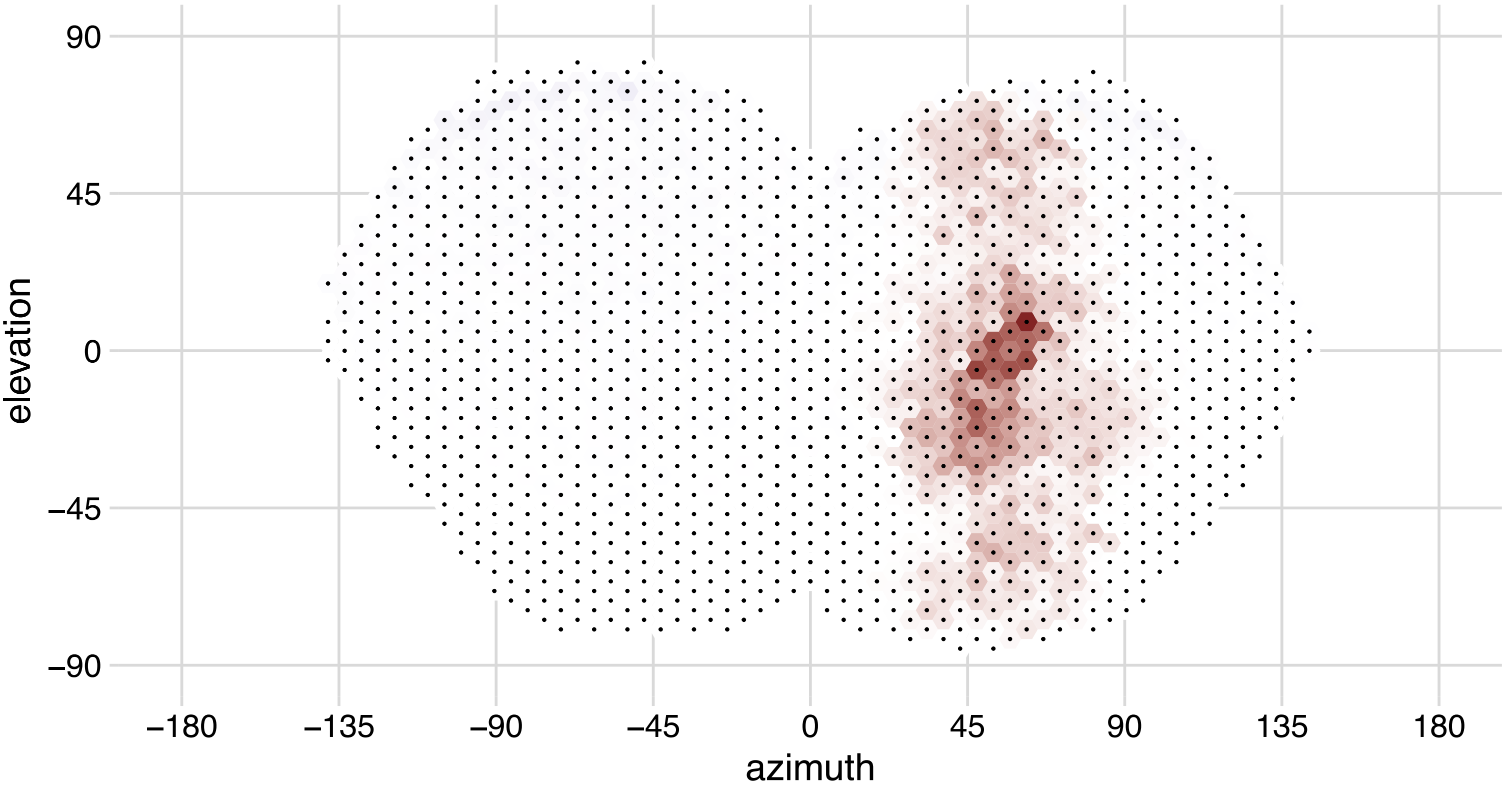

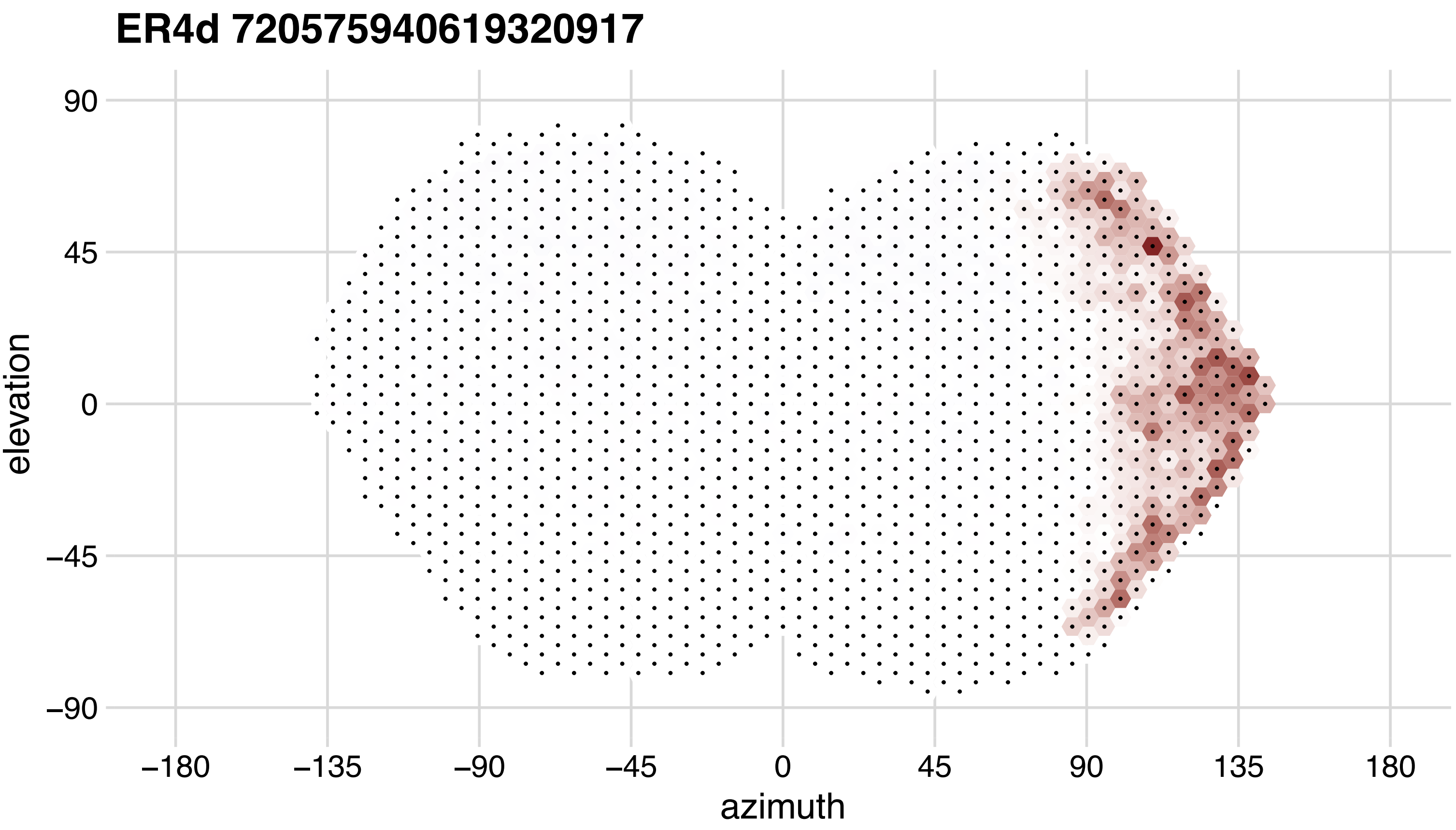

ER4d 720575940619320917
90
45
0
−45
−90
elevation
−180
−135
−90
−45
0
45
90
135
180
azimuth

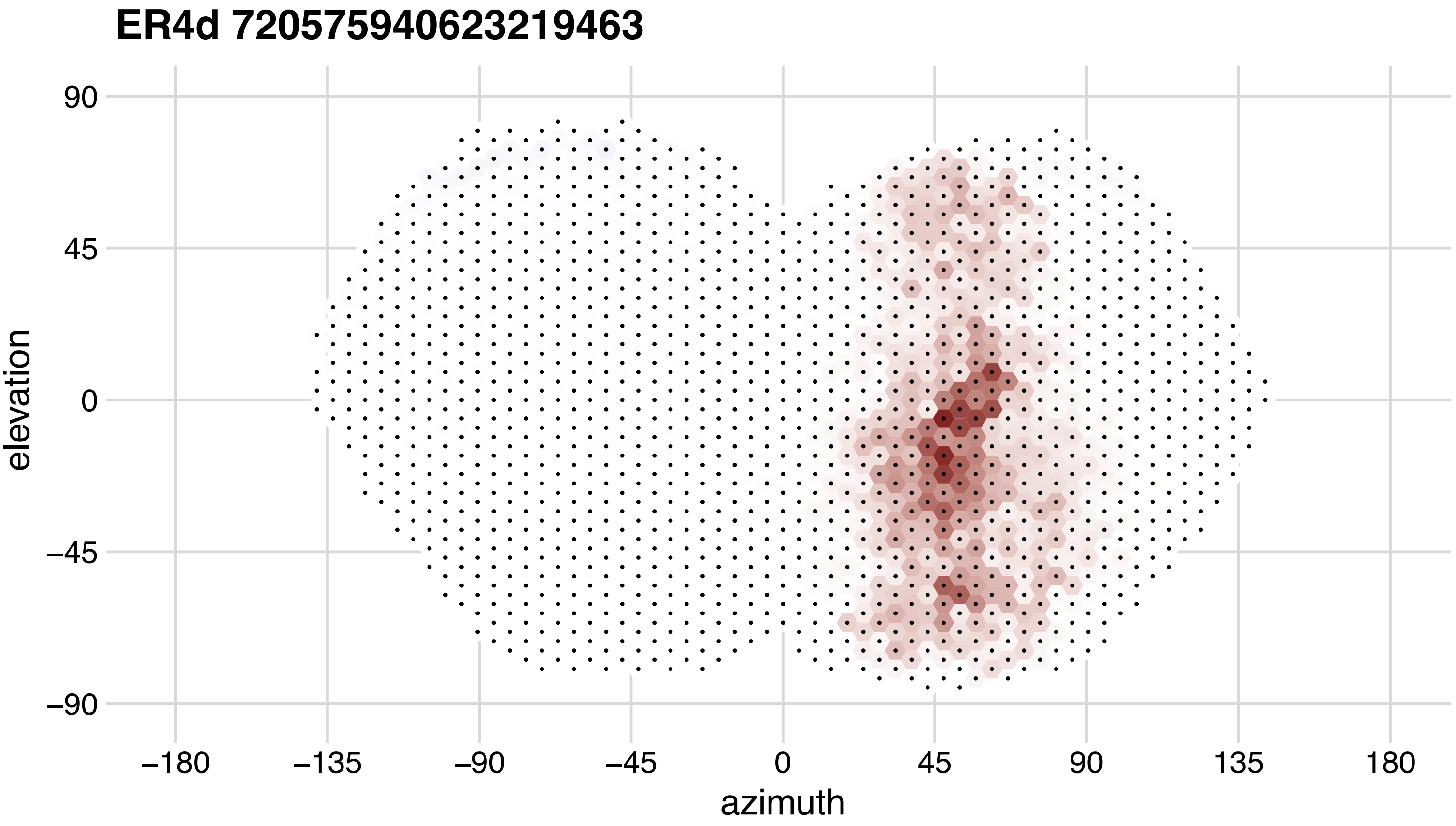

ER4d 720575940623219463
90
45
0
−45
−90
elevation
−180
−135
−90
−45
0
45
90
135
180
azimuth

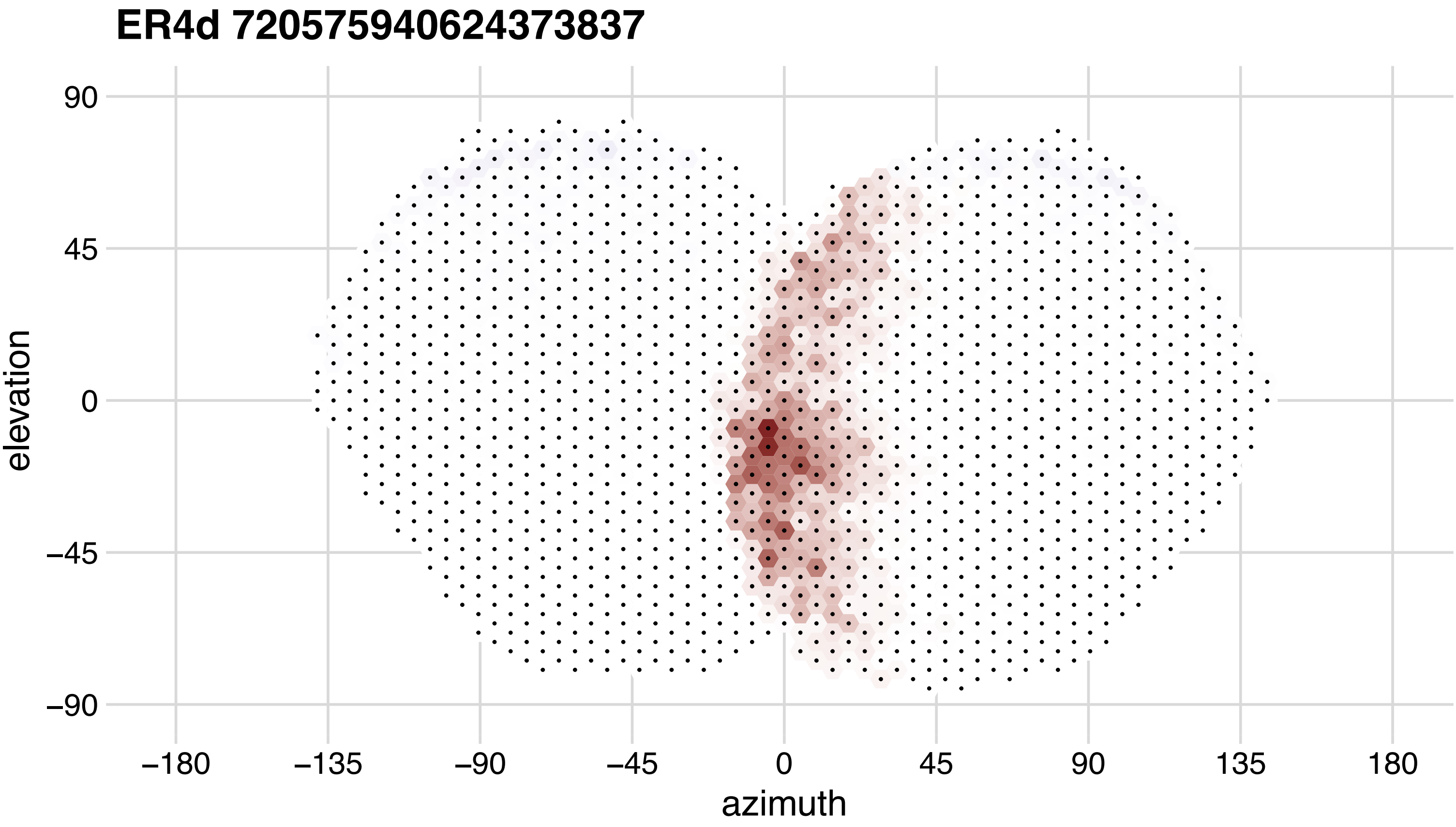
ER4d 720575940624373837
90
45
0
−45
−90
elevation
−180
−135
−90
−45
0
45
90
135
180
azimuth

## ER4d 720575940625511762

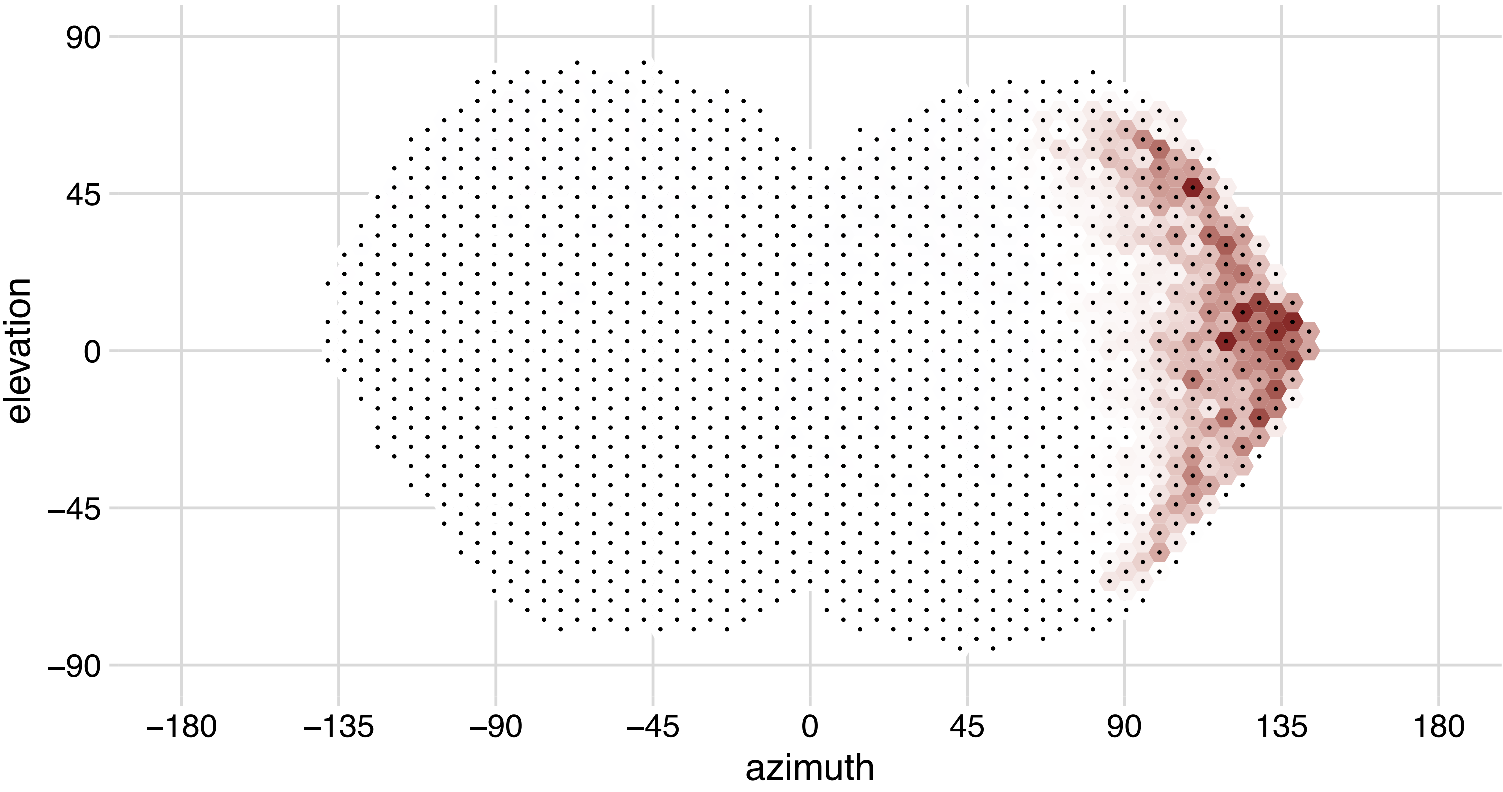

## ER4d 720575940629720848

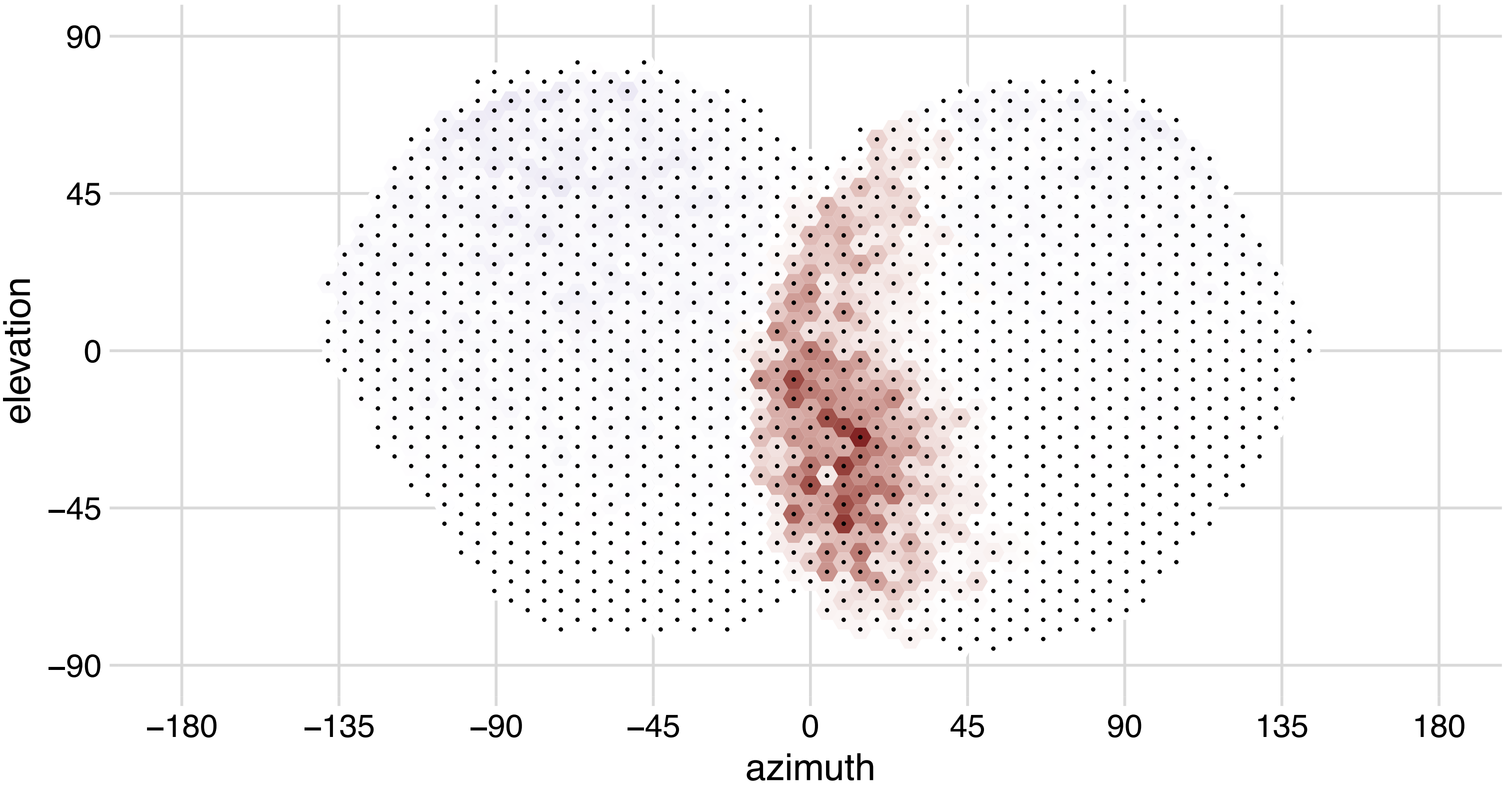

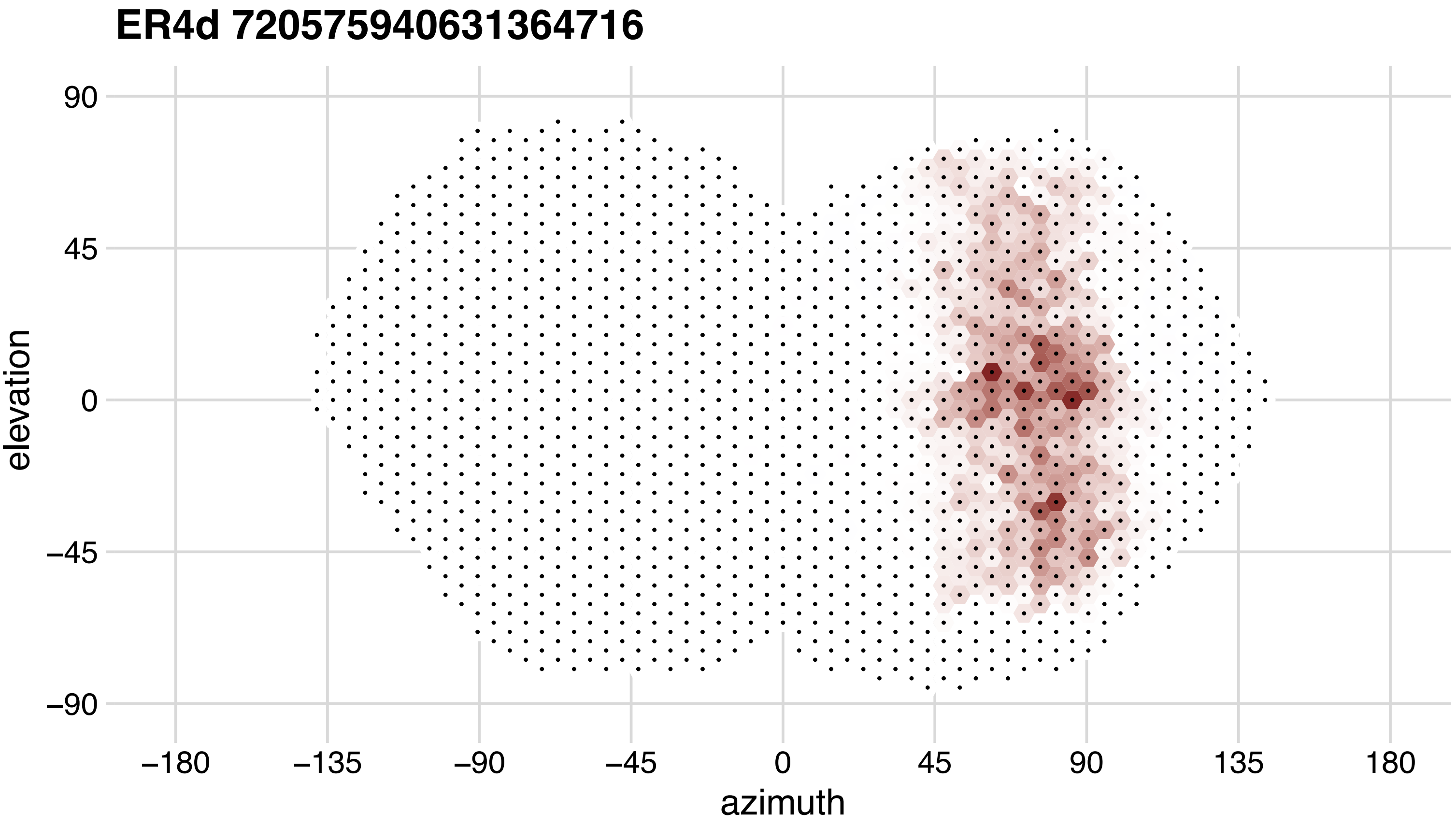
ER4d 720575940631364716
90
45
0
−45
−90
elevation
−180
−135
−90
−45
0
45
90
135
180
azimuth

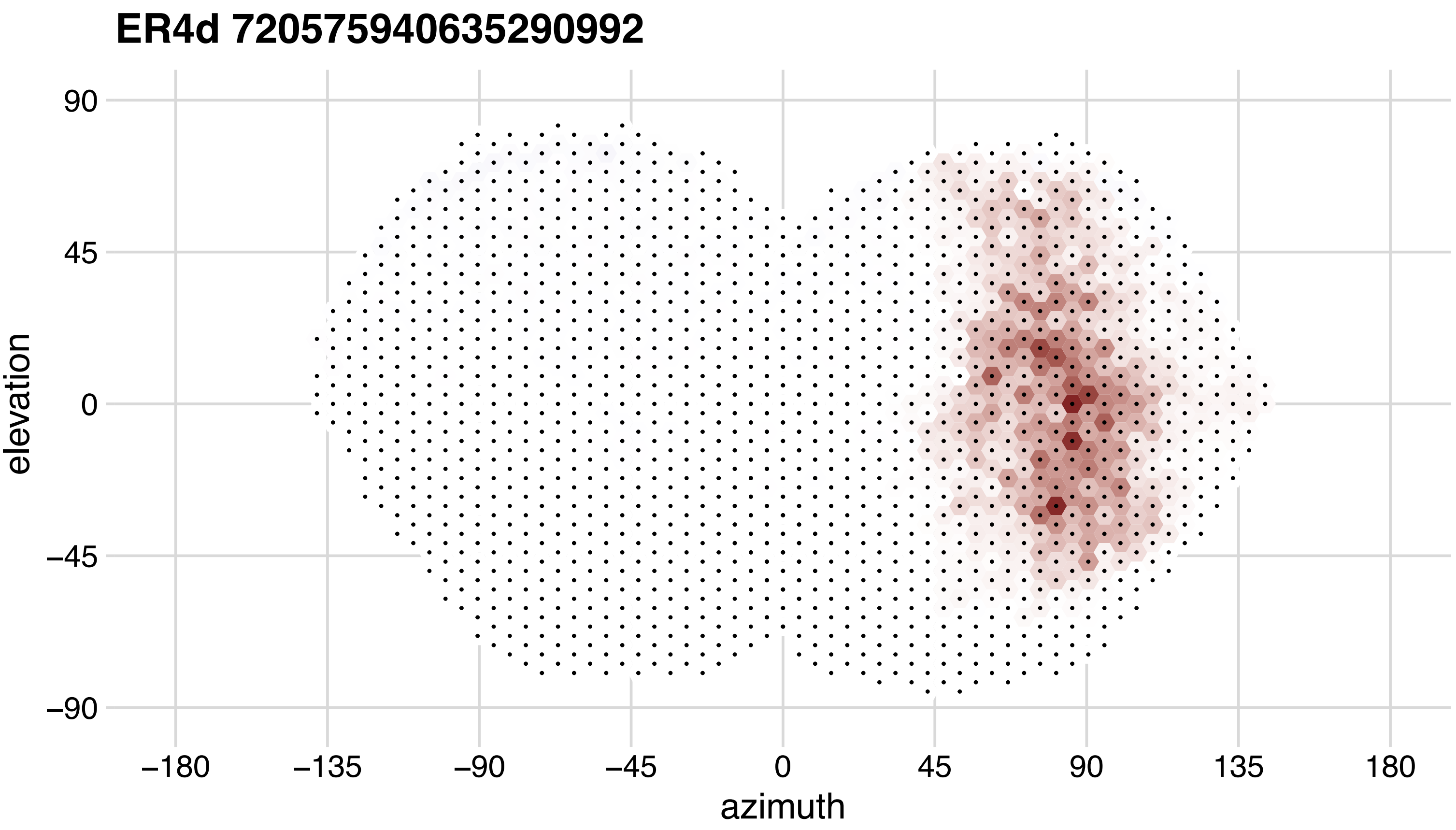

ER4d 720575940635290992
90
45
0
−45
−90
elevation
−180
−135
−90
−45
0
45
90
135
180
azimuth

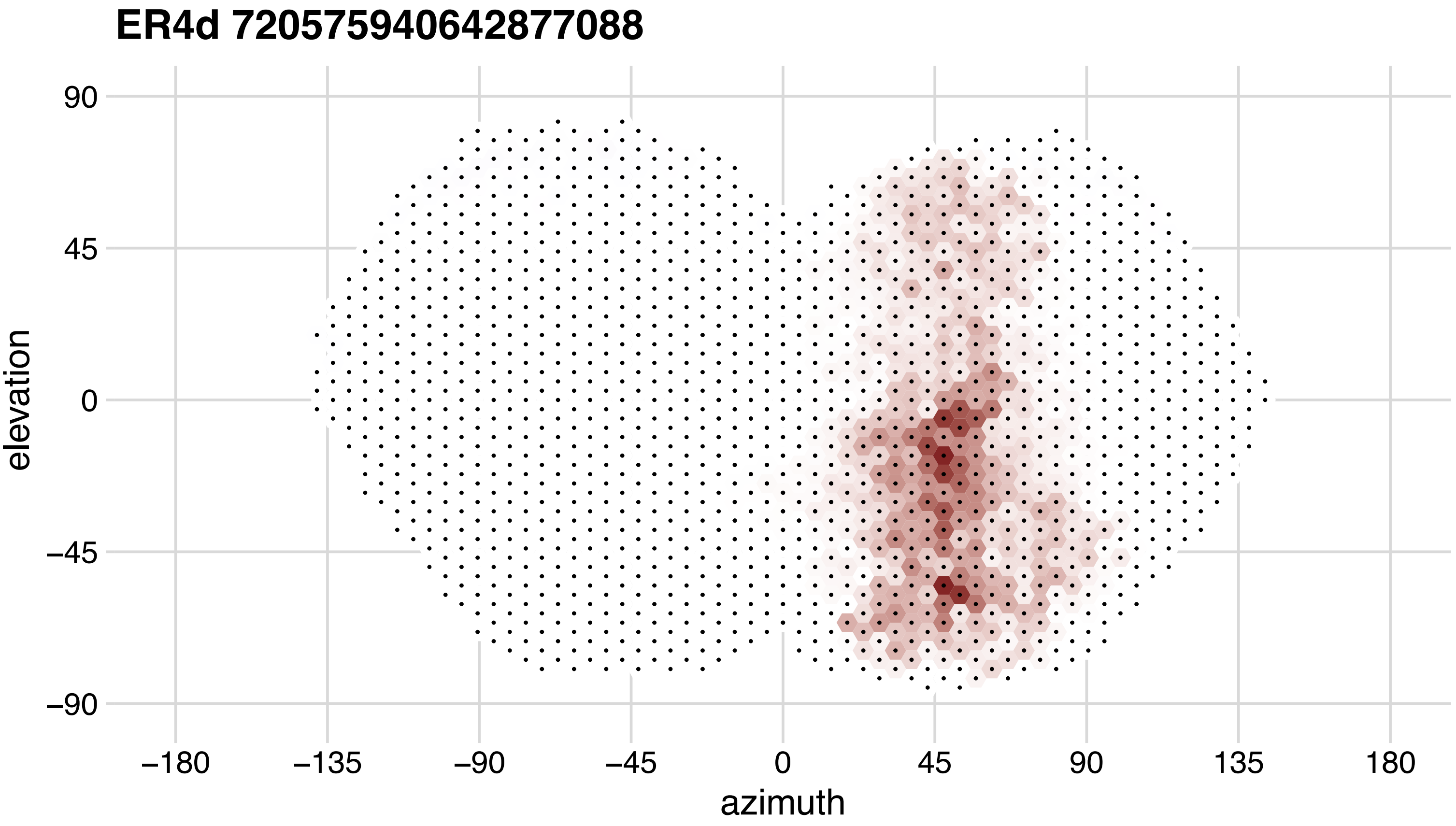
ER4d 720575940642877088
90
45
0
−45
−90
elevation
−180
−135
−90
−45
0
45
90
135
180
azimuth

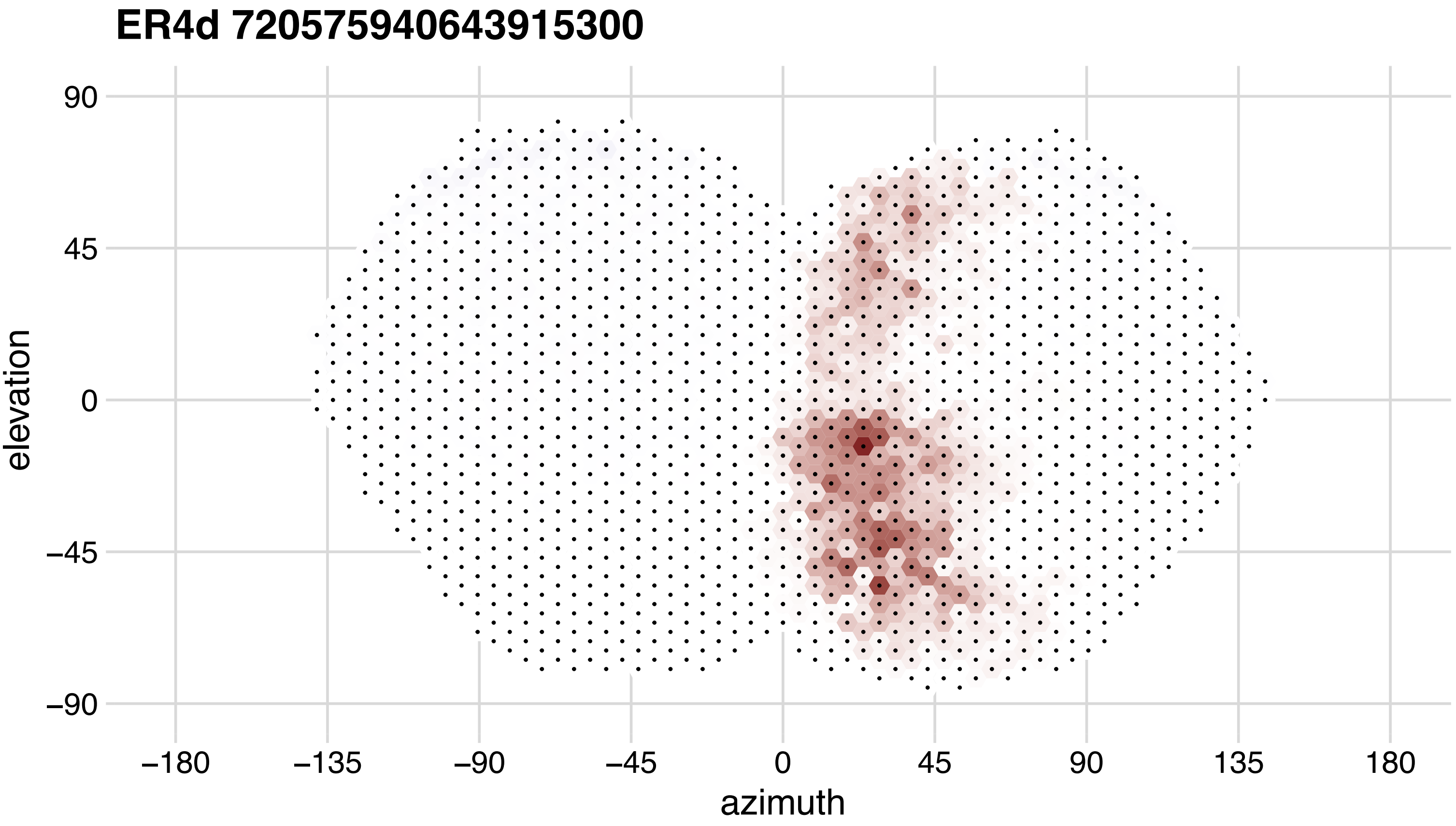

ER4d 720575940643915300
90
45
0
−45
−90
elevation
−180
−135
−90
−45
0
45
90
135
180
azimuth

## ER4m 720575940615712735

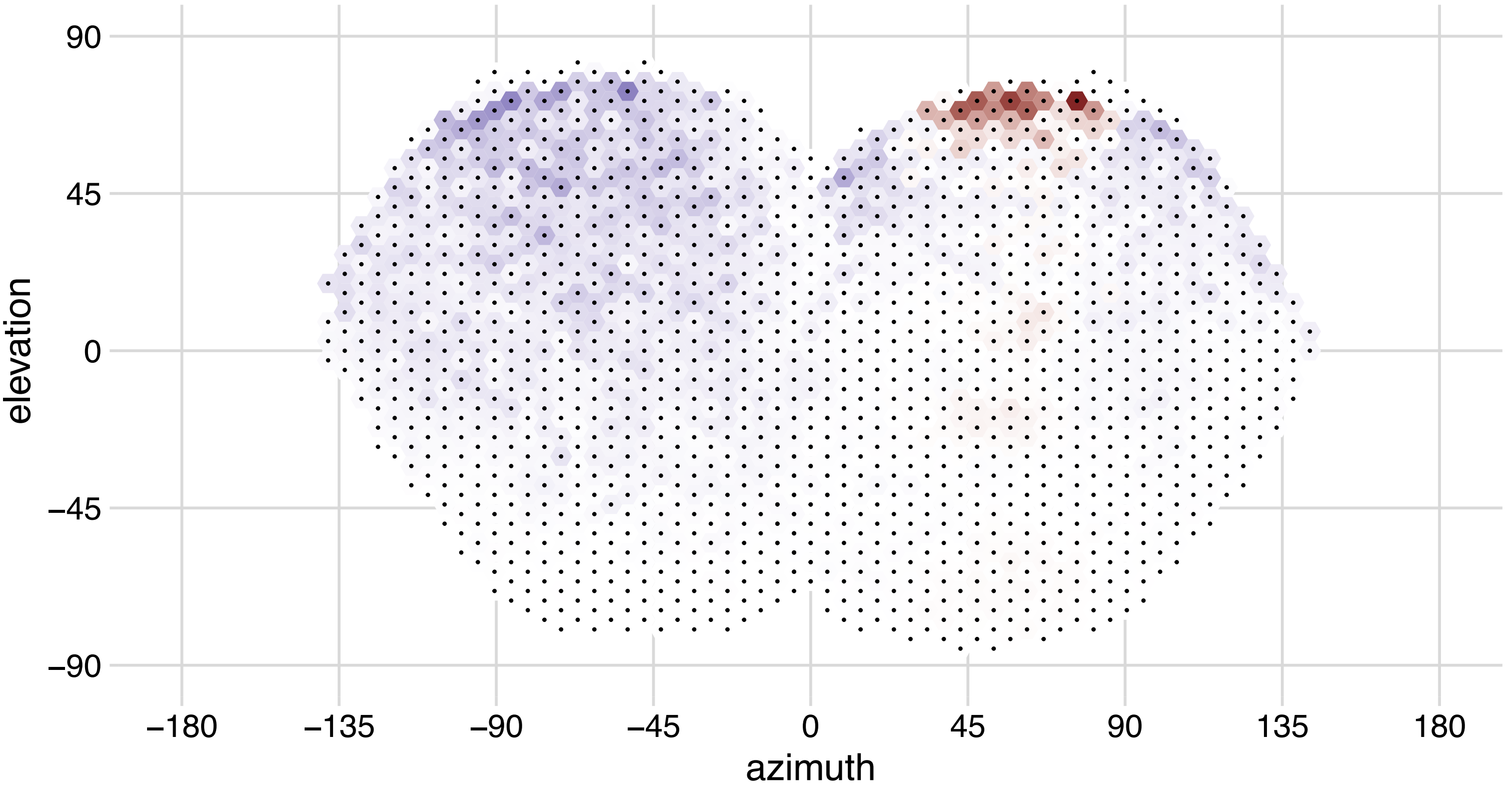

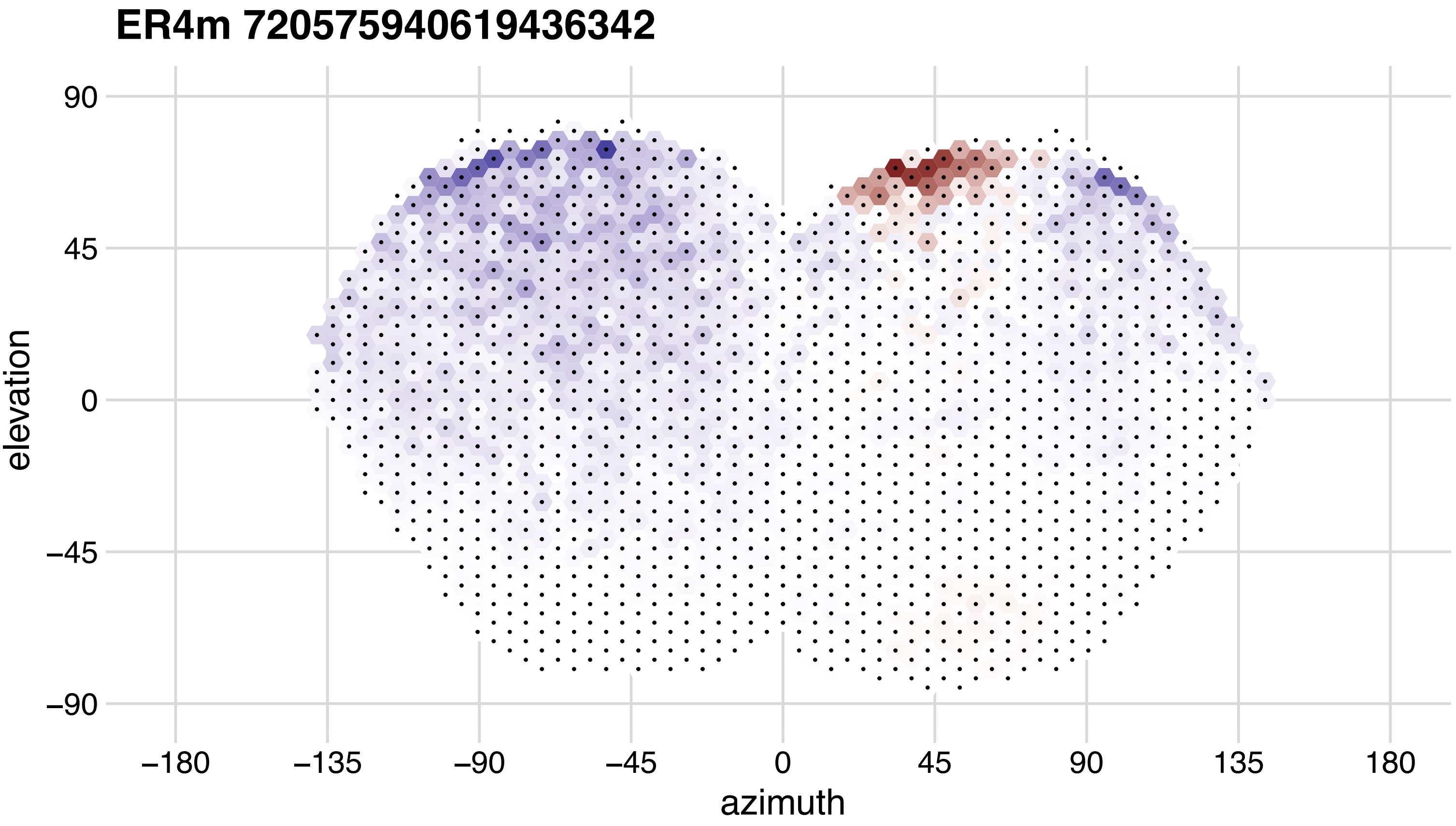
ER4m 720575940619436342
90
45
0
−45
−90
elevation
−180
−135
−90
−45
0
45
90
135
180
azimuth

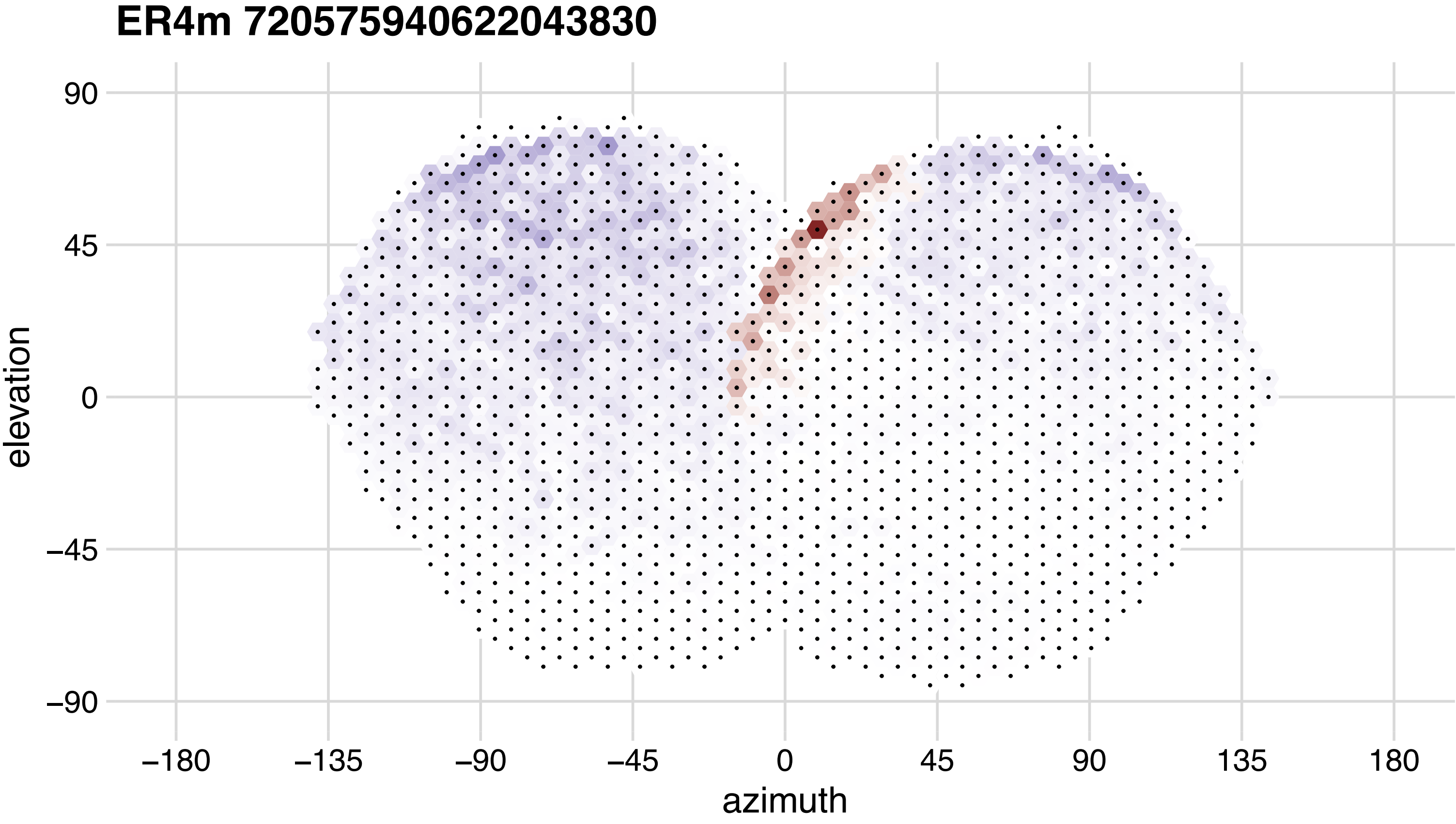
ER4m 720575940622043830
90
45
0
−45
−90
elevation
−180
−135
−90
−45
0
45
90
135
180
azimuth

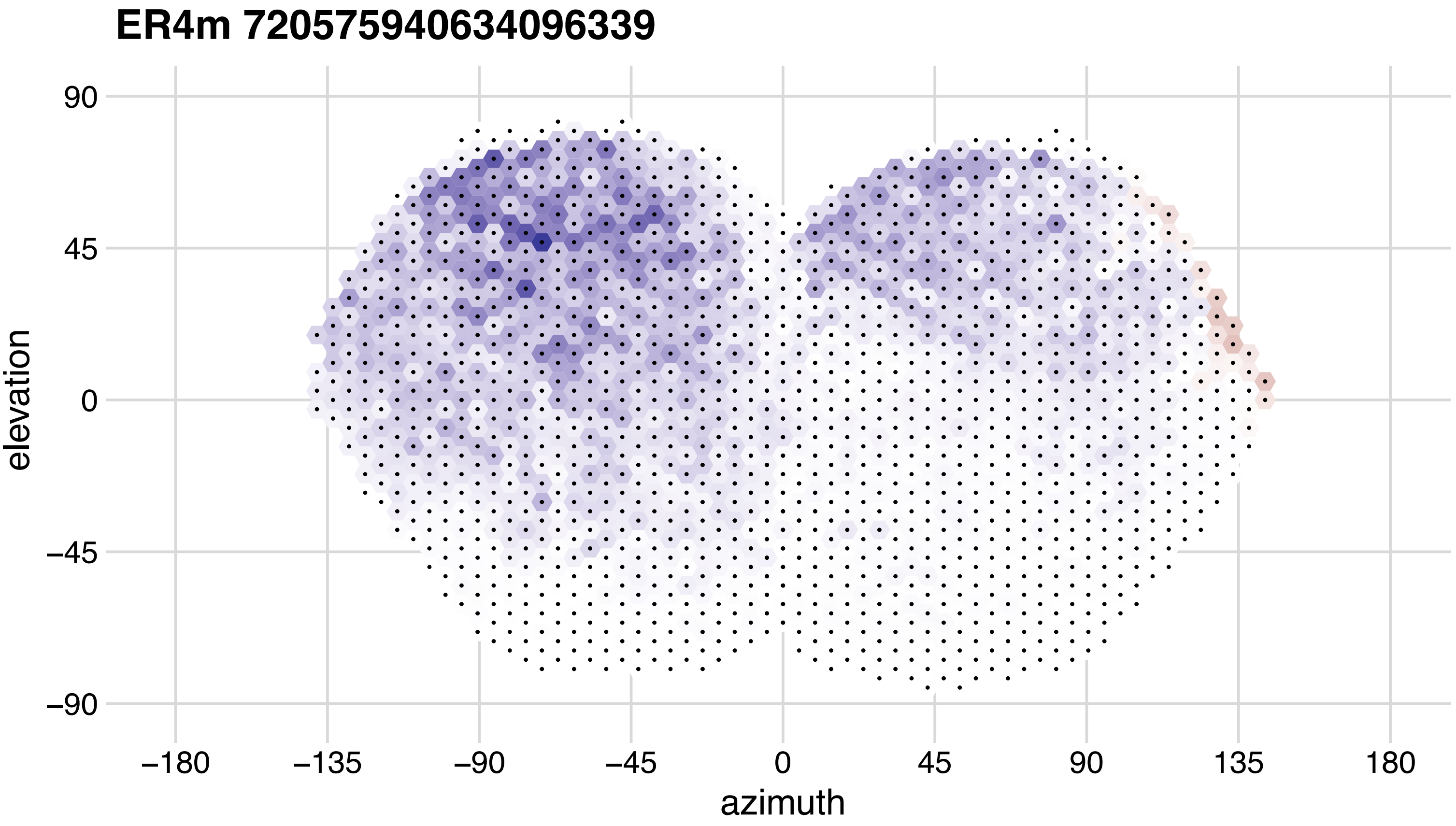

ER4m 720575940634096339
90
45
0
−45
−90
elevation
−180
−135
−90
−45
0
45
90
135
180
azimuth

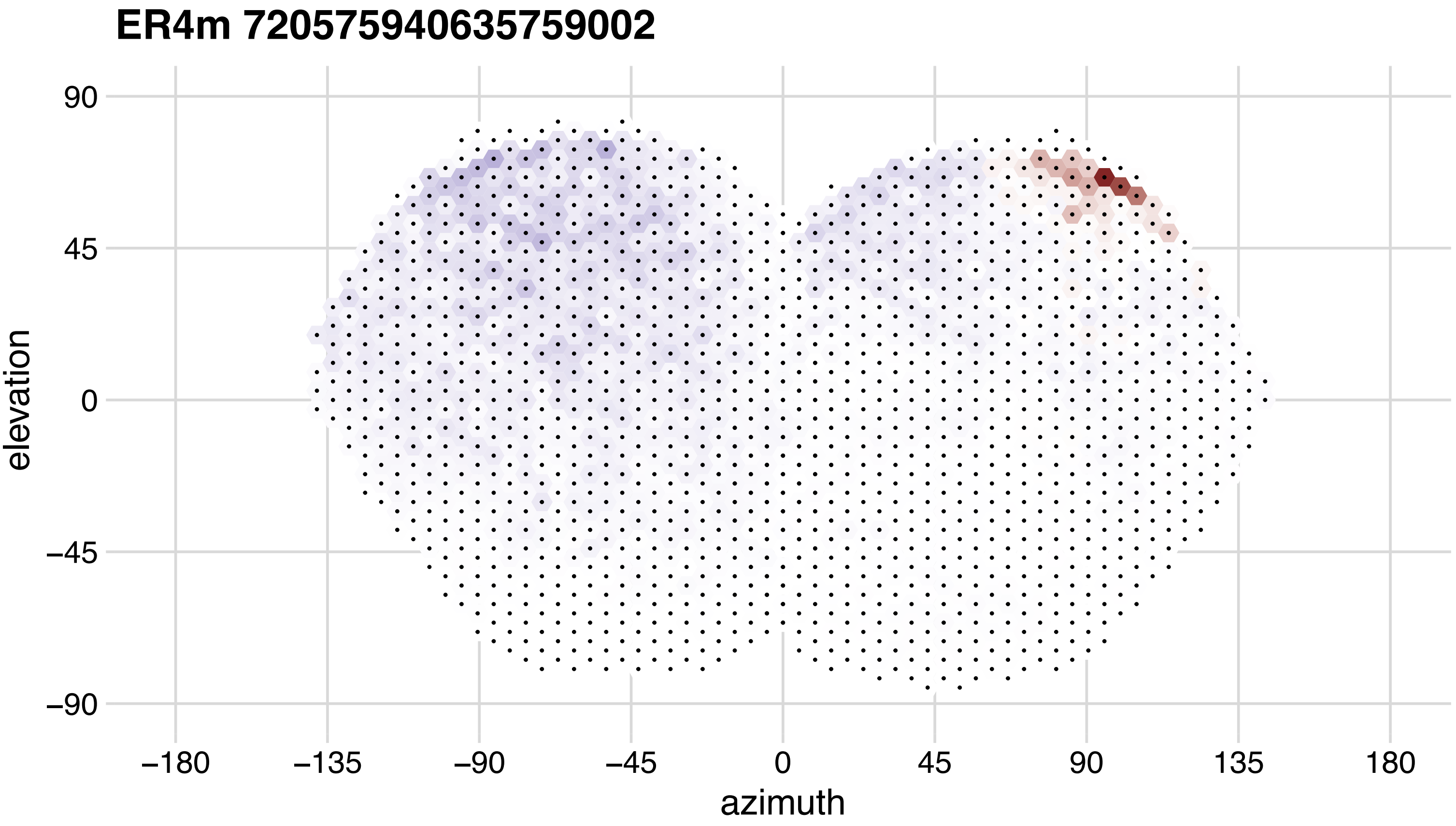

ER4m 720575940635759002
90
45
0
−45
−90
elevation
−180
−135
−90
−45
0
45
90
135
180
azimuth

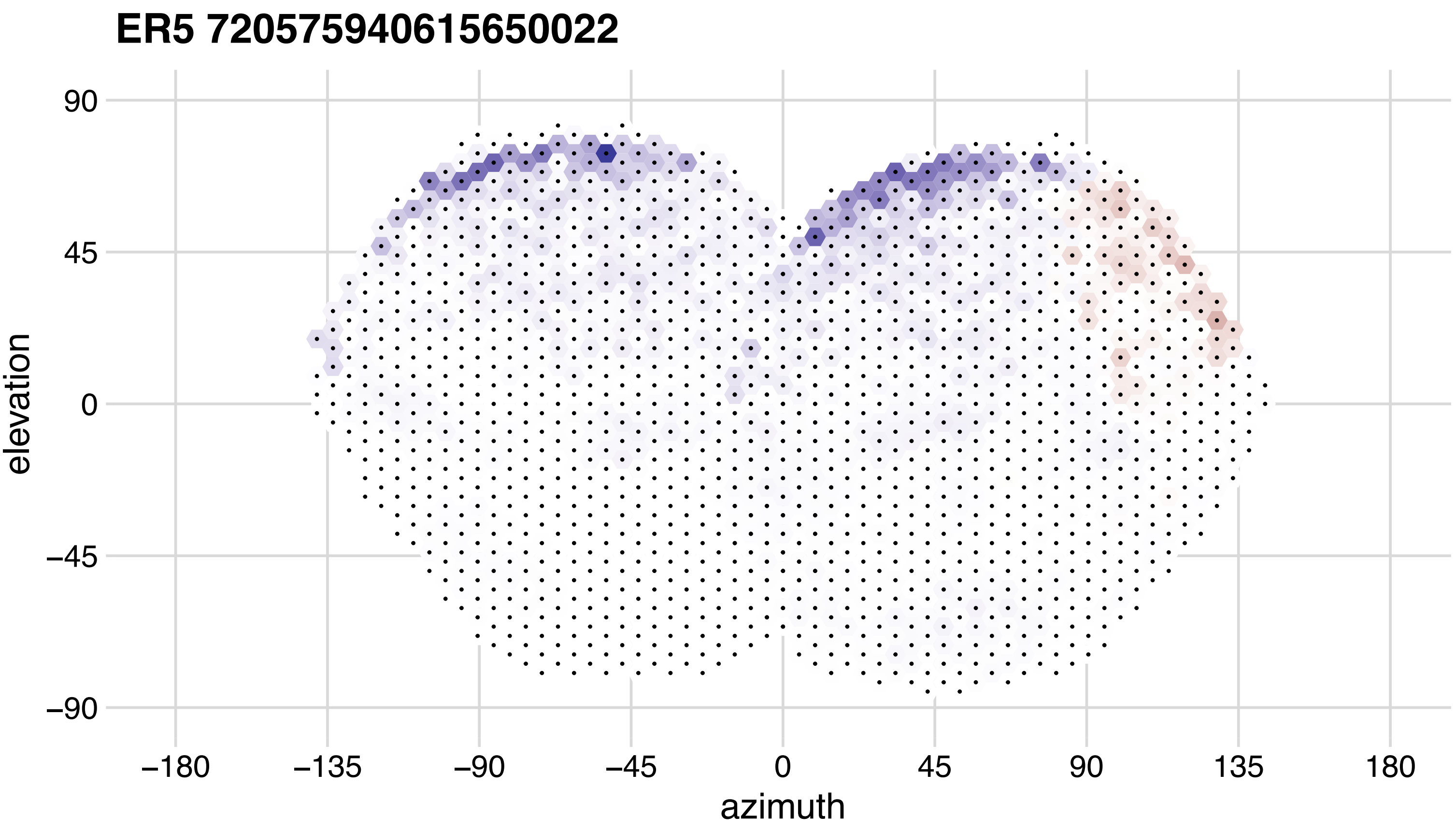
ER5 720575940615650022
elevation
90
45
0
−45
−90
−180
−135
−90
−45
0
45
90
135
180
azimuth

## ER5 720575940620940045

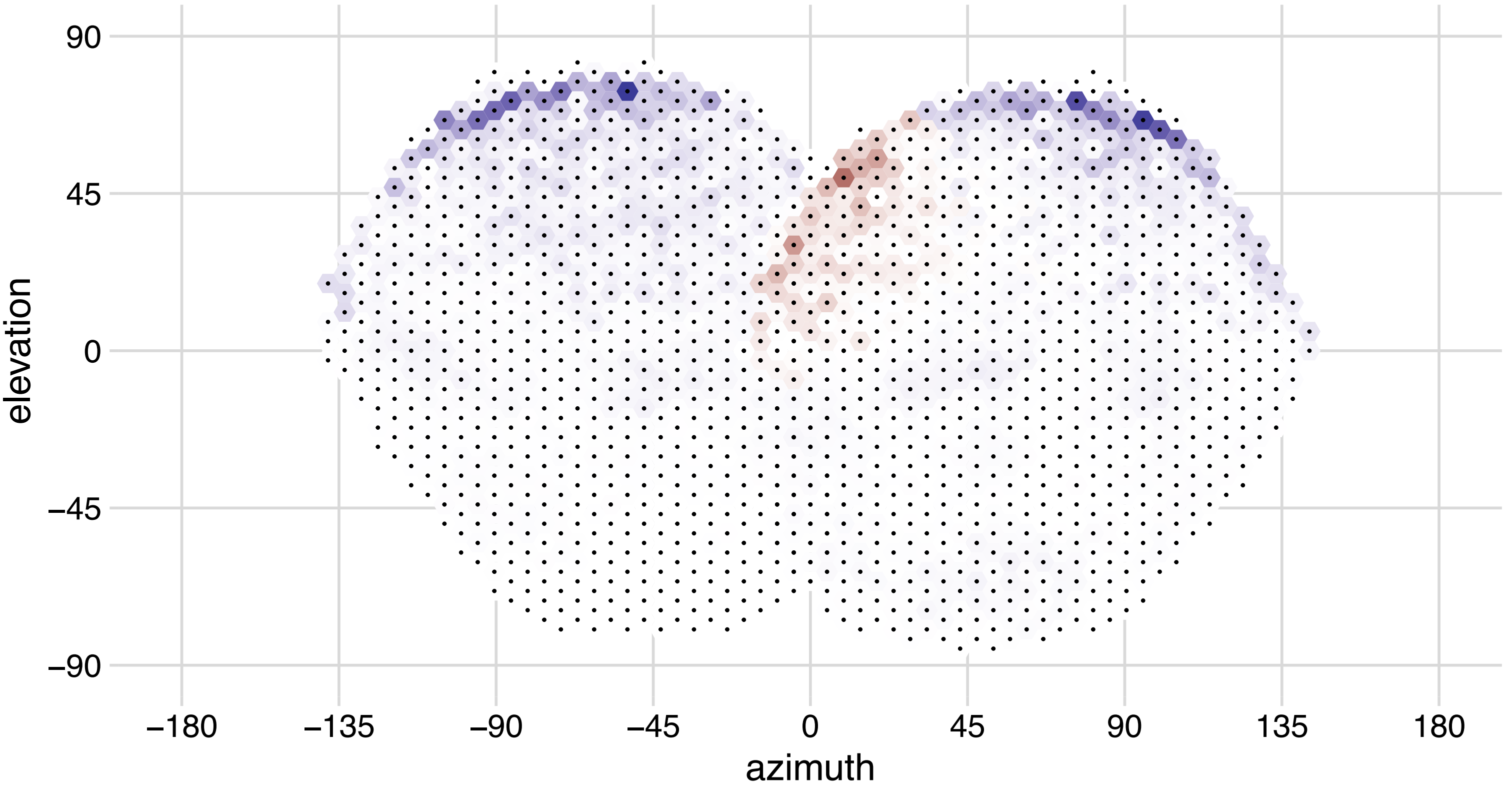

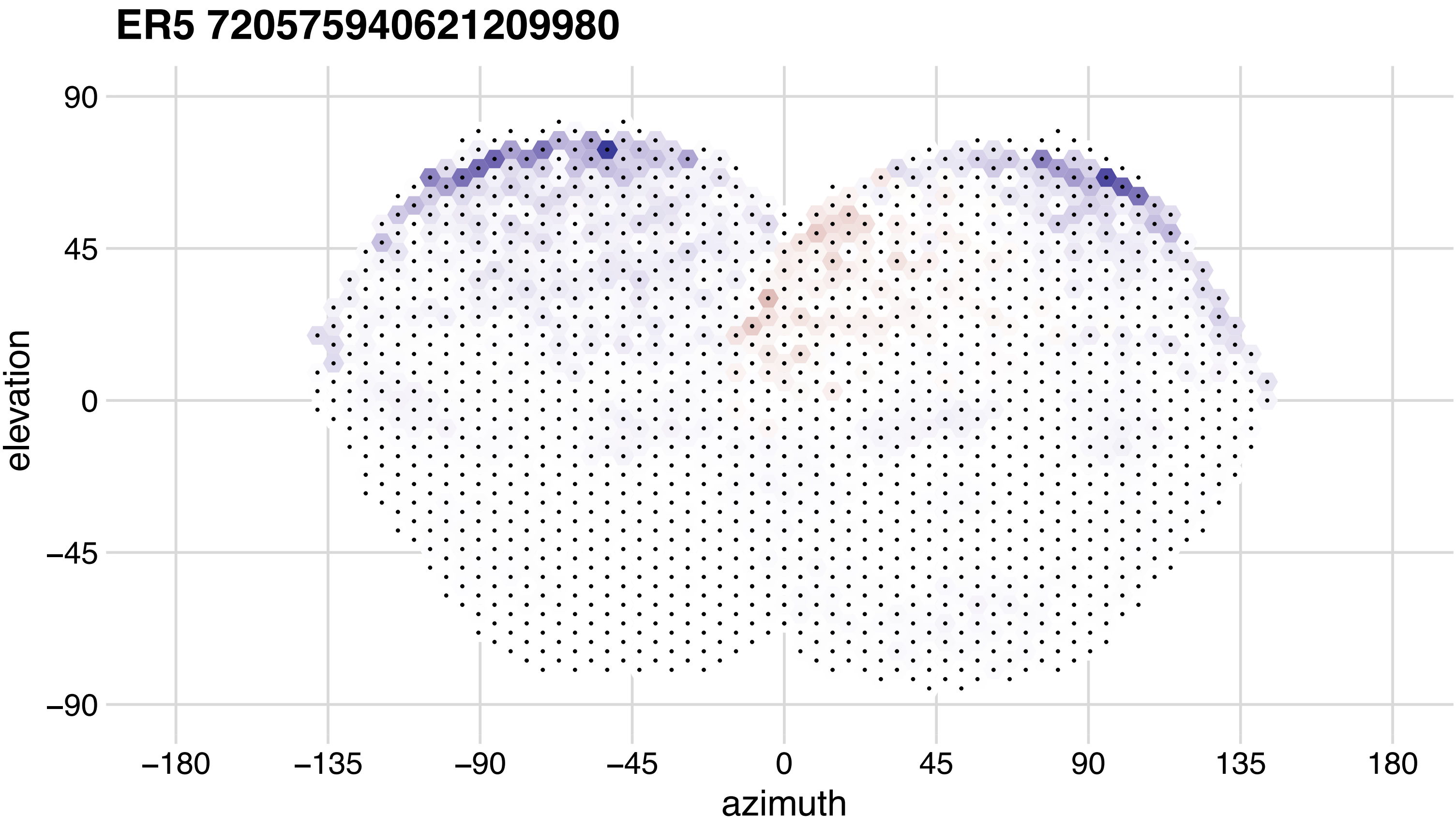
ER5 720575940621209980
90
45
0
−45
−90
elevation
−180
−135
−90
−45
0
45
90
135
180
azimuth

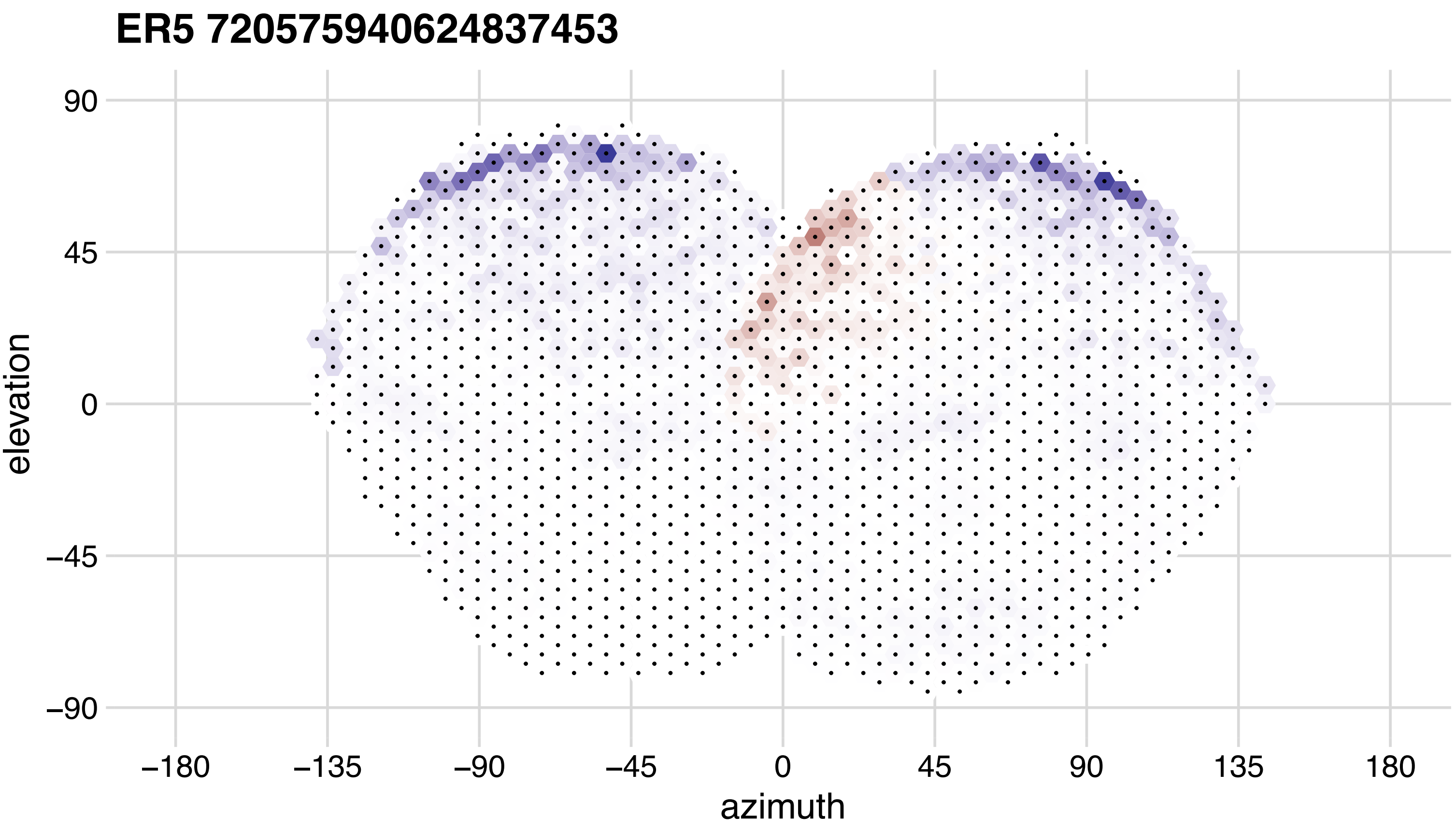
ER5 720575940624837453
elevation
90
45
0
−45
−90
−180
−135
−90
−45
0
45
90
135
180
azimuth

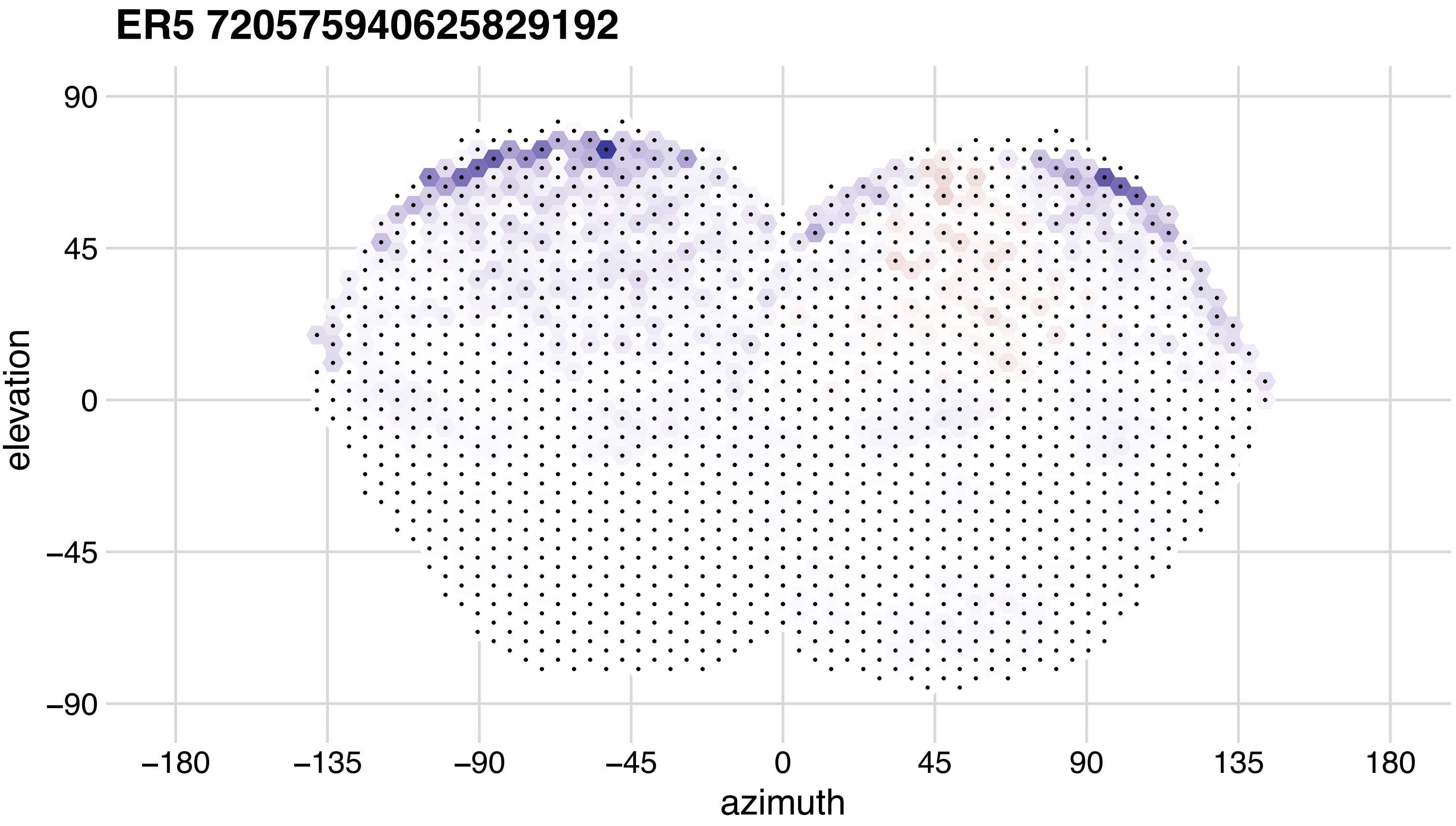
ER5 720575940625829192
90
45
0
−45
−90
elevation
−180
−135
−90
−45
0
45
90
135
180
azimuth

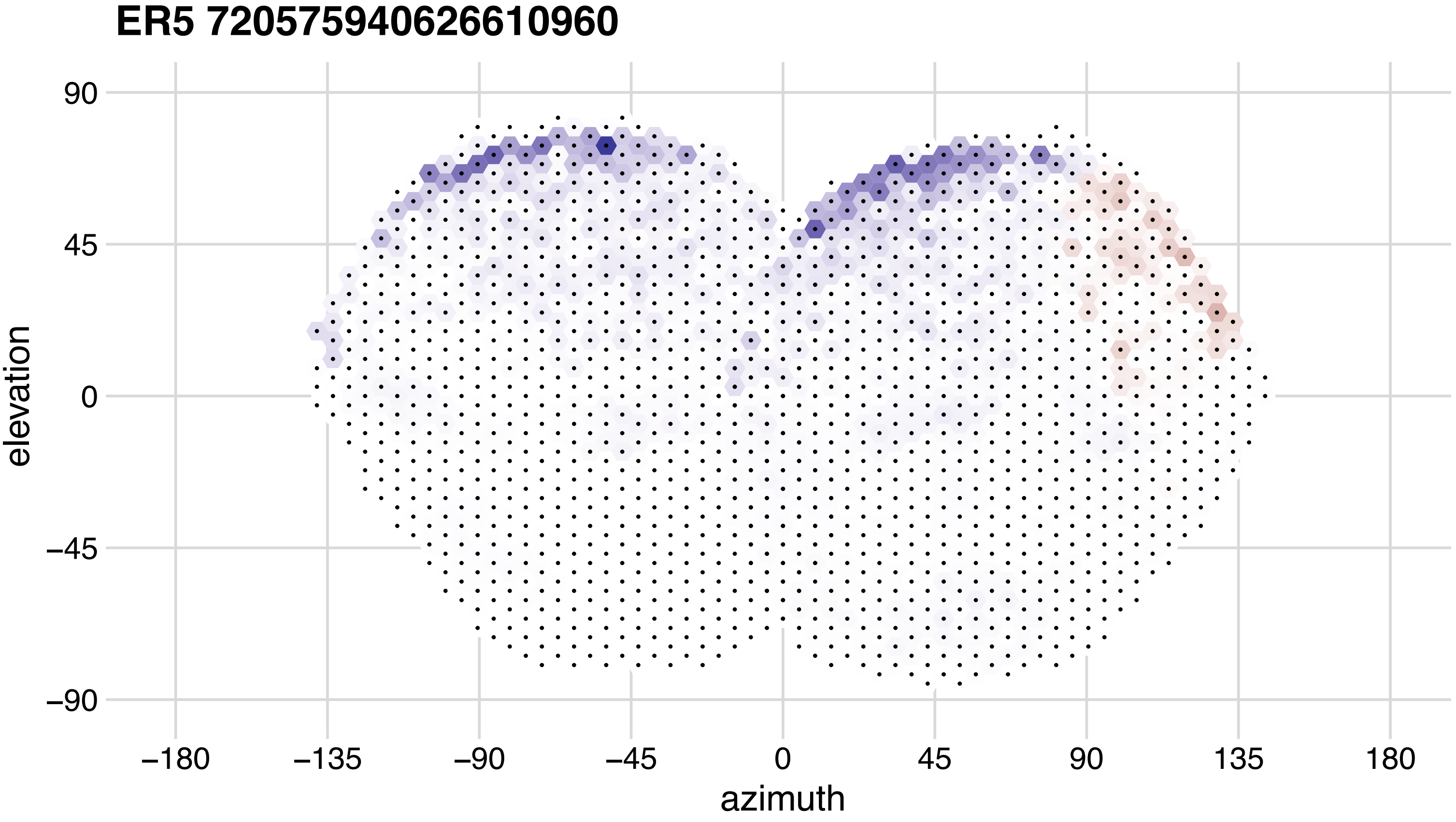
ER5 720575940626610960
90
45
0
−45
−90
elevation
−180
−135
−90
−45
0
45
90
135
180
azimuth

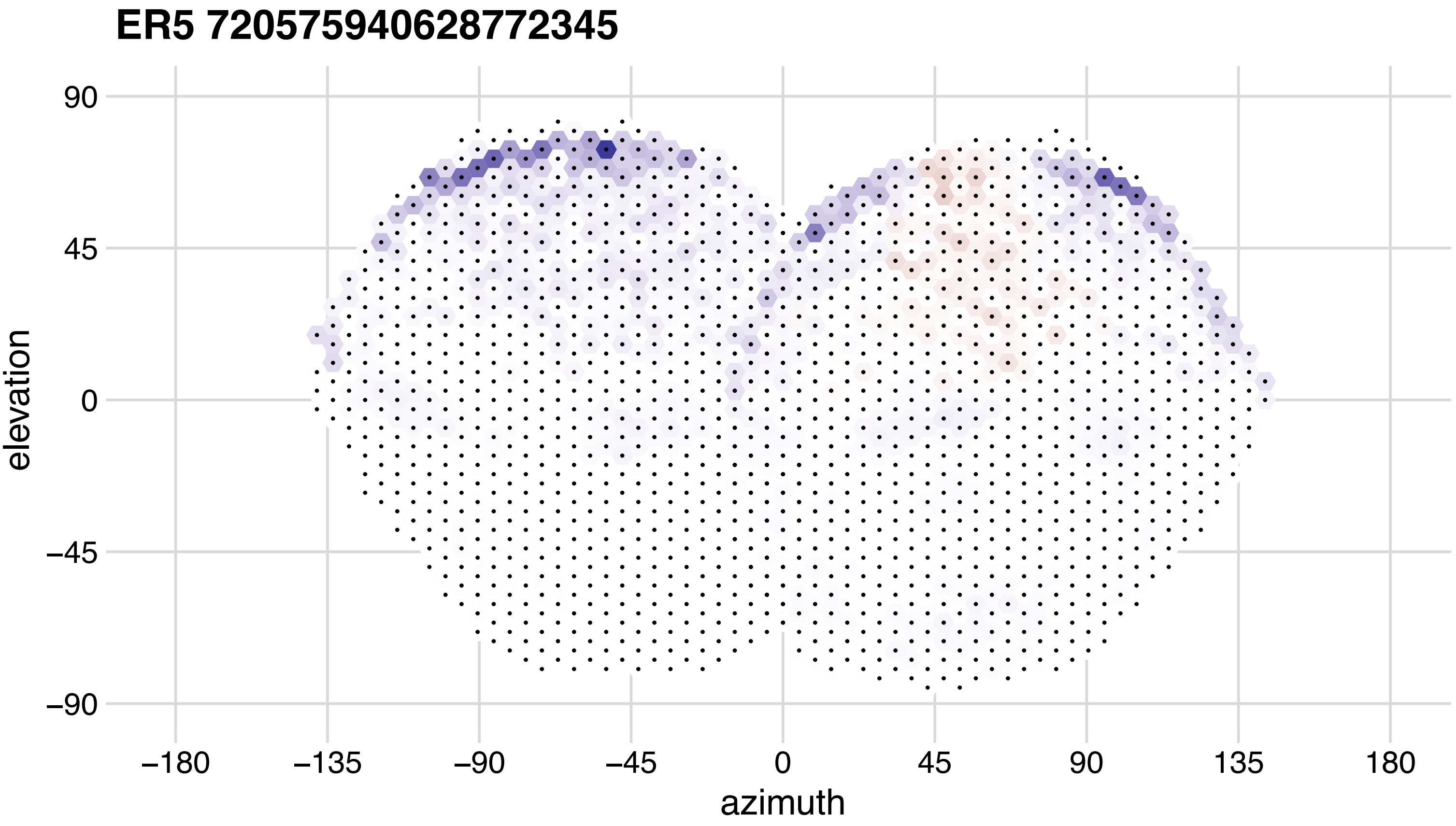
ER5 720575940628772345
90
45
0
−45
−90
elevation
−180
−135
−90
−45
0
45
90
135
180
azimuth

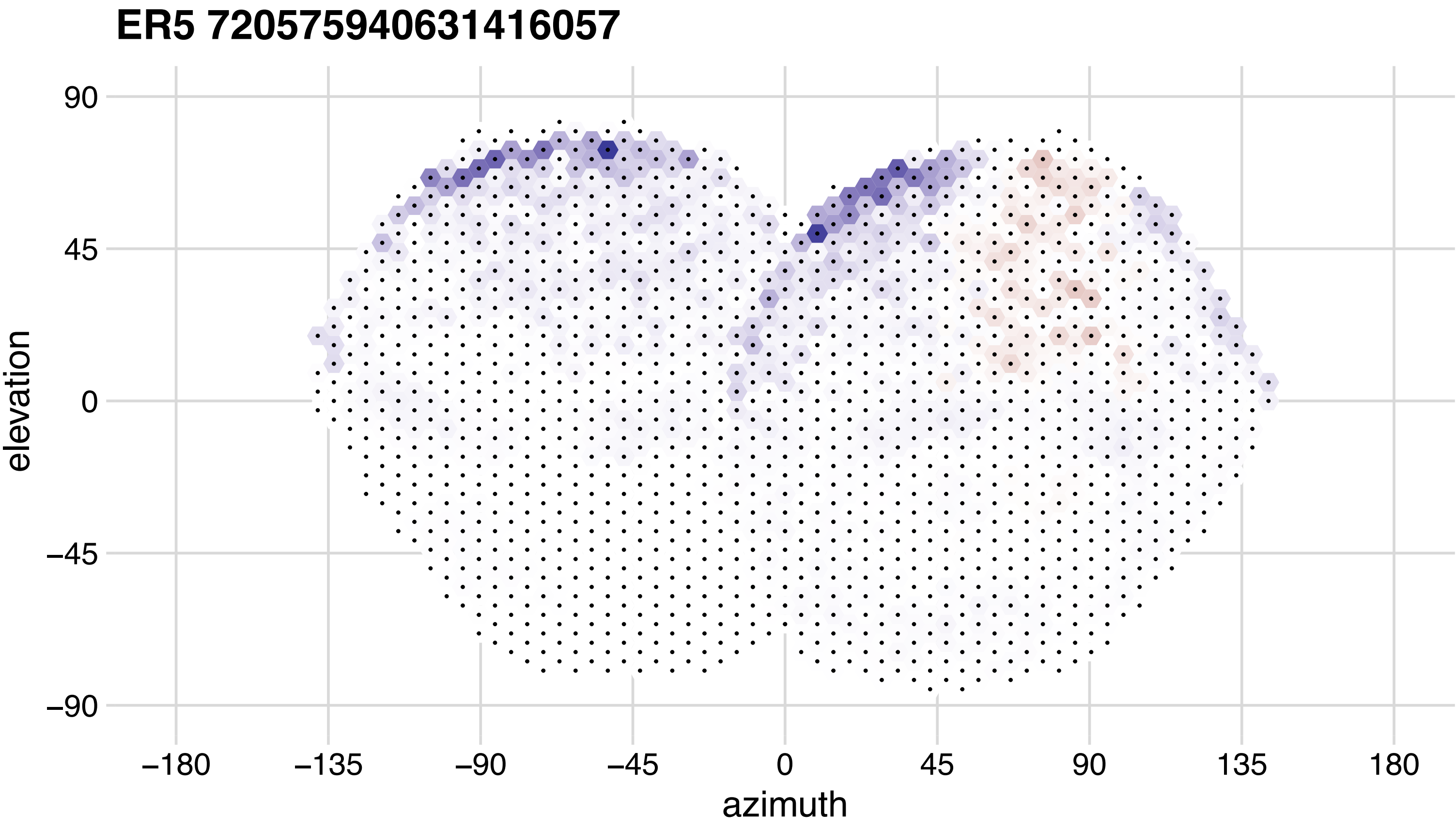

ER5 720575940631416057
90
45
0
−45
−90
elevation
−180
−135
−90
−45
0
45
90
135
180
azimuth

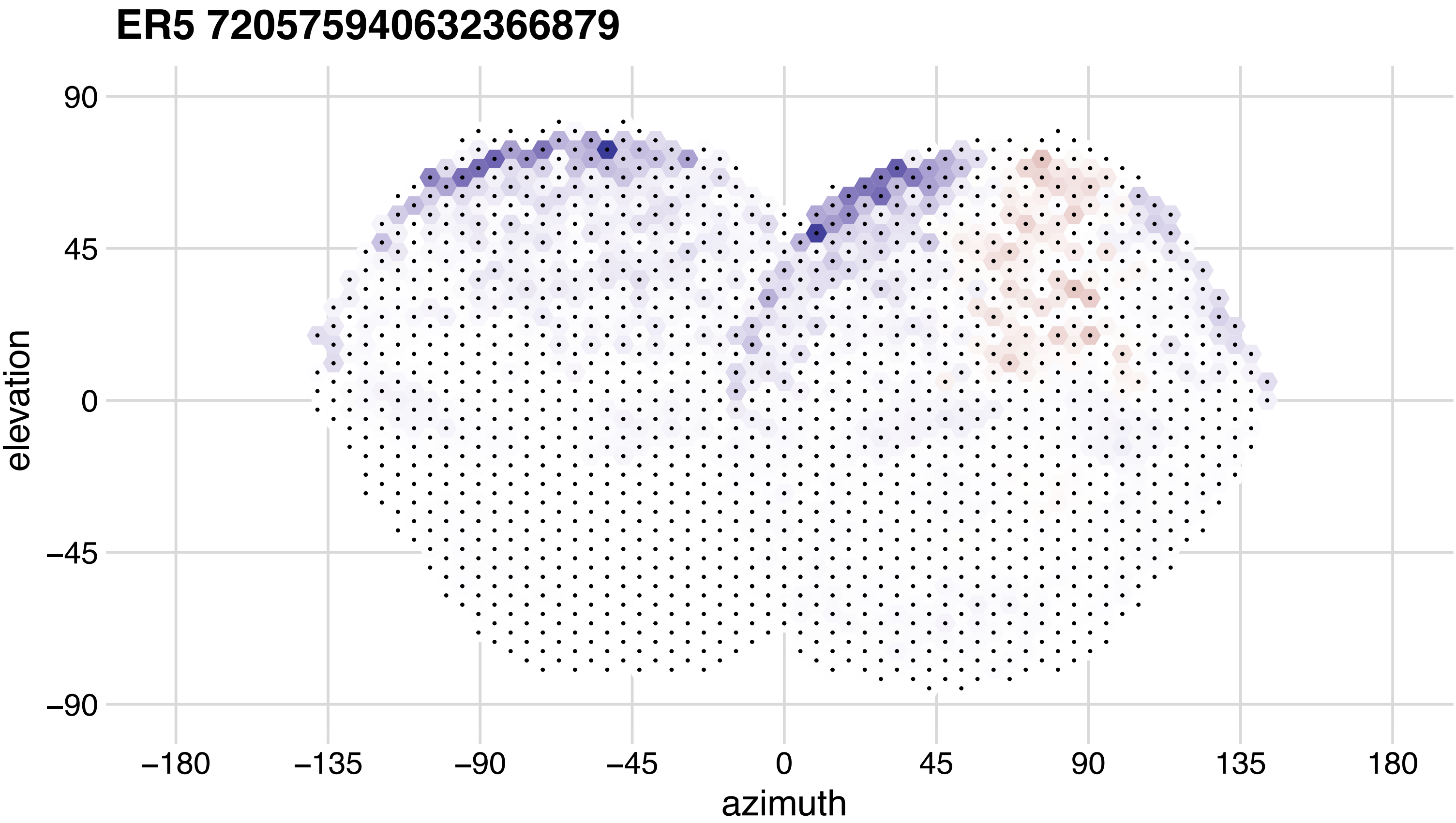
ER5 720575940632366879
90
45
0
−45
−90
elevation
−180
−135
−90
−45
0
45
90
135
180
azimuth

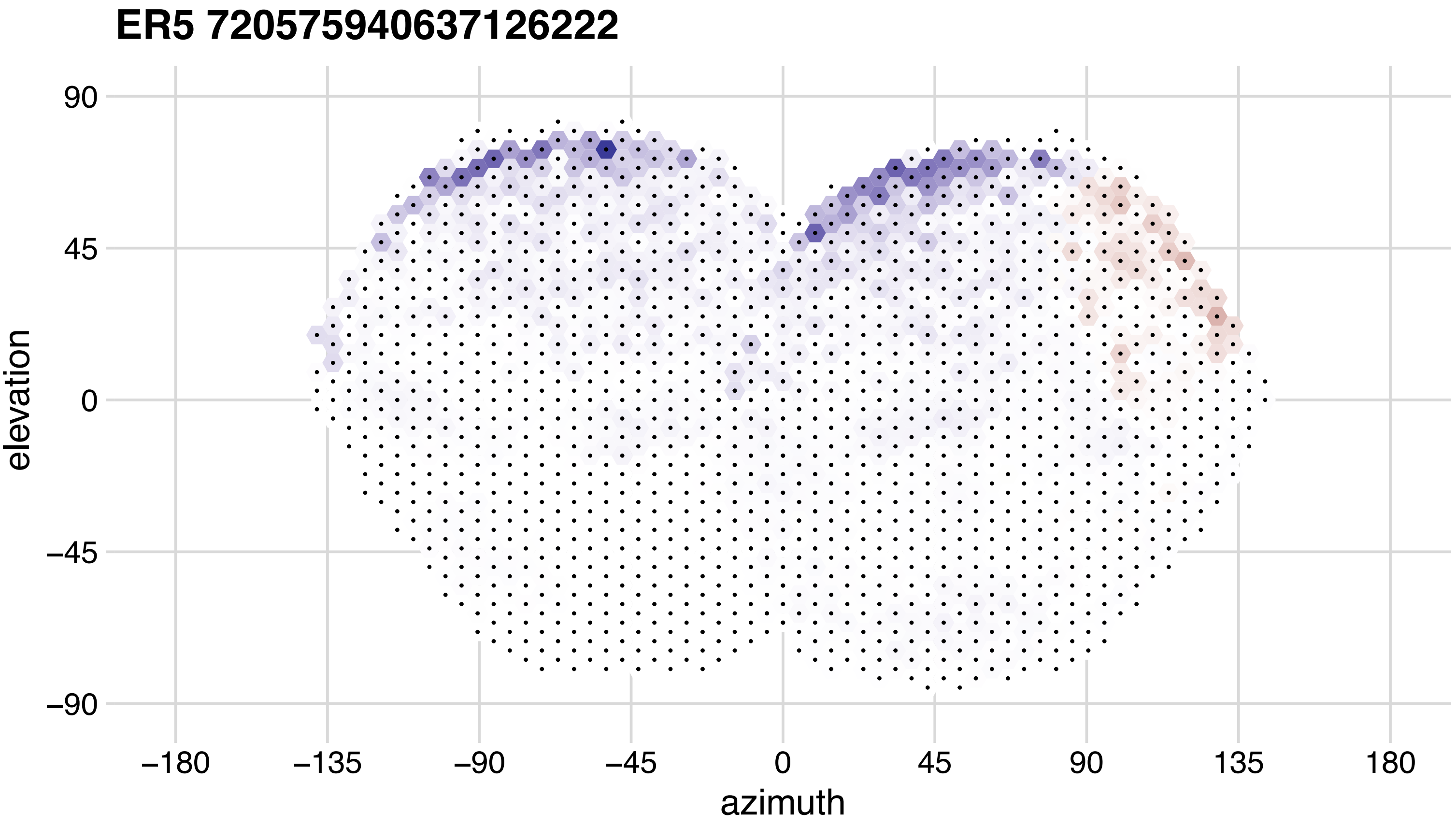

ER5 720575940637126222
90
45
0
−45
−90
elevation
−180
−135
−90
−45
0
45
90
135
180
azimuth

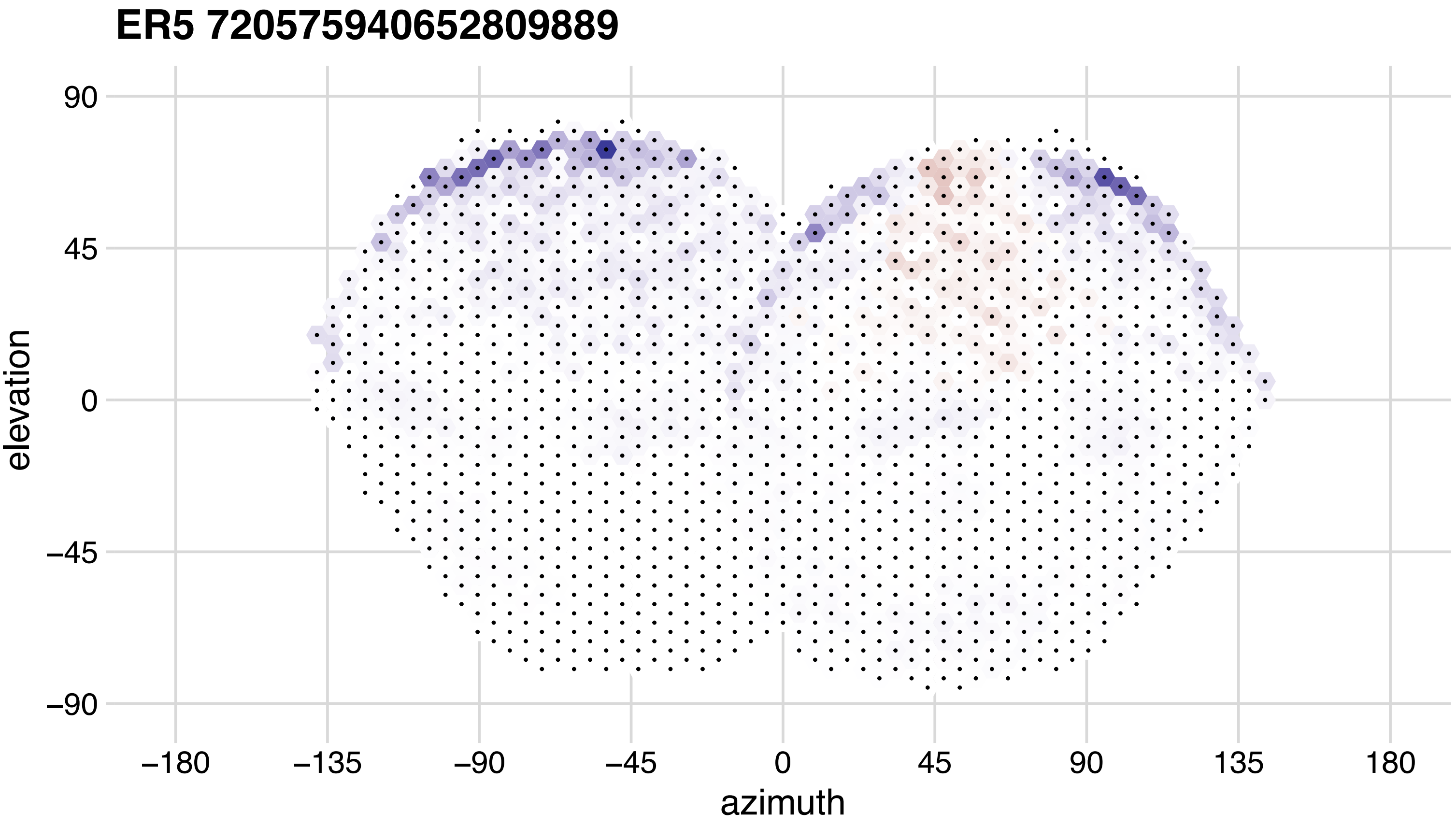
ER5 720575940652809889
90
45
0
−45
−90
elevation
−180
−135
−90
−45
0
45
90
135
180
azimuth

## ER6 720575940617523174

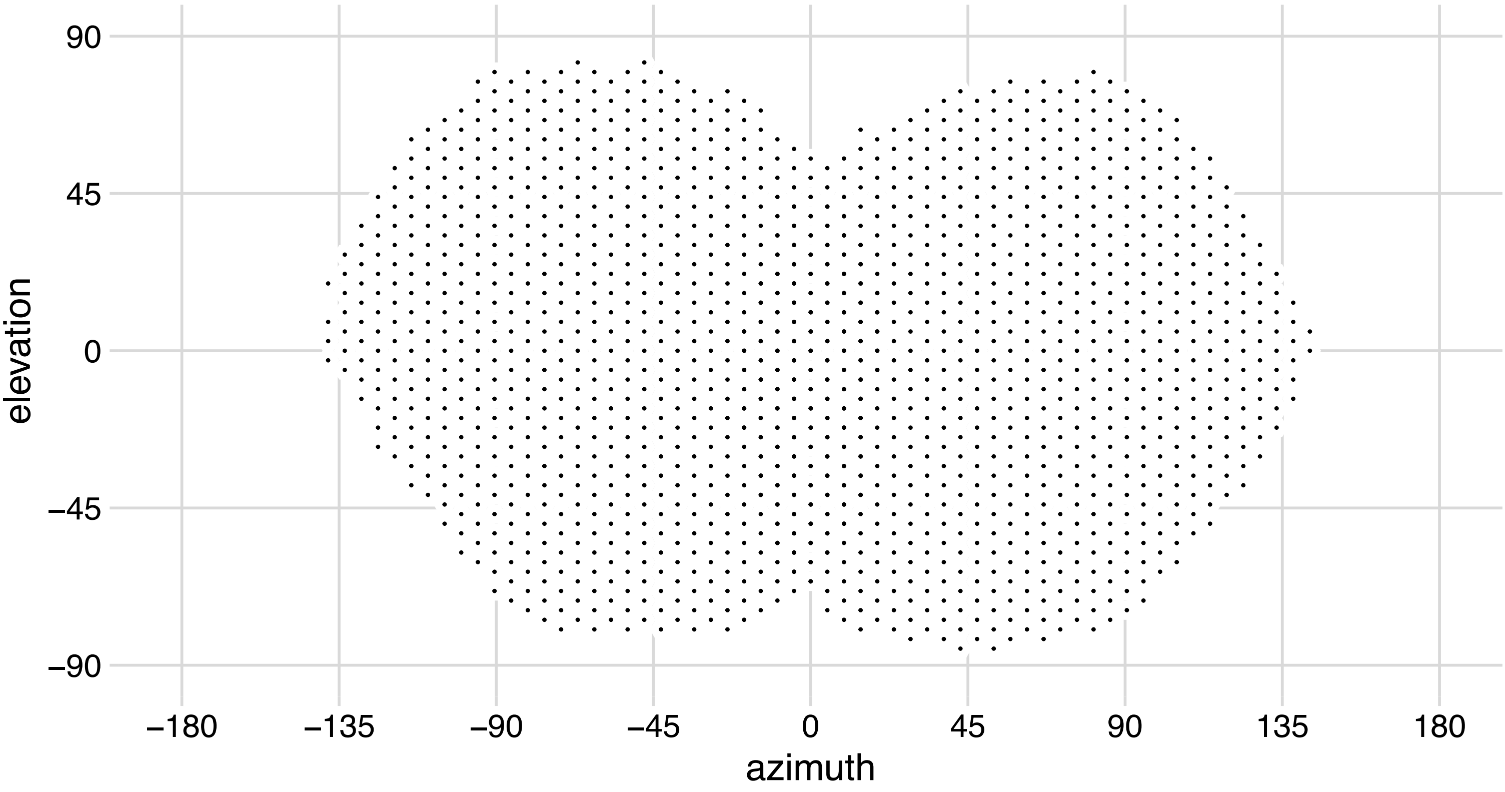

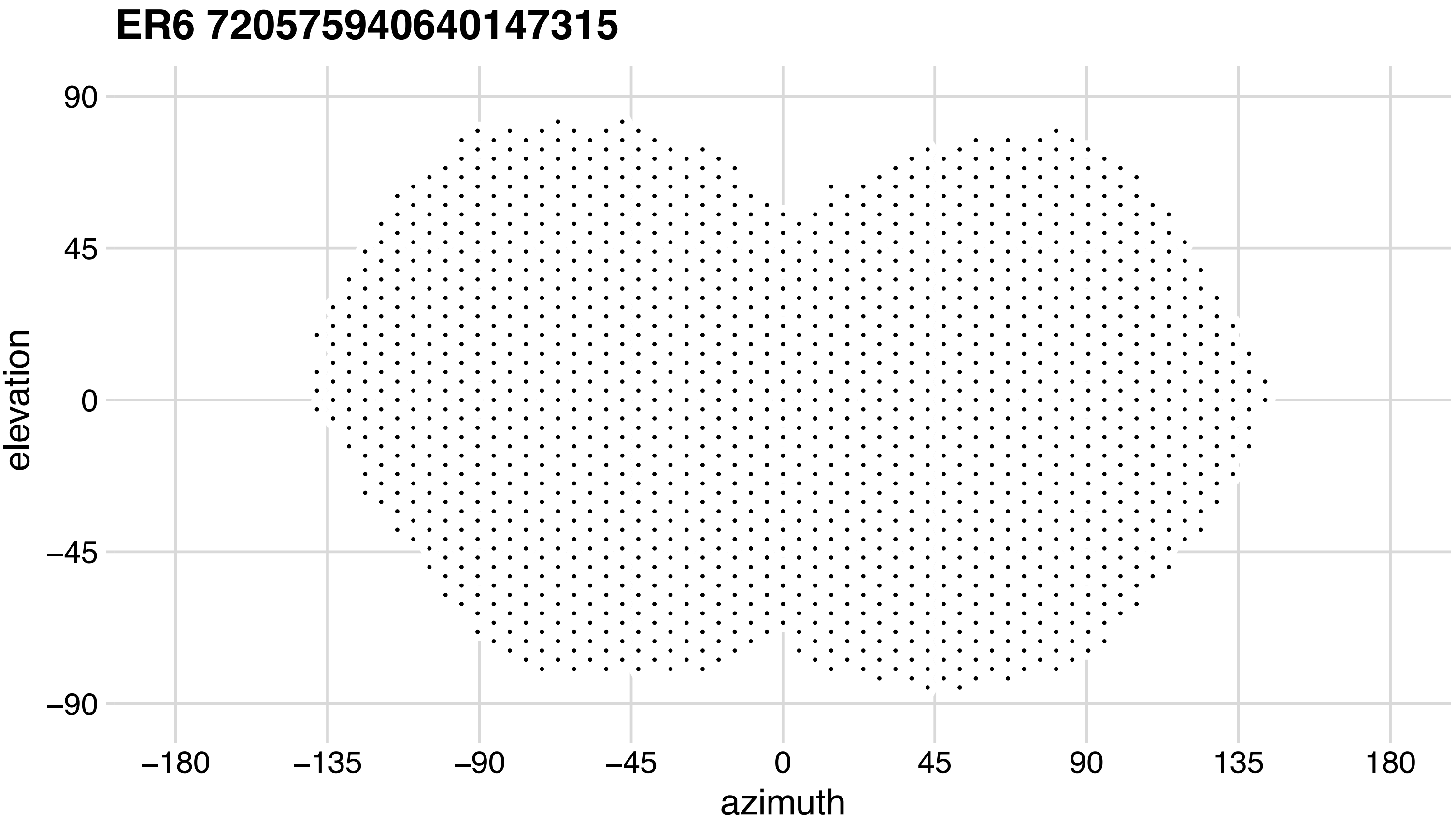

ER6 720575940640147315
90
45
0
−45
−90
elevation
−180
−135
−90
−45
0
45
90
135
180
azimuth
